# Multi-omics analysis reveals integrin α3-dependent mechanisms of Zika virus oncolytic activity in pediatric neural tumors

**DOI:** 10.64898/2026.05.01.722162

**Authors:** Yi Sui, Matthew Sherwood, Oswaldo K. Okamoto, Yihua Wang, Kevin Maringer, Rob. M. Ewing

**Author notes:** To whom correspondence should be addressed: Kevin Maringer, Rob M. Ewing.

## Abstract

Oncolytic virotherapy is an innovative approach to cancer treatment that uses replication-competent viruses to selectively target and destroy cancer cells while leaving healthy tissues largely unaffected. Zika virus (ZIKV), a neurotropic orthoflavivirus, has recently gained attention as a potential oncolytic agent due to its ability to infect neural-derived cells and suppress tumor growth in preclinical models. Although existing studies have examined ZIKV’s oncolytic effects, the mechanisms underlying these effects remain largely unexplored. Additionally, the roles of individual ZIKV proteins and their interactions with host factors remain incompletely understood. Here, we used RNA sequencing, affinity purification–mass spectrometry, and functional assays to uncover previously unidentified mechanisms underlying ZIKV’s oncolytic activity in pediatric neural tumors. We found that the ZIKV non-structural proteins NS4A and NS5 exert oncolytic effects, reducing tumorsphere size. ZIKV-host protein-protein interaction networks were characterized and showed that integrin α3 (gene: *ITGA3*), a mediator of cell-matrix adhesion, interacts with ZIKV NS2B and NS4A. Integrin α3 was further shown to be involved in ZIKV- and NS4A-induced tumorsphere size reduction, while *ITGA3* knockdown and ZIKV infection additively inhibited 3D invasion. These findings provide critical mechanistic insights that could inform the rational design of ZIKV-based virotherapies and highlight opportunities for combination treatment strategies.

## Introduction

Oncolytic virotherapy is an innovative cancer treatment strategy that employs replication-competent viruses to selectively infect and destroy malignant cells while sparing normal tissues^1,2^. These viruses exert their therapeutic effects through two complementary mechanisms: direct lysis of tumor cells and the induction of systemic anti-tumor immunity^3,4^. Modern oncolytic viruses (OVs) are often genetically engineered to enhance tumor specificity, improve safety, and incorporate transgenes that stimulate immune responses or deliver additional therapeutic payloads^3,5–8^. One emerging candidate for oncolytic virotherapy is Zika virus (ZIKV), which has recently gained attention and is the focus of this study. ZIKV is a mosquito-borne virus belonging to the *Orthoflavivirus* genus in the Flaviviridae family, which is a family of positive-sense, single-stranded, enveloped RNA viruses^9^. The RNA genome of ZIKV encodes three structural proteins: capsid, precursor membrane (prM), and envelope (E), as well as seven non-structural proteins (NS1, NS2A, NS2B, NS3, NS4A, NS4B, and NS5)^10^. ZIKV non-structural proteins play important roles in viral genome replication, polyprotein processing, and modulation of host immune responses, while structural proteins are essential for viral entry, assembly, and particle formation^11,12^. Most adults infected with ZIKV are asymptomatic, however, ZIKV infection can cause microcephaly and other congenital abnormalities in the developing fetus and newborn and is also a trigger for Guillain-Barré syndrome, neuropathy, and myelitis, particularly in adults and older children^13–15^.

Existing studies on ZIKV-based oncolytic virotherapy mainly focus on treating central nervous system (CNS) tumors due to ZIKV’s neurotropism^16,17^. CNS tumors are abnormal growths of cells originating from the brain and spinal cord^18,19^. The most studied CNS tumors in this context include glioblastoma, atypical teratoid/rhabdoid tumor (AT/RT), and medulloblastoma^6,16,17,20,21^. AT/RT and medulloblastoma are pediatric brain tumors, and studies have shown that ZIKV induces more pronounced cell death in AT/RT cells and tumorspheres compared to medulloblastoma^16^. AT/RT is a rare and highly aggressive CNS tumor, especially in children under three years old^22,23^. It can occur throughout the brain and spinal cord, although it most commonly arises in the cerebellum or brainstem^24^.

Beyond CNS tumors, another pediatric neural tumor, neuroblastoma, has begun to receive attention for ZIKV-based oncolytic virotherapy. Neuroblastoma, a malignancy of the sympathetic nervous system that originates from neural crest-derived cells, primarily affects infants and young children^25^. It most commonly occurs in the adrenal glands or paravertebral ganglia^26^. As the most common tumor in children younger than five years old, neuroblastoma accounts for approximately 15% of all pediatric cancer deaths^27^. Although ZIKV’s oncolytic effects have been well studied in multiple models of pediatric brain tumors^16,28,29^, its application to neuroblastoma remains relatively unexplored. Previous studies have investigated ZIKV’s oncolytic potential in neuroblastoma using two-dimensional (2D) cell cultures and mouse models^30,31^, however, gaps remain regarding its effects in three-dimensional (3D) tumor models and the mechanisms underlying ZIKV’s oncolytic activity. Moreover, while studies have examined ZIKV’s oncolytic effects in various tumors, the roles of individual ZIKV proteins in these processes have rarely been studied. This knowledge could provide deeper mechanistic insights and inform strategies to enhance tumor selectivity and anti-tumor efficacy.

In this study, we aimed to identify key mechanisms underlying ZIKV’s oncolytic effects in pediatric neural tumors by investigating common mechanisms in both neuroblastoma and pediatric brain tumors through a multi-omics approach. AT/RT was chosen as a representative pediatric brain tumor model due to its heightened susceptibility to ZIKV-induced cell death^16^. Both 2D and 3D models were used in this study. Since ZIKV has already been shown to exert oncolytic effects in an AT/RT 3D model^16^ but has not been investigated in a neuroblastoma 3D model, we first employed neuroblastoma tumorsphere 3D models to confirm ZIKV’s oncolytic activity. RNA sequencing (RNA-Seq) was then performed to analyze the underlying mechanisms. We further examined the roles of ZIKV non-structural proteins in tumorsphere formation and conducted affinity proteomics to identify host factors interacting with these proteins. Comparative multi-omics analysis of the two tumor models was then performed, enabling a broader understanding of potential mechanisms underlying ZIKV-mediated oncolysis. Our findings demonstrate that ZIKV exerts potent oncolytic effects in 3D neuroblastoma models. We identified integrin α3 (gene name: *ITGA3*) as a key interactor with NS2B and NS4A in both neuroblastoma and pediatric brain tumor models and found that it is involved in ZIKV- and NS4A-induced reductions in tumorsphere size. Moreover, *ITGA3* knockdown and ZIKV infection additively suppressed 3D invasion, highlighting the therapeutic potential of combining ZIKV-based virotherapy with integrin-targeting strategies. Collectively, these results provide mechanistic insights into ZIKV-mediated oncolysis, offering promising avenues for novel treatments in pediatric neural tumors.

## Results

### ZIKV exhibits oncolytic effects in neuroblastoma models

To investigate the common mechanisms underlying ZIKV’s oncolytic effects in both neuroblastoma and pediatric brain tumors, we first examined its oncolytic effects in neuroblastoma, which have not yet been well demonstrated. SH-SY5Y cells, derived from a metastatic bone tumor in a four-year-old patient with non-MYCN-amplified neuroblastoma, and IMR-32 cells, established from an abdominal mass in a 13-month-old patient with MYCN-amplified neuroblastoma, were infected with either the ZIKV PRVABC59 or BeH819015 strain at a multiplicity of infection (MOI) of 2 in monolayer cultures. Cell viability decreased over time in both cell lines following infection with either strain (**Figure S1A and B**). Interestingly, both strains induced earlier virus-mediated cell death in SH-SY5Y cells compared with IMR-32 cells. Virus growth kinetics also revealed a delay in replication in IMR-32 cells compared with SH-SY5Y cells (**Figure S1C and D**).

SH-SY5Y and IMR-32 tumorspheres were subsequently infected with either PRVABC59 or BeH819015 strain at an MOI of 2 or 10. Tumorsphere size was significantly reduced at 8 days post-infection (dpi) following infection with the PRVABC59 strain at both MOIs in both SH-SY5Y (**Figure 1A and B**) and IMR-32 (**Figure 1C and D**) tumorspheres. In contrast, BeH819015 only caused a significant reduction in SH-SY5Y tumorspheres at an MOI of 10. These findings confirm the strong oncolytic effect of the PRVABC59 strain in these experimental systems, which was therefore selected for further investigation. The viability of SH-SY5Y and IMR-32 tumorspheres infected with the PRVABC59 strain at an MOI of 2 or 10 is shown in **Figure 1E and F**. At an MOI of 10, viability dropped to nearly zero by 4 dpi in almost all tumorsphere replicates. However, not all replicates infected at an MOI of 2 exhibited reduced viability even at 6 dpi. To compare viability differences between MOIs, tumorspheres with less or more than 1% viability were counted and replotted (**Figure S1E and F**). This analysis confirmed that a greater proportion of tumorspheres infected at an MOI of 10 had a viability below 1% compared with those infected at an MOI of 2. Virus replication kinetics differed between the two MOIs as well. Virus titers in supernatants from tumorspheres infected at an MOI of 2 or 10 are shown in **Figure 1G and H**. The proportion of tumorspheres with no detectable virus were calculated and replotted (**Figure S1G and H**), showing that all tumorspheres became infected at an MOI of 10 even from the earliest time points tested, whereas at an MOI of 2 a proportion of tumorspheres failed to get infected even at later time points. These results indicate a binary outcome of infection at an MOI of 2: either the virus established a productive infection and reduced cell viability, or it failed to replicate and viability remained high. In contrast, infection at an MOI of 10 established robust infection in all samples, leading to a consistent drop in cell viability across all tumorspheres.

**Figure 1.**
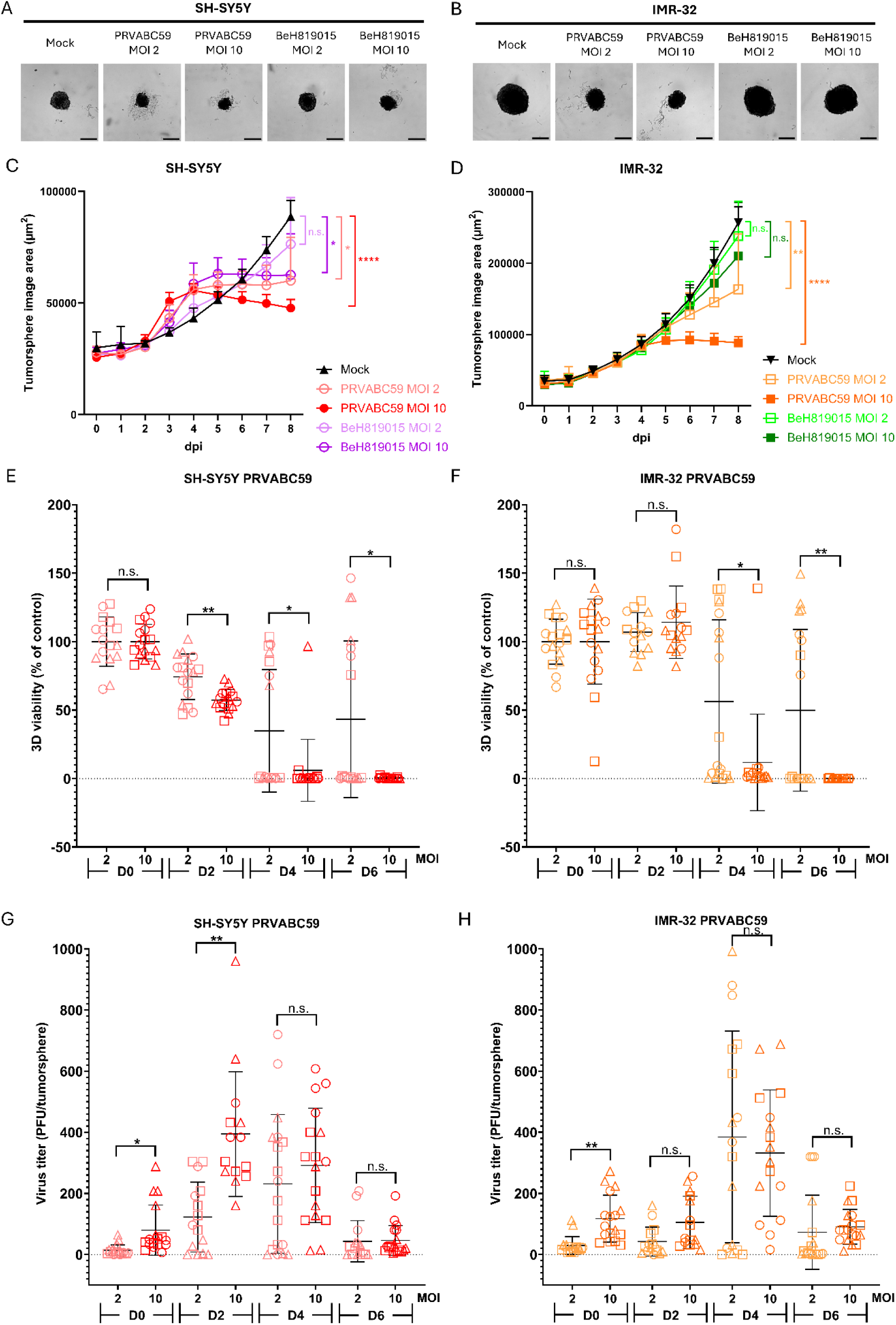
Oncolytic effects of ZIKV infection in neuroblastoma tumorspheres. (A-B) Representative images at 8 dpi for SH-SY5Y (A) and IMR-32 (B) control tumorspheres or tumorspheres following infection with the ZIKV PRVABC59 or BeH819015 strain at an MOI of 2 or 10. (C-D) Quantification of tumorsphere area for SH-SY5Y (C) and IMR-32 (D) tumorspheres under the same conditions as in (A-B). Scale bar = 300 μm. *N* = 3, *n* ≥ 2. Statistical significance indicators (asterisks) are shown for comparisons at 8 dpi. (E-F) 3D viability of individual SH-SY5Y (E) and IMR-32 (F) tumorspheres infected with the ZIKV PRVABC59 strain at an MOI of 2 or 10. *N* = 3, *n* ≥ 3. (G-H) ZIKV titers in supernatants from individual SH-SY5Y (G) and IMR-32 (H) tumorspheres infected with the ZIKV PRVABC59 strain at an MOI of 2 or 10. *N* = 3, *n* ≥ 3. Different symbol shapes represent different biological replicates. D0, D2, D4, and D6 represent 0, 2, 4, and 6 dpi, respectively. Data are presented as mean ± SD. Statistical significance was calculated using two-way ANOVA with Holm-Šidák multiple comparisons test. \**p* < 0.05; \*\**p* < 0.01; \*\*\*\**p* < 0.0001. n.s. indicates no significance.

### Transcriptome responses induced by ZIKV infection

To elucidate the mechanisms underlying the ZIKV-induced oncolytic effect, RNA-Seq was performed on SH-SY5Y and IMR-32 neuroblastoma tumorspheres with or without ZIKV infection at an MOI of 10, as this MOI consistently produced a strong and reproducible oncolytic effect. Among the top 30 significantly differentially expressed genes (DEGs), only three were downregulated in SH-SY5Y tumorspheres, while the remainder were upregulated (**Figure 2A, Table S1**). In ZIKV-infected IMR-32 tumorspheres, all 30 of the most significantly altered genes were upregulated (**Figure 2B, Table S2**). These top 30 regulated genes are predominantly associated with endoplasmic reticulum (ER) stress, the unfolded protein response (UPR), oxidative stress, amino acid transport, and immune responses in both cell models. The expression profiles of these genes are consistent with known cellular responses to ZIKV infection and replication^32–35^.

**Figure 2.**
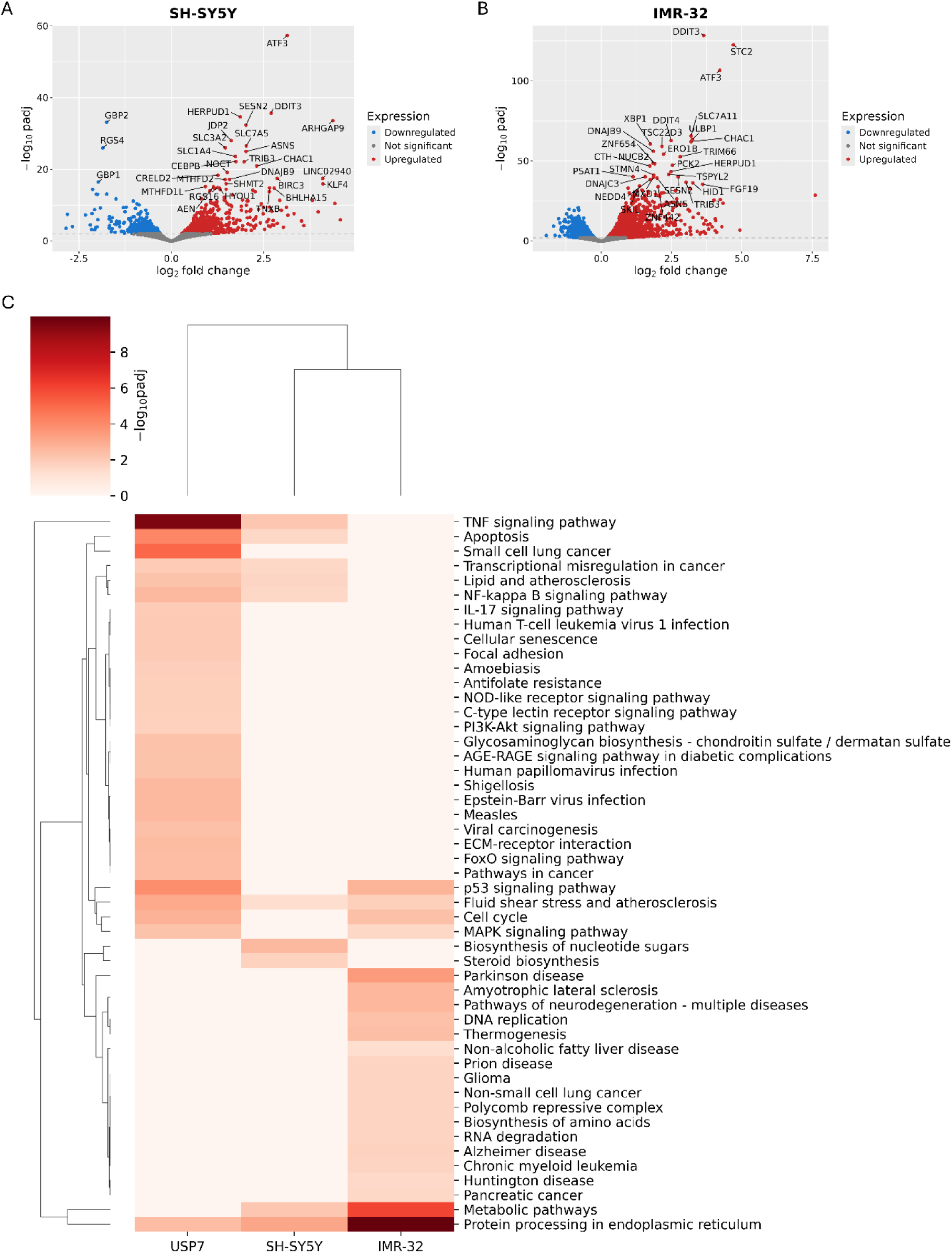
Transcriptome responses induced by ZIKV infection. (A-B) Volcano plots showing DEGs in SH-SY5Y (A) and IMR-32 (B) tumorspheres following infection with the ZIKV PRVABC59 strain at an MOI of 10. “padj” represents the adjusted *p*- value calculated using the Benjamini-Hochberg method. The dotted gray lines indicate the significance thresholds: -log_10_padj = 2 (equivalent to padj = 0.01). Red points represent significantly upregulated genes (log_2_ fold change > 0 and -log_10_padj ≥ 2), while blue points represent significantly downregulated genes (log_2_fold change < 0 and -log_10_padj ≥ 2). Gray points represent genes that are not significantly differentially expressed. The top 30 upregulated or downregulated genes, ranked by -log_10_padj, are labelled with their gene symbols. (C) Heatmap showing significant KEGG pathways regulated by ZIKV infection in USP7, SH-SY5Y and IMR-32 models.

To explore common pathways regulated by ZIKV infection in both neuroblastoma and pediatric brain tumor cells, a KEGG pathway analysis was performed (**Figure 2C**). Since ZIKV induces more pronounced cell death in USP7-AT/RT cells compared with other tested pediatric brain tumor cell lines^16^, published RNA-Seq data from ZIKV-infected USP7 cells^29^ were also reanalyzed and presented within the heatmap. The most significant common pathway across all three cell lines is “Protein processing in endoplasmic reticulum”, which aligns with the utilisation of the ER for ZIKV replication, protein folding, and assembly^36^. Among the two neuroblastoma cell lines, the altered pathways in SH-SY5Y cells were more similar to the pathways altered in pediatric brain tumor USP7 cells. The common significantly regulated pathways include “TNF signaling pathway”, “Apoptosis”, “Transcriptional misregulation in cancer”, “Lipid and atherosclerosis”, “NF-kappa B signaling pathway”, “Fluid shear stress and atherosclerosis”, and “Protein processing in endoplasmic reticulum”. Overall, these data identify cell line-specific transcriptional changes following ZIKV infection that do not necessarily correlate with the tumor origin of the cell line.

### ZIKV NS4A and NS5 suppress tumorsphere formation

Neuroblastoma SH-SY5Y and pediatric brain tumor USP7 cells were then transduced with each of the ZIKV non-structural proteins to investigate each protein’s role in mediating ZIKV’s oncolytic effects. SH-SY5Y cells were selected as they were more permissive to ZIKV infection compared to IMR-32 cells (**Figure 1**). Stable cell lines of SH-SY5Y or USP7 cells expressing GFP (control), NS1-GFP, NS2B-GFP, NS3-GFP, NS4A-GFP, NS4B-GFP, or NS5- GFP were established using lentivirus transduction. Immunofluorescence images (**Figure S2**) and western blot analysis (**Figure S3A and B**) confirmed successful transduction and target protein expression and demonstrated the subcellular localization of each of the non-structural proteins.

Tumorspheres from all established cell lines were then generated and cultured for 12 days. On day 12, GFP alone did not significantly affect the size or viability of SH-SY5Y or USP7 tumorspheres (**Figure 3**). Interestingly, NS1-GFP, NS3-GFP, NS4A-GFP, and NS4B-GFP significantly increased SH-SY5Y tumorsphere area and viability (**Figure 3A-C**), indicating that these non-structural proteins promote tumorsphere formation in neuroblastoma SH-SY5Y cells. In contrast, NS5-GFP significantly reduced both SH-SY5Y tumorsphere area and viability, suggesting an inhibitory effect of ZIKV NS5 on tumorsphere formation. In USP7 tumorspheres, only ZIKV NS4A-GFP significantly reduced both tumorsphere area and viability, while other non-structural proteins had no significant effect (**Figure 3D-F**). These findings suggest that ZIKV non-structural proteins exert distinct functions across different tumor cell types. NS5 and NS4A are the only non-structural proteins that suppressed tumorsphere formation in neuroblastoma SH-SY5Y and pediatric brain tumor USP7 cells under these experimental conditions, respectively, indicating potential roles for these proteins in ZIKV’s oncolytic effects.

**Figure 3.**
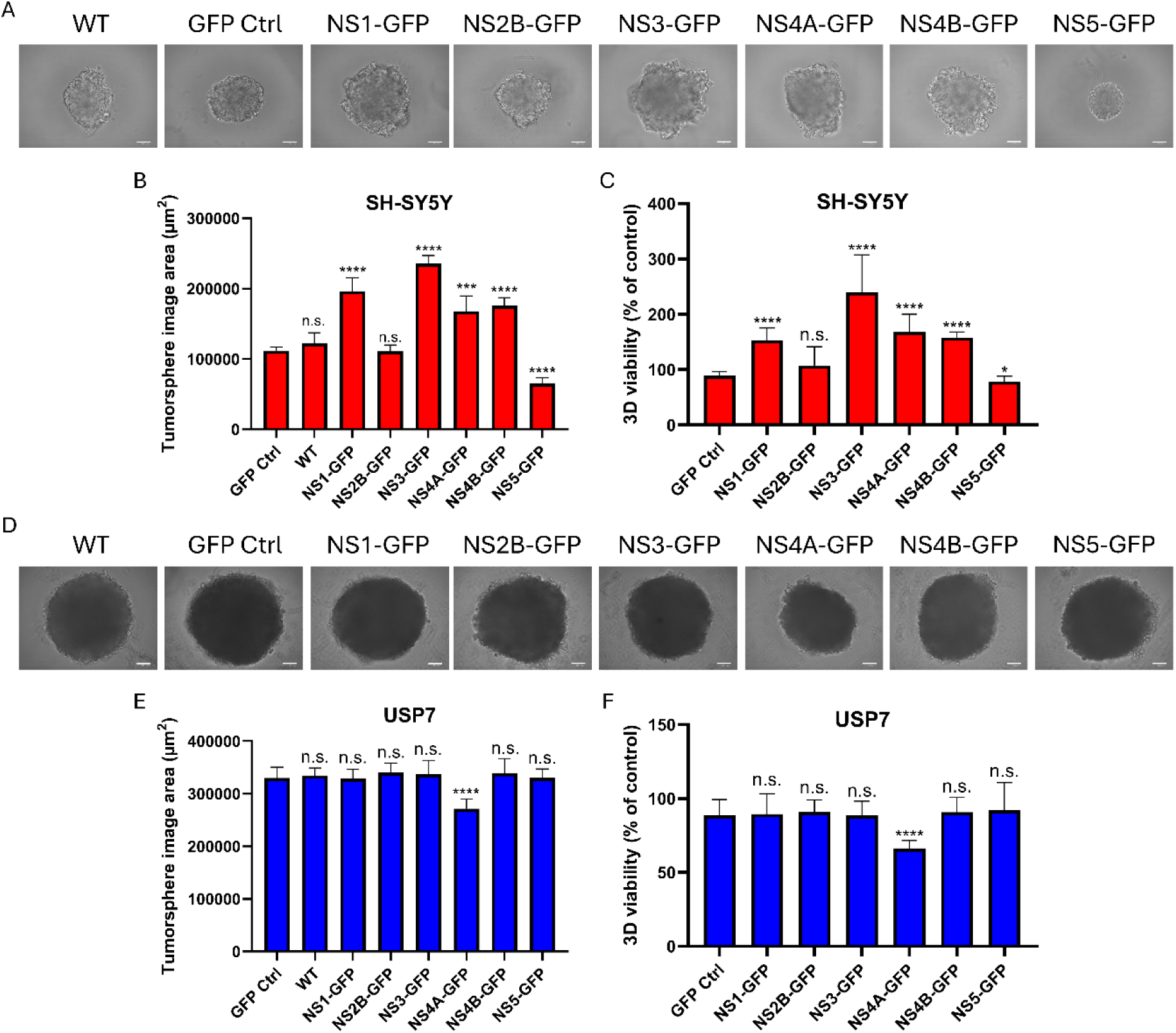
Effect of ZIKV non-structural proteins on tumorsphere formation. (A-C) Representative images (A), tumorsphere area (B), and 3D viability measurements (C) for SH-SY5Y tumorspheres formed by wild-type cells and cells transduced with GFP, NS1-GFP, NS2B-GFP, NS3-GFP, NS4A-GFP, NS4B-GFP, or NS5-GFP. (D-F) Representative images (D), tumorsphere area (E), and 3D viability measurements (F) for USP7 tumorspheres formed by the same cell lines. Tumorspheres were cultured for 12 days after seeding at 1 × 10³ cells per well. Scale bar = 100 µm. *N* = 3, *n* = 3. Data are presented as mean ± SD. Statistical significance was calculated using one-way ANOVA with Dunnett’s T3 multiple comparisons test, all compared to GFP. \**p* < 0.05; \*\*\**p* < 0.001; ****p < 0.0001. n.s. indicates no significance.

### Integrated transcriptomics and affinity proteomics analysis identifies integrin α3 as a key factor modulated by ZIKV

To further investigate the mechanisms by which ZIKV non-structural proteins regulate tumorsphere size and contribute to ZIKV’s oncolytic effects, affinity proteomics was performed to construct the ZIKV-host protein-protein interaction (PPI) networks for these tumor cells (**Figure 4A**). Mass spectrometry data confirmed that non-structural protein spectral counts were detected in all sample replicates (**Figure S3C**). A total of 345 protein targets and 516 interactions were identified in SH-SY5Y cells (**Figure 4B**; **Table S3**), while 269 protein targets and 344 interactions were identified in USP7 cells (**Figure 4C**; **Table S4**). In total, 79 common interaction partners were shared between the two cell lines (**Figure 5A**), suggesting mostly distinct ZIKV-host interaction profiles across tumor types. NS2B was found to interact with the most common proteins shared between the two cell lines (**Table S5**). To identify key interaction partners common to both tumor types that may contribute to ZIKV’s oncolytic effects, comparative analysis of RNA-Seq and NS2B affinity proteomics datasets was performed for NS2B. Overlapping Gene Ontology (GO) cellular component terms between the two datasets revealed that most overlapping terms had previously already been implicated in ZIKV replication, such as “Extracellular exosome”^37^, “Cytoplasm”^38^, “Endoplasmic reticulum”^36^, and “Golgi apparatus”^39^ (**Figure 5B**). Interestingly, “Focal adhesion”, which plays an important role in connecting tumor cells to the extracellular matrix and facilitating tumor invasion^40^, was enriched in both datasets. To further identify the key interaction partners within the “Focal adhesion” cellular component, NS2B’s interaction partners enriched in this term were plotted (**Figure 5C**). Among these partners, integrin α3 stood out because it interacts with NS2B in both tumor types and has previously been implicated in regulating cancer progression and metastasis^40–42^. In addition to integrin α3, integrin-linked kinase (gene name: *ILK*), which acts as a scaffold protein at focal adhesions to help link integrins to the actin cytoskeleton^43^, also interacts with NS2B in both tumor types. The co-detection of integrin α3 and integrin-linked kinase in the affinity proteomics data strengthens the evidence for an interaction between integrin α3 and NS2B, highlighting integrin α3’s potential role in the regulation of ZIKV’s oncolytic effects.

**Figure 4.**
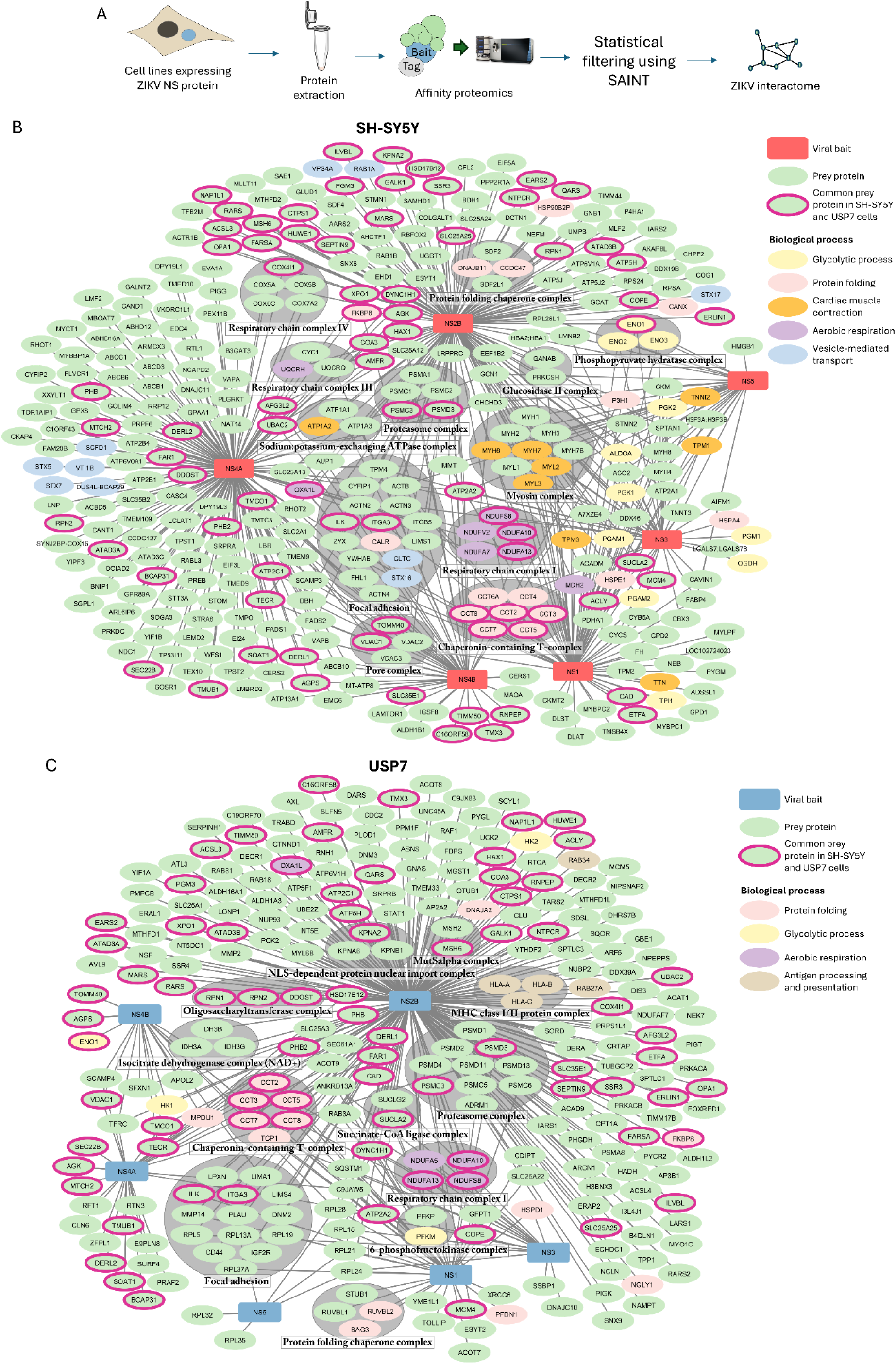
Summary of the ZIKV-host PPI networks. (A) Overall workflow of affinity proteomics approach used to define ZIKV-host PPIs. (B-C) High-confidence ZIKV-host PPIs identified in SH-SY5Y (B) and USP7 cells (C). Viral baits (red or blue rectangles), host prey proteins (green circles), common prey shared between the two tumor types (pink borders), and virus-host interactions (gray lines) are shown. Proteins involved in different biological processes are colored as indicated. Protein complexes are highlighted with gray clusters on the map.

**Figure 5.**
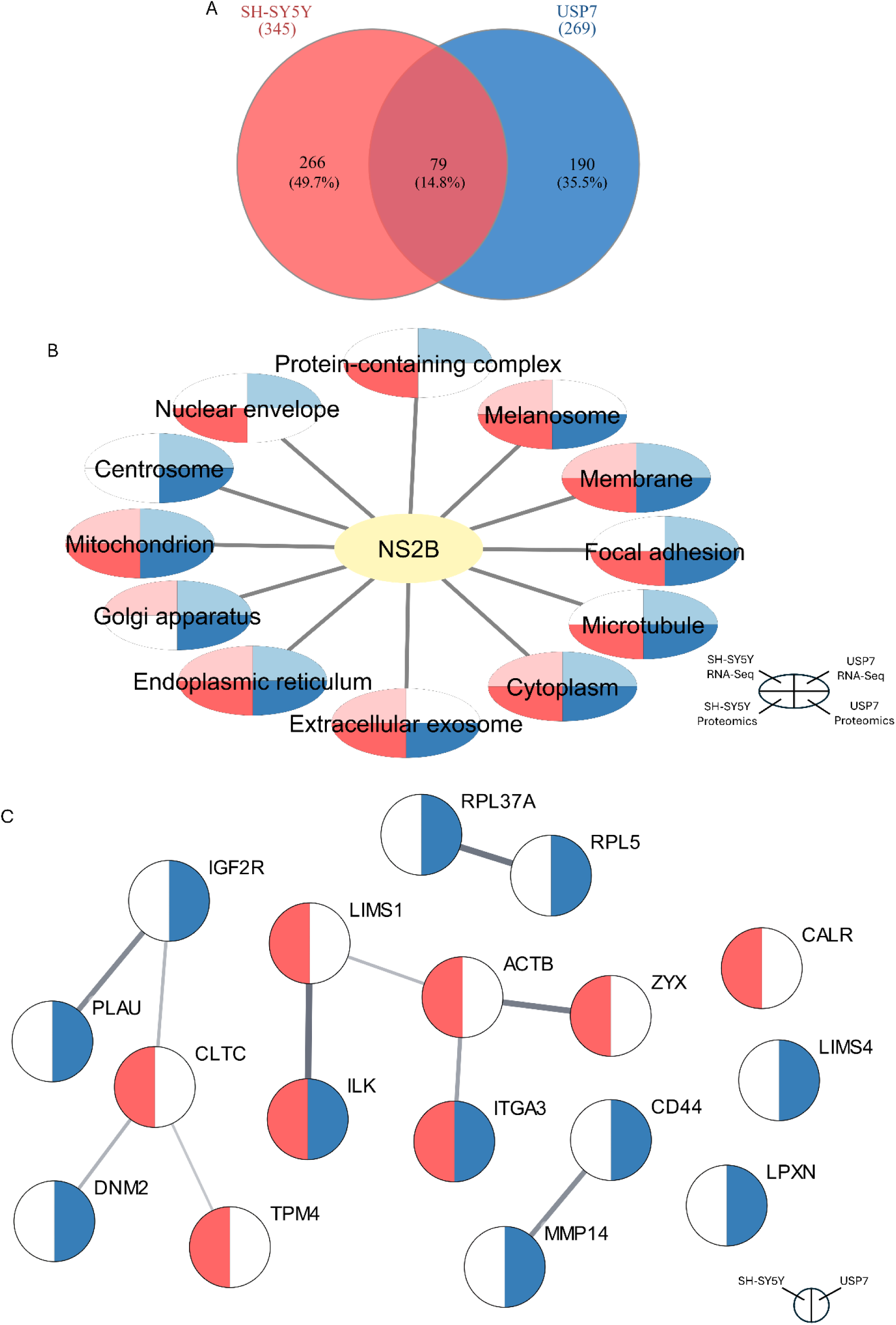
Comparative transcriptomics and affinity proteomics analysis. (A) Overlap of ZIKV-host PPI networks in SH-SY5Y and USP7 cells. (B) Overlap of GO cellular component terms between RNA-Seq and NS2B affinity proteomics datasets. (C) Interaction partners of NS2B enriched in focal adhesion GO cellular component term; gray lines represent physical interactions exported from STRING, and line thickness indicates the confidence level of each interaction.

### Integrin α3 interacts with NS2B and NS4A in both neuroblastoma SH-SY5Y and pediatric brain tumor USP7 cells

To investigate the potential roles of integrin α3 in regulating the changes in tumorsphere size induced by ZIKV non-structural proteins and ZIKV’s oncolytic effects, interactions between integrin α3 and ZIKV non-structural proteins were first validated by co-immunoprecipitation (Co-IP). All USP7 cell samples transduced with individual ZIKV non-structural proteins were subjected to immunoprecipitation using GFP-Trap beads, followed by western blotting. The results confirmed an interaction between integrin α3 and NS2B, and NS4A was also found to interact with integrin α3 in USP7 cells (**Figure S4**). Further validation showed that both NS2B and NS4A interacted with integrin α3 and integrin β1, which forms a heterodimer with integrin α3 to constitute the integrin α3β1 complex, in SH-SY5Y and USP7 cells (**Figure 6A-B**).

**Figure 6.**
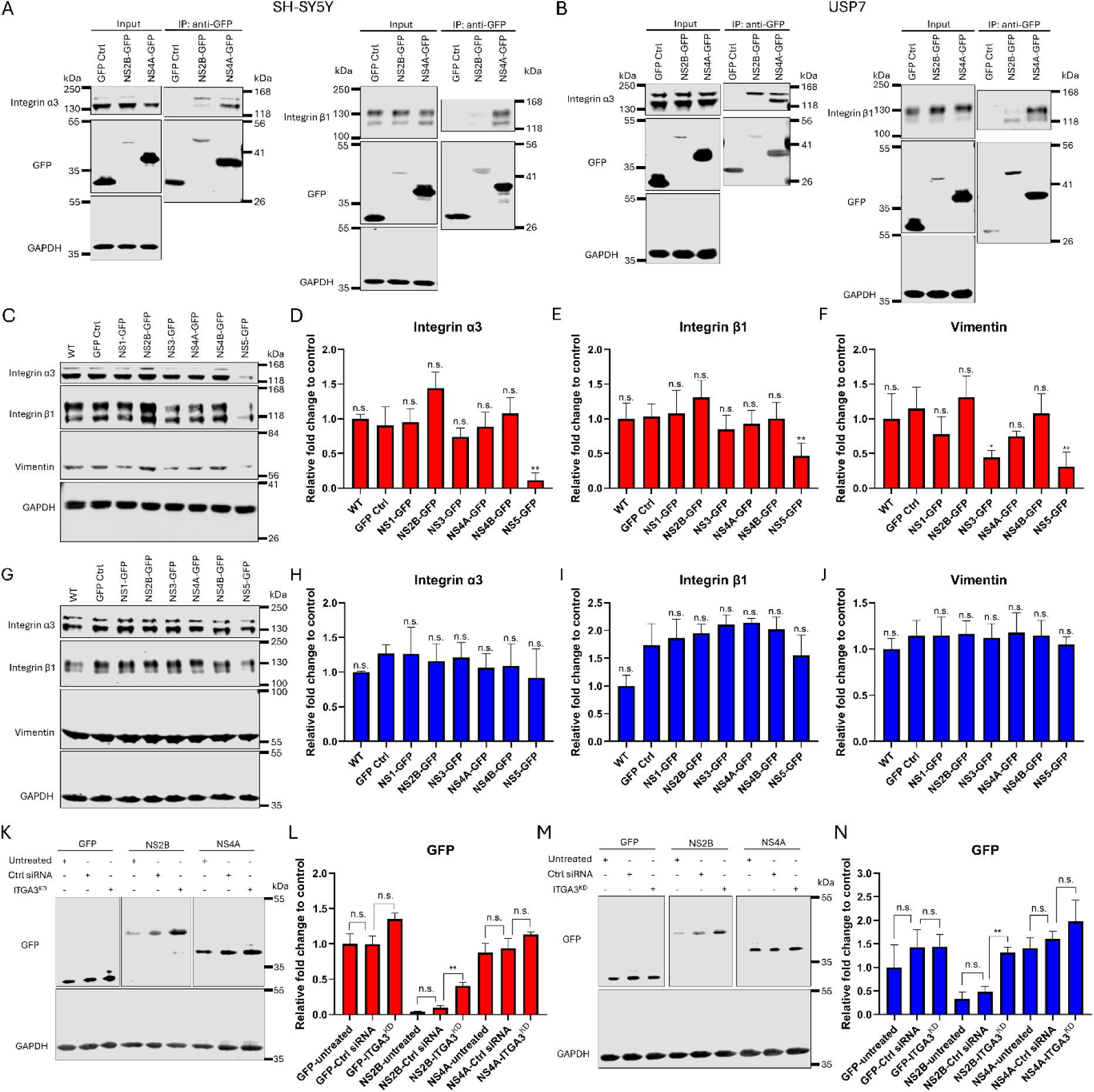
Integrin α3 interacts with NS2B and NS4A. (A-B) Co-IP analysis confirming that integrin α3 and integrin β1 interact with NS2B and NS4A in SH-SY5Y (A) and USP7 cells (B). (C-F) Western blot images (C) and relative fold changes of integrin α3 (D), integrin β1 (E), and vimentin (F) expression regulated by GFP or ZIKV non-structural proteins in SH-SY5Y cells. *N* = 5, *n* = 1. (G-J) Western blot images (G) and relative fold changes of integrin α3 (H), integrin β1 (I), and vimentin (J) expression regulated by the same proteins as in (C-F) in USP7 cells. *N* = 3, *n* = 1. (K-L) Western blot images showing GFP expression in SH-SY5Y cells transduced with GFP, NS2B-GFP or NS4A-GFP under the following conditions: untreated, RISC-Free control (Ctrl siRNA), and *ITGA3* knockdown (ITGA3^KD^) (K), with corresponding relative fold changes (L). *N* = 3, *n* = 1. (M-N) Western blot images (M) and corresponding relative fold changes (N) under the same conditions as in (K-L) for USP7 cells. *N* = 3, *n* = 1. All values are normalised to GAPDH control. Data are presented as mean ± SD. Statistical significance was calculated using one-way ANOVA with Dunnett’s T3 multiple comparisons test. \**p* < 0.05; \*\**p* < 0.01. n.s. indicates no significance.

To determine whether ZIKV non-structural proteins affect the expression levels of these integrins, western blotting was performed. In SH-SY5Y cells, only NS5 significantly reduced the expression of integrin α3 and integrin β1 (**Figure 6C-E**). This finding is consistent with previous research showing that ZIKV NS5 inhibits the migration and invasion of U87 glioma cells^44^. To further explore the role of NS5 in tumor metastasis, the expression level of the epithelial–mesenchymal transition marker vimentin, which has been shown to contribute to tumor invasion and metastasis^45–47^, was assessed. Consistent with previous results, NS5 significantly reduced vimentin expression in SH-SY5Y cells (**Figure 6C and F**). However, in USP7 cells, none of the non-structural proteins significantly affected the expression of integrin α3, integrin β1, or vimentin (**Figure 6G-J**). These results suggest that NS5 downregulates integrin α3, integrin β1, and vimentin specifically in SH-SY5Y cells, indicating a potential role for NS5 in inhibiting neuroblastoma invasion. In contrast, NS2B and NS4A do not affect the expression levels of integrin α3 or integrin β1 in either neuroblastoma or pediatric brain tumor cells, despite their physical interactions with these proteins.

To further investigate how integrin α3 influences the protein levels of its interacting partners NS2B and NS4A, *ITGA3* was silenced using siRNA, followed by western blot analysis. siRNA transfection consistently and effectively reduced integrin α3 protein expression in all GFP-expressing, NS2B-expressing, and NS4A-expressing SH-SY5Y and USP7 cells (**Figure S5A-D**). The expression levels of GFP and NS4A remained unchanged in both RISC-Free control and *ITGA3*-knockdown SH-SY5Y and USP7 cells (**Figure 6K-N**). However, NS2B protein levels were significantly increased in *ITGA3*-knockdown cells in both cell lines. No change in NS2B expression was observed in RISC-Free control cells, indicating that the observed effect is not due to off-target effects of the siRNA. These findings suggest that integrin α3 is involved in suppressing NS2B protein levels in both neuroblastoma SH-SY5Y and pediatric brain tumor USP7 cells.

### Knockdown of *ITGA3* rescues NS4A-induced reduction in USP7 tumorsphere size

To explore whether integrin α3 is involved in tumorsphere formation regulated by its interaction partners NS2B and NS4A, *ITGA3* was knocked down in wild-type, GFP-expressing, NS2B-expressing, and NS4A-expressing SH-SY5Y and USP7 cells, followed by tumorsphere formation assays. **Figure S5E-H** shows a consistent and effective reduction of integrin α3 protein expression levels in wild-type SH-SY5Y and USP7 cells. In SH-SY5Y cells, *ITGA3* knockdown significantly reduced tumorsphere size in both wild-type and GFP-expressing cells (**Figure 7A**). *ITGA3* knockdown also significantly reduced tumorsphere size of NS2B- and NS4A-expressing SH-SY5Y cells (**Figure 7B**). However, it is unclear whether these effects are related to interactions between integrin α3 and the non-structural proteins, as *ITGA3* knockdown also reduced tumorsphere size in control cells. In USP7 cells, *ITGA3* knockdown did not affect the tumorsphere size of either wild-type or GFP-expressing cells (**Figure 7C**). There was also no change in the tumorsphere size of NS2B-expressing USP7 cells following *ITGA3* knockdown (**Figure 7D**). However, *ITGA3* knockdown significantly increased the tumorsphere size of NS4A-expressing USP7 cells to the same level observed in untreated wild-type cells. Representative images of tumorspheres from control and NS4A-expressing USP7 cells under different treatments are shown in **Figure 7E and F**. The RISC-Free control had no such effects, indicating that the rescue of NS4A-induced tumorsphere size reduction is specifically due to *ITGA3* knockdown.

**Figure 7.**
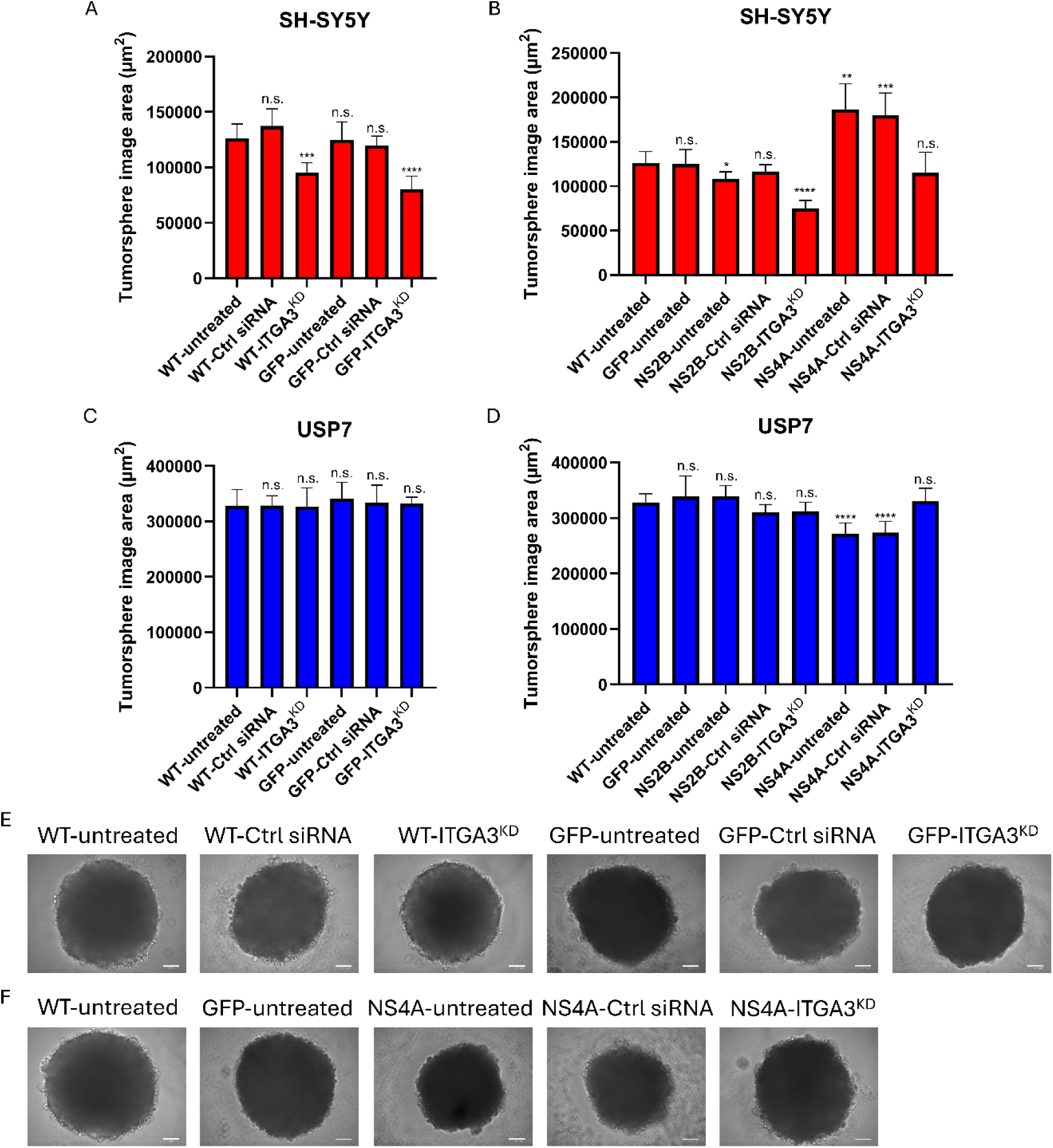
Effects of *ITGA3* knockdown on the regulation of tumorsphere formation by NS2B and NS4A. Tumorsphere formation assays were performed using wild-type cells and cells expressing GFP, NS2B, or NS4A under the following conditions: untreated, RISC-Free control (Ctrl siRNA), and *ITGA3* knockdown (ITGA3^KD^). Tumorspheres were cultured for 12 days after seeding at 1 × 10³ cells per well. (A-B) Quantification of tumorsphere area for wild-type and GFP-expressing (A), or NS2B- and NS4A-expressing (B) SH-SY5Y tumorspheres. (C-D) Quantification of tumorsphere area for wild-type and GFP-expressing (C), or NS2B- and NS4A-expressing (D) USP7 tumorspheres. (E-F) Representative images of corresponding wild-type and GFP-expressing (E), or NS4A-expressing (F) USP7 tumorspheres. *N* = 3, *n* = 3. Scale bar = 100 µm. Data are presented as mean ± SD. Statistical significance was calculated using one-way ANOVA with Dunnett’s T3 multiple comparisons test, all are compared to untreated wild-type cells. \**p* < 0.05; \*\**p* < 0.01; \*\*\**p* < 0.001; \*\*\*\**p* < 0.0001. n.s. indicates no significance.

### Integrin α3 plays different roles in ZIKV’s effects on SH-SY5Y and USP7 tumorsphere size

Given integrin α3’s interactions with NS2B and NS4A and its role in NS4A-mediated reduction of tumorsphere formation, we hypothesised that integrin α3 may be involved in ZIKV’s oncolytic effects. To determine whether ZIKV infection alters integrin α3 protein levels, we assessed its expression and found that ZIKV infection did not significantly change integrin α3 protein levels in either SH-SY5Y or USP7 cells (**Figure S6A-D**).

*ITGA3* was then knocked down (**Figure S6E-H**) followed by ZIKV infection to examine changes in cell viability, virus titer and tumorsphere size. *ITGA3* knockdown significantly reduced uninfected cell viability and virus titer in 2D-cultured SH-SY5Y cells (**Figure 8A and B**), but had no effect on viability in the context of ZIKV infection. In line with this finding, in SH-SY5Y tumorspheres *ITGA3* knockdown (confirmed in **Figure S6I-L**) significantly reduced tumorsphere size in non-infected tumorspheres but not during ZIKV infection (**Figure 8C**). Similar corresponding trends in the effects of *ITGA3* knockdown on viability were also observed in tumorspheres, although these differences were not significant (**Figure 8D**). Virus titers in tumorspheres showed no significant change following *ITGA3* knockdown as well (**Figure 8E**). Overall, these findings suggest that while *ITGA3* knockdown has a similar (though less pronounced) phenotypic effect to ZIKV infection in SH-SY5Y cell monolayers and tumorspheres, the effects of ZIKV infection and *ITGA3* are not additive, and integrin α3 is not required for the ability of ZIKV to shrink tumorsphere size. Nevertheless, ZIKV infection and *ITGA3* knockdown may reduce cell viability and tumorsphere size through shared pathways that are already maximally disrupted during ZIKV infection.

**Figure 8.**
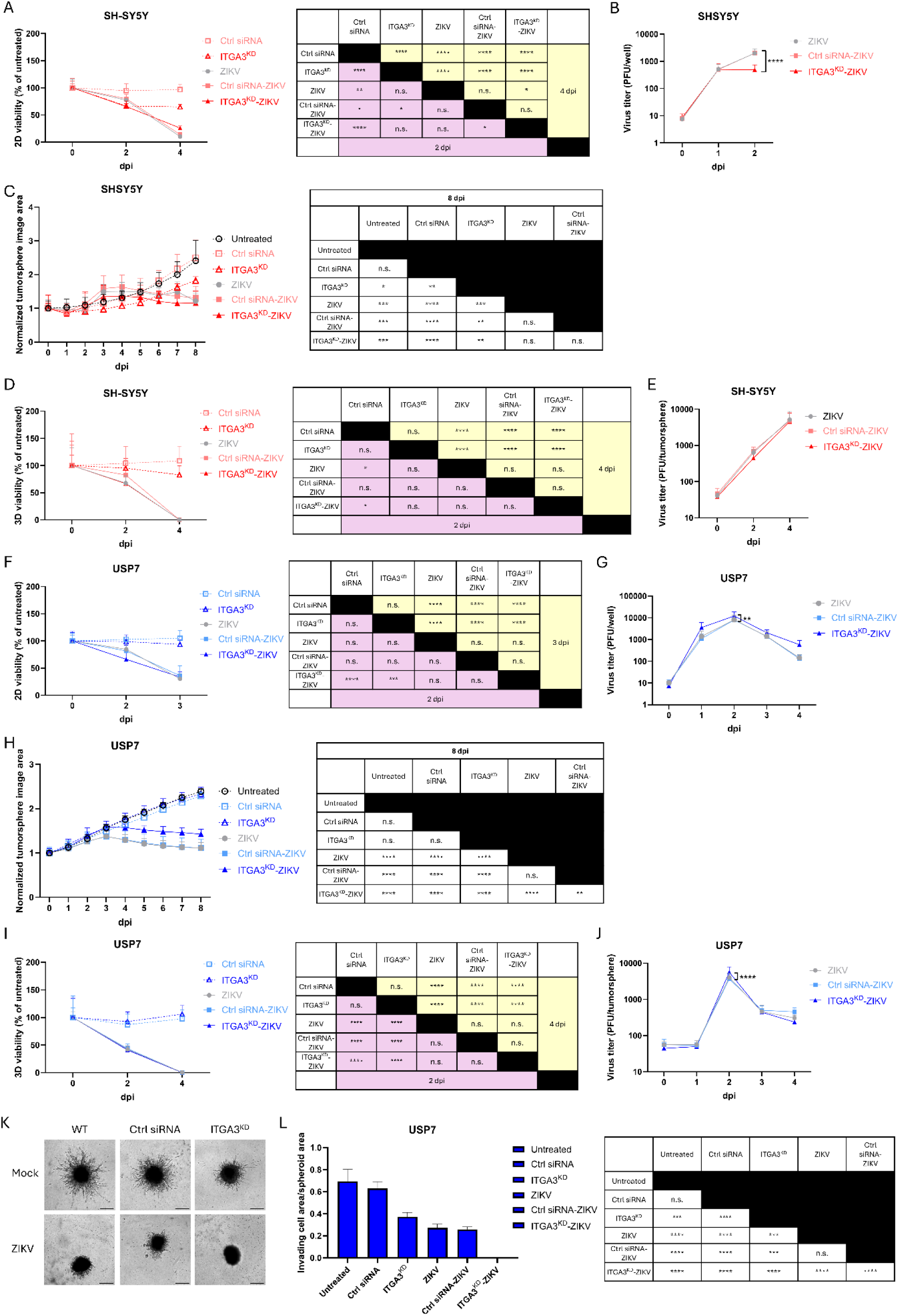
Effects of *ITGA3* knockdown on ZIKV oncolytic activity. (A-B) Cell viability (A) and extracellular viral titres (B) in SH-SY5Y cell 2D cultures transfected with RISC-Free control (Ctrl siRNA) or *ITGA3* siRNA (ITGA3^KD^) in the presence or absence of ZIKV infection. (C-E) Tumorsphere area normalized to d0 (C), 3D viability (D) and extracellular viral titres (E) in SH-SY5Y tumorspheres under the same treatments as in (A-B). (F-G) Cell viability (F) and extracellular viral titers (G) in USP7 cell 2D cultures under the same treatments as in (A-B). (H-J) Tumorsphere area normalized to d0 (H), 3D viability (I) and extracellular viral titres (J) in USP7 tumorspheres under the same treatments as in (A-B). (K-L) 3D invasion assay in USP7 tumorspheres treated as in (A), showing representative images (K) and the ratio of invading cell area to spheroid area (L) 4 days post-treatment. For (A-J): *N* = 3, *n* ≥ 3; statistical significance was calculated using two-way ANOVA with Holm-Šidák multiple comparisons test. For (K-L): *N* = 3, *n* ≥ 2; statistical significance was calculated using one-way ANOVA with Dunnett’s T3 multiple comparisons test. Data are presented as mean ± SD. Each table shows the statistical significance for the figure immediately to the left. For figures without a statistical significance table, asterisks in the figure indicate all significant comparisons between conditions, specifically highlighting differences between *ITGA3* knockdown in the presence of ZIKV infection and either ZIKV infection alone or ZIKV infection without knockdown. \**p* < 0.05; \*\**p* < 0.01; \*\*\**p* < 0.001; \*\*\*\**p* < 0.0001. n.s. indicates no significance.

In USP7 cells, *ITGA3* knockdown did not influence cell viability under either ZIKV-infected or non-infected conditions in 2D culture (**Figure 8F**). Integrin α3 also had no notable effects on virus titer, although a marginal statistically significant increase was observed at 2 dpi (**Figure 8G**). Interestingly, in ZIKV-infected USP7 tumorspheres, *ITGA3* knockdown, which had no effect on uninfected tumorspheres, reduced the capacity of ZIKV infection to shrink tumorsphere size (**Figure 8H**). Similarly to our observations in 2D culture, *ITGA3* knockdown did not influence 3D viability under either ZIKV-infected or non-infected conditions (**Figure 8I**), and also did not affect ZIKV titers except for a marginal statistically significant increase at 2 dpi (**Figure 8J**). Taken together, these results suggest that, in contrast to neuroblastoma SH-SY5Y cells, integrin α3 contributes to the ZIKV-induced reduction in pediatric brain tumor USP7 tumorsphere size. These effects are not mediated through impacts on cell viability or viral replication.

### *ITGA3* knockdown and ZIKV infection additively suppress 3D invasion in USP7 tumorspheres

Integrin α3 is involved in the focal adhesion pathway and mediates cell-matrix interactions. Given its important role in invasion and its interactions with ZIKV non-structural proteins, we investigated whether integrin α3 and ZIKV also functionally interact in a tumor invasion assay. Our transcriptomics data showed that the focal adhesion KEGG pathway was significantly regulated in pediatric brain tumor USP7 cells by ZIKV infection (**Figure 2C**), therefore, *ITGA3* knockdown was performed in USP7 tumorspheres. The results showed that ZIKV infection significantly inhibited USP7 tumorspheres invasion (**Figure 8K and L**), consistent with the transcriptomics data. *ITGA3* knockdown also significantly inhibited tumorsphere invasion, indicating its important role in this process. When *ITGA3* knockdown was combined with ZIKV infection, invasion was further suppressed compared with either treatment alone. Based on the area ratio of invading cells to spheroid (**Figure 8L**), *ITGA3* knockdown and ZIKV act additively to suppress 3D invasion of USP7 tumorspheres. This was also confirmed by the intensity ratio of invading cells to spheroid when the data were reanalyzed using an alternative method (**Figure S6M**). Therefore, ZIKV-induced suppression of invasion in pediatric brain tumor USP7 tumorspheres is unlikely to be mediated by integrin α3 and the two treatments likely function via independent pathways.

## Discussion

Although ZIKV exhibits neural tropism and promising oncolytic properties, its translation into clinical applications requires careful optimization^48^. Strategies such as genetic engineering and combination therapy need to be developed to enhance safety, improve tumor selectivity, and maximize therapeutic potency for effective and controlled treatment^48^. Therefore, understanding the mechanisms underlying ZIKV’s oncolytic effects is critical. In this study, we demonstrated that ZIKV infection reduces the size of both SH-SY5Y and IMR-32 neuroblastoma tumorspheres. We also performed a multi-omics analysis to uncover interactions between individual ZIKV non-structural proteins and host factors, as well as the mechanisms driving ZIKV’s oncolytic activity in neuroblastoma and AT/RT cell lines. We show that integrin α3 interacts with NS2B and NS4A, and demonstrate a contribution of integrin α3 to ZIKV-induced oncolytic effects in our USP7 AT/RT model. We also identified a synergistic function of *ITGA3* knockdown combined with ZIKV infection in inhibiting USP7 tumor invasion, providing new insights for genetic engineering and potential combination therapies. Interestingly, the functional interaction between integrin α3 and ZIKV differed between SH-SY5Y and USP7 models, with *ITGA3* knockdown phenocopying the effects of ZIKV infection in SH-SY5Y models, but integrin α3 does not contribute to ZIKV-induced impacts on SH-SY5Y cell viability and tumorsphere size.

Before exploring the mechanisms behind ZIKV’s oncolytic effects, we first examined whether ZIKV exhibits oncolytic activity in neuroblastoma 3D models. The PRVABC59 ZIKV strain was selected because our previous meta-analysis indicated it as the most promising candidate for neuroblastoma oncolytic virotherapy^49^. Our findings demonstrate its potent oncolytic effect in neuroblastoma 3D models generated from both SH-SY5Y and IMR-32 cell lines, especially compared to the BeH819015 strain, which only significantly reduced SH-SY5Y tumorspheres at an MOI of 10. Interestingly, infection with PRVABC59 at an MOI of 2 produced a binary outcome: either the virus established productive infection and reduced cell viability, or it failed to infect and viability remained high. In contrast, infection at an MOI of 10 consistently established robust infection across all samples, with a consistent decrease in cell viability. These findings indicate that, for future *in vivo* applications or neuroblastoma treatments, selecting the most effective strain and dose is critical.

We also investigated whether individual ZIKV non-structural proteins can exert oncolytic effects in isolation and found that NS4A and NS5 inhibit tumorsphere formation of AT/RT and neuroblastoma cell lines, respectively. NS4A has been reported to induce autophagy and apoptosis^50^, and to cause microcephaly by interacting with and inhibiting ANKLE2 as shown in HEK293T cells and *Drosophila* larvae^51^. ZIKV NS5 meanwhile has been reported to significantly inhibit proliferation of the human glioma cell line U87 and suppress tumorigenicity of mouse GL261 glioma cells *in vivo*.^44^ Our results show the oncolytic effects of NS4A and NS5 in pediatric neural tumors, consistent with these findings. The differences in non-structural protein function across tumor types suggest the importance of studying their effects in specific target tumors.

Furthermore, we show that integrin α3 interacts with NS2B and NS4A in cell lines of both neuroblastoma and AT/RT, and that it contributes to NS4A-induced oncolytic activity in AT/RT. However, it plays distinct roles in ZIKV-induced oncolytic activities in the two tumor types, which is another interesting finding that underscores the importance of studying the mechanisms in specific target tumors. Integrin α3 has been implicated in several viral infections. For example, it is an essential host factor for hepatitis E virus (HEV) infection^52^. In hepatocarcinoma cells, *ITGA3* knockout reduces HEV permissibility, as integrin α3 is critical for cellular attachment and entry of the non-enveloped HEV form^52^. Another study showed that the integrin α3β1 heterodimer serves as a cellular receptor for Kaposi’s sarcoma-associated herpesvirus entry into Chinese hamster ovary cells^53^. Here, we show for the first time that integrin α3 is also involved in the oncolytic effects induced by ZIKV and its protein NS4A. Knockdown of *ITGA3* did not significantly affect cell viability in either neuroblastoma or AT/RT cells in 2D or 3D culture models. Although some changes in viral titers were observed following *ITGA3* knockdown, these differences were not notable. These findings indicate that integrin α3 does not play a major role in ZIKV infection and that its roles in ZIKV-induced reduction in tumorsphere size is independent of alterations in cell viability or viral replication. In pediatric brain tumor cells, NS4A may act as a key effector of ZIKV’s oncolytic activity, with integrin α3 functioning as a downstream effector of NS4A. However, although integrin α3 interacts with NS4A, *ITGA3* knockdown does not affect NS4A protein levels, and altered NS4A expression does not change integrin α3 protein abundance.

Moreover, there is currently no evidence in the literature suggesting that integrin α3 plays a role in ANKLE2-related pathways, and our affinity proteomics data showed no interaction between NS4A and ANKLE2 in either USP7 or SH-SY5Y cells. These findings indicate that the interaction between NS4A and integrin α3 is not regulatory at the level of protein expression but may instead influence functional activities or downstream signaling pathways, and NS4A-induced tumorsphere size reduction may not be related to ANKLE2 regulation.

Integrin α3 forms a heterodimer with integrin β1, which interacts with various extracellular matrix proteins^54^, suggesting the potential role of integrin α3 in cancer metastasis. Previous studies have shown that integrin α3 facilitates glioma cell migration and invasion^42^, and our results show that integrin α3 contributes to 3D invasion in pediatric brain tumor AT/RT. Moreover, ZIKV infection has been reported to reduce metastasis in AT/RT tumor-bearing mouse models^16^, and our findings show that ZIKV infection reduces 3D invasion in AT/RT tumorspheres. These observations support our transcriptomic and affinity proteomic data, which revealed that ZIKV infection significantly regulates the focal adhesion KEGG pathway (**Figure 2C**) and that its non-structural protein NS2B interacts with host factors involved in focal adhesion cellular component term in AT/RT (**Figure 5B and C**). Interestingly, the combination of ZIKV infection with *ITGA3* knockdown additively suppresses 3D invasion in AT/RT tumorspheres. This indicates that ZIKV-induced reduction in invasion is unlikely to be mediated through integrin α3-mediated signaling pathways and suggests a potential route to combination therapy.

Taken together, our findings provide new insights for the improvement of ZIKV’s oncolytic effects. We demonstrate the oncolytic roles of ZIKV proteins NS4A and NS5 in pediatric neural tumors and highlight the importance of integrin α3 in ZIKV- and NS4A-induced oncolytic activities. These results suggest that it may be possible to genetically engineer these non-structural proteins to enhance ZIKV’s antitumor activity, and indicate that ZIKV’s oncolytic potency may vary depending on the *ITGA3* expression level in patients’ tumors. Additionally, our findings indicate a synergistic effect of ZIKV infection and *ITGA3* knockdown in reducing invasion within our AT/RT tumor model, suggesting the potential of combination strategies involving ZIKV infection and integrin-targeting approaches to inhibit tumor metastasis. However, inhibiting integrin α3 may also reduce ZIKV’s oncolytic effects, particularly its effect on tumor size reduction. Therefore, combinatorial treatment strategies involving integrin α3 need to be carefully balanced. Key factors including the sequence of administration and the temporal scheduling of treatments should be evaluated. Further work is needed to explore the potential benefits of exploiting the functional interactions between ZIKV infection and integrin α3 for oncotherapeutics.

## Materials and methods

### Cell culture

Neuroblastoma SH-SY5Y (CRL-2266) and IMR-32 (CCL-127) cells were purchased from the American Type Culture Collection (ATCC, Manassas, USA). USP7 AT/RT cells were provided and established in-house^16^ at the University of São Paulo. HEK293T cells were provided by Nullin Divecha (University of Southampton, Southampton, UK), and Vero cells were provided by Trevor Sweeney (The Pirbright Institute, Pirbright, UK). All cell lines were cultured in medium supplemented with 10% FBS (Thermo Fisher, Waltham, USA, cat. no. 10270106) and 1% penicillin/streptomycin (Thermo Fisher, cat. no. 15140122). The media used for these cells were as follows: SH-SY5Y, HEK293T and Vero: Dulbecco’s modified Eagle medium (DMEM) (Thermo Fisher, cat. no. 41966029); IMR-32: Roswell Park Memorial Institute (RPMI) Medium 1640 with GlutaMAX™ (Thermo Fisher, cat. no. 61870010); USP7: high-glucose DMEM with GlutaMAX™ (Thermo Fisher, cat. no. 61965026). All cell lines were cultured at 37 ℃ in a humidified atmosphere containing 5% CO_2_ and were confirmed to be mycoplasma free.

### siRNA transfection

For each well of a 6-well plate, siRNAs (QIAGEN, Santa Clarita, CA, USA, cat. no. SI00034174) or RISC-Free control siRNAs (Horizon, Cambridge, UK, cat. no. D-001220-01- 05) were transfected into 4 × 10^5^ SH-SY5Y cells or 3 × 10^5^ USP7 cells using Lipofectamine™ RNAiMAX Transfection Reagent (Thermo Fisher, cat. no. 13778075) at a final concentration of 12.5 nM siRNA per well. Untreated control cells were cultured in complete growth medium. All wells were gently mixed by rocking the plate back and forth and incubated at 37 ℃ in a CO_2_ incubator for 42 h before harvesting for downstream assays.

### Tumorsphere formation assay

Cells were seeded at 1×10^3^ cells per well in 200 µL of medium in 96-well clear ultra-low attachment microplates (Thermo Fisher, cat. no. 10023683) to generate one tumorsphere per well. Images of the tumorspheres were taken daily using an Incucyte® S3 Live-Cell Analysis System (Sartorius, Göttingen, Germany) (Figure 1A and C) and a ZOE Fluorescent Cell Imager (Bio Rad, Hercules, California, USA) (Figure 3A and D; Figure 7E and F), and the area of the tumorspheres was measured using Image J software (version 1.54f; National Institutes of Health, USA; http://imagej.org).

### Virus techniques

The ZIKV BeH819015 strain from Brazil was rescued from the infectious cDNA (icDNA) clone icBeH819015 ^55^, which was provided by Trevor Sweeney (The Pirbright Institute). The ZIKV PRVABC59 (KU501215.1) strain from Puerto Rico was a kind gift from Matt Evans (Icahn School of Medicine at Mount Sinai, New York, USA). Both viral stocks were produced by low MOI infection in Vero cells. For plaque assay titration, Vero cells were seeded in a 48-well plate at 1.5 x 10^5^ cells per well and incubated overnight prior to infection. Ten-fold serial dilutions of viral stocks/samples in serum-free medium was added to each well after the medium was removed, and incubated for 1.5 h at 37 ℃ with shaking every 15-20 min. Vero cells were washed with PBS, and an overlay containing 1% agarose (Invitrogen, Thermo Fisher, cat. no. 16520100) mixed with medium supplemented with 2.5% FBS was added. The Vero cells were then incubated at 37 ℃ in a humidified atmosphere supplemented with 5% CO_2_ for 3 days before fixation in 10% formalin (Sigma-Aldrich, Merck, St. Louis, USA, cat. no. HT501128). Cells were stained with 0.1% (w/v) crystal violet (Sigma-Aldrich, cat. no. 61135) for counting.

For infection of 2D cell cultures, cell monolayers were seeded in 96-well plates at 1×10^4^ cells per well in 200 µL of medium one day prior to infection. Cells were infected with ZIKV at an MOI of 2 in serum-free medium for 1 h at 37 ℃, then the inoculum was removed and replaced with complete culture medium. Supernatant and cell lysates were collected at the time points as indicated in figures. 3D tumorsphere infection was performed one day after tumorspheres formation for SH-SY5Y and IMR-32 cells, and three days post-tumorsphere formation for USP7 cells. All tumorspheres were infected with ZIKV at an MOI of 2 or 10 for 2 h at 37 ℃. The virus inoculum was removed, tumorspheres were washed with PBS, and then 200 µL of complete culture medium was added to each well. Supernatant and tumorspheres were collected at time points as indicated in figures

### Lentivirus transduction

ZIKV non-structural protein plasmid DNA with GFP-Spark tag was purchased from Sino Biological company (Paoli, PA, USA; cat. no. LVCV-35 for vector with GFP-Spark tag only; cat. no. VG40544-ACGLN for NS1-GFP; cat. no. VG40612-ACGLN for NS2B-GFP; cat. no. VG40545-ACGLN for NS3-GFP; cat. no. VG40613-ACGLN for NS4A-GFP; cat. no. VG40614-ACGLN for NS4B-GFP; cat. no. VG40584-ACGLN for NS5-GFP). Bacterial transformation was performed using One Shot™ TOP10 Chemically Competent *E. coli* (Invitrogen, Thermo Fisher, cat. no. C404003) according to the manufacturer’s instruction. Plasmid DNA was isolated using QIAGEN plasmid kits. Lentiviral particles were generated in HEK293T cells using a third-generation packaging system. Briefly, HEK293T cells (1 × 10⁶ cells per well) were seeded in 6-well plates with complete culture media. Viral plasmid and packaging plasmids (Gag-Pol and VSV-G plasmids) purchased from Addgene (Watertown, MA, USA) were mixed at a 4:2:1 ratio in serum-free DMEM and transfected into cells using polyethylenimine (PEI; Merck, Burlington, MA, USA, cat. no. 408719). The culture medium was replaced with fresh complete culture media following overnight incubation at 37 ℃ and viral supernatants were collected at 24, 48 and 72 h post-transfection, clarified by centrifugation at 3000 × g for 15 min, filtered through a 0.45 µm syringe filter, aliquoted, and stored at -80 ℃ until use.

Target cells were plated in 1.5 mL of medium in a 6-well plate and incubated at 37 ℃ prior to infection. Lentivirus stock was mixed with 5 µg/mL polybrene (Sigma-Aldrich, cat. no. 107689-10G) for infection and cells were incubated overnight, after which the medium was replaced with complete culture medium. After an additional 48 h, the cells were passaged and collected into a flask. Cells expressing GFP were selected using the Bigfoot cell sorter (Thermo Fisher) and bulked up to generate stable cell lines expressing GFP negative control, NS1-GFP, NS2B-GFP, NS3-GFP, NS4A-GFP, NS4B-GFP, or NS5-GFP.

### Viability assay

Viability of 2D cell monolayers and 3D tumorspheres was measured using the CellTiter-Glo® Luminescent Cell Viability Assay kit (Promega, Madison, WI, USA, cat. no. G7570) or the CellTiter-Glo® 3D Cell Viability Assay (Promega, cat. no. G9683), following the manufacturer’s instructions. Luminescence was measured using the GloMax®-Multi + Detection System (Promega). Cell (3D) viability was calculated using the following equation:

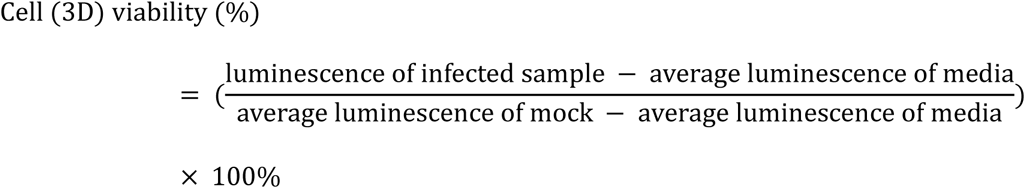

All viability data were normalized to the average data at 0 dpi.

### 3D invasion assay

Tumorspheres were formed and infected as described above prior to the addition of invasion matrix (Bio-Techne, Minneapolis, MN, USA, cat. no. 3500-096-03) on ice. Plates were centrifuged at 300 × g at 4 ℃ for 5 min and incubated at 37 ℃ for 1 h. Plates were incubated at 37 ℃ in a humidified atmosphere with 5% CO_2_ for 4 days prior to imaging. Images were captured using the Incucyte® S3 Live-Cell Analysis System (Sartorius). Spheroid area and invading cell area were quantified using the system. Mean intensity of spheroids and invading cells was quantified using image J software (version 1.54f; National Institutes of Health, USA; http://imagej.org).

### Confocal microscopy

Wild-type or lentivirus-transduced SH-SY5Y or USP7 cells were seeded on coverslips (0.16-0.19 mm thickness) overnight to allow attachment. Coverslips were then washed with PBS and fixed with 4% PFA (diluted in PBS) at room temperature for 15 min. After washing with PBS, 0.1% Triton X-100 (Sigma Aldrich, cat. no. T8787) (diluted in PBS) was added for permeabilization at room temperature for 15 min. After washing again with PBS, coverslips were stained with DAPI (Invitrogen, Thermo Fisher, cat. no. D1306, 1:2000) at room temperature for 10 min, protected from light, then washed three times with PBS for 10 min in the dark. Rhodamine phalloidin (Invitrogen, Thermo Fisher, cat. no. R415, 1:1000) was added to stain F-actin at room temperature for 1 h, protected from light. Coverslips were then washed three times with PBS for 10 min in the dark, mounted with Fluoromount-G™ mounting medium (Invitrogen, Thermo Fisher, cat. no. 00-4958-02), and stored in the dark at 4 ℃ after dry until imaging. Images were captured using a Leica SP8 confocal microscope (Leica Microsystems, Wetzlar, Germany).

### Co-immunoprecipitation

Wild-type or lentivirus-transduced SH-SY5Y or USP7 cells were seeded in 100 mm dishes at a density of 1 x 10^7^ cells per dish. Cells were harvested by scraping into ice-cold PBS and centrifuged at 500 × g for 3 min at 4 ℃. The supernatant was discarded, and the cell pellet was washed twice more by gentle resuspension in ice-cold PBS. Cell lysis and immunoprecipitation were conducted using the ChromoTek GFP-Trap® Agarose kit (Proteintech, Manchester, UK, cat. no. gta) according to the manufacturer’s instruction.

### Western blot

Wild-type or lentivirus-transduced SH-SY5Y or USP7 cells were seeded in 6-well plates at a density of 6 x 10^5^ cells per well. All samples were kept on ice throughout the procedure. Cells were washed twice with PBS, and 120 μl of ice-cold RIPA lysis buffer (Thermo Fisher, cat. no. 89900), supplemented with 1× Halt™ Protease Inhibitor Cocktail and 1× EDTA (Thermo Fisher, cat. no. 87786), was added to each well. Plates were kept on ice for 5 min with intermittent pipetting, followed by sonication for 5 min. Lysates were then centrifuged at 14,000 × g for 15 min at 4 ℃, and the supernatant was collected and mixed thoroughly. Protein concentration was determined using the Bradford assay. LDS sample buffer (Thermo Fisher, cat. no. NP0007), supplemented with 5% 2-mercaptoethanol (Sigma Aldrich, cat. no. M6250), was added to lysates and samples were boiled at 100 ℃ for 5 min. Samples were subjected to SDS-PAGE and then transferred on to 0.45 µm nitrocellulose membranes (Thermo Fisher, cat. no. 88018).

Membranes were stained with Ponceau S solution (Sigma Aldrich, cat. no. P7170-1L), washed with Tris-buffered saline with Tween 20 (TBST), and blocked with 5% milk powder (Marvel, Premier Foods, UK) for 1 h. Membranes were then incubated overnight at 4 ℃ with the following primary antibodies: GFP (Cell Signaling Technology, Danvers, MA, USA, cat. no. 2956, 1:1000), integrin α3 (Proteintech, cat. no. 66070-1-Ig, 1:1000), integrin β1 (Proteintech, cat. no. 12594-1-AP, 1:1000), vimentin (Cell Signaling Technology, cat. no. 5741, 1:1000), and GAPDH (Cell Signaling Technology, cat. no. 2118, 1:1000). After three 10-min washes with TBST, membranes were incubated for 1 h at room temperature with one of the following secondary antibodies: IRDye® 680LT Goat anti-Rabbit IgG (LI-COR Biosciences, Lincoln, NE, USA, cat. no. 926-68021, 1:5000), IRDye® 800CW Goat anti-Rabbit IgG (LI-COR Biosciences, cat. no. 926-32211, 1:5000), IRDye® 680LT Goat anti-Mouse IgG (LI-COR Biosciences, cat. no. 926-68020, 1:5000), and IRDye® 800CW Goat anti-Mouse IgG (LI-COR Biosciences, cat. no. 926-32210, 1:5000). Membranes were washed again three times with TBST and imaged using the LI-COR Odyssey Western Blot imager (LI-COR Biosciences). Quantification was performed using Image J software (version 1.54f; National Institutes of Health, USA; http://imagej.org).

### RNA-Seq and data analysis

Tumorsphere samples were collected into TRIzol (Thermo Fisher, cat. no. 15596018), and total RNA was extracted according to the manufacturer’s instruction. mRNA sequencing was conducted by Novogene company (Beijing, China). Raw RNA-Seq reads were quality-checked using FastQC (version 0.11.9) and trimmed with Trim Galore (version 0.6.10). Post-trim quality was checked again with FastQC. Trimmed reads were then aligned against the *Homo sapiens* GRCh38 reference genome using HISAT2 (version 2.2.1). Samtools (version 1.16.1) and Subread (featureCounts, version 2.0.6) were used to generate count tables. DESeq2 (version 1.42.1) was used for differential gene expression analysis. Significance was adjusted using the Benjamini-Hochberg method. Official gene symbols of significant DEGs (padj < 0.05) were submitted to the Database for Annotation, Visualisation and Integration Discovery (DAVID) online bioinformatics resources (https://david.ncifcrf.gov/) for KEGG pathway analysis.

### Affinity purification-mass spectrometry (AP-MS) and data analysis

SH-SY5Y or USP7 cells expressing GFP alone or GFP-tagged non-structural protein were collected, washed with PBS, and lysed as described in the Co-IP method. Immunoprecipitation, washing, and subsequent on-bead digestion were performed as previously described^56^ using GFP-Trap Agarose beads (Chromotex, Proteintech, cat. no. gtma). Reversed-phase nano-liquid chromatography-tandem mass spectrometry was performed as previously described^56^ using a nano-Ultimate 3000 liquid chromatography system coupled to a Lumos Fusion mass spectrometer (both Thermo Fisher).

Raw mass spectrometry data were processed using MSFragger within the FragPipe platform and searched against the UniProtKB human proteome (proteome ID: UP000005640) spiked with GFP and ZIKV non-structural protein sequences. Quantitative outputs from MSFragger were analyzed using the LIMMA package (https://github.com/wasimaftab/LIMMA-pipeline-proteomics)^57^ in R. Protein lists after the analysis were then submitted to Significance Analysis of Interactome (SAINT) and Contaminant Repository for Affinity Purification (CRAPome) online analysis (http://www.crapome.org/). Common nonspecific GFP-tag interactors were removed using a CRAPome frequency cutoff > 60%. SAINT probability (SP) score = 1 for SH-SY5Y and SP score ≥ 0.85 for USP7 were set as thresholds for high-confidence interactions. Official gene symbols of high-confidence interacting proteins were then submitted to the DAVID for GO analysis. High-confidence interacting proteins were mapped using Cytoscape software (version 3.9.1).

### Statistical analysis

Statistical analyses for plaque assay titration and functional assays were conducted in GraphPad Prism (version 10.3.1; GraphPad Software, San Diego, CA, USA) using one-way ANOVA with Dunnett’s T3 multiple comparisons test, two-way ANOVA with Holm-Šidák multiple comparisons test, or Student’s *t*-test. padj < 0.05 for RNA-Seq analysis and *p* < 0.05 for all other analyses were considered statistically significant. “padj” represents the adjusted *p*-value calculated using the Benjamini-Hochberg method.

## Data availability

RNA-Seq data have been deposited in the NCBI Gene Expression Omnibus (GEO) under accession number GSE313773. The mass spectrometry proteomics data have been deposited to the ProteomeXchange Consortium via the PRIDE^58^ partner repository with the dataset identifier PXD077091.

## Acknowledgements

We thank Jinhui Gao for technical support with the experiments. We also thank Dr Alexander von Kriegsheim’s research group at the University of Edinburgh for performing the mass spectrometry work. This work was supported by grants from Neuroblastoma UK, The Rosetrees Trust, The Little Princess Trust and MRC MR/S01411X/1. Y.S. was supported by the China Scholarship Council (grant no. 202106350031). Work performed at The Pirbright Institute was supported by BBSRC grants BBS/E/PI/230002B and BBS/E/PI/23NB0003.

## Author contributions

Y.S. and R.M.E. conceptualized the approaches used, with input from K.M., O.K.O., and Y.W. Experiments and data analysis were conducted by Y.S., with assistance from M.S., who prepared lentivirus stocks and performed lentivirus transduction in USP7 cells, as well as cell lysis for transduced USP7 cells before AP-MS. Y.S. wrote the original manuscript, and all authors reviewed and edited it.

