## Supplementary Figures for "Multi-omics analysis reveals integrin α3-dependent mechanisms of Zika virus oncolytic activity in pediatric neural tumors"

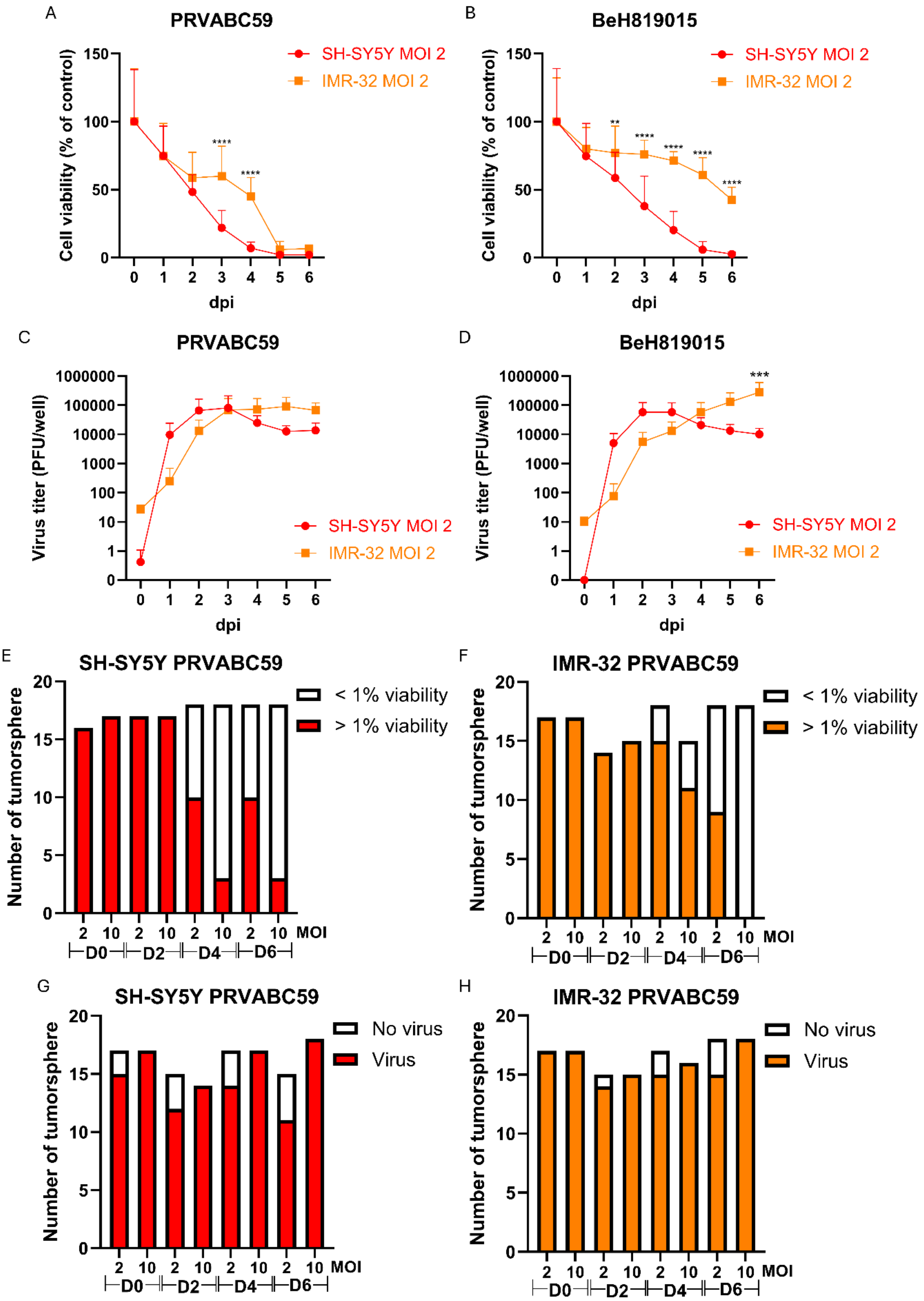

**Figure S1. Cell viability and viral titers in neuroblastoma models**

(A-B) Average cell viability of SH-SY5Y and IMR-32 cells infected with the ZIKV PRVABC59 (A) or BeH819015 (B) strains at an MOI of 2. $N \geq 2$, $n = 6$. (C-D) Average viral titers in supernatants from SH-SY5Y and IMR-32 cells infected with the ZIKV PRVABC59 (C) or BeH819015 (D) strain at an MOI of 2. For 0 dpi: $N \geq 1$, $n = 3$; for 1-6 dpi: $N \geq 2$, $n = 3$. (E-F) Data from Figure 1E and 1F replotted to show the proportion of PRVABC59-infected SH-SY5Y (E) and IMR-32 (F) tumorspheres with viability > 1% versus < 1%. $N = 3$, $n \geq 3$. (G-H) Data from Figure 1G and 1H replotted to show the proportion of PRVABC59-infected SH-SY5Y (G) and IMR-32 (H) tumorspheres with virus detected versus no virus detected in the supernatant. $N = 3$, $n \geq 3$. D0, D2, D4, and D6 represent 0, 2, 4, and 6 dpi, respectively. Data are presented as mean ± SD. Asterisks indicate significant differences between SH-SY5Y and IMR-32 cells. Statistical significance was calculated using two-way ANOVA with Holm-Šidák multiple comparisons test. **$p < 0.01$; ***$p < 0.001$; ****$p < 0.0001$. n.s. indicates no significance.

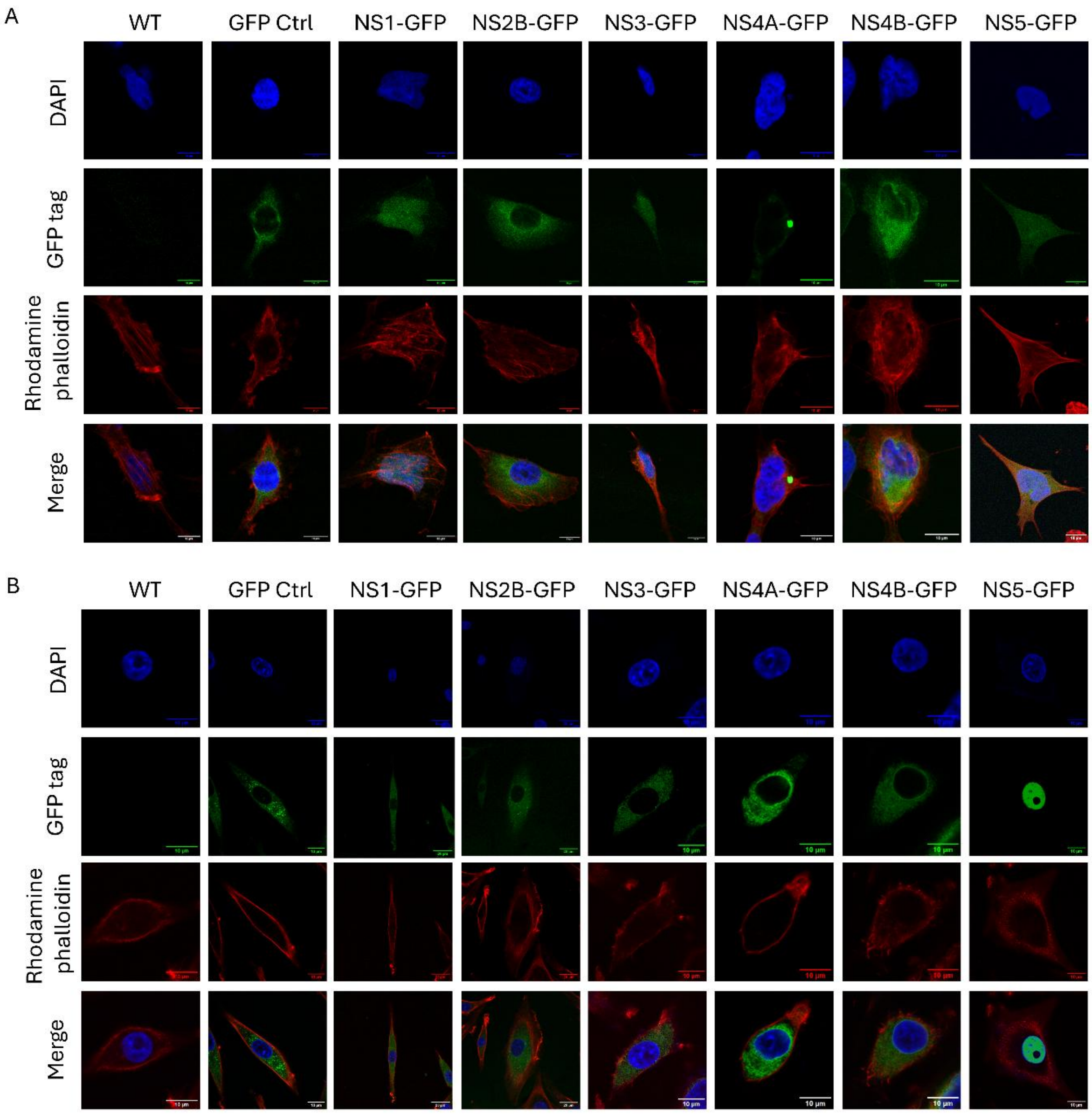


**Figure S2. Subcellular localization of ZIKV non-structural proteins**

Subcellular localization of ZIKV non-structural proteins in SH-SY5Y (A) and USP7 (B) cells. The nucleus, stained with DAPI, is shown in blue; the cytoskeleton, stained with rhodamine phalloidin, is shown in red; and ZIKV proteins tagged with GFP are shown in green. Scale bar = 10 μm.

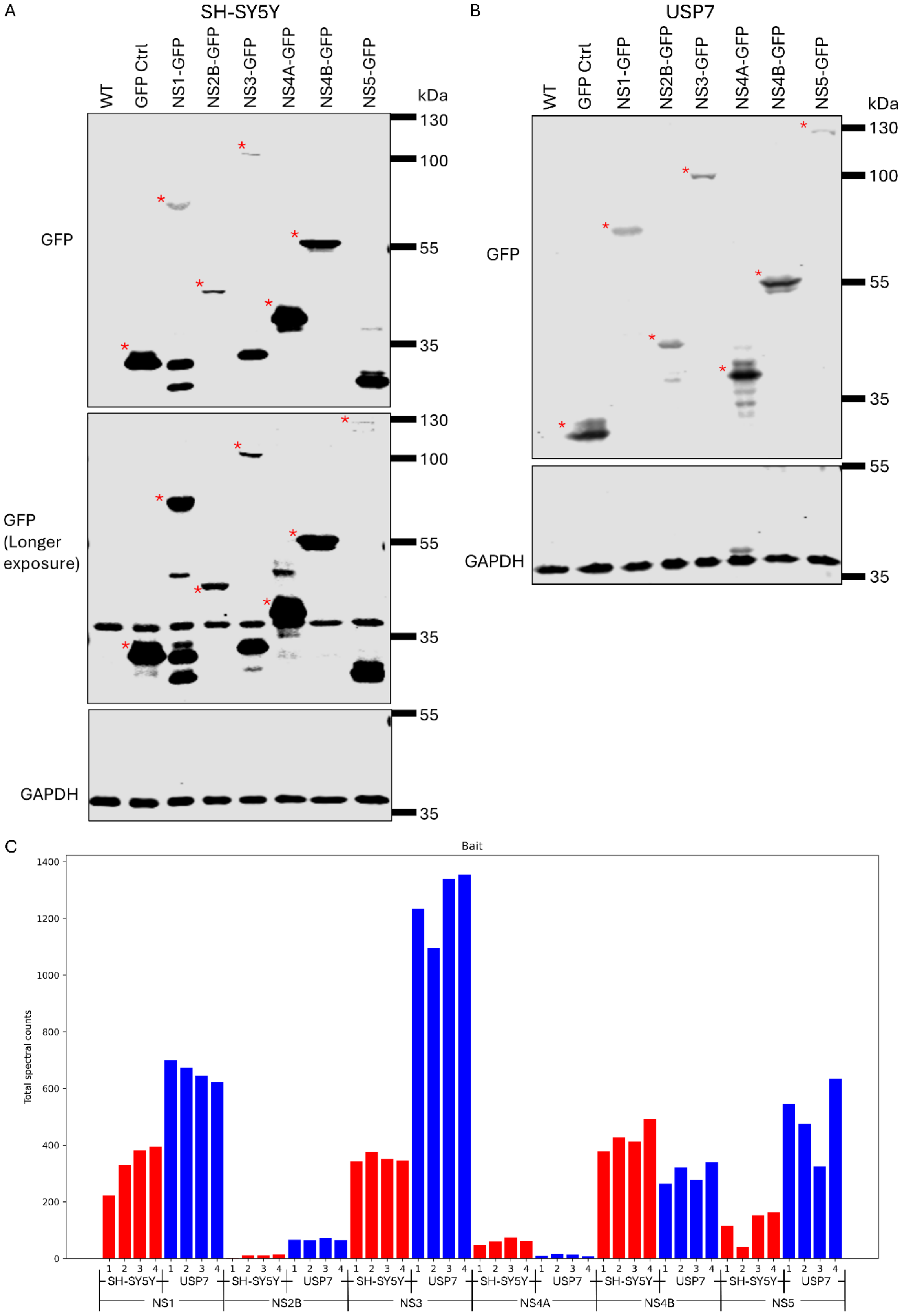
A
SH-SY5Y
WT
GFP Ctrl
NS1-GFP
NS2B-GFP
NS3-GFP
NS4A-GFP
NS4B-GFP
NS5-GFP
kDa
130
100
55
35
GFP
GFP (Longer exposure)
GAPDH
B
USP7
C
Bait
Total spectral counts
0
200
400
600
800
1000
1200
1400
1 2 3 4
SH-SY5Y
USP7
NS1
NS2B
NS3
NS4A
NS4B
NS5

**Figure S3. Detection of ZIKV non-structural proteins**

(A-B) Western blot analysis using GFP and GAPDH primary antibodies in SH-SY5Y (A) and USP7 (B) cells transduced with ZIKV non-structural proteins. Red asterisks indicate bands corresponding to GFP-tagged non-structural proteins at the expected molecular weights. NS5-GFP in SH-SY5Y cells was only detectable after longer exposure. (C) Total spectral counts of all ZIKV non-structural proteins (baits) in SH-SY5Y and USP7 affinity proteomics samples. Replicates for each sample are labelled as 1, 2, 3, and 4.

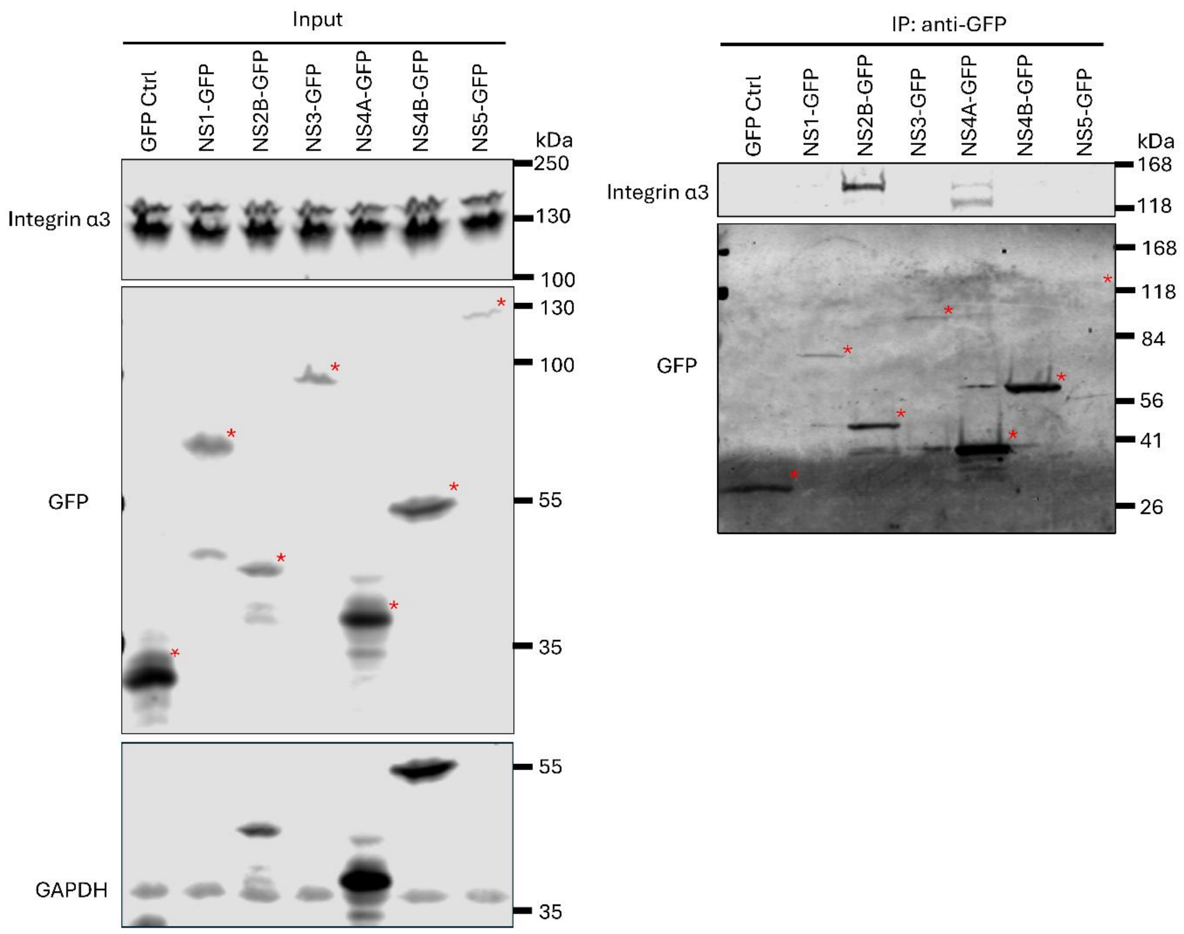


**Figure S4. Co-IP reveals ZIKV non-structural proteins interacting with integrin α3**

Co-IP analysis showing interactions between integrin α3 and ZIKV non-structural proteins in USP7 cells. Red asterisks indicate bands corresponding to GFP-tagged non-structural proteins at the expected molecular weights.

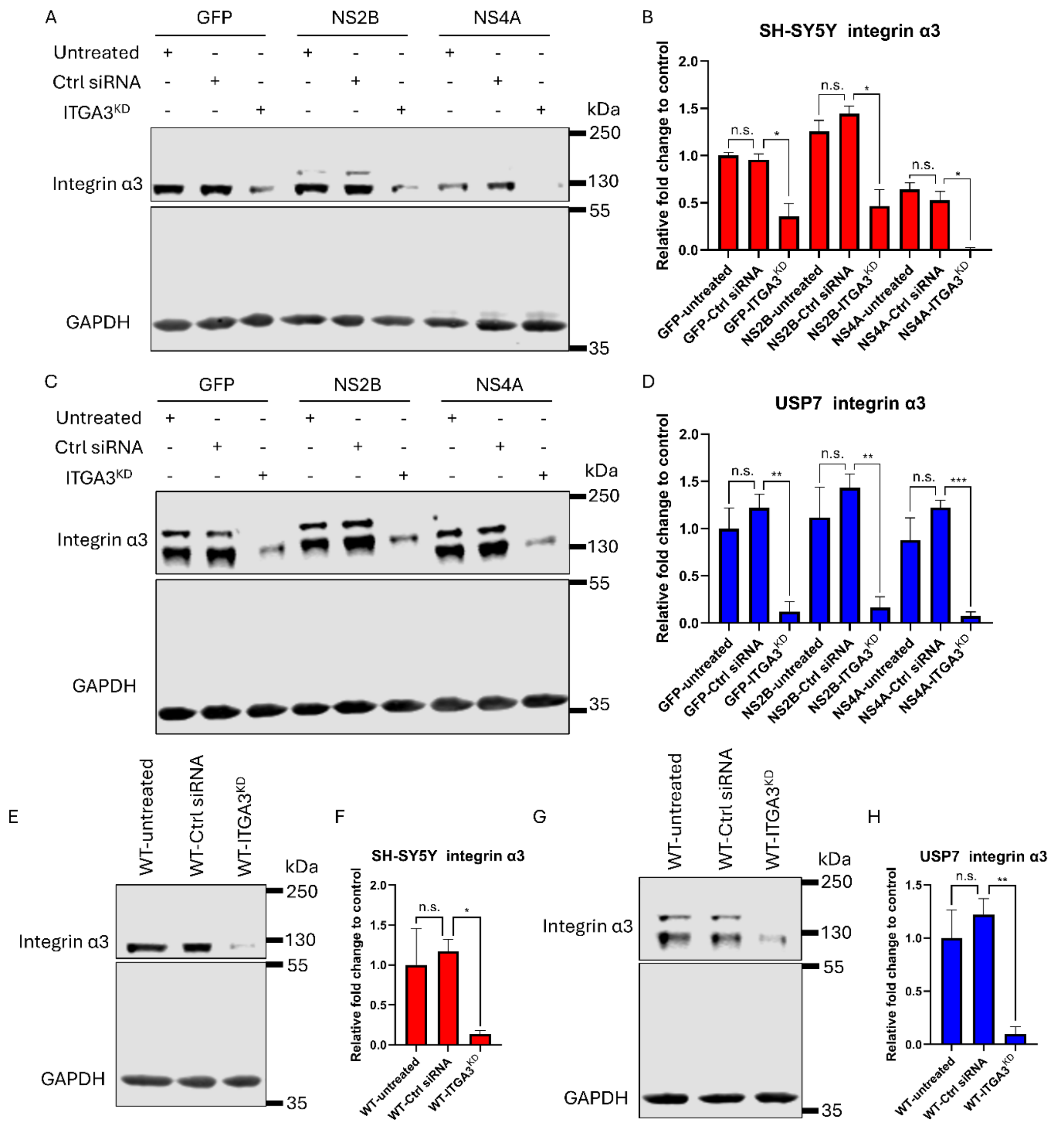


**Figure S5. Knockdown of *ITGA3* in SH-SY5Y and USP7 wild-type cells or cells expressing GFP, NS2B, or NS4A**

(A-B) Western blot images showing integrin α3 expression in SH-SY5Y cells transduced with GFP, NS2B-GFP or NS4A-GFP under the following conditions: untreated, RISC-Free control (Ctrl siRNA), and *ITGA3* knockdown ($ITGA3^{KD}$) (A), with corresponding relative fold changes (B). (C-D) Western blot images (C) and corresponding relative fold changes (D) under the same conditions as in (A-B) for USP7 cells. (E-F) Western blot images showing integrin α3 expression in SH-SY5Y wild-type cells under the same treatments (E), with corresponding relative fold changes (F). (G-H) Western blot images (G) and corresponding relative fold changes (H) under the same conditions as in (E-F) for USP7 cells. $N = 3$, $n = 1$. All values are normalized to GAPDH control. Data are presented as mean ± SD. Statistical significance was calculated using one-way ANOVA with Dunnett's T3 multiple comparisons test. *$p < 0.05$; **$p < 0.01$; ***$p < 0.001$. n.s. indicates no significance.

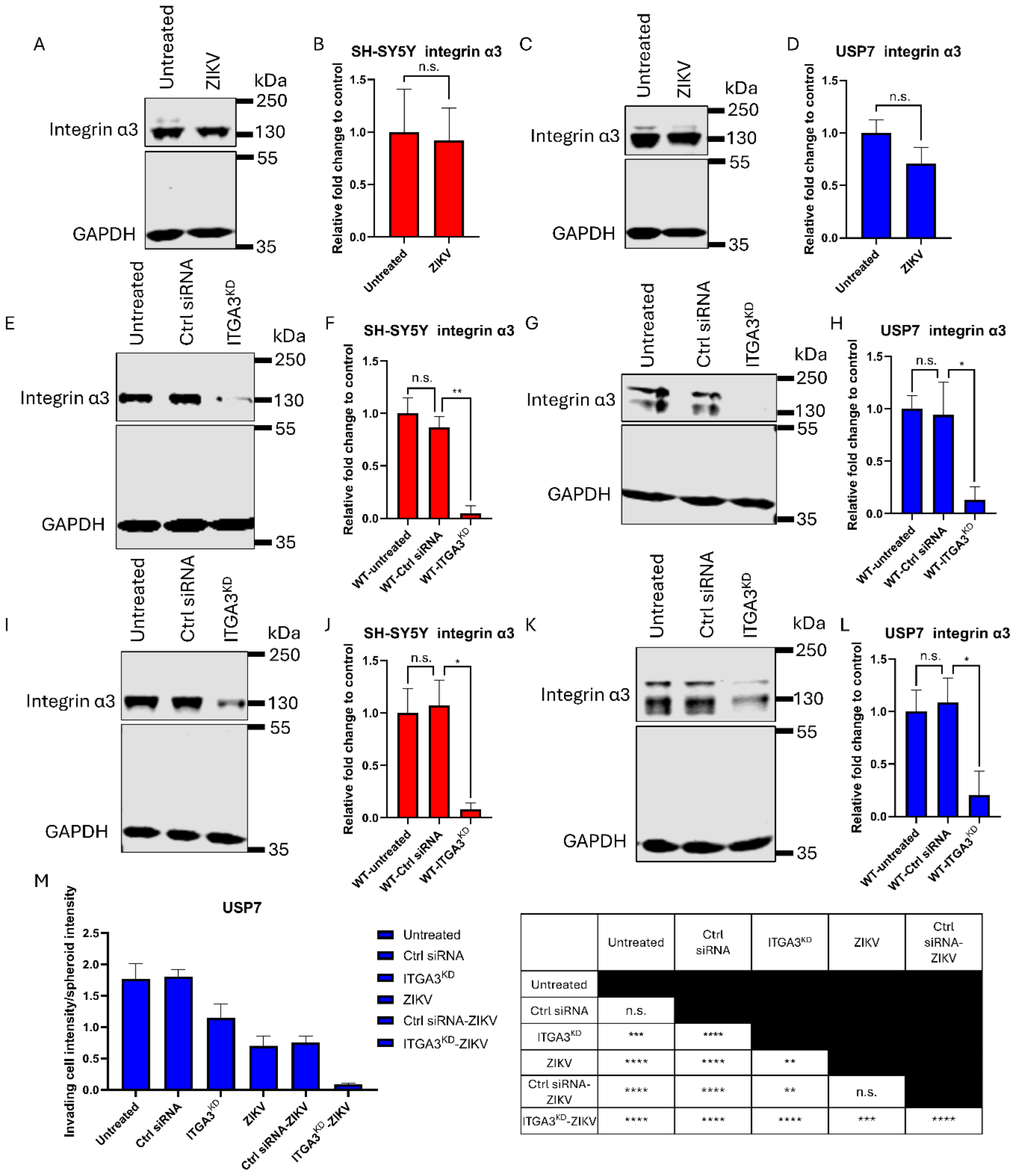


| | Untreated | Ctrl siRNA | ITGA3$^{KD}$ | ZIKV | Ctrl siRNA-ZIKV |
|---|---|---|---|---|---|
| Untreated | | | | | |
| Ctrl siRNA | n.s. | | | | |
| ITGA3$^{KD}$ | *** | **** | | | |
| ZIKV | **** | **** | ** | | |
| Ctrl siRNA-ZIKV | **** | **** | ** | n.s. | |
| ITGA3$^{KD}$-ZIKV | **** | **** | **** | *** | **** |

**Figure S6. Integrin α3 expression levels and 3D invasion analysis, related to Figure 8**

(A-B) Western blot images showing integrin α3 expression in SH-SY5Y cells following ZIKV infection (A), with corresponding relative fold changes (B). (C-D) Western blot images (C) and corresponding relative fold changes (D) under the same conditions as in (A-B) for USP7 cells. (E-F) Western blot images showing integrin α3 expression in SH-SY5Y cells under the following conditions: untreated, RISC-Free control (Ctrl siRNA), and *ITGA3* knockdown

(ITGA3$^{KD}$) (E), with corresponding relative fold changes (F). (G-H) Western blot images (G) and corresponding relative fold changes (H) under the same conditions as in (E-F) for USP7 cells. (I-J) Western blot images showing integrin α3 expression in SH-SY5Y cells 8 days after the same treatments as in (E-F) (I), with corresponding relative fold changes (J). (K-L) Western blot images (K) and corresponding relative fold changes (L) under the same conditions as in (I-J) for USP7 cells. (M) 3D invasion assay in USP7 tumorspheres under the following conditions: untreated, RISC-Free control (Ctrl siRNA) or *ITGA3* knockdown (ITGA3$^{KD}$) in the presence or absence of ZIKV infection, showing the ratio of invading cell intensity to spheroid intensity 4 days post-treatment. For (A-D): $N = 3$, $n = 1$; statistical significance was calculated using student's $t$-test. For (E-L): $N = 3$, $n = 1$; statistical significance was calculated using one-way ANOVA with Dunnett's T3 multiple comparisons test. For (M): $N = 3$, $n \geq 2$; statistical significance was calculated using one-way ANOVA with Dunnett's T3 multiple comparisons test; the table shows the statistical significance. Data are presented as mean ± SD. *$p < 0.05$; **$p < 0.01$; ***$p < 0.001$; ****$p < 0.0001$. n.s. indicates no significance.
