## Supplementary Tables for "Multi-omics analysis reveals integrin α3-dependent mechanisms of Zika virus oncolytic activity in pediatric neural tumors"

### Summary of supplementary tables

**Table S1. List of DEGs in SH-SY5Y tumorspheres**
**Table S2. List of DEGs in IMR-32 tumorspheres**
**Table S3. List of interactomes in SH-SY5Y cells**
**Table S4. List of interactomes in USP7 cells**
**Table S5. Number and percentage of interacting proteins identified in SH-SY5Y and USP7 cells**

**Table S1. List of DEGs in SH-SY5Y tumorspheres**

| gene | baseMean | log2FoldChange | lfcSE | pvalue | padj |
|---|---|---|---|---|---|
| IFNB1 | 9.380886605 | 4.57039368 | 1.183977102 | 1.11E-08 | 1.27E-06 |
| SLC5A10 | 30.96858279 | 4.421729151 | 0.665206988 | 9.81E-14 | 3.36E-11 |
| ARHGAP9 | 231.928379 | 4.36845269 | 0.348969676 | 6.22E-38 | 2.60E-34 |
| KLF4 | 113.338439 | 4.09425509 | 0.479475145 | 1.36E-19 | 1.14E-16 |
| LINC02940 | 51.59302275 | 4.092278219 | 0.456461962 | 3.35E-21 | 3.50E-18 |
| CXCL10 | 25.25637694 | 3.970319511 | 0.687502362 | 3.76E-11 | 7.40E-09 |
| CCL5 | 39.15921262 | 3.817147914 | 0.537404212 | 1.10E-14 | 4.60E-12 |
| OSM | 8.623665211 | 3.580490215 | 1.293984871 | 8.24E-07 | 5.11E-05 |
| OASL | 73.58948831 | 3.522273563 | 0.791093155 | 2.36E-08 | 2.49E-06 |
| IL21R | 32.55118616 | 3.317501687 | 0.592322653 | 1.37E-10 | 2.36E-08 |
| ULBP1 | 62.2330267 | 3.149248812 | 0.582015502 | 3.84E-10 | 6.22E-08 |
| ATF3 | 910.6396086 | 3.13335766 | 0.191518536 | 4.44E-62 | 4.63E-58 |
| DMGDH | 53.27293075 | 3.112780586 | 0.488547841 | 1.42E-12 | 3.74E-10 |
| ZNF473CR | 10.41374569 | 3.028806826 | 1.167301672 | 3.03E-06 | 0.000151601 |
| COL20A1 | 25.19204958 | 2.974088154 | 0.49511373 | 1.35E-11 | 2.93E-09 |
| QPCT | 44.34644786 | 2.930689177 | 0.413207756 | 1.05E-14 | 4.46E-12 |
| GRAP | 18.2470821 | 2.862995521 | 0.711290636 | 1.60E-07 | 1.27E-05 |
| BIRC3 | 220.9806534 | 2.861267665 | 0.319974161 | 3.20E-21 | 3.50E-18 |
| BHLHA15 | 73.02804346 | 2.777384966 | 0.339891225 | 2.50E-18 | 1.74E-15 |
| CASS4 | 14.223046 | 2.7737017 | 0.7690247 | 5.76E-07 | 3.76E-05 |
| MIOX | 25.02769683 | 2.751606086 | 0.595556593 | 1.85E-08 | 1.97E-06 |
| LSMEM2 | 17.62827302 | 2.72854539 | 0.734010409 | 4.41E-07 | 3.08E-05 |
| AOC3 | 72.63962932 | 2.713987188 | 0.459590465 | 2.61E-11 | 5.40E-09 |
| DDIT3 | 1334.326188 | 2.69803278 | 0.208854165 | 2.97E-40 | 2.07E-36 |
| HEATR9 | 12.45308922 | 2.681309674 | 0.833420654 | 1.60E-06 | 8.97E-05 |
| TNXB | 71.14266382 | 2.654540806 | 0.323875215 | 2.11E-18 | 1.52E-15 |
| PIWIL2 | 56.7774651 | 2.653180564 | 0.414764204 | 1.28E-12 | 3.46E-10 |
| LVRN | 60.28269674 | 2.644557782 | 0.336054258 | 3.04E-17 | 1.67E-14 |
| LHFPL3-AS2 | 13.4687522 | 2.626423842 | 0.761422129 | 9.72E-07 | 5.90E-05 |
| SLC6A13 | 18.56113255 | 2.624516876 | 0.546051981 | 8.70E-09 | 1.03E-06 |
| ADM2 | 342.4106912 | 2.584445622 | 0.588002389 | 4.98E-08 | 4.71E-06 |
| R3HDML | 8.818357058 | 2.523820444 | 1.497588858 | 1.57E-05 | 0.00056898 |
| PHF24 | 38.77760652 | 2.493498629 | 0.389776506 | 1.43E-12 | 3.74E-10 |
| ALDH8A1 | 71.12915439 | 2.483946216 | 0.442019869 | 1.56E-10 | 2.64E-08 |
| ARC | 255.6249378 | 2.429417253 | 0.358970015 | 1.31E-13 | 4.20E-11 |
| RAB44 | 22.16137442 | 2.406038659 | 0.571970821 | 1.12E-07 | 9.50E-06 |
| VLDLR-AS1 | 84.95259208 | 2.352921405 | 0.38038246 | 6.32E-12 | 1.50E-09 |
| CHAC1 | 486.0714556 | 2.310056226 | 0.234438813 | 8.65E-25 | 1.13E-21 |
| ACVR1C | 29.29089092 | 2.290897667 | 0.433992609 | 1.12E-09 | 1.68E-07 |
| PPEF2 | 16.88729822 | 2.282665131 | 0.62384276 | 7.07E-07 | 4.46E-05 |
| SELPLG | 65.1067255 | 2.269546759 | 0.452982248 | 4.23E-09 | 5.32E-07 |
| PSD2 | 282.0804687 | 2.268248185 | 0.286344176 | 3.17E-17 | 1.70E-14 |
| CARS1-AS1 | 12.86519791 | 2.242151923 | 0.81049141 | 6.41E-06 | 0.000275705 |
| TGFA | 14.30514698 | 2.225912034 | 0.659797997 | 1.61E-06 | 8.98E-05 |
| TMEM253 | 37.1197976 | 2.217442937 | 0.466862219 | 1.42E-08 | 1.56E-06 |
| SLC7A11 | 1400.181204 | 2.199741276 | 0.273038962 | 1.20E-17 | 6.99E-15 |
| NEURL3 | 12.62126671 | 2.159799131 | 1.232167285 | 3.46E-05 | 0.001093728 |
| C1S | 53.92857803 | 2.145386139 | 0.345642285 | 7.00E-12 | 1.64E-09 |

| | | | | | |
|---|---|---|---|---|---|
| HSPA6 | 35.71083489 | 2.142836508 | 0.563092005 | 4.96E-07 | 3.36E-05 |
| TNFRSF9 | 529.7202433 | 2.126663956 | 0.35399831 | 2.36E-11 | 4.92E-09 |
| TINCR | 31.77275532 | 2.122822184 | 0.408261722 | 1.87E-09 | 2.69E-07 |
| NXPH4 | 100.495833 | 2.069550893 | 0.290080241 | 1.64E-14 | 6.48E-12 |
| NCF1 | 23.12344682 | 2.06796185 | 0.506001604 | 2.03E-07 | 1.57E-05 |
| CDH15 | 35.31712666 | 2.055196145 | 0.399254719 | 2.48E-09 | 3.48E-07 |
| TNFAIP3 | 179.5298466 | 2.048764531 | 0.286882257 | 1.60E-14 | 6.43E-12 |
| FAM83G | 929.5361554 | 2.026711387 | 0.302385811 | 3.33E-13 | 1.01E-10 |
| RND1 | 115.7779907 | 2.025769455 | 0.338893576 | 3.02E-11 | 6.19E-09 |
| SLC7A5 | 5223.05766 | 2.024124243 | 0.18105518 | 1.27E-30 | 2.95E-27 |
| SESN2 | 1871.081139 | 2.017801846 | 0.16357017 | 1.50E-36 | 4.48E-33 |
| ASNS | 458.2039082 | 2.016406586 | 0.186092653 | 5.80E-29 | 1.01E-25 |
| SLC52A1 | 9.325933884 | 2.014398701 | 0.955806039 | 2.80E-05 | 0.000922189 |
| TRIB3 | 2317.278487 | 1.964971142 | 0.192929601 | 6.05E-26 | 8.42E-23 |
| DHX58 | 18.2608922 | 1.92786983 | 0.528868188 | 8.96E-07 | 5.49E-05 |
| SLC22A31 | 13.7449898 | 1.927765826 | 0.748192698 | 1.31E-05 | 0.000487293 |
| BEST1 | 160.8015372 | 1.925207174 | 0.265906525 | 8.66E-15 | 3.77E-12 |
| ELL3 | 10.48727893 | 1.911028816 | 0.813494502 | 2.10E-05 | 0.000725358 |
| ALPK3 | 159.8918558 | 1.894491023 | 0.309922323 | 1.41E-11 | 3.03E-09 |
| ISG20 | 118.8745544 | 1.889837302 | 0.287240021 | 7.83E-13 | 2.18E-10 |
| HERPUD1 | 5059.846672 | 1.8643188 | 0.145464533 | 4.35E-39 | 2.27E-35 |
| PHLDA1-DT | 52.8980646 | 1.854707394 | 0.393336274 | 1.79E-08 | 1.92E-06 |
| GZMM | 18.5682462 | 1.812805862 | 0.585811473 | 4.38E-06 | 0.000207885 |
| FCGR1CP | 18.26628256 | 1.80697411 | 0.575595171 | 3.92E-06 | 0.000188848 |
| RRAD | 68.20224127 | 1.77975927 | 0.369983075 | 1.14E-08 | 1.29E-06 |
| CHILL1 | 12.40382415 | 1.775662162 | 0.758974627 | 2.51E-05 | 0.000840767 |
| SLC5A2 | 28.8787785 | 1.749595147 | 0.389092891 | 4.29E-08 | 4.11E-06 |
| CEBPB | 1034.546687 | 1.744914594 | 0.170782501 | 4.49E-26 | 6.70E-23 |
| SLCO4C1 | 64.56366082 | 1.735304594 | 0.416505214 | 1.56E-07 | 1.25E-05 |
| INSC | 35.78246872 | 1.735007371 | 0.801677692 | 3.71E-05 | 0.001144712 |
| TG | 40.41765759 | 1.731377801 | 0.406459861 | 1.09E-07 | 9.37E-06 |
| SLC1A4 | 1924.606322 | 1.728008379 | 0.163890973 | 1.47E-27 | 2.37E-24 |
| NPC1L1 | 19.324703 | 1.691275549 | 0.765635595 | 3.60E-05 | 0.001122009 |
| GARIN5B | 8.157204927 | 1.689884315 | 1.266334283 | 0.000115255 | 0.002812632 |
| RPL32P27 | 7.428236776 | 1.688604237 | 1.266230446 | 0.000115615 | 0.002817532 |
| FES | 55.53028811 | 1.682336455 | 0.403653768 | 1.51E-07 | 1.23E-05 |
| TMEM215 | 61.96196973 | 1.677457061 | 0.70533207 | 2.60E-05 | 0.000865235 |
| STC2 | 919.1343275 | 1.671576038 | 0.336738231 | 5.08E-09 | 6.32E-07 |
| SPX | 11.93143177 | 1.65664035 | 0.949801375 | 8.50E-05 | 0.002213807 |
| SCD | 29986.2574 | 1.654211825 | 0.215808975 | 2.86E-16 | 1.39E-13 |
| IFIT2 | 173.3944875 | 1.634449571 | 0.250764996 | 8.41E-13 | 2.31E-10 |
| JDP2 | 557.3845234 | 1.616301638 | 0.140798594 | 3.68E-32 | 9.61E-29 |
| PCK2 | 1733.016727 | 1.602567246 | 0.241805057 | 3.81E-13 | 1.12E-10 |
| PCDH19 | 20.75918431 | 1.598891266 | 0.558542938 | 9.09E-06 | 0.000361039 |
| HID1 | 1024.040881 | 1.582164715 | 0.259585764 | 1.01E-11 | 2.29E-09 |
| GABRR2 | 273.4595915 | 1.574422775 | 0.275707669 | 9.16E-11 | 1.68E-08 |
| DNAJB9 | 3206.965465 | 1.574214816 | 0.176434869 | 6.89E-21 | 6.60E-18 |
| HLA-B | 65.99118335 | 1.572384398 | 0.364797328 | 7.92E-08 | 7.22E-06 |
| ANKK1 | 11.70267427 | 1.565439429 | 0.765641049 | 5.87E-05 | 0.001651025 |
| TNFSF13B | 167.2444435 | 1.560637999 | 0.308956043 | 2.81E-09 | 3.84E-07 |

| | | | | | |
|---|---|---|---|---|---|
| MUC20 | 119.0863496 | 1.558310753 | 0.234066849 | 2.75E-13 | 8.51E-11 |
| FCRL6 | 8.151487903 | 1.555350937 | 1.183735851 | 0.000160028 | 0.003613169 |
| AOC2 | 132.1407897 | 1.550896942 | 0.298794514 | 1.38E-09 | 2.03E-07 |
| PYGM | 75.75309286 | 1.550452648 | 0.396992917 | 3.71E-07 | 2.62E-05 |
| ZC3H12D | 40.71493081 | 1.538246504 | 0.495705557 | 4.93E-06 | 0.000227463 |
| CFAP43 | 182.6382954 | 1.534576967 | 0.265334463 | 5.50E-11 | 1.05E-08 |
| AHRR | 31.90799688 | 1.518116829 | 0.550814759 | 1.26E-05 | 0.000475833 |
| NOCT | 543.6869319 | 1.515535035 | 0.161021052 | 6.09E-23 | 7.49E-20 |
| LSMEM1 | 131.1133193 | 1.503680046 | 0.303253335 | 3.98E-09 | 5.10E-07 |
| INSIG1 | 5140.666632 | 1.503421463 | 0.220670575 | 8.16E-14 | 2.94E-11 |
| ALDH1L2 | 724.5613213 | 1.48254854 | 0.376039497 | 3.11E-07 | 2.25E-05 |
| TIGAR | 737.6520763 | 1.481420899 | 0.183268714 | 5.94E-18 | 3.76E-15 |
| SHMT2 | 5954.511375 | 1.480464914 | 0.1730515 | 1.16E-19 | 1.01E-16 |
| C1orf162 | 81.56930089 | 1.476395489 | 0.262344746 | 1.14E-10 | 2.02E-08 |
| MSANTD1 | 41.42948712 | 1.469941051 | 0.397948402 | 7.64E-07 | 4.78E-05 |
| BTLA | 4.902391572 | 1.459709825 | 1.482036362 | 0.000134982 | 0.003171089 |
| SLC3A2 | 6186.295548 | 1.457145528 | 0.132682832 | 4.92E-30 | 1.03E-26 |
| MTHFD2 | 6130.774132 | 1.454643972 | 0.164092414 | 6.95E-21 | 6.60E-18 |
| RELB | 1452.876878 | 1.45206161 | 0.206010671 | 1.33E-14 | 5.46E-12 |
| NFKBIE | 843.640057 | 1.432551017 | 0.194289296 | 1.19E-15 | 5.64E-13 |
| ENO3 | 2350.011742 | 1.429454096 | 0.201207012 | 8.26E-15 | 3.67E-12 |
| LINC02804 | 35.06543006 | 1.422704326 | 0.562910819 | 2.44E-05 | 0.000816632 |
| DNAJB5-DT | 14.83538937 | 1.417668574 | 0.988021789 | 0.000199635 | 0.004352159 |
| SAA2 | 3.820203325 | 1.401883335 | 1.526393818 | 0.000123066 | 0.002944151 |
| MSTN | 33.58951323 | 1.389705005 | 0.524575955 | 1.85E-05 | 0.000652083 |
| HHATL | 3.365954632 | 1.376496072 | 1.492355164 | 0.000142672 | 0.003329269 |
| MIR616 | 30.96840868 | 1.372433337 | 0.385928625 | 1.22E-06 | 7.19E-05 |
| H2BC15 | 126.7130148 | 1.368684432 | 0.360406925 | 5.07E-07 | 3.41E-05 |
| FOS | 814.4077417 | 1.359399503 | 0.282131783 | 6.16E-09 | 7.44E-07 |
| LINC00184 | 24.06291135 | 1.355686322 | 0.553670344 | 3.18E-05 | 0.001015128 |
| LINC01068 | 35.32662451 | 1.345945365 | 0.513669096 | 2.08E-05 | 0.000722221 |
| H2BC18 | 48.12215626 | 1.345354923 | 0.372981779 | 1.02E-06 | 6.13E-05 |
| NKPD1 | 71.74466521 | 1.341869889 | 0.325104802 | 1.36E-07 | 1.12E-05 |
| NHLH1 | 66.42497151 | 1.339447639 | 0.337852807 | 2.64E-07 | 1.95E-05 |
| GALR2 | 18.53478352 | 1.339216567 | 0.582159832 | 4.62E-05 | 0.001362531 |
| SCN11A | 77.10798853 | 1.332476762 | 0.366328834 | 9.11E-07 | 5.56E-05 |
| SDF2L1 | 1088.159062 | 1.321935129 | 0.165747499 | 7.45E-18 | 4.45E-15 |
| CMYA5 | 83.11467542 | 1.306937963 | 0.309189651 | 8.81E-08 | 7.92E-06 |
| PRR18 | 28.02905617 | 1.305986294 | 0.490776281 | 1.95E-05 | 0.000680567 |
| HYOU1 | 17788.71034 | 1.300287903 | 0.16075624 | 2.77E-18 | 1.87E-15 |
| DDIT4 | 2376.813615 | 1.29507048 | 0.708557363 | 0.000136171 | 0.00319543 |
| ABCG1 | 145.7023805 | 1.292665455 | 0.325644791 | 2.56E-07 | 1.89E-05 |
| ZNF460 | 108.6727134 | 1.289797573 | 0.450258007 | 1.13E-05 | 0.000434379 |
| HPX | 289.1987688 | 1.289573279 | 0.212739657 | 5.62E-12 | 1.35E-09 |
| C3orf20 | 21.27197597 | 1.28755391 | 0.554656472 | 4.71E-05 | 0.00137722 |
| ATP5MFP4 | 7.26326292 | 1.283650535 | 1.031804318 | 0.000319642 | 0.006268279 |
| TMEM225B | 23.63912827 | 1.274901629 | 0.522348192 | 3.57E-05 | 0.001120483 |
| C20orf202 | 10.64407388 | 1.266951157 | 1.005244646 | 0.000329932 | 0.006433835 |
| LINC03016 | 19.67683816 | 1.263775079 | 0.66397819 | 0.000123784 | 0.002957929 |
| DEF6 | 35.89335321 | 1.262255986 | 0.437985039 | 1.10E-05 | 0.000426217 |

| | | | | | |
|---|---|---|---|---|---|
| CRELD2 | 3398.465737 | 1.261781244 | 0.138751457 | 3.97E-22 | 4.61E-19 |
| ADAMTS18 | 18.2635742 | 1.26149848 | 0.906978199 | 0.000295154 | 0.005872999 |
| ZNF442 | 93.86302962 | 1.260899556 | 0.443330063 | 1.23E-05 | 0.0004649 |
| WARS1 | 5660.873696 | 1.255687209 | 0.211964076 | 1.24E-11 | 2.75E-09 |
| BBC3 | 1091.78268 | 1.255000413 | 0.257531986 | 4.22E-09 | 5.32E-07 |
| AKNA | 293.4496816 | 1.254296198 | 0.425747761 | 9.18E-06 | 0.000363111 |
| UQCC3 | 92.24785848 | 1.25315296 | 0.269854738 | 1.30E-08 | 1.44E-06 |
| REELD1 | 19.73334626 | 1.250896246 | 0.613554079 | 9.50E-05 | 0.002400158 |
| RGS16 | 1756.801379 | 1.249813623 | 0.153830493 | 1.79E-18 | 1.33E-15 |
| NR1D1 | 531.7784767 | 1.249214291 | 0.196108711 | 7.42E-13 | 2.09E-10 |
| NDUFA4L2 | 107.3051091 | 1.240836767 | 0.265889938 | 1.16E-08 | 1.31E-06 |
| RPL17-C18orf32 | 15.49413408 | 1.238384243 | 0.729685362 | 0.000193252 | 0.004244019 |
| ARPP21 | 39.32950776 | 1.234220951 | 0.553780996 | 6.34E-05 | 0.001755746 |
| GOLPH3-DT | 65.16680032 | 1.231849753 | 0.305292524 | 1.96E-07 | 1.52E-05 |
| MYLIP | 875.2350378 | 1.231844433 | 0.178725019 | 2.10E-14 | 7.98E-12 |
| ADCY10 | 22.28666642 | 1.231843156 | 0.559083088 | 6.75E-05 | 0.001850897 |
| PROC | 406.4557001 | 1.228717551 | 0.388928257 | 4.87E-06 | 0.000225989 |
| BHLHE40-AS1 | 42.24757686 | 1.225902354 | 0.577013649 | 8.18E-05 | 0.002139925 |
| SLC6A9 | 491.5387109 | 1.215914302 | 0.24815142 | 3.64E-09 | 4.78E-07 |
| PPP1R15A | 2860.281503 | 1.214121877 | 0.181986692 | 9.51E-14 | 3.31E-11 |
| NFKB2 | 2645.170478 | 1.208288149 | 0.188606149 | 5.58E-13 | 1.60E-10 |
| DNAJA4 | 85.50502672 | 1.204267157 | 0.251094933 | 6.13E-09 | 7.44E-07 |
| ERO1B | 1336.509391 | 1.203860435 | 0.186561909 | 4.10E-13 | 1.19E-10 |
| MYLK3 | 38.42537873 | 1.201792634 | 0.424401513 | 1.36E-05 | 0.000500786 |
| KLHL15 | 1247.345169 | 1.201662757 | 0.180423019 | 1.02E-13 | 3.42E-11 |
| NLRP3 | 5.88253289 | 1.199336803 | 1.219081932 | 0.000371287 | 0.007011147 |
| EAF2 | 42.59692203 | 1.192138975 | 0.337794779 | 1.43E-06 | 8.19E-05 |
| NUP153-AS1 | 24.1419247 | 1.191516294 | 0.524970013 | 6.14E-05 | 0.001708824 |
| ZNF703 | 255.5971201 | 1.191516185 | 0.270870969 | 4.09E-08 | 3.95E-06 |
| BTG2 | 1048.034956 | 1.191354592 | 0.31041025 | 4.45E-07 | 3.10E-05 |
| LINC02680 | 10.29903525 | 1.190915342 | 0.870026029 | 0.000356713 | 0.006841089 |
| TAS1R3 | 71.43773513 | 1.182434622 | 0.270960146 | 4.82E-08 | 4.57E-06 |
| CCDC163 | 149.1758041 | 1.182196822 | 0.380009625 | 5.96E-06 | 0.000260457 |
| IQCN | 1725.452436 | 1.181491358 | 0.291953348 | 1.92E-07 | 1.50E-05 |
| RAPSN | 25.74127392 | 1.176888717 | 0.620810152 | 0.000147097 | 0.003388893 |
| RANP4 | 60.15256313 | 1.17492629 | 0.384670054 | 7.19E-06 | 0.000301491 |
| NUPR1 | 22.81473872 | 1.172701657 | 0.924967767 | 0.00041535 | 0.007690229 |
| ETV7 | 19.5830279 | 1.169875214 | 0.585032437 | 0.000119159 | 0.002880369 |
| GSDMD | 9.925680452 | 1.168538142 | 0.902597003 | 0.000409133 | 0.007608851 |
| ZFAND2A | 988.0381984 | 1.165761159 | 0.145002295 | 3.22E-18 | 2.10E-15 |
| FICD | 361.0152095 | 1.16496107 | 0.174246775 | 8.45E-14 | 2.99E-11 |
| PITPNM3 | 139.9800663 | 1.16436812 | 0.214626321 | 2.21E-10 | 3.66E-08 |
| TMIE | 62.71848467 | 1.151469521 | 0.360518363 | 4.74E-06 | 0.000222551 |
| SLC37A2 | 82.33755471 | 1.149645654 | 0.335527673 | 2.16E-06 | 0.00011474 |
| DHCR7 | 3746.34968 | 1.144125672 | 0.218300864 | 6.22E-10 | 9.77E-08 |
| ZBTB2 | 1842.546512 | 1.137917604 | 0.171776199 | 1.30E-13 | 4.20E-11 |
| AEN | 2627.849174 | 1.136859816 | 0.138835717 | 9.45E-19 | 7.31E-16 |
| FOSL1 | 490.8777783 | 1.136832336 | 0.369532719 | 7.07E-06 | 0.000297608 |
| IL17RE | 129.2542461 | 1.136251107 | 0.227980885 | 2.46E-09 | 3.47E-07 |
| GRIN2C | 213.1400636 | 1.131020567 | 0.237914135 | 7.96E-09 | 9.50E-07 |

| | | | | | |
|---|---|---|---|---|---|
| CEBPG | 3658.380665 | 1.131016064 | 0.186128024 | 4.71E-12 | 1.14E-09 |
| SREBF1 | 4321.456602 | 1.129584623 | 0.228377335 | 3.02E-09 | 4.07E-07 |
| IL20RB | 96.2193083 | 1.120034123 | 0.259524627 | 6.40E-08 | 6.02E-06 |
| PIK3CD | 774.2107406 | 1.107621403 | 0.322578774 | 2.24E-06 | 0.000118753 |
| ZNF654 | 1659.843804 | 1.106181641 | 0.188828909 | 1.88E-11 | 4.00E-09 |
| CT62 | 6.397794502 | 1.095982756 | 1.304869601 | 0.000354473 | 0.006810642 |
| IGHV1-46 | 25.44094997 | 1.094586885 | 0.56323509 | 0.000156171 | 0.003545257 |
| PHGDH | 4235.026967 | 1.093134522 | 0.272029284 | 2.43E-07 | 1.82E-05 |
| TNFRSF10A | 79.26501563 | 1.092720135 | 0.245543761 | 3.63E-08 | 3.61E-06 |
| SDCBP2 | 96.57690952 | 1.083879445 | 0.302784414 | 1.38E-06 | 8.05E-05 |
| IRF1 | 493.7345815 | 1.083637426 | 0.231499751 | 1.24E-08 | 1.39E-06 |
| SAXO4 | 133.2972516 | 1.078761374 | 0.239520658 | 2.91E-08 | 2.95E-06 |
| DNAH12 | 29.72718519 | 1.078402871 | 0.542649377 | 0.000146574 | 0.00338787 |
| GADD45G | 118.9387702 | 1.078286497 | 0.348562158 | 7.40E-06 | 0.000309164 |
| TRIM72 | 6.092672412 | 1.07777473 | 1.191299343 | 0.000495052 | 0.008762006 |
| RASSF9 | 69.32075162 | 1.072930577 | 0.425377724 | 3.97E-05 | 0.001211886 |
| YARS1 | 5452.517075 | 1.069424071 | 0.158725685 | 6.74E-14 | 2.47E-11 |
| FGF8 | 11.46594294 | 1.063816926 | 0.905426454 | 0.00059845 | 0.010286941 |
| SLC7A1 | 5941.752086 | 1.062987113 | 0.154421608 | 2.47E-14 | 9.21E-12 |
| MKNK2 | 2682.603546 | 1.062564902 | 0.187906226 | 6.96E-11 | 1.30E-08 |
| FCGR1A | 12.06883721 | 1.062350164 | 0.765352046 | 0.000475446 | 0.008530669 |
| MXD1 | 848.9203596 | 1.060129058 | 0.176441147 | 8.29E-12 | 1.90E-09 |
| LIF | 287.9777688 | 1.059317387 | 0.373120822 | 1.68E-05 | 0.000600022 |
| DNHD1 | 2812.72369 | 1.057965213 | 0.217245702 | 5.14E-09 | 6.35E-07 |
| SPTBN5 | 144.54778 | 1.054368053 | 0.355365051 | 1.15E-05 | 0.000443954 |
| TSPYL2 | 1933.077197 | 1.051009362 | 0.230799049 | 2.46E-08 | 2.57E-06 |
| GDF15 | 1551.189652 | 1.047792778 | 0.181966896 | 3.95E-11 | 7.70E-09 |
| RPL7P24 | 11.94598954 | 1.047613499 | 0.736372227 | 0.000468938 | 0.00844291 |
| POLR2A | 15459.34111 | 1.045749089 | 0.203979536 | 1.40E-09 | 2.05E-07 |
| HSD3BP5 | 124.6724536 | 1.044152011 | 0.421153317 | 4.73E-05 | 0.001382973 |
| VEGFA | 7032.918366 | 1.041489745 | 0.132618024 | 1.78E-17 | 1.00E-14 |
| ODC1-DT | 46.62800273 | 1.032003938 | 0.348084876 | 1.22E-05 | 0.000464552 |
| PLA2G4C | 192.9617553 | 1.027655449 | 0.26640546 | 5.43E-07 | 3.60E-05 |
| HYPK | 178.6775631 | 1.025646575 | 0.224485669 | 2.47E-08 | 2.57E-06 |
| SEMA4A | 198.7874824 | 1.022255941 | 0.252479866 | 2.53E-07 | 1.88E-05 |
| IFIH1 | 161.644924 | 1.01038401 | 0.349182224 | 1.60E-05 | 0.000577812 |
| BLTP3A | 2214.511216 | 1.009312533 | 0.174078781 | 3.57E-11 | 7.17E-09 |
| CYP51A1 | 569.945179 | 1.003648757 | 0.197357261 | 2.02E-09 | 2.88E-07 |
| FAM184A | 672.9984463 | 1.001707381 | 0.239453398 | 1.53E-07 | 1.23E-05 |
| SLC15A2 | 125.2454131 | 1.000791487 | 0.280648282 | 1.77E-06 | 9.73E-05 |
| TSC22D3 | 1584.037734 | 0.989453814 | 0.17293 | 6.12E-11 | 1.16E-08 |
| TDRKH-AS1 | 127.2504119 | 0.989295691 | 0.232587189 | 1.17E-07 | 9.89E-06 |
| GPT2 | 3178.979586 | 0.989195409 | 0.209172407 | 1.30E-08 | 1.44E-06 |
| ESPNL | 11.19037308 | 0.987916442 | 0.711534448 | 0.000577188 | 0.010009379 |
| TMEM182 | 291.7385736 | 0.985298025 | 0.189004571 | 1.10E-09 | 1.67E-07 |
| ITGB7 | 39.53773714 | 0.97541816 | 0.427154158 | 9.22E-05 | 0.002346655 |
| ACRBP | 113.4672587 | 0.971992164 | 0.257109647 | 8.70E-07 | 5.36E-05 |
| LRIT3 | 50.68362898 | 0.971448413 | 0.391604669 | 5.63E-05 | 0.001599229 |
| CES3 | 64.10096504 | 0.970495178 | 0.344190509 | 2.21E-05 | 0.000750753 |
| ACHE | 242.6053781 | 0.968571821 | 0.272346293 | 2.03E-06 | 0.000108374 |

| | | | | | |
|---|---|---|---|---|---|
| AARD | 3.392044795 | 0.962893635 | 1.313310962 | 0.000352386 | 0.006783032 |
| CITED4 | 189.3083939 | 0.961712695 | 0.425560549 | 0.00010143 | 0.002540007 |
| HSD17B7 | 1249.537935 | 0.961234261 | 0.171527498 | 1.39E-10 | 2.38E-08 |
| PTGES | 203.8809187 | 0.960457055 | 0.399827258 | 7.11E-05 | 0.001929663 |
| FASN | 27356.79182 | 0.957205419 | 0.247339959 | 6.47E-07 | 4.16E-05 |
| RNA5SP39 | 10.75254701 | 0.955046892 | 0.759548582 | 0.000755343 | 0.012363111 |
| TMEM269 | 114.3462561 | 0.95440804 | 0.302795813 | 8.40E-06 | 0.000337921 |
| CBX4 | 2124.32241 | 0.95124161 | 0.165491902 | 6.31E-11 | 1.19E-08 |
| CREBRF | 1764.472368 | 0.949751698 | 0.224701084 | 1.53E-07 | 1.24E-05 |
| MSH4 | 12.34406142 | 0.949080076 | 0.711291419 | 0.000695617 | 0.011569585 |
| HSPA5 | 41631.46493 | 0.943441747 | 0.142184948 | 2.35E-13 | 7.43E-11 |
| KRTAP5-AS1 | 88.09143108 | 0.943215792 | 0.35611205 | 3.88E-05 | 0.001188728 |
| MIDEAS | 2410.192121 | 0.938748709 | 0.222656409 | 1.68E-07 | 1.33E-05 |
| IP6K3 | 22.0615169 | 0.936728606 | 0.521844832 | 0.000313769 | 0.006182136 |
| ALS2CL | 151.777155 | 0.934523649 | 0.306913496 | 1.25E-05 | 0.000472364 |
| SGK2 | 212.3538415 | 0.933895473 | 0.313976535 | 1.56E-05 | 0.000565657 |
| DNAJB11 | 4039.761736 | 0.933235615 | 0.14167897 | 3.46E-13 | 1.03E-10 |
| IDI2 | 8.147479128 | 0.927118229 | 0.877009905 | 0.000964689 | 0.01497958 |
| BHMT2 | 3.06893474 | 0.922642621 | 1.275759179 | 0.000420892 | 0.007765306 |
| MTHFD1L | 2827.031671 | 0.921826839 | 0.111009486 | 8.18E-19 | 6.57E-16 |
| RPL13AP20 | 19.64470144 | 0.919477901 | 0.547197571 | 0.00041583 | 0.007692303 |
| EML2 | 455.854653 | 0.919150878 | 0.17599959 | 1.44E-09 | 2.09E-07 |
| ACSL3 | 6845.400004 | 0.917180065 | 0.146949745 | 3.69E-12 | 9.06E-10 |
| ADAMTSL2 | 1239.7379 | 0.915389254 | 0.197307078 | 2.77E-08 | 2.85E-06 |
| EPGN | 5.235702746 | 0.915358471 | 1.18480326 | 0.000633017 | 0.010730973 |
| ELAPOR1 | 116.3815426 | 0.915320308 | 0.225472457 | 3.58E-07 | 2.55E-05 |
| TRAF1 | 228.6026182 | 0.915089017 | 0.334857338 | 3.35E-05 | 0.00105931 |
| LINC02328 | 39.46397773 | 0.912164465 | 0.462551538 | 0.000229447 | 0.004835525 |
| C1orf116 | 12.44125733 | 0.905928985 | 0.729521518 | 0.000891558 | 0.014127612 |
| MYO7A | 63.47356817 | 0.902225925 | 0.282077641 | 8.64E-06 | 0.000347188 |
| OAZ3 | 195.7687729 | 0.90124858 | 0.214553513 | 2.12E-07 | 1.62E-05 |
| CARS1 | 4511.836844 | 0.901181546 | 0.14810842 | 1.10E-11 | 2.46E-09 |
| SIGLEC14 | 2.870390449 | 0.900012485 | 1.251386138 | 0.000472969 | 0.008500823 |
| PHLDB3 | 470.6125576 | 0.899721558 | 0.220633462 | 3.55E-07 | 2.55E-05 |
| SHISA2 | 8.540201896 | 0.899193298 | 0.775957727 | 0.000992854 | 0.015325762 |
| TLR7 | 17.60936342 | 0.898399833 | 0.656026123 | 0.000766783 | 0.012511139 |
| MVD | 2101.676484 | 0.896372901 | 0.219701812 | 3.58E-07 | 2.55E-05 |
| SERPINF2 | 72.26203223 | 0.895428818 | 0.308025021 | 2.17E-05 | 0.00074339 |
| MVK | 1628.582607 | 0.89528063 | 0.257935452 | 3.58E-06 | 0.000175188 |
| PSMD10P1 | 7.894416386 | 0.895224581 | 0.842755712 | 0.001071794 | 0.016220588 |
| OVGP1 | 115.5866463 | 0.894599282 | 0.269749429 | 6.06E-06 | 0.000264152 |
| SARS1 | 6395.283504 | 0.893650368 | 0.149514531 | 2.23E-11 | 4.71E-09 |
| PSAT1 | 4779.147382 | 0.892376826 | 0.257265315 | 3.65E-06 | 0.000178132 |
| FGF17 | 28.70628824 | 0.891965867 | 0.461742142 | 0.000266558 | 0.005452565 |
| ZNF425 | 651.0166422 | 0.889686477 | 0.175615508 | 3.86E-09 | 4.98E-07 |
| SMOX | 916.8164785 | 0.889036304 | 0.30520702 | 2.18E-05 | 0.000745263 |
| GLIS2 | 731.7527291 | 0.888100892 | 0.272225499 | 7.42E-06 | 0.00030919 |
| KCTD16 | 328.1454185 | 0.884646359 | 0.416110302 | 0.000175522 | 0.003908081 |
| NUAK2 | 297.7889057 | 0.883074555 | 0.416193932 | 0.000178148 | 0.003958112 |
| ARHGAP27 | 370.3256228 | 0.882918604 | 0.210982221 | 2.47E-07 | 1.85E-05 |

| | | | | | |
|---|---|---|---|---|---|
| DNAJC3 | 6359.867173 | 0.882653562 | 0.246233281 | 2.53E-06 | 0.000130706 |
| LYSMD3 | 2125.309914 | 0.882527083 | 0.175346335 | 4.77E-09 | 5.96E-07 |
| ARHGEF33 | 61.95654758 | 0.877305282 | 0.363231194 | 8.87E-05 | 0.002276429 |
| TBX19 | 639.7584211 | 0.877263487 | 0.253932364 | 4.07E-06 | 0.000195058 |
| OTUD1 | 475.9101693 | 0.87283575 | 0.209411918 | 2.78E-07 | 2.03E-05 |
| PLD5P1 | 133.5489269 | 0.87236439 | 0.359548981 | 8.77E-05 | 0.002259567 |
| HRH2 | 88.31718009 | 0.872290688 | 0.325183826 | 4.46E-05 | 0.001333377 |
| ZBTB7B | 2373.799913 | 0.869411835 | 0.169388664 | 3.08E-09 | 4.11E-07 |
| ICAM1 | 421.8612952 | 0.866775807 | 0.230159401 | 1.40E-06 | 8.06E-05 |
| TMEM41B | 2870.658034 | 0.865902601 | 0.185719776 | 3.22E-08 | 3.22E-06 |
| VWA3A | 14.71072424 | 0.861958751 | 0.657233493 | 0.000934637 | 0.014664986 |
| ADGRG1 | 161.5111675 | 0.861106085 | 0.32747265 | 5.36E-05 | 0.001539313 |
| BNIPL | 69.79928136 | 0.859472564 | 0.271208607 | 1.12E-05 | 0.000432774 |
| SEC11C | 1337.617153 | 0.858684027 | 0.156575995 | 4.95E-10 | 7.89E-08 |
| SELENOKP1 | 9.434345627 | 0.856900748 | 0.753998905 | 0.001163219 | 0.017205264 |
| NAMPT | 3904.46178 | 0.854477244 | 0.147895834 | 9.49E-11 | 1.71E-08 |
| KLHL26 | 533.3197878 | 0.854170836 | 0.21552916 | 6.96E-07 | 4.43E-05 |
| FAS | 515.8425394 | 0.853357441 | 0.230872785 | 1.92E-06 | 0.000103328 |
| TNFRSF14-AS1 | 8.575730746 | 0.849522329 | 0.73731207 | 0.001176011 | 0.017369861 |
| TUFT1 | 1073.776144 | 0.848731717 | 0.157273664 | 8.54E-10 | 1.32E-07 |
| GLI1 | 250.9932822 | 0.846915602 | 0.232549344 | 2.41E-06 | 0.00012592 |
| SLFNL1 | 50.08875533 | 0.844341497 | 0.331605849 | 7.11E-05 | 0.001929663 |
| PPM1D | 2391.805795 | 0.841884896 | 0.188348036 | 8.84E-08 | 7.92E-06 |
| FAM182A | 16.86054984 | 0.841575594 | 0.594050171 | 0.00085059 | 0.013591868 |
| KIF1A | 4552.2469 | 0.838303527 | 0.290011159 | 2.82E-05 | 0.00092625 |
| ODC1 | 8354.379958 | 0.837336902 | 0.188241052 | 9.97E-08 | 8.86E-06 |
| CPEB4 | 2732.15336 | 0.832181764 | 0.174274387 | 2.28E-08 | 2.41E-06 |
| GARS1 | 11176.50673 | 0.826288672 | 0.158649201 | 2.69E-09 | 3.72E-07 |
| MGAM | 6.585679069 | 0.819325657 | 0.98753592 | 0.001251847 | 0.018257553 |
| FBXO16 | 160.8423987 | 0.818032161 | 0.235078146 | 4.83E-06 | 0.0002251 |
| XBP1 | 5092.750818 | 0.816991177 | 0.158447068 | 3.73E-09 | 4.87E-07 |
| LINC00910 | 399.8771656 | 0.815414343 | 0.223473864 | 2.71E-06 | 0.00013943 |
| DNAJC1 | 1036.434633 | 0.814660661 | 0.148916342 | 7.05E-10 | 1.10E-07 |
| NANS | 1164.254124 | 0.813866067 | 0.126359266 | 2.06E-12 | 5.25E-10 |
| MATN1 | 25.24510247 | 0.813158071 | 0.450375444 | 0.000452747 | 0.00821514 |
| PARP10 | 149.3378442 | 0.812897059 | 0.253443708 | 1.21E-05 | 0.000459942 |
| SOX30 | 33.83662268 | 0.810902509 | 0.442180292 | 0.00043039 | 0.007919548 |
| INSIG1-DT | 184.6924562 | 0.808129897 | 0.299615242 | 5.48E-05 | 0.001569757 |
| TNFRSF10B | 6906.220376 | 0.804531834 | 0.19738844 | 5.68E-07 | 3.72E-05 |
| HMGCS1 | 6877.074688 | 0.803050091 | 0.186446889 | 2.20E-07 | 1.67E-05 |
| LINC01748 | 45.32726806 | 0.80286252 | 0.309038395 | 7.30E-05 | 0.001973439 |
| SLC1A5 | 4031.318209 | 0.802645894 | 0.108762962 | 3.14E-15 | 1.46E-12 |
| UCN2 | 29.6744722 | 0.799931627 | 0.609554231 | 0.00114856 | 0.017049373 |
| ZGLP1 | 68.4584078 | 0.799643358 | 0.276841867 | 3.33E-05 | 0.001056651 |
| MIR34AHG | 486.2612024 | 0.798571946 | 0.20792197 | 1.45E-06 | 8.30E-05 |
| SDK2 | 438.3175742 | 0.798110471 | 0.384867168 | 0.000268201 | 0.005480804 |
| NFKB1 | 2356.480952 | 0.797090035 | 0.133221512 | 4.10E-11 | 7.94E-09 |
| MFNG | 8.153178169 | 0.795800551 | 0.738542489 | 0.001508474 | 0.020836301 |
| NFAM1 | 13.99094107 | 0.794824381 | 0.681765996 | 0.001403533 | 0.019825112 |
| FAM117A | 522.9487668 | 0.791628774 | 0.227674331 | 5.50E-06 | 0.000245399 |

| | | | | | |
|---|---|---|---|---|---|
| PMEPA1 | 4646.985387 | 0.791607513 | 0.230000917 | 6.19E-06 | 0.000268919 |
| PYCR1 | 3830.7268 | 0.7897863 | 0.150503801 | 2.73E-09 | 3.76E-07 |
| RN7SL600P | 16.00057301 | 0.788458837 | 0.586739621 | 0.001142262 | 0.016979462 |
| KRTAP5-9 | 15.83263838 | 0.786722556 | 0.601082406 | 0.001209475 | 0.017713798 |
| UPP2 | 7.761107712 | 0.786546546 | 0.787293351 | 0.001618283 | 0.021960904 |
| LARP1B | 1315.661104 | 0.784172689 | 0.177812768 | 1.56E-07 | 1.25E-05 |
| TXK | 28.37792973 | 0.783433564 | 0.417735032 | 0.000437017 | 0.008020297 |
| TNFSF4 | 220.8357206 | 0.783172317 | 0.201237391 | 1.29E-06 | 7.60E-05 |
| CAPN9 | 8.987584306 | 0.781751639 | 0.699472449 | 0.001540717 | 0.021155742 |
| PLEKHF2 | 642.788387 | 0.778336914 | 0.21373927 | 3.31E-06 | 0.000164606 |
| RPS12P3 | 22.70332297 | 0.774832998 | 0.491127084 | 0.000811306 | 0.013084269 |
| GDNF | 58.92998139 | 0.774774735 | 0.502576588 | 0.000866007 | 0.013774979 |
| SEC24A | 3625.434142 | 0.770595666 | 0.170895116 | 1.09E-07 | 9.37E-06 |
| ANKRD33B | 285.3851118 | 0.770019407 | 0.450217602 | 0.000643004 | 0.010882605 |
| CALCR | 31.87394867 | 0.76876399 | 0.432148955 | 0.000564002 | 0.009824179 |
| EFNA1 | 940.4656955 | 0.768564691 | 0.249517643 | 2.17E-05 | 0.00074339 |
| LINC02934 | 24.26796953 | 0.765621727 | 0.463183984 | 0.000730297 | 0.012047588 |
| MANF | 4272.863451 | 0.76506857 | 0.129442954 | 7.83E-11 | 1.45E-08 |
| DNAJC25 | 668.5255407 | 0.764239762 | 0.184946208 | 5.54E-07 | 3.65E-05 |
| ZFPM1 | 2080.292209 | 0.763508648 | 0.217397255 | 5.56E-06 | 0.000247696 |
| EGR1 | 3524.25378 | 0.763150923 | 0.209051063 | 3.44E-06 | 0.000170455 |
| IRAK2 | 200.2617805 | 0.762241825 | 0.307529831 | 0.000117958 | 0.002857956 |
| MIA | 5.366604471 | 0.760765115 | 1.009696427 | 0.001343754 | 0.019169103 |
| CHIC2 | 788.4131906 | 0.757050866 | 0.149222015 | 8.10E-09 | 9.61E-07 |
| JAZF1 | 377.0712092 | 0.756757184 | 0.195214651 | 1.56E-06 | 8.81E-05 |
| ZNF773 | 464.0335411 | 0.756023725 | 0.231572017 | 1.30E-05 | 0.000485186 |
| BCAN-AS1 | 92.1950276 | 0.754052109 | 0.35269477 | 0.000277038 | 0.005612675 |
| TUBE1 | 1559.765288 | 0.753939517 | 0.182977142 | 6.11E-07 | 3.95E-05 |
| VWCE | 403.6602008 | 0.753194606 | 0.172775708 | 2.28E-07 | 1.72E-05 |
| KLHL28 | 1497.319408 | 0.751643537 | 0.179853907 | 4.87E-07 | 3.34E-05 |
| WDR49 | 2.736536259 | 0.748501801 | 1.145239822 | 0.000753295 | 0.012339268 |
| FAM174B | 329.1129744 | 0.747616223 | 0.248846263 | 2.99E-05 | 0.00097033 |
| CDK2AP2 | 1689.968097 | 0.747379082 | 0.169440603 | 1.88E-07 | 1.48E-05 |
| TMEM63C | 314.0049189 | 0.746499542 | 0.190516253 | 1.40E-06 | 8.06E-05 |
| FBLL1 | 646.8371798 | 0.745476233 | 0.203167416 | 3.50E-06 | 0.00017266 |
| OSGIN2 | 1383.275893 | 0.744688712 | 0.186239589 | 1.03E-06 | 6.21E-05 |
| POLR2MP1 | 38.54569821 | 0.744147926 | 0.333010073 | 0.00023125 | 0.004863712 |
| TIFA | 752.4680786 | 0.742304298 | 0.334011792 | 0.000239878 | 0.005004852 |
| SYVN1 | 3384.993459 | 0.737675809 | 0.175434032 | 4.70E-07 | 3.25E-05 |
| TLL2 | 33.75829222 | 0.736322174 | 0.436177018 | 0.000767422 | 0.012511793 |
| ARRDC3 | 3405.741691 | 0.735862525 | 0.239597129 | 2.59E-05 | 0.000861839 |
| KCNE2 | 3.602009569 | 0.73432355 | 1.098624916 | 0.000966707 | 0.01499976 |
| PHKG1 | 191.3813115 | 0.733506336 | 0.203733981 | 4.72E-06 | 0.000221937 |
| TMEM38A | 236.1576232 | 0.7327895 | 0.206840595 | 5.78E-06 | 0.000254995 |
| CEACAM19 | 117.9001554 | 0.732722016 | 0.443894719 | 0.00083603 | 0.013389948 |
| LEISA1 | 199.8993268 | 0.729039797 | 0.328491384 | 0.00025617 | 0.005281445 |
| PTPN22 | 5.658906445 | 0.728247377 | 0.881569274 | 0.001896458 | 0.024879096 |
| E2F7 | 1751.819989 | 0.727852627 | 0.209909973 | 7.63E-06 | 0.000314144 |
| ZDBF2 | 1857.328753 | 0.726953973 | 0.162442495 | 1.58E-07 | 1.26E-05 |
| ESRRB | 11.99066364 | 0.724915147 | 0.633007643 | 0.00186658 | 0.024626357 |

| | | | | | |
|---|---|---|---|---|---|
| PNLDC1 | 23.10101017 | 0.724250723 | 0.616217853 | 0.001813203 | 0.024043644 |
| RAB39B | 957.1555805 | 0.720125248 | 0.184282795 | 1.64E-06 | 9.15E-05 |
| KCNG1 | 1075.562128 | 0.719799105 | 0.198148499 | 4.49E-06 | 0.000212036 |
| PLEK | 14.77663603 | 0.718903333 | 0.616371013 | 0.001870446 | 0.024646218 |
| CYP27A1 | 47.58358777 | 0.717558174 | 0.442484257 | 0.000939771 | 0.014701955 |
| REL | 388.7146352 | 0.717547911 | 0.347647615 | 0.000383503 | 0.007202756 |
| SYT14 | 2304.306523 | 0.716915675 | 0.153796304 | 7.18E-08 | 6.64E-06 |
| MAP3K2 | 6826.205165 | 0.714990625 | 0.235319496 | 3.14E-05 | 0.001006056 |
| TGDS | 1191.806834 | 0.714564246 | 0.161061038 | 1.97E-07 | 1.53E-05 |
| DRAM1 | 485.793116 | 0.714358417 | 0.249915026 | 5.31E-05 | 0.0015304 |
| IARS1 | 11671.56378 | 0.712713289 | 0.140171146 | 9.65E-09 | 1.13E-06 |
| NOTUM | 90.99759405 | 0.709395003 | 0.417979602 | 0.00084532 | 0.013528357 |
| TEX14 | 37.16893609 | 0.708760003 | 0.465307484 | 0.001161953 | 0.017198715 |
| HKDC1 | 22.29787326 | 0.708391853 | 0.487990068 | 0.001312182 | 0.018860923 |
| KCNT2 | 1811.329412 | 0.706548898 | 0.190804159 | 3.71E-06 | 0.000180248 |
| ZNF441 | 629.9295451 | 0.705592563 | 0.196235479 | 5.44E-06 | 0.000243564 |
| ACSBG2 | 28.72062724 | 0.70557234 | 0.455894813 | 0.001128933 | 0.016842013 |
| LGALS9 | 110.5264663 | 0.705206269 | 0.283688172 | 0.000149781 | 0.003437561 |
| ERN1 | 2074.317555 | 0.702918335 | 0.201823454 | 8.11E-06 | 0.000329349 |
| KDM3AP1 | 2.582382957 | 0.696569428 | 1.099633805 | 0.000927165 | 0.014581208 |
| ZNF408 | 759.7703794 | 0.696326659 | 0.184552837 | 3.01E-06 | 0.000151283 |
| SLC16A9 | 1009.464291 | 0.694002452 | 0.308845186 | 0.000282156 | 0.005699054 |
| DCAF4L1 | 159.0383705 | 0.693017096 | 0.392085158 | 0.000794194 | 0.012847987 |
| MSMO1 | 3264.087363 | 0.691708628 | 0.196405474 | 7.47E-06 | 0.00031001 |
| LRRC66 | 37.65542727 | 0.691580791 | 0.343479291 | 0.000482429 | 0.008589547 |
| SEC23B | 5070.055628 | 0.691464426 | 0.11618014 | 9.26E-11 | 1.68E-08 |
| SNX16 | 728.30325 | 0.689423206 | 0.190171103 | 5.27E-06 | 0.000238288 |
| CCDC174 | 1511.034764 | 0.686606185 | 0.14971414 | 1.15E-07 | 9.76E-06 |
| ZNF550 | 1281.72108 | 0.685853896 | 0.164811497 | 7.07E-07 | 4.46E-05 |
| ZNF584 | 741.0664309 | 0.685775877 | 0.12866518 | 3.09E-09 | 4.11E-07 |
| LST1 | 9.193690872 | 0.685206239 | 0.828575255 | 0.002333262 | 0.029023333 |
| STK32A | 78.34300706 | 0.685193745 | 0.367340599 | 0.00067781 | 0.011333915 |
| TICAM2 | 17.49161949 | 0.683626697 | 0.490643921 | 0.00159979 | 0.021752354 |
| CCDC30 | 326.2359498 | 0.683383338 | 0.190343377 | 6.11E-06 | 0.000266063 |
| GALK2 | 553.7140216 | 0.682535878 | 0.148348776 | 1.10E-07 | 9.40E-06 |
| PARD6B | 220.237924 | 0.681705678 | 0.249512156 | 8.77E-05 | 0.002259567 |
| RAB32 | 2090.351666 | 0.680590853 | 0.140839738 | 3.80E-08 | 3.73E-06 |
| HLA-DMB | 2.741042458 | 0.677600573 | 1.091854434 | 0.0009439 | 0.014744472 |
| FADS2 | 14485.05111 | 0.675050727 | 0.186774483 | 5.83E-06 | 0.000256165 |
| UNC13A | 2439.349078 | 0.674036642 | 0.177413208 | 3.01E-06 | 0.000151283 |
| TMEM135 | 1588.907917 | 0.672744501 | 0.171257896 | 1.86E-06 | 0.000100863 |
| ALG2 | 1706.041061 | 0.672612118 | 0.139345296 | 4.02E-08 | 3.91E-06 |
| PLD6 | 316.3907936 | 0.669664443 | 0.273365784 | 0.000194255 | 0.004252647 |
| TRAF3 | 2905.440735 | 0.664470463 | 0.153183233 | 3.71E-07 | 2.62E-05 |
| DDX59-AS1 | 17.61434841 | 0.664434452 | 0.49030638 | 0.001844429 | 0.024380319 |
| ANO9 | 124.7796868 | 0.66438308 | 0.716461441 | 0.002722893 | 0.032663763 |
| IDI2-AS1 | 258.6105941 | 0.663778708 | 0.215771718 | 3.54E-05 | 0.001115407 |
| STX11 | 8.159122055 | 0.662616446 | 0.85503388 | 0.002405675 | 0.029690671 |
| ARID3A | 851.4029254 | 0.662543691 | 0.21156652 | 3.01E-05 | 0.000972324 |
| CRISPLD2 | 553.3769218 | 0.660128321 | 0.215844785 | 3.80E-05 | 0.001168824 |

| | | | | | |
|---|---|---|---|---|---|
| NXPE2 | 3.179044155 | 0.659056719 | 1.0929445 | 0.000898511 | 0.014194699 |
| SLC38A1 | 10224.02944 | 0.658590092 | 0.162074527 | 1.16E-06 | 6.95E-05 |
| TECTA | 318.4704225 | 0.65804732 | 0.255193252 | 0.000147794 | 0.003395686 |
| UCN | 178.4429808 | 0.657836115 | 0.23463821 | 8.05E-05 | 0.002120714 |
| DNAH8 | 11.25890809 | 0.656751117 | 0.588946273 | 0.002560938 | 0.031096035 |
| NFKBIB | 1261.223423 | 0.655992922 | 0.165125019 | 1.68E-06 | 9.29E-05 |
| SRPRB | 2210.232174 | 0.653879857 | 0.114401707 | 4.21E-10 | 6.77E-08 |
| CDKN2AIP | 1744.046346 | 0.65378788 | 0.21451675 | 4.04E-05 | 0.001228962 |
| GOT1 | 1535.59364 | 0.653082597 | 0.174894206 | 4.15E-06 | 0.000198327 |
| MFSD9 | 846.8770055 | 0.650532303 | 0.149522834 | 3.69E-07 | 2.62E-05 |
| CPEB3 | 342.6980189 | 0.650241733 | 0.176987876 | 5.21E-06 | 0.000237273 |
| CPM | 36.16367146 | 0.650060699 | 0.429493298 | 0.001537897 | 0.021130913 |
| ARHGEF2 | 4822.306789 | 0.647430734 | 0.191919594 | 1.48E-05 | 0.000542753 |
| EEF1A2 | 3559.92865 | 0.646651241 | 0.182037565 | 8.12E-06 | 0.000329349 |
| EPIC1 | 239.9391347 | 0.646073505 | 0.198106913 | 2.15E-05 | 0.000739878 |
| SLC26A4 | 27.64367985 | 0.645198682 | 0.423811208 | 0.001551263 | 0.021245755 |
| SLC26A7 | 15.45379336 | 0.645184342 | 0.538015801 | 0.002471906 | 0.030243557 |
| ASS1 | 109.6454751 | 0.642372171 | 0.234225492 | 0.000102578 | 0.002556491 |
| PER1 | 1742.613251 | 0.642118107 | 0.181707802 | 8.82E-06 | 0.000352691 |
| MID1IP1 | 2171.422917 | 0.64201991 | 0.160017173 | 1.51E-06 | 8.59E-05 |
| NR1I2 | 9.784549845 | 0.641627134 | 0.752778235 | 0.002957213 | 0.034669486 |
| TBL2 | 2595.691794 | 0.639673202 | 0.134904193 | 6.70E-08 | 6.25E-06 |
| NPIPB6 | 80.5740434 | 0.639112152 | 0.277567115 | 0.000323731 | 0.006330631 |
| C14orf28 | 139.8876602 | 0.638828882 | 0.269364758 | 0.000274264 | 0.005588302 |
| RHBDD3 | 1075.227749 | 0.638069844 | 0.225235126 | 8.13E-05 | 0.002134917 |
| INPP4B | 152.9185431 | 0.637481383 | 0.263160592 | 0.000243526 | 0.005070827 |
| SERPINB8 | 272.7816545 | 0.637026158 | 0.238061486 | 0.000126237 | 0.003002803 |
| GPR35 | 188.9676962 | 0.636266991 | 0.278212262 | 0.000340833 | 0.00660324 |
| GOLGA4-AS1 | 35.10352836 | 0.636114939 | 0.406146771 | 0.001505905 | 0.020814573 |
| IDI1 | 5535.975719 | 0.636020885 | 0.157119476 | 1.34E-06 | 7.87E-05 |
| PMAIP1 | 3296.664551 | 0.634210056 | 0.24735512 | 0.000172371 | 0.003855197 |
| PSPH | 1900.443065 | 0.634049513 | 0.222739859 | 7.96E-05 | 0.002109182 |
| SDE2 | 3888.473578 | 0.633924945 | 0.136472515 | 1.06E-07 | 9.29E-06 |
| MAPT-AS1 | 16.34511494 | 0.633597771 | 0.553815334 | 0.002767989 | 0.03293986 |
| CBARP | 1437.328521 | 0.633539205 | 0.245497081 | 0.000165534 | 0.00372942 |
| FOXN2 | 2066.713095 | 0.631626913 | 0.168230561 | 4.17E-06 | 0.000198936 |
| LINC02995 | 59.80769212 | 0.631338529 | 0.3257461 | 0.000761133 | 0.012438388 |
| ZFAND3 | 4131.723586 | 0.630268774 | 0.111306144 | 6.07E-10 | 9.60E-08 |
| NAMPTP1 | 215.8668076 | 0.630172814 | 0.232800281 | 0.000119847 | 0.002888968 |
| BRINP2 | 1012.067863 | 0.630087287 | 0.189316745 | 1.85E-05 | 0.000652083 |
| SLC30A1 | 4166.149599 | 0.629904074 | 0.171601963 | 5.69E-06 | 0.000251972 |
| H3P4 | 30.83595904 | 0.629311897 | 0.402680468 | 0.00156701 | 0.021418197 |
| PDXK | 3521.260233 | 0.629245356 | 0.162902362 | 2.81E-06 | 0.000143488 |
| BHLHE41 | 102.7996917 | 0.628992238 | 0.30592135 | 0.000599283 | 0.01029279 |
| PHC3 | 2842.845302 | 0.628691608 | 0.172673187 | 6.36E-06 | 0.000274644 |
| PEG3 | 376.2044537 | 0.627690058 | 0.29551194 | 0.000519005 | 0.009116409 |
| SQSTM1 | 9929.046522 | 0.626616527 | 0.180464088 | 1.16E-05 | 0.000444002 |
| PURA | 1488.791441 | 0.625890011 | 0.148708894 | 7.23E-07 | 4.55E-05 |
| CA5B | 679.5131872 | 0.621970228 | 0.166893253 | 4.78E-06 | 0.000223282 |
| GARIN1A | 321.2249135 | 0.620524369 | 0.251986526 | 0.000239499 | 0.005001936 |

| | | | | | |
|---|---|---|---|---|---|
| XPOT | 11240.43814 | 0.620074681 | 0.14363194 | 4.67E-07 | 3.24E-05 |
| THEM5 | 27.87562014 | 0.618997156 | 0.400900353 | 0.001704357 | 0.02290573 |
| LINC01554 | 15.8384634 | 0.618639822 | 0.599455969 | 0.003258266 | 0.037225868 |
| NECTIN4 | 47.42987275 | 0.61855985 | 0.355664337 | 0.001206693 | 0.017685467 |
| RFX2 | 1076.614834 | 0.618532789 | 0.252059865 | 0.000247236 | 0.005132736 |
| ZNF503 | 1372.99485 | 0.618500727 | 0.15118592 | 1.19E-06 | 7.08E-05 |
| ZNF844 | 177.454349 | 0.618029904 | 0.444752539 | 0.002185219 | 0.027794338 |
| TAF4B | 417.7141685 | 0.617647306 | 0.162638544 | 3.73E-06 | 0.00018077 |
| CDKN1A | 7748.204446 | 0.615510645 | 0.249948348 | 0.00024554 | 0.005107682 |
| KDM7A | 2787.961188 | 0.614569157 | 0.151565173 | 1.40E-06 | 8.06E-05 |
| STARD10 | 712.9519716 | 0.614195057 | 0.182089619 | 1.70E-05 | 0.000605686 |
| TMEM236 | 12.17797843 | 0.613833675 | 0.572918205 | 0.003244024 | 0.037109785 |
| SCDP1 | 26.08319511 | 0.613666927 | 0.496201183 | 0.002745526 | 0.032828692 |
| C5orf67 | 8.542488371 | 0.613236252 | 0.742574735 | 0.003345104 | 0.038051468 |
| KTN1-AS1 | 181.0523899 | 0.612353329 | 0.237097585 | 0.000182488 | 0.004024574 |
| FOXO6 | 246.147189 | 0.612068554 | 0.253895612 | 0.000284751 | 0.005729313 |
| FDFT1 | 8383.527776 | 0.611032352 | 0.158054362 | 2.95E-06 | 0.000149294 |
| OPTN | 3875.042745 | 0.610810628 | 0.176600344 | 1.29E-05 | 0.000482033 |
| DUSP26 | 447.5778627 | 0.610390678 | 0.261815482 | 0.000349361 | 0.00673721 |
| SAMD11 | 5096.828958 | 0.610187517 | 0.254922082 | 0.000300001 | 0.005938884 |
| CARD14 | 79.18115901 | 0.609785369 | 0.251266864 | 0.000276601 | 0.005612675 |
| MEX3B | 3737.803301 | 0.60783474 | 0.190638113 | 3.18E-05 | 0.001015128 |
| SMIM14 | 2584.779321 | 0.607235551 | 0.141570379 | 5.44E-07 | 3.60E-05 |
| OSBP | 3694.058727 | 0.607219601 | 0.128138628 | 7.35E-08 | 6.77E-06 |
| NUCB2 | 2873.332768 | 0.606641691 | 0.195003355 | 4.08E-05 | 0.001239949 |
| GPCPD1 | 1482.289874 | 0.606004405 | 0.149395863 | 1.42E-06 | 8.18E-05 |
| PTGFR | 380.4674403 | 0.605121974 | 0.306891985 | 0.000814257 | 0.013111615 |
| AKIRIN1 | 5542.63862 | 0.604445428 | 0.117381376 | 9.93E-09 | 1.15E-06 |
| KLHDC7B | 6.011331215 | 0.603879748 | 0.905323326 | 0.002388863 | 0.029574036 |
| AARS1 | 5647.577565 | 0.603687156 | 0.123757819 | 3.82E-08 | 3.73E-06 |
| SERP1 | 7700.108951 | 0.601911685 | 0.127524967 | 8.11E-08 | 7.37E-06 |
| NINJ1 | 655.0421565 | 0.601681201 | 0.199396452 | 5.55E-05 | 0.0015826 |
| LINC00623 | 7.19344644 | 0.601178752 | 0.671702525 | 0.003683705 | 0.040878943 |
| POU5F1P3 | 20.97278943 | 0.599414286 | 0.541535165 | 0.003380464 | 0.038411858 |
| VCPIP1 | 2152.384333 | 0.599269428 | 0.147342388 | 1.39E-06 | 8.06E-05 |
| ZNF555 | 684.9829384 | 0.598253088 | 0.173013947 | 1.35E-05 | 0.000497852 |
| EPS8L2 | 96.73124908 | 0.597444414 | 0.280856138 | 0.000603917 | 0.010329903 |
| MSTO1 | 1277.565163 | 0.597106307 | 0.175539843 | 1.64E-05 | 0.000591419 |
| PROKR1 | 10.76841155 | 0.597001125 | 0.632712871 | 0.003758907 | 0.041405466 |
| CADM3 | 93.21467993 | 0.596200989 | 0.259403329 | 0.000407219 | 0.007586773 |
| ZBTB49 | 711.7841211 | 0.595816786 | 0.138039499 | 4.98E-07 | 3.37E-05 |
| ZNF853 | 3425.274984 | 0.595475027 | 0.144943295 | 1.19E-06 | 7.05E-05 |
| CLK4 | 1217.251621 | 0.595366999 | 0.197381549 | 5.73E-05 | 0.001618158 |
| ADRA2A | 100.5788792 | 0.595316822 | 0.265894216 | 0.00047163 | 0.008484051 |
| PRKAA2 | 1162.438483 | 0.594853117 | 0.161856022 | 6.31E-06 | 0.000273406 |
| CYLD | 2897.045598 | 0.59416095 | 0.156265215 | 3.95E-06 | 0.000189532 |
| SELENOK | 2117.735107 | 0.59414406 | 0.166808193 | 9.51E-06 | 0.00037481 |
| ZBTB34 | 2439.741807 | 0.594102989 | 0.111599761 | 4.08E-09 | 5.19E-07 |
| PVT1 | 464.7114118 | 0.593701405 | 0.177540853 | 2.02E-05 | 0.000702761 |
| TMED7-TICAM2 | 10.05721427 | 0.592753397 | 0.617892244 | 0.003826393 | 0.041992334 |

| | | | | | |
|---|---|---|---|---|---|
| NPC1 | 2243.753912 | 0.592310897 | 0.158015535 | 4.85E-06 | 0.000225776 |
| CBS | 585.0407021 | 0.59218694 | 0.281395071 | 0.000653804 | 0.011011848 |
| HIF1A-AS1 | 33.79032123 | 0.590598517 | 0.389569428 | 0.002072104 | 0.026631322 |
| BMAL1 | 1990.606733 | 0.59055382 | 0.15727891 | 4.76E-06 | 0.000223109 |
| LARP6 | 1783.439244 | 0.589545825 | 0.189430586 | 4.36E-05 | 0.001309154 |
| ZNF597 | 786.9342243 | 0.588939356 | 0.157893224 | 5.25E-06 | 0.000238288 |
| ZNF770 | 2424.357205 | 0.58794327 | 0.167845741 | 1.20E-05 | 0.000457255 |
| ARL5B | 5960.798372 | 0.587301897 | 0.153226227 | 3.58E-06 | 0.000175188 |
| IER5 | 2907.405188 | 0.587291545 | 0.201483322 | 8.05E-05 | 0.002120714 |
| LDLR | 5066.97208 | 0.587250987 | 0.253155682 | 0.00040495 | 0.007557988 |
| APBA3 | 682.6738205 | 0.587078863 | 0.17390642 | 1.86E-05 | 0.000652083 |
| RFPL3S | 54.27769477 | 0.586857163 | 0.409495299 | 0.002407182 | 0.029690671 |
| TIMM44 | 1822.717087 | 0.586792268 | 0.132866284 | 3.29E-07 | 2.37E-05 |
| TTLL9 | 92.69308027 | 0.586057788 | 0.231435826 | 0.000237171 | 0.004963245 |
| ZNF324 | 1022.030408 | 0.585499854 | 0.164572366 | 9.93E-06 | 0.000388862 |
| NIPAL1 | 67.26822417 | 0.584949527 | 0.288577518 | 0.000806818 | 0.01303201 |
| PPIEL | 12.73617456 | 0.584424477 | 0.627705142 | 0.004015879 | 0.043501884 |
| RBM15 | 919.1951091 | 0.582458025 | 0.156720275 | 5.63E-06 | 0.00025 |
| GFPT1 | 5141.339374 | 0.582171624 | 0.205549457 | 0.000104987 | 0.002607208 |
| APLF | 509.0096483 | 0.58158518 | 0.202951486 | 9.55E-05 | 0.002403905 |
| LCMT1-AS2 | 27.0072519 | 0.580549143 | 0.43505464 | 0.002868895 | 0.033908807 |
| SERPINC1 | 11.21471512 | 0.580477571 | 0.569972667 | 0.003970615 | 0.043188749 |
| ACER2 | 171.5260188 | 0.577748493 | 0.25460973 | 0.000481232 | 0.008582861 |
| STX5 | 1693.701233 | 0.574612332 | 0.135662156 | 7.32E-07 | 4.59E-05 |
| ANKRD11 | 10360.03496 | 0.574441815 | 0.189488985 | 5.96E-05 | 0.001672819 |
| TEKT5 | 44.87987108 | 0.573422714 | 0.341652498 | 0.001725383 | 0.023069542 |
| ALAS1 | 1654.946413 | 0.572427439 | 0.167124623 | 1.65E-05 | 0.000593163 |
| H4C12 | 13.95506185 | 0.571816579 | 0.52849527 | 0.003975232 | 0.043195999 |
| RNF19B | 1031.323883 | 0.571246956 | 0.14178553 | 1.73E-06 | 9.55E-05 |
| MAGT1 | 5044.948275 | 0.570753446 | 0.157109203 | 7.92E-06 | 0.000324442 |
| CLCN6 | 2891.543451 | 0.569425447 | 0.170619765 | 2.25E-05 | 0.000764528 |
| LDLRAD4 | 1806.569295 | 0.56915871 | 0.209197844 | 0.000153472 | 0.003499203 |
| WDR74 | 1831.139183 | 0.567987168 | 0.157688992 | 8.94E-06 | 0.000356275 |
| TMEM151A | 196.3611747 | 0.567279421 | 0.305691764 | 0.001273822 | 0.018462027 |
| BLOC1S2 | 1911.814476 | 0.567214399 | 0.167374987 | 1.90E-05 | 0.000666784 |
| YOD1 | 1068.233663 | 0.566925315 | 0.201669574 | 0.000119484 | 0.002884888 |
| SLC2A6 | 1110.811586 | 0.566633862 | 0.217707748 | 0.000215698 | 0.004587428 |
| LINC00930 | 3.230878696 | 0.566243647 | 0.953344898 | 0.001910423 | 0.025030857 |
| LINC01338 | 28.05219171 | 0.565700474 | 0.432740702 | 0.003224385 | 0.03693981 |
| SLC35B1 | 4096.491565 | 0.56553354 | 0.145480514 | 3.07E-06 | 0.000153406 |
| CTH | 282.0139526 | 0.565442101 | 0.229752334 | 0.000316726 | 0.006222787 |
| MGAT4A | 150.5020375 | 0.565156714 | 0.25969025 | 0.000641795 | 0.010870951 |
| HSD17B13 | 75.98948729 | 0.563808028 | 0.265216347 | 0.000726567 | 0.012005027 |
| RAB11FIP1 | 507.4853259 | 0.563757034 | 0.172189587 | 2.84E-05 | 0.000931721 |
| CANX | 39192.00762 | 0.563339109 | 0.173282267 | 3.07E-05 | 0.000990484 |
| SEC63 | 8048.137005 | 0.562972159 | 0.175706991 | 3.59E-05 | 0.001121656 |
| DRD4 | 108.0454731 | 0.561397197 | 0.289065568 | 0.001102786 | 0.016547944 |
| ABTB2 | 383.3736374 | 0.56124176 | 0.202426396 | 0.000137108 | 0.003213816 |
| NDUFAF4P2 | 7.818293477 | 0.560906996 | 0.740106577 | 0.004107633 | 0.044152302 |
| CCDC33 | 178.1428206 | 0.560383168 | 0.242865481 | 0.000480314 | 0.008573806 |

| | | | | | |
|---|---|---|---|---|---|
| PDIA2 | 106.5703368 | 0.559732773 | 0.25064732 | 0.00057751 | 0.010009379 |
| PTPRH | 135.0963739 | 0.55950785 | 0.260802148 | 0.000711228 | 0.011788891 |
| ESAM-AS1 | 120.9091786 | 0.559027596 | 0.27179092 | 0.000871813 | 0.01383572 |
| PPP1R15B | 7039.361932 | 0.55793079 | 0.131014014 | 6.84E-07 | 4.37E-05 |
| SMTNL1 | 18.92070371 | 0.557554944 | 0.550511226 | 0.004494392 | 0.047121177 |
| ABCC2 | 108.6351769 | 0.557432358 | 0.297810983 | 0.00130449 | 0.018789149 |
| RN7SL34P | 12.85749267 | 0.556902743 | 0.6770305 | 0.004526066 | 0.047334449 |
| ASB7 | 1001.853922 | 0.556479137 | 0.197898677 | 0.000124843 | 0.002979821 |
| MTHFR | 1569.628718 | 0.555742498 | 0.186983221 | 7.73E-05 | 0.002058765 |
| TCIM | 327.2310041 | 0.55568486 | 0.53671638 | 0.004479795 | 0.047015332 |
| PPM1K | 1250.633348 | 0.555181971 | 0.156804872 | 1.16E-05 | 0.000444002 |
| SBSN | 8.873668923 | 0.555167618 | 0.588400314 | 0.004678764 | 0.048514836 |
| SLC22A20P | 43.24113907 | 0.554696617 | 0.336085578 | 0.002019458 | 0.026083101 |
| RAB33A | 356.0594718 | 0.554590746 | 0.33807403 | 0.002056772 | 0.026490066 |
| SCAND3 | 565.3906667 | 0.553851569 | 0.245860492 | 0.000568501 | 0.009877818 |
| PTPRN | 1390.461398 | 0.55344124 | 0.205640146 | 0.000179336 | 0.003976052 |
| GAPDHP48 | 29.68705166 | 0.552969625 | 0.538281412 | 0.004569037 | 0.047688327 |
| NXF1 | 4416.966653 | 0.552699963 | 0.122091434 | 2.11E-07 | 1.62E-05 |
| CNST | 3564.145933 | 0.552697882 | 0.145424799 | 4.43E-06 | 0.000209796 |
| NFXL1 | 1437.737693 | 0.551685736 | 0.205165184 | 0.000181994 | 0.004018022 |
| SLC25A41 | 12.80326256 | 0.551460327 | 0.548968057 | 0.004665815 | 0.048408116 |
| GRM3-AS1 | 58.0565855 | 0.551308014 | 0.295623661 | 0.001370264 | 0.019474724 |
| CCDC62 | 56.12138075 | 0.550771607 | 0.327168761 | 0.00194772 | 0.02545565 |
| PEX5L | 93.2359254 | 0.550150232 | 0.29160862 | 0.001319754 | 0.018956721 |
| IGIP | 381.9386348 | 0.549751015 | 0.215929266 | 0.000273159 | 0.005576654 |
| EIF5 | 11669.11265 | 0.54898328 | 0.174999221 | 4.71E-05 | 0.00137722 |
| SNORC | 57.90565481 | 0.548566546 | 0.391628393 | 0.003126685 | 0.036157707 |
| RIOK3 | 4425.223112 | 0.548558113 | 0.150357032 | 7.96E-06 | 0.000324796 |
| SLC12A7 | 188.1799764 | 0.548306044 | 0.358620492 | 0.002575661 | 0.031238487 |
| CCNP | 62.96786851 | 0.548239372 | 0.359983216 | 0.00260156 | 0.031515997 |
| ZNF274 | 574.0616078 | 0.547216193 | 0.182571795 | 7.40E-05 | 0.001991061 |
| TRIM11 | 2183.475288 | 0.546734185 | 0.186790556 | 9.21E-05 | 0.002346655 |
| GMPPB | 1185.757553 | 0.546005435 | 0.196614463 | 0.000144861 | 0.003365316 |
| SRP19 | 1437.028083 | 0.545491394 | 0.15523118 | 1.31E-05 | 0.000487293 |
| ZNF319 | 633.2700914 | 0.545154643 | 0.169249839 | 3.62E-05 | 0.001123465 |
| TOB1 | 2106.497883 | 0.544927467 | 0.200382867 | 0.000172072 | 0.003855197 |
| LINC00622 | 212.3347541 | 0.544249993 | 0.325625198 | 0.002068569 | 0.026617823 |
| SREBF2 | 5416.341018 | 0.544007737 | 0.189599399 | 0.000111088 | 0.002735943 |
| PIM2 | 456.9251824 | 0.543528715 | 0.139438217 | 3.09E-06 | 0.000154105 |
| DNAI4 | 314.7857826 | 0.541716815 | 0.225661035 | 0.000418099 | 0.007727435 |
| NCR3LG1 | 938.2898656 | 0.539599926 | 0.253979497 | 0.00083295 | 0.013361103 |
| RHBDD2 | 4129.693755 | 0.539577814 | 0.179275813 | 7.33E-05 | 0.001977276 |
| SLC25A33 | 1361.454425 | 0.539272419 | 0.188316836 | 0.00011528 | 0.002812632 |
| MRPL49 | 2495.89522 | 0.539135803 | 0.127620699 | 8.12E-07 | 5.05E-05 |
| ZDHHC11 | 136.6812086 | 0.538544041 | 0.240051659 | 0.000631299 | 0.010715949 |
| PIM3 | 1447.965572 | 0.538147399 | 0.203334878 | 0.000218531 | 0.004642959 |
| TRABD2A | 92.57818761 | 0.537030371 | 0.329269713 | 0.002320113 | 0.028963279 |
| ARID3B | 895.2765008 | 0.536841383 | 0.205277729 | 0.00024019 | 0.005006347 |
| SYNDIG1L | 3.075651877 | 0.536498906 | 0.921638912 | 0.002237978 | 0.028253958 |
| ID2-AS1 | 632.2950693 | 0.536448493 | 0.169516076 | 4.50E-05 | 0.001343859 |

| | | | | | |
|---|---|---|---|---|---|
| ARHGEF16 | 97.10130056 | 0.536436168 | 0.271785361 | 0.001189098 | 0.017488947 |
| WBP1LP2 | 24.0587767 | 0.535437711 | 0.421858027 | 0.004015579 | 0.043501884 |
| PLPBP | 1904.333675 | 0.533861389 | 0.133551965 | 2.11E-06 | 0.000112602 |
| RNU6-890P | 5.971055964 | 0.533622368 | 0.716908164 | 0.004735544 | 0.048840412 |
| TMEM39A | 3153.084636 | 0.533596501 | 0.141248256 | 5.06E-06 | 0.000231571 |
| GCC1 | 2413.58226 | 0.532570246 | 0.135941234 | 2.92E-06 | 0.00014852 |
| FADS1 | 7491.204833 | 0.532400439 | 0.152141775 | 1.44E-05 | 0.000526012 |
| IL17C | 4.516149397 | 0.530632643 | 0.783704533 | 0.004098111 | 0.044095945 |
| MAP2K3 | 1111.533226 | 0.530608726 | 0.226927912 | 0.000519876 | 0.009124037 |
| BRPF3 | 2614.194256 | 0.528788712 | 0.163241216 | 3.60E-05 | 0.001122009 |
| PTPN6 | 67.53595011 | 0.528621751 | 0.289069918 | 0.001684008 | 0.022676015 |
| DHDH | 34.50103706 | 0.528125903 | 0.373739786 | 0.003467554 | 0.039152224 |
| LONP1 | 6454.680439 | 0.527730905 | 0.16379782 | 3.84E-05 | 0.001178667 |
| BEND3P3 | 84.76135912 | 0.527332542 | 0.306380785 | 0.002089787 | 0.026825571 |
| OAS2 | 7.373102156 | 0.527210997 | 0.786636625 | 0.004098168 | 0.044095945 |
| GPR179 | 16.25657565 | 0.527067695 | 0.450198376 | 0.00470131 | 0.048631427 |
| SLC22A4 | 198.407716 | 0.525698051 | 0.298850956 | 0.001964453 | 0.025578306 |
| CHURC1-FNTB | 3.190539275 | 0.522411676 | 0.914193264 | 0.002299846 | 0.028813612 |
| LAMB2P1 | 61.73315746 | 0.522096991 | 0.257181498 | 0.001141138 | 0.016974832 |
| CDRT4 | 5.244178451 | 0.521998769 | 0.748965728 | 0.004608599 | 0.048029234 |
| COX19 | 986.6174424 | 0.521748272 | 0.167247115 | 5.51E-05 | 0.00157402 |
| H3-7 | 463.1997084 | 0.521677805 | 0.218245004 | 0.000476094 | 0.008534964 |
| CDK20 | 602.7012712 | 0.521530454 | 0.16625659 | 5.21E-05 | 0.001506577 |
| CCDC81 | 70.29812197 | 0.520931756 | 0.266612439 | 0.001358548 | 0.019354217 |
| SH2D5 | 42.84519374 | 0.520850047 | 0.335490089 | 0.002924774 | 0.034432864 |
| ABHD4 | 799.8272513 | 0.520748498 | 0.168868437 | 6.22E-05 | 0.001724856 |
| DEDD2 | 750.4238643 | 0.519753197 | 0.234072099 | 0.000739608 | 0.012172358 |
| ZNF620 | 439.8516703 | 0.519327494 | 0.193650393 | 0.000215632 | 0.004587428 |
| TRIM23 | 1586.947076 | 0.519311643 | 0.156553737 | 2.85E-05 | 0.000932455 |
| AACS | 2159.929362 | 0.519139182 | 0.173847442 | 8.56E-05 | 0.002224874 |
| AKAP8L | 2480.524496 | 0.518711592 | 0.123209153 | 8.88E-07 | 5.45E-05 |
| RAB5A | 3842.98523 | 0.518266335 | 0.162606395 | 4.46E-05 | 0.001333377 |
| LYG1 | 87.35212797 | 0.517604393 | 0.247577535 | 0.001019891 | 0.015662073 |
| LNX2 | 829.8052392 | 0.517044708 | 0.146110027 | 1.30E-05 | 0.000485186 |
| ZNF467 | 298.7013426 | 0.516701821 | 0.32443397 | 0.002806303 | 0.033338817 |
| MIA3 | 6799.750043 | 0.51657355 | 0.181373249 | 0.000133056 | 0.00313499 |
| SCYL1 | 2626.936619 | 0.516182804 | 0.159854794 | 3.90E-05 | 0.001193039 |
| GOLPH3L | 1554.800585 | 0.514952267 | 0.158793136 | 3.73E-05 | 0.001149556 |
| SQLE | 5021.266176 | 0.514885368 | 0.147055858 | 1.50E-05 | 0.000545889 |
| TMEM198 | 668.1634027 | 0.514835651 | 0.225114831 | 0.000643793 | 0.010887142 |
| MROH8 | 180.3776225 | 0.514006355 | 0.177536147 | 0.000116341 | 0.002831921 |
| MYLK-AS2 | 4.439163089 | 0.513627052 | 0.766462239 | 0.004505159 | 0.047186681 |
| LSAMP-AS1 | 11.59198312 | 0.513494565 | 0.743526743 | 0.004797432 | 0.049308255 |
| ANKRD11P2 | 103.1526468 | 0.513172299 | 0.205206567 | 0.000369853 | 0.006996712 |
| ELF4 | 713.9881519 | 0.513149106 | 0.175415224 | 0.000106287 | 0.002630103 |
| FYTTD1P1 | 2.786812437 | 0.512730748 | 0.948250746 | 0.001737335 | 0.023214487 |
| ODAD4 | 298.1860789 | 0.512550454 | 0.189425266 | 0.000207367 | 0.004463559 |
| ZNF805 | 878.4017021 | 0.512305173 | 0.129656646 | 2.65E-06 | 0.000136639 |
| FAM9C | 2.355000604 | 0.512166335 | 0.931436003 | 0.001986529 | 0.025801401 |
| DIRAS3 | 94.69043908 | 0.511538988 | 0.25516094 | 0.001283979 | 0.018544878 |

| | | | | | |
|---|---|---|---|---|---|
| CSF1R | 36.74645256 | 0.511427368 | 0.356421226 | 0.003723841 | 0.041149427 |
| HAPSTR1 | 4539.832261 | 0.51114782 | 0.170514136 | 8.52E-05 | 0.002215895 |
| PREB | 3224.675442 | 0.511063305 | 0.150721582 | 2.26E-05 | 0.00076487 |
| CDC14A | 476.4228846 | 0.511043287 | 0.159001095 | 4.17E-05 | 0.001261023 |
| RAP1GAP | 894.9363042 | 0.510602311 | 0.195733569 | 0.000277073 | 0.005612675 |
| JUN | 2655.045712 | 0.510114459 | 0.202372947 | 0.000355482 | 0.006823742 |
| CALCB | 173.4315049 | 0.50996921 | 0.256366504 | 0.001341406 | 0.019160831 |
| FNIP1 | 2969.908072 | 0.508135024 | 0.153201765 | 2.96E-05 | 0.000963355 |
| KCTD5 | 2538.122606 | 0.50810313 | 0.1468414 | 1.77E-05 | 0.000631546 |
| RHBDF2 | 1078.014606 | 0.507782156 | 0.180668362 | 0.00015491 | 0.003520457 |
| ARK2N | 2117.304176 | 0.507684136 | 0.1375546 | 7.50E-06 | 0.000310192 |
| BRD2 | 12234.92742 | 0.507385038 | 0.137886149 | 7.83E-06 | 0.000321306 |
| KIF26A | 4730.631317 | 0.506966624 | 0.266764485 | 0.00165482 | 0.022369527 |
| WHAMM | 664.1165561 | 0.506519939 | 0.148700372 | 2.16E-05 | 0.000743228 |
| RWDD2A | 862.1770868 | 0.505996151 | 0.16020545 | 5.13E-05 | 0.001484698 |
| RIPK2 | 1082.638448 | 0.50543108 | 0.210411644 | 0.000502 | 0.008869943 |
| STARD7-AS1 | 344.7856647 | 0.50454842 | 0.172992643 | 0.000113154 | 0.002770481 |
| MDM2 | 6399.665737 | 0.504003912 | 0.185191527 | 0.000205801 | 0.004444838 |
| PLEKHM2 | 3998.020357 | 0.503022606 | 0.17115347 | 0.000106082 | 0.002628132 |
| MCEE | 300.1208888 | 0.502952592 | 0.19896113 | 0.000360942 | 0.006896866 |
| NAPA-AS1 | 33.3323373 | 0.500186205 | 0.337134066 | 0.003716681 | 0.041135604 |
| ASB3 | 1644.196255 | 0.5001391 | 0.160777578 | 6.14E-05 | 0.001708824 |
| TIGD3 | 59.14727909 | 0.499599722 | 0.279707045 | 0.002187093 | 0.027801241 |
| PARD6A | 186.3615632 | 0.499228474 | 0.191949414 | 0.000299115 | 0.005932595 |
| GOLGA5 | 3526.171908 | 0.498979031 | 0.18578401 | 0.000234013 | 0.004916852 |
| RPN1 | 8704.694206 | 0.497692934 | 0.125322088 | 2.50E-06 | 0.000130099 |
| SPTY2D1 | 2284.724277 | 0.497011454 | 0.129506256 | 4.31E-06 | 0.000205275 |
| PINLYP | 83.58189507 | 0.496256442 | 0.242025167 | 0.001266061 | 0.018387821 |
| ATG101 | 1068.673952 | 0.495676484 | 0.14951154 | 3.09E-05 | 0.0009938 |
| PI4K2A | 1643.578322 | 0.495292633 | 0.138727731 | 1.22E-05 | 0.000464552 |
| REM2 | 62.92871656 | 0.495160793 | 0.285606388 | 0.002490119 | 0.030399668 |
| IPMK | 1233.487301 | 0.495052872 | 0.20117064 | 0.000451841 | 0.00821514 |
| CD14 | 40.28175236 | 0.494344901 | 0.339957512 | 0.004035808 | 0.043648921 |
| SPATA17 | 189.4881852 | 0.493870888 | 0.272223444 | 0.002142779 | 0.027354489 |
| BST2 | 57.51936826 | 0.492178845 | 0.321962009 | 0.003645402 | 0.040626585 |
| DYM | 2307.884777 | 0.491907073 | 0.123343008 | 2.36E-06 | 0.000123828 |
| CSNK2A2 | 1797.056169 | 0.491124084 | 0.126925419 | 3.84E-06 | 0.00018588 |
| CES2 | 1523.310573 | 0.490618378 | 0.152855 | 4.54E-05 | 0.001350235 |
| EPB41L4A-AS1 | 850.3724136 | 0.489959586 | 0.148186947 | 3.25E-05 | 0.001035381 |
| LPXN | 307.6579407 | 0.489526056 | 0.227521232 | 0.001035948 | 0.015827188 |
| CPEB2 | 1601.2831 | 0.488873874 | 0.162065324 | 8.75E-05 | 0.002257814 |
| HM13 | 5260.080581 | 0.488330382 | 0.128202093 | 4.93E-06 | 0.000227463 |
| SMAP1 | 698.4339876 | 0.486301006 | 0.123394828 | 2.90E-06 | 0.000147584 |
| GADD45A | 3121.630554 | 0.485696226 | 0.164016977 | 0.000105986 | 0.002628132 |
| EPB41 | 4171.45852 | 0.485021232 | 0.135437044 | 1.21E-05 | 0.000459942 |
| H2BC26 | 98.94809617 | 0.484724636 | 0.308720793 | 0.003574431 | 0.039984993 |
| NFASC | 485.0395068 | 0.482713133 | 0.202988832 | 0.000602822 | 0.010319629 |
| SERTAD1 | 392.8312893 | 0.482605483 | 0.235268186 | 0.001369084 | 0.019474724 |
| ITPKC | 735.476948 | 0.480062312 | 0.170595476 | 0.000172934 | 0.003858683 |
| POLR3C | 3290.536895 | 0.479055203 | 0.156126083 | 7.66E-05 | 0.002044752 |

| | | | | | |
|---|---|---|---|---|---|
| R3HDML-AS1 | 3.066067648 | 0.47902798 | 0.870124045 | 0.002864618 | 0.033896621 |
| LMCD1-AS1 | 66.25484516 | 0.478024549 | 0.256023619 | 0.002120245 | 0.027149793 |
| C2CD2L | 1258.77876 | 0.477859586 | 0.141699096 | 2.67E-05 | 0.000884363 |
| MINAR1 | 540.9442254 | 0.477586481 | 0.173622967 | 0.000212457 | 0.004537554 |
| AP3D1 | 10362.77652 | 0.477222205 | 0.156400103 | 8.16E-05 | 0.002139077 |
| NPAS3 | 166.9644587 | 0.477134706 | 0.255773437 | 0.002139514 | 0.02734624 |
| LINC01473 | 51.10494259 | 0.47703624 | 0.286315937 | 0.003174728 | 0.036531239 |
| GRPEL2 | 2177.644111 | 0.476419304 | 0.196451346 | 0.000548447 | 0.009593236 |
| CCDC154 | 106.7707292 | 0.47408479 | 0.252579936 | 0.002128056 | 0.027233124 |
| LPIN1 | 5855.983185 | 0.472978642 | 0.158214162 | 0.000101827 | 0.002543842 |
| CDHR1 | 144.0968442 | 0.472600358 | 0.327446969 | 0.004734787 | 0.048840412 |
| IRF5 | 119.3087841 | 0.472075444 | 0.229506374 | 0.001438197 | 0.02018598 |
| AIRIM | 1531.475062 | 0.470972224 | 0.151134428 | 6.72E-05 | 0.001847404 |
| PGPEP1 | 1898.80323 | 0.470826089 | 0.200010083 | 0.000684992 | 0.011426556 |
| PKD1P5 | 258.1725152 | 0.470555276 | 0.208938901 | 0.000896511 | 0.014184567 |
| LMBRD2 | 1976.945189 | 0.469917328 | 0.16405916 | 0.00015443 | 0.003513369 |
| ZNF697 | 1862.14553 | 0.469721201 | 0.136817332 | 2.20E-05 | 0.00074818 |
| PER2 | 256.9864237 | 0.469348381 | 0.252568083 | 0.0022809 | 0.028644983 |
| DERL2 | 3255.676128 | 0.469217939 | 0.138370124 | 2.57E-05 | 0.00085669 |
| UBQLNL | 2.675090567 | 0.468578702 | 0.899945356 | 0.002192781 | 0.027822747 |
| STMN4 | 240.7211836 | 0.468520011 | 0.223057507 | 0.001324011 | 0.018991741 |
| YAF2 | 1492.351467 | 0.468471171 | 0.184492074 | 0.000414871 | 0.007690229 |
| ZNF622 | 1752.560514 | 0.468270761 | 0.152577263 | 7.97E-05 | 0.002109182 |
| C1orf52 | 1984.052996 | 0.467876524 | 0.135807587 | 2.11E-05 | 0.000729619 |
| POLH | 1601.321397 | 0.467073817 | 0.138066913 | 2.66E-05 | 0.000882996 |
| SLC29A3 | 694.7407785 | 0.46646778 | 0.157527939 | 0.000114765 | 0.002806632 |
| RYBP | 4584.102918 | 0.465841576 | 0.18014793 | 0.000367332 | 0.006969604 |
| TFEB | 215.1447943 | 0.465520673 | 0.223906547 | 0.00141866 | 0.019938571 |
| MED26 | 235.7838035 | 0.465302959 | 0.209802848 | 0.001008227 | 0.015528625 |
| ZNF800 | 2221.573371 | 0.465166389 | 0.154051217 | 9.52E-05 | 0.002401323 |
| PKN1 | 6568.954143 | 0.464577404 | 0.17327443 | 0.000278688 | 0.005639913 |
| AJUBA | 1264.262372 | 0.464131308 | 0.160682815 | 0.000146643 | 0.00338787 |
| PDP2 | 1280.753747 | 0.463000209 | 0.202140843 | 0.000846245 | 0.0135328 |
| SRP68 | 6078.724795 | 0.462221485 | 0.118644709 | 3.67E-06 | 0.000178889 |
| PACSIN2 | 1379.130146 | 0.462221348 | 0.205946765 | 0.000958421 | 0.014915517 |
| SAV1 | 1604.445024 | 0.461798767 | 0.14324559 | 4.80E-05 | 0.001399727 |
| SLC25A34 | 239.5980015 | 0.460911448 | 0.206264302 | 0.000990268 | 0.015297143 |
| MAP3K2-DT | 224.5810267 | 0.460856793 | 0.280773824 | 0.003695863 | 0.040992088 |
| MMAB | 1293.216858 | 0.460287806 | 0.208399888 | 0.001061901 | 0.01611759 |
| COG3 | 2981.046773 | 0.459884666 | 0.161567158 | 0.000170896 | 0.003833696 |
| GSDMB | 513.8018263 | 0.459770704 | 0.156672115 | 0.000128778 | 0.003045902 |
| SLC8A2 | 252.5859552 | 0.459763711 | 0.202123779 | 0.000898175 | 0.014194699 |
| DHCR24 | 4485.411296 | 0.458520353 | 0.227192555 | 0.001708706 | 0.022949404 |
| PCYT2 | 2595.233002 | 0.458408668 | 0.208630966 | 0.001105138 | 0.016569143 |
| PRKAB1 | 2053.944804 | 0.458297003 | 0.121128388 | 5.87E-06 | 0.000257526 |
| TRMT44 | 570.4703364 | 0.458067408 | 0.152236259 | 0.000101579 | 0.00254068 |
| PFN1P2 | 62.31065345 | 0.45762554 | 0.300064794 | 0.004616492 | 0.048050145 |
| CACNA1B | 779.2737851 | 0.457468601 | 0.21064008 | 0.001184826 | 0.017450706 |
| ZNF501 | 255.2305509 | 0.456671075 | 0.261623138 | 0.003118007 | 0.036097322 |
| SOWAHC | 979.9643987 | 0.456436283 | 0.150385771 | 9.37E-05 | 0.002378156 |

| | | | | | |
|---|---|---|---|---|---|
| HMGCR | 6709.07046 | 0.455620458 | 0.159894123 | 0.00017251 | 0.003855197 |
| DNAJC6 | 2511.168109 | 0.454991187 | 0.156327311 | 0.000142214 | 0.003322296 |
| SYP | 944.0642035 | 0.453450383 | 0.162608451 | 0.000211203 | 0.004524074 |
| ADPRHL1 | 395.9334816 | 0.452395031 | 0.178615734 | 0.000458796 | 0.008296067 |
| TAB2 | 5885.021413 | 0.450892658 | 0.160251504 | 0.000197548 | 0.004320193 |
| EXOC8 | 1246.042472 | 0.450687181 | 0.160822373 | 0.000204891 | 0.00442977 |
| ZMAT3 | 6758.467597 | 0.450485911 | 0.165586134 | 0.000264565 | 0.005417102 |
| KCNJ14 | 189.7524333 | 0.44998864 | 0.186431087 | 0.0006513 | 0.010987736 |
| ELL | 559.6454675 | 0.448218813 | 0.171415809 | 0.000368448 | 0.006982792 |
| ANKRA2 | 1060.573999 | 0.447363075 | 0.153533392 | 0.000145606 | 0.003375129 |
| KLHL36 | 2351.372948 | 0.446665708 | 0.186031135 | 0.000686927 | 0.01144969 |
| NANP | 706.8433517 | 0.446574006 | 0.189392728 | 0.000773764 | 0.012566142 |
| INKA2 | 452.02626 | 0.446417004 | 0.197887932 | 0.001017552 | 0.01563765 |
| AUH | 553.1641266 | 0.445448993 | 0.148350069 | 0.000109507 | 0.002703363 |
| NUFIP2 | 6392.794747 | 0.444564607 | 0.142926805 | 7.63E-05 | 0.002041891 |
| RINL | 97.70407744 | 0.444162519 | 0.24613231 | 0.003002125 | 0.035008034 |
| CXXC4 | 989.1315725 | 0.444090388 | 0.214503298 | 0.001648045 | 0.022306813 |
| LINC01415 | 141.6658531 | 0.443015714 | 0.245314577 | 0.003016344 | 0.035153315 |
| RNF111 | 3474.310105 | 0.442394772 | 0.195254492 | 0.001014738 | 0.0156174 |
| SNHG15 | 1296.255034 | 0.442388528 | 0.195572024 | 0.001024759 | 0.015713723 |
| PDIA4 | 20707.61355 | 0.442153229 | 0.142041079 | 7.64E-05 | 0.00204324 |
| RBM44 | 118.2785678 | 0.441859677 | 0.243206679 | 0.002970625 | 0.034737689 |
| TRIM4 | 997.8562646 | 0.441097353 | 0.124867846 | 1.66E-05 | 0.000596012 |
| SNHG32 | 1669.7112 | 0.439682912 | 0.152894964 | 0.000170741 | 0.003833696 |
| EBI3 | 3.842228794 | 0.439584478 | 0.795050289 | 0.004422492 | 0.046595371 |
| CALR | 30137.5899 | 0.43902698 | 0.117624637 | 7.64E-06 | 0.000314144 |
| BCOR | 5260.938348 | 0.438820027 | 0.206140333 | 0.001473655 | 0.020518186 |
| ZNF205 | 395.5306165 | 0.43764569 | 0.226696921 | 0.002380837 | 0.029527185 |
| RNF113A | 556.0549395 | 0.43679591 | 0.155276692 | 0.000211734 | 0.004530811 |
| IGSF1 | 1153.796786 | 0.436338587 | 0.211580173 | 0.001762105 | 0.023500364 |
| TRIM35 | 1356.520759 | 0.435500176 | 0.172414835 | 0.000511678 | 0.00902567 |
| FAM151B | 170.9267656 | 0.435201745 | 0.194947154 | 0.001154931 | 0.017119047 |
| LINC03009 | 371.5640946 | 0.434980184 | 0.160228563 | 0.000290871 | 0.005807688 |
| NFIL3 | 811.6695271 | 0.434360995 | 0.134558068 | 5.27E-05 | 0.001523113 |
| FAM43A | 333.1718344 | 0.433854696 | 0.247087089 | 0.003553481 | 0.039840365 |
| HNF4G | 316.7787995 | 0.433607135 | 0.187715268 | 0.000948513 | 0.014805453 |
| TDRKH | 1930.726101 | 0.433149986 | 0.134942704 | 5.65E-05 | 0.001602413 |
| AFTPH | 2153.991343 | 0.433100142 | 0.139554408 | 8.21E-05 | 0.002146817 |
| NEK10 | 111.4511118 | 0.432769607 | 0.255263689 | 0.004037134 | 0.043648921 |
| HSP90B1 | 46666.02203 | 0.432089238 | 0.165384359 | 0.00040311 | 0.007530372 |
| ZNF70 | 747.5127408 | 0.431973087 | 0.164821205 | 0.000393507 | 0.007364159 |
| DNAJB5 | 1154.394844 | 0.431808895 | 0.144220591 | 0.00011993 | 0.002888968 |
| BEND7 | 1325.679067 | 0.431761139 | 0.134583817 | 5.72E-05 | 0.001618158 |
| GZF1 | 1434.780082 | 0.431578869 | 0.130276716 | 3.93E-05 | 0.001199921 |
| SNHG17 | 1351.126977 | 0.431035591 | 0.156409331 | 0.000261097 | 0.005367139 |
| SYNGR2 | 1742.368602 | 0.429721514 | 0.18915463 | 0.001074729 | 0.016241474 |
| TYW5 | 1305.360703 | 0.429150665 | 0.174939483 | 0.000653458 | 0.011011848 |
| INCA1 | 145.9368841 | 0.429128661 | 0.251439215 | 0.004048269 | 0.043739321 |
| THUMPD2 | 1160.537246 | 0.428582213 | 0.20370952 | 0.001668618 | 0.022526886 |
| CDK17 | 2280.112052 | 0.427709764 | 0.152635159 | 0.000229325 | 0.004835525 |

| | | | | | |
|---|---|---|---|---|---|
| UAP1L1 | 1880.247309 | 0.427626792 | 0.223686545 | 0.002649612 | 0.032023814 |
| MPIG6B | 61.19516033 | 0.427477881 | 0.258072683 | 0.004511338 | 0.047227716 |
| ESAM | 321.6483421 | 0.426270412 | 0.18291658 | 0.000936003 | 0.014664986 |
| RIMOC1 | 4523.55493 | 0.423561183 | 0.166662156 | 0.000521755 | 0.009149334 |
| OLMALINC | 1079.743233 | 0.423106962 | 0.206993972 | 0.001999987 | 0.025927832 |
| SPACA4 | 2.83529612 | 0.422769519 | 0.805049332 | 0.004057455 | 0.043770639 |
| SLMAP | 2996.226097 | 0.422121136 | 0.15396526 | 0.000285855 | 0.005740461 |
| RCHY1 | 1433.679669 | 0.422115432 | 0.188826896 | 0.001238236 | 0.018084309 |
| TENT4B | 1467.959713 | 0.420709555 | 0.179294235 | 0.000925112 | 0.01455988 |
| ATP6V1C2 | 192.656447 | 0.420687804 | 0.218747717 | 0.002700839 | 0.03248347 |
| DDB2 | 1561.900823 | 0.420415574 | 0.128067721 | 4.61E-05 | 0.001360515 |
| DNAJA3 | 2203.863891 | 0.419884817 | 0.136754004 | 9.80E-05 | 0.002458979 |
| HES6 | 724.740887 | 0.419549563 | 0.236083901 | 0.00374594 | 0.041306206 |
| ZSWIM6 | 2542.752468 | 0.418385318 | 0.132373285 | 7.20E-05 | 0.001951053 |
| TOGARAM1 | 940.0807348 | 0.418258184 | 0.184951837 | 0.001182686 | 0.017436063 |
| BLTP3B | 1777.943901 | 0.417867969 | 0.187121879 | 0.001278225 | 0.018503797 |
| IFI27 | 94.45169257 | 0.417552464 | 0.25206476 | 0.004833426 | 0.049459142 |
| NSDHL | 1504.516822 | 0.417111171 | 0.130634064 | 6.46E-05 | 0.001787668 |
| PRSS57 | 4.186795384 | 0.417043743 | 0.784767286 | 0.00462905 | 0.048146266 |
| TRIM8 | 3722.714946 | 0.414564131 | 0.199778728 | 0.001956021 | 0.02552872 |
| EMC9 | 510.7589452 | 0.413844127 | 0.22115985 | 0.003176671 | 0.036533462 |
| ZCCHC7 | 1633.716936 | 0.413803354 | 0.158960477 | 0.000458225 | 0.008292917 |
| SELENOS | 2982.053229 | 0.413758631 | 0.128923919 | 6.20E-05 | 0.001720971 |
| FBXL17 | 2173.853628 | 0.413302097 | 0.13163372 | 7.95E-05 | 0.002109182 |
| MAFF | 359.3258439 | 0.411014544 | 0.210676455 | 0.002698576 | 0.03248347 |
| ZNF451 | 3305.732087 | 0.410875738 | 0.148968086 | 0.000288338 | 0.005773676 |
| RAB11FIP5 | 991.3438537 | 0.410697845 | 0.217963038 | 0.00315091 | 0.036341478 |
| PCNX2 | 2158.743211 | 0.407907875 | 0.183678162 | 0.001403943 | 0.019825112 |
| RASGRF2 | 293.7999963 | 0.406710615 | 0.220799589 | 0.003555779 | 0.039840365 |
| RIMS4 | 2264.771299 | 0.405644383 | 0.207181881 | 0.002750325 | 0.032841929 |
| N4BP2 | 2709.684498 | 0.404874144 | 0.145509579 | 0.000275955 | 0.005611808 |
| KMT5A | 3016.949021 | 0.404637821 | 0.168588584 | 0.000875966 | 0.013891071 |
| CHKA | 1740.140972 | 0.402723334 | 0.144921327 | 0.000282533 | 0.005701162 |
| DNAH10 | 422.157818 | 0.402240923 | 0.215960351 | 0.003501723 | 0.039446324 |
| YTHDF1 | 3368.267078 | 0.402195302 | 0.143470374 | 0.000262032 | 0.005381052 |
| FAM163B | 891.1450454 | 0.402189119 | 0.231958827 | 0.004617507 | 0.048050145 |
| SCMH1 | 1831.564229 | 0.401866435 | 0.148103575 | 0.00034936 | 0.00673721 |
| DDX49 | 3022.705954 | 0.401794911 | 0.170928701 | 0.001024333 | 0.015713723 |
| GFUS | 2146.157438 | 0.401617267 | 0.152362981 | 0.000444874 | 0.008121674 |
| ZNF888 | 451.6145755 | 0.401591259 | 0.219807865 | 0.003805083 | 0.041803872 |
| CACNB1 | 820.1411004 | 0.401378829 | 0.172509818 | 0.001097587 | 0.016515199 |
| LRRC59 | 10153.07685 | 0.400949507 | 0.118034591 | 3.32E-05 | 0.001056065 |
| PARP8 | 1526.326746 | 0.400634499 | 0.222646063 | 0.004066521 | 0.043821982 |
| NT5DC2 | 13116.38261 | 0.400394003 | 0.143688676 | 0.000279431 | 0.005649484 |
| SLC22A17 | 3139.755069 | 0.400238529 | 0.161680002 | 0.00072448 | 0.011980024 |
| CRCP | 2712.422107 | 0.399924189 | 0.161710984 | 0.000730991 | 0.01204952 |
| SPATA2 | 649.2557196 | 0.399747614 | 0.16051801 | 0.000695782 | 0.011569585 |
| KCNA3 | 147.1072129 | 0.399392347 | 0.203992018 | 0.002867155 | 0.033907436 |
| MTO1 | 806.8916943 | 0.398134194 | 0.206445568 | 0.003097423 | 0.035918757 |
| VPS11 | 2613.076693 | 0.397692185 | 0.125872872 | 8.05E-05 | 0.002120714 |

| | | | | | |
|---|---|---|---|---|---|
| JADE1 | 1775.400082 | 0.397675075 | 0.20087112 | 0.002753381 | 0.032859635 |
| MPV17L2 | 610.3921259 | 0.395884355 | 0.167915444 | 0.001042097 | 0.015897843 |
| KICS2 | 486.7018811 | 0.393449551 | 0.175790892 | 0.0014685 | 0.020498653 |
| SLC39A14 | 7543.138167 | 0.392513173 | 0.171919783 | 0.001308185 | 0.018824739 |
| LINC02958 | 2.019507259 | 0.392126883 | 0.793282776 | 0.00406851 | 0.043821982 |
| WDR43 | 5034.216004 | 0.392093959 | 0.153715923 | 0.000609777 | 0.010413084 |
| PPP1R3F | 367.469844 | 0.391757373 | 0.161181672 | 0.000869551 | 0.013810316 |
| CDK5R2 | 423.4152009 | 0.390435379 | 0.205285419 | 0.003471116 | 0.039164911 |
| ATXN2L | 6713.08908 | 0.389832357 | 0.182394982 | 0.001960348 | 0.025556725 |
| RNF138 | 1667.055101 | 0.389743347 | 0.209671208 | 0.003847105 | 0.042176787 |
| ZNF696 | 1119.923512 | 0.388444928 | 0.171290627 | 0.001400333 | 0.019800922 |
| SMURF1 | 3153.993383 | 0.387928288 | 0.161166475 | 0.000953092 | 0.014854722 |
| ZBTB48 | 1094.239439 | 0.387704636 | 0.141467556 | 0.000349053 | 0.00673721 |
| CNOT3 | 2162.932534 | 0.387680551 | 0.163170027 | 0.001042711 | 0.015897843 |
| PARD3B | 522.2327661 | 0.387281662 | 0.180593972 | 0.001956978 | 0.02552872 |
| YDJC | 789.0102966 | 0.387278085 | 0.184808565 | 0.002218383 | 0.028079353 |
| ZNF777 | 1136.445624 | 0.386715675 | 0.182471427 | 0.002095884 | 0.026887304 |
| UGDH | 3784.840758 | 0.386544862 | 0.137946747 | 0.000288722 | 0.005775833 |
| IFRD1 | 1472.441173 | 0.386516394 | 0.187392895 | 0.002422663 | 0.029841537 |
| UBN2 | 4348.961474 | 0.386222348 | 0.143474886 | 0.000410665 | 0.007630544 |
| BRICD5 | 229.8423772 | 0.385965989 | 0.178489794 | 0.001884342 | 0.02476682 |
| PITPNM1 | 2568.688917 | 0.38595852 | 0.210049495 | 0.004143253 | 0.044406326 |
| GAN | 999.0472377 | 0.385758163 | 0.145385726 | 0.000464354 | 0.008367585 |
| IFI6 | 562.4909926 | 0.385320302 | 0.211331614 | 0.004294634 | 0.045622289 |
| SND1 | 14376.89951 | 0.385008831 | 0.140425798 | 0.000353705 | 0.006802145 |
| WDR25 | 501.0040812 | 0.383937704 | 0.15423186 | 0.000770268 | 0.012538616 |
| UBE2J1 | 4899.809484 | 0.382834326 | 0.153148917 | 0.000752394 | 0.012334186 |
| RAB21 | 3198.181456 | 0.382643026 | 0.138545222 | 0.000336739 | 0.006536054 |
| FAM53C | 1819.963993 | 0.381533246 | 0.183441857 | 0.002399275 | 0.029650217 |
| ADCY5 | 173.9581123 | 0.381409612 | 0.214771297 | 0.00489559 | 0.049899655 |
| PFKFB4 | 612.0347276 | 0.381138255 | 0.177800306 | 0.002043522 | 0.026361314 |
| METTL22 | 1038.036564 | 0.381006795 | 0.168335634 | 0.001488107 | 0.020664299 |
| TMED2 | 10369.62205 | 0.38010582 | 0.152252468 | 0.000773146 | 0.01256588 |
| PARS2 | 323.0745478 | 0.379881538 | 0.191808075 | 0.003100758 | 0.035937474 |
| TULP4 | 4392.734531 | 0.379729105 | 0.130280603 | 0.000208313 | 0.004475949 |
| EPRS1 | 10363.19611 | 0.379696359 | 0.160379697 | 0.001124984 | 0.016818383 |
| KDM4B | 2898.428499 | 0.379615809 | 0.176201475 | 0.002007414 | 0.026007968 |
| ACACA | 6874.671814 | 0.379339078 | 0.186188446 | 0.00270835 | 0.03254539 |
| NCKAP5L | 2463.697338 | 0.379082647 | 0.192100426 | 0.003172264 | 0.036523004 |
| ANKRD18B | 237.5296376 | 0.378467941 | 0.194481653 | 0.003400664 | 0.038557479 |
| FBRS | 3465.642305 | 0.377495773 | 0.204927606 | 0.004352585 | 0.046068857 |
| EAF1 | 2472.592894 | 0.377357879 | 0.152822702 | 0.000852945 | 0.013599371 |
| GABRA3 | 729.9756364 | 0.376923404 | 0.142829068 | 0.000513876 | 0.009056798 |
| C21orf91 | 1296.411024 | 0.376901623 | 0.186365621 | 0.002858301 | 0.033857205 |
| UVRAG | 1297.432805 | 0.376231412 | 0.152328937 | 0.000857836 | 0.013655412 |
| ZNF416 | 390.1329659 | 0.376201769 | 0.199165558 | 0.003949167 | 0.043002265 |
| GOLGA4 | 5087.492613 | 0.376154673 | 0.180233344 | 0.002444232 | 0.02998531 |
| GPR157 | 369.4261404 | 0.37604267 | 0.189195712 | 0.003130532 | 0.036182158 |
| ITPR1 | 5745.72271 | 0.375816933 | 0.191489286 | 0.003330012 | 0.037900436 |
| UPP1 | 912.4934754 | 0.375812606 | 0.193037294 | 0.003457696 | 0.03911917 |

| | | | | | |
|---|---|---|---|---|---|
| MSANTD7 | 1606.292459 | 0.375796921 | 0.174573011 | 0.00206978 | 0.026617823 |
| COLQ | 302.5906196 | 0.375463591 | 0.170573333 | 0.001825757 | 0.024194753 |
| ZNF398 | 1576.206591 | 0.375315045 | 0.145668591 | 0.000628549 | 0.010683602 |
| ADPRM | 350.3742264 | 0.375149694 | 0.201441611 | 0.004228073 | 0.045191049 |
| PDIA3 | 19565.89349 | 0.375050018 | 0.147042621 | 0.000680704 | 0.011373205 |
| IER2 | 2108.041827 | 0.374979033 | 0.144708394 | 0.000602372 | 0.010319629 |
| CALN1 | 2.01483215 | 0.374808183 | 0.77398903 | 0.004500064 | 0.047156968 |
| SCARB1 | 1330.901215 | 0.374652407 | 0.18434553 | 0.002832656 | 0.033613653 |
| UAP1 | 3770.547101 | 0.374103584 | 0.155196127 | 0.00103327 | 0.015797843 |
| BCL2L13 | 2277.479701 | 0.374036916 | 0.125135423 | 0.000168305 | 0.003785999 |
| HELB | 160.6658325 | 0.372927757 | 0.199920809 | 0.004265721 | 0.045421337 |
| SMAD6 | 2001.355111 | 0.372600784 | 0.167074303 | 0.001723121 | 0.023069542 |
| TNIP1 | 2534.559 | 0.37219749 | 0.145186711 | 0.000667384 | 0.01121344 |
| GTPBP1 | 1217.027162 | 0.371897937 | 0.174803948 | 0.002269439 | 0.028586994 |
| FUBP3 | 6393.196767 | 0.371733533 | 0.107129396 | 3.00E-05 | 0.000970434 |
| ZNF256 | 324.2600481 | 0.371238965 | 0.169047994 | 0.001905231 | 0.024978498 |
| MAP3K14 | 383.126293 | 0.371142961 | 0.191764905 | 0.003671539 | 0.040765595 |
| EFNB1 | 567.2034144 | 0.370961694 | 0.18292232 | 0.002938141 | 0.034531832 |
| FRS3 | 372.7551709 | 0.370376992 | 0.188489691 | 0.003440196 | 0.038963396 |
| HSF1 | 3004.030582 | 0.370282245 | 0.156010235 | 0.001178679 | 0.017396968 |
| TMEM50B | 2142.984372 | 0.369213085 | 0.17483995 | 0.002408235 | 0.029690671 |
| JUND | 3994.928618 | 0.368571398 | 0.172867442 | 0.00229523 | 0.028790313 |
| PNMA2 | 2015.394397 | 0.368056651 | 0.172574309 | 0.002299712 | 0.028813612 |
| AP5Z1 | 3517.303385 | 0.3678228 | 0.18844121 | 0.003614397 | 0.04035614 |
| KLHL17 | 1619.869639 | 0.36737927 | 0.159715902 | 0.001475965 | 0.020536665 |
| MED13 | 9812.336032 | 0.367345532 | 0.157955064 | 0.001375779 | 0.019533074 |
| DOHH | 694.6396159 | 0.367294712 | 0.172686679 | 0.002346998 | 0.029159461 |
| IQCH | 158.8959431 | 0.366593739 | 0.186424616 | 0.003521422 | 0.039625484 |
| USP32 | 6240.371057 | 0.366465851 | 0.167305866 | 0.001999439 | 0.025927832 |
| PPP1R16B | 2441.068543 | 0.366195148 | 0.184269397 | 0.003357843 | 0.038175587 |
| RNF6 | 2121.928363 | 0.366143532 | 0.194930146 | 0.004347476 | 0.046043126 |
| PLCXD2 | 511.0821797 | 0.36605272 | 0.129305103 | 0.000300991 | 0.00595284 |
| FAM135A | 957.3356462 | 0.365065973 | 0.16391611 | 0.001830486 | 0.024242046 |
| PHF10 | 6630.96976 | 0.364943554 | 0.136027428 | 0.000486552 | 0.008640847 |
| SMCR8 | 2709.8603 | 0.362909938 | 0.150796887 | 0.001128983 | 0.016842013 |
| WDR45B | 7220.029253 | 0.362006989 | 0.109673733 | 6.06E-05 | 0.001697892 |
| ZNF12 | 4167.03721 | 0.361982245 | 0.145549417 | 0.000899882 | 0.014205617 |
| TICAM1 | 776.9161053 | 0.361758473 | 0.184588567 | 0.003704491 | 0.041044193 |
| DR1 | 5442.226458 | 0.360848102 | 0.116114991 | 0.000122462 | 0.002936412 |
| ZBTB21 | 1510.917135 | 0.360812228 | 0.175314245 | 0.002929256 | 0.034466203 |
| TPRN | 684.9181065 | 0.360471729 | 0.19082638 | 0.004419784 | 0.046595371 |
| SLC25A38 | 1681.434687 | 0.360166403 | 0.116114111 | 0.000125624 | 0.00299163 |
| TYRO3 | 1365.733329 | 0.360100326 | 0.192644555 | 0.004638509 | 0.048220632 |
| ZSWIM9 | 510.6850628 | 0.35962095 | 0.175250132 | 0.002999694 | 0.034999224 |
| TVP23B | 2223.692516 | 0.359168272 | 0.167869419 | 0.002406649 | 0.029690671 |
| ARL4D | 220.5458166 | 0.358188846 | 0.157936741 | 0.001721378 | 0.023069542 |
| GRINA | 3991.74748 | 0.357614517 | 0.176482374 | 0.003245045 | 0.037109785 |
| ZMYM5 | 1062.335476 | 0.357584926 | 0.159524977 | 0.001857311 | 0.024522127 |
| TCEA1 | 5657.162842 | 0.357497026 | 0.174575223 | 0.003078698 | 0.03576118 |
| ZEB1 | 3441.499648 | 0.357280255 | 0.15425222 | 0.00151699 | 0.020940081 |

| | | | | | |
|---|---|---|---|---|---|
| LARP4B | 4871.30898 | 0.356825999 | 0.114734202 | 0.000125139 | 0.002983471 |
| ZNF79 | 490.2369459 | 0.356653287 | 0.156508 | 0.001689524 | 0.022735641 |
| BRAF | 2793.093089 | 0.356479758 | 0.130519725 | 0.000444158 | 0.008121674 |
| KDSR | 3261.515449 | 0.356080686 | 0.165965672 | 0.00242333 | 0.029841537 |
| EDEM2 | 990.9217629 | 0.356074013 | 0.125515811 | 0.000317241 | 0.006227043 |
| CDC37L1 | 922.1494701 | 0.355204857 | 0.160572536 | 0.002045462 | 0.026370041 |
| UBOX5 | 687.3868553 | 0.355153512 | 0.184406766 | 0.004214613 | 0.04507025 |
| MTMR14 | 2006.798812 | 0.35467819 | 0.12442965 | 0.00030669 | 0.006059812 |
| GTF2IRD1 | 2661.591155 | 0.354580687 | 0.151371438 | 0.001437886 | 0.02018598 |
| RNF10 | 5053.371009 | 0.353491055 | 0.130258118 | 0.000479879 | 0.00857337 |
| CDC34 | 1974.336836 | 0.353170206 | 0.172118339 | 0.003145305 | 0.036332795 |
| ILRUN | 3826.675449 | 0.352806583 | 0.118234789 | 0.000199343 | 0.004350336 |
| EIF4EBP1 | 2436.231538 | 0.352806481 | 0.161446134 | 0.002235798 | 0.028253958 |
| PPTC7 | 2615.423641 | 0.352690988 | 0.181734896 | 0.004144023 | 0.044406326 |
| ETV3 | 3252.331934 | 0.352671671 | 0.138950716 | 0.000827631 | 0.013285991 |
| ZNF282 | 2271.893604 | 0.35135957 | 0.150346157 | 0.001495126 | 0.020716969 |
| DYRK4 | 535.1711289 | 0.350740883 | 0.144083441 | 0.001139102 | 0.016956629 |
| TMEM231 | 596.8720415 | 0.349806261 | 0.151323728 | 0.001623202 | 0.022013361 |
| NTMT1 | 637.3023685 | 0.347871996 | 0.140239468 | 0.001017394 | 0.01563765 |
| MADD | 4508.487099 | 0.347518505 | 0.119814476 | 0.000274816 | 0.005594082 |
| AVL9 | 5641.270884 | 0.346527774 | 0.120468409 | 0.000299793 | 0.005938884 |
| SLC33A1 | 1075.784269 | 0.346495177 | 0.178927806 | 0.004383819 | 0.046310604 |
| URB2 | 2377.093971 | 0.346271794 | 0.123018212 | 0.000367418 | 0.006969604 |
| ZNF791 | 2163.330115 | 0.345526571 | 0.157773063 | 0.002331229 | 0.029017122 |
| YIPF5 | 5459.478814 | 0.345285454 | 0.18066337 | 0.004697191 | 0.048612898 |
| ZRANB1 | 1260.612015 | 0.345251681 | 0.134950159 | 0.000824116 | 0.013239742 |
| CRELD1 | 263.1343424 | 0.344515935 | 0.170457968 | 0.003622849 | 0.040418374 |
| VPS37A | 1693.885854 | 0.344255561 | 0.132463547 | 0.000734908 | 0.01210453 |
| AFF4 | 11650.81226 | 0.343737724 | 0.148365905 | 0.001675262 | 0.022579815 |
| RUNDC3A | 1158.014653 | 0.343697146 | 0.163590811 | 0.002981357 | 0.034843669 |
| LPCAT3 | 431.7715886 | 0.343564917 | 0.17842482 | 0.00460041 | 0.047967826 |
| NR1D2 | 4413.134362 | 0.343033421 | 0.13519993 | 0.000893192 | 0.01414277 |
| BRPF1 | 1682.479534 | 0.341873732 | 0.153759311 | 0.002192694 | 0.027822747 |
| EIF1 | 18982.27482 | 0.340793544 | 0.1332715 | 0.000856577 | 0.013645778 |
| KRCC1 | 2141.424226 | 0.340714109 | 0.141012115 | 0.001295715 | 0.018675649 |
| CFAP97 | 4221.631674 | 0.340608302 | 0.153590829 | 0.002248857 | 0.028344827 |
| NUP58 | 5401.300337 | 0.340556377 | 0.117264593 | 0.00028713 | 0.005756547 |
| ZNF264 | 1650.340062 | 0.340524409 | 0.146096097 | 0.001651808 | 0.022343265 |
| TBC1D22B | 690.4268686 | 0.340118842 | 0.123646939 | 0.000473931 | 0.008510783 |
| IRS2 | 9373.307953 | 0.339825194 | 0.146494568 | 0.001712953 | 0.02299166 |
| ZBTB10 | 2382.439352 | 0.339357831 | 0.134514501 | 0.000959688 | 0.014924117 |
| SEC61A1 | 16761.68884 | 0.338440855 | 0.138285292 | 0.001205744 | 0.017685467 |
| TRMT61A | 934.4715448 | 0.337903305 | 0.176173702 | 0.00489066 | 0.049873751 |
| HAX1 | 2839.678856 | 0.337412862 | 0.121778263 | 0.0004547 | 0.00824342 |
| ENC1 | 8316.355384 | 0.336440876 | 0.165325947 | 0.003723476 | 0.041149427 |
| RAD52 | 871.9126462 | 0.336303058 | 0.139193789 | 0.001342221 | 0.019160831 |
| KAT7 | 3328.512627 | 0.335891614 | 0.121924627 | 0.000484209 | 0.008606557 |
| PRKCI | 3870.307094 | 0.335843709 | 0.14928893 | 0.002141525 | 0.027354489 |
| ZNF419 | 840.9538025 | 0.335797365 | 0.153052739 | 0.00248646 | 0.030386027 |
| STARD4 | 11624.79826 | 0.334999799 | 0.162777643 | 0.003554905 | 0.039840365 |

| | | | | | |
|---|---|---|---|---|---|
| KDM6A | 2429.70467 | 0.334370832 | 0.147167442 | 0.002038668 | 0.02631494 |
| RCOR1 | 3642.441236 | 0.334172978 | 0.140277137 | 0.001503434 | 0.02080797 |
| SIRT1 | 2898.009786 | 0.334158982 | 0.159231071 | 0.003228853 | 0.036970718 |
| TMEM170B | 1793.121396 | 0.334075713 | 0.138557491 | 0.001386056 | 0.019652254 |
| FBXW4 | 1201.693858 | 0.333847372 | 0.163977359 | 0.003790893 | 0.041691841 |
| BET1 | 1839.689398 | 0.333182944 | 0.150906921 | 0.002446491 | 0.02998531 |
| FDXR | 1066.335636 | 0.333030759 | 0.15170998 | 0.002533442 | 0.030797987 |
| GORASP2 | 4925.734852 | 0.332376402 | 0.122420289 | 0.000565486 | 0.009833611 |
| RIMKLA | 575.7429826 | 0.332348203 | 0.135904216 | 0.001273544 | 0.018462027 |
| FXR2 | 3616.394072 | 0.332184355 | 0.140420834 | 0.001601023 | 0.021754958 |
| SERTAD2 | 1541.052345 | 0.331402093 | 0.135159055 | 0.001259633 | 0.01833271 |
| EIF1B | 1435.419556 | 0.331278324 | 0.152359996 | 0.002714525 | 0.032600837 |
| SHB | 648.6321521 | 0.33085261 | 0.159762383 | 0.003555586 | 0.039840365 |
| CLPTM1L | 6730.442158 | 0.3308409 | 0.114041205 | 0.000314994 | 0.006194581 |
| NHERF1 | 1448.494555 | 0.330781195 | 0.155638819 | 0.003096284 | 0.035918757 |
| PATL1 | 2518.445813 | 0.330396829 | 0.134874159 | 0.001278476 | 0.018503797 |
| NFKBIA | 2124.437644 | 0.329695724 | 0.166186644 | 0.004452482 | 0.046846395 |
| TARS1 | 9514.050079 | 0.329489814 | 0.158437487 | 0.003515973 | 0.039585501 |
| BRSK1 | 1996.404046 | 0.329031467 | 0.146716784 | 0.002304526 | 0.028837642 |
| GTF2A1 | 4989.037031 | 0.328399475 | 0.148384762 | 0.002506645 | 0.030561162 |
| PDHX | 1453.819857 | 0.327917916 | 0.147377023 | 0.002437983 | 0.02998531 |
| TADA2B | 1338.648523 | 0.327819329 | 0.120039668 | 0.000559386 | 0.009760048 |
| PDIA6 | 14424.45356 | 0.327592473 | 0.124839833 | 0.000780014 | 0.012657798 |
| ACVR1 | 3389.726564 | 0.326180101 | 0.135452131 | 0.001491778 | 0.020701522 |
| ATF1 | 1221.42239 | 0.3258705 | 0.147623657 | 0.002598047 | 0.03149171 |
| SURF4 | 10626.7914 | 0.325846065 | 0.144413704 | 0.002279371 | 0.028644983 |
| RAD9A | 923.3025366 | 0.325446271 | 0.161900556 | 0.004319388 | 0.045815346 |
| NIPA1 | 3050.193345 | 0.324209896 | 0.14056364 | 0.002015107 | 0.026064854 |
| SC5D | 2850.53898 | 0.323793031 | 0.156071789 | 0.00372774 | 0.041170728 |
| NEK4 | 1066.764024 | 0.323686893 | 0.14397779 | 0.002372094 | 0.029453731 |
| SNX11 | 1676.861752 | 0.323235084 | 0.106943845 | 0.000224598 | 0.004757322 |
| FARSA | 4013.245275 | 0.32144123 | 0.14209575 | 0.002327478 | 0.029017122 |
| NEU1 | 1900.758135 | 0.3195224 | 0.15189162 | 0.003587125 | 0.040105518 |
| ESCO1 | 1713.486464 | 0.319213281 | 0.156749392 | 0.004258416 | 0.04538027 |
| BMP8B | 298.5850091 | 0.318645461 | 0.158494249 | 0.004562015 | 0.047662675 |
| KLHDC4 | 2245.943737 | 0.318059547 | 0.151830642 | 0.003713122 | 0.041118003 |
| PRKRIP1 | 2569.857737 | 0.31803256 | 0.124238134 | 0.001028788 | 0.015752371 |
| MED10 | 2267.903655 | 0.317382328 | 0.130119747 | 0.001471917 | 0.020507667 |
| BMT2 | 1793.491286 | 0.31729065 | 0.150958786 | 0.003671041 | 0.040765595 |
| ZBTB5 | 2046.182108 | 0.31702371 | 0.103765098 | 0.000213036 | 0.004544687 |
| WWC1 | 863.4119515 | 0.315965888 | 0.152905776 | 0.004059657 | 0.043771774 |
| ELAVL3 | 2016.738567 | 0.314823448 | 0.152534795 | 0.004124112 | 0.044261091 |
| TTBK2 | 2130.334151 | 0.313685749 | 0.140430225 | 0.002683352 | 0.032349879 |
| DPF2 | 2570.091183 | 0.310988236 | 0.131011232 | 0.001874182 | 0.024676318 |
| PRKACA | 2682.105589 | 0.31076827 | 0.147810143 | 0.003872948 | 0.042393357 |
| TP53 | 6022.996405 | 0.310599426 | 0.148574246 | 0.003997761 | 0.043373114 |
| INO80D | 1455.246798 | 0.30987215 | 0.124834832 | 0.001390453 | 0.019701229 |
| MTDH | 7739.02984 | 0.309520731 | 0.128963833 | 0.001763864 | 0.023508808 |
| TMEM222 | 1565.257213 | 0.308766089 | 0.147320651 | 0.004006684 | 0.043447349 |
| REV3L | 2401.733151 | 0.307222339 | 0.121760526 | 0.001264932 | 0.018387821 |

| | | | | | |
|---|---|---|---|---|---|
| ARIH2 | 5952.415962 | 0.307199117 | 0.103500296 | 0.000314812 | 0.006194581 |
| DCP1B | 931.8506277 | 0.307174109 | 0.150649014 | 0.00468307 | 0.048514836 |
| CYFIP2 | 2781.398629 | 0.306942484 | 0.137519154 | 0.002859665 | 0.033857205 |
| PIK3C2A | 6864.133495 | 0.306775832 | 0.148683664 | 0.004424162 | 0.046595371 |
| FAF1 | 2727.503004 | 0.306406505 | 0.135336787 | 0.002637589 | 0.031896953 |
| ZCCHC2 | 1055.495669 | 0.306161101 | 0.143328771 | 0.003699356 | 0.041009053 |
| ADIPOR2 | 3754.398588 | 0.30510609 | 0.116076036 | 0.000943613 | 0.014744472 |
| RRP9 | 1206.086003 | 0.304996501 | 0.149595319 | 0.004774272 | 0.049142769 |
| ZXDB | 659.2576003 | 0.304505649 | 0.13249542 | 0.002446226 | 0.02998531 |
| TNPO1 | 16004.50855 | 0.304350425 | 0.133404272 | 0.002563833 | 0.031113108 |
| KMT2A | 6168.74197 | 0.304049249 | 0.144611282 | 0.004104885 | 0.044145485 |
| WASL | 7055.498075 | 0.30389384 | 0.138924942 | 0.00330399 | 0.037665842 |
| TERF2IP | 3789.251346 | 0.302958382 | 0.141815682 | 0.003803228 | 0.041803872 |
| POLR1H | 363.2758402 | 0.3022664 | 0.131348532 | 0.002476187 | 0.030278197 |
| INAFM2 | 945.9295158 | 0.300910653 | 0.124869271 | 0.001857506 | 0.024522127 |
| OSBPL2 | 2239.374506 | 0.29982107 | 0.124065765 | 0.001840925 | 0.024349416 |
| FDPS | 3066.70767 | 0.299504009 | 0.124609644 | 0.001915643 | 0.025083517 |
| UBIAD1 | 1075.285591 | 0.299300587 | 0.127946427 | 0.00229403 | 0.028790313 |
| INPP5B | 1165.034583 | 0.29859102 | 0.117576751 | 0.001308763 | 0.018824739 |
| FOXJ2 | 1374.559519 | 0.298400016 | 0.130527245 | 0.002673155 | 0.032271006 |
| RLF | 2058.033922 | 0.296921992 | 0.13578784 | 0.003526068 | 0.039635048 |
| KCMF1 | 3861.688447 | 0.296719244 | 0.118685128 | 0.001494818 | 0.020716969 |
| SWI5 | 487.7511032 | 0.295275617 | 0.142644795 | 0.004815712 | 0.049374643 |
| HAUS2 | 1230.775671 | 0.29462058 | 0.142181256 | 0.004818556 | 0.049379559 |
| KMT2C | 6658.509359 | 0.290763691 | 0.132494505 | 0.003657502 | 0.040652972 |
| CSNK1A1 | 5882.478693 | 0.28997636 | 0.117956092 | 0.001799388 | 0.02390599 |
| PANK2 | 1752.37589 | 0.289936174 | 0.107922733 | 0.00092035 | 0.014506798 |
| KLF3 | 3367.06649 | 0.289903147 | 0.119407476 | 0.001962575 | 0.025569799 |
| PTAR1 | 5262.619289 | 0.28771163 | 0.137433458 | 0.004868295 | 0.049754295 |
| TOP3A | 3847.396792 | 0.28708074 | 0.133193924 | 0.004188445 | 0.044813353 |
| PIKFYVE | 5186.933475 | 0.285573239 | 0.132963591 | 0.004331731 | 0.045899644 |
| MARS1 | 7873.402338 | 0.2843851 | 0.098439729 | 0.00051814 | 0.009108891 |
| FOXK2 | 5300.591069 | 0.283950927 | 0.128408926 | 0.003736008 | 0.041218446 |
| TRMT1 | 1633.125418 | 0.283402763 | 0.123344248 | 0.002990477 | 0.034930715 |
| GTF2F1 | 4775.839269 | 0.282938728 | 0.124487323 | 0.003209732 | 0.036832559 |
| FLCN | 2642.966923 | 0.282557192 | 0.132408999 | 0.004616308 | 0.048050145 |
| HIPK1 | 4771.520936 | 0.28149723 | 0.132682569 | 0.004811129 | 0.04936812 |
| SETD2 | 6147.019914 | 0.279567703 | 0.125576716 | 0.003749925 | 0.041328332 |
| YY1AP1 | 2856.713753 | 0.279400564 | 0.128399147 | 0.004276357 | 0.045474399 |
| ZSCAN32 | 887.2924348 | 0.278957224 | 0.126287019 | 0.0039464 | 0.042994555 |
| H3-3B | 38196.27899 | 0.278794689 | 0.126046344 | 0.003922843 | 0.042804897 |
| GOSR2 | 4965.871177 | 0.277356464 | 0.129809988 | 0.004821135 | 0.049381761 |
| EIF1AD | 1286.934684 | 0.274680657 | 0.116305789 | 0.002746073 | 0.032828692 |
| ATP6V0A2 | 3701.424697 | 0.271525391 | 0.115778615 | 0.002966423 | 0.034707978 |
| YY1 | 7384.837344 | 0.271416916 | 0.104339184 | 0.001444447 | 0.020219355 |
| DZIP1 | 5972.390145 | 0.269139924 | 0.114269252 | 0.002958968 | 0.034669486 |
| NSD3 | 6497.751824 | 0.267107473 | 0.120552556 | 0.004366618 | 0.046175601 |
| TSC22D2 | 1868.99106 | 0.2621626 | 0.113537138 | 0.003603036 | 0.04026185 |
| XPC | 1975.569715 | 0.255009392 | 0.110831321 | 0.003972496 | 0.043188749 |
| NAXD | 3855.931098 | 0.25382787 | 0.112775753 | 0.004583369 | 0.04781402 |

| | | | | | |
|---|---|---|---|---|---|
| ARFIP2 | 3902.687828 | 0.25325955 | 0.113331303 | 0.004805211 | 0.049345291 |
| AUP1 | 4672.453621 | 0.249108474 | 0.106011662 | 0.003721312 | 0.041149427 |
| XPR1 | 12154.79798 | -0.218415801 | 0.091906737 | 0.004884998 | 0.049840339 |
| CREBZF | 5396.786249 | -0.235923457 | 0.10225151 | 0.004812712 | 0.04936812 |
| SORT1 | 4656.925042 | -0.242758179 | 0.105269076 | 0.004476881 | 0.04700838 |
| SETD6 | 1574.495834 | -0.2563642 | 0.113923188 | 0.004466223 | 0.046920049 |
| SSR2 | 12103.18419 | -0.256963955 | 0.111819554 | 0.003919584 | 0.042791694 |
| SHOX2 | 2947.586421 | -0.257913051 | 0.112979703 | 0.004037813 | 0.043648921 |
| HPS1 | 612.6904191 | -0.260185022 | 0.116886202 | 0.004567945 | 0.047688327 |
| UBE2L3 | 4092.057225 | -0.260633231 | 0.11629128 | 0.004371704 | 0.046205991 |
| RSAD1 | 2318.828009 | -0.270658465 | 0.122693833 | 0.004318732 | 0.045815346 |
| TMEM18 | 1553.316406 | -0.272308551 | 0.122065558 | 0.003983271 | 0.043238364 |
| ATXN7L3B | 5154.4581 | -0.27305907 | 0.111525545 | 0.002198897 | 0.0278834 |
| BEX4 | 2875.327743 | -0.273625542 | 0.122573268 | 0.003915533 | 0.042778977 |
| TTC5 | 954.369754 | -0.274036486 | 0.117769103 | 0.003037018 | 0.035355698 |
| IP6K2 | 3869.340546 | -0.274188128 | 0.116859301 | 0.002877703 | 0.033993676 |
| NCOA5 | 2201.441485 | -0.2747951 | 0.119688904 | 0.00327394 | 0.037384497 |
| RABIF | 1147.06572 | -0.278008476 | 0.125375808 | 0.003896756 | 0.042610016 |
| NAGA | 620.1740185 | -0.279822096 | 0.126941389 | 0.003959689 | 0.043094376 |
| GLT8D1 | 3049.221959 | -0.281060653 | 0.118112914 | 0.00245078 | 0.030020255 |
| CCDC144NL-AS1 | 2593.023611 | -0.28194948 | 0.112681662 | 0.001722044 | 0.023069542 |
| RAD51D | 655.1775606 | -0.284574006 | 0.114973781 | 0.001811009 | 0.024029814 |
| SFXN2 | 523.7142338 | -0.287676263 | 0.135055739 | 0.004441838 | 0.04675796 |
| SYNJ2BP | 3817.854791 | -0.289340125 | 0.132010376 | 0.003733187 | 0.041209098 |
| PAN2 | 1588.597298 | -0.28960919 | 0.13831128 | 0.004778183 | 0.049158797 |
| GEMIN8 | 1500.036028 | -0.292053478 | 0.104564851 | 0.000647632 | 0.010943207 |
| GSDME | 3594.991115 | -0.292421447 | 0.120700768 | 0.001942716 | 0.025406152 |
| LINC00667 | 774.4062567 | -0.293547109 | 0.116341736 | 0.001442243 | 0.020215599 |
| TCEAL8 | 2702.344565 | -0.294100408 | 0.131814809 | 0.003221069 | 0.036922082 |
| APEX1 | 5936.757842 | -0.294127179 | 0.131617308 | 0.00319023 | 0.036669213 |
| DNALI1 | 1000.627771 | -0.295549821 | 0.127534497 | 0.002522277 | 0.030701371 |
| INSR | 929.4712646 | -0.295972446 | 0.134176307 | 0.003383323 | 0.038423439 |
| MGAT5 | 6821.48136 | -0.296987476 | 0.118357536 | 0.001451853 | 0.020309412 |
| CCNJL | 1143.661828 | -0.297143025 | 0.139768172 | 0.004113424 | 0.044191805 |
| NFS1 | 766.1759761 | -0.29725611 | 0.115291394 | 0.001182997 | 0.017436063 |
| CEP89 | 1721.115523 | -0.297664126 | 0.107702964 | 0.000670137 | 0.011250645 |
| ZSCAN2 | 476.5909408 | -0.297855692 | 0.139010705 | 0.003916371 | 0.042778977 |
| DYNC1I2 | 6272.340261 | -0.298213196 | 0.144747065 | 0.004805761 | 0.049345291 |
| ZBED5 | 3632.508734 | -0.299147981 | 0.12791723 | 0.002301455 | 0.028816478 |
| H2AZ2 | 12024.50817 | -0.300086462 | 0.116511651 | 0.001160055 | 0.017182803 |
| APPL2 | 4002.114636 | -0.300501661 | 0.125798851 | 0.00197799 | 0.02570648 |
| HIBCH | 1035.712524 | -0.303026494 | 0.13290527 | 0.002604551 | 0.031533947 |
| AP1S1 | 4260.15812 | -0.303312642 | 0.139385017 | 0.003421197 | 0.038769238 |
| HACD3 | 6599.681496 | -0.303768496 | 0.124449743 | 0.001656302 | 0.02237508 |
| GDF11 | 1649.837415 | -0.304187357 | 0.123259267 | 0.001527433 | 0.021028639 |
| NEDD8 | 1856.362497 | -0.304688572 | 0.137482364 | 0.003042247 | 0.035396841 |
| ZNF789 | 1206.847706 | -0.305309862 | 0.147850045 | 0.004461567 | 0.046894734 |
| TPM3P9 | 401.2367395 | -0.30544167 | 0.141134843 | 0.003462506 | 0.039131192 |
| CLCN4 | 943.6009925 | -0.305657071 | 0.124378315 | 0.001552247 | 0.021245755 |
| PIGC | 1406.338432 | -0.305854771 | 0.132321501 | 0.00233115 | 0.029017122 |

| | | | | | |
|---|---|---|---|---|---|
| TMEM109 | 2046.505018 | -0.306256961 | 0.131273411 | 0.002189682 | 0.027817224 |
| SCARB2 | 20714.85487 | -0.307299498 | 0.125248972 | 0.001546383 | 0.021219592 |
| EXTL2 | 4662.312796 | -0.307606021 | 0.137118445 | 0.002758146 | 0.03286796 |
| COL11A1 | 1911.810358 | -0.308349333 | 0.140311763 | 0.003093205 | 0.035909714 |
| SMARCA1 | 6517.654065 | -0.309016155 | 0.147070525 | 0.003945087 | 0.042994555 |
| TXNDC15 | 2042.221544 | -0.30906502 | 0.116063395 | 0.000818645 | 0.013172117 |
| TIMM8B | 1420.013809 | -0.309378997 | 0.138151585 | 0.002744064 | 0.032828692 |
| ABHD11 | 533.4219973 | -0.30958643 | 0.151394844 | 0.004520824 | 0.047303306 |
| CNN3 | 14814.35456 | -0.309884885 | 0.144747921 | 0.003537669 | 0.03974406 |
| KNOP1 | 1659.164777 | -0.310683557 | 0.104784254 | 0.000307274 | 0.006065609 |
| EPM2AIP1 | 2412.254263 | -0.310693645 | 0.130403749 | 0.001833401 | 0.02426526 |
| SEM1 | 3275.002027 | -0.311037553 | 0.15157049 | 0.004386038 | 0.046310623 |
| TARDBP | 8377.412616 | -0.311316582 | 0.150966664 | 0.004267218 | 0.045421337 |
| COQ7 | 734.38382 | -0.311332834 | 0.144085243 | 0.00331854 | 0.03779046 |
| CALM1 | 23051.86622 | -0.311393582 | 0.154462945 | 0.004781297 | 0.049166607 |
| IQCG | 662.0204829 | -0.311417647 | 0.13556245 | 0.002306224 | 0.028841611 |
| SLC35B4 | 5663.562294 | -0.311724061 | 0.106054645 | 0.000331243 | 0.006453372 |
| ZNF69 | 410.3554562 | -0.312122234 | 0.136740659 | 0.002385097 | 0.029552575 |
| TCEAL4 | 2118.718789 | -0.312719878 | 0.146830097 | 0.003552129 | 0.039840365 |
| YWHAH | 4709.95208 | -0.31280368 | 0.146365525 | 0.003483276 | 0.039259698 |
| ENOSF1 | 961.7609852 | -0.313321691 | 0.148024182 | 0.003654744 | 0.040652972 |
| SKP1 | 606.8638496 | -0.313509641 | 0.146174094 | 0.003394892 | 0.038533864 |
| PIGN | 1744.083055 | -0.313523931 | 0.149068811 | 0.00377602 | 0.041550146 |
| TRIM2 | 4704.287273 | -0.313955881 | 0.147050402 | 0.003468117 | 0.039152224 |
| CNTRL | 1911.571635 | -0.314872259 | 0.153024667 | 0.004187813 | 0.044813353 |
| C2orf68 | 2213.483623 | -0.315720158 | 0.113724406 | 0.000540656 | 0.009464881 |
| TMBIM4 | 550.9954985 | -0.31644565 | 0.157841454 | 0.004710191 | 0.04869918 |
| CEP152 | 1016.542952 | -0.316917677 | 0.158341906 | 0.00473047 | 0.04883631 |
| ALAD | 1466.529914 | -0.317100022 | 0.147149228 | 0.003207413 | 0.036826184 |
| BBS2 | 1153.893795 | -0.317479511 | 0.150662993 | 0.003615338 | 0.04035614 |
| ZNF302 | 2037.403961 | -0.318355993 | 0.143597527 | 0.002696114 | 0.032473086 |
| CCDC14 | 7593.202022 | -0.318444111 | 0.133225017 | 0.001675782 | 0.022579815 |
| ARMCX6 | 863.3321933 | -0.319255603 | 0.132456303 | 0.001572484 | 0.021478956 |
| RGL1 | 2189.823066 | -0.319714379 | 0.105587648 | 0.000227846 | 0.00481637 |
| PIGU | 865.8284543 | -0.31991661 | 0.136400844 | 0.001876242 | 0.024676318 |
| POMK | 3702.277289 | -0.320316753 | 0.107389061 | 0.000263932 | 0.005409443 |
| MYCBP | 750.0276086 | -0.320582606 | 0.148292943 | 0.003060156 | 0.035585392 |
| MATN3 | 667.6053935 | -0.320729333 | 0.161951129 | 0.004825944 | 0.049406781 |
| UNC50 | 1296.225251 | -0.320939295 | 0.113622106 | 0.000441443 | 0.008081904 |
| VMA21 | 2417.719492 | -0.321913062 | 0.128254782 | 0.00115237 | 0.017093214 |
| TMEM254 | 446.8563571 | -0.322069697 | 0.13430359 | 0.001589007 | 0.021662142 |
| MED20 | 723.7534397 | -0.322177093 | 0.148442554 | 0.002953159 | 0.03466932 |
| TDP1 | 1099.120014 | -0.322575386 | 0.11816504 | 0.000587728 | 0.010161169 |
| TCEAL3 | 560.6290607 | -0.322759243 | 0.142420059 | 0.002276523 | 0.028641681 |
| PUDP | 568.0906812 | -0.323684198 | 0.128227233 | 0.001087952 | 0.016398278 |
| CDC14B | 2064.142905 | -0.32418475 | 0.137454745 | 0.001744874 | 0.023285423 |
| FAHD1 | 1340.869025 | -0.324827213 | 0.119668093 | 0.000604704 | 0.010334894 |
| GVQW3 | 660.5645314 | -0.325682953 | 0.153075595 | 0.003206952 | 0.036826184 |
| HNRNPA3 | 8154.255584 | -0.325948251 | 0.143437918 | 0.002180398 | 0.02774991 |
| GALNT1 | 4995.34344 | -0.326541395 | 0.134520979 | 0.001407304 | 0.019840541 |

| | | | | | |
|---|---|---|---|---|---|
| SPA17 | 283.461778 | -0.326628247 | 0.157394346 | 0.003637807 | 0.040563592 |
| SCPEP1 | 3109.701319 | -0.326796726 | 0.135430147 | 0.001463349 | 0.020442834 |
| TRIM66 | 1473.90195 | -0.327232887 | 0.15646289 | 0.003474299 | 0.039179666 |
| FZD1 | 1345.153733 | -0.327290117 | 0.134940597 | 0.001406554 | 0.019840541 |
| ITGB1BP1 | 2308.88767 | -0.327681065 | 0.150650583 | 0.002788355 | 0.033163322 |
| TOX-DT | 257.1753433 | -0.32774979 | 0.163270394 | 0.004269192 | 0.045421337 |
| PBDC1 | 765.6926753 | -0.327835954 | 0.163789042 | 0.004327754 | 0.045880787 |
| GRAMD1B | 1150.287639 | -0.327893179 | 0.167821173 | 0.004857122 | 0.049677272 |
| BPHL | 1035.199325 | -0.327928176 | 0.113905612 | 0.00034753 | 0.006720523 |
| USP16 | 2285.933359 | -0.328443603 | 0.158863312 | 0.003656101 | 0.040652972 |
| MRPL27 | 1536.515343 | -0.329901826 | 0.136310171 | 0.001397097 | 0.019772849 |
| TMEM44 | 1190.794485 | -0.331149524 | 0.152710607 | 0.002759278 | 0.03286796 |
| RRAGA | 3508.573601 | -0.331591229 | 0.124241135 | 0.000657078 | 0.011058085 |
| TUBG2 | 1398.821542 | -0.332207382 | 0.166034185 | 0.004187995 | 0.044813353 |
| ETFRF1 | 415.0502716 | -0.332445512 | 0.152255564 | 0.002625309 | 0.031766841 |
| MPLKIP | 1162.590967 | -0.332674627 | 0.15571465 | 0.002963574 | 0.034694087 |
| CMTM3 | 2632.984947 | -0.333688026 | 0.147655877 | 0.002118587 | 0.027145213 |
| DAZAP2 | 4804.325515 | -0.335496397 | 0.150167492 | 0.002238942 | 0.028253958 |
| ADGRA3 | 4057.217884 | -0.335657706 | 0.115575628 | 0.000299071 | 0.005932595 |
| FMO5 | 176.3378867 | -0.335685237 | 0.169159885 | 0.004242443 | 0.04529243 |
| DUSP6 | 2249.807821 | -0.336514635 | 0.142348304 | 0.001553376 | 0.021245755 |
| ZNF737 | 794.7400845 | -0.336638079 | 0.158746214 | 0.00299276 | 0.034937833 |
| PGAP4 | 1230.145873 | -0.337738285 | 0.133189216 | 0.000935537 | 0.014664986 |
| NIPBL-DT | 934.2706734 | -0.338272037 | 0.135500492 | 0.001046403 | 0.015940282 |
| HARS2 | 2591.364103 | -0.338923202 | 0.12779066 | 0.00065132 | 0.010987736 |
| TCF4 | 20319.50502 | -0.338923995 | 0.119508047 | 0.000363959 | 0.006935486 |
| PPP1R3E | 331.840985 | -0.339463097 | 0.16227179 | 0.003151277 | 0.036341478 |
| TMEM80 | 538.8996298 | -0.339789643 | 0.165338273 | 0.003450435 | 0.039058175 |
| TMEM25 | 652.8476335 | -0.340826938 | 0.178432313 | 0.0048718 | 0.049754295 |
| HNRNPR | 10707.4784 | -0.340971759 | 0.144511683 | 0.001519182 | 0.020956479 |
| TRDMT1 | 656.0647313 | -0.341842538 | 0.174096126 | 0.004256279 | 0.04538027 |
| KLHL9 | 3059.34187 | -0.342747864 | 0.11367256 | 0.000194116 | 0.004252647 |
| TRAPPC1 | 2289.981438 | -0.342914528 | 0.135332745 | 0.00090304 | 0.014244709 |
| TMEM186 | 454.9308977 | -0.343343569 | 0.141911855 | 0.001257826 | 0.018331954 |
| TTC31 | 1439.613632 | -0.344318868 | 0.151411333 | 0.001876274 | 0.024676318 |
| DOCK10 | 1206.193103 | -0.344992948 | 0.138852292 | 0.001027816 | 0.01574904 |
| PCGF3 | 3622.572177 | -0.345575641 | 0.125579627 | 0.000452095 | 0.00821514 |
| RABL2A | 760.3186721 | -0.34570242 | 0.125659291 | 0.000452702 | 0.00821514 |
| TEAD1 | 8935.685494 | -0.346698777 | 0.166099383 | 0.00301795 | 0.035153315 |
| RESF1 | 1274.399774 | -0.347004001 | 0.145216788 | 0.001331979 | 0.019066739 |
| BEND3 | 580.3806267 | -0.348785836 | 0.163718816 | 0.002659709 | 0.032127254 |
| TCTN3 | 1664.953537 | -0.348805245 | 0.115125928 | 0.000175884 | 0.003911976 |
| ERLIN2 | 2465.219131 | -0.349395473 | 0.110650162 | 0.000112043 | 0.002752964 |
| BDH2 | 1600.759874 | -0.349458869 | 0.162676234 | 0.00252674 | 0.030734398 |
| KRIT1 | 1467.118131 | -0.349504519 | 0.140583269 | 0.000988665 | 0.015295017 |
| DELE1 | 2491.288669 | -0.349864549 | 0.156731306 | 0.00201564 | 0.026064854 |
| NLN | 2175.425876 | -0.350268848 | 0.138226436 | 0.000853013 | 0.013599371 |
| MIR4458HG | 245.9235241 | -0.35205555 | 0.177656392 | 0.003772257 | 0.041530625 |
| GOLM1 | 4788.935449 | -0.352672898 | 0.118113245 | 0.000198239 | 0.004330768 |
| SOX4 | 17454.5699 | -0.352817329 | 0.108164212 | 7.50E-05 | 0.00201272 |

| | | | | | |
|---|---|---|---|---|---|
| MIR1915HG | 265.1819136 | -0.353136124 | 0.167637772 | 0.002738731 | 0.032797246 |
| EFHC1 | 2109.445452 | -0.354902374 | 0.136725977 | 0.000684283 | 0.011423856 |
| CHROMR | 1480.071741 | -0.355373987 | 0.120877287 | 0.000226771 | 0.004798487 |
| FAM217B | 1514.800506 | -0.356404876 | 0.130690755 | 0.000450273 | 0.008205889 |
| NDUFB3 | 899.9740401 | -0.356574042 | 0.172385742 | 0.002942576 | 0.034564515 |
| ABRAXAS1 | 497.6371822 | -0.357782953 | 0.186852658 | 0.004244064 | 0.04529243 |
| SLC5A3 | 1900.349299 | -0.357795175 | 0.19089613 | 0.004663199 | 0.048405029 |
| RBIS | 696.2842181 | -0.358556253 | 0.173755312 | 0.002936563 | 0.034531832 |
| RELA-DT | 165.3223791 | -0.359863079 | 0.1928569 | 0.004681749 | 0.048514836 |
| KCNMB4 | 714.3912206 | -0.360247268 | 0.133765096 | 0.000487195 | 0.008644923 |
| DGCR8 | 2459.417716 | -0.360542628 | 0.124738982 | 0.000257499 | 0.005303615 |
| SLC38A6 | 372.2622765 | -0.360848211 | 0.179343315 | 0.003283875 | 0.037477452 |
| FAM200B | 691.8892194 | -0.360937627 | 0.177011927 | 0.003069377 | 0.03567275 |
| CTPS2 | 1874.899792 | -0.361051813 | 0.129925323 | 0.000369008 | 0.006987059 |
| BBIP1 | 1132.97979 | -0.361366473 | 0.145986459 | 0.000935321 | 0.014664986 |
| OGFOD3 | 1691.39337 | -0.36151189 | 0.175521968 | 0.002903222 | 0.034198417 |
| ZBED5-AS1 | 133.9317033 | -0.361657552 | 0.186923921 | 0.003935501 | 0.042920599 |
| DCTN6 | 969.8694434 | -0.36200265 | 0.178402412 | 0.003122046 | 0.036124066 |
| ANLN | 4321.391474 | -0.362079959 | 0.164101481 | 0.001976266 | 0.025700074 |
| BORCS8 | 374.3406574 | -0.363021931 | 0.174208655 | 0.002701642 | 0.03248347 |
| RAB9B | 205.9893695 | -0.363433967 | 0.182592589 | 0.003398847 | 0.038557479 |
| TMEM19 | 1126.45075 | -0.363553476 | 0.151243197 | 0.001133057 | 0.01689072 |
| PBX1 | 2711.246974 | -0.363583498 | 0.107029745 | 4.19E-05 | 0.001263988 |
| SDF2 | 1707.25494 | -0.363767279 | 0.134625119 | 0.000461957 | 0.00833879 |
| LINC01238 | 208.2550082 | -0.365585445 | 0.177532067 | 0.00282267 | 0.033514193 |
| ANPEP | 524.8821368 | -0.366504305 | 0.171123777 | 0.002271213 | 0.028592098 |
| MIR137HG | 394.7716577 | -0.368071682 | 0.156469038 | 0.001270184 | 0.018434882 |
| BBS10 | 1296.936097 | -0.368629892 | 0.180015195 | 0.002843256 | 0.033720268 |
| SMC4 | 8952.542984 | -0.369419199 | 0.151610176 | 0.000989492 | 0.015296483 |
| ARL4C | 1429.414901 | -0.369481675 | 0.176901745 | 0.002549787 | 0.030978646 |
| AOPEP | 1256.886835 | -0.370014534 | 0.134027864 | 0.000367311 | 0.006969604 |
| DMAC1 | 1103.947265 | -0.370406834 | 0.159703652 | 0.001370737 | 0.019474724 |
| NXT2 | 2338.761838 | -0.370812168 | 0.179527873 | 0.002678418 | 0.032315864 |
| PLK2 | 4173.498875 | -0.371314254 | 0.19867048 | 0.004279662 | 0.045486378 |
| FAM104B | 564.9578425 | -0.371738789 | 0.155504416 | 0.001111703 | 0.016631742 |
| TCTN2 | 516.065721 | -0.371799103 | 0.156718003 | 0.001169938 | 0.017292394 |
| KLHL13 | 7052.512023 | -0.371970533 | 0.145695753 | 0.000689887 | 0.011489863 |
| LEPROT | 4335.169277 | -0.372759436 | 0.195657765 | 0.003896822 | 0.042610016 |
| TMTC2 | 3382.98848 | -0.373946312 | 0.132068959 | 0.000284695 | 0.005729313 |
| TSC22D1 | 5129.228666 | -0.374017606 | 0.11873467 | 9.62E-05 | 0.002417713 |
| ELAC1 | 357.9401533 | -0.374186519 | 0.189951518 | 0.003309684 | 0.037710178 |
| RBL2 | 2915.05465 | -0.37471367 | 0.168079604 | 0.001701914 | 0.022887627 |
| TCTA | 751.8598917 | -0.37514941 | 0.163257417 | 0.001408199 | 0.019840541 |
| PLS3 | 5190.994983 | -0.375996863 | 0.17917426 | 0.002376654 | 0.029492822 |
| GAL3ST4 | 750.8106867 | -0.375999548 | 0.144857209 | 0.000590375 | 0.010190061 |
| CTSA | 4214.035369 | -0.376895128 | 0.146571192 | 0.000631618 | 0.010715949 |
| VANGL1 | 1496.888963 | -0.377669971 | 0.170343289 | 0.001723886 | 0.023069542 |
| STAT5B | 2240.753544 | -0.379081723 | 0.125083968 | 0.000141257 | 0.003303637 |
| GSTM4 | 674.2631872 | -0.37933609 | 0.165614976 | 0.001397401 | 0.019772849 |
| ARSB | 2703.572679 | -0.379574199 | 0.136931166 | 0.00033247 | 0.006471238 |

| | | | | | |
|---|---|---|---|---|---|
| ZNRF3-AS1 | 2.457279567 | -0.381454524 | 0.773069974 | 0.004657321 | 0.048384724 |
| SFPQ | 22062.39123 | -0.381946989 | 0.131181338 | 0.000207523 | 0.004463559 |
| SETMAR | 754.2954595 | -0.382466964 | 0.180503239 | 0.002158757 | 0.027508022 |
| MST1P2 | 151.1719424 | -0.38304691 | 0.191565394 | 0.002897366 | 0.03416325 |
| AMD1 | 4466.068339 | -0.383564448 | 0.140045148 | 0.000360202 | 0.006894456 |
| TRIL | 2642.575253 | -0.383699566 | 0.195512575 | 0.003153545 | 0.036347569 |
| HNRNPA2B1 | 52300.82598 | -0.383714725 | 0.172700862 | 0.001634729 | 0.022155293 |
| GPRC5D-AS1 | 91.67029895 | -0.383855128 | 0.216945081 | 0.004878663 | 0.049800033 |
| SEMA5A | 4776.971017 | -0.384418484 | 0.210071042 | 0.004258814 | 0.04538027 |
| EEPD1 | 420.1706368 | -0.384699309 | 0.186591592 | 0.002457902 | 0.030089854 |
| LRRC8B | 3796.334543 | -0.384754486 | 0.201771527 | 0.003568491 | 0.039961361 |
| LUC7L3 | 14989.96681 | -0.385290546 | 0.145706954 | 0.000478939 | 0.008563902 |
| MTATP6P1 | 13390.36951 | -0.385965752 | 0.197246278 | 0.003147804 | 0.036341478 |
| NDUFS5 | 5554.949749 | -0.386075193 | 0.167655404 | 0.001292119 | 0.01864955 |
| FAM20B | 4941.35235 | -0.38780273 | 0.101265394 | 6.43E-06 | 0.000275705 |
| NLGN1 | 345.9853047 | -0.3880308 | 0.164422475 | 0.001088246 | 0.016398278 |
| NFYA | 4091.857395 | -0.388252911 | 0.102983495 | 8.20E-06 | 0.000331887 |
| CPT1A | 4713.71948 | -0.389526366 | 0.139370795 | 0.000290164 | 0.005799108 |
| ARHGEF3 | 606.7681748 | -0.389799224 | 0.18564465 | 0.002158684 | 0.027508022 |
| ATP8B3 | 348.2183192 | -0.390041345 | 0.19149683 | 0.002522555 | 0.030701371 |
| CASP2 | 3336.992069 | -0.390307645 | 0.115168145 | 3.63E-05 | 0.001123465 |
| NQO1 | 1961.223721 | -0.391554387 | 0.1958509 | 0.002737096 | 0.032796467 |
| SUOX | 963.1228851 | -0.391634609 | 0.144611428 | 0.000377407 | 0.007113843 |
| ZNF180 | 634.9574146 | -0.39320023 | 0.197398623 | 0.002755428 | 0.032865279 |
| POGLUT3 | 3813.133026 | -0.395201055 | 0.141681461 | 0.000285452 | 0.005737882 |
| ZNF33A | 2552.952426 | -0.395530503 | 0.14573575 | 0.000361937 | 0.006909561 |
| PIN4 | 1175.498048 | -0.396176145 | 0.16008034 | 0.000744061 | 0.012226366 |
| FKBP9 | 10054.46599 | -0.39648472 | 0.149296011 | 0.000431572 | 0.007934314 |
| OGG1 | 750.2926303 | -0.397299525 | 0.123697615 | 6.69E-05 | 0.001844019 |
| ARSA | 1408.622893 | -0.398422515 | 0.229413728 | 0.004713703 | 0.048711374 |
| LARP7 | 1442.961927 | -0.400624564 | 0.211392777 | 0.00328905 | 0.037516006 |
| ASB16-AS1 | 440.9451291 | -0.401573091 | 0.168863758 | 0.000949854 | 0.014815306 |
| SLC35A3 | 1149.748122 | -0.403362034 | 0.166675995 | 0.000834204 | 0.013370946 |
| IFT70B | 408.460658 | -0.403461897 | 0.1705177 | 0.00097109 | 0.015056573 |
| TUBB | 38207.79297 | -0.404168418 | 0.164373766 | 0.000742574 | 0.012211543 |
| SAMD15 | 195.7100872 | -0.404310392 | 0.211865445 | 0.003113365 | 0.036063578 |
| EFNB3 | 1579.530938 | -0.404504523 | 0.138962318 | 0.000181999 | 0.004018022 |
| SH3PXD2B | 4945.486918 | -0.405188971 | 0.143827652 | 0.00024638 | 0.005120046 |
| EXOC3-AS1 | 157.7496239 | -0.405547679 | 0.182956884 | 0.001440461 | 0.020204177 |
| FAM210B | 3315.809479 | -0.405649252 | 0.12737634 | 7.05E-05 | 0.00192033 |
| CAT | 1202.195031 | -0.405678197 | 0.135132425 | 0.000133145 | 0.00313499 |
| RBM4B | 1369.045528 | -0.406161023 | 0.13729186 | 0.000153923 | 0.003505658 |
| B4GALT5 | 5401.278808 | -0.406471029 | 0.125196013 | 5.62E-05 | 0.001598997 |
| PDCL3 | 1148.070871 | -0.406842955 | 0.157960877 | 0.00051814 | 0.009108891 |
| MIATNB | 252.498052 | -0.407076849 | 0.166718458 | 0.000766728 | 0.012511139 |
| EPCAM-DT | 205.2348311 | -0.40713541 | 0.166830198 | 0.000769292 | 0.012532502 |
| N6AMT1 | 906.6364777 | -0.407622479 | 0.124534301 | 5.07E-05 | 0.001471828 |
| LDHA | 12926.86853 | -0.407816132 | 0.152500001 | 0.0003815 | 0.007171578 |
| GPC5 | 171.5468293 | -0.408009396 | 0.230569614 | 0.00411792 | 0.044217355 |
| ZNF43 | 1915.851518 | -0.408056796 | 0.196227091 | 0.002016635 | 0.026064854 |

| | | | | | |
|---|---|---|---|---|---|
| TWIST1 | 9048.849753 | -0.40819918 | 0.179229733 | 0.001203958 | 0.0176826 |
| HMGN2P5 | 139.5527147 | -0.408460712 | 0.192026951 | 0.001784275 | 0.023765675 |
| HAGLR | 653.1789742 | -0.409085399 | 0.162057218 | 0.000595718 | 0.010248402 |
| NIT2 | 1476.700318 | -0.40929918 | 0.138230506 | 0.000150019 | 0.003439247 |
| TUBAP2 | 84.96875597 | -0.409315288 | 0.234918754 | 0.00431803 | 0.045815346 |
| EFCAB13 | 117.4251776 | -0.410757306 | 0.242621364 | 0.004722359 | 0.048776692 |
| SFT2D2 | 6429.900965 | -0.410964206 | 0.129924309 | 7.41E-05 | 0.001991061 |
| NDRG4 | 2376.549438 | -0.411056328 | 0.131753307 | 8.62E-05 | 0.002236481 |
| BIVM | 2092.280501 | -0.411137093 | 0.118326148 | 2.35E-05 | 0.000791983 |
| MAP3K5 | 276.9005442 | -0.411185488 | 0.206372267 | 0.002443962 | 0.02998531 |
| BTN3A2 | 1004.955989 | -0.411990087 | 0.13715229 | 0.000127967 | 0.003033577 |
| GLRX | 585.2645964 | -0.412078912 | 0.201961166 | 0.002164436 | 0.027563569 |
| FAM200C | 267.457998 | -0.412682248 | 0.183902138 | 0.001281577 | 0.018535832 |
| BST1 | 146.8611236 | -0.412911814 | 0.205686471 | 0.002331373 | 0.029017122 |
| GALC | 1543.930436 | -0.414622503 | 0.137422553 | 0.000120681 | 0.002903724 |
| RASL10B | 1077.201082 | -0.415168713 | 0.176967955 | 0.000957068 | 0.014905569 |
| PRKCH | 253.6936981 | -0.415219519 | 0.192748614 | 0.001595829 | 0.02172679 |
| CC2D2A | 1080.291602 | -0.415253209 | 0.140809546 | 0.000151331 | 0.003465506 |
| WAC-AS1 | 1299.855549 | -0.416112093 | 0.143730562 | 0.000180159 | 0.003985819 |
| COPG2 | 2657.10198 | -0.416422657 | 0.12545055 | 4.10E-05 | 0.001243839 |
| MYADM | 3284.330697 | -0.417169479 | 0.223946497 | 0.003165805 | 0.036468752 |
| C6orf120 | 1732.772946 | -0.4176933 | 0.138714926 | 0.000121251 | 0.002910719 |
| DOCK11 | 3594.10292 | -0.417992348 | 0.211733004 | 0.002442116 | 0.02998531 |
| MAP3K9-DT | 219.9519423 | -0.418831054 | 0.210219963 | 0.002326735 | 0.029017122 |
| FKBP7 | 1631.641775 | -0.419539416 | 0.146830462 | 0.000200244 | 0.004360897 |
| CLCN3 | 8489.784171 | -0.421060707 | 0.127058487 | 4.10E-05 | 0.001243839 |
| DMAC2L | 297.3581083 | -0.421946142 | 0.165519239 | 0.000514472 | 0.00905965 |
| PRDM10 | 848.0068413 | -0.423163137 | 0.1289787 | 4.58E-05 | 0.00135573 |
| PRAME | 14007.30026 | -0.423663407 | 0.136910013 | 8.85E-05 | 0.002273616 |
| PBX3 | 4553.810213 | -0.424238186 | 0.182986596 | 0.000978696 | 0.015152013 |
| DCBLD2 | 8746.116595 | -0.42448528 | 0.216758724 | 0.002430221 | 0.029908765 |
| MYH10 | 23551.33341 | -0.425002488 | 0.153230614 | 0.000254607 | 0.005254411 |
| FRMD4A | 2472.477755 | -0.425758381 | 0.23837407 | 0.003525476 | 0.039635048 |
| RNF145 | 5139.424104 | -0.426778421 | 0.14125482 | 0.000112032 | 0.002752964 |
| NES | 12565.38086 | -0.42718192 | 0.171804291 | 0.00060041 | 0.010303661 |
| GALNT11 | 2731.989466 | -0.427413463 | 0.111647995 | 5.38E-06 | 0.000241861 |
| SPRY4-AS1 | 110.7008397 | -0.427941226 | 0.196988537 | 0.001408835 | 0.019840541 |
| ATPSCKMT | 663.262946 | -0.428236913 | 0.150786134 | 0.000202583 | 0.004388957 |
| RAB40B | 630.8815081 | -0.428980246 | 0.1772884 | 0.000719217 | 0.01190241 |
| IQCD | 87.25641521 | -0.429108936 | 0.254809445 | 0.004231565 | 0.045205232 |
| PCDHB16 | 265.8602832 | -0.430238006 | 0.20212557 | 0.001552953 | 0.021245755 |
| SLC25A53 | 422.4685833 | -0.430357176 | 0.196719189 | 0.001335087 | 0.019085073 |
| TXNIP | 2759.032319 | -0.430607456 | 0.203148225 | 0.00158441 | 0.021613586 |
| PCDHB9 | 547.8148297 | -0.430638578 | 0.146727302 | 0.000146863 | 0.003388893 |
| PLP1 | 122.3654522 | -0.431765803 | 0.261216653 | 0.004408693 | 0.046526299 |
| ZFHX4-AS1 | 172.1219719 | -0.431961431 | 0.218373843 | 0.002212855 | 0.028026371 |
| HSPA8 | 53528.11352 | -0.432117363 | 0.180186214 | 0.00075059 | 0.012314273 |
| CD302 | 161.2961495 | -0.433233414 | 0.24467513 | 0.003462213 | 0.039131192 |
| VMP1 | 8340.628751 | -0.433799965 | 0.220497921 | 0.0022424 | 0.028280509 |
| DTX3L | 1016.926351 | -0.434623914 | 0.146323655 | 0.000128164 | 0.003034808 |

| | | | | | |
|---|---|---|---|---|---|
| FGD6 | 1070.851805 | -0.434856511 | 0.203651991 | 0.001484744 | 0.020631328 |
| SARS2 | 142.1805565 | -0.436081888 | 0.236220934 | 0.002898601 | 0.03416325 |
| PLD2 | 1248.842731 | -0.436351907 | 0.218460858 | 0.002057308 | 0.026490066 |
| COX16 | 191.9927743 | -0.436387558 | 0.22843781 | 0.002511905 | 0.030607436 |
| DICER1-AS1 | 179.3450958 | -0.436607243 | 0.224473323 | 0.002318918 | 0.028963279 |
| IFT70A | 664.3911857 | -0.436748258 | 0.13486232 | 5.03E-05 | 0.00146487 |
| MTCL1 | 2399.374123 | -0.436801685 | 0.198633916 | 0.001247758 | 0.018210647 |
| EIF4E2 | 3472.929579 | -0.437184422 | 0.153115214 | 0.000184589 | 0.004062324 |
| LINC01237 | 180.3697262 | -0.437264992 | 0.252647251 | 0.003653065 | 0.040652972 |
| TMOD2 | 2744.717995 | -0.437419104 | 0.172424872 | 0.000490474 | 0.008695713 |
| ZNF33B | 2144.646906 | -0.438645752 | 0.136908786 | 5.64E-05 | 0.0016008 |
| MYLK | 3249.946761 | -0.439525332 | 0.228543051 | 0.002391927 | 0.029594428 |
| SLC13A4 | 1077.393443 | -0.439551351 | 0.221607492 | 0.002083164 | 0.026756998 |
| CILK1 | 2133.897017 | -0.439640196 | 0.131990224 | 3.56E-05 | 0.001119307 |
| PCCA-DT | 318.5850553 | -0.440891543 | 0.161065552 | 0.000263272 | 0.005401223 |
| FZD2 | 1504.976433 | -0.441228549 | 0.17005212 | 0.000405467 | 0.007560866 |
| CPPED1 | 1235.762489 | -0.441667997 | 0.130950681 | 3.03E-05 | 0.000977288 |
| PABIR2 | 2567.658545 | -0.441759708 | 0.14857882 | 0.000122814 | 0.002941473 |
| LGR4 | 2372.15377 | -0.441971526 | 0.267922798 | 0.004135718 | 0.04436285 |
| AKAP6 | 1736.362203 | -0.442429786 | 0.159229509 | 0.000229719 | 0.004836367 |
| NUDT13 | 306.2133774 | -0.44291467 | 0.155462345 | 0.000183258 | 0.004037286 |
| B3GNT5 | 1373.830242 | -0.443392907 | 0.148713709 | 0.000118599 | 0.002870156 |
| LAMA4-AS1 | 4.349310827 | -0.443515358 | 0.797965756 | 0.004354318 | 0.046068857 |
| RABL3 | 774.733018 | -0.443975555 | 0.169239885 | 0.000367001 | 0.006969604 |
| ZNF84 | 3702.214546 | -0.444322874 | 0.146866302 | 0.000101992 | 0.002544919 |
| EID1 | 6973.951924 | -0.444944201 | 0.17097599 | 0.000388759 | 0.007281818 |
| GPRASP1 | 754.123736 | -0.445437651 | 0.168186508 | 0.000337807 | 0.006550701 |
| MIR762HG | 95.57680077 | -0.445776716 | 0.246789068 | 0.002959813 | 0.034669486 |
| KRBA2 | 1082.297386 | -0.446903091 | 0.131539834 | 2.71E-05 | 0.000893993 |
| POP5 | 869.8120808 | -0.447566173 | 0.122309369 | 9.94E-06 | 0.000388862 |
| CCDC191 | 680.443028 | -0.447988547 | 0.141371529 | 6.14E-05 | 0.001708824 |
| TOMM5 | 184.2479708 | -0.448827886 | 0.213468416 | 0.001480611 | 0.020587592 |
| PLPP2 | 894.2426301 | -0.44884123 | 0.16038716 | 0.000209239 | 0.004491221 |
| SUMF1 | 1063.778623 | -0.449411822 | 0.163791715 | 0.000247535 | 0.005133838 |
| MLLT3 | 868.0618337 | -0.44950051 | 0.173081858 | 0.000385876 | 0.007234315 |
| CBR4 | 1231.063199 | -0.450538604 | 0.16046295 | 0.000201605 | 0.004372292 |
| SGMS1-AS1 | 239.2976035 | -0.45163176 | 0.156833464 | 0.000159486 | 0.003607877 |
| SLC35A5 | 1986.087328 | -0.451831032 | 0.140333641 | 5.06E-05 | 0.001469643 |
| GLMP | 1712.003632 | -0.451853773 | 0.158430531 | 0.000174082 | 0.003880162 |
| FMN2 | 914.4640785 | -0.452105898 | 0.17628886 | 0.000418922 | 0.007735796 |
| METTL3 | 3172.872922 | -0.452669842 | 0.118940133 | 5.45E-06 | 0.000243564 |
| PDGFD | 1899.530754 | -0.454161994 | 0.21177236 | 0.001293042 | 0.018649992 |
| MYL12A | 8034.020098 | -0.454588302 | 0.213222009 | 0.001330686 | 0.019061307 |
| GYG2 | 320.529694 | -0.4549842 | 0.17990724 | 0.00045768 | 0.008290236 |
| FOXR2 | 594.6412824 | -0.455222257 | 0.164132679 | 0.000219569 | 0.004660261 |
| FAM13A | 377.4927492 | -0.455273202 | 0.172730973 | 0.000334136 | 0.006491567 |
| CASP6 | 698.137168 | -0.455477552 | 0.155338532 | 0.000132261 | 0.003121204 |
| MATCAP1 | 2942.937662 | -0.456122968 | 0.176990078 | 0.000395669 | 0.007397991 |
| SVIL | 6917.424595 | -0.45618046 | 0.283816416 | 0.004054153 | 0.043758034 |
| RUSC1 | 4159.690519 | -0.456785886 | 0.197029518 | 0.000813575 | 0.013110743 |

| | | | | | |
|---|---|---|---|---|---|
| ZNF226 | 1600.002682 | -0.457091933 | 0.136345274 | 3.08E-05 | 0.000991442 |
| CTHRC1 | 2303.860413 | -0.457171601 | 0.30342259 | 0.004770247 | 0.049125547 |
| TSPYL4 | 3137.892252 | -0.457502565 | 0.120991724 | 5.94E-06 | 0.000259933 |
| LINC03015 | 213.0635362 | -0.458795787 | 0.184156043 | 0.000499766 | 0.008837948 |
| NIFK-AS1 | 273.894221 | -0.460201171 | 0.167887063 | 0.000237085 | 0.004963245 |
| CFHR3 | 3.579659202 | -0.460219673 | 0.780376196 | 0.004744966 | 0.048913437 |
| TTN | 1196.63008 | -0.460246053 | 0.172477832 | 0.000295667 | 0.005875352 |
| FAM47E | 82.06038373 | -0.461214849 | 0.23548096 | 0.00192713 | 0.025218119 |
| SAYSD1 | 544.0194917 | -0.461592335 | 0.145617393 | 5.80E-05 | 0.001636114 |
| RPL7AP22 | 2.835640171 | -0.46171672 | 0.869906798 | 0.002748568 | 0.032839729 |
| ABRACL | 812.0666397 | -0.46212234 | 0.191953455 | 0.00062078 | 0.010575036 |
| ERICD | 53.08306676 | -0.462885998 | 0.290799823 | 0.003981749 | 0.043238364 |
| MALAT1 | 282.2574125 | -0.464476779 | 0.288605531 | 0.003819997 | 0.041945658 |
| FANCL | 1589.99332 | -0.465688358 | 0.169092913 | 0.000222102 | 0.004709232 |
| HIF3A | 90.0318822 | -0.465695997 | 0.23995445 | 0.001950954 | 0.02548197 |
| SNAI2 | 1339.109705 | -0.465937068 | 0.221464925 | 0.001333168 | 0.019070697 |
| ALDH1A2 | 48.33698357 | -0.466117995 | 0.307280974 | 0.004415348 | 0.046573002 |
| LINC01963 | 1468.636799 | -0.466305566 | 0.143830287 | 4.43E-05 | 0.001328246 |
| SMPDL3A | 212.2519087 | -0.46665498 | 0.230390066 | 0.001599079 | 0.021752354 |
| TMEM200B | 82.73378819 | -0.466757869 | 0.318123177 | 0.004751876 | 0.048960495 |
| HDAC10 | 114.5639204 | -0.466834123 | 0.309611765 | 0.004458516 | 0.046886262 |
| ING4 | 1769.061115 | -0.468599671 | 0.131311772 | 1.33E-05 | 0.000491776 |
| PARP9 | 713.8074512 | -0.468639141 | 0.185321854 | 0.000427117 | 0.00786626 |
| ITGB5 | 3515.063225 | -0.46887527 | 0.214233989 | 0.001063589 | 0.016126595 |
| FMN1 | 4421.111041 | -0.469116121 | 0.269091168 | 0.002887251 | 0.034067925 |
| TAGAP-AS1 | 138.7095928 | -0.469333673 | 0.215836515 | 0.001098943 | 0.016523711 |
| ZFTA | 2741.544932 | -0.469527926 | 0.150917991 | 6.88E-05 | 0.001880293 |
| COQ10A | 803.6643658 | -0.469563289 | 0.145192348 | 4.50E-05 | 0.001343859 |
| ANK3 | 1235.985327 | -0.469584421 | 0.21610627 | 0.001101619 | 0.016547944 |
| KBTBD3 | 155.4648237 | -0.469811469 | 0.193186187 | 0.000557246 | 0.009738975 |
| TRIM45 | 437.0705501 | -0.47064051 | 0.15163319 | 7.02E-05 | 0.001914383 |
| METTL25 | 154.8962685 | -0.47209605 | 0.221616034 | 0.001206213 | 0.017685467 |
| ZBTB14 | 676.5736173 | -0.473767292 | 0.132942564 | 1.33E-05 | 0.000491848 |
| MPHOSPH6 | 701.3693392 | -0.473822747 | 0.19909001 | 0.000628688 | 0.010683602 |
| TPM4 | 29693.95239 | -0.473844336 | 0.151941459 | 6.60E-05 | 0.001822282 |
| ANKRD42-DT | 141.9909463 | -0.474140674 | 0.222819688 | 0.001198759 | 0.017618629 |
| TMEM131L | 1188.420956 | -0.474678259 | 0.172296506 | 0.000212484 | 0.004537554 |
| FAR2 | 816.704102 | -0.474973906 | 0.143382164 | 3.34E-05 | 0.00105931 |
| BAG2 | 1187.370141 | -0.475184164 | 0.190282511 | 0.000451801 | 0.00821514 |
| PCDHB14 | 1241.944813 | -0.475700054 | 0.221315893 | 0.001127203 | 0.016839514 |
| FDXACB1 | 126.315091 | -0.475970756 | 0.265089259 | 0.002490488 | 0.030399668 |
| BCKDHB | 1208.000545 | -0.476060604 | 0.124925557 | 5.03E-06 | 0.000230987 |
| ZFHX4 | 2798.704458 | -0.476657056 | 0.172941189 | 0.000209825 | 0.004499178 |
| MTURN | 2593.322452 | -0.47668694 | 0.175367602 | 0.00023544 | 0.004941881 |
| DLC1 | 1563.856958 | -0.476768613 | 0.25831853 | 0.002235427 | 0.028253958 |
| MT-TE | 58.00514889 | -0.476980297 | 0.312842216 | 0.004054192 | 0.043758034 |
| ACCS | 813.7649843 | -0.477108178 | 0.200765206 | 0.00062364 | 0.010615096 |
| STMP1 | 2881.630815 | -0.478164435 | 0.144026532 | 3.22E-05 | 0.001027338 |
| CEBPZOS | 2224.212354 | -0.479136703 | 0.166875533 | 0.000145183 | 0.003369042 |
| TOX2 | 438.7950987 | -0.479269476 | 0.193100997 | 0.000461872 | 0.00833879 |

| | | | | | |
|---|---|---|---|---|---|
| MEIS2 | 10445.53045 | -0.479386116 | 0.154324365 | 6.73E-05 | 0.001847575 |
| LTBP1 | 3251.600277 | -0.481579931 | 0.313784887 | 0.003868889 | 0.042371135 |
| DNAAF10 | 437.7288224 | -0.481593285 | 0.188119261 | 0.000366169 | 0.006969604 |
| OLFM3 | 214.236592 | -0.482505172 | 0.238325948 | 0.001457358 | 0.020372772 |
| PNISR | 9204.631702 | -0.482947484 | 0.158617493 | 8.16E-05 | 0.002139077 |
| CEND1 | 726.0757791 | -0.483545798 | 0.214074412 | 0.000820014 | 0.013183989 |
| CHST6 | 103.9215205 | -0.484651345 | 0.34029941 | 0.004492957 | 0.047121177 |
| MARCKS | 18375.17654 | -0.485709962 | 0.15489251 | 5.95E-05 | 0.001672819 |
| FANK1 | 293.5336207 | -0.486175846 | 0.266615484 | 0.002206055 | 0.027957193 |
| SNHG3 | 1728.402355 | -0.486360667 | 0.166007066 | 0.000116945 | 0.002843291 |
| RTCA-AS1 | 301.8512123 | -0.486630197 | 0.209156208 | 0.000676485 | 0.011320828 |
| DACH1 | 604.3786336 | -0.487238939 | 0.270115207 | 0.002280769 | 0.028644983 |
| SHFL | 402.8697232 | -0.487904825 | 0.204881165 | 0.000581263 | 0.010066072 |
| OSGEPL1 | 620.892828 | -0.488277724 | 0.204812365 | 0.000576074 | 0.01000109 |
| SPINT1 | 44.38692447 | -0.489257335 | 0.355369427 | 0.004658926 | 0.048384724 |
| HERPUD2-AS1 | 1162.184396 | -0.489381837 | 0.154682288 | 5.34E-05 | 0.001536566 |
| FUCA1 | 822.7517959 | -0.490693543 | 0.15090287 | 3.93E-05 | 0.001199921 |
| UTRN | 2344.307337 | -0.491240845 | 0.176249865 | 0.000178808 | 0.003968543 |
| CCDC89 | 54.50433637 | -0.491568677 | 0.281968048 | 0.002500562 | 0.030504812 |
| L3HYPDH | 686.7579179 | -0.492050812 | 0.248608426 | 0.001525443 | 0.02101509 |
| SMIM7 | 1173.6656 | -0.492667785 | 0.129981649 | 5.26E-06 | 0.000238288 |
| UXS1 | 1689.885196 | -0.493130467 | 0.121625476 | 1.78E-06 | 9.73E-05 |
| IPO5P1 | 366.9285551 | -0.493241627 | 0.201806081 | 0.00047734 | 0.008549948 |
| MT-ND5 | 91006.00004 | -0.493598842 | 0.191490545 | 0.000328365 | 0.006409252 |
| TMEM108-AS1 | 86.09650161 | -0.495271872 | 0.334843436 | 0.003862889 | 0.042327621 |
| SDC3 | 2372.507334 | -0.495630449 | 0.236172197 | 0.00113562 | 0.016916852 |
| USP13 | 1329.901135 | -0.497864044 | 0.121371131 | 1.45E-06 | 8.29E-05 |
| PM20D1-AS1 | 918.9105347 | -0.49791715 | 0.148848581 | 2.76E-05 | 0.000910349 |
| ANKRD20A17P | 69.82357213 | -0.498203114 | 0.276425741 | 0.002136419 | 0.027323399 |
| TRAF5 | 2660.014554 | -0.500197455 | 0.166047656 | 8.48E-05 | 0.002211325 |
| H1-10-AS1 | 120.1175442 | -0.500307367 | 0.224624783 | 0.000809348 | 0.013062784 |
| TPM4P2 | 60.70579329 | -0.501154101 | 0.260892808 | 0.001640672 | 0.022221424 |
| PLEKHG2 | 5484.01678 | -0.502454097 | 0.237722353 | 0.001053903 | 0.016007827 |
| THG1L | 998.6447279 | -0.502946944 | 0.135358906 | 6.89E-06 | 0.00029128 |
| MT2A | 959.3207057 | -0.503066721 | 0.308362637 | 0.002858404 | 0.033857205 |
| IRF6 | 387.1219581 | -0.50412399 | 0.245981553 | 0.001212035 | 0.017738857 |
| SNRNP35 | 561.971369 | -0.504686212 | 0.169719922 | 9.44E-05 | 0.002386479 |
| ECM1 | 478.9720598 | -0.50512267 | 0.240781861 | 0.001077192 | 0.016266928 |
| UNC5C-AS1 | 396.3979578 | -0.507031347 | 0.22169969 | 0.000671367 | 0.011258073 |
| PTPN18 | 1052.555426 | -0.507229471 | 0.169187173 | 8.64E-05 | 0.002239264 |
| CNPY4 | 1117.054221 | -0.507236044 | 0.143576309 | 1.36E-05 | 0.000500786 |
| ZNF248 | 1463.177562 | -0.50749527 | 0.155803023 | 3.64E-05 | 0.00112526 |
| CTSO | 673.6578278 | -0.507595938 | 0.205068488 | 0.000407751 | 0.007589911 |
| SGIP1 | 5498.177539 | -0.507753135 | 0.224299381 | 0.000709297 | 0.011766211 |
| LINC02614 | 94.06613509 | -0.508291538 | 0.296749118 | 0.002385714 | 0.029552575 |
| ACYP1 | 370.9490767 | -0.508868366 | 0.249943969 | 0.001215904 | 0.017783025 |
| PRDM15 | 804.0765227 | -0.509434357 | 0.146479338 | 1.66E-05 | 0.000594457 |
| AK4 | 278.9467024 | -0.509761315 | 0.196997201 | 0.000295267 | 0.005872999 |
| LAMB1 | 55757.31153 | -0.509913696 | 0.245921393 | 0.001109042 | 0.016615734 |
| PROS1 | 1736.701082 | -0.510396809 | 0.128582648 | 2.47E-06 | 0.000128565 |

| | | | | | |
|---|---|---|---|---|---|
| C11orf71 | 151.8357061 | -0.510633051 | 0.181532338 | 0.000151935 | 0.003467929 |
| MGP | 9416.739799 | -0.511415014 | 0.276455228 | 0.001789307 | 0.023802346 |
| TMSB10 | 23369.33819 | -0.513598891 | 0.182001808 | 0.000145944 | 0.003379202 |
| LBH | 1092.885853 | -0.513871792 | 0.276208817 | 0.001725217 | 0.023069542 |
| ST3GAL5 | 647.1890552 | -0.514034776 | 0.245495558 | 0.001033194 | 0.015797843 |
| SCAMP1-AS1 | 322.2923635 | -0.514249006 | 0.190186732 | 0.000206921 | 0.004459803 |
| LRRIQ1 | 239.0784325 | -0.514796744 | 0.263787219 | 0.001415711 | 0.019923944 |
| TMEM107 | 263.0247504 | -0.515038496 | 0.178131034 | 0.000117245 | 0.00284727 |
| DNER | 270.1399724 | -0.515389094 | 0.250321611 | 0.001111203 | 0.016631742 |
| CNIH3 | 1192.973336 | -0.515749207 | 0.24832456 | 0.001064039 | 0.016126595 |
| ZNF738 | 975.4634192 | -0.515961126 | 0.18005785 | 0.000126475 | 0.003005033 |
| TUBA1A | 36029.88015 | -0.516226749 | 0.158855047 | 3.63E-05 | 0.001123465 |
| SPEF2 | 233.4725928 | -0.517266806 | 0.200680978 | 0.00029303 | 0.00584521 |
| URB1-AS1 | 171.757192 | -0.517446538 | 0.196948389 | 0.000254365 | 0.005254411 |
| KIAA1217 | 1823.458769 | -0.517719234 | 0.262396941 | 0.001328077 | 0.01903699 |
| TFIP11-DT | 175.8602151 | -0.517976506 | 0.223247564 | 0.0005819 | 0.010068743 |
| ZFC3H1 | 3462.53819 | -0.518575602 | 0.178365732 | 0.000109941 | 0.002710892 |
| SYNPO2 | 13902.68751 | -0.519152325 | 0.224884962 | 0.000595708 | 0.010248402 |
| RTL8B | 1090.059518 | -0.519574857 | 0.137477903 | 5.18E-06 | 0.000236332 |
| INSYN2B | 740.6612875 | -0.519651582 | 0.299989427 | 0.002144774 | 0.027363224 |
| C1orf216 | 1288.088287 | -0.519849044 | 0.15987659 | 3.56E-05 | 0.001119307 |
| PRR29 | 59.48729268 | -0.520210678 | 0.289610061 | 0.001889254 | 0.024800171 |
| ABHD15 | 364.3140178 | -0.520435739 | 0.191581785 | 0.000193908 | 0.004252647 |
| MLLT11 | 8254.720034 | -0.522049673 | 0.173147407 | 7.72E-05 | 0.002058765 |
| DNAAF11 | 246.9737411 | -0.523174518 | 0.202173913 | 0.000276824 | 0.005612675 |
| PROCA1 | 251.7724594 | -0.523829168 | 0.167906984 | 5.47E-05 | 0.001567834 |
| ATAT1 | 2299.347022 | -0.524605084 | 0.144431159 | 8.99E-06 | 0.000357634 |
| TMEM17 | 221.0034172 | -0.525185896 | 0.18622323 | 0.000139581 | 0.003268101 |
| TRIB1 | 782.1121718 | -0.525899235 | 0.218144508 | 0.000441535 | 0.008081904 |
| CTTN-DT | 67.6305751 | -0.525900195 | 0.249640042 | 0.000937684 | 0.014680312 |
| GMEB1 | 882.1270399 | -0.527222893 | 0.152047025 | 1.63E-05 | 0.000589552 |
| TEAD2 | 3330.508351 | -0.527281162 | 0.144412035 | 8.31E-06 | 0.000335227 |
| BRPF3-AS1 | 56.28418954 | -0.5274832 | 0.287318222 | 0.001671225 | 0.022547509 |
| SMARCE1 | 623.5406342 | -0.529261832 | 0.141360498 | 5.81E-06 | 0.000256158 |
| MYOM1 | 115.8597037 | -0.529670977 | 0.396951973 | 0.003828269 | 0.041992334 |
| LRRC20 | 868.3762114 | -0.529711237 | 0.191282154 | 0.000159302 | 0.003607877 |
| PCDH10 | 12634.78916 | -0.530069541 | 0.341245709 | 0.002760374 | 0.03286796 |
| SNRPEP4 | 117.1964342 | -0.532725654 | 0.254143267 | 0.000924357 | 0.014558977 |
| PLAT | 7771.672355 | -0.535810535 | 0.333709397 | 0.002442068 | 0.02998531 |
| FUT11 | 1352.13221 | -0.536083733 | 0.118788434 | 2.28E-07 | 1.72E-05 |
| CD59 | 4969.677974 | -0.536200202 | 0.257429063 | 0.000934361 | 0.014664986 |
| CHRM2 | 307.746011 | -0.537376214 | 0.289949318 | 0.001521478 | 0.020974306 |
| SAMD9L | 451.3339096 | -0.537748529 | 0.247155815 | 0.000746317 | 0.012253804 |
| LINC01852 | 33.6259634 | -0.539226006 | 0.379708439 | 0.003211673 | 0.036834588 |
| ENDOD1 | 1494.494601 | -0.540525098 | 0.289025747 | 0.001443446 | 0.020218898 |
| ZNF341 | 213.0035113 | -0.541416659 | 0.181177091 | 7.79E-05 | 0.002070595 |
| MAGEA10 | 46.31967882 | -0.543058231 | 0.442404497 | 0.004034282 | 0.043648921 |
| HECW2-AS1 | 51.72341843 | -0.54312832 | 0.507997193 | 0.004688853 | 0.048550669 |
| TNFSF12 | 235.4280437 | -0.54456117 | 0.221318527 | 0.000352292 | 0.006783032 |
| ZNF471 | 699.0249305 | -0.544827051 | 0.249149544 | 0.00069939 | 0.011611097 |

| | | | | | |
|---|---|---|---|---|---|
| NLK | 872.9082961 | -0.545961948 | 0.172473379 | 4.34E-05 | 0.001305683 |
| CBLN4 | 9.238665015 | -0.546231497 | 0.561857633 | 0.004871188 | 0.049754295 |
| LINC00342 | 1456.201443 | -0.547057758 | 0.225085941 | 0.000376468 | 0.007102562 |
| CYP1B1-AS1 | 48.0219697 | -0.547741503 | 0.363915651 | 0.002684335 | 0.032349879 |
| MAP2K6 | 1859.405626 | -0.548577506 | 0.19896282 | 0.000151747 | 0.003467929 |
| MMP16 | 3706.670877 | -0.549296494 | 0.173346915 | 4.24E-05 | 0.001277913 |
| NELL2 | 48.22423056 | -0.550992467 | 0.371085787 | 0.002719037 | 0.03263626 |
| GUCY1B1 | 1015.498755 | -0.551434184 | 0.289329924 | 0.001258941 | 0.01833271 |
| ALDH5A1 | 1186.583375 | -0.551650126 | 0.121392472 | 1.95E-07 | 1.52E-05 |
| SNCAIP | 70.51630441 | -0.553243263 | 0.304754634 | 0.001495867 | 0.020716969 |
| DDX5 | 54971.15643 | -0.55354324 | 0.154000847 | 9.58E-06 | 0.000375945 |
| FKBP5 | 1651.84047 | -0.55385701 | 0.159835027 | 1.52E-05 | 0.000553905 |
| FNDC11 | 198.2113426 | -0.554219671 | 0.299074004 | 0.001379829 | 0.019577261 |
| PLAUR | 507.4979975 | -0.554821282 | 0.176772461 | 4.58E-05 | 0.00135573 |
| LINC02909 | 7.069942219 | -0.554932597 | 0.68425135 | 0.004535631 | 0.047410741 |
| ZFAND2A-DT | 227.4419127 | -0.555886646 | 0.217020024 | 0.00025422 | 0.005254411 |
| ACTBP2 | 63.2256012 | -0.556527078 | 0.380595834 | 0.002726441 | 0.032687557 |
| SRD5A3 | 120.5136087 | -0.556533018 | 0.259884882 | 0.000729428 | 0.012042772 |
| HDAC9 | 9779.168566 | -0.558138543 | 0.171679662 | 3.13E-05 | 0.001004178 |
| SRSF5 | 7690.455654 | -0.558700176 | 0.130053673 | 5.82E-07 | 3.78E-05 |
| SEMA6A | 1697.142275 | -0.558731032 | 0.183504556 | 6.09E-05 | 0.00170039 |
| NIPSNAP3B | 370.9429599 | -0.55894428 | 0.157596585 | 1.12E-05 | 0.000432774 |
| PXYLP1 | 1112.897373 | -0.559095776 | 0.136587169 | 1.36E-06 | 7.98E-05 |
| PCDHGA10 | 58.6127604 | -0.559288299 | 0.302078176 | 0.001344637 | 0.019169103 |
| CHML | 3257.692153 | -0.559535404 | 0.161476045 | 1.50E-05 | 0.000545889 |
| HNRNPH2 | 3880.6372 | -0.560131117 | 0.159694467 | 1.29E-05 | 0.000482033 |
| GUCY1A1 | 989.8348611 | -0.560243783 | 0.231379526 | 0.000360486 | 0.006894456 |
| ZNF217 | 3750.98883 | -0.560461972 | 0.329852332 | 0.001789123 | 0.023802346 |
| ASIC1 | 433.3632891 | -0.562215657 | 0.199119099 | 0.000117659 | 0.002854005 |
| RCAN3 | 675.0818932 | -0.562597693 | 0.123637877 | 1.88E-07 | 1.48E-05 |
| FTX | 5305.718588 | -0.563381867 | 0.194809928 | 9.55E-05 | 0.002403905 |
| ARSJ | 1901.768239 | -0.564292201 | 0.282287615 | 0.000961026 | 0.014933803 |
| GTF2H5 | 844.5830649 | -0.569567848 | 0.15468849 | 6.63E-06 | 0.000282574 |
| PJVK | 219.5438568 | -0.569880081 | 0.175217313 | 2.99E-05 | 0.000970315 |
| ARPC5 | 13241.09577 | -0.570073281 | 0.191019196 | 7.02E-05 | 0.001914383 |
| PDE3A | 2741.511051 | -0.570401929 | 0.217534365 | 0.000200881 | 0.004370198 |
| GM2A | 1429.810389 | -0.570816042 | 0.115467628 | 2.89E-08 | 2.95E-06 |
| LINC01410 | 492.4070722 | -0.571825728 | 0.202542005 | 0.00011286 | 0.00276977 |
| GLB1L | 248.8781926 | -0.572328897 | 0.201568534 | 0.000107412 | 0.002654789 |
| MEGF6 | 1414.229335 | -0.573240814 | 0.277612793 | 0.000791564 | 0.012815363 |
| ARPIN | 1205.687701 | -0.575184868 | 0.116130749 | 2.75E-08 | 2.85E-06 |
| CNTNAP2 | 12790.07048 | -0.576319744 | 0.354895663 | 0.001869769 | 0.024646218 |
| PPFIA4 | 426.5416687 | -0.576889656 | 0.200543846 | 9.43E-05 | 0.002386479 |
| RNF207 | 452.5202624 | -0.577221613 | 0.214954936 | 0.000163295 | 0.003682965 |
| ZNF493 | 698.0789065 | -0.578172887 | 0.157285173 | 6.62E-06 | 0.000282574 |
| SLC9A3-AS1 | 454.3283019 | -0.579422724 | 0.302031095 | 0.001047211 | 0.015940969 |
| ARGLU1 | 14655.23915 | -0.580018242 | 0.157388846 | 6.36E-06 | 0.000274644 |
| LINC00460 | 81.35296694 | -0.580122538 | 0.342322821 | 0.001613239 | 0.021906696 |
| PCDHB11 | 54.63166966 | -0.580237636 | 0.297347718 | 0.000973284 | 0.015079399 |
| C2orf16 | 197.5938829 | -0.580567059 | 0.175425652 | 2.38E-05 | 0.000799539 |

| | | | | | |
|---|---|---|---|---|---|
| LINC00562 | 19.65754431 | -0.581720591 | 0.44516884 | 0.002958376 | 0.034669486 |
| SDC2 | 5101.305205 | -0.583593935 | 0.302733508 | 0.001003719 | 0.015482038 |
| FGF14-AS2 | 83.0179311 | -0.583700289 | 0.258394602 | 0.000477969 | 0.00855388 |
| MT-ND6 | 23139.85599 | -0.583706724 | 0.211344645 | 0.000127702 | 0.003030741 |
| TMEM35A | 339.7048778 | -0.583756179 | 0.283738282 | 0.000759854 | 0.012427213 |
| RBPMS | 1377.946267 | -0.584042958 | 0.158743743 | 6.43E-06 | 0.000275705 |
| SHTN1 | 2066.596233 | -0.587854132 | 0.217342058 | 0.000147174 | 0.003388893 |
| ANXA2P2 | 2207.026966 | -0.587855503 | 0.345989416 | 0.001533122 | 0.021093057 |
| PAN3-AS1 | 50.74100202 | -0.588696592 | 0.273960701 | 0.000602426 | 0.010319629 |
| LINC01011 | 58.27041743 | -0.590507017 | 0.266218288 | 0.000508039 | 0.008969063 |
| SNHG25 | 41.2400567 | -0.592978807 | 0.339871984 | 0.00136937 | 0.019474724 |
| COL1A2 | 344.7747412 | -0.593039449 | 0.406333919 | 0.002236813 | 0.028253958 |
| LINC00938 | 551.3685698 | -0.593171217 | 0.137689531 | 5.19E-07 | 3.47E-05 |
| ZNF559 | 829.9187086 | -0.594797535 | 0.172113187 | 1.38E-05 | 0.000504721 |
| ZFP14 | 1008.722783 | -0.59534445 | 0.155476781 | 3.55E-06 | 0.000174507 |
| LINC01285 | 63.5691673 | -0.595615069 | 0.301646386 | 0.000852028 | 0.013599371 |
| GABPB1-IT1 | 985.4933837 | -0.595775477 | 0.139792444 | 6.27E-07 | 4.04E-05 |
| ZFPM2 | 117.1697419 | -0.596114014 | 0.371910703 | 0.001739296 | 0.023225831 |
| SPTSSB | 744.361124 | -0.596865736 | 0.176710769 | 1.78E-05 | 0.000633091 |
| LINC00899 | 26.53972674 | -0.597204747 | 0.421928826 | 0.00233978 | 0.029087086 |
| TTC14 | 2117.538664 | -0.597824376 | 0.12885515 | 1.19E-07 | 9.95E-06 |
| ITGB3 | 404.3690567 | -0.598250216 | 0.392854521 | 0.001966958 | 0.025594965 |
| CNIH3-AS2 | 66.95895423 | -0.598628087 | 0.380845287 | 0.001809357 | 0.024023148 |
| SNN | 929.3630812 | -0.598661778 | 0.181480733 | 2.30E-05 | 0.00077729 |
| MOXD1 | 10818.62056 | -0.601082012 | 0.259585933 | 0.000380861 | 0.007166026 |
| AADAT | 1795.687016 | -0.601559827 | 0.154724393 | 2.79E-06 | 0.00014261 |
| ANGPTL4 | 386.3127597 | -0.601768288 | 0.690129671 | 0.003650862 | 0.040652972 |
| MIR4453HG | 218.893998 | -0.602348595 | 0.165673524 | 7.11E-06 | 0.000298633 |
| OSBPL3 | 2081.59977 | -0.602626809 | 0.266906609 | 0.000434065 | 0.007973137 |
| TMEM276 | 166.1527464 | -0.603652354 | 0.209070224 | 8.12E-05 | 0.002134917 |
| BDNF | 1968.965429 | -0.604523004 | 0.538055202 | 0.003246329 | 0.037109785 |
| PDE5A | 823.4470868 | -0.604812947 | 0.167085087 | 7.46E-06 | 0.00031001 |
| F2R | 24620.41124 | -0.604966777 | 0.405737628 | 0.001998623 | 0.025927832 |
| ADAMTSL1 | 152.3174227 | -0.605165671 | 0.29361693 | 0.000671657 | 0.011258073 |
| PRXL2C | 808.3282071 | -0.607407151 | 0.182970452 | 2.08E-05 | 0.000722221 |
| LECT2 | 811.1997577 | -0.607584858 | 0.660224557 | 0.003573047 | 0.039984993 |
| PCDHGA6 | 138.8380485 | -0.608100559 | 0.251774348 | 0.000287207 | 0.005756547 |
| DCAF16 | 2936.502674 | -0.610867161 | 0.140447971 | 4.17E-07 | 2.92E-05 |
| TCAF1 | 6067.847173 | -0.61096252 | 0.130938845 | 1.02E-07 | 9.05E-06 |
| HOATZ | 84.35127736 | -0.613252035 | 0.408740701 | 0.001886131 | 0.02477475 |
| GRAMD1C | 170.6290162 | -0.613374727 | 0.184109095 | 1.95E-05 | 0.000681134 |
| RPL7AP43 | 35.22104373 | -0.613717375 | 0.351929413 | 0.00122688 | 0.017930996 |
| TMEM198B | 1266.081892 | -0.614878444 | 0.213862119 | 8.01E-05 | 0.002116273 |
| EMB | 1733.983366 | -0.615510937 | 0.262270866 | 0.00032774 | 0.006403036 |
| S100A4 | 328.9045032 | -0.615729748 | 0.261679835 | 0.000322494 | 0.006318279 |
| ANXA2 | 40061.85361 | -0.61778755 | 0.342873293 | 0.00107466 | 0.016241474 |
| ILF3-DT | 572.7838321 | -0.617886312 | 0.16637785 | 5.08E-06 | 0.000232269 |
| TTC34 | 42.50201594 | -0.618345673 | 0.382092538 | 0.001505425 | 0.020814573 |
| PFN2 | 9518.034996 | -0.619435929 | 0.133005241 | 1.04E-07 | 9.16E-06 |
| LINC03013 | 22.96145835 | -0.620255292 | 0.519862624 | 0.002798225 | 0.033261769 |

| | | | | | |
|---|---|---|---|---|---|
| PCDHB7 | 174.0078768 | -0.620558285 | 0.203517632 | 4.65E-05 | 0.001366514 |
| PYROXD2 | 357.1258159 | -0.6232374 | 0.271960241 | 0.000360097 | 0.006894456 |
| SMIM30 | 1649.277871 | -0.624847475 | 0.175970753 | 8.87E-06 | 0.00035438 |
| FJX1 | 909.8089856 | -0.625129514 | 0.274680285 | 0.000370336 | 0.00699951 |
| PCDH18 | 4757.328634 | -0.625390606 | 0.221086798 | 8.72E-05 | 0.002256671 |
| SEMA6C | 1296.351641 | -0.626781309 | 0.257821117 | 0.000251341 | 0.005207599 |
| ADGRB3-DT | 338.8646163 | -0.626849101 | 0.249909506 | 0.000206468 | 0.004454642 |
| OBSCN-AS1 | 102.0544068 | -0.627271795 | 0.249577118 | 0.000203384 | 0.004401731 |
| PLCB1 | 1474.787817 | -0.628611038 | 0.153146907 | 1.10E-06 | 6.56E-05 |
| MDH1B | 111.3514349 | -0.629633776 | 0.216774183 | 6.87E-05 | 0.001880293 |
| LINC03072 | 1017.070359 | -0.629732831 | 0.168831614 | 4.60E-06 | 0.000216701 |
| TMEM60 | 1130.0986 | -0.629803126 | 0.183753553 | 1.33E-05 | 0.000491848 |
| THBS2-AS1 | 62.10456567 | -0.630166865 | 0.40096644 | 0.001536983 | 0.021130913 |
| PAXIP1-AS2 | 355.7587588 | -0.631327257 | 0.176783836 | 8.07E-06 | 0.000328438 |
| PRSS35 | 999.2871331 | -0.631954061 | 0.57012258 | 0.002885588 | 0.034067553 |
| GNG2 | 3141.700397 | -0.634607533 | 0.136772592 | 1.09E-07 | 9.37E-06 |
| TNFRSF11A | 31.74335913 | -0.636443588 | 0.414417922 | 0.001583129 | 0.021610231 |
| AP3M2 | 2618.20214 | -0.636547037 | 0.182365838 | 1.05E-05 | 0.000407469 |
| APOL6 | 101.321354 | -0.636842818 | 0.275966672 | 0.000323449 | 0.006330631 |
| GSTM2 | 862.1511436 | -0.637047549 | 0.14922384 | 5.44E-07 | 3.60E-05 |
| CCDC9B | 546.5439129 | -0.63796439 | 0.243843156 | 0.000147454 | 0.003391608 |
| MEST | 135.5317744 | -0.638940644 | 0.201528754 | 2.96E-05 | 0.000963355 |
| RNU6-1053P | 39.1768027 | -0.639158556 | 0.322011273 | 0.000662865 | 0.011146493 |
| MYL6 | 11901.96452 | -0.639332943 | 0.205934718 | 3.62E-05 | 0.001123465 |
| CHURC1 | 1367.455941 | -0.639780467 | 0.198843323 | 2.54E-05 | 0.000848421 |
| YWHAZP4 | 59.48297321 | -0.641139744 | 0.267136748 | 0.000252797 | 0.005232581 |
| COPS8-DT | 64.41627809 | -0.642242668 | 0.294302334 | 0.000422303 | 0.007784466 |
| MAGI2 | 997.2267378 | -0.642668372 | 0.177851111 | 6.66E-06 | 0.00028259 |
| PLXNC1 | 947.4886521 | -0.643494384 | 0.342141185 | 0.000804619 | 0.013006549 |
| PCDHB10 | 968.4833846 | -0.644320139 | 0.153384839 | 7.05E-07 | 4.46E-05 |
| DUBR | 638.0691271 | -0.645307934 | 0.163691794 | 1.96E-06 | 0.000105058 |
| PLEKHA8P1 | 389.8327278 | -0.647153642 | 0.21827651 | 5.32E-05 | 0.00153343 |
| TMTC4 | 2613.44808 | -0.648053447 | 0.133992512 | 4.18E-08 | 4.03E-06 |
| RBM43 | 404.3430423 | -0.648987219 | 0.179695447 | 6.54E-06 | 0.000279877 |
| VWA1 | 267.2188445 | -0.649674919 | 0.326242795 | 0.000619911 | 0.010568846 |
| HMGXB4 | 1772.979115 | -0.650220377 | 0.109698912 | 1.25E-10 | 2.18E-08 |
| RAB7B | 126.9837876 | -0.654199672 | 0.493630886 | 0.002016797 | 0.026064854 |
| MBNL1-AS1 | 104.0310867 | -0.654620144 | 0.351147485 | 0.000786972 | 0.012760792 |
| ST6GALNAC5 | 948.302676 | -0.655872356 | 0.225419501 | 6.01E-05 | 0.001684943 |
| TMEM168 | 2714.125891 | -0.655959313 | 0.142970355 | 1.28E-07 | 1.06E-05 |
| H2BC21 | 533.3604214 | -0.656070843 | 0.140812431 | 9.31E-08 | 8.31E-06 |
| CD226 | 39.42415247 | -0.657859066 | 0.34034972 | 0.000672959 | 0.011270855 |
| SYT4 | 2640.906133 | -0.658863675 | 0.167311069 | 1.90E-06 | 0.000102353 |
| ZMYND8 | 2172.424453 | -0.662854744 | 0.120532797 | 1.36E-09 | 2.02E-07 |
| PSMA3-AS1 | 2596.318858 | -0.663851728 | 0.155347968 | 4.87E-07 | 3.34E-05 |
| ACTBP11 | 58.06657007 | -0.663976054 | 0.28997892 | 0.000294959 | 0.005872999 |
| HIC1 | 593.1161215 | -0.664496199 | 0.409211648 | 0.001186559 | 0.0174639 |
| ABCG4 | 101.1725767 | -0.66468467 | 0.251566996 | 0.000121081 | 0.002909991 |
| HS2ST1 | 4151.141738 | -0.665477589 | 0.145035237 | 1.24E-07 | 1.03E-05 |
| PCDHB8 | 238.3592898 | -0.665596651 | 0.203516128 | 1.92E-05 | 0.000671529 |

| | | | | | |
|---|---|---|---|---|---|
| VCL | 12122.14259 | -0.665963396 | 0.351156395 | 0.000697161 | 0.011583304 |
| NRBP2 | 1924.115087 | -0.666132659 | 0.311684862 | 0.000415224 | 0.007690229 |
| LYPD6 | 655.6849371 | -0.666536939 | 0.145027475 | 1.19E-07 | 9.98E-06 |
| CNN2 | 2330.152775 | -0.66714955 | 0.190218484 | 8.72E-06 | 0.000349572 |
| RAD51-AS1 | 489.9328837 | -0.667306764 | 0.150751063 | 2.51E-07 | 1.87E-05 |
| PAM | 17005.61522 | -0.667899667 | 0.288809847 | 0.000274013 | 0.005588302 |
| MRNIP-DT | 53.02599538 | -0.66813675 | 0.334464545 | 0.000558309 | 0.00974941 |
| OLFML3 | 315.8345044 | -0.668458148 | 0.248046871 | 0.000103255 | 0.002567251 |
| EDNRA | 11528.36701 | -0.669629393 | 0.124557848 | 2.58E-09 | 3.59E-07 |
| DARS1-AS1 | 22.01620929 | -0.670784647 | 0.452395189 | 0.001469307 | 0.020498653 |
| SGCD | 300.2155085 | -0.670943129 | 0.235737591 | 6.72E-05 | 0.001847404 |
| CHST7 | 349.9533301 | -0.675747181 | 0.216781225 | 2.96E-05 | 0.000963917 |
| ATP2B4 | 6650.140917 | -0.676609367 | 0.347348187 | 0.000595297 | 0.010248402 |
| NIPAL4-DT | 89.41543391 | -0.676811264 | 0.410438374 | 0.001070347 | 0.016210443 |
| WWTR1 | 2790.391704 | -0.677788569 | 0.269823454 | 0.000159621 | 0.003607877 |
| TLE4 | 6022.14486 | -0.678521492 | 0.205184655 | 1.61E-05 | 0.000583155 |
| ZNF345 | 342.8924256 | -0.678715374 | 0.178732777 | 2.97E-06 | 0.000150011 |
| SCARA3 | 575.0894362 | -0.679316275 | 0.266267272 | 0.000143271 | 0.003339525 |
| TOR4A | 348.8514364 | -0.679928972 | 0.327943993 | 0.000447593 | 0.008164169 |
| ADAMTS1 | 2977.049688 | -0.679950666 | 0.262550701 | 0.000129244 | 0.003053464 |
| NDN | 473.1117907 | -0.680626097 | 0.236265609 | 5.82E-05 | 0.001640985 |
| HYAL2 | 1542.496957 | -0.681041484 | 0.160840872 | 5.36E-07 | 3.58E-05 |
| BDNF-AS | 180.9201684 | -0.683528596 | 0.348643203 | 0.000560926 | 0.009778747 |
| COPG2IT1 | 59.71779137 | -0.683594168 | 0.349347442 | 0.000565217 | 0.009833611 |
| NLGN3 | 438.9529207 | -0.687898277 | 0.188675261 | 4.93E-06 | 0.000227463 |
| SUCNR1 | 4000.483003 | -0.689405772 | 0.684781652 | 0.002395822 | 0.029625065 |
| SNHG4 | 705.5054353 | -0.689711126 | 0.142847841 | 3.73E-08 | 3.70E-06 |
| CEP192-DT | 67.34251925 | -0.692163177 | 0.274040937 | 0.000143901 | 0.003346751 |
| C1orf115 | 207.7816836 | -0.692181591 | 0.240195128 | 5.50E-05 | 0.00157402 |
| MIR4477B | 195.0852196 | -0.692438592 | 0.255054772 | 8.74E-05 | 0.002257814 |
| RBM12 | 4044.382593 | -0.693256395 | 0.110561797 | 1.32E-11 | 2.89E-09 |
| FPGT | 510.1845911 | -0.693712723 | 0.163574967 | 4.95E-07 | 3.36E-05 |
| NIPSNAP3A | 1190.260918 | -0.696916991 | 0.208277954 | 1.31E-05 | 0.000487293 |
| ZNF491 | 217.6366055 | -0.69800743 | 0.183863386 | 2.75E-06 | 0.000140946 |
| TAS2R14 | 143.9500938 | -0.698053734 | 0.238093189 | 4.64E-05 | 0.001366492 |
| BAALC | 345.971894 | -0.698953182 | 0.215967263 | 1.84E-05 | 0.000652083 |
| C1QL4 | 80.02683706 | -0.69967029 | 0.302758472 | 0.000236207 | 0.004952992 |
| FZD7 | 2874.365229 | -0.699950031 | 0.194865407 | 5.65E-06 | 0.000250745 |
| MIR221 | 11.53998566 | -0.700588392 | 0.722001056 | 0.002310058 | 0.028872274 |
| RNASEL | 293.4320779 | -0.70081611 | 0.41136874 | 0.000867171 | 0.013783009 |
| MIR222HG | 146.5299837 | -0.700848253 | 0.519860462 | 0.001591044 | 0.021675764 |
| ANKRD29 | 39.34286214 | -0.702273921 | 0.515363325 | 0.001547765 | 0.021224604 |
| LINC00324 | 37.4862217 | -0.705661254 | 0.352348739 | 0.000462531 | 0.00834193 |
| AHSA2P | 3321.544641 | -0.708481731 | 0.181584885 | 1.77E-06 | 9.73E-05 |
| MYCT1 | 1529.044407 | -0.712284967 | 0.362021705 | 0.000483967 | 0.008606557 |
| C1QTNF6 | 2152.742319 | -0.713517729 | 0.205382041 | 7.96E-06 | 0.000324796 |
| EOLA2-DT | 151.6226411 | -0.71357476 | 0.258936968 | 7.08E-05 | 0.001925901 |
| ADAM23 | 1650.379877 | -0.713983133 | 0.145935136 | 2.47E-08 | 2.57E-06 |
| MIR23B | 9.35985969 | -0.714351087 | 0.683205548 | 0.002113384 | 0.027095166 |
| NRAD1 | 5.145535335 | -0.715540469 | 0.931848428 | 0.001791101 | 0.023811041 |

| | | | | | |
|---|---|---|---|---|---|
| DEPP1 | 114.0751157 | -0.71650316 | 0.358537612 | 0.000444517 | 0.008121674 |
| PRECSIT | 139.7836523 | -0.720016478 | 0.508186377 | 0.001323001 | 0.018990298 |
| GJA1 | 1588.775634 | -0.722039192 | 0.20760321 | 7.56E-06 | 0.000312058 |
| SYTL4 | 142.4141485 | -0.722219309 | 0.271972241 | 8.92E-05 | 0.002282886 |
| EFNA4 | 513.9211208 | -0.723256756 | 0.184836134 | 1.59E-06 | 8.94E-05 |
| GLCE | 1548.429626 | -0.724900085 | 0.206385895 | 6.64E-06 | 0.000282574 |
| PDCD6IP-DT | 187.1808926 | -0.726675595 | 0.236621435 | 2.70E-05 | 0.000892295 |
| CEACAM1 | 65.46006001 | -0.727186922 | 0.37522893 | 0.000482349 | 0.008589547 |
| BMP5 | 106.0384553 | -0.72766359 | 0.279299301 | 9.93E-05 | 0.002489866 |
| GRIA4 | 124.1753465 | -0.727907007 | 0.261187371 | 6.09E-05 | 0.00170039 |
| ZNF436 | 1985.356094 | -0.730051173 | 0.142356137 | 7.10E-09 | 8.52E-07 |
| BTD | 869.7246078 | -0.731471416 | 0.160842515 | 1.12E-07 | 9.50E-06 |
| CDH18 | 176.4813494 | -0.731509458 | 0.241167316 | 2.96E-05 | 0.000963355 |
| ADGRL3 | 489.2431831 | -0.731783401 | 0.187364509 | 1.57E-06 | 8.86E-05 |
| LINC00173 | 18.83716175 | -0.732755508 | 0.489831901 | 0.001102932 | 0.016547944 |
| ACTG1 | 68051.51547 | -0.73464694 | 0.187511867 | 1.48E-06 | 8.41E-05 |
| SEMA3D | 446.872196 | -0.737688317 | 0.256695901 | 4.56E-05 | 0.001354019 |
| IQGAP2 | 4766.148044 | -0.738130148 | 0.206406864 | 5.02E-06 | 0.000230987 |
| PON2 | 2550.151706 | -0.739795388 | 0.158889263 | 6.59E-08 | 6.17E-06 |
| TRAM1L1 | 345.3189763 | -0.741780623 | 0.184649728 | 9.75E-07 | 5.90E-05 |
| C1GALT1C1L | 175.9064171 | -0.742065709 | 0.23295178 | 1.78E-05 | 0.000631546 |
| ITGB8 | 498.8247132 | -0.742486519 | 0.214524623 | 7.28E-06 | 0.000304876 |
| CCDC18-AS1 | 875.0717834 | -0.745486138 | 0.20066927 | 2.97E-06 | 0.000150011 |
| SLC40A1 | 417.3881139 | -0.746938512 | 0.197121875 | 2.26E-06 | 0.00011924 |
| ZNF778-DT | 233.0031242 | -0.749635571 | 0.157366826 | 3.80E-08 | 3.73E-06 |
| ADAMTS5 | 3503.193861 | -0.752121516 | 0.376073326 | 0.000379757 | 0.007151686 |
| ST6GAL2-IT1 | 44.29195967 | -0.752196667 | 0.389261441 | 0.000438947 | 0.00804865 |
| ATP8B1-AS1 | 73.38354916 | -0.753765044 | 0.257003659 | 3.58E-05 | 0.001121656 |
| TMSB4XP4 | 138.0430687 | -0.757606996 | 0.426597878 | 0.000592945 | 0.010225982 |
| SEMA3A | 1031.139717 | -0.758878735 | 0.167741463 | 1.08E-07 | 9.37E-06 |
| ADAM21 | 107.152286 | -0.761200822 | 0.246515396 | 2.19E-05 | 0.00074818 |
| DHFR2 | 409.9011529 | -0.766761942 | 0.154660322 | 1.36E-08 | 1.51E-06 |
| C4orf3 | 2721.85904 | -0.769120656 | 0.156842118 | 1.75E-08 | 1.89E-06 |
| NEAT1 | 19506.13181 | -0.770727968 | 0.310699118 | 0.000113152 | 0.002770481 |
| PGBD2 | 292.1804919 | -0.771141265 | 0.200736208 | 1.66E-06 | 9.19E-05 |
| LINC01165 | 28.75726494 | -0.772207496 | 0.394754189 | 0.000384614 | 0.007217126 |
| ROBO2 | 147.5291916 | -0.775014595 | 0.229536869 | 8.30E-06 | 0.000335227 |
| EDIL3-DT | 243.9190153 | -0.777473613 | 0.340897079 | 0.000180018 | 0.003985819 |
| MIR3117 | 22.07186958 | -0.777956485 | 0.603352864 | 0.001282781 | 0.0185404 |
| PTPRD-AS1 | 33.55012828 | -0.778197163 | 0.394816859 | 0.000363723 | 0.006935486 |
| HENMT1 | 24.63479014 | -0.78054192 | 0.490171777 | 0.000773142 | 0.01256588 |
| TRERF1 | 1298.433804 | -0.783145991 | 0.200860411 | 1.26E-06 | 7.42E-05 |
| TMSB4XP8 | 565.2555905 | -0.784215626 | 0.364666086 | 0.000237755 | 0.004970478 |
| ARID5B | 2699.648154 | -0.784816971 | 0.189978966 | 5.02E-07 | 3.38E-05 |
| MED11 | 207.0914435 | -0.788374763 | 0.212303797 | 2.44E-06 | 0.000127437 |
| CD248 | 282.2018331 | -0.78849819 | 0.282039401 | 4.54E-05 | 0.00134991 |
| ZMPSTE24-DT | 232.9969532 | -0.791130288 | 0.195249097 | 6.68E-07 | 4.28E-05 |
| ARHGAP42 | 66.53506701 | -0.79273842 | 0.312235588 | 8.88E-05 | 0.002276809 |
| PBXIP1 | 4156.984772 | -0.794466689 | 0.190609682 | 4.11E-07 | 2.89E-05 |
| CCN4 | 20.59844633 | -0.794797 | 0.585795748 | 0.001095265 | 0.016492153 |

| | | | | | |
|---|---|---|---|---|---|
| ARHGAP18 | 844.4449837 | -0.800585142 | 0.364769493 | 0.000201107 | 0.004370569 |
| MAGEE1 | 249.8144165 | -0.803607552 | 0.155121762 | 3.57E-09 | 4.72E-07 |
| LASTR | 9.387150859 | -0.812006955 | 0.82592083 | 0.001470725 | 0.020504741 |
| MINDY2-DT | 21.02697854 | -0.813097287 | 0.492450651 | 0.000615862 | 0.010508391 |
| HSPA2 | 406.1064571 | -0.816007914 | 0.182769946 | 1.04E-07 | 9.16E-06 |
| TNS2 | 1240.403957 | -0.817755755 | 0.364750181 | 0.000168408 | 0.003785999 |
| DNAJC22 | 141.944414 | -0.82006444 | 0.321974884 | 7.79E-05 | 0.002070595 |
| SCD5 | 1506.872699 | -0.820244232 | 0.136997531 | 3.42E-11 | 6.94E-09 |
| PECAM1 | 153.1301349 | -0.821585738 | 0.824007369 | 0.001416955 | 0.01992802 |
| WNT5A-AS1 | 129.7101826 | -0.826659018 | 0.487672161 | 0.000539533 | 0.009453147 |
| LINC01291 | 139.9226881 | -0.836275175 | 0.257421064 | 9.57E-06 | 0.000375945 |
| ADORA1 | 78.72788859 | -0.837943946 | 0.283019794 | 2.30E-05 | 0.00077729 |
| AGTR1 | 304.7017296 | -0.840941166 | 0.213893114 | 8.38E-07 | 5.18E-05 |
| ADAMTS3 | 2530.637513 | -0.841195252 | 0.162564455 | 2.93E-09 | 3.97E-07 |
| DMP1 | 127.225162 | -0.84615453 | 0.792117127 | 0.001265815 | 0.018387821 |
| MATN2 | 842.2945483 | -0.847443973 | 0.313824511 | 4.65E-05 | 0.001366514 |
| LGALS8-AS1 | 5.432990672 | -0.848755915 | 0.995727552 | 0.001148598 | 0.017049373 |
| CRYZ | 1898.398709 | -0.851842791 | 0.148087084 | 1.12E-10 | 2.00E-08 |
| AGGF1P2 | 55.60843776 | -0.853026479 | 0.324493257 | 5.53E-05 | 0.001578146 |
| SEMA3F | 438.3539622 | -0.853188355 | 0.283063482 | 1.85E-05 | 0.000652083 |
| RPS23P8 | 159.3502904 | -0.855297476 | 0.259200634 | 7.49E-06 | 0.000310192 |
| FZD4 | 557.7993049 | -0.856003877 | 0.459606509 | 0.000347038 | 0.006717226 |
| DPT | 65.68920051 | -0.857737247 | 0.307515206 | 3.49E-05 | 0.001102049 |
| LRATD1 | 303.5662653 | -0.860509566 | 0.153087724 | 2.26E-10 | 3.72E-08 |
| ZFPM2-AS1 | 262.4682675 | -0.862938625 | 0.299796837 | 2.65E-05 | 0.000881505 |
| TMSB4X | 29640.42706 | -0.863705373 | 0.322962979 | 4.70E-05 | 0.00137722 |
| RGS2 | 1818.62347 | -0.870541658 | 0.17714456 | 9.28E-09 | 1.09E-06 |
| C1GALT1C1 | 596.8579148 | -0.872329744 | 0.194226575 | 6.84E-08 | 6.35E-06 |
| GAPDHP14 | 20.93106512 | -0.872740411 | 0.555981857 | 0.000590344 | 0.010190061 |
| CD93 | 8.959128632 | -0.873076067 | 0.752052937 | 0.00107854 | 0.016275514 |
| ACTB | 120863.6195 | -0.873129786 | 0.130743122 | 2.77E-13 | 8.51E-11 |
| MT1E | 62.80332039 | -0.874336893 | 0.453121902 | 0.000282977 | 0.005704612 |
| EHMT2-AS1 | 29.60090939 | -0.876677688 | 0.464402496 | 0.000307603 | 0.006066364 |
| ITGB8-AS1 | 49.65530793 | -0.877031653 | 0.351998202 | 7.30E-05 | 0.001973439 |
| TMEM150C | 597.1425231 | -0.878115915 | 0.179773066 | 1.03E-08 | 1.19E-06 |
| PLA2G4A | 79.58049267 | -0.879960555 | 0.63461072 | 0.000790028 | 0.01280041 |
| F2RL2 | 4590.918646 | -0.880845449 | 0.757401686 | 0.001049479 | 0.015956106 |
| GLIPR1 | 251.7664944 | -0.880855359 | 0.245496197 | 2.52E-06 | 0.000130411 |
| ANKRD1 | 537.3993485 | -0.883154817 | 0.760221436 | 0.001042856 | 0.015897843 |
| TMEM244 | 10.73230861 | -0.890561947 | 0.791641087 | 0.001049734 | 0.015956106 |
| PLEKHH2 | 3570.57264 | -0.892725013 | 0.301183992 | 1.85E-05 | 0.000652083 |
| KCNAB1 | 8188.576891 | -0.898266505 | 0.409650394 | 0.000143561 | 0.003342554 |
| MEGF9 | 1726.109843 | -0.899187379 | 0.159890271 | 1.75E-10 | 2.94E-08 |
| GPR88 | 33.09601996 | -0.899207598 | 0.435138842 | 0.000192544 | 0.004232919 |
| SEMA3E | 77.28233284 | -0.90134682 | 0.379797586 | 9.12E-05 | 0.002328775 |
| ZNF385D | 215.4944071 | -0.901655647 | 0.228211959 | 5.88E-07 | 3.81E-05 |
| LINC00920 | 112.202777 | -0.904135441 | 0.270599804 | 5.38E-06 | 0.000241861 |
| DKK1 | 16939.93162 | -0.905374107 | 0.19069082 | 1.74E-08 | 1.88E-06 |
| FAM13A-AS1 | 101.5980511 | -0.909935311 | 0.279608074 | 7.02E-06 | 0.000296004 |
| TM4SF4 | 16.12150872 | -0.913950629 | 0.861523316 | 0.00100684 | 0.015518715 |

| | | | | | |
|---|---|---|---|---|---|
| TSLP | 242.6858124 | -0.914635085 | 0.248800865 | 1.59E-06 | 8.94E-05 |
| OSMR | 1251.757449 | -0.920099489 | 0.399800629 | 0.000103037 | 0.002564883 |
| IKZF2 | 246.7139238 | -0.920304074 | 0.233443294 | 5.62E-07 | 3.69E-05 |
| SHC4 | 249.1330942 | -0.928708303 | 0.299908223 | 1.09E-05 | 0.000422982 |
| NREP | 2158.476173 | -0.930551555 | 0.138126606 | 1.27E-13 | 4.20E-11 |
| NT5E | 8417.538592 | -0.936303012 | 0.229290863 | 2.97E-07 | 2.17E-05 |
| MPP4 | 150.4473764 | -0.940178415 | 0.406304583 | 9.42E-05 | 0.002386336 |
| SLITRK6 | 6770.213443 | -0.940719036 | 0.248107999 | 9.33E-07 | 5.68E-05 |
| NOL12 | 149.602031 | -0.942539163 | 0.237982329 | 4.77E-07 | 3.29E-05 |
| MMP3 | 241.1713962 | -0.944138351 | 0.266452953 | 2.31E-06 | 0.00012185 |
| LMOD1 | 202.6839125 | -0.946183654 | 0.33113142 | 2.12E-05 | 0.00073154 |
| BNC2 | 588.3835214 | -0.949233185 | 0.388734173 | 6.62E-05 | 0.001827515 |
| RASSF5 | 237.8902131 | -0.950462889 | 0.408266817 | 8.83E-05 | 0.002270763 |
| PCOLCE2 | 416.1050914 | -0.953324777 | 0.20230996 | 1.64E-08 | 1.78E-06 |
| B3GNT8 | 22.06870398 | -0.958061773 | 0.499896795 | 0.000228211 | 0.004819194 |
| PCDHGC5 | 168.7617357 | -0.963499484 | 0.41877412 | 9.12E-05 | 0.002328775 |
| RMDN2 | 509.1434761 | -0.974761621 | 0.15700382 | 3.29E-12 | 8.19E-10 |
| TPBG | 6614.326232 | -0.976802548 | 0.261640623 | 1.03E-06 | 6.17E-05 |
| GREM1 | 13238.5794 | -0.982266898 | 0.414913336 | 7.34E-05 | 0.001977276 |
| ERAP2 | 824.6175683 | -0.98467013 | 0.275179445 | 1.78E-06 | 9.73E-05 |
| GRIA2 | 215.2311297 | -0.991564305 | 0.277591925 | 1.78E-06 | 9.73E-05 |
| ASB9 | 94.55239775 | -0.995122478 | 0.235459993 | 1.30E-07 | 1.07E-05 |
| DNM3OS | 4370.287846 | -1.001419427 | 0.331039625 | 1.09E-05 | 0.000424806 |
| CYP1B1 | 231.8061362 | -1.006301915 | 0.194069479 | 1.17E-09 | 1.75E-07 |
| ARHGDIB | 2622.679657 | -1.014111336 | 0.396529766 | 4.11E-05 | 0.001245338 |
| OLFML2B | 331.3094191 | -1.014350882 | 0.700913908 | 0.000491373 | 0.008704261 |
| ADAMTS12 | 136.9275034 | -1.016531347 | 0.410231288 | 5.06E-05 | 0.001469643 |
| ACTG2 | 54.02208046 | -1.017647636 | 0.534108261 | 0.000201346 | 0.004371223 |
| MT1G | 12.14383585 | -1.017760121 | 0.925287992 | 0.000712648 | 0.011803051 |
| ANGPTL2 | 1532.412307 | -1.027223232 | 0.339164276 | 1.01E-05 | 0.000395967 |
| RARB | 809.0106276 | -1.030953803 | 0.148589647 | 1.86E-14 | 7.21E-12 |
| HGF | 382.4311787 | -1.031369451 | 0.199098327 | 1.10E-09 | 1.67E-07 |
| ARHGAP29 | 808.8977721 | -1.035706527 | 0.293648545 | 1.86E-06 | 0.000100863 |
| BCAM | 35.67600435 | -1.036878037 | 0.519303499 | 0.000157179 | 0.003564264 |
| NRP1 | 9045.022997 | -1.039008774 | 0.415613756 | 4.53E-05 | 0.00134991 |
| ARHGAP5-AS1 | 129.9673144 | -1.039708285 | 0.351667 | 1.23E-05 | 0.0004649 |
| GAS1RR | 18.04887499 | -1.043768636 | 0.564624502 | 0.000214078 | 0.004562272 |
| MILR1 | 12.77125167 | -1.064609191 | 0.721415421 | 0.000413682 | 0.007679778 |
| G0S2 | 124.9042072 | -1.078353308 | 0.5636231 | 0.000172593 | 0.003855197 |
| MAB21L2 | 500.8577693 | -1.082036019 | 0.273456429 | 3.20E-07 | 2.31E-05 |
| LINC02412 | 56.35040817 | -1.087539731 | 0.539550606 | 0.000134918 | 0.003171089 |
| ANGPT2 | 64.65127335 | -1.094914401 | 0.430321517 | 3.59E-05 | 0.001121656 |
| SLC8A1 | 366.8507567 | -1.097394072 | 0.223421806 | 3.79E-09 | 4.92E-07 |
| MIRLET7BHG | 542.3676957 | -1.098351891 | 0.330584611 | 3.35E-06 | 0.00016617 |
| DUSP5P1 | 106.264304 | -1.099311606 | 0.244418877 | 2.88E-08 | 2.95E-06 |
| TCAF2 | 238.3863583 | -1.114114844 | 0.383046643 | 1.23E-05 | 0.0004649 |
| SLITRK5 | 246.6272102 | -1.115334184 | 0.240218888 | 1.40E-08 | 1.54E-06 |
| GAS1 | 330.8080886 | -1.120963828 | 0.339893743 | 3.51E-06 | 0.00017266 |
| HTR7P1 | 155.5660371 | -1.160697209 | 0.240518738 | 5.39E-09 | 6.62E-07 |
| ST8SIA4 | 59.31914507 | -1.170089917 | 0.541751203 | 8.28E-05 | 0.002160309 |

| | | | | | |
|---|---|---|---|---|---|
| CAMK1G | 164.4121311 | -1.170668102 | 0.442402032 | 2.40E-05 | 0.000804615 |
| PRRX1 | 11440.06314 | -1.183709792 | 0.444978341 | 2.26E-05 | 0.000765946 |
| STARD4-AS1 | 1967.655106 | -1.221012525 | 0.194719938 | 1.36E-12 | 3.65E-10 |
| CA12 | 147.4733558 | -1.251585091 | 0.217947165 | 3.65E-11 | 7.25E-09 |
| HAPLN1 | 5.278887999 | -1.264809142 | 1.184486829 | 0.000333694 | 0.006489014 |
| GALNT5 | 195.4246297 | -1.270373778 | 0.298869534 | 7.85E-08 | 7.19E-06 |
| LINC01050 | 12.0969178 | -1.271082622 | 0.860396405 | 0.000258401 | 0.005316945 |
| CABP7 | 35.57559022 | -1.288472442 | 0.638455004 | 9.35E-05 | 0.002375327 |
| WNT5A | 3082.964963 | -1.288631349 | 0.312675583 | 1.37E-07 | 1.12E-05 |
| CCDC80 | 2161.978369 | -1.299464586 | 0.208414161 | 1.94E-12 | 5.00E-10 |
| CALHM5 | 34.66180524 | -1.349948561 | 0.650504518 | 7.53E-05 | 0.002017731 |
| ACTA2 | 2996.548451 | -1.397885122 | 0.408968872 | 1.94E-06 | 0.000104128 |
| PAPPA | 2137.966266 | -1.399863188 | 0.317728106 | 4.37E-08 | 4.17E-06 |
| MIR503HG | 228.5039242 | -1.403406453 | 0.338398762 | 1.31E-07 | 1.08E-05 |
| PRDM1 | 196.37357 | -1.426220196 | 0.334799437 | 8.40E-08 | 7.59E-06 |
| TGFB3 | 338.1952856 | -1.479954496 | 0.328634572 | 3.14E-08 | 3.17E-06 |
| NXPH2 | 109.6950106 | -1.486571936 | 0.376532426 | 3.05E-07 | 2.22E-05 |
| PTGER3 | 821.4901072 | -1.497022202 | 0.206742612 | 3.99E-15 | 1.81E-12 |
| PLN | 72.35986552 | -1.505729138 | 0.372440033 | 2.15E-07 | 1.64E-05 |
| RUBCNL | 13.71440462 | -1.520347584 | 1.019543991 | 0.000151868 | 0.003467929 |
| IBSP | 84.22177553 | -1.521459542 | 0.382224056 | 2.76E-07 | 2.03E-05 |
| LPL | 19.11634816 | -1.597124035 | 0.55985867 | 9.34E-06 | 0.000368595 |
| REM1 | 122.5402578 | -1.598450816 | 0.313241296 | 2.33E-09 | 3.32E-07 |
| SALRNA1 | 60.20520523 | -1.602313632 | 0.560219049 | 9.13E-06 | 0.000361763 |
| DIO2 | 1606.163018 | -1.627719273 | 0.943377149 | 9.18E-05 | 0.002340826 |
| LPP-AS2 | 84.31364906 | -1.627773649 | 0.358377381 | 3.19E-08 | 3.21E-06 |
| COLEC12 | 323.9443506 | -1.670468316 | 0.452285662 | 8.02E-07 | 5.00E-05 |
| SLC5A7 | 29331.37622 | -1.670842894 | 0.315608998 | 1.01E-09 | 1.56E-07 |
| PDE1C | 49.08747546 | -1.677578372 | 0.339285377 | 5.61E-09 | 6.85E-07 |
| FZD4-DT | 17.58194645 | -1.684587314 | 0.552652762 | 5.33E-06 | 0.000240542 |
| HAS2 | 390.2272961 | -1.747226379 | 0.224757926 | 1.50E-16 | 7.84E-14 |
| GBP2 | 457.1601252 | -1.748741928 | 0.139810169 | 2.13E-37 | 7.42E-34 |
| RGS4 | 24541.5035 | -1.853704815 | 0.167501365 | 5.52E-30 | 1.05E-26 |
| SPP1 | 726.0646712 | -1.918883875 | 0.300018449 | 2.51E-12 | 6.32E-10 |
| GBP1 | 235.8692874 | -1.977948953 | 0.225389058 | 3.93E-20 | 3.56E-17 |
| ANO3 | 71.39577234 | -1.989311773 | 0.410795546 | 1.04E-08 | 1.19E-06 |
| MAF | 111.7514408 | -1.995705869 | 0.257580357 | 1.88E-16 | 9.35E-14 |
| CD109-AS1 | 16.85903439 | -2.00692335 | 0.610172223 | 2.34E-06 | 0.000122968 |
| DDIT4L | 83.97576824 | -2.027101687 | 0.261328928 | 1.69E-16 | 8.63E-14 |
| NPAS4 | 14.88401778 | -2.049409789 | 0.668278478 | 3.95E-06 | 0.000189532 |
| CARMN | 938.5955915 | -2.073096547 | 0.925977401 | 2.09E-05 | 0.000722221 |
| RNF144B | 85.72876649 | -2.074774172 | 0.334442475 | 7.62E-12 | 1.77E-09 |
| TM4SF1 | 699.5641989 | -2.130527988 | 0.261723428 | 6.86E-18 | 4.21E-15 |
| EMCN | 33.34840222 | -2.143006314 | 0.374853572 | 1.23E-10 | 2.15E-08 |
| OACYLP | 9.315039474 | -2.154550138 | 0.960285644 | 1.86E-05 | 0.000652083 |
| NDP | 22.7836487 | -2.193392361 | 0.577317008 | 4.91E-07 | 3.35E-05 |
| OLFM4 | 33.73714459 | -2.322933264 | 0.422064818 | 3.36E-10 | 5.48E-08 |
| MIR145 | 15.77605707 | -2.674759472 | 0.848455733 | 1.86E-06 | 0.000100863 |
| LRRC53 | 28.33485828 | -2.809017734 | 0.508177965 | 2.18E-10 | 3.64E-08 |
| HSD11B1 | 9.586746547 | -2.847855764 | 1.34715932 | 6.78E-06 | 0.000287033 |

**Table S2. List of DEGs in IMR-32 tumorspheres**

| gene | baseMean | log2FoldChange | lfcSE | pvalue | padj |
|---|---|---|---|---|---|
| HSPA6 | 298.3937367 | 7.607395786 | 0.729002417 | 5.64E-32 | 2.57E-29 |
| GPA33 | 10.86780014 | 4.930315115 | 1.168132244 | 7.41E-09 | 1.66E-07 |
| STC2 | 934.4911765 | 4.698717045 | 0.197722501 | 2.81E-127 | 2.89E-123 |
| NIBAN1 | 77.58828167 | 4.332878099 | 0.425374993 | 1.20E-26 | 3.11E-24 |
| KLHDC7B | 241.6194738 | 4.291168258 | 0.752437761 | 3.59E-11 | 1.29E-09 |
| CNOT6LP1 | 84.9829434 | 4.232433439 | 0.394891752 | 5.19E-29 | 1.78E-26 |
| ATF3 | 1849.241148 | 4.216919991 | 0.190102454 | 3.68E-111 | 2.52E-107 |
| GRAP | 16.26553774 | 4.101488093 | 1.177013683 | 2.19E-07 | 3.54E-06 |
| PTPRH | 36.09881004 | 4.067146573 | 0.5615208 | 2.66E-15 | 1.81E-13 |
| GDF15 | 35.11379882 | 4.054990304 | 0.531031744 | 1.46E-16 | 1.21E-14 |
| FGF7 | 16.67097379 | 4.053993594 | 0.917576249 | 1.76E-08 | 3.67E-07 |
| DMGDH | 86.95465818 | 4.007783812 | 0.374553771 | 8.65E-29 | 2.87E-26 |
| FCGR1A | 22.44690035 | 3.944593098 | 0.686375193 | 4.50E-11 | 1.59E-09 |
| LGI1 | 26.06636789 | 3.867329653 | 0.59186374 | 4.37E-13 | 2.16E-11 |
| CD300E | 18.65875559 | 3.81703546 | 0.74497348 | 1.26E-09 | 3.32E-08 |
| FUT1 | 107.5900525 | 3.784901809 | 0.597553282 | 1.70E-12 | 7.74E-11 |
| ITGA8 | 33.02754871 | 3.715679411 | 0.509249467 | 2.74E-15 | 1.85E-13 |
| SLC7A3 | 5.997349894 | 3.708794652 | 1.383044015 | 1.55E-06 | 1.97E-05 |
| NXPH4 | 289.2453236 | 3.649125741 | 0.552059739 | 3.33E-13 | 1.69E-11 |
| SLC5A12 | 16.12623217 | 3.647792107 | 0.912718429 | 1.08E-07 | 1.89E-06 |
| DDIT3 | 1136.019559 | 3.645411026 | 0.149357444 | 2.26E-133 | 4.64E-129 |
| FGF19 | 137.9987015 | 3.607626033 | 0.283256037 | 5.51E-39 | 4.05E-36 |
| RGS16 | 300.869674 | 3.58830778 | 0.45593222 | 4.01E-17 | 3.64E-15 |
| LAMP3 | 15.33362726 | 3.561412795 | 0.935076312 | 1.99E-07 | 3.25E-06 |
| FGF21 | 15.34936104 | 3.555361983 | 0.945258088 | 2.27E-07 | 3.64E-06 |
| DMBX1 | 9.853012242 | 3.511661557 | 1.503920077 | 3.20E-06 | 3.71E-05 |
| LVRN | 54.89511323 | 3.450240032 | 0.397412418 | 5.38E-20 | 6.92E-18 |
| HCLS1 | 14.6189309 | 3.413963963 | 0.795937046 | 5.17E-08 | 9.80E-07 |
| SELPLG | 47.20936797 | 3.272971649 | 0.410127254 | 2.02E-17 | 1.93E-15 |
| HID1 | 650.2648372 | 3.256542863 | 0.252102229 | 6.97E-40 | 5.31E-37 |
| ULBP1 | 559.1831746 | 3.242103463 | 0.188604729 | 6.73E-68 | 2.77E-64 |
| SLC7A11 | 2480.926755 | 3.194380404 | 0.182749214 | 4.46E-70 | 2.29E-66 |
| CHAC1 | 748.9119426 | 3.191328013 | 0.187967394 | 2.58E-66 | 7.58E-63 |
| DLX2 | 8.248912327 | 3.187569501 | 1.529015648 | 7.01E-06 | 7.39E-05 |
| FOS | 593.6535842 | 3.164618417 | 0.702631732 | 2.69E-08 | 5.41E-07 |
| SLC6A13 | 23.54325293 | 3.162221576 | 0.560940079 | 1.41E-10 | 4.48E-09 |
| H2BC15 | 198.2936942 | 3.148502647 | 0.256361657 | 2.22E-36 | 1.43E-33 |
| ZNF467 | 97.45296906 | 3.141592769 | 0.344835293 | 1.32E-21 | 1.99E-19 |
| C2orf66 | 66.18154415 | 3.092643272 | 0.331042775 | 1.54E-22 | 2.66E-20 |
| SLCO2A1 | 15.38588514 | 3.023131785 | 0.939597636 | 1.34E-06 | 1.73E-05 |
| TRIB3 | 1527.719676 | 3.015143619 | 0.232006074 | 2.53E-40 | 2.00E-37 |
| PDLIM3 | 231.2907889 | 2.973329871 | 0.456929572 | 8.09E-13 | 3.83E-11 |
| TEX14 | 125.7279509 | 2.938279715 | 0.288436291 | 3.73E-26 | 9.36E-24 |
| MTDHP4 | 10.31756266 | 2.839351625 | 1.038481368 | 4.36E-06 | 4.86E-05 |
| ACHE | 52.05323329 | 2.834434742 | 0.387585226 | 3.01E-15 | 2.02E-13 |
| TRIM66 | 2883.339311 | 2.812514262 | 0.180243176 | 1.24E-56 | 2.13E-53 |
| TRIM55 | 20.5486683 | 2.792952887 | 0.560756806 | 3.69E-09 | 8.87E-08 |
| CEBPA | 109.176058 | 2.77925394 | 0.273772751 | 4.70E-26 | 1.16E-23 |

| | | | | | |
|---|---|---|---|---|---|
| BHLHA15 | 173.3242076 | 2.771552075 | 0.329388788 | 4.96E-19 | 5.66E-17 |
| ADRA1B | 11.55999417 | 2.762805076 | 0.821817438 | 1.11E-06 | 1.46E-05 |
| TBX21 | 19.53317669 | 2.750589688 | 0.561581198 | 5.33E-09 | 1.23E-07 |
| LINC01686 | 11.27072247 | 2.742857746 | 1.153455158 | 9.29E-06 | 9.45E-05 |
| SESN2 | 1885.698721 | 2.736624837 | 0.201902237 | 1.18E-43 | 1.11E-40 |
| ARID5A | 446.7374707 | 2.68532294 | 0.246385945 | 1.57E-29 | 5.68E-27 |
| HAVCR1 | 6.322827002 | 2.685057009 | 1.531737836 | 2.17E-05 | 0.000200334 |
| H4C15 | 23.19283621 | 2.679817508 | 0.532323723 | 2.79E-09 | 6.85E-08 |
| ARHGAP9 | 243.3613704 | 2.65518751 | 0.269151401 | 7.29E-25 | 1.61E-22 |
| ALPK2 | 54.71390894 | 2.63870843 | 0.422791262 | 3.72E-12 | 1.60E-10 |
| NTS | 47.44124602 | 2.638025185 | 0.398711743 | 3.36E-13 | 1.70E-11 |
| ULBP2 | 175.7659036 | 2.637579373 | 0.220352412 | 6.74E-35 | 3.96E-32 |
| PNLDC1 | 45.72431977 | 2.63384463 | 0.364041969 | 4.61E-15 | 3.05E-13 |
| BCL6 | 324.4212346 | 2.622016966 | 0.25222826 | 3.17E-27 | 8.92E-25 |
| INSC | 35.37206883 | 2.5783826 | 0.408196875 | 2.24E-12 | 1.01E-10 |
| TIGAR | 44.38923141 | 2.577560993 | 0.414442985 | 4.11E-12 | 1.76E-10 |
| HAS1 | 13.8776986 | 2.56859733 | 0.726626177 | 7.38E-07 | 1.01E-05 |
| KNDC1 | 295.6413931 | 2.55365793 | 0.268889205 | 2.32E-23 | 4.34E-21 |
| SLC6A12 | 20.73036162 | 2.540254954 | 0.620471761 | 1.26E-07 | 2.17E-06 |
| WWTR1 | 97.43571 | 2.538564868 | 0.277346453 | 5.76E-22 | 9.41E-20 |
| PCK2 | 1683.560331 | 2.535532234 | 0.171851922 | 3.48E-51 | 4.77E-48 |
| ADRA1A | 62.347778 | 2.529019259 | 0.373835453 | 1.16E-13 | 6.23E-12 |
| MGAM2 | 347.3900716 | 2.514688737 | 0.240951221 | 1.80E-27 | 5.20E-25 |
| RPL23AP15 | 10.54661036 | 2.493658619 | 0.897166907 | 5.23E-06 | 5.71E-05 |
| DDIT4 | 4573.410121 | 2.482270173 | 0.145770253 | 5.69E-67 | 1.95E-63 |
| HERPUD1 | 5174.958131 | 2.479795329 | 0.175639074 | 3.18E-47 | 3.85E-44 |
| DNAJB5-DT | 11.75759343 | 2.478241336 | 0.861937844 | 4.24E-06 | 4.73E-05 |
| HSD17B13 | 39.10545625 | 2.45973294 | 0.350207529 | 1.85E-14 | 1.12E-12 |
| MSTN | 46.98388645 | 2.455634263 | 0.355531113 | 4.17E-14 | 2.40E-12 |
| ALB | 56.85859408 | 2.448624021 | 0.316211296 | 8.64E-17 | 7.43E-15 |
| LINC00664 | 16.96112695 | 2.432614869 | 0.67419443 | 6.07E-07 | 8.60E-06 |
| ATF5 | 3843.423496 | 2.416946831 | 0.235642746 | 1.04E-26 | 2.75E-24 |
| H2BC18 | 44.76780447 | 2.414145786 | 0.387945556 | 3.68E-12 | 1.59E-10 |
| AKNA | 625.7935837 | 2.404257839 | 0.388080769 | 4.35E-12 | 1.85E-10 |
| SYT5 | 165.3748051 | 2.402199542 | 0.278340046 | 5.48E-20 | 7.01E-18 |
| TSPYL2 | 2312.414978 | 2.394459223 | 0.173405183 | 2.19E-45 | 2.50E-42 |
| APOL6 | 10.20336841 | 2.364157138 | 1.098835517 | 2.13E-05 | 0.000197448 |
| CASS4 | 16.16172724 | 2.359994528 | 0.603360944 | 2.30E-07 | 3.70E-06 |
| CORIN | 162.5943454 | 2.349066559 | 0.231756455 | 3.40E-26 | 8.64E-24 |
| HSPA8P13 | 74.02138969 | 2.346484017 | 0.309754756 | 2.99E-16 | 2.32E-14 |
| RANP4 | 64.01831952 | 2.332943244 | 0.324501764 | 5.28E-15 | 3.47E-13 |
| NR4A3 | 79.33767047 | 2.323963845 | 0.356256356 | 5.22E-13 | 2.56E-11 |
| LINC01068 | 10.94501275 | 2.321466768 | 1.007340627 | 1.67E-05 | 0.000159596 |
| DRGX | 997.452484 | 2.314157837 | 0.221866949 | 1.57E-27 | 4.67E-25 |
| TNFSF13B | 52.58926011 | 2.312754734 | 0.355323576 | 5.72E-13 | 2.80E-11 |
| GCM1 | 22.65641029 | 2.297202624 | 0.508592189 | 2.38E-08 | 4.84E-07 |
| LINC02881 | 9.519621056 | 2.2692308 | 1.1997134 | 3.71E-05 | 0.00031698 |
| H4C8 | 119.2197895 | 2.24379125 | 0.278138691 | 5.94E-18 | 6.10E-16 |
| H2AC6 | 353.982899 | 2.220563661 | 0.190177851 | 1.42E-33 | 7.67E-31 |
| HSPD1P10 | 10.93469375 | 2.220224154 | 0.819313423 | 7.33E-06 | 7.67E-05 |

| | | | | | |
|---|---|---|---|---|---|
| PNMA8C | 201.5700809 | 2.218231152 | 0.22063612 | 7.43E-26 | 1.76E-23 |
| ERO1B | 1424.63896 | 2.214472928 | 0.140282974 | 3.25E-58 | 6.08E-55 |
| MIR616 | 29.0579832 | 2.208933784 | 0.551374283 | 1.62E-07 | 2.72E-06 |
| PPP1R15A | 1670.235115 | 2.207504338 | 0.223266906 | 3.97E-25 | 8.98E-23 |
| MYOCD | 27.28322976 | 2.204372923 | 0.492831107 | 2.77E-08 | 5.55E-07 |
| CD200R1 | 9.978495284 | 2.196878439 | 1.001549472 | 2.36E-05 | 0.000215664 |
| CFAP43 | 57.19014514 | 2.187208991 | 0.363105686 | 1.16E-11 | 4.58E-10 |
| ZBTB7B | 552.6632116 | 2.17104904 | 0.219345448 | 3.59E-25 | 8.21E-23 |
| TSC22D3 | 2100.795055 | 2.160616366 | 0.131043523 | 3.73E-63 | 8.53E-60 |
| WNT4 | 17.90663265 | 2.158552706 | 0.652596297 | 1.52E-06 | 1.94E-05 |
| KIAA1671 | 504.595425 | 2.142751104 | 0.236586622 | 1.14E-21 | 1.74E-19 |
| OPTN | 493.1269203 | 2.13522097 | 0.183663098 | 2.60E-33 | 1.34E-30 |
| BEST1 | 268.1214615 | 2.134895214 | 0.216293765 | 4.79E-25 | 1.07E-22 |
| MOB3B | 156.8581806 | 2.123779717 | 0.248757281 | 1.17E-19 | 1.45E-17 |
| MERTK | 55.16543176 | 2.106679579 | 0.407606987 | 1.19E-09 | 3.13E-08 |
| CBX4 | 4596.382283 | 2.08398297 | 0.186355097 | 4.38E-31 | 1.77E-28 |
| AJUBA | 158.2263485 | 2.077975931 | 0.273666301 | 2.63E-16 | 2.07E-14 |
| TCP11L2 | 180.2505162 | 2.073154341 | 0.207997474 | 1.90E-25 | 4.38E-23 |
| TNFRSF12A | 129.1429448 | 2.062317497 | 0.372817191 | 1.83E-10 | 5.67E-09 |
| GLI1 | 45.22498661 | 2.050829349 | 0.402684743 | 1.68E-09 | 4.29E-08 |
| OVGP1 | 244.7915293 | 2.046414442 | 0.180841951 | 1.00E-31 | 4.30E-29 |
| CTRB2 | 5.224129997 | 2.044802932 | 1.534498675 | 8.64E-05 | 0.000650872 |
| MIR7-3HG | 49.34648332 | 2.034441525 | 0.439716508 | 1.38E-08 | 2.95E-07 |
| ALOXE3 | 7.137483952 | 2.031181873 | 1.454946164 | 9.03E-05 | 0.000676234 |
| TEKT3 | 51.86867038 | 2.030546262 | 0.406514134 | 2.66E-09 | 6.57E-08 |
| DNAI4 | 206.169484 | 2.026380425 | 0.209952806 | 4.53E-24 | 9.04E-22 |
| TPTE2 | 14.43099908 | 2.020444252 | 0.761528858 | 9.52E-06 | 9.67E-05 |
| CCT6B | 45.1518235 | 2.019118906 | 0.346533776 | 3.58E-11 | 1.29E-09 |
| ZNF442 | 387.2997901 | 2.018076756 | 0.162343733 | 1.69E-37 | 1.16E-34 |
| MIOX | 39.64887179 | 2.015760783 | 0.406754851 | 3.18E-09 | 7.68E-08 |
| HSPA1A | 213.1040082 | 2.013813521 | 0.745563561 | 8.45E-06 | 8.69E-05 |
| BHLHE41 | 12.19107468 | 2.000606844 | 0.895492062 | 2.64E-05 | 0.000236977 |
| NUTM2F | 24.46176867 | 1.998154961 | 0.479481336 | 8.58E-08 | 1.55E-06 |
| TG | 34.52274798 | 1.997021597 | 0.388291711 | 1.30E-09 | 3.39E-08 |
| PHF11 | 23.02394133 | 1.990509848 | 0.610282334 | 1.79E-06 | 2.23E-05 |
| LINC00877 | 81.94978749 | 1.989525293 | 0.254631047 | 5.05E-17 | 4.49E-15 |
| LGALS9 | 98.69906085 | 1.987160088 | 0.317798659 | 2.88E-12 | 1.28E-10 |
| ASNS | 670.0561518 | 1.984347979 | 0.148058501 | 5.85E-43 | 5.01E-40 |
| CACNA1G | 43.72764179 | 1.984310988 | 0.393827114 | 2.13E-09 | 5.33E-08 |
| ANKRD34C-AS1 | 9.610216846 | 1.979599564 | 1.038055524 | 5.33E-05 | 0.000430068 |
| PARD3B | 202.8176358 | 1.970992731 | 0.245499565 | 9.23E-18 | 9.31E-16 |
| CCDC30 | 206.1703156 | 1.956607335 | 0.202012244 | 3.55E-24 | 7.16E-22 |
| BHLHE40 | 121.1981159 | 1.940117868 | 0.401273551 | 5.35E-09 | 1.23E-07 |
| HFE | 7.44874344 | 1.938583742 | 1.30182754 | 0.00010446 | 0.000768024 |
| PTGFR | 766.6470542 | 1.933854726 | 0.167943397 | 1.21E-32 | 5.78E-30 |
| SPAG1 | 188.436224 | 1.919170591 | 0.238930841 | 9.39E-18 | 9.39E-16 |
| AMDHD1 | 52.1224964 | 1.917260535 | 0.374895045 | 1.45E-09 | 3.74E-08 |
| ZNF425 | 317.513297 | 1.916753083 | 0.202197415 | 2.75E-23 | 5.04E-21 |
| TSHZ3-AS1 | 13.06249601 | 1.908880209 | 0.723804882 | 1.06E-05 | 0.00010686 |
| ZNF654 | 2213.174177 | 1.907010257 | 0.127877786 | 3.17E-52 | 4.66E-49 |

| | | | | | |
|---|---|---|---|---|---|
| CFAP107 | 3.566147982 | 1.905207339 | 1.579411015 | 0.00010992 | 0.000803524 |
| NBPF13P | 4.042912471 | 1.897481867 | 1.601468363 | 0.00010864 | 0.000795623 |
| GABRR2 | 208.857459 | 1.892929375 | 0.214279976 | 1.09E-20 | 1.51E-18 |
| INHBE | 315.6798701 | 1.884474804 | 0.180784494 | 2.25E-27 | 6.43E-25 |
| ERN1 | 2183.327957 | 1.882001625 | 0.193329897 | 2.45E-24 | 5.05E-22 |
| STMN4 | 1249.386077 | 1.865594845 | 0.135553183 | 5.43E-45 | 5.88E-42 |
| EFEMP1 | 11.4556075 | 1.86366315 | 0.828393467 | 2.91E-05 | 0.000257904 |
| MXD1 | 2056.259315 | 1.858221523 | 0.137545182 | 1.77E-43 | 1.58E-40 |
| DNAJB9 | 1799.381623 | 1.85089091 | 0.115040496 | 4.06E-60 | 8.35E-57 |
| NUCB2 | 1770.933907 | 1.849360382 | 0.123627403 | 1.81E-52 | 2.86E-49 |
| IQCN | 780.6471921 | 1.842871477 | 0.220242361 | 6.40E-19 | 7.12E-17 |
| LINC01160 | 23.5567882 | 1.84042783 | 0.500351713 | 4.76E-07 | 6.99E-06 |
| KLF9 | 698.5114816 | 1.820851484 | 0.172347051 | 5.64E-28 | 1.76E-25 |
| NFATC2 | 14.32661648 | 1.816951449 | 0.729746476 | 1.67E-05 | 0.000159822 |
| GKN3P | 47.2405422 | 1.811166691 | 0.366200619 | 3.00E-09 | 7.32E-08 |
| SDCBP2 | 64.67339381 | 1.807575086 | 0.289924468 | 3.15E-12 | 1.38E-10 |
| SOX30 | 8.449697649 | 1.807475659 | 0.990343064 | 7.86E-05 | 0.0006008 |
| MYRFL | 9.447263725 | 1.806787379 | 0.947646196 | 6.68E-05 | 0.000523244 |
| SPATA17 | 84.88433293 | 1.797032253 | 0.287491334 | 2.85E-12 | 1.26E-10 |
| MAP3K7CL | 60.48939678 | 1.793467511 | 0.31035433 | 4.36E-11 | 1.55E-09 |
| LYZ | 13.59025606 | 1.779441656 | 0.754813474 | 2.42E-05 | 0.000220469 |
| BBC3 | 602.0350044 | 1.768255334 | 0.193677056 | 8.59E-22 | 1.34E-19 |
| MYC | 47.31165026 | 1.767265889 | 0.364227689 | 4.45E-09 | 1.05E-07 |
| H2BC6 | 61.6725422 | 1.765267561 | 0.299414317 | 2.22E-11 | 8.34E-10 |
| SERTAD1 | 149.6153159 | 1.760890087 | 0.209839106 | 5.49E-19 | 6.24E-17 |
| SLC6A9 | 447.7216726 | 1.7574296 | 0.277719665 | 1.73E-12 | 7.88E-11 |
| PIPOX | 54.24435405 | 1.756935638 | 0.340997354 | 1.08E-09 | 2.89E-08 |
| XBP1 | 5446.493562 | 1.749149147 | 0.104427255 | 9.67E-65 | 2.49E-61 |
| TNFSF4 | 95.91116475 | 1.747661378 | 0.238758158 | 2.31E-15 | 1.59E-13 |
| PGPEP1 | 422.1263928 | 1.746548188 | 0.205591282 | 2.30E-19 | 2.75E-17 |
| PLA2G4C | 35.15892979 | 1.746524913 | 0.427356679 | 1.06E-07 | 1.86E-06 |
| GPAT3 | 9.690680507 | 1.74401867 | 0.885653704 | 6.35E-05 | 0.000500119 |
| CREB3L2 | 2491.524305 | 1.739874362 | 0.173556538 | 1.65E-25 | 3.85E-23 |
| KRTAP5-AS1 | 61.38543449 | 1.739788567 | 0.356202272 | 3.75E-09 | 8.97E-08 |
| NEDD4 | 1297.034521 | 1.739238926 | 0.130934032 | 4.72E-42 | 3.89E-39 |
| IRAK2 | 45.63805941 | 1.739197473 | 0.351396818 | 2.78E-09 | 6.85E-08 |
| FAM133A | 116.696056 | 1.733343937 | 0.267030231 | 6.16E-13 | 2.98E-11 |
| CTH | 1259.903845 | 1.729575096 | 0.11760985 | 1.01E-50 | 1.30E-47 |
| ADM2 | 1080.221984 | 1.725147254 | 0.241217387 | 7.53E-15 | 4.81E-13 |
| ITGA11 | 169.4099666 | 1.724650201 | 0.323154898 | 4.20E-10 | 1.22E-08 |
| SGK3 | 21.54060167 | 1.713251293 | 0.627308103 | 9.34E-06 | 9.50E-05 |
| DNAH8 | 4.973857529 | 1.707553412 | 1.482660055 | 0.00017678 | 0.001212178 |
| CREB3L1 | 10.64329558 | 1.703679018 | 1.160726365 | 0.00016687 | 0.001154966 |
| SLC1A4 | 3056.114831 | 1.70184172 | 0.186721135 | 1.01E-21 | 1.55E-19 |
| C10orf71-AS1 | 16.19925189 | 1.697199369 | 0.68002782 | 1.82E-05 | 0.00017219 |
| TMEM236 | 24.67831231 | 1.694216193 | 0.47925795 | 7.62E-07 | 1.04E-05 |
| STAC2 | 590.9279778 | 1.691989346 | 0.253429887 | 1.80E-13 | 9.40E-12 |
| VEGFA | 3752.646682 | 1.691552326 | 0.163853412 | 8.02E-27 | 2.14E-24 |
| ATOSA | 1823.362527 | 1.680826001 | 0.144008372 | 2.87E-33 | 1.44E-30 |
| H3C6 | 105.3501695 | 1.678375491 | 0.242690359 | 3.63E-14 | 2.12E-12 |

| | | | | | |
|---|---|---|---|---|---|
| ZNF433 | 320.4690047 | 1.665712575 | 0.139865394 | 1.74E-34 | 9.65E-32 |
| CEBPB | 894.9823047 | 1.657315932 | 0.226938888 | 2.38E-15 | 1.63E-13 |
| CPEB3 | 428.5314116 | 1.653764145 | 0.157913183 | 1.69E-27 | 4.96E-25 |
| SRGAP3 | 287.1253411 | 1.65083663 | 0.274516287 | 9.76E-12 | 3.90E-10 |
| ALDH8A1 | 319.0301565 | 1.648952243 | 0.169768505 | 3.55E-24 | 7.16E-22 |
| CARS1-AS1 | 21.01762497 | 1.647305009 | 0.601689553 | 9.58E-06 | 9.72E-05 |
| NEK11 | 74.77506235 | 1.645443234 | 0.295023979 | 1.08E-10 | 3.54E-09 |
| L3MBTL2-AS1 | 48.1374911 | 1.643169165 | 0.442284016 | 4.01E-07 | 6.00E-06 |
| FBXO16 | 281.5900239 | 1.643119709 | 0.173981072 | 4.59E-23 | 8.21E-21 |
| H2AC19 | 8.784555731 | 1.641110398 | 0.94173724 | 0.00011845 | 0.000856446 |
| H2BC11 | 51.75125219 | 1.637947488 | 0.30540781 | 3.26E-10 | 9.72E-09 |
| HSPA1B | 1581.884096 | 1.62518085 | 0.225883768 | 4.83E-15 | 3.19E-13 |
| SMIM14 | 1535.134644 | 1.618364764 | 0.167533678 | 5.70E-24 | 1.13E-21 |
| BDKRB2 | 146.0327971 | 1.607500457 | 0.745116205 | 4.83E-05 | 0.000397815 |
| TBX20 | 275.7549258 | 1.604988986 | 0.167034059 | 9.08E-24 | 1.73E-21 |
| CREBRF | 528.7485074 | 1.604531233 | 0.149460115 | 9.85E-29 | 3.22E-26 |
| APLF | 176.6626286 | 1.600565304 | 0.183052046 | 2.39E-20 | 3.16E-18 |
| P4HA2 | 414.5017706 | 1.599561494 | 0.175284273 | 8.09E-22 | 1.28E-19 |
| ID4 | 1444.007279 | 1.598675807 | 0.146825709 | 1.87E-29 | 6.64E-27 |
| FBXO34-AS1 | 9.109858875 | 1.597223259 | 0.883750643 | 0.00011109 | 0.000810627 |
| ADAMTSL3 | 65.24463202 | 1.592748165 | 0.30841971 | 8.13E-10 | 2.22E-08 |
| ELAPOR1 | 293.8315141 | 1.586987137 | 0.175594049 | 1.73E-21 | 2.56E-19 |
| DUSP6 | 412.1390893 | 1.581571834 | 0.251641464 | 1.68E-12 | 7.68E-11 |
| LARP6 | 1515.112185 | 1.572169981 | 0.146642431 | 1.07E-28 | 3.44E-26 |
| DNAJC3 | 3009.982668 | 1.568130712 | 0.115353405 | 7.15E-44 | 7.00E-41 |
| STX5 | 1272.983121 | 1.568079625 | 0.137530769 | 5.76E-32 | 2.57E-29 |
| ZFAND2A | 687.7082319 | 1.567860375 | 0.14303201 | 7.85E-30 | 3.04E-27 |
| POU5F1P3 | 44.80595999 | 1.564338608 | 0.39117277 | 1.33E-07 | 2.27E-06 |
| IRF1 | 131.912721 | 1.562831395 | 0.275937884 | 5.71E-11 | 1.98E-09 |
| ZC3H12D | 22.47703205 | 1.5624478 | 0.571757323 | 1.04E-05 | 0.000104419 |
| FAS | 445.9883021 | 1.562254965 | 0.198067811 | 2.43E-17 | 2.28E-15 |
| TRPS1 | 688.4740651 | 1.561546696 | 0.18481042 | 2.62E-19 | 3.10E-17 |
| TINCR | 52.60800849 | 1.55608531 | 0.3294124 | 6.13E-09 | 1.40E-07 |
| S100A1 | 50.88202525 | 1.550329633 | 0.297437022 | 5.87E-10 | 1.65E-08 |
| SLC3A2 | 9559.489144 | 1.546112629 | 0.165004085 | 7.32E-23 | 1.30E-20 |
| LYSMD3 | 1322.548694 | 1.538973909 | 0.136688375 | 2.64E-31 | 1.09E-28 |
| NHS | 615.6179064 | 1.538456273 | 0.175613418 | 1.71E-20 | 2.32E-18 |
| ZMYND12 | 22.58181894 | 1.53684442 | 0.547993278 | 8.50E-06 | 8.73E-05 |
| NKX6-2 | 6.265259182 | 1.5340986 | 1.368223781 | 0.00027908 | 0.001776588 |
| MIR22HG | 90.83635769 | 1.529757622 | 0.266590195 | 3.47E-11 | 1.26E-09 |
| CLIC2 | 140.6505895 | 1.527509141 | 0.204594254 | 5.14E-16 | 3.87E-14 |
| RELB | 57.54363795 | 1.527040656 | 0.375194421 | 9.71E-08 | 1.73E-06 |
| ARC | 1232.3501 | 1.524077786 | 0.56385388 | 1.17E-05 | 0.000116485 |
| MCHR1 | 178.0230694 | 1.522520185 | 0.285928247 | 3.10E-10 | 9.28E-09 |
| LINC00910 | 768.5053304 | 1.520459458 | 0.167251047 | 8.54E-22 | 1.34E-19 |
| PLEK | 8.048920643 | 1.518190262 | 1.002302565 | 0.00022247 | 0.001468199 |
| CSTA | 3.281528395 | 1.516120604 | 1.469002925 | 0.00025503 | 0.001650443 |
| GBP2 | 11.377196 | 1.516111414 | 0.73862325 | 7.11E-05 | 0.00055112 |
| KLF10 | 2899.911089 | 1.506482327 | 0.194573326 | 5.97E-17 | 5.25E-15 |
| LINC02995 | 42.03264686 | 1.500513019 | 0.34809164 | 3.49E-08 | 6.85E-07 |

| | | | | | |
|---|---|---|---|---|---|
| SV2B | 89.60672773 | 1.496273552 | 0.482948444 | 3.41E-06 | 3.92E-05 |
| ADAMTS6 | 42.44680406 | 1.49292752 | 0.511968018 | 6.18E-06 | 6.60E-05 |
| SYTL2 | 103.2964541 | 1.492530136 | 0.278378549 | 2.37E-10 | 7.26E-09 |
| PCLO | 1182.438314 | 1.492147813 | 0.152431086 | 1.07E-24 | 2.32E-22 |
| MIR325HG | 30.15920168 | 1.490753007 | 0.444494898 | 1.42E-06 | 1.83E-05 |
| HPX | 131.0298264 | 1.490065763 | 0.262408896 | 4.29E-11 | 1.53E-09 |
| NIM1K | 47.74485358 | 1.488055205 | 0.379309727 | 1.69E-07 | 2.82E-06 |
| LINC02940 | 63.21963434 | 1.486063031 | 0.398227006 | 3.56E-07 | 5.39E-06 |
| GLRA4 | 16.33108347 | 1.482643583 | 0.561660931 | 1.47E-05 | 0.00014279 |
| C1S | 89.6125294 | 1.4826344 | 0.268093845 | 9.41E-11 | 3.10E-09 |
| BAIAP3 | 119.2540792 | 1.482561016 | 0.358333495 | 7.08E-08 | 1.30E-06 |
| LDLRAD4 | 2162.569755 | 1.480462455 | 0.18748875 | 1.61E-17 | 1.56E-15 |
| DENND3 | 1045.236197 | 1.47854248 | 0.165791614 | 3.26E-21 | 4.75E-19 |
| CHRNA9 | 12.22631926 | 1.470287819 | 0.817892663 | 0.00013979 | 0.000993275 |
| CDC42EP1 | 3.964786374 | 1.468761657 | 1.516297904 | 0.00024871 | 0.00161442 |
| TBX19 | 323.4718185 | 1.466748841 | 0.173279304 | 1.52E-19 | 1.83E-17 |
| EPHA4 | 180.3032569 | 1.466527109 | 0.193740833 | 1.82E-16 | 1.48E-14 |
| CARS1 | 4268.86982 | 1.462456605 | 0.122228415 | 5.05E-35 | 3.05E-32 |
| AMY2A | 4.809129081 | 1.462288751 | 1.417585955 | 0.00030407 | 0.001912215 |
| CFAP221 | 27.59740018 | 1.460299415 | 0.454043822 | 2.32E-06 | 2.78E-05 |
| LYST | 1173.118684 | 1.454920836 | 0.132725448 | 4.67E-30 | 1.85E-27 |
| ODAD3 | 27.93865452 | 1.454887726 | 0.460916103 | 2.85E-06 | 3.36E-05 |
| AOC2 | 185.9703412 | 1.453569742 | 0.208927364 | 1.36E-14 | 8.32E-13 |
| SERPINB6 | 309.8709261 | 1.452012713 | 0.166085525 | 1.30E-20 | 1.80E-18 |
| ARID4A | 1937.090273 | 1.445684961 | 0.120275191 | 2.34E-35 | 1.46E-32 |
| ACAD11 | 131.6978028 | 1.445591998 | 0.21813617 | 1.19E-13 | 6.38E-12 |
| LPXN | 96.90187636 | 1.44478063 | 0.245425979 | 1.13E-11 | 4.49E-10 |
| BAG3 | 563.7859259 | 1.443854141 | 0.489299267 | 5.72E-06 | 6.18E-05 |
| TRPM6 | 12.22764684 | 1.441060039 | 0.831531801 | 0.00016961 | 0.001170418 |
| SEC14L3 | 7.733255999 | 1.4337584 | 1.177875473 | 0.00037206 | 0.002279151 |
| ELFN1 | 417.4720005 | 1.431308771 | 0.631321806 | 4.59E-05 | 0.000380675 |
| CFAP53 | 29.623943 | 1.4304899 | 0.434643673 | 1.80E-06 | 2.23E-05 |
| PAH | 17.9004124 | 1.41667085 | 0.595595059 | 3.43E-05 | 0.000297017 |
| RASGRP3 | 37.74090824 | 1.416168059 | 0.55895351 | 2.18E-05 | 0.000200816 |
| CFAP69 | 15.76921763 | 1.413990156 | 0.587842126 | 3.19E-05 | 0.000278362 |
| IL21R | 67.9419017 | 1.413542676 | 0.375057179 | 2.99E-07 | 4.64E-06 |
| PLEKHG1 | 132.3605243 | 1.413285236 | 0.227706325 | 1.48E-12 | 6.80E-11 |
| NDUFA4L2 | 164.4516615 | 1.413270088 | 0.291220233 | 2.52E-09 | 6.26E-08 |
| GPRC5A | 73.82285602 | 1.409839576 | 0.322080765 | 2.31E-08 | 4.70E-07 |
| CMYA5 | 50.58292581 | 1.409595166 | 0.315429333 | 1.53E-08 | 3.23E-07 |
| HSPA1L | 20.36609753 | 1.405787233 | 0.633148333 | 5.42E-05 | 0.000436589 |
| SCN11A | 210.29358 | 1.40007054 | 0.23799959 | 9.75E-12 | 3.90E-10 |
| SERPINI1 | 31.45154999 | 1.399375947 | 0.431557372 | 2.16E-06 | 2.62E-05 |
| NCEH1 | 363.4503374 | 1.399015063 | 0.158760403 | 5.01E-21 | 7.21E-19 |
| CLK4 | 1464.006704 | 1.393426211 | 0.179387206 | 2.66E-17 | 2.48E-15 |
| LINC03080 | 11.8758747 | 1.392857496 | 0.757698452 | 0.00014652 | 0.00103364 |
| ALDH1L2 | 5290.010568 | 1.388920921 | 0.168359687 | 5.54E-19 | 6.26E-17 |
| XDH | 43.22256616 | 1.385011273 | 0.368819042 | 3.13E-07 | 4.83E-06 |
| CCPG1 | 379.7026627 | 1.37651637 | 0.137484253 | 5.41E-26 | 1.31E-23 |
| FAM131B-AS2 | 72.56798277 | 1.373311747 | 0.319227988 | 3.08E-08 | 6.11E-07 |

| | | | | | |
|---|---|---|---|---|---|
| PCSK1 | 13.41566897 | 1.373082644 | 0.804520186 | 0.00020444 | 0.001365387 |
| SAMD4A | 551.5203308 | 1.370905861 | 0.228206959 | 4.14E-12 | 1.77E-10 |
| KCND1 | 175.4713171 | 1.37002503 | 0.234978008 | 1.18E-11 | 4.66E-10 |
| ODAD4 | 215.5838975 | 1.369359513 | 0.168006045 | 1.09E-18 | 1.19E-16 |
| WHAMM | 713.9709349 | 1.369181714 | 0.200940836 | 2.34E-14 | 1.39E-12 |
| SKIL | 4463.495191 | 1.364665211 | 0.10962489 | 6.35E-38 | 4.50E-35 |
| ARID5B | 180.1806782 | 1.36379501 | 0.168334109 | 1.56E-18 | 1.68E-16 |
| RAB17 | 10.6129428 | 1.363496459 | 0.845926323 | 0.0002537 | 0.001643159 |
| BMAL1 | 433.5729779 | 1.363217691 | 0.136713804 | 7.28E-26 | 1.74E-23 |
| THAP10 | 806.0456295 | 1.362738542 | 0.15293975 | 1.61E-21 | 2.40E-19 |
| DNASE2 | 415.3102336 | 1.361938413 | 0.184969899 | 4.61E-16 | 3.49E-14 |
| ZBTB16 | 72.21953171 | 1.361018572 | 0.270618532 | 9.25E-10 | 2.50E-08 |
| TMEM87B | 494.290064 | 1.359813163 | 0.15256429 | 1.54E-21 | 2.31E-19 |
| DUSP10 | 722.7944613 | 1.357271177 | 0.204501486 | 7.22E-14 | 4.01E-12 |
| RHBDF1 | 215.9186699 | 1.350428441 | 0.248507833 | 1.04E-10 | 3.42E-09 |
| LINC01518 | 302.761123 | 1.344982367 | 0.183499216 | 5.32E-16 | 3.96E-14 |
| PLCD1 | 223.6897204 | 1.343285182 | 0.214572917 | 7.73E-13 | 3.67E-11 |
| FICD | 308.0838012 | 1.342627515 | 0.166651664 | 1.94E-18 | 2.07E-16 |
| DEPP1 | 24.96299124 | 1.340547429 | 0.515403332 | 1.96E-05 | 0.00018404 |
| GPR139 | 68.09588862 | 1.339346388 | 0.34132633 | 1.54E-07 | 2.59E-06 |
| EFCAB6 | 40.85084295 | 1.337789841 | 0.3893796 | 1.09E-06 | 1.44E-05 |
| LINC02934 | 10.00267105 | 1.33756598 | 0.836392265 | 0.00027477 | 0.001754753 |
| JAK3 | 52.86301218 | 1.336315483 | 0.392825056 | 1.24E-06 | 1.62E-05 |
| MRC2 | 1834.854111 | 1.336113748 | 0.22634541 | 6.72E-12 | 2.77E-10 |
| H3P4 | 7.587336225 | 1.335537233 | 0.964064257 | 0.00039937 | 0.002418327 |
| PLAC9 | 8.190055779 | 1.332166059 | 0.935775474 | 0.00038018 | 0.002319907 |
| BAALC-AS1 | 68.68868965 | 1.332102221 | 0.265977369 | 9.61E-10 | 2.59E-08 |
| SH2D3C | 1089.137335 | 1.331784747 | 0.203630687 | 1.21E-13 | 6.44E-12 |
| PAK6 | 122.1125584 | 1.330108713 | 0.2592774 | 5.07E-10 | 1.44E-08 |
| CCDC13-AS1 | 9.21464759 | 1.328292482 | 0.826822162 | 0.00027627 | 0.001762163 |
| ELL2 | 264.7533254 | 1.326884616 | 0.180461464 | 4.02E-16 | 3.05E-14 |
| DLGAP1-AS2 | 28.17331463 | 1.323770381 | 0.445060435 | 5.87E-06 | 6.31E-05 |
| CHIC2 | 770.3336418 | 1.322104754 | 0.141651617 | 2.45E-23 | 4.55E-21 |
| MAFK | 193.0305931 | 1.312825303 | 0.350057914 | 3.15E-07 | 4.86E-06 |
| FAIM2 | 1099.08499 | 1.31058762 | 0.265023328 | 1.27E-09 | 3.34E-08 |
| KAT2B | 708.6381259 | 1.307700924 | 0.122300134 | 2.56E-29 | 8.93E-27 |
| CATSPERE | 70.9258605 | 1.306484346 | 0.280572535 | 5.36E-09 | 1.24E-07 |
| CCDC81 | 90.36024796 | 1.303332801 | 0.252864279 | 4.18E-10 | 1.22E-08 |
| TOB1 | 2403.482044 | 1.303195541 | 0.135177126 | 1.13E-24 | 2.42E-22 |
| BTK | 17.92276739 | 1.302149209 | 0.579773891 | 6.03E-05 | 0.000477996 |
| SH3KBP1 | 1333.740301 | 1.301851022 | 0.203488916 | 2.67E-13 | 1.37E-11 |
| ABCC11 | 28.92146273 | 1.300226196 | 0.473797488 | 1.31E-05 | 0.000129195 |
| PSPH | 1428.087999 | 1.295915302 | 0.136996674 | 6.04E-24 | 1.18E-21 |
| RHBDD2 | 2097.252586 | 1.294812252 | 0.238434985 | 9.03E-11 | 2.97E-09 |
| FLACC1 | 65.93801359 | 1.294669256 | 0.249560897 | 3.42E-10 | 1.02E-08 |
| NYAP1 | 413.0918547 | 1.293517204 | 0.236527488 | 7.25E-11 | 2.44E-09 |
| CYP46A1 | 38.30115815 | 1.291079652 | 0.375917412 | 1.12E-06 | 1.48E-05 |
| NUPR1 | 5.425356173 | 1.289938535 | 1.270541926 | 0.00051941 | 0.003021513 |
| PI15 | 206.2459584 | 1.289857252 | 0.258379345 | 9.56E-10 | 2.58E-08 |
| EML2 | 727.8126966 | 1.283559086 | 0.202168081 | 3.38E-13 | 1.71E-11 |

| | | | | | |
|---|---|---|---|---|---|
| COL11A1 | 69.65957953 | 1.283475573 | 0.341061373 | 3.00E-07 | 4.64E-06 |
| CDS1 | 87.7274917 | 1.28005911 | 0.283163583 | 1.00E-08 | 2.20E-07 |
| ADCY10 | 32.01969235 | 1.276896371 | 0.497133579 | 2.40E-05 | 0.00021867 |
| SPATA31F1 | 12.35681022 | 1.2642277 | 0.66706617 | 0.00016581 | 0.001148437 |
| GALK2 | 789.2919425 | 1.263427245 | 0.10741615 | 9.83E-35 | 5.61E-32 |
| ZNF860 | 11.36083756 | 1.26312341 | 0.73745116 | 0.00025506 | 0.001650443 |
| EFCAB5 | 61.49330907 | 1.259382058 | 0.34386104 | 4.59E-07 | 6.79E-06 |
| RWDD2A | 668.3343397 | 1.259041032 | 0.141770583 | 9.57E-22 | 1.48E-19 |
| HYPK | 229.1867586 | 1.258677081 | 0.246008738 | 4.67E-10 | 1.34E-08 |
| HMGB3P10 | 3.920621769 | 1.257738177 | 1.3091789 | 0.00052479 | 0.003046784 |
| PHF24 | 129.5421315 | 1.256554489 | 0.260204154 | 2.11E-09 | 5.28E-08 |
| UBA7 | 145.3153039 | 1.255920902 | 0.266361055 | 3.75E-09 | 8.97E-08 |
| ALG2 | 1534.567965 | 1.253753357 | 0.11229013 | 8.74E-32 | 3.83E-29 |
| TMEM39A | 2731.3857 | 1.252698318 | 0.131340829 | 2.04E-24 | 4.27E-22 |
| GLYAT | 2.828300176 | 1.250523532 | 1.348948344 | 0.0004922 | 0.002890219 |
| NACC2 | 172.1421592 | 1.249936488 | 0.23072178 | 8.71E-11 | 2.88E-09 |
| TMIE | 71.99518607 | 1.248772869 | 0.316149791 | 1.36E-07 | 2.32E-06 |
| LINC02641 | 75.20173602 | 1.247073416 | 0.286987209 | 2.26E-08 | 4.61E-07 |
| CFAP251 | 90.73257663 | 1.241962963 | 0.287207585 | 2.49E-08 | 5.03E-07 |
| GALR2 | 26.47132464 | 1.241959377 | 0.584502811 | 9.56E-05 | 0.000710637 |
| JDP2 | 428.9651417 | 1.238701879 | 0.201757639 | 1.10E-12 | 5.14E-11 |
| IER5L | 515.0041327 | 1.237933969 | 0.248801744 | 9.61E-10 | 2.59E-08 |
| ISG20 | 162.2898903 | 1.235834565 | 0.2261456 | 6.47E-11 | 2.21E-09 |
| WARS1 | 4352.391381 | 1.232897962 | 0.151308711 | 4.51E-19 | 5.18E-17 |
| SNPH | 142.2386665 | 1.229424279 | 0.321516333 | 2.37E-07 | 3.78E-06 |
| TRPM4 | 111.9594282 | 1.22910665 | 0.230448169 | 1.35E-10 | 4.29E-09 |
| AGA | 280.0232339 | 1.228971171 | 0.14470907 | 2.40E-20 | 3.16E-18 |
| SMIM32 | 19.0279093 | 1.227328933 | 0.554934592 | 7.74E-05 | 0.00059278 |
| ZFP36 | 66.16751117 | 1.226342286 | 0.463664329 | 2.06E-05 | 0.000191638 |
| RSPH3 | 400.9090133 | 1.225316473 | 0.161499507 | 3.90E-17 | 3.55E-15 |
| EPB41L1-AS1 | 11.51019612 | 1.222383288 | 0.812376312 | 0.00042986 | 0.002575639 |
| ADSS1 | 191.3147948 | 1.222259137 | 0.256345006 | 2.79E-09 | 6.85E-08 |
| CREB5 | 1723.127968 | 1.221429127 | 0.15251326 | 1.34E-18 | 1.46E-16 |
| LINC01252 | 85.13264532 | 1.218849238 | 0.248018933 | 1.30E-09 | 3.40E-08 |
| EDEM1 | 2302.172362 | 1.213200285 | 0.117261429 | 4.59E-28 | 1.45E-25 |
| IMPG2 | 41.39663615 | 1.212720059 | 0.355021803 | 1.30E-06 | 1.69E-05 |
| LINC00930 | 21.74306336 | 1.211706254 | 0.50962443 | 4.92E-05 | 0.000403272 |
| DNAJC1 | 522.9179201 | 1.206501688 | 0.170509815 | 1.69E-15 | 1.18E-13 |
| INSL5 | 5.280981095 | 1.202954781 | 1.233112828 | 0.00065667 | 0.003685526 |
| SERPINB8 | 54.70273282 | 1.201295697 | 0.340559711 | 8.42E-07 | 1.14E-05 |
| ZNF441 | 1072.273624 | 1.199586761 | 0.128276605 | 8.57E-24 | 1.65E-21 |
| KCNJ3 | 91.28949112 | 1.199017618 | 0.331778566 | 5.92E-07 | 8.44E-06 |
| MAP1LC3B | 2481.891289 | 1.197992112 | 0.164318485 | 3.36E-16 | 2.59E-14 |
| RGPD8 | 109.0854809 | 1.197497855 | 0.393205508 | 5.44E-06 | 5.90E-05 |
| SAT1 | 422.6925925 | 1.197395181 | 0.18395532 | 8.77E-14 | 4.82E-12 |
| ACSL6 | 160.2032244 | 1.197069141 | 0.224534535 | 1.31E-10 | 4.20E-09 |
| GRHL1 | 100.3243064 | 1.196633693 | 0.240136491 | 8.94E-10 | 2.43E-08 |
| TFAMP1 | 9.0779342 | 1.195646857 | 0.895823126 | 0.000607 | 0.00344338 |
| PPARG | 255.1749686 | 1.194289406 | 0.158682803 | 5.52E-17 | 4.87E-15 |
| ERICH2 | 133.2436318 | 1.19391975 | 0.209957186 | 1.65E-11 | 6.36E-10 |

| | | | | | |
|---|---|---|---|---|---|
| SHMT2 | 12391.64314 | 1.193381891 | 0.155208152 | 1.55E-17 | 1.50E-15 |
| KRCC1 | 858.8367428 | 1.192661599 | 0.163675309 | 3.40E-16 | 2.62E-14 |
| DNAJC25 | 653.4913283 | 1.189411047 | 0.152387693 | 6.07E-18 | 6.18E-16 |
| CHRM4 | 243.1280801 | 1.187507356 | 0.342617564 | 1.10E-06 | 1.46E-05 |
| ARL9 | 19.13126586 | 1.186217151 | 0.509633627 | 6.07E-05 | 0.000480728 |
| GOLGA5 | 1731.279481 | 1.18604537 | 0.104872205 | 1.03E-32 | 5.04E-30 |
| ZNF563 | 127.0968557 | 1.185344964 | 0.25366111 | 4.51E-09 | 1.06E-07 |
| SERPINH1 | 4153.023623 | 1.185027689 | 0.223197435 | 1.48E-10 | 4.66E-09 |
| RFPL2 | 35.86984805 | 1.18449274 | 0.391228321 | 5.95E-06 | 6.38E-05 |
| HYOU1 | 11434.29521 | 1.183250136 | 0.128893162 | 3.98E-23 | 7.18E-21 |
| JDP2-AS1 | 35.02490796 | 1.182492121 | 0.430500588 | 1.61E-05 | 0.00015416 |
| ALPK3 | 45.59296728 | 1.181810755 | 0.347992652 | 1.47E-06 | 1.88E-05 |
| SLC41A2 | 236.3814805 | 1.180307737 | 0.175746561 | 2.05E-14 | 1.23E-12 |
| JAZF1 | 688.7857193 | 1.178310752 | 0.132658173 | 6.02E-22 | 9.75E-20 |
| RND1 | 35.14592411 | 1.175730633 | 0.423595229 | 1.48E-05 | 0.000143415 |
| KCNT2 | 3361.325787 | 1.170374762 | 0.109792658 | 1.29E-29 | 4.74E-27 |
| PPP1R3C | 180.5795821 | 1.16963327 | 0.212209549 | 4.58E-11 | 1.61E-09 |
| TFEB | 249.071378 | 1.167973599 | 0.261577496 | 1.29E-08 | 2.78E-07 |
| CDKN2AIP | 2887.249279 | 1.166312482 | 0.104432409 | 4.63E-32 | 2.16E-29 |
| MTMR11 | 190.0254205 | 1.166085467 | 0.203857627 | 1.32E-11 | 5.17E-10 |
| AK8 | 10.62197961 | 1.163841255 | 0.792476509 | 0.0005342 | 0.003094467 |
| LRRC66 | 125.2386121 | 1.162255854 | 0.188278716 | 7.71E-13 | 3.67E-11 |
| H2BC5 | 174.9380863 | 1.159078872 | 0.195239273 | 3.47E-12 | 1.51E-10 |
| CT62 | 11.95805808 | 1.158071394 | 0.689366211 | 0.00035594 | 0.002194755 |
| ATXN1 | 1108.591615 | 1.156477666 | 0.158314782 | 2.74E-16 | 2.15E-14 |
| MIA3 | 8299.070773 | 1.154519396 | 0.108115252 | 9.89E-30 | 3.77E-27 |
| CPEB2 | 474.7793107 | 1.152558854 | 0.167768461 | 6.66E-15 | 4.29E-13 |
| MED8-AS1 | 51.14152195 | 1.151535006 | 0.305406277 | 3.28E-07 | 5.03E-06 |
| AK7 | 97.29780875 | 1.150307141 | 0.250788948 | 7.17E-09 | 1.63E-07 |
| KLHL32 | 191.6168634 | 1.147722007 | 0.173785272 | 4.28E-14 | 2.46E-12 |
| LARP1B | 1212.32754 | 1.144940385 | 0.1117756 | 9.74E-28 | 2.94E-25 |
| ANKRD31 | 32.70279249 | 1.144198187 | 0.448907547 | 3.39E-05 | 0.000293937 |
| TDRKH-AS1 | 79.64590194 | 1.142272897 | 0.263180715 | 2.45E-08 | 4.96E-07 |
| ALDH1A2 | 11.23557467 | 1.142093255 | 0.693516863 | 0.00039937 | 0.002418327 |
| UTY | 1459.420912 | 1.134192561 | 0.112242608 | 3.98E-27 | 1.11E-24 |
| CALCB | 82.87554516 | 1.13400158 | 0.353832495 | 3.42E-06 | 3.92E-05 |
| TUFT1 | 726.8452964 | 1.133903576 | 0.16456007 | 5.71E-15 | 3.71E-13 |
| MYOZ2 | 9.918525413 | 1.133110473 | 0.765762272 | 0.00056996 | 0.003260235 |
| ORAI3 | 22.0006288 | 1.131819775 | 0.453207216 | 4.13E-05 | 0.000348345 |
| SYNGR3 | 137.3505896 | 1.131761488 | 0.337386672 | 1.92E-06 | 2.36E-05 |
| ARFGAP3 | 1735.782174 | 1.13145498 | 0.110371361 | 8.72E-28 | 2.68E-25 |
| GADD45B | 8.297576527 | 1.131346174 | 1.004988367 | 0.00088795 | 0.00475517 |
| PSAT1 | 7518.742012 | 1.130604153 | 0.0846154 | 6.59E-44 | 6.78E-41 |
| PDLIM7 | 2613.828126 | 1.130105572 | 0.254004718 | 1.47E-08 | 3.12E-07 |
| CDKL5 | 548.180483 | 1.129970353 | 0.136139996 | 9.03E-20 | 1.14E-17 |
| MSX2 | 2215.228443 | 1.129287609 | 0.184906391 | 1.18E-12 | 5.49E-11 |
| COG3 | 2645.046883 | 1.129095245 | 0.113197772 | 1.49E-26 | 3.84E-24 |
| KCNA3 | 392.1748153 | 1.12781765 | 0.159504767 | 1.55E-15 | 1.09E-13 |
| SGO1-AS1 | 8.695600895 | 1.125015184 | 0.796690789 | 0.00065541 | 0.003679456 |
| SLC15A2 | 187.8201708 | 1.12300101 | 0.240515637 | 4.92E-09 | 1.15E-07 |

| | | | | | |
|---|---|---|---|---|---|
| SARNP | 33.58253393 | 1.122129812 | 0.41248153 | 2.04E-05 | 0.000190059 |
| ARHGAP31 | 224.7768132 | 1.121887496 | 0.172487675 | 8.58E-14 | 4.74E-12 |
| CNTN5 | 56.08927179 | 1.120991484 | 0.400065086 | 1.54E-05 | 0.000149011 |
| HCFC2 | 1326.626648 | 1.119849931 | 0.157407126 | 1.13E-15 | 8.01E-14 |
| RAB5A | 2216.116912 | 1.119253426 | 0.106822388 | 7.96E-29 | 2.68E-26 |
| OSCP1 | 222.4727944 | 1.119074414 | 0.180415694 | 6.44E-13 | 3.09E-11 |
| KIFC3 | 449.4462504 | 1.118792161 | 0.302724291 | 4.85E-07 | 7.10E-06 |
| ADAMTSL2 | 345.2635021 | 1.117508079 | 0.590760916 | 0.00024231 | 0.001578876 |
| GPT2 | 5899.562896 | 1.114778126 | 0.170603534 | 7.04E-14 | 3.92E-12 |
| TICAM2 | 9.589293852 | 1.114278882 | 0.736210451 | 0.00056498 | 0.003237189 |
| MYH15 | 252.7192373 | 1.112435607 | 0.213855586 | 2.86E-10 | 8.61E-09 |
| EIF2AK3 | 2080.40645 | 1.106376481 | 0.099895962 | 1.16E-31 | 4.88E-29 |
| HRH4 | 7.68548649 | 1.105829028 | 0.796071483 | 0.00071902 | 0.003977901 |
| INPP5J | 413.7688583 | 1.102600497 | 0.224610068 | 1.46E-09 | 3.77E-08 |
| ARRDC3-AS1 | 11.40928904 | 1.097435481 | 0.684698011 | 0.00049776 | 0.002917055 |
| FAM241A | 195.7369367 | 1.097422688 | 0.210235372 | 2.67E-10 | 8.06E-09 |
| LINC02878 | 42.48063135 | 1.095085485 | 0.367022425 | 8.73E-06 | 8.93E-05 |
| UCHL1-DT | 3.237568128 | 1.094903108 | 1.304445738 | 0.00067738 | 0.00378829 |
| CPO | 52.63218363 | 1.092944474 | 0.345078075 | 4.43E-06 | 4.92E-05 |
| ASB3 | 1550.533206 | 1.090023192 | 0.110574737 | 4.94E-26 | 1.21E-23 |
| NT5C1A | 28.08623329 | 1.088427603 | 0.457698363 | 6.81E-05 | 0.000531845 |
| HSD17B7P2 | 6.711327984 | 1.088003152 | 0.965976779 | 0.00101616 | 0.005329497 |
| ISL2 | 576.6922421 | 1.086205432 | 0.198785167 | 6.76E-11 | 2.29E-09 |
| RGPD5 | 130.7438367 | 1.085692018 | 0.242396724 | 1.42E-08 | 3.02E-07 |
| GARIN1A | 150.1227774 | 1.084589891 | 0.265268636 | 9.28E-08 | 1.66E-06 |
| AOC3 | 93.84850872 | 1.084261583 | 0.292337193 | 5.06E-07 | 7.38E-06 |
| ARMCX3 | 2756.420925 | 1.083732781 | 0.107352191 | 4.51E-27 | 1.24E-24 |
| ZNF555 | 812.8576789 | 1.083020012 | 0.147011173 | 1.81E-16 | 1.47E-14 |
| TVP23C-CDRT4 | 72.05934496 | 1.081967795 | 0.323075643 | 2.26E-06 | 2.72E-05 |
| TRIM23 | 1521.786521 | 1.081932642 | 0.130958312 | 1.33E-19 | 1.64E-17 |
| PLK2 | 579.0188572 | 1.079496084 | 0.135697091 | 1.73E-18 | 1.85E-16 |
| ARHGEF2 | 8850.575938 | 1.077845033 | 0.216215812 | 1.04E-09 | 2.79E-08 |
| YPEL5 | 1137.363135 | 1.073803503 | 0.100792569 | 1.29E-29 | 4.74E-27 |
| CCDC184 | 148.5956119 | 1.073344179 | 0.31203996 | 1.62E-06 | 2.04E-05 |
| MLC1 | 6.199044082 | 1.072744514 | 1.12832083 | 0.0010113 | 0.005310594 |
| RFPL3S | 105.7530417 | 1.070581188 | 0.2604402 | 8.79E-08 | 1.59E-06 |
| TMEM151A | 258.1108852 | 1.069392736 | 0.295170104 | 7.68E-07 | 1.05E-05 |
| SHISA2 | 68.08485951 | 1.069010546 | 0.335157933 | 4.40E-06 | 4.89E-05 |
| WDR35-DT | 25.47734805 | 1.068147265 | 0.470361658 | 0.00010023 | 0.000740308 |
| BNC1 | 20.92697025 | 1.068000866 | 0.483338629 | 0.00011983 | 0.000865253 |
| PLEKHF2 | 323.1370082 | 1.064339774 | 0.14492559 | 2.30E-16 | 1.84E-14 |
| TTC21A | 257.6212853 | 1.064043568 | 0.239933503 | 1.90E-08 | 3.94E-07 |
| ALLC | 7.585982305 | 1.060274497 | 0.786616211 | 0.00087892 | 0.004711713 |
| ARL4D | 1906.421694 | 1.056346607 | 0.214171822 | 1.51E-09 | 3.87E-08 |
| EML6 | 696.7964539 | 1.056123974 | 0.158701261 | 3.64E-14 | 2.12E-12 |
| PPP2R5B | 694.553181 | 1.055299803 | 0.219958422 | 3.09E-09 | 7.49E-08 |
| MDGA2 | 26.31210008 | 1.051474396 | 0.444806466 | 8.03E-05 | 0.000612268 |
| MYO7A | 23.53924973 | 1.050139247 | 0.512428053 | 0.00019865 | 0.001332769 |
| DDX39B-AS1 | 41.04089475 | 1.049646183 | 0.381872646 | 2.33E-05 | 0.000213088 |
| DNAJB2 | 1204.488242 | 1.046702328 | 0.186682397 | 3.39E-11 | 1.23E-09 |

| | | | | | |
|---|---|---|---|---|---|
| CCNG2 | 2298.792711 | 1.044376493 | 0.128946166 | 6.13E-19 | 6.85E-17 |
| KCTD11 | 131.1778208 | 1.043387633 | 0.239535488 | 3.05E-08 | 6.07E-07 |
| AKAP1-DT | 30.43118076 | 1.042781467 | 0.463653508 | 0.00011653 | 0.000844362 |
| PARP8 | 303.77031 | 1.042693381 | 0.176833339 | 5.86E-12 | 2.43E-10 |
| SLC37A2 | 155.6987397 | 1.040926488 | 0.264588479 | 2.22E-07 | 3.57E-06 |
| CRY2 | 1377.128091 | 1.040382726 | 0.154604078 | 2.34E-14 | 1.39E-12 |
| RNU6-30P | 31.44511244 | 1.039868226 | 0.449277952 | 9.71E-05 | 0.000719829 |
| BBS12 | 160.3237769 | 1.039121127 | 0.184350495 | 2.94E-11 | 1.08E-09 |
| SPATS2L | 1261.01748 | 1.03653846 | 0.209958093 | 1.61E-09 | 4.12E-08 |
| FOXO6 | 1030.130042 | 1.033988171 | 0.257510386 | 1.58E-07 | 2.65E-06 |
| FNDC3B | 2085.998637 | 1.031083112 | 0.120909672 | 1.72E-20 | 2.32E-18 |
| SH2B3 | 907.7920011 | 1.030577221 | 0.20013066 | 5.18E-10 | 1.47E-08 |
| SLC22A14 | 15.4376856 | 1.028763106 | 0.555740638 | 0.00035467 | 0.002187606 |
| ZMAT1 | 260.3080622 | 1.027585854 | 0.15572187 | 6.27E-14 | 3.52E-12 |
| LTO1P1 | 14.46657161 | 1.027186142 | 0.61225064 | 0.00053085 | 0.003078529 |
| SGK2 | 107.9718279 | 1.026367147 | 0.354606285 | 1.54E-05 | 0.000148404 |
| ABCG1 | 214.6264326 | 1.025685033 | 0.216740484 | 5.02E-09 | 1.17E-07 |
| PDZK1 | 120.1563652 | 1.024255388 | 0.248873155 | 1.05E-07 | 1.85E-06 |
| LINC01363 | 4.852733842 | 1.023232017 | 1.131411541 | 0.00113167 | 0.005853044 |
| CCDC174 | 1448.700157 | 1.022629759 | 0.118716586 | 8.54E-21 | 1.20E-18 |
| KLHDC1 | 121.5480469 | 1.022174672 | 0.227388892 | 1.71E-08 | 3.57E-07 |
| USP53 | 528.5049429 | 1.019377008 | 0.140220604 | 5.21E-16 | 3.91E-14 |
| GOLPH3-DT | 106.7174009 | 1.019122059 | 0.276401444 | 7.21E-07 | 9.95E-06 |
| MOBP | 30.88989466 | 1.018759013 | 0.393418365 | 4.47E-05 | 0.000371731 |
| LEFTY3P | 3.844384186 | 1.017756501 | 1.192217593 | 0.00101332 | 0.005318492 |
| RHCE | 16.42422546 | 1.017681485 | 0.533388661 | 0.00032079 | 0.002003264 |
| SRP19 | 1541.155611 | 1.015703009 | 0.124962664 | 5.85E-19 | 6.58E-17 |
| RPL7AP2 | 8.10592093 | 1.015083695 | 0.755524401 | 0.00102307 | 0.005358731 |
| GOLGA8N | 38.25367738 | 1.014982088 | 0.373714517 | 2.99E-05 | 0.000263888 |
| PTPDC1 | 1146.16417 | 1.010380987 | 0.112122986 | 2.62E-22 | 4.49E-20 |
| NECTIN4 | 15.35547189 | 1.009830069 | 0.531293568 | 0.00033602 | 0.002085684 |
| MCEE | 446.4590859 | 1.009465231 | 0.154282521 | 1.04E-13 | 5.65E-12 |
| ANGPT4 | 3.728895777 | 1.009450479 | 1.184665511 | 0.00104587 | 0.005460074 |
| TMEM14B-DT | 31.05514331 | 1.008510336 | 0.444815555 | 0.00012554 | 0.000900458 |
| RGS8 | 88.21114583 | 1.00690305 | 0.266571024 | 5.21E-07 | 7.56E-06 |
| TMEM130 | 903.4240756 | 1.006493492 | 0.229471844 | 3.18E-08 | 6.30E-07 |
| NAMA | 39.89081276 | 1.006084701 | 0.330259397 | 9.71E-06 | 9.84E-05 |
| STXBP5-AS1 | 6.651420149 | 1.006050248 | 0.846767327 | 0.00127061 | 0.00644678 |
| KCNJ2 | 23.8991324 | 1.005822525 | 0.454913713 | 0.00014915 | 0.001049286 |
| CRYL1 | 143.4240813 | 1.00544371 | 0.188169612 | 2.02E-10 | 6.22E-09 |
| DNAH10 | 395.5705481 | 1.000456379 | 0.188041081 | 2.37E-10 | 7.26E-09 |
| SPSB1 | 141.5118725 | 0.9987031 | 0.266972534 | 6.36E-07 | 8.98E-06 |
| SEC11C | 2031.834742 | 0.99810872 | 0.110824573 | 3.07E-22 | 5.21E-20 |
| RN7SL825P | 5.285616551 | 0.995876848 | 1.033825026 | 0.00136837 | 0.006856504 |
| NFXL1 | 2134.264894 | 0.995038926 | 0.116913465 | 2.66E-20 | 3.46E-18 |
| JUN | 7723.650032 | 0.994355611 | 0.157987357 | 6.23E-13 | 3.00E-11 |
| ARHGAP27 | 502.1019189 | 0.993783166 | 0.211571998 | 7.27E-09 | 1.64E-07 |
| OSBPL10 | 391.4430223 | 0.993764888 | 0.245988541 | 1.75E-07 | 2.90E-06 |
| CCNP | 26.762482 | 0.99286482 | 0.451322087 | 0.00016203 | 0.001127538 |
| PID1 | 167.593605 | 0.992518135 | 0.547039786 | 0.00044105 | 0.002633494 |

| | | | | | |
|---|---|---|---|---|---|
| DNAJB11 | 3213.926565 | 0.99214048 | 0.08640434 | 2.13E-33 | 1.12E-30 |
| COL20A1 | 16.81655249 | 0.990743609 | 0.579819722 | 0.00056415 | 0.003235157 |
| TLE6 | 115.8274092 | 0.990047009 | 0.23795114 | 1.03E-07 | 1.81E-06 |
| CLDN9 | 5.196975655 | 0.988199951 | 1.144809186 | 0.00118753 | 0.006095948 |
| ABHD4 | 32.08034537 | 0.986081522 | 0.408138979 | 8.83E-05 | 0.000662832 |
| TGDS | 809.7105024 | 0.985786952 | 0.131206103 | 1.07E-16 | 9.07E-15 |
| SYT17 | 744.6616273 | 0.985674779 | 0.157547823 | 8.56E-13 | 4.05E-11 |
| TRAPPC6B | 487.2176793 | 0.985128569 | 0.133071784 | 2.51E-16 | 1.99E-14 |
| FGFR3 | 626.4691369 | 0.985128301 | 0.199267847 | 2.12E-09 | 5.31E-08 |
| SLC38A4-AS1 | 22.48050458 | 0.984924321 | 0.42790778 | 0.00012492 | 0.000897267 |
| SERTM2 | 27.11665952 | 0.983658097 | 0.455343344 | 0.00018792 | 0.001272325 |
| RMST | 13.59222472 | 0.980953711 | 0.599038209 | 0.00068041 | 0.003802463 |
| ADAMTSL4 | 123.7877474 | 0.980531363 | 0.23907071 | 1.43E-07 | 2.43E-06 |
| ITSN2 | 630.6107826 | 0.980362348 | 0.127163289 | 2.41E-17 | 2.28E-15 |
| FSCN3 | 33.23456381 | 0.979487532 | 0.356976112 | 3.18E-05 | 0.000278271 |
| ID2-AS1 | 182.6537362 | 0.978750454 | 0.182975945 | 2.36E-10 | 7.24E-09 |
| MEF2A | 4130.847567 | 0.978314041 | 0.10881774 | 4.32E-22 | 7.23E-20 |
| FILIP1L | 61.30830444 | 0.978003156 | 0.341185789 | 2.11E-05 | 0.000195398 |
| AMER2 | 1194.302596 | 0.977837596 | 0.133863031 | 5.67E-16 | 4.16E-14 |
| CA5B | 228.5406669 | 0.976009744 | 0.219937944 | 3.00E-08 | 6.00E-07 |
| ABCC12 | 3.609972928 | 0.974190269 | 1.169670732 | 0.00114891 | 0.005922815 |
| KLHDC8B | 672.3841379 | 0.973871483 | 0.179996878 | 1.73E-10 | 5.41E-09 |
| PRICKLE1 | 855.833734 | 0.973846093 | 0.156715033 | 1.25E-12 | 5.77E-11 |
| RAPGEF3 | 80.96229631 | 0.973796553 | 0.269790628 | 1.27E-06 | 1.66E-05 |
| PROC | 84.84278382 | 0.973702317 | 0.24027346 | 1.87E-07 | 3.08E-06 |
| GOLGB1 | 5646.959663 | 0.972918354 | 0.113629699 | 2.12E-20 | 2.85E-18 |
| RPL35AP16 | 4.886727698 | 0.971978613 | 1.111696721 | 0.00131085 | 0.00661667 |
| DCT | 27.22899638 | 0.971323043 | 0.372744472 | 5.21E-05 | 0.000422492 |
| NANS | 1195.265151 | 0.970711737 | 0.132848951 | 5.98E-16 | 4.38E-14 |
| KIZ | 763.6614987 | 0.970221409 | 0.122929029 | 6.16E-18 | 6.24E-16 |
| CPEB4 | 1486.761823 | 0.969701439 | 0.1195133 | 1.01E-18 | 1.11E-16 |
| KCNJ9 | 174.3973839 | 0.968890348 | 0.189057735 | 8.97E-10 | 2.43E-08 |
| C14orf28 | 180.4694075 | 0.968152149 | 0.180080252 | 2.20E-10 | 6.78E-09 |
| SYVN1 | 2692.300507 | 0.966903893 | 0.1928703 | 1.67E-09 | 4.27E-08 |
| ZNF763 | 95.01992041 | 0.965891468 | 0.223092118 | 5.43E-08 | 1.02E-06 |
| HMGCL | 243.539479 | 0.96554596 | 0.173449356 | 7.45E-11 | 2.49E-09 |
| ZNF432 | 1441.906005 | 0.963050092 | 0.146274208 | 1.17E-13 | 6.28E-12 |
| DNAJA4 | 149.0635633 | 0.962951699 | 0.182504548 | 4.06E-10 | 1.19E-08 |
| COPZ2 | 54.28751765 | 0.959493632 | 0.395051102 | 9.58E-05 | 0.000711727 |
| GTF2IP14 | 36.78719451 | 0.959256653 | 0.359129346 | 4.47E-05 | 0.000371743 |
| TATDN2P3 | 2.494358715 | 0.958873887 | 1.189723153 | 0.00111085 | 0.005759811 |
| KAZALD1 | 39.22656557 | 0.956795721 | 0.420636902 | 0.00015278 | 0.001070794 |
| SEL1L | 5609.571232 | 0.956319335 | 0.07881202 | 1.38E-36 | 9.17E-34 |
| PMEPA1 | 1809.715884 | 0.956239564 | 0.204591414 | 1.07E-08 | 2.34E-07 |
| LURAP1L | 219.1891465 | 0.950554703 | 0.235419684 | 2.34E-07 | 3.75E-06 |
| HSD17B7 | 1296.719159 | 0.95015757 | 0.141754442 | 5.87E-14 | 3.32E-12 |
| SNAI2 | 1008.394755 | 0.949630053 | 0.113541782 | 1.54E-19 | 1.85E-17 |
| ATP13A4 | 17.25137526 | 0.949527835 | 0.50042411 | 0.00042853 | 0.002569925 |
| WNT8B | 18.70540037 | 0.948178929 | 0.567579371 | 0.00071973 | 0.003980774 |
| MGAM | 379.6100113 | 0.947984501 | 0.169469625 | 7.43E-11 | 2.49E-09 |

| | | | | | |
|---|---|---|---|---|---|
| DDR2 | 17887.31749 | 0.947968613 | 0.098979887 | 2.41E-24 | 5.00E-22 |
| STYXL1 | 642.0153574 | 0.947529594 | 0.108741698 | 7.48E-21 | 1.05E-18 |
| TMEM132B | 100.8190456 | 0.946670775 | 0.320236129 | 1.82E-05 | 0.00017219 |
| NRG2 | 182.2800125 | 0.94588024 | 0.207120459 | 1.99E-08 | 4.11E-07 |
| PCDH1 | 409.1946158 | 0.942909449 | 0.204431576 | 1.63E-08 | 3.42E-07 |
| FAXDC2 | 113.7369253 | 0.939770558 | 0.227662409 | 1.68E-07 | 2.80E-06 |
| TNFAIP3 | 14.21201785 | 0.936238925 | 0.719425732 | 0.00143181 | 0.007106848 |
| HBA2 | 5.751139418 | 0.934423845 | 0.94853689 | 0.00175645 | 0.008449032 |
| NR3C1 | 697.5359465 | 0.934208462 | 0.121500485 | 4.71E-17 | 4.21E-15 |
| LINC00941 | 55.81622487 | 0.932692409 | 0.370358407 | 8.31E-05 | 0.000629835 |
| SMIM38 | 27.2887698 | 0.932138267 | 0.519904602 | 0.00058872 | 0.003355418 |
| ZNF791 | 1986.981111 | 0.93116103 | 0.135265607 | 2.03E-14 | 1.22E-12 |
| CCNYL1 | 707.465627 | 0.930811913 | 0.170042632 | 1.77E-10 | 5.51E-09 |
| PPM1D | 4834.227903 | 0.930662411 | 0.098006223 | 6.64E-24 | 1.29E-21 |
| HLA-DMA | 31.79345543 | 0.929844755 | 0.411839105 | 0.00018186 | 0.001241196 |
| DMP1 | 6.159696288 | 0.926878557 | 0.963244425 | 0.00178701 | 0.008575969 |
| STXBP5L | 703.8390521 | 0.926122095 | 0.131639065 | 7.25E-15 | 4.65E-13 |
| NFKBIZ | 202.9651211 | 0.925695674 | 0.186911188 | 3.30E-09 | 7.98E-08 |
| LRRC43 | 104.4220534 | 0.925299287 | 0.319596414 | 2.54E-05 | 0.000229816 |
| GOLGA8Q | 11.09843171 | 0.925076906 | 0.711025468 | 0.00149388 | 0.007361659 |
| C4orf33 | 541.2471322 | 0.924729874 | 0.109359114 | 9.50E-20 | 1.19E-17 |
| ZBTB38 | 481.7795356 | 0.923677403 | 0.153623834 | 7.44E-12 | 3.02E-10 |
| H2BC26 | 129.230087 | 0.923151413 | 0.252839301 | 1.50E-06 | 1.91E-05 |
| UIMC1 | 1787.925484 | 0.922540102 | 0.105213006 | 6.32E-21 | 9.02E-19 |
| EXOC8 | 2077.729153 | 0.922524313 | 0.138527657 | 1.07E-13 | 5.81E-12 |
| STARD4 | 8302.726984 | 0.921793165 | 0.102162242 | 6.39E-22 | 1.02E-19 |
| ZBTB49 | 408.5171006 | 0.92133209 | 0.133943695 | 2.36E-14 | 1.39E-12 |
| ZDHHC2 | 63.04535194 | 0.921228061 | 0.37741451 | 0.00011165 | 0.000813889 |
| FRAT1 | 70.15296754 | 0.920518679 | 0.285981211 | 8.25E-06 | 8.53E-05 |
| KIAA0825 | 76.65377967 | 0.91968242 | 0.286762239 | 8.67E-06 | 8.88E-05 |
| CDC14A | 308.1070158 | 0.91953779 | 0.203304167 | 3.09E-08 | 6.13E-07 |
| SHC4 | 97.94488048 | 0.9151205 | 0.266621779 | 3.81E-06 | 4.31E-05 |
| MICU3 | 742.9229341 | 0.914828248 | 0.129908081 | 7.91E-15 | 4.96E-13 |
| ZC3H6 | 793.1956188 | 0.914501025 | 0.155329037 | 1.79E-11 | 6.85E-10 |
| ANKRA2 | 948.8293194 | 0.914240777 | 0.125996773 | 1.66E-15 | 1.16E-13 |
| LINC01376 | 62.29687101 | 0.913967989 | 0.273067234 | 5.33E-06 | 5.80E-05 |
| H2BC4 | 10.70262005 | 0.912807486 | 0.656417875 | 0.00137983 | 0.006893763 |
| PLCL1 | 583.9734923 | 0.912766239 | 0.169580087 | 3.57E-10 | 1.06E-08 |
| GSC | 82.57416079 | 0.912750358 | 0.245178087 | 1.21E-06 | 1.58E-05 |
| RAB39B | 1307.082864 | 0.912412743 | 0.11202513 | 1.55E-18 | 1.68E-16 |
| HEXIM1 | 2781.767275 | 0.911652984 | 0.137591546 | 1.54E-13 | 8.11E-12 |
| MYO1A | 27.85173376 | 0.91120534 | 0.403179199 | 0.00019828 | 0.001331653 |
| TPST1 | 1080.382318 | 0.910656717 | 0.109328522 | 3.37E-19 | 3.93E-17 |
| SRGAP2B | 585.7083859 | 0.910648141 | 0.15126971 | 8.20E-12 | 3.31E-10 |
| SERPINE2 | 2651.22486 | 0.907809991 | 0.167434537 | 3.01E-10 | 9.03E-09 |
| ABTB1 | 346.7508843 | 0.907229728 | 0.176630259 | 1.48E-09 | 3.82E-08 |
| TIPARP | 883.0612296 | 0.907218842 | 0.140933811 | 5.78E-13 | 2.82E-11 |
| SCIRT | 11.75697398 | 0.906734113 | 0.590130162 | 0.00110689 | 0.005740737 |
| ERF | 2328.65976 | 0.906721856 | 0.206625037 | 6.63E-08 | 1.22E-06 |
| WIPI1 | 394.2453684 | 0.906530249 | 0.125810289 | 2.65E-15 | 1.81E-13 |

| | | | | | |
|---|---|---|---|---|---|
| FGF14 | 308.7415126 | 0.906264557 | 0.213326398 | 1.28E-07 | 2.20E-06 |
| ATP11B | 2034.44094 | 0.903936664 | 0.114085847 | 1.06E-17 | 1.06E-15 |
| RAB30 | 4761.923159 | 0.900789358 | 0.11500051 | 2.30E-17 | 2.18E-15 |
| ABCA1 | 39.75449697 | 0.900629155 | 0.354609811 | 9.28E-05 | 0.000693486 |
| IRS2 | 1854.001318 | 0.900556246 | 0.211129313 | 1.24E-07 | 2.15E-06 |
| DTNA | 1348.65371 | 0.897509051 | 0.131724976 | 4.99E-14 | 2.84E-12 |
| STXBP3 | 1450.996963 | 0.897173383 | 0.123487653 | 1.92E-15 | 1.33E-13 |
| SMOX | 423.9348231 | 0.895357106 | 0.161644527 | 1.76E-10 | 5.50E-09 |
| ABCC2 | 222.8603252 | 0.895063618 | 0.193693073 | 2.40E-08 | 4.87E-07 |
| RSAD2 | 84.34762333 | 0.894039381 | 0.216892954 | 2.53E-07 | 3.99E-06 |
| ERRFI1 | 722.0964231 | 0.893891381 | 0.151931845 | 2.31E-11 | 8.65E-10 |
| CCDC96 | 15.38020893 | 0.89332997 | 0.575763358 | 0.00113925 | 0.005880415 |
| THRB | 138.8677201 | 0.892992922 | 0.199450463 | 4.93E-08 | 9.39E-07 |
| LRP2BP | 129.8541166 | 0.892879095 | 0.188789019 | 1.43E-08 | 3.04E-07 |
| GLT8D2 | 84.59989479 | 0.892739228 | 0.222550437 | 4.17E-07 | 6.23E-06 |
| LINC01176 | 186.0579464 | 0.892627157 | 0.206372251 | 1.01E-07 | 1.79E-06 |
| IZUMO1 | 25.3658181 | 0.892528393 | 0.407810032 | 0.00026554 | 0.001708051 |
| PLP1 | 33.91401984 | 0.892526034 | 0.539966453 | 0.00094127 | 0.005004178 |
| LINC01508 | 5.203781285 | 0.892321327 | 0.99351533 | 0.0019295 | 0.009180546 |
| MTO1 | 1869.926751 | 0.892188494 | 0.097947355 | 4.37E-22 | 7.24E-20 |
| TXNDC11 | 925.4045951 | 0.891847053 | 0.14456824 | 3.98E-12 | 1.71E-10 |
| LRIF1 | 2024.718373 | 0.891129172 | 0.10528524 | 1.39E-19 | 1.71E-17 |
| LCA5 | 497.2331101 | 0.890403165 | 0.137060781 | 4.77E-13 | 2.35E-11 |
| ZBTB2 | 2242.900911 | 0.888925425 | 0.114114042 | 3.79E-17 | 3.47E-15 |
| TRAM1 | 3582.350651 | 0.888588735 | 0.119312155 | 5.45E-16 | 4.05E-14 |
| ACVR1C | 46.71300352 | 0.888170794 | 0.290412412 | 1.84E-05 | 0.000173438 |
| NR4A1 | 1387.57989 | 0.888004086 | 0.404744412 | 0.00026805 | 0.001722578 |
| GOT1 | 2585.574607 | 0.887935765 | 0.095772495 | 1.02E-22 | 1.78E-20 |
| SLC6A1 | 35.45022985 | 0.887719023 | 0.406358296 | 0.0002753 | 0.001757621 |
| C1orf162 | 231.9072817 | 0.887274822 | 0.241206039 | 1.77E-06 | 2.21E-05 |
| ITGA3 | 250.7917737 | 0.88577577 | 0.183686054 | 9.63E-09 | 2.11E-07 |
| SEC63 | 10134.52178 | 0.884521285 | 0.086370589 | 7.50E-27 | 2.03E-24 |
| LINC00662 | 2990.472546 | 0.884408438 | 0.094304614 | 3.92E-23 | 7.13E-21 |
| CDK2AP2 | 1042.087179 | 0.8842212 | 0.217767023 | 3.63E-07 | 5.49E-06 |
| DNAH6 | 150.2947539 | 0.884042202 | 0.198881587 | 6.27E-08 | 1.16E-06 |
| SLC30A1 | 2292.411166 | 0.883994986 | 0.13461695 | 3.22E-13 | 1.64E-11 |
| RIOK3 | 3671.15503 | 0.883844149 | 0.09933 | 3.37E-21 | 4.88E-19 |
| ARHGEF2-AS2 | 26.88446964 | 0.883825838 | 0.486688523 | 0.00069363 | 0.003859237 |
| SLC22A1 | 4.922696947 | 0.883675087 | 1.064007976 | 0.00174141 | 0.008388447 |
| LY6G6C | 48.86928556 | 0.883598737 | 0.346576934 | 9.96E-05 | 0.000736202 |
| EGR1 | 977.8529964 | 0.883391055 | 0.783866071 | 0.00208837 | 0.009811553 |
| BEAN1-AS1 | 3.649253872 | 0.882693773 | 1.10807174 | 0.00156169 | 0.007649965 |
| UPP2 | 6.113426002 | 0.882408307 | 0.784867042 | 0.00210012 | 0.009853284 |
| LONRF3 | 43.00499691 | 0.881072468 | 0.294560129 | 2.43E-05 | 0.000221473 |
| FMN1 | 1399.454841 | 0.878962771 | 0.174929536 | 3.62E-09 | 8.72E-08 |
| CCDC163 | 264.5238705 | 0.878777586 | 0.186204522 | 1.74E-08 | 3.63E-07 |
| LINGO3 | 10.13733436 | 0.878332262 | 0.699748244 | 0.00189809 | 0.009058344 |
| ARG2 | 409.5310859 | 0.87831241 | 0.114841965 | 1.34E-16 | 1.12E-14 |
| LINC02951 | 38.31549104 | 0.877991141 | 0.398743457 | 0.00027699 | 0.001766231 |
| PAPPA2 | 19.41203196 | 0.877809346 | 0.51013994 | 0.00087597 | 0.004698338 |

| | | | | | |
|---|---|---|---|---|---|
| EEF1DP4 | 29.84038173 | 0.877722014 | 0.360156247 | 0.000144 | 0.00101757 |
| LINC00158 | 37.79600327 | 0.877155713 | 0.310366631 | 4.34E-05 | 0.000363394 |
| EAF2 | 84.32509803 | 0.876472512 | 0.211650714 | 2.73E-07 | 4.27E-06 |
| LSMEM1 | 44.82085838 | 0.876291272 | 0.310694653 | 4.45E-05 | 0.000370804 |
| SIAH3 | 435.0826546 | 0.875163868 | 0.209579526 | 2.36E-07 | 3.77E-06 |
| LRRC9 | 38.91135225 | 0.874317953 | 0.367874601 | 0.00017474 | 0.001200978 |
| WEE2-AS1 | 31.10723698 | 0.873912129 | 0.403222593 | 0.00031037 | 0.001950017 |
| WNT11 | 399.8558096 | 0.873625235 | 0.241874887 | 2.60E-06 | 3.08E-05 |
| KRT8P11 | 15.77455505 | 0.871875545 | 0.48101703 | 0.00074006 | 0.004078969 |
| DNAAF3-AS1 | 6.272977781 | 0.871769351 | 1.028549574 | 0.0019246 | 0.009163561 |
| KLHL15 | 2323.119797 | 0.871036544 | 0.110268408 | 2.02E-17 | 1.93E-15 |
| SCAND3 | 935.2112029 | 0.870621458 | 0.142170386 | 6.87E-12 | 2.81E-10 |
| UGDH | 3187.748221 | 0.870539326 | 0.088855025 | 8.25E-25 | 1.80E-22 |
| N4BP2L1 | 431.7851637 | 0.870145485 | 0.21051129 | 3.00E-07 | 4.64E-06 |
| NEXN | 29.54656989 | 0.870023772 | 0.380986534 | 0.00023223 | 0.001523773 |
| RNU6-446P | 6.368182621 | 0.869535101 | 0.812060079 | 0.00227026 | 0.010486419 |
| FAM161B | 303.5377411 | 0.869343311 | 0.174381358 | 4.94E-09 | 1.15E-07 |
| AFG1L | 280.8217777 | 0.869006133 | 0.13355671 | 5.84E-13 | 2.84E-11 |
| HOATZ | 26.38188062 | 0.868432864 | 0.457157061 | 0.0006186 | 0.003498582 |
| BLTP3A | 4321.148327 | 0.868396588 | 0.099816778 | 2.46E-20 | 3.23E-18 |
| SCG5 | 20.61029863 | 0.867772201 | 0.442325648 | 0.00053646 | 0.003106192 |
| SYNE3 | 681.3123863 | 0.867664899 | 0.155505962 | 1.91E-10 | 5.93E-09 |
| COLQ | 623.9882574 | 0.866437095 | 0.143436245 | 1.22E-11 | 4.79E-10 |
| JAG1 | 435.2006236 | 0.866205755 | 0.187652269 | 3.30E-08 | 6.50E-07 |
| CLUL1 | 16.7640926 | 0.865945633 | 0.500099566 | 0.00090567 | 0.004836213 |
| ZNF844 | 728.9580066 | 0.8655004 | 0.192879149 | 6.17E-08 | 1.15E-06 |
| NFIL3 | 719.2624835 | 0.864663749 | 0.149632098 | 6.16E-11 | 2.12E-09 |
| ADARB2 | 292.5435858 | 0.864064245 | 0.205067465 | 2.22E-07 | 3.58E-06 |
| HSD17B14 | 45.11779545 | 0.861934419 | 0.324701148 | 8.29E-05 | 0.000629227 |
| NBR2 | 1453.293867 | 0.861138841 | 0.144800435 | 2.32E-11 | 8.67E-10 |
| HEY1 | 236.4151332 | 0.860868638 | 0.207132431 | 2.96E-07 | 4.60E-06 |
| RIPK4 | 107.6655103 | 0.860059417 | 0.275924174 | 1.84E-05 | 0.000173547 |
| CALCOCO2 | 1742.077261 | 0.85980006 | 0.114711479 | 5.59E-16 | 4.14E-14 |
| PPIC | 19.11363573 | 0.859553215 | 0.49774783 | 0.00094299 | 0.005008172 |
| ST7L | 1304.501533 | 0.859384509 | 0.112084501 | 1.49E-16 | 1.23E-14 |
| BLTP3B | 2654.757724 | 0.859317779 | 0.111531325 | 1.11E-16 | 9.43E-15 |
| OTUD1 | 190.87024 | 0.857567467 | 0.179294309 | 1.58E-08 | 3.33E-07 |
| RPS12P3 | 20.35995755 | 0.857524792 | 0.408047612 | 0.0004023 | 0.002432479 |
| GADD45G | 1302.224246 | 0.857489183 | 0.20206204 | 2.07E-07 | 3.36E-06 |
| JAK2 | 378.3984324 | 0.856414387 | 0.134305036 | 1.62E-12 | 7.44E-11 |
| SEC24D | 5091.518096 | 0.856105443 | 0.130099065 | 4.20E-13 | 2.09E-11 |
| CRELD2 | 1199.727313 | 0.856059435 | 0.151266587 | 1.38E-10 | 4.39E-09 |
| ZNF420 | 1276.086637 | 0.855858875 | 0.111620998 | 1.56E-16 | 1.29E-14 |
| FOXO1 | 206.736371 | 0.855559486 | 0.176735591 | 1.21E-08 | 2.62E-07 |
| C12orf50 | 11.00890498 | 0.855385779 | 0.694336742 | 0.00214028 | 0.010009706 |
| TTC39A | 211.5347385 | 0.85468295 | 0.248973381 | 6.09E-06 | 6.52E-05 |
| A2ML1 | 6.454876965 | 0.853730944 | 0.740217161 | 0.00231321 | 0.010653697 |
| ST7 | 828.9157746 | 0.853645751 | 0.174750969 | 9.87E-09 | 2.17E-07 |
| PALMD | 206.6397287 | 0.85229374 | 0.456094456 | 0.00072064 | 0.003984751 |
| TACC1 | 1439.072431 | 0.850842392 | 0.120381815 | 1.51E-14 | 9.17E-13 |

| | | | | | |
|---|---|---|---|---|---|
| SELENOK | 2073.135855 | 0.850054856 | 0.103405058 | 1.95E-18 | 2.07E-16 |
| VCPIP1 | 1552.888407 | 0.849224989 | 0.111501092 | 2.55E-16 | 2.02E-14 |
| MAP1LC3B2 | 30.21473214 | 0.847483821 | 0.351522417 | 0.00018684 | 0.001266371 |
| UBR2 | 3213.557451 | 0.846623779 | 0.094189463 | 2.54E-21 | 3.74E-19 |
| HPCAL4 | 42.24702808 | 0.846508876 | 0.300356415 | 5.52E-05 | 0.000442288 |
| SLC7A5 | 7805.210733 | 0.845437259 | 0.236358218 | 3.81E-06 | 4.31E-05 |
| RGS5 | 347.059863 | 0.844983345 | 0.177925601 | 2.15E-08 | 4.41E-07 |
| RCHY1 | 1524.71908 | 0.843679712 | 0.096480304 | 2.37E-20 | 3.16E-18 |
| SNORC | 74.0674609 | 0.843387331 | 0.262602329 | 1.50E-05 | 0.000145443 |
| SDF2L1 | 865.3127555 | 0.842950729 | 0.154487693 | 5.19E-10 | 1.47E-08 |
| ERP44 | 2127.876894 | 0.842722731 | 0.0864523 | 2.03E-24 | 4.27E-22 |
| HSPA5 | 22068.47376 | 0.842379284 | 0.099526954 | 2.79E-19 | 3.28E-17 |
| NOG | 152.4277522 | 0.841941914 | 0.221217289 | 1.58E-06 | 2.00E-05 |
| SLC25A42 | 335.5109863 | 0.841411077 | 0.198854179 | 2.58E-07 | 4.05E-06 |
| WAS | 17.18015277 | 0.84087237 | 0.475419834 | 0.00094878 | 0.005035011 |
| PIWIL2 | 62.87536711 | 0.840654156 | 0.380114585 | 0.00033451 | 0.002078206 |
| PPIL4 | 2581.327299 | 0.840510454 | 0.095479446 | 1.48E-20 | 2.02E-18 |
| PKD1L3 | 7.062635418 | 0.839853018 | 0.707758758 | 0.00238717 | 0.010920475 |
| TMEFF2 | 114.7381223 | 0.839475303 | 0.209064547 | 6.75E-07 | 9.43E-06 |
| CDO1 | 603.3430566 | 0.838615625 | 0.116727908 | 7.63E-15 | 4.86E-13 |
| CASP8 | 100.4103795 | 0.838566377 | 0.275616881 | 2.81E-05 | 0.000249867 |
| MCF2L2 | 105.3919574 | 0.83817092 | 0.3074828 | 7.85E-05 | 0.00059978 |
| TMF1 | 1781.329469 | 0.836529148 | 0.107037855 | 6.40E-17 | 5.58E-15 |
| SPMIP6 | 57.38932562 | 0.835345876 | 0.306111569 | 7.93E-05 | 0.000605453 |
| KIF13B | 836.5058885 | 0.835187803 | 0.174801925 | 2.08E-08 | 4.28E-07 |
| SYT9 | 204.6951912 | 0.834569188 | 0.205036315 | 5.60E-07 | 8.08E-06 |
| SLC22A15 | 819.6099399 | 0.83412114 | 0.122939227 | 1.39E-13 | 7.36E-12 |
| ZCCHC8 | 3325.782291 | 0.833600433 | 0.099825001 | 8.28E-19 | 9.16E-17 |
| HERC2P8 | 116.6957647 | 0.833264555 | 0.237357902 | 5.48E-06 | 5.94E-05 |
| UBE2Q2P1 | 4.757434866 | 0.831283979 | 0.932851087 | 0.00250479 | 0.0113555 |
| MAP2K5 | 969.3554617 | 0.830958527 | 0.18198843 | 6.13E-08 | 1.14E-06 |
| UNC13D | 202.0878678 | 0.829680611 | 0.245017889 | 9.01E-06 | 9.18E-05 |
| NRG3 | 73.92107999 | 0.82859504 | 0.277240628 | 3.64E-05 | 0.000311458 |
| KCND3 | 29.82430384 | 0.828416604 | 0.337103701 | 0.00018569 | 0.001261062 |
| ADAMTS20 | 1922.480845 | 0.828276048 | 0.115230894 | 8.55E-15 | 5.33E-13 |
| THSD4 | 57.96738717 | 0.826908786 | 0.366203794 | 0.0003225 | 0.00201268 |
| GOLPH3L | 1389.876869 | 0.826268847 | 0.103043385 | 1.44E-17 | 1.41E-15 |
| PLEKHH3 | 2587.823368 | 0.826070979 | 0.210377864 | 1.12E-06 | 1.48E-05 |
| PDIA5 | 518.7108386 | 0.825071856 | 0.161226103 | 4.10E-09 | 9.69E-08 |
| RAB37 | 16.32277966 | 0.824867637 | 0.508442211 | 0.00137234 | 0.006869699 |
| SLC16A6 | 164.5548115 | 0.824652974 | 0.171908916 | 2.13E-08 | 4.37E-07 |
| ZDBF2 | 2999.804685 | 0.823529512 | 0.133625934 | 9.76E-12 | 3.90E-10 |
| MAFA | 6.802263287 | 0.82315738 | 0.82223363 | 0.00278456 | 0.012437385 |
| PPM1K | 1967.567647 | 0.822893185 | 0.136195924 | 2.09E-11 | 7.89E-10 |
| LMNTD1 | 8.505128683 | 0.822438141 | 0.703879638 | 0.00261336 | 0.01175835 |
| ATXN7L1 | 580.0803637 | 0.822173053 | 0.155021059 | 1.56E-09 | 4.00E-08 |
| ZNF773 | 338.6234159 | 0.822046928 | 0.185215155 | 1.23E-07 | 2.14E-06 |
| AUH | 626.6873011 | 0.821706992 | 0.113925773 | 7.80E-15 | 4.91E-13 |
| SDSL | 23.97292026 | 0.821318579 | 0.460485936 | 0.00101899 | 0.005341417 |
| RPL17P47 | 94.92874231 | 0.820849121 | 0.205107653 | 8.62E-07 | 1.17E-05 |

| | | | | | |
|---|---|---|---|---|---|
| SAMD8 | 2216.12682 | 0.820651193 | 0.108492797 | 5.66E-16 | 4.16E-14 |
| NRIP3 | 55.40162512 | 0.81986226 | 0.292423619 | 7.07E-05 | 0.000549148 |
| RIMS2 | 1356.860338 | 0.819822431 | 0.117980331 | 5.35E-14 | 3.04E-12 |
| WDR25 | 257.8917088 | 0.81973971 | 0.147881621 | 4.21E-10 | 1.22E-08 |
| LIG4 | 1380.833452 | 0.818183623 | 0.110782901 | 2.27E-15 | 1.57E-13 |
| CCDC185 | 6.874711633 | 0.818050993 | 0.822206815 | 0.00284541 | 0.012642065 |
| RNF113A | 610.2976397 | 0.817132745 | 0.103499815 | 4.44E-17 | 3.99E-15 |
| SELENOS | 2477.944031 | 0.816639268 | 0.106663901 | 2.94E-16 | 2.29E-14 |
| SPATA18 | 323.2495949 | 0.816202341 | 0.185941432 | 1.65E-07 | 2.74E-06 |
| MCTP2 | 32.00821717 | 0.815728874 | 0.399132422 | 0.00058974 | 0.003359389 |
| SQSTM1 | 1837.337649 | 0.815270751 | 0.132448437 | 1.14E-11 | 4.52E-10 |
| TPPP3 | 101.1848413 | 0.814379461 | 0.206824678 | 1.21E-06 | 1.58E-05 |
| PRPH2 | 162.6638706 | 0.813504443 | 0.243717079 | 1.24E-05 | 0.000122654 |
| MANF | 2929.321275 | 0.811945612 | 0.114862793 | 2.53E-14 | 1.49E-12 |
| BTBD8 | 262.8301801 | 0.811944943 | 0.211924904 | 1.91E-06 | 2.36E-05 |
| DRP2 | 150.0575472 | 0.811547625 | 0.16314566 | 1.01E-08 | 2.22E-07 |
| DUSP1 | 960.8336353 | 0.811455911 | 0.179555421 | 9.50E-08 | 1.69E-06 |
| DRC1 | 5.352687814 | 0.811406171 | 0.872338402 | 0.00285143 | 0.012659082 |
| PPP1R13B | 1451.627414 | 0.811278624 | 0.16923758 | 2.53E-08 | 5.09E-07 |
| GAS7 | 265.55081 | 0.810715677 | 0.224533722 | 4.62E-06 | 5.11E-05 |
| SLC33A1 | 1242.14495 | 0.810614248 | 0.132413401 | 1.49E-11 | 5.76E-10 |
| LINC01512 | 5.627712866 | 0.81035117 | 0.891548077 | 0.00281316 | 0.012536367 |
| IL5RA | 6.308357401 | 0.809806748 | 0.786027361 | 0.00295201 | 0.013024069 |
| MTARC2 | 247.3510725 | 0.809516598 | 0.170622087 | 3.29E-08 | 6.49E-07 |
| MYDGF | 1879.882397 | 0.808938654 | 0.150312139 | 1.19E-09 | 3.14E-08 |
| PLIN1 | 18.16208992 | 0.807684478 | 0.517415349 | 0.00166935 | 0.008100622 |
| DNAJC10 | 7991.2003 | 0.806325567 | 0.090251124 | 7.40E-21 | 1.05E-18 |
| BACH2 | 1422.180815 | 0.80553034 | 0.127488137 | 4.57E-12 | 1.94E-10 |
| ZFAS1 | 1988.7597 | 0.805379485 | 0.092364278 | 5.09E-20 | 6.58E-18 |
| PTHLH | 19.31777671 | 0.80428975 | 0.442364269 | 0.00104157 | 0.005444528 |
| TENT4B | 1909.037161 | 0.804168396 | 0.097184045 | 2.37E-18 | 2.50E-16 |
| NFAT5 | 3204.330579 | 0.803593053 | 0.13165145 | 1.82E-11 | 6.97E-10 |
| ATP9B | 968.4225887 | 0.802811083 | 0.146331758 | 7.17E-10 | 1.98E-08 |
| P4HA1 | 736.7660595 | 0.802734248 | 0.133103915 | 2.90E-11 | 1.07E-09 |
| GLIPR1 | 38.83128143 | 0.802584707 | 0.344598923 | 0.00031417 | 0.001970867 |
| CANX | 37705.46856 | 0.802137075 | 0.086906622 | 5.20E-22 | 8.56E-20 |
| FAM151B | 79.4754028 | 0.801625976 | 0.218944999 | 4.15E-06 | 4.65E-05 |
| EVI5L | 1093.180245 | 0.800632307 | 0.175124019 | 8.37E-08 | 1.52E-06 |
| TNXB | 46.48810066 | 0.799909438 | 0.412843231 | 0.00082678 | 0.00446128 |
| B3GNT2 | 956.2777673 | 0.799430177 | 0.108076865 | 2.69E-15 | 1.82E-13 |
| VLDLR-AS1 | 683.3104872 | 0.799408927 | 0.217743475 | 4.08E-06 | 4.59E-05 |
| SAXO4 | 108.9884075 | 0.799077892 | 0.215223805 | 3.49E-06 | 3.98E-05 |
| RWDD2B | 1146.127954 | 0.799009796 | 0.103190778 | 1.90E-16 | 1.53E-14 |
| TMEM38A | 255.4523879 | 0.798814962 | 0.19368703 | 6.45E-07 | 9.09E-06 |
| CSGALNACT2 | 2346.215086 | 0.798539696 | 0.09332217 | 2.32E-19 | 2.75E-17 |
| USP50 | 155.9991236 | 0.798490295 | 0.183810232 | 2.46E-07 | 3.90E-06 |
| DEPDC4 | 168.2432709 | 0.798389796 | 0.174751597 | 8.72E-08 | 1.57E-06 |
| CSDC2 | 452.6469058 | 0.797893202 | 0.471940189 | 0.00139037 | 0.006939686 |
| VWC2L | 4.453266954 | 0.7966083 | 1.000234522 | 0.00242696 | 0.011068544 |
| LMOD1 | 10.94892059 | 0.796605134 | 0.590970411 | 0.00243217 | 0.01108984 |

| | | | | | |
|---|---|---|---|---|---|
| TGFB2 | 32.98147129 | 0.796337631 | 0.317315191 | 0.00020132 | 0.001347223 |
| ATF6 | 4279.217912 | 0.796157021 | 0.097372522 | 5.99E-18 | 6.13E-16 |
| TMEM135 | 1185.832392 | 0.795994218 | 0.11503834 | 9.04E-14 | 4.96E-12 |
| CA8 | 17.26823658 | 0.795586344 | 0.421329845 | 0.00094443 | 0.005014545 |
| HAPSTR1 | 3627.629549 | 0.795411037 | 0.109532171 | 7.73E-15 | 4.90E-13 |
| SLC12A4 | 255.4698717 | 0.795188592 | 0.441248095 | 0.00113355 | 0.005859809 |
| DNHD1 | 1747.078567 | 0.794965826 | 0.168219978 | 4.26E-08 | 8.26E-07 |
| RNF6 | 1195.172549 | 0.794545977 | 0.118645892 | 4.30E-13 | 2.13E-11 |
| PLA2G10 | 11.83730912 | 0.793339541 | 0.547069539 | 0.00214421 | 0.01002581 |
| ZNF10 | 2117.742051 | 0.792938718 | 0.105578248 | 1.24E-15 | 8.80E-14 |
| RABGAP1L | 1486.072618 | 0.791839762 | 0.112026408 | 3.31E-14 | 1.94E-12 |
| WNK4 | 46.95771704 | 0.791781634 | 0.288915226 | 0.00010693 | 0.000784219 |
| LRIT3 | 120.1704249 | 0.791734626 | 0.193598874 | 8.07E-07 | 1.10E-05 |
| DNAH12 | 11.96409383 | 0.790117284 | 0.561376111 | 0.00231782 | 0.01067017 |
| IGFBP6 | 28.76693752 | 0.789766165 | 0.351892438 | 0.00042654 | 0.00256174 |
| SERPINE3 | 136.0972001 | 0.789433452 | 0.197914844 | 1.26E-06 | 1.65E-05 |
| NBEAL1 | 782.4898604 | 0.789428179 | 0.109742311 | 1.38E-14 | 8.44E-13 |
| XYLT1 | 611.9234702 | 0.788849145 | 0.171140624 | 8.01E-08 | 1.46E-06 |
| MROH8 | 361.1219745 | 0.788582145 | 0.193455333 | 8.84E-07 | 1.19E-05 |
| ZNF572 | 278.4607299 | 0.788448816 | 0.135579032 | 1.28E-10 | 4.11E-09 |
| OLMALINC | 611.3426463 | 0.788400998 | 0.140468309 | 4.17E-10 | 1.22E-08 |
| PDE4DIPP2 | 602.9504991 | 0.788179764 | 0.153357295 | 5.66E-09 | 1.30E-07 |
| CPEB1 | 119.1319035 | 0.788050617 | 0.260464625 | 4.53E-05 | 0.000376217 |
| CACNG3 | 44.22997181 | 0.787821043 | 0.392577411 | 0.00076246 | 0.004181145 |
| ZKSCAN4 | 818.5331764 | 0.787668884 | 0.104516323 | 1.09E-15 | 7.80E-14 |
| EPHX1 | 568.9846402 | 0.787393881 | 0.190160138 | 6.80E-07 | 9.48E-06 |
| LINC02572 | 35.18195736 | 0.787290066 | 0.308868767 | 0.00019304 | 0.001301942 |
| LMNA | 2346.402664 | 0.786872872 | 0.156256778 | 9.88E-09 | 2.17E-07 |
| CEBPG | 6335.301161 | 0.786807756 | 0.092414478 | 3.94E-19 | 4.55E-17 |
| KCNK9 | 51.17664742 | 0.786691593 | 0.317529294 | 0.00023632 | 0.001547169 |
| UFL1 | 1931.570008 | 0.786419653 | 0.104069149 | 9.51E-16 | 6.86E-14 |
| ACE | 393.9828752 | 0.785732401 | 0.199332143 | 1.60E-06 | 2.02E-05 |
| RNU6-548P | 10.69930996 | 0.785387525 | 0.559328875 | 0.00238418 | 0.010910491 |
| GDNF | 6.119760857 | 0.785170054 | 0.778053627 | 0.00330205 | 0.014253428 |
| NUDT17 | 313.4202824 | 0.783160591 | 0.148732607 | 3.07E-09 | 7.45E-08 |
| EFHB | 53.70839153 | 0.781519793 | 0.307304564 | 0.00020507 | 0.001368292 |
| LZTFL1 | 1152.571869 | 0.781465572 | 0.11054926 | 3.78E-14 | 2.19E-12 |
| ZNF630 | 164.4289495 | 0.780823597 | 0.158600142 | 1.89E-08 | 3.92E-07 |
| PIBF1 | 1397.491606 | 0.780141119 | 0.099326181 | 1.01E-16 | 8.63E-15 |
| CXXC4 | 494.3232637 | 0.779996603 | 0.154755364 | 1.05E-08 | 2.29E-07 |
| LEAP2 | 127.6694357 | 0.779489192 | 0.196088808 | 1.49E-06 | 1.90E-05 |
| RHO | 5.824382278 | 0.779473846 | 0.811677352 | 0.00335764 | 0.01445394 |
| CSNK1G1 | 2689.277291 | 0.779330176 | 0.115572246 | 3.81E-13 | 1.91E-11 |
| SGMS1 | 485.4898674 | 0.779148694 | 0.152377875 | 7.24E-09 | 1.64E-07 |
| ABTB2 | 461.3846807 | 0.779078956 | 0.192484791 | 1.10E-06 | 1.46E-05 |
| GRIA3 | 168.7500938 | 0.778922678 | 0.180159766 | 3.35E-07 | 5.10E-06 |
| YAF2 | 1850.360718 | 0.778640814 | 0.096768722 | 2.20E-17 | 2.09E-15 |
| LMBRD2 | 2039.46345 | 0.778590544 | 0.131777213 | 8.28E-11 | 2.75E-09 |
| PNPLA8 | 1411.145976 | 0.778504787 | 0.114249033 | 2.36E-13 | 1.22E-11 |
| HELQ | 718.8583067 | 0.778393226 | 0.136747773 | 2.98E-10 | 8.96E-09 |

| | | | | | |
|---|---|---|---|---|---|
| SPRY4 | 583.1247777 | 0.776313079 | 0.471503401 | 0.00169882 | 0.00821409 |
| CAMSAP3 | 136.9237182 | 0.775783258 | 0.298649358 | 0.00018409 | 0.001252288 |
| CASP3 | 2703.486047 | 0.775419551 | 0.104258916 | 2.72E-15 | 1.84E-13 |
| ST8SIA5 | 211.1248312 | 0.775368483 | 0.184495118 | 5.93E-07 | 8.44E-06 |
| ZDHHC11B | 554.3425283 | 0.774723001 | 0.171716321 | 1.49E-07 | 2.51E-06 |
| LINC01012 | 264.206008 | 0.774449741 | 0.174207619 | 2.03E-07 | 3.30E-06 |
| SEC63P1 | 28.95544981 | 0.77401678 | 0.35205172 | 0.00053244 | 0.003085129 |
| PIP4P2 | 641.5548149 | 0.773338018 | 0.119908821 | 2.95E-12 | 1.30E-10 |
| ATXN1-AS1 | 45.67184356 | 0.772890363 | 0.314891055 | 0.00027914 | 0.001776588 |
| DUSP3 | 2265.394482 | 0.772222211 | 0.125557603 | 2.03E-11 | 7.67E-10 |
| TOB1-AS1 | 34.78740054 | 0.77188466 | 0.356921037 | 0.00058928 | 0.003357693 |
| TTC41P | 73.20183403 | 0.771828156 | 0.217140825 | 8.38E-06 | 8.63E-05 |
| RAB28P5 | 34.87426103 | 0.771712256 | 0.304438127 | 0.00022643 | 0.001488608 |
| RORA | 513.7402665 | 0.771659696 | 0.147180942 | 3.98E-09 | 9.45E-08 |
| ELOVL4 | 1164.808651 | 0.771031371 | 0.101224937 | 7.31E-16 | 5.31E-14 |
| MSANTD1 | 39.12208935 | 0.769698324 | 0.305609067 | 0.00024055 | 0.001569851 |
| COG6 | 1346.289412 | 0.768656459 | 0.149686217 | 7.31E-09 | 1.64E-07 |
| KDM3A | 4174.309168 | 0.768445383 | 0.088385907 | 1.05E-19 | 1.30E-17 |
| COQ10B | 498.3528187 | 0.767795383 | 0.136318653 | 4.80E-10 | 1.37E-08 |
| ACBD3 | 3521.997012 | 0.767544359 | 0.089735867 | 3.61E-19 | 4.19E-17 |
| PRUNE2 | 1981.958113 | 0.767296108 | 0.158537962 | 3.35E-08 | 6.59E-07 |
| PEG13 | 45.43945665 | 0.767100283 | 0.31571166 | 0.00031156 | 0.001956322 |
| PIK3IP1 | 305.0340401 | 0.766765002 | 0.180046105 | 5.10E-07 | 7.43E-06 |
| CCDC186 | 933.5692596 | 0.765397597 | 0.114442314 | 6.62E-13 | 3.17E-11 |
| ZNF597 | 417.3912306 | 0.765144213 | 0.134764 | 3.82E-10 | 1.12E-08 |
| SCFD1 | 2393.130741 | 0.764402192 | 0.091919725 | 2.84E-18 | 2.98E-16 |
| TAMALIN | 263.5517384 | 0.76333591 | 0.23571455 | 2.80E-05 | 0.000249452 |
| SCYL2 | 5103.454189 | 0.763227687 | 0.098075456 | 2.23E-16 | 1.79E-14 |
| PARD6B | 215.5823591 | 0.763123316 | 0.172850648 | 2.65E-07 | 4.15E-06 |
| UBASH3B | 130.8975389 | 0.762360896 | 0.312644283 | 0.0003158 | 0.001979899 |
| PPP2R2B | 929.5426664 | 0.762065318 | 0.126661636 | 5.27E-11 | 1.84E-09 |
| DCP1B | 1149.194121 | 0.761480609 | 0.100444994 | 1.09E-15 | 7.80E-14 |
| TECTA | 344.0074565 | 0.761395813 | 0.15791543 | 3.93E-08 | 7.66E-07 |
| NOXRED1 | 96.89229135 | 0.761359188 | 0.228624411 | 2.08E-05 | 0.000193191 |
| SDCCAG8 | 1152.555262 | 0.761258792 | 0.113382013 | 5.84E-13 | 2.84E-11 |
| ECEL1 | 172.2767443 | 0.760656127 | 0.608040484 | 0.00323853 | 0.014020462 |
| MORC4 | 132.8593819 | 0.760092087 | 0.219152167 | 1.29E-05 | 0.00012731 |
| MIA2 | 767.6009389 | 0.759512771 | 0.094762057 | 3.63E-17 | 3.34E-15 |
| DSEL | 200.1620666 | 0.759322983 | 0.162745393 | 8.59E-08 | 1.55E-06 |
| SLC10A3 | 275.8830251 | 0.758937779 | 0.246442483 | 4.94E-05 | 0.000404549 |
| PDIA4 | 5083.809587 | 0.75822943 | 0.113816382 | 8.64E-13 | 4.08E-11 |
| SHISA4 | 117.1474225 | 0.758193569 | 0.199519306 | 3.77E-06 | 4.28E-05 |
| NPHP1 | 25.61663632 | 0.757580407 | 0.361858625 | 0.00076421 | 0.004187393 |
| YPEL2 | 756.2951587 | 0.757107222 | 0.128862653 | 1.32E-10 | 4.20E-09 |
| TMEM41B | 2873.423026 | 0.756660965 | 0.102623256 | 5.58E-15 | 3.66E-13 |
| CNOT4 | 1392.344429 | 0.755963085 | 0.125541656 | 5.50E-11 | 1.92E-09 |
| SERPINC1 | 11.30197855 | 0.755475807 | 0.55198747 | 0.00291313 | 0.012883556 |
| CEP112 | 549.6767617 | 0.755075723 | 0.146231681 | 7.39E-09 | 1.66E-07 |
| CFP | 64.54176286 | 0.754447242 | 0.240766771 | 4.32E-05 | 0.000362046 |
| FILIP1 | 34.76868581 | 0.754391622 | 0.364853304 | 0.00082843 | 0.004468975 |

| | | | | | |
|---|---|---|---|---|---|
| ZMYM5 | 1159.490566 | 0.75434204 | 0.106431749 | 4.66E-14 | 2.66E-12 |
| ZFHX4-AS1 | 63.58429427 | 0.753615177 | 0.293883159 | 0.00024137 | 0.001574716 |
| SDE2 | 3167.411165 | 0.753540497 | 0.098023844 | 5.29E-16 | 3.95E-14 |
| EGFEM1P | 85.53367489 | 0.752332247 | 0.275090077 | 0.00015094 | 0.001060067 |
| FOSB | 23.81353315 | 0.752149357 | 0.463364261 | 0.00203054 | 0.009568279 |
| ZBTB47-AS1 | 17.02433365 | 0.751885349 | 0.436415891 | 0.00170555 | 0.008239617 |
| GRIN2C | 283.4489397 | 0.751776814 | 0.216456929 | 1.38E-05 | 0.000135202 |
| PSMD5 | 36.19914604 | 0.751001964 | 0.37008626 | 0.00092339 | 0.004924425 |
| HSD17B1P1 | 217.3048077 | 0.750826732 | 0.193363264 | 2.93E-06 | 3.44E-05 |
| DCAF8L1 | 2.489496435 | 0.750732461 | 1.051401664 | 0.00220957 | 0.010270694 |
| SPRY2 | 644.4986479 | 0.750507254 | 0.400604787 | 0.00128675 | 0.006514208 |
| DNAH7 | 19.82253087 | 0.750486237 | 0.46659999 | 0.00210206 | 0.00986013 |
| C5orf34 | 733.8828454 | 0.750176749 | 0.1151282 | 2.53E-12 | 1.13E-10 |
| MN1 | 413.752325 | 0.749520142 | 0.184490669 | 1.43E-06 | 1.83E-05 |
| TMEM272 | 43.57385717 | 0.749315371 | 0.265665373 | 0.00012052 | 0.000869602 |
| HOXD1 | 6.378070761 | 0.748534389 | 0.70930368 | 0.00390361 | 0.016382422 |
| ZNF181 | 1050.674492 | 0.748501775 | 0.12527982 | 8.01E-11 | 2.68E-09 |
| SLC26A7 | 58.56895953 | 0.748278759 | 0.255732278 | 8.85E-05 | 0.000663109 |
| CREG2 | 76.62658519 | 0.747359493 | 0.351314127 | 0.00076022 | 0.004171046 |
| ZNF416 | 326.7960209 | 0.747109608 | 0.138000115 | 2.10E-09 | 5.28E-08 |
| PITPNM3 | 214.2485559 | 0.747021023 | 0.462429214 | 0.00212003 | 0.009926294 |
| ELP4 | 703.9033099 | 0.745949282 | 0.108939479 | 2.82E-13 | 1.44E-11 |
| PPIAP53 | 44.29119621 | 0.745910931 | 0.262866656 | 0.00011805 | 0.000854186 |
| ZNF14 | 571.7947547 | 0.745314115 | 0.185331642 | 1.77E-06 | 2.21E-05 |
| GPCPD1 | 3016.990073 | 0.745256167 | 0.107497049 | 1.56E-13 | 8.22E-12 |
| PALLD | 2023.132265 | 0.744952773 | 0.134720912 | 1.13E-09 | 3.00E-08 |
| MCF2 | 27.0414645 | 0.744745308 | 0.357994182 | 0.00086131 | 0.004628165 |
| TMEM214 | 2248.521272 | 0.744665528 | 0.1392186 | 3.09E-09 | 7.49E-08 |
| IFNGR1 | 1406.102892 | 0.744580507 | 0.101116578 | 6.96E-15 | 4.47E-13 |
| YOD1 | 2175.678895 | 0.74421845 | 0.098501371 | 1.64E-15 | 1.15E-13 |
| ZNF569 | 1472.581759 | 0.744148751 | 0.098131733 | 1.33E-15 | 9.38E-14 |
| PPM1E | 6464.198343 | 0.743970966 | 0.092251134 | 2.94E-17 | 2.73E-15 |
| SARS1 | 14539.42968 | 0.743636506 | 0.093984762 | 1.01E-16 | 8.63E-15 |
| SCG2 | 5082.51814 | 0.743032571 | 0.131537873 | 5.87E-10 | 1.65E-08 |
| TMEM232 | 56.54450996 | 0.742597992 | 0.230346344 | 3.59E-05 | 0.000308339 |
| BTG2 | 889.6069095 | 0.742596458 | 0.116912913 | 8.11E-12 | 3.28E-10 |
| TTLL11 | 2044.985636 | 0.741666856 | 0.154451686 | 5.43E-08 | 1.02E-06 |
| MIPEPP1 | 3.183320829 | 0.741146144 | 0.997169658 | 0.00271428 | 0.012157793 |
| DUOX2 | 41.94418281 | 0.740986478 | 0.31724788 | 0.00048483 | 0.002854286 |
| PPM1M | 575.7559402 | 0.740579153 | 0.172811083 | 6.06E-07 | 8.59E-06 |
| PRSS58 | 3.091287627 | 0.740494149 | 0.989106659 | 0.00278768 | 0.0124486 |
| PARD6G-AS1 | 231.2798384 | 0.739824681 | 0.198676276 | 6.14E-06 | 6.56E-05 |
| LINC02680 | 6.755828785 | 0.739117169 | 0.678304894 | 0.0040432 | 0.016872777 |
| CACNG5 | 186.5158597 | 0.739081005 | 0.208731745 | 1.23E-05 | 0.000121586 |
| IL20RB | 70.73563683 | 0.738309583 | 0.2158346 | 1.90E-05 | 0.000179097 |
| CYBA | 9.87626987 | 0.738018408 | 0.544592921 | 0.00325006 | 0.014061474 |
| HDAC5 | 6186.077456 | 0.736556583 | 0.17667854 | 1.05E-06 | 1.39E-05 |
| PCP4L1 | 92.81303337 | 0.736245483 | 0.236677513 | 5.53E-05 | 0.00044277 |
| CZIB-DT | 19.5877828 | 0.735868926 | 0.431915291 | 0.00194779 | 0.009248279 |
| ACSL3 | 5294.024997 | 0.735720268 | 0.088550636 | 4.32E-18 | 4.46E-16 |

| | | | | | |
|---|---|---|---|---|---|
| ZNF451 | 4749.091015 | 0.735652758 | 0.107171241 | 2.83E-13 | 1.45E-11 |
| SOCS3 | 34.74040138 | 0.735482809 | 0.37559648 | 0.00119551 | 0.006133861 |
| STON1 | 297.3308252 | 0.735263735 | 0.175587027 | 9.82E-07 | 1.32E-05 |
| RHOQ-AS1 | 50.71708046 | 0.734932646 | 0.283353261 | 0.00026024 | 0.001679127 |
| DNM3 | 531.7165365 | 0.734301215 | 0.136645205 | 3.04E-09 | 7.39E-08 |
| PPP1R21 | 836.6929975 | 0.734120549 | 0.106752566 | 2.64E-13 | 1.36E-11 |
| LINC02202 | 57.63232493 | 0.73350544 | 0.304202843 | 0.00042548 | 0.002556122 |
| CFAP300 | 131.7479911 | 0.733349845 | 0.16987197 | 5.71E-07 | 8.19E-06 |
| APBB2 | 1435.856985 | 0.731376075 | 0.107126892 | 3.85E-13 | 1.92E-11 |
| IFT74 | 709.9116591 | 0.730815818 | 0.109369533 | 1.05E-12 | 4.89E-11 |
| LBH | 842.579091 | 0.73064492 | 0.111494272 | 2.49E-12 | 1.12E-10 |
| LINC00640 | 8.336349596 | 0.73013417 | 0.648586245 | 0.00415889 | 0.017267011 |
| HULC | 72.47203898 | 0.729816145 | 0.22995661 | 4.79E-05 | 0.000395043 |
| CCDC181 | 245.1220224 | 0.729340439 | 0.188489311 | 3.89E-06 | 4.39E-05 |
| FZD6 | 177.8359727 | 0.729016839 | 0.159183202 | 1.81E-07 | 3.00E-06 |
| SLC10A7 | 266.3527477 | 0.728863303 | 0.161466869 | 2.47E-07 | 3.90E-06 |
| NDRG1 | 484.3499411 | 0.728680049 | 0.185198509 | 3.02E-06 | 3.52E-05 |
| OGFRL1 | 1037.587453 | 0.727580936 | 0.112147653 | 3.99E-12 | 1.71E-10 |
| EPHB3 | 649.5455412 | 0.727267158 | 0.229113931 | 4.91E-05 | 0.000402251 |
| NEU1 | 1066.962701 | 0.727093315 | 0.140349899 | 9.33E-09 | 2.06E-07 |
| INTS6-AS1 | 138.2665734 | 0.726789467 | 0.231125257 | 5.43E-05 | 0.0004366 |
| CCDC74B | 340.9497882 | 0.726738504 | 0.160452785 | 2.35E-07 | 3.76E-06 |
| GABRG2 | 37.75595726 | 0.726350097 | 0.381473438 | 0.00142096 | 0.007061519 |
| ARMH1 | 65.99154023 | 0.72601492 | 0.270796923 | 0.00022098 | 0.001460221 |
| NIPAL1 | 55.65652863 | 0.725332727 | 0.24905214 | 0.00011373 | 0.000826532 |
| TYW5 | 1239.83243 | 0.725250068 | 0.111065998 | 3.10E-12 | 1.36E-10 |
| FCGBP | 51.44806281 | 0.725230114 | 0.255174695 | 0.00014032 | 0.000995694 |
| CFAP90 | 17.05286246 | 0.725190573 | 0.443126693 | 0.00233466 | 0.010723705 |
| DNAI7 | 15.77338029 | 0.725117934 | 0.447190113 | 0.00239401 | 0.01094033 |
| PCMTD1-DT | 305.9162066 | 0.725111502 | 0.170750909 | 8.50E-07 | 1.16E-05 |
| ZNF841 | 1617.340148 | 0.724846873 | 0.139365654 | 8.59E-09 | 1.90E-07 |
| ZNF534 | 29.97034236 | 0.724310676 | 0.314290652 | 0.00059618 | 0.003387643 |
| SPOCK2 | 181.4448505 | 0.723764543 | 0.229308136 | 5.38E-05 | 0.00043359 |
| MEG9 | 208.1475032 | 0.723616293 | 0.236638667 | 7.38E-05 | 0.000569259 |
| TAOK3 | 1448.787121 | 0.72349601 | 0.097432151 | 5.61E-15 | 3.66E-13 |
| KCNB1 | 2052.30608 | 0.72321737 | 0.146985531 | 3.72E-08 | 7.27E-07 |
| TDH | 140.8200246 | 0.722265288 | 0.198027195 | 9.84E-06 | 9.96E-05 |
| DHRS3 | 4.55895555 | 0.72204929 | 0.868225353 | 0.003907 | 0.016393311 |
| HLF | 727.9848402 | 0.721870618 | 0.125760951 | 4.42E-10 | 1.27E-08 |
| IGIP | 407.2075021 | 0.720591687 | 0.179820711 | 2.45E-06 | 2.92E-05 |
| SGIP1 | 4507.873344 | 0.720544651 | 0.131721921 | 2.09E-09 | 5.25E-08 |
| STOML1 | 477.7098552 | 0.720263738 | 0.164274705 | 4.88E-07 | 7.14E-06 |
| AHCYL2 | 1319.605738 | 0.720162518 | 0.107380474 | 1.00E-12 | 4.69E-11 |
| SPATA7 | 326.0276857 | 0.720048718 | 0.127267718 | 7.27E-10 | 2.00E-08 |
| ITGA7 | 142.9301943 | 0.719887364 | 0.25092806 | 0.00013615 | 0.0009698 |
| UBC | 25474.80284 | 0.719642816 | 0.097462982 | 8.02E-15 | 5.01E-13 |
| BBS7 | 1653.138883 | 0.719310985 | 0.110412369 | 3.67E-12 | 1.59E-10 |
| PGM3 | 3922.839664 | 0.719237905 | 0.101888636 | 8.66E-14 | 4.78E-12 |
| TTC39B | 213.4467982 | 0.718906563 | 0.154654302 | 1.47E-07 | 2.48E-06 |
| GMDS-DT | 128.1909054 | 0.718766752 | 0.192400826 | 7.31E-06 | 7.65E-05 |

| | | | | | |
|---|---|---|---|---|---|
| ARL13B | 1168.263707 | 0.718367232 | 0.100179835 | 3.91E-14 | 2.27E-12 |
| EAPP | 968.1686292 | 0.717625266 | 0.097089787 | 7.75E-15 | 4.90E-13 |
| TOGARAM1 | 1555.315217 | 0.716655454 | 0.092884395 | 6.57E-16 | 4.79E-14 |
| VAMP4 | 1645.245654 | 0.716551841 | 0.10999155 | 3.79E-12 | 1.63E-10 |
| ERLEC1 | 1826.812635 | 0.716515358 | 0.10563287 | 6.19E-13 | 2.99E-11 |
| LARGE2 | 45.28258542 | 0.716140477 | 0.331725459 | 0.00089174 | 0.004772985 |
| HSP90B1 | 37024.24109 | 0.71581909 | 0.08880892 | 4.24E-17 | 3.82E-15 |
| ITPKC | 424.8126955 | 0.715743646 | 0.186049413 | 4.90E-06 | 5.39E-05 |
| CHRD | 118.9204272 | 0.715613068 | 0.227250778 | 5.96E-05 | 0.000473409 |
| SLC12A1 | 33.81999189 | 0.715510594 | 0.314934775 | 0.00069097 | 0.003845486 |
| LINC01605 | 11.24265554 | 0.715467951 | 0.558281682 | 0.00398161 | 0.016658756 |
| VPS37A | 1295.219722 | 0.715385904 | 0.100799627 | 6.89E-14 | 3.86E-12 |
| ANKRD34A | 121.9045379 | 0.713719756 | 0.200194369 | 1.45E-05 | 0.000141528 |
| MYLK4 | 33.36633823 | 0.71305698 | 0.334130405 | 0.0009647 | 0.005103712 |
| GRB10 | 2666.111652 | 0.712913449 | 0.198153093 | 1.30E-05 | 0.000128205 |
| CGRRF1 | 653.6407775 | 0.712778403 | 0.110455364 | 5.92E-12 | 2.45E-10 |
| CFL1P5 | 11.8446132 | 0.712753087 | 0.529500668 | 0.00377675 | 0.01595093 |
| OSBP | 3415.195298 | 0.710762368 | 0.123769121 | 4.93E-10 | 1.41E-08 |
| SYNJ1 | 1665.406507 | 0.71044742 | 0.110572813 | 7.29E-12 | 2.97E-10 |
| H4C14 | 10.28792686 | 0.710271004 | 0.533438189 | 0.0038923 | 0.016344975 |
| ZNF44 | 1631.957603 | 0.710036304 | 0.157587721 | 3.15E-07 | 4.85E-06 |
| PLEKHA5 | 2596.156869 | 0.709381223 | 0.112878117 | 1.83E-11 | 6.97E-10 |
| KRT18P55 | 10.27842675 | 0.708899187 | 0.586272626 | 0.00439902 | 0.0180742 |
| RHBDD3 | 2458.695354 | 0.708838852 | 0.151976519 | 1.52E-07 | 2.56E-06 |
| EPS15 | 3413.326049 | 0.708670915 | 0.086405284 | 1.43E-17 | 1.40E-15 |
| RPL26L1-AS1 | 23.44784137 | 0.708536233 | 0.397568518 | 0.00202167 | 0.009533068 |
| SSR3 | 7585.389764 | 0.708499488 | 0.094310607 | 3.44E-15 | 2.30E-13 |
| WHRN | 28.65028164 | 0.708465109 | 0.413106342 | 0.00227639 | 0.010510013 |
| ARID4B | 4281.650139 | 0.708419122 | 0.111795449 | 1.32E-11 | 5.17E-10 |
| HERPUD2 | 823.6787071 | 0.70773831 | 0.125107174 | 8.37E-10 | 2.28E-08 |
| ERICH2-DT | 11.773628 | 0.70710792 | 0.570382287 | 0.00433259 | 0.017858358 |
| ATOSB | 304.7450864 | 0.707015425 | 0.209338933 | 3.03E-05 | 0.000267138 |
| PDGFA | 12.51805873 | 0.706829302 | 0.495798455 | 0.00355344 | 0.015172969 |
| TAPT1 | 1493.076765 | 0.706804596 | 0.08996639 | 2.42E-16 | 1.93E-14 |
| TMED2 | 10086.88341 | 0.706163523 | 0.09211057 | 1.08E-15 | 7.78E-14 |
| TM6SF1 | 65.25575723 | 0.705793694 | 0.257044509 | 0.00022085 | 0.001459862 |
| SLC9A1 | 1182.787994 | 0.705245636 | 0.228970147 | 8.21E-05 | 0.000623881 |
| ARMCX1 | 341.5433468 | 0.704917679 | 0.205788316 | 2.62E-05 | 0.000235658 |
| ERAP1 | 739.8295918 | 0.703578677 | 0.122720129 | 5.64E-10 | 1.59E-08 |
| PITPNM1 | 2547.894312 | 0.703435559 | 0.224477268 | 7.05E-05 | 0.000548303 |
| LIPG | 57.31678684 | 0.702511438 | 0.265057296 | 0.00029592 | 0.001866107 |
| CASC18 | 5.343179962 | 0.70245912 | 0.755818716 | 0.0048387 | 0.019575639 |
| EDN1 | 24.90102303 | 0.70237973 | 0.348485936 | 0.00134616 | 0.006761683 |
| CADM2 | 209.2914471 | 0.702039212 | 0.249341667 | 0.00018749 | 0.001270367 |
| RAPGEF2 | 2799.818633 | 0.701417265 | 0.113174798 | 3.46E-11 | 1.25E-09 |
| IRF2BPL | 1230.109094 | 0.701129781 | 0.221499811 | 6.51E-05 | 0.000511513 |
| UGGT2 | 1761.136457 | 0.700455733 | 0.111159272 | 1.81E-11 | 6.93E-10 |
| TMED7 | 2657.726765 | 0.699986246 | 0.093852805 | 5.69E-15 | 3.70E-13 |
| SNX31 | 4.24732913 | 0.699782418 | 0.83979537 | 0.00441817 | 0.01813839 |
| PLD5P1 | 104.131096 | 0.699489483 | 0.202576454 | 2.51E-05 | 0.000227715 |

| | | | | | |
|---|---|---|---|---|---|
| SBDS | 2273.504432 | 0.699433956 | 0.087071326 | 6.34E-17 | 5.54E-15 |
| PLCD4 | 43.78412527 | 0.699133601 | 0.295134955 | 0.00062669 | 0.003540425 |
| PACSIN1 | 122.5116334 | 0.698885854 | 0.285031169 | 0.00051201 | 0.002986964 |
| FRMD5 | 1304.949291 | 0.69885144 | 0.099343441 | 1.29E-13 | 6.88E-12 |
| UBE2QL1 | 1029.704761 | 0.698807811 | 0.158409053 | 5.45E-07 | 7.88E-06 |
| DUSP14 | 893.2446481 | 0.69866066 | 0.129721458 | 4.23E-09 | 1.00E-07 |
| GFPT1 | 5742.255506 | 0.698482945 | 0.079792594 | 1.41E-19 | 1.72E-17 |
| KLHL28 | 1562.998634 | 0.698177642 | 0.141531284 | 4.59E-08 | 8.79E-07 |
| GRM3-AS1 | 8.533028605 | 0.695698621 | 0.593380019 | 0.00485326 | 0.01962296 |
| MTM1 | 211.3414338 | 0.695346143 | 0.138908719 | 3.27E-08 | 6.46E-07 |
| EFNA1 | 107.0877915 | 0.695100118 | 0.233689213 | 0.00012529 | 0.000899298 |
| ENO3 | 1232.525138 | 0.695078143 | 0.175936746 | 4.03E-06 | 4.54E-05 |
| SPACA6 | 295.36099 | 0.69470478 | 0.197339222 | 2.08E-05 | 0.000192753 |
| KCTD16 | 1014.302986 | 0.694552191 | 0.176028746 | 4.13E-06 | 4.63E-05 |
| AGL | 2827.748338 | 0.694270572 | 0.111671077 | 3.30E-11 | 1.20E-09 |
| USP51 | 20.08768015 | 0.693185581 | 0.435286853 | 0.00306472 | 0.013446339 |
| RBM5-AS1 | 56.33951294 | 0.693075592 | 0.28490282 | 0.00056451 | 0.003235569 |
| ZNF396 | 47.72802076 | 0.693072008 | 0.290903113 | 0.00063811 | 0.003598054 |
| GNAT2 | 72.33289962 | 0.692122325 | 0.229966465 | 0.00011583 | 0.000839579 |
| CLK1 | 5338.668255 | 0.691764522 | 0.152789466 | 3.46E-07 | 5.25E-06 |
| KLHL11 | 972.6786465 | 0.691662236 | 0.120184405 | 5.64E-10 | 1.59E-08 |
| MFAP3L | 724.7857035 | 0.691456275 | 0.128200815 | 4.38E-09 | 1.03E-07 |
| ITGA1 | 1735.473102 | 0.691260494 | 0.129672582 | 6.18E-09 | 1.41E-07 |
| ABTB3 | 1145.133966 | 0.690889771 | 0.425962804 | 0.00297424 | 0.013113675 |
| FBXL20 | 1670.094718 | 0.690641218 | 0.11051693 | 2.80E-11 | 1.03E-09 |
| NPHP3 | 477.3267849 | 0.690054017 | 0.14081613 | 5.89E-08 | 1.10E-06 |
| KCNJ4 | 91.15139052 | 0.689855142 | 0.585917101 | 0.00499674 | 0.020080714 |
| NALCN | 96.50171752 | 0.689411596 | 0.219461806 | 7.88E-05 | 0.000602016 |
| RNF19A | 1543.445801 | 0.688572609 | 0.120126959 | 6.66E-10 | 1.85E-08 |
| AVIL | 360.3194507 | 0.687874865 | 0.151696453 | 3.49E-07 | 5.29E-06 |
| LINC01352 | 114.4962949 | 0.687817001 | 0.186353023 | 1.19E-05 | 0.000118373 |
| HIVEP1 | 1260.992904 | 0.687259354 | 0.162298092 | 1.34E-06 | 1.73E-05 |
| SEC61A1 | 11719.60945 | 0.687114296 | 0.107732183 | 1.27E-11 | 4.99E-10 |
| ATP1B2 | 63.69002367 | 0.686402345 | 0.322831938 | 0.0012075 | 0.006181474 |
| ADAMTSL4-AS2 | 16.84563743 | 0.686022556 | 0.423643611 | 0.00308374 | 0.013521125 |
| FAM199X | 2491.489783 | 0.685841722 | 0.111610245 | 5.67E-11 | 1.97E-09 |
| PLCXD2 | 965.6870272 | 0.685737185 | 0.108767684 | 2.07E-11 | 7.81E-10 |
| NPIPB6 | 151.9161551 | 0.685662892 | 0.200065299 | 3.16E-05 | 0.000276514 |
| KDM5B | 5864.651097 | 0.685535765 | 0.098454936 | 2.47E-13 | 1.28E-11 |
| ARHGEF2-AS1 | 5.998226712 | 0.685348681 | 0.73352534 | 0.00531322 | 0.02110103 |
| CEBPB-AS1 | 56.5900799 | 0.685266561 | 0.284583936 | 0.00064122 | 0.003612627 |
| SYNPO2L-AS1 | 34.65230964 | 0.685210952 | 0.359110896 | 0.0018902 | 0.009022741 |
| SMG1P2 | 147.5784945 | 0.68506497 | 0.263807044 | 0.00039892 | 0.002417025 |
| LY75 | 4.444353192 | 0.684941253 | 0.849589283 | 0.00455507 | 0.018596457 |
| DUSP16 | 2312.186942 | 0.684285222 | 0.103644023 | 2.99E-12 | 1.32E-10 |
| CYP51A1 | 512.624667 | 0.684273328 | 0.118835268 | 5.98E-10 | 1.68E-08 |
| RNFT1 | 601.0105369 | 0.683879196 | 0.108817564 | 2.40E-11 | 8.93E-10 |
| ZNF267 | 333.4679498 | 0.682943476 | 0.131293464 | 1.35E-08 | 2.89E-07 |
| UBE2H | 2641.003601 | 0.682174069 | 0.112847235 | 1.09E-10 | 3.57E-09 |
| MYORG | 193.5960946 | 0.682119161 | 0.214059086 | 7.32E-05 | 0.000565487 |

| | | | | | |
|---|---|---|---|---|---|
| SNCAIP | 154.6485419 | 0.681884589 | 0.182975021 | 1.11E-05 | 0.000110804 |
| TAB2 | 5272.878131 | 0.681704492 | 0.090071311 | 2.98E-15 | 2.00E-13 |
| DPP8 | 5410.78058 | 0.68086919 | 0.087967778 | 7.95E-16 | 5.76E-14 |
| STT3A | 4460.644036 | 0.679765679 | 0.116941851 | 4.54E-10 | 1.31E-08 |
| CDYL2 | 545.5986698 | 0.67933113 | 0.136299105 | 4.32E-08 | 8.35E-07 |
| QPCT | 2.39871876 | 0.679294105 | 0.996383098 | 0.00291389 | 0.012883556 |
| APRG1 | 87.8307923 | 0.679070954 | 0.208426587 | 5.98E-05 | 0.000474311 |
| LRRC49 | 1658.73462 | 0.679048707 | 0.098024898 | 3.39E-13 | 1.71E-11 |
| DACT3-AS1 | 44.59742983 | 0.679002818 | 0.338381803 | 0.00163676 | 0.007962717 |
| THSD7A | 674.7659158 | 0.678573708 | 0.177954213 | 8.21E-06 | 8.49E-05 |
| IQSEC3-AS1 | 19.56346594 | 0.678475857 | 0.434202864 | 0.0035248 | 0.015066293 |
| LINC02901 | 11.53893387 | 0.67812163 | 0.551817525 | 0.00514873 | 0.020554952 |
| ZNF610 | 776.8770679 | 0.67811509 | 0.112543073 | 1.29E-10 | 4.13E-09 |
| ZNF622 | 1258.632384 | 0.677753624 | 0.112207749 | 1.18E-10 | 3.81E-09 |
| SLC22A20P | 22.78367488 | 0.67753915 | 0.460719626 | 0.0040157 | 0.016777511 |
| BTBD19 | 101.6753684 | 0.677381224 | 0.198658691 | 3.64E-05 | 0.000311538 |
| IL11 | 393.1038634 | 0.676833588 | 0.207672832 | 6.09E-05 | 0.000482051 |
| S100A13 | 72.26290512 | 0.676750171 | 0.244487135 | 0.00027104 | 0.001736607 |
| SLC26A11 | 1419.664516 | 0.676600203 | 0.163237888 | 2.19E-06 | 2.64E-05 |
| FBXO34 | 1785.180924 | 0.676575741 | 0.092391451 | 2.00E-14 | 1.20E-12 |
| SLC51A | 113.2223383 | 0.676382379 | 0.203630086 | 4.97E-05 | 0.000405936 |
| MYO5B | 100.9453787 | 0.676273739 | 0.199539699 | 3.95E-05 | 0.000334373 |
| CCDC62 | 136.6181207 | 0.675841205 | 0.235178675 | 0.00020252 | 0.001353496 |
| PUS10 | 257.9416831 | 0.674896599 | 0.144877098 | 2.23E-07 | 3.59E-06 |
| P2RX4 | 357.9617908 | 0.673761416 | 0.130868274 | 1.96E-08 | 4.05E-07 |
| EVX2 | 4.793324562 | 0.673093981 | 0.853208357 | 0.00468824 | 0.019068229 |
| BMT2 | 968.2111527 | 0.672593154 | 0.134868423 | 4.56E-08 | 8.75E-07 |
| EHBP1 | 2046.255529 | 0.672457184 | 0.084749537 | 1.85E-16 | 1.49E-14 |
| HHIP | 541.5114983 | 0.672307004 | 0.123143656 | 3.71E-09 | 8.90E-08 |
| COG1 | 2855.622729 | 0.671638111 | 0.119227031 | 1.40E-09 | 3.64E-08 |
| SLC30A7 | 3416.422172 | 0.67100814 | 0.109467642 | 7.25E-11 | 2.44E-09 |
| SPRED3 | 165.9937091 | 0.670744557 | 0.274661071 | 0.00066957 | 0.003748698 |
| RIPK2 | 400.8831991 | 0.670305461 | 0.13005768 | 1.97E-08 | 4.07E-07 |
| PDCD2L | 252.5622174 | 0.67002475 | 0.153291683 | 8.79E-07 | 1.19E-05 |
| LINC00963 | 577.0341735 | 0.669905274 | 0.233657009 | 0.00021835 | 0.001445146 |
| DEDD2 | 1046.258549 | 0.66972042 | 0.147861043 | 4.30E-07 | 6.42E-06 |
| MACROD2 | 203.1652362 | 0.669536906 | 0.206670072 | 6.97E-05 | 0.000543166 |
| SYN3 | 100.1311365 | 0.669014742 | 0.235578232 | 0.00023799 | 0.001556428 |
| USO1 | 4182.393051 | 0.668570729 | 0.107413728 | 4.10E-11 | 1.47E-09 |
| PPP1R2 | 1752.252362 | 0.668374551 | 0.105503922 | 2.03E-11 | 7.67E-10 |
| BET1 | 589.7611883 | 0.668084473 | 0.133502973 | 4.37E-08 | 8.43E-07 |
| RENBP | 132.7129909 | 0.667889559 | 0.196492206 | 4.15E-05 | 0.000349643 |
| ZNF862 | 280.5415311 | 0.667583126 | 0.170974733 | 6.41E-06 | 6.82E-05 |
| ETV5-AS1 | 8.492797496 | 0.667374181 | 0.66299339 | 0.00598209 | 0.023276383 |
| IQUB | 20.78223195 | 0.667079371 | 0.41831754 | 0.00363324 | 0.015446329 |
| CYLD | 2116.162433 | 0.667006412 | 0.098020675 | 8.97E-13 | 4.21E-11 |
| CCDC66 | 1873.016225 | 0.666952698 | 0.128179669 | 1.57E-08 | 3.31E-07 |
| HRK | 3574.649359 | 0.66692281 | 0.158126793 | 1.77E-06 | 2.21E-05 |
| EGR2 | 9.428388499 | 0.666830028 | 0.776302382 | 0.00550714 | 0.021710289 |
| ZNF211 | 788.8364956 | 0.666139818 | 0.134304482 | 5.58E-08 | 1.05E-06 |

| | | | | | |
|---|---|---|---|---|---|
| HECA | 955.8097311 | 0.664540955 | 0.115294322 | 7.05E-10 | 1.95E-08 |
| TRPC1 | 1101.163358 | 0.664471874 | 0.150097354 | 7.23E-07 | 9.96E-06 |
| DERL2 | 2589.948673 | 0.664168718 | 0.087944228 | 4.01E-15 | 2.67E-13 |
| TRMT1L | 1598.238571 | 0.66407225 | 0.101226336 | 4.84E-12 | 2.04E-10 |
| NHLH1 | 1541.576663 | 0.663682376 | 0.156100226 | 1.58E-06 | 2.00E-05 |
| KSR2 | 60.4188606 | 0.663562982 | 0.264747037 | 0.00060913 | 0.003453602 |
| PAK6-AS1 | 7.135703223 | 0.663294054 | 0.623647519 | 0.00609529 | 0.023631122 |
| ARRDC3 | 4083.312334 | 0.663179645 | 0.115119772 | 7.28E-10 | 2.00E-08 |
| SND1 | 10169.45579 | 0.663146212 | 0.140823806 | 1.97E-07 | 3.23E-06 |
| DNAJC6 | 5311.714186 | 0.663069884 | 0.095456989 | 3.47E-13 | 1.75E-11 |
| COL9A1 | 12.69855323 | 0.662584742 | 0.530349821 | 0.00555433 | 0.021851772 |
| PAIP2B | 402.7190755 | 0.662528094 | 0.148479479 | 6.29E-07 | 8.90E-06 |
| HMOX1 | 38.23154684 | 0.662508088 | 0.326569776 | 0.00178358 | 0.008561492 |
| ORC5 | 1184.769499 | 0.662253397 | 0.095405281 | 3.62E-13 | 1.82E-11 |
| TRIM61 | 6.610057782 | 0.662199582 | 0.647832533 | 0.00616052 | 0.023857815 |
| RASGRF2 | 1022.068084 | 0.661991857 | 0.152359433 | 1.07E-06 | 1.42E-05 |
| ITGA2 | 120.572903 | 0.661603417 | 0.166805467 | 5.31E-06 | 5.79E-05 |
| RBPMS2 | 85.31478155 | 0.661378254 | 0.335927552 | 0.00202051 | 0.009529756 |
| CARMIL1 | 446.1126359 | 0.660632355 | 0.128637083 | 2.39E-08 | 4.84E-07 |
| TMEM200A | 125.4835533 | 0.660548255 | 0.204628172 | 7.90E-05 | 0.000603087 |
| SLC9A6 | 1144.179965 | 0.660341266 | 0.089023105 | 1.15E-14 | 7.07E-13 |
| RAB23 | 838.8597132 | 0.660269313 | 0.112123215 | 3.51E-10 | 1.04E-08 |
| IFI27 | 200.9963569 | 0.660004468 | 0.183645371 | 2.25E-05 | 0.00020632 |
| MGAT2 | 74.62071802 | 0.659855454 | 0.201137975 | 6.69E-05 | 0.00052403 |
| JMJD1C | 4205.600516 | 0.659602583 | 0.127966961 | 2.18E-08 | 4.46E-07 |
| ALAS1 | 1825.944681 | 0.659339894 | 0.169413055 | 7.31E-06 | 7.66E-05 |
| FREM1 | 36.28634846 | 0.659328442 | 0.293251549 | 0.00115783 | 0.00596132 |
| ELAPOR2 | 914.2541423 | 0.658924466 | 0.136071178 | 1.08E-07 | 1.89E-06 |
| TTC8 | 2104.57338 | 0.658438631 | 0.14200029 | 2.92E-07 | 4.53E-06 |
| USP44 | 357.0338549 | 0.65837279 | 0.128294545 | 2.49E-08 | 5.03E-07 |
| C10orf143 | 55.17102402 | 0.658285983 | 0.263207036 | 0.00064716 | 0.003641067 |
| ZFP69B | 739.0502153 | 0.658054038 | 0.117402712 | 1.88E-09 | 4.76E-08 |
| SERTAD3 | 278.4260081 | 0.657767389 | 0.164104645 | 4.66E-06 | 5.15E-05 |
| SLC4A5 | 217.0739657 | 0.657765081 | 0.241124634 | 0.00036093 | 0.002220875 |
| MAFF | 129.943054 | 0.657375124 | 0.314285085 | 0.00163931 | 0.0079732 |
| CNIH3 | 143.1525518 | 0.656401367 | 0.165810338 | 5.77E-06 | 6.21E-05 |
| AZIN2 | 78.46791274 | 0.656283076 | 0.220882582 | 0.00018316 | 0.001247587 |
| EDN2 | 23.6094627 | 0.655723298 | 0.417343891 | 0.00405972 | 0.01692701 |
| FAM153A | 4.139281151 | 0.655537546 | 0.895839055 | 0.00437917 | 0.018007043 |
| TMEM267 | 508.5913546 | 0.655353516 | 0.113552253 | 7.37E-10 | 2.02E-08 |
| ZNF365 | 138.6319215 | 0.655125831 | 0.177365052 | 1.63E-05 | 0.000156604 |
| LMX1B | 2.321678071 | 0.655032087 | 0.983347896 | 0.00312511 | 0.013665243 |
| PURA | 1662.859892 | 0.654772406 | 0.116625527 | 1.84E-09 | 4.66E-08 |
| PEG3 | 1904.199746 | 0.654196597 | 0.115466801 | 1.38E-09 | 3.59E-08 |
| IRX5 | 338.4614738 | 0.654061892 | 0.172798219 | 1.17E-05 | 0.000116373 |
| MIR155HG | 18.18303312 | 0.652822728 | 0.370582442 | 0.00311033 | 0.013626072 |
| CCDC18 | 1677.231919 | 0.652474084 | 0.111826414 | 5.21E-10 | 1.48E-08 |
| GGT1 | 30.49579155 | 0.652385631 | 0.327275459 | 0.0020707 | 0.009735206 |
| TBCK | 1697.737166 | 0.652134481 | 0.112416477 | 6.38E-10 | 1.79E-08 |
| LINC01666 | 7.923287349 | 0.651988405 | 0.568236463 | 0.00631957 | 0.024381904 |

| | | | | | |
|---|---|---|---|---|---|
| SPATA2L | 162.0557263 | 0.651983938 | 0.243807709 | 0.00043978 | 0.002628202 |
| HELZ2 | 1051.874292 | 0.651485753 | 0.195590403 | 6.14E-05 | 0.000485838 |
| LINC02774 | 39.41933571 | 0.650677825 | 0.284358867 | 0.00114139 | 0.005889986 |
| IQCH | 122.3825065 | 0.650109972 | 0.178136527 | 2.02E-05 | 0.000189034 |
| POPDC2 | 175.7434649 | 0.650071878 | 0.152912536 | 1.81E-06 | 2.24E-05 |
| CASP9 | 192.7991845 | 0.649437578 | 0.181136848 | 2.57E-05 | 0.000232326 |
| SEC23B | 4435.126571 | 0.649249896 | 0.07923343 | 2.76E-17 | 2.57E-15 |
| CACNA1I | 140.3021901 | 0.649125358 | 0.257245832 | 0.00066485 | 0.003727353 |
| MSS51 | 291.0743891 | 0.649122871 | 0.13797435 | 2.31E-07 | 3.71E-06 |
| LRP1 | 7248.408375 | 0.648942884 | 0.181815806 | 2.74E-05 | 0.000244986 |
| EMC9 | 728.2190947 | 0.647815551 | 0.120353704 | 7.19E-09 | 1.63E-07 |
| LINC00629 | 17.92038741 | 0.647397938 | 0.382041983 | 0.0035991 | 0.015320179 |
| TRIM11 | 2684.18107 | 0.6473094 | 0.152547191 | 1.92E-06 | 2.36E-05 |
| PLEKHA6 | 1113.774276 | 0.647011513 | 0.230331472 | 0.00031905 | 0.001994793 |
| TTTY14 | 348.9665698 | 0.646825291 | 0.134753414 | 1.49E-07 | 2.51E-06 |
| NEDD9 | 1510.687729 | 0.646716281 | 0.132409019 | 9.84E-08 | 1.75E-06 |
| SYTL3 | 337.9080956 | 0.646075847 | 0.147991854 | 1.14E-06 | 1.50E-05 |
| ZFAND1 | 2176.295716 | 0.646037362 | 0.113871923 | 1.42E-09 | 3.69E-08 |
| SLC17A6-DT | 5.080990157 | 0.6458223 | 0.72003361 | 0.00645399 | 0.024801274 |
| STK40 | 2832.512952 | 0.64578416 | 0.173570312 | 1.62E-05 | 0.000155101 |
| LINC01550 | 58.79513559 | 0.64508512 | 0.248262576 | 0.00057077 | 0.003263107 |
| PPP1R15B | 8776.241717 | 0.644817078 | 0.090297839 | 1.02E-13 | 5.57E-12 |
| ESCO1 | 1344.206258 | 0.644348977 | 0.132479168 | 1.11E-07 | 1.95E-06 |
| HRC | 16.21075048 | 0.644231437 | 0.46219586 | 0.00544966 | 0.021543019 |
| SEC24A | 2958.239729 | 0.643584617 | 0.097976853 | 5.54E-12 | 2.32E-10 |
| LRRC37B | 861.1293697 | 0.643416991 | 0.132828094 | 1.24E-07 | 2.14E-06 |
| KLF15 | 583.484823 | 0.643341083 | 0.181934105 | 3.26E-05 | 0.000284072 |
| ELFN1-AS1 | 2.139810301 | 0.643135298 | 0.958807074 | 0.00351695 | 0.015048393 |
| C2orf50 | 22.75861531 | 0.642747139 | 0.374341962 | 0.00359849 | 0.015320179 |
| TRIT1 | 1645.56144 | 0.642241064 | 0.113959268 | 1.83E-09 | 4.65E-08 |
| YIPF7 | 45.99714156 | 0.642089004 | 0.260992247 | 0.00082656 | 0.004461256 |
| TMED10 | 7181.180613 | 0.641846414 | 0.077044344 | 9.41E-18 | 9.39E-16 |
| KDM7A | 1634.445974 | 0.641830455 | 0.121475711 | 1.30E-08 | 2.80E-07 |
| PROX2 | 71.18848367 | 0.64171626 | 0.255399533 | 0.00073002 | 0.00402724 |
| MATN1-AS1 | 8.120135862 | 0.641675334 | 0.602762921 | 0.0069069 | 0.026171657 |
| GSX2 | 19.6060574 | 0.641662159 | 0.381069734 | 0.00381554 | 0.016093826 |
| CRTC2 | 2506.86963 | 0.641329575 | 0.158111567 | 4.49E-06 | 4.98E-05 |
| KATNA1 | 1021.289319 | 0.641064712 | 0.097166237 | 4.67E-12 | 1.98E-10 |
| ODR4 | 2847.067572 | 0.640623416 | 0.099571169 | 1.39E-11 | 5.40E-10 |
| UEVLD | 548.6787485 | 0.639990649 | 0.115211379 | 2.96E-09 | 7.23E-08 |
| LRRC74B | 22.65037348 | 0.639950197 | 0.415891182 | 0.00473102 | 0.019203628 |
| AGK-DT | 18.88803418 | 0.639482879 | 0.39734972 | 0.00431772 | 0.017819733 |
| CIR1 | 1370.091691 | 0.639433717 | 0.108297667 | 3.88E-10 | 1.14E-08 |
| DYRK3 | 1430.346336 | 0.639369194 | 0.148613321 | 1.60E-06 | 2.03E-05 |
| ASB5 | 6.767764419 | 0.638751894 | 0.59932619 | 0.00702621 | 0.026525961 |
| CLIC4 | 11230.21568 | 0.63833193 | 0.083220125 | 2.04E-15 | 1.41E-13 |
| LINC02986 | 303.9654509 | 0.638062833 | 0.163291199 | 8.44E-06 | 8.68E-05 |
| LMBRD1 | 990.1248958 | 0.637727462 | 0.131063665 | 1.17E-07 | 2.04E-06 |
| CLBA1 | 357.0671543 | 0.637110137 | 0.15514628 | 3.79E-06 | 4.29E-05 |
| RCOR1 | 4231.929214 | 0.636962424 | 0.114620792 | 3.01E-09 | 7.32E-08 |

| | | | | | |
|---|---|---|---|---|---|
| FAM149A | 585.2343073 | 0.636521839 | 0.196656322 | 9.67E-05 | 0.000717466 |
| TUBA3FP | 114.5105069 | 0.636393294 | 0.238830433 | 0.0005206 | 0.003027001 |
| RBM15 | 1950.18103 | 0.636387763 | 0.102637105 | 6.45E-11 | 2.21E-09 |
| RUFY1 | 1679.669634 | 0.635937345 | 0.102565183 | 6.47E-11 | 2.21E-09 |
| CCDC110 | 15.82599241 | 0.635884372 | 0.41764218 | 0.00496931 | 0.020009572 |
| LINC02970 | 2.40114875 | 0.63557321 | 0.971358253 | 0.00332606 | 0.014339008 |
| ASL | 528.8563948 | 0.635028387 | 0.160108668 | 6.87E-06 | 7.24E-05 |
| DLGAP1-AS1 | 30.47631319 | 0.634901823 | 0.364962537 | 0.00368543 | 0.015613325 |
| EIF1B | 2019.672975 | 0.634888023 | 0.113745978 | 2.67E-09 | 6.58E-08 |
| SARAF | 7209.401036 | 0.634372533 | 0.079130921 | 1.35E-16 | 1.13E-14 |
| RASSF1 | 1198.221195 | 0.634239961 | 0.107839948 | 4.67E-10 | 1.34E-08 |
| LINC00663 | 71.51661083 | 0.634212978 | 0.246296985 | 0.00066891 | 0.003746015 |
| SPMAP2L | 5.730198678 | 0.633775445 | 0.640663787 | 0.00723679 | 0.02720112 |
| AKAP8L | 2628.405382 | 0.633443095 | 0.143093785 | 9.75E-07 | 1.31E-05 |
| IDI1 | 5619.336854 | 0.63341691 | 0.107291388 | 4.11E-10 | 1.20E-08 |
| KIF9-AS1 | 153.2892824 | 0.633317813 | 0.216974559 | 0.00026526 | 0.001707208 |
| FRG1DP | 140.1644834 | 0.633297258 | 0.188365175 | 6.58E-05 | 0.000515918 |
| ALKBH3 | 714.7710834 | 0.633034507 | 0.113826504 | 3.04E-09 | 7.39E-08 |
| FBXW4 | 1009.127864 | 0.632507881 | 0.153205518 | 3.61E-06 | 4.10E-05 |
| SOCS5 | 1694.716871 | 0.632372165 | 0.096033204 | 5.49E-12 | 2.30E-10 |
| TSTD3 | 342.5392956 | 0.632043144 | 0.14640399 | 1.61E-06 | 2.03E-05 |
| PHGDH | 14334.59694 | 0.631903011 | 0.115258737 | 4.79E-09 | 1.12E-07 |
| DDX59-AS1 | 4.140284721 | 0.631843777 | 0.793552773 | 0.00609624 | 0.023631122 |
| ZNF385C | 23.25704294 | 0.631791668 | 0.382164226 | 0.00429323 | 0.017745931 |
| ISL1-DT | 1213.074632 | 0.631646817 | 0.124545444 | 4.37E-08 | 8.43E-07 |
| BTBD10 | 1309.117428 | 0.630930348 | 0.10337509 | 1.24E-10 | 4.01E-09 |
| LINC02795 | 46.92127181 | 0.630925979 | 0.283118761 | 0.00154244 | 0.00756288 |
| LACTB | 485.2992046 | 0.630652526 | 0.130746753 | 1.54E-07 | 2.59E-06 |
| BTLA | 4.808954475 | 0.630342762 | 0.773803112 | 0.00637283 | 0.02455054 |
| CLCF1 | 23.12932404 | 0.62981967 | 0.391656708 | 0.00464002 | 0.018898287 |
| GZF1 | 1998.206856 | 0.62899375 | 0.096791596 | 1.00E-11 | 4.00E-10 |
| LINC00222 | 4.29016513 | 0.628839586 | 0.79531916 | 0.00613719 | 0.023771929 |
| EFCAB2 | 1033.663826 | 0.628083133 | 0.13860825 | 6.35E-07 | 8.97E-06 |
| DHX29 | 2893.201144 | 0.627838441 | 0.088084014 | 1.31E-13 | 6.94E-12 |
| GUCY1B2 | 27.64547771 | 0.627735549 | 0.348790952 | 0.00354063 | 0.015124536 |
| SLFNL1 | 130.5781881 | 0.62759543 | 0.187030001 | 7.09E-05 | 0.0005503 |
| SREBF1 | 2555.330639 | 0.627160181 | 0.222635698 | 0.0003754 | 0.002298931 |
| SPTY2D1 | 1584.575085 | 0.627045615 | 0.093954503 | 3.16E-12 | 1.38E-10 |
| SLC1A5 | 5201.736776 | 0.626763215 | 0.153762023 | 4.73E-06 | 5.22E-05 |
| RIMS1 | 266.2997636 | 0.626437671 | 0.13303577 | 2.79E-07 | 4.36E-06 |
| TJP1 | 3093.427495 | 0.626359364 | 0.087037137 | 8.03E-14 | 4.45E-12 |
| KIAA0753 | 1707.876851 | 0.626245474 | 0.099109178 | 3.32E-11 | 1.21E-09 |
| RPL7AP45 | 61.28642271 | 0.625787603 | 0.226898052 | 0.00044726 | 0.002663431 |
| RMDN3 | 3059.065447 | 0.625450779 | 0.130870853 | 2.01E-07 | 3.28E-06 |
| CALCA | 4.154609545 | 0.625332987 | 0.79412758 | 0.00622767 | 0.024095176 |
| KIAA0513 | 159.4278967 | 0.625185306 | 0.226659003 | 0.00044945 | 0.002673546 |
| LGALS1 | 114.7847176 | 0.624827394 | 0.25450912 | 0.00097713 | 0.005162797 |
| ADGRA1 | 25.79841868 | 0.624775077 | 0.378867306 | 0.00455052 | 0.01858624 |
| DNAL4 | 511.8696168 | 0.624604411 | 0.12844126 | 1.34E-07 | 2.29E-06 |
| SIDT2 | 622.6539131 | 0.624370402 | 0.159327512 | 9.17E-06 | 9.33E-05 |

| | | | | | |
|---|---|---|---|---|---|
| CHORDC1 | 3234.169671 | 0.624341695 | 0.123839122 | 5.44E-08 | 1.03E-06 |
| HSPA4L | 1759.46929 | 0.623628373 | 0.106823454 | 6.63E-10 | 1.84E-08 |
| WDPCP | 534.829391 | 0.623583787 | 0.110244678 | 1.92E-09 | 4.84E-08 |
| XBP1P1 | 16.84703182 | 0.623151687 | 0.395643167 | 0.00509195 | 0.020395565 |
| TOM1L2 | 1390.199385 | 0.623098618 | 0.164495043 | 1.55E-05 | 0.000149337 |
| HES1 | 41.87905854 | 0.622256835 | 0.306332905 | 0.00246742 | 0.011214621 |
| ZNF823 | 503.3412922 | 0.622222981 | 0.146121649 | 2.28E-06 | 2.74E-05 |
| BEND7 | 903.9385467 | 0.621878138 | 0.127610318 | 1.30E-07 | 2.23E-06 |
| ZNF790 | 538.3988403 | 0.621865372 | 0.140777678 | 1.13E-06 | 1.49E-05 |
| MAP2K5-DT | 47.37218121 | 0.621254246 | 0.264721665 | 0.00129595 | 0.006555927 |
| GGCX | 1703.380004 | 0.621063696 | 0.125507559 | 9.01E-08 | 1.62E-06 |
| WDR45 | 781.1033334 | 0.620614201 | 0.106346446 | 6.89E-10 | 1.91E-08 |
| MAGT1 | 2400.795509 | 0.6206116 | 0.107512099 | 1.00E-09 | 2.68E-08 |
| MOK | 279.9068777 | 0.620384172 | 0.176536745 | 4.37E-05 | 0.000365815 |
| PTCH2 | 250.236676 | 0.619983879 | 0.179500483 | 5.43E-05 | 0.0004366 |
| PHLDB3 | 125.6202698 | 0.619945057 | 0.261501395 | 0.0012421 | 0.006329253 |
| GRPEL2 | 2232.704473 | 0.619834278 | 0.089748975 | 6.76E-13 | 3.23E-11 |
| IRAG1 | 68.06732245 | 0.619805517 | 0.223450964 | 0.0004527 | 0.00269049 |
| DHX58 | 13.55334695 | 0.619804022 | 0.451171235 | 0.00655416 | 0.02509863 |
| PATJ | 497.6790849 | 0.619677916 | 0.146280448 | 2.56E-06 | 3.04E-05 |
| NBAT1 | 102.4207792 | 0.619588293 | 0.214159451 | 0.00032462 | 0.002023898 |
| INMT | 3.999086166 | 0.619436528 | 0.784900601 | 0.00648511 | 0.024899151 |
| MDM1 | 2049.885526 | 0.619095954 | 0.096376195 | 1.78E-11 | 6.85E-10 |
| ACSBG1 | 23.92968717 | 0.618563409 | 0.358104034 | 0.00425754 | 0.017612555 |
| PRSS46P | 6.115770266 | 0.618160883 | 0.697171404 | 0.00753094 | 0.028116604 |
| ZCCHC7 | 1083.727259 | 0.618110313 | 0.102947358 | 2.55E-10 | 7.71E-09 |
| EFNA3 | 24.9898678 | 0.617958658 | 0.32872891 | 0.00335267 | 0.014438576 |
| XRN1 | 3296.563704 | 0.617816397 | 0.127478703 | 1.54E-07 | 2.59E-06 |
| DOK6 | 1175.101745 | 0.617761798 | 0.101533584 | 1.56E-10 | 4.93E-09 |
| ZNF627 | 1493.697662 | 0.617681894 | 0.122751935 | 6.06E-08 | 1.13E-06 |
| ZSWIM3 | 371.1860316 | 0.617355174 | 0.133410111 | 4.46E-07 | 6.63E-06 |
| DENND4A | 2397.348984 | 0.616866129 | 0.108501651 | 1.72E-09 | 4.38E-08 |
| REM2 | 5.775589985 | 0.616759633 | 0.628338325 | 0.00800108 | 0.029576522 |
| PTPRT | 44.9691743 | 0.61675284 | 0.287774785 | 0.00207983 | 0.009773673 |
| TSHZ3 | 1250.7896 | 0.616710425 | 0.226064353 | 0.00052724 | 0.003059312 |
| BUD31 | 1272.092214 | 0.615666348 | 0.096943072 | 2.95E-11 | 1.08E-09 |
| ITGAV | 4748.849159 | 0.615653451 | 0.090881919 | 1.75E-12 | 7.94E-11 |
| RETREG1 | 260.2697602 | 0.615472924 | 0.158862762 | 1.19E-05 | 0.000117974 |
| SLC38A3 | 103.1349947 | 0.614896696 | 0.236040033 | 0.00073919 | 0.004075261 |
| LTA | 4.459273377 | 0.614695694 | 0.735088082 | 0.00725637 | 0.02724981 |
| ZNF641 | 1477.29089 | 0.614560182 | 0.120863292 | 4.74E-08 | 9.04E-07 |
| LINC02798 | 7.037681211 | 0.614241333 | 0.603309127 | 0.0081718 | 0.030083225 |
| GCC1 | 1344.249692 | 0.614087545 | 0.11586102 | 1.52E-08 | 3.22E-07 |
| TMED9 | 2340.288426 | 0.6139708 | 0.095004281 | 1.44E-11 | 5.60E-10 |
| ABCD1 | 278.9067 | 0.61396698 | 0.232867602 | 0.00068773 | 0.003832519 |
| RINT1 | 1278.093694 | 0.613279008 | 0.112406319 | 6.52E-09 | 1.49E-07 |
| PRMT9 | 869.371974 | 0.613195308 | 0.101236467 | 1.92E-10 | 5.95E-09 |
| PCCA | 1084.001575 | 0.613172695 | 0.107326271 | 1.51E-09 | 3.88E-08 |
| HACL1 | 631.3478696 | 0.612923134 | 0.118461783 | 3.01E-08 | 6.02E-07 |
| BCL10 | 901.1841651 | 0.612221407 | 0.097334421 | 4.48E-11 | 1.58E-09 |

| | | | | | |
|---|---|---|---|---|---|
| METTL22 | 1543.963619 | 0.612170426 | 0.11516861 | 1.42E-08 | 3.03E-07 |
| AP4E1 | 2935.171945 | 0.612152534 | 0.091516853 | 3.23E-12 | 1.41E-10 |
| KIF6 | 59.28209824 | 0.612042391 | 0.22750958 | 0.00060854 | 0.003451182 |
| MINDY2 | 3316.206532 | 0.611693638 | 0.112057074 | 6.50E-09 | 1.48E-07 |
| YIPF2 | 622.3897229 | 0.61160115 | 0.162129236 | 1.83E-05 | 0.000172763 |
| CLEC2D | 332.8247351 | 0.611518553 | 0.145590458 | 3.22E-06 | 3.72E-05 |
| TUBB8 | 4.266285422 | 0.611505839 | 0.790081943 | 0.00658715 | 0.02520162 |
| DCST2 | 24.92654393 | 0.61149322 | 0.319586784 | 0.00333809 | 0.01438486 |
| NPAS3 | 36.02988297 | 0.611463852 | 0.30604896 | 0.00287405 | 0.012742969 |
| CFAP276 | 26.80522968 | 0.611289489 | 0.374587942 | 0.0051598 | 0.020595135 |
| LIN54 | 2199.391173 | 0.610438143 | 0.085808037 | 1.66E-13 | 8.73E-12 |
| RND2 | 981.8527055 | 0.610141516 | 0.184655912 | 9.90E-05 | 0.00073208 |
| PSMD10P1 | 6.084258906 | 0.609887585 | 0.653245767 | 0.00818997 | 0.030144725 |
| FAAH2 | 6.69050953 | 0.609825598 | 0.623598664 | 0.00834067 | 0.030604314 |
| LETM2 | 346.3730397 | 0.60939457 | 0.153514023 | 8.56E-06 | 8.78E-05 |
| SRGAP2C | 883.970028 | 0.609241947 | 0.134933472 | 8.09E-07 | 1.10E-05 |
| HMBOX1 | 1759.991375 | 0.609241866 | 0.114483461 | 1.41E-08 | 3.01E-07 |
| BTN3A3 | 47.16212117 | 0.608940547 | 0.263042712 | 0.0015517 | 0.00760464 |
| GALNT3 | 124.4778143 | 0.608673147 | 0.226161702 | 0.00062564 | 0.00353547 |
| MNAT1 | 747.0191856 | 0.608518724 | 0.111663502 | 7.02E-09 | 1.60E-07 |
| LEPR | 324.9703882 | 0.608317386 | 0.186770713 | 0.00011759 | 0.000851451 |
| PFN1P2 | 61.30830676 | 0.60818004 | 0.229300432 | 0.00069594 | 0.003868954 |
| ETV5 | 1153.033352 | 0.60756857 | 0.530844298 | 0.00833557 | 0.030593031 |
| FAM13B | 4056.043703 | 0.607532048 | 0.090183163 | 2.42E-12 | 1.09E-10 |
| POC5 | 1312.012992 | 0.607012992 | 0.092468896 | 7.77E-12 | 3.15E-10 |
| MOSPD2 | 745.6994716 | 0.606783165 | 0.145541965 | 3.83E-06 | 4.33E-05 |
| SLC2A10 | 87.85078471 | 0.606675641 | 0.202496868 | 0.0002704 | 0.001733894 |
| RNASEH2B-AS1 | 19.21172005 | 0.606188921 | 0.376255488 | 0.00551517 | 0.021726804 |
| ERC2 | 365.3444359 | 0.606156389 | 0.116932351 | 3.03E-08 | 6.04E-07 |
| LNX2 | 562.7737313 | 0.606050383 | 0.113349978 | 1.26E-08 | 2.73E-07 |
| MYLK3 | 55.29524891 | 0.605812361 | 0.233439655 | 0.00082312 | 0.004448515 |
| SHOC2 | 2224.878768 | 0.605489642 | 0.100413448 | 2.42E-10 | 7.35E-09 |
| APOE | 47.06099603 | 0.605472543 | 0.289375636 | 0.00253325 | 0.01146177 |
| ARHGAP26 | 731.8489004 | 0.605064306 | 0.149667829 | 6.60E-06 | 7.00E-05 |
| CCDC113 | 421.6877583 | 0.60503896 | 0.150947732 | 7.60E-06 | 7.94E-05 |
| ZNF461 | 672.3076426 | 0.604988966 | 0.127829777 | 3.00E-07 | 4.64E-06 |
| TMEM45B | 64.87867209 | 0.604874564 | 0.257799058 | 0.00150092 | 0.007391026 |
| EMC7 | 1653.761298 | 0.604816643 | 0.089803392 | 2.50E-12 | 1.12E-10 |
| GPATCH2 | 1713.727924 | 0.604183404 | 0.098095488 | 1.10E-10 | 3.57E-09 |
| MBOAT4 | 21.82863944 | 0.604164223 | 0.380416808 | 0.00576943 | 0.022577094 |
| RGPD6 | 80.41587699 | 0.604131071 | 0.336733279 | 0.0043105 | 0.017806121 |
| USP15 | 4284.34012 | 0.603944038 | 0.098240806 | 1.18E-10 | 3.81E-09 |
| PNPLA3 | 630.4503932 | 0.603834364 | 0.163624342 | 2.67E-05 | 0.000239235 |
| HM13 | 4544.329718 | 0.603657212 | 0.128273439 | 3.45E-07 | 5.24E-06 |
| RDH16 | 45.52277996 | 0.603372951 | 0.267721228 | 0.00186063 | 0.008898154 |
| ERMARD | 1040.400784 | 0.603131329 | 0.120061842 | 7.16E-08 | 1.31E-06 |
| ZBTB21 | 2113.694942 | 0.602788674 | 0.113573991 | 1.61E-08 | 3.37E-07 |
| DACT3 | 797.3404206 | 0.602098244 | 0.208483232 | 0.00038659 | 0.002354118 |
| FSIP2LP | 10.82202193 | 0.602034275 | 0.497391731 | 0.00837909 | 0.030725336 |
| VCAM1 | 5.739778821 | 0.601673384 | 0.787810349 | 0.00684366 | 0.025975084 |

| | | | | | |
|---|---|---|---|---|---|
| GCC2 | 3424.997663 | 0.60134409 | 0.106860383 | 2.73E-09 | 6.73E-08 |
| SLC39A13-AS1 | 6.198717659 | 0.601112658 | 0.598687644 | 0.00885559 | 0.032231216 |
| BRAF | 2385.296773 | 0.600116648 | 0.110409744 | 8.14E-09 | 1.82E-07 |
| RANBP17 | 1600.054292 | 0.600102118 | 0.125669879 | 2.54E-07 | 4.00E-06 |
| KDM6A | 1490.169066 | 0.599758486 | 0.1021345 | 6.58E-10 | 1.83E-08 |
| GOLGA4 | 3765.833365 | 0.599713028 | 0.109976821 | 7.40E-09 | 1.66E-07 |
| NRTN | 7.763389937 | 0.599515119 | 0.566551664 | 0.00896233 | 0.032521843 |
| ENTPD3-AS1 | 52.00225595 | 0.599258406 | 0.261133031 | 0.0017668 | 0.00849087 |
| EIF1B-AS1 | 52.00225595 | 0.599258406 | 0.261133031 | 0.0017668 | 0.00849087 |
| CCS | 781.8162425 | 0.599212949 | 0.121077399 | 1.08E-07 | 1.90E-06 |
| FBRS | 3117.023648 | 0.599137018 | 0.189768974 | 0.00017622 | 0.001209543 |
| TUBE1 | 4502.668912 | 0.598951696 | 0.144756311 | 4.68E-06 | 5.17E-05 |
| TMEM50B | 1106.703436 | 0.597811165 | 0.105055585 | 1.95E-09 | 4.91E-08 |
| STYX | 769.4579718 | 0.59700753 | 0.10386064 | 1.40E-09 | 3.64E-08 |
| BMPER | 21.80715124 | 0.596779957 | 0.326779998 | 0.00435153 | 0.017925647 |
| EXOC6B | 1899.074474 | 0.596649414 | 0.095532892 | 6.74E-11 | 2.28E-09 |
| NPPC | 13.78183635 | 0.596369313 | 0.4513416 | 0.00807746 | 0.029805278 |
| MDGA1 | 1523.305136 | 0.596173102 | 0.207772656 | 0.00043011 | 0.002576381 |
| USP8 | 3403.306327 | 0.595628354 | 0.089885519 | 5.60E-12 | 2.34E-10 |
| SNAP91 | 2943.263523 | 0.595358383 | 0.129945098 | 6.67E-07 | 9.32E-06 |
| NPM1P13 | 17.48684344 | 0.59505231 | 0.400271597 | 0.00696941 | 0.026355097 |
| MSTO2P | 1074.554397 | 0.59491167 | 0.143466919 | 4.66E-06 | 5.15E-05 |
| PTPN6 | 47.67443389 | 0.594816376 | 0.304908682 | 0.00359695 | 0.015318795 |
| MAB21L3 | 19.82778901 | 0.594520446 | 0.377139844 | 0.00630352 | 0.024342839 |
| PDHX | 2027.936313 | 0.594486041 | 0.085513605 | 5.99E-13 | 2.90E-11 |
| ARHGEF33 | 23.83663332 | 0.594408365 | 0.342856057 | 0.00510587 | 0.02042311 |
| TCEA3 | 13.54678595 | 0.594345172 | 0.432100724 | 0.00782719 | 0.029048607 |
| GPRC5C | 360.5260176 | 0.594201919 | 0.16329634 | 3.52E-05 | 0.000303384 |
| ABAT | 1500.995609 | 0.594077771 | 0.130808595 | 8.12E-07 | 1.11E-05 |
| SHC2 | 1689.063024 | 0.59371659 | 0.167455976 | 4.96E-05 | 0.000405912 |
| ACRBP | 114.90071 | 0.593340293 | 0.203371698 | 0.00038418 | 0.002342209 |
| ZNF585A | 1228.720876 | 0.593136466 | 0.124745239 | 2.97E-07 | 4.61E-06 |
| SETD7 | 4968.721658 | 0.59302059 | 0.097175105 | 1.70E-10 | 5.34E-09 |
| RFFL | 402.9149027 | 0.592904507 | 0.135840952 | 1.84E-06 | 2.27E-05 |
| RIT1 | 1635.273061 | 0.592153522 | 0.092395111 | 2.43E-11 | 9.05E-10 |
| PLK3 | 488.3189091 | 0.592022874 | 0.201025897 | 0.00035878 | 0.002209626 |
| MINDY3 | 1709.524313 | 0.591641676 | 0.08924666 | 5.67E-12 | 2.36E-10 |
| TRPV6 | 24.24148453 | 0.591630705 | 0.318355277 | 0.00431345 | 0.017811611 |
| H3C13 | 21.99457215 | 0.591600029 | 0.397590708 | 0.00714174 | 0.026887302 |
| MAML2 | 120.8184783 | 0.591345695 | 0.250853801 | 0.0016518 | 0.008024498 |
| FAM149B1 | 1281.183767 | 0.591237423 | 0.12196017 | 1.91E-07 | 3.14E-06 |
| PHKG1 | 139.7525452 | 0.591114441 | 0.224775399 | 0.00086041 | 0.004625739 |
| ARHGAP20 | 25.36351964 | 0.590898409 | 0.302355086 | 0.00369573 | 0.015644394 |
| PIWIL4 | 82.74045868 | 0.59080059 | 0.207002929 | 0.0004713 | 0.002785011 |
| PDXK | 2116.772447 | 0.590526032 | 0.135132454 | 1.83E-06 | 2.26E-05 |
| HRH1 | 8.189322261 | 0.590035134 | 0.512521777 | 0.00933556 | 0.033597563 |
| PHC3 | 3662.990829 | 0.58986346 | 0.111595068 | 2.00E-08 | 4.12E-07 |
| ATAD2B | 1752.166423 | 0.58979235 | 0.095314447 | 1.02E-10 | 3.35E-09 |
| UBN2 | 3758.040858 | 0.589654712 | 0.101067969 | 8.92E-10 | 2.43E-08 |
| RAB6A | 4803.592608 | 0.589442933 | 0.10570547 | 4.01E-09 | 9.50E-08 |

| | | | | | |
|---|---|---|---|---|---|
| INSIG1 | 2993.30788 | 0.588880647 | 0.086473704 | 1.68E-12 | 7.70E-11 |
| LRRC39 | 386.3122627 | 0.588866945 | 0.128410111 | 6.89E-07 | 9.59E-06 |
| IDI2-AS1 | 222.3664334 | 0.588820058 | 0.165857967 | 5.09E-05 | 0.000414896 |
| DHDH | 10.75241824 | 0.588732852 | 0.500004499 | 0.00931877 | 0.033560624 |
| PTPN13 | 7235.536435 | 0.588567687 | 0.116794848 | 7.42E-08 | 1.36E-06 |
| UVRAG | 1309.327377 | 0.588557332 | 0.108200308 | 8.71E-09 | 1.92E-07 |
| CCDC92 | 2096.17737 | 0.588199585 | 0.149229119 | 1.15E-05 | 0.000114773 |
| ARK2N | 2242.329799 | 0.587938142 | 0.09981624 | 6.47E-10 | 1.81E-08 |
| ME1 | 638.9019239 | 0.587934444 | 0.144416211 | 6.78E-06 | 7.16E-05 |
| GBX2 | 17.16154303 | 0.587610461 | 0.430685842 | 0.00829901 | 0.030496946 |
| BNIPL | 88.45760634 | 0.587421358 | 0.226423963 | 0.00097323 | 0.005144876 |
| SCUBE2 | 28.85029506 | 0.587133498 | 0.297632477 | 0.0036958 | 0.015644394 |
| RPL10P19 | 22.54473766 | 0.586966438 | 0.37091949 | 0.00663407 | 0.025333886 |
| NONOP2 | 51.04907683 | 0.586959201 | 0.254090328 | 0.00190634 | 0.009091349 |
| PLGLB2 | 41.34071433 | 0.586756107 | 0.266109596 | 0.00238435 | 0.010910491 |
| LRRC4C | 17.97667856 | 0.585870252 | 0.370982758 | 0.00671747 | 0.025581128 |
| IL23A | 21.65617656 | 0.585514144 | 0.339583244 | 0.00556227 | 0.021869914 |
| SNX16 | 328.9596382 | 0.585334551 | 0.127220457 | 6.60E-07 | 9.26E-06 |
| FRY | 845.8620149 | 0.585166481 | 0.148746357 | 1.22E-05 | 0.000120587 |
| TYRP1 | 6.610743325 | 0.58504772 | 0.565876828 | 0.00984873 | 0.035070006 |
| FAM200A | 854.9764853 | 0.584904556 | 0.108363787 | 1.13E-08 | 2.46E-07 |
| FOXJ1 | 21.79944753 | 0.584812278 | 0.451607988 | 0.00894385 | 0.03248943 |
| NKD1 | 1017.704076 | 0.584591424 | 0.158386584 | 3.14E-05 | 0.000275452 |
| RNU1-91P | 6.709951005 | 0.5845097 | 0.577912585 | 0.00986281 | 0.035101901 |
| SCAPER | 1793.309268 | 0.584457632 | 0.105296 | 4.82E-09 | 1.13E-07 |
| RNU1-129P | 2.552370658 | 0.584374869 | 0.856861901 | 0.0057647 | 0.022569163 |
| GPR35 | 145.4755264 | 0.584343118 | 0.168831906 | 7.26E-05 | 0.00056159 |
| KCNC4 | 1169.85505 | 0.583965685 | 0.164668739 | 5.38E-05 | 0.00043359 |
| ABI3BP | 21.40607943 | 0.583923471 | 0.360090532 | 0.00647064 | 0.02485288 |
| EFCAB7 | 845.3404381 | 0.58373301 | 0.123131968 | 3.43E-07 | 5.21E-06 |
| ALKBH3-AS1 | 21.69966006 | 0.583633569 | 0.351721016 | 0.00617454 | 0.023903078 |
| KLF6 | 2213.803547 | 0.583105174 | 0.089704295 | 1.43E-11 | 5.55E-10 |
| BCAS3 | 871.5815144 | 0.583088383 | 0.131960251 | 1.56E-06 | 1.98E-05 |
| REL-DT | 8.876116304 | 0.582815352 | 0.610038756 | 0.00981717 | 0.0349879 |
| CLEC19A | 4.960527229 | 0.582441815 | 0.745643015 | 0.00808846 | 0.029835156 |
| TMEM217 | 14.74212323 | 0.581481673 | 0.417219831 | 0.00846017 | 0.030967416 |
| LYG1 | 128.0127095 | 0.58146916 | 0.181448581 | 0.00017684 | 0.001212178 |
| GKN2 | 7.723990012 | 0.581115241 | 0.658750752 | 0.0094554 | 0.033963478 |
| TMEM241 | 641.2690615 | 0.581010829 | 0.125312592 | 5.78E-07 | 8.28E-06 |
| PINK1 | 158.6186351 | 0.580920334 | 0.168545229 | 7.87E-05 | 0.000600885 |
| LNP1 | 371.4485982 | 0.580558374 | 0.118202436 | 1.51E-07 | 2.55E-06 |
| SAP18P3 | 7.155845932 | 0.58039195 | 0.522452745 | 0.0100845 | 0.035779432 |
| TBL2 | 2521.237463 | 0.580282406 | 0.133213103 | 2.11E-06 | 2.56E-05 |
| RELCH | 2031.710393 | 0.57991942 | 0.092282193 | 5.97E-11 | 2.06E-09 |
| CYP2F2P | 16.31630758 | 0.579652452 | 0.419428737 | 0.00866532 | 0.031610859 |
| DNAJB5 | 1768.062437 | 0.57953818 | 0.112236277 | 4.16E-08 | 8.07E-07 |
| LINC03040 | 7.181280377 | 0.578636628 | 0.597107296 | 0.0101426 | 0.0359236 |
| ITPR1 | 3704.52204 | 0.578338115 | 0.115785259 | 1.01E-07 | 1.79E-06 |
| SEPSECS | 1057.897831 | 0.578246246 | 0.114671143 | 7.90E-08 | 1.44E-06 |
| ZNF79 | 433.9984321 | 0.578007217 | 0.121878551 | 3.55E-07 | 5.38E-06 |

| | | | | | |
|---|---|---|---|---|---|
| NECTIN2 | 4.765359408 | 0.577833701 | 0.666199585 | 0.0095183 | 0.034123845 |
| KANSL2 | 3350.980317 | 0.577797221 | 0.090005898 | 2.52E-11 | 9.37E-10 |
| RHOXF1 | 151.7123901 | 0.577514257 | 0.180205901 | 0.00018285 | 0.001245886 |
| SLC7A6OS | 423.8396146 | 0.577148276 | 0.131002383 | 1.73E-06 | 2.17E-05 |
| RRM2B | 1086.18186 | 0.576998296 | 0.123976041 | 5.48E-07 | 7.92E-06 |
| PKMP1 | 29.69025232 | 0.576895451 | 0.278043041 | 0.00335441 | 0.014443073 |
| NOTUM | 143.8981864 | 0.57601526 | 0.192056522 | 0.0003516 | 0.00216932 |
| GJB5 | 5.741583559 | 0.575768596 | 0.625429786 | 0.01008927 | 0.035784026 |
| CRIP2 | 1370.877766 | 0.575735095 | 0.175295054 | 0.00014297 | 0.001011382 |
| NRAV | 217.7729051 | 0.575392176 | 0.143036484 | 9.17E-06 | 9.33E-05 |
| KRBOX5 | 1084.625066 | 0.57529416 | 0.118813581 | 2.23E-07 | 3.58E-06 |
| KRTAP5-1 | 52.90348652 | 0.575084861 | 0.285454012 | 0.00380534 | 0.016061797 |
| DENND1C | 2.697797746 | 0.574943033 | 0.84998689 | 0.00602 | 0.023406165 |
| C2CD2L | 1260.13797 | 0.574626039 | 0.160658282 | 5.21E-05 | 0.000422492 |
| UBR1 | 4257.228714 | 0.574214101 | 0.104779227 | 7.73E-09 | 1.73E-07 |
| TNN | 14.3031501 | 0.573981385 | 0.423269156 | 0.00923799 | 0.033356919 |
| DUSP5 | 134.2156147 | 0.573820356 | 0.251573326 | 0.00227984 | 0.010523579 |
| MAPK8IP1 | 2632.219455 | 0.573757461 | 0.207549944 | 0.00070273 | 0.003899307 |
| ZBTB43 | 1607.691806 | 0.572976622 | 0.086461909 | 6.60E-12 | 2.72E-10 |
| RPS29P5 | 29.04205149 | 0.572539918 | 0.32387525 | 0.0058128 | 0.022709254 |
| PURG | 751.0741049 | 0.571258386 | 0.132856224 | 2.91E-06 | 3.42E-05 |
| INTS6 | 3547.182773 | 0.571255729 | 0.100678873 | 2.62E-09 | 6.48E-08 |
| HABP4 | 827.2585825 | 0.571224422 | 0.126634565 | 1.12E-06 | 1.48E-05 |
| BRAP | 2718.156231 | 0.571083356 | 0.099153263 | 1.59E-09 | 4.08E-08 |
| HEYL | 100.3347186 | 0.571030962 | 0.297651201 | 0.0046531 | 0.018940309 |
| ZNF319 | 713.7839191 | 0.57091334 | 0.208999524 | 0.00078729 | 0.004288719 |
| NEK10 | 31.10981033 | 0.570745137 | 0.323127436 | 0.00591307 | 0.023057161 |
| USP27X | 178.9677289 | 0.570554992 | 0.153982902 | 3.34E-05 | 0.000290182 |
| ZNF134 | 1639.86666 | 0.570429428 | 0.084666044 | 3.17E-12 | 1.39E-10 |
| RNF217 | 961.9257863 | 0.570090366 | 0.103682736 | 7.20E-09 | 1.63E-07 |
| H2BC21 | 144.9456856 | 0.569951032 | 0.175606352 | 0.0001706 | 0.00117641 |
| HMGCS1 | 12303.80193 | 0.569873858 | 0.112762879 | 7.94E-08 | 1.44E-06 |
| CHL1 | 321.9533999 | 0.56931617 | 0.178592944 | 0.000207 | 0.001378056 |
| CBX8 | 2199.554967 | 0.568951248 | 0.11140984 | 6.06E-08 | 1.13E-06 |
| CEP295 | 1729.16835 | 0.5689309 | 0.126953027 | 1.31E-06 | 1.70E-05 |
| REELD1 | 62.3730464 | 0.568920534 | 0.23448919 | 0.00172251 | 0.008311027 |
| ACSS2 | 867.5373413 | 0.568824331 | 0.103544907 | 7.47E-09 | 1.67E-07 |
| KCNH3 | 206.1077935 | 0.568610685 | 0.195466781 | 0.00048905 | 0.002875843 |
| INPP5B | 1187.608428 | 0.568501933 | 0.116568458 | 1.97E-07 | 3.23E-06 |
| DYRK1B | 629.1898926 | 0.568097388 | 0.172543733 | 0.00014834 | 0.001043984 |
| PIERCE2 | 50.09839534 | 0.568089455 | 0.258039297 | 0.00282895 | 0.012591428 |
| GARS1 | 8542.491501 | 0.567713762 | 0.108527279 | 3.17E-08 | 6.28E-07 |
| SNAP23 | 317.2835383 | 0.56764027 | 0.140010471 | 8.55E-06 | 8.78E-05 |
| LINC00242 | 54.1075611 | 0.567294449 | 0.24828717 | 0.00239459 | 0.01094033 |
| DNAH2 | 39.86598942 | 0.567283143 | 0.295029317 | 0.00477045 | 0.019329967 |
| ARCN1 | 6081.490948 | 0.567226412 | 0.079684827 | 2.22E-13 | 1.15E-11 |
| RYR2 | 2064.164051 | 0.566753927 | 0.128195349 | 1.75E-06 | 2.19E-05 |
| FAM186A | 28.33298516 | 0.566702245 | 0.333189004 | 0.00670435 | 0.02554534 |
| TLL2 | 15.5918658 | 0.566510026 | 0.400306521 | 0.00929179 | 0.0334928 |
| SLC22A12 | 3.1483069 | 0.566060237 | 0.86798015 | 0.00565083 | 0.022184729 |

| | | | | | |
|---|---|---|---|---|---|
| NGLY1 | 1448.792211 | 0.565369402 | 0.1014299 | 4.85E-09 | 1.14E-07 |
| GPR89B | 420.0560454 | 0.564997618 | 0.175736966 | 0.00019647 | 0.001320781 |
| C4A | 52.29298907 | 0.564930972 | 0.270782968 | 0.00364399 | 0.015460094 |
| PRKRIP1 | 1271.772998 | 0.564863003 | 0.120394275 | 5.01E-07 | 7.32E-06 |
| MACO1 | 2412.600841 | 0.564749072 | 0.095071564 | 5.66E-10 | 1.60E-08 |
| SCLT1 | 1603.305349 | 0.564691623 | 0.095173011 | 5.91E-10 | 1.66E-08 |
| TBC1D23 | 2049.916051 | 0.564242373 | 0.101360026 | 5.10E-09 | 1.18E-07 |
| GNG7 | 9.345201712 | 0.563906105 | 0.463837852 | 0.01090067 | 0.038090278 |
| LRP10 | 138.2345891 | 0.563865188 | 0.258042757 | 0.00303463 | 0.013339935 |
| EPB41L4A-AS1 | 1649.573409 | 0.563584092 | 0.111964597 | 9.22E-08 | 1.65E-06 |
| NARS1 | 11568.95827 | 0.563490418 | 0.088982111 | 4.91E-11 | 1.72E-09 |
| OXCT2P1 | 6.793411022 | 0.562808206 | 0.56042653 | 0.01138688 | 0.039487329 |
| ZNF501 | 630.815174 | 0.562791594 | 0.110919015 | 7.53E-08 | 1.37E-06 |
| PAOX | 21.71454279 | 0.562664594 | 0.348669968 | 0.00775294 | 0.028817001 |
| OR7D2 | 84.92112697 | 0.562267049 | 0.222095951 | 0.00141988 | 0.007060091 |
| SIX4 | 517.1996326 | 0.561634191 | 0.141273348 | 1.24E-05 | 0.000122525 |
| TMEM202-AS1 | 32.92197645 | 0.561580543 | 0.27935441 | 0.00431126 | 0.017806121 |
| RBAK | 2377.369759 | 0.561323916 | 0.11227689 | 1.12E-07 | 1.95E-06 |
| RIMOC1 | 3110.869099 | 0.561258358 | 0.102431635 | 8.53E-09 | 1.89E-07 |
| GLIPR1-AS1 | 75.71524405 | 0.560946954 | 0.266422702 | 0.00363842 | 0.015454609 |
| UBXN2B | 3321.539094 | 0.56062187 | 0.106400124 | 2.72E-08 | 5.46E-07 |
| MVK | 986.7214804 | 0.560554165 | 0.130181701 | 3.07E-06 | 3.58E-05 |
| KDM4C | 861.5150477 | 0.560473326 | 0.11150901 | 9.79E-08 | 1.74E-06 |
| TRIM72 | 8.54989124 | 0.560306234 | 0.47087447 | 0.01131798 | 0.039301424 |
| SGTB | 1316.694914 | 0.560129854 | 0.122182257 | 8.65E-07 | 1.17E-05 |
| CEP350 | 4981.534985 | 0.559916384 | 0.099600696 | 3.84E-09 | 9.15E-08 |
| H1-7 | 10.91392676 | 0.559722191 | 0.43862023 | 0.01087529 | 0.038027453 |
| GAN | 926.4209444 | 0.559569355 | 0.111245035 | 9.66E-08 | 1.72E-06 |
| REST | 1740.129024 | 0.559534561 | 0.128662936 | 2.56E-06 | 3.04E-05 |
| PDP1 | 1444.550173 | 0.559325899 | 0.094705747 | 7.24E-10 | 1.99E-08 |
| ARIH2OS | 41.81269916 | 0.55920727 | 0.243981936 | 0.00253137 | 0.011455801 |
| LINC01964 | 2.53487661 | 0.55911062 | 0.837422784 | 0.00650638 | 0.024959849 |
| ARHGAP40 | 4.150435278 | 0.558729133 | 0.755202357 | 0.00859868 | 0.031418492 |
| ALDH6A1 | 717.9032644 | 0.558496743 | 0.11536297 | 2.53E-07 | 3.99E-06 |
| MADD-AS1 | 35.01447999 | 0.558416435 | 0.342094154 | 0.00784612 | 0.029085511 |
| DUSP19 | 130.9069868 | 0.558218766 | 0.173373186 | 0.00020452 | 0.001365514 |
| LINC00174 | 477.6620879 | 0.55816696 | 0.164558446 | 0.00011473 | 0.000832206 |
| PCSK5 | 134.4281064 | 0.55792893 | 0.194220276 | 0.0005946 | 0.003381452 |
| RASGRF2-AS1 | 21.67098597 | 0.557546528 | 0.372743369 | 0.00922304 | 0.033315046 |
| ZNF536 | 1258.322771 | 0.557244147 | 0.112766558 | 1.54E-07 | 2.59E-06 |
| UNC5CL | 150.9876144 | 0.55722964 | 0.213171298 | 0.00120667 | 0.006181474 |
| AFAP1L2 | 25.61151125 | 0.557179481 | 0.314723054 | 0.0065884 | 0.02520162 |
| LINC00173 | 13.5740648 | 0.556948891 | 0.391573388 | 0.00994283 | 0.035331658 |
| COPA | 16914.15903 | 0.556654168 | 0.082220719 | 2.78E-12 | 1.24E-10 |
| AKIRIN1 | 6361.989844 | 0.556631518 | 0.11405313 | 2.11E-07 | 3.41E-06 |
| SPEF1 | 25.54201803 | 0.556517503 | 0.320795618 | 0.00696711 | 0.026351243 |
| KLHL24 | 3385.792555 | 0.556385425 | 0.125461195 | 1.78E-06 | 2.21E-05 |
| RRBP1 | 1473.479446 | 0.556295017 | 0.162844465 | 0.00010707 | 0.000784963 |
| DLGAP2 | 1443.146326 | 0.556257081 | 0.166119704 | 0.00013519 | 0.000963975 |
| PLPPR4 | 1362.252442 | 0.556206433 | 0.127697734 | 2.54E-06 | 3.02E-05 |

| | | | | | |
|---|---|---|---|---|---|
| SERINC3 | 2574.126919 | 0.555911871 | 0.084345518 | 9.45E-12 | 3.79E-10 |
| TRIP4 | 1227.772246 | 0.555734844 | 0.091134086 | 2.29E-10 | 7.03E-09 |
| NEFM | 2105.173887 | 0.555698178 | 0.196931664 | 0.00069997 | 0.00388718 |
| FGF18 | 19.58927414 | 0.555621888 | 0.332414518 | 0.00764164 | 0.028467885 |
| TMEM196 | 444.8691521 | 0.555064194 | 0.179781786 | 0.00032012 | 0.001999706 |
| HSD17B12 | 2522.017479 | 0.554969545 | 0.087085562 | 4.01E-11 | 1.44E-09 |
| TXNDC11-AS1 | 3.623991056 | 0.554750067 | 0.802851254 | 0.00748964 | 0.027987823 |
| HIVEP3 | 410.9485224 | 0.554729641 | 0.238564373 | 0.00244881 | 0.011153427 |
| RCOR3 | 4767.368537 | 0.554679405 | 0.092421546 | 4.18E-10 | 1.22E-08 |
| EIF2B2 | 1703.660864 | 0.554568015 | 0.083634827 | 7.28E-12 | 2.97E-10 |
| RBM20 | 666.0304913 | 0.554386367 | 0.126335551 | 2.23E-06 | 2.68E-05 |
| RPS15AP24 | 5.312899894 | 0.554216135 | 0.619636309 | 0.01142918 | 0.039615467 |
| ITGA10 | 188.9896646 | 0.554143205 | 0.182562251 | 0.00037883 | 0.002315061 |
| SORBS1 | 1185.020594 | 0.553744301 | 0.091456882 | 3.03E-10 | 9.07E-09 |
| NAP1L3 | 1865.315137 | 0.553662816 | 0.095878294 | 1.65E-09 | 4.20E-08 |
| SC5D | 2615.092789 | 0.5529804 | 0.13160362 | 5.11E-06 | 5.60E-05 |
| PMAIP1 | 3486.634445 | 0.552764389 | 0.091485079 | 3.30E-10 | 9.81E-09 |
| C10orf88 | 1247.31334 | 0.552261128 | 0.099084769 | 5.33E-09 | 1.23E-07 |
| CCDC71L | 437.2797269 | 0.552160736 | 0.194282046 | 0.00068089 | 0.003803821 |
| TRIM4 | 1477.629333 | 0.552036194 | 0.10059484 | 8.67E-09 | 1.92E-07 |
| CNTN4 | 422.4764186 | 0.55170805 | 0.151048399 | 4.72E-05 | 0.000390538 |
| FERMT2 | 5229.981386 | 0.551339564 | 0.111041323 | 1.43E-07 | 2.43E-06 |
| CPLANE1 | 1500.195069 | 0.551246416 | 0.120885949 | 1.03E-06 | 1.38E-05 |
| ZNF354A | 1384.340404 | 0.551040563 | 0.120177509 | 9.20E-07 | 1.24E-05 |
| MAPK8IP1P2 | 58.10056921 | 0.55086496 | 0.269916084 | 0.0044794 | 0.018354421 |
| CCDC146 | 147.5044739 | 0.550733447 | 0.159248552 | 9.65E-05 | 0.000716288 |
| PTPN4 | 1962.282421 | 0.550027579 | 0.110154401 | 1.25E-07 | 2.15E-06 |
| RNF182 | 1893.357599 | 0.549862956 | 0.111290954 | 1.63E-07 | 2.72E-06 |
| ETV3 | 2391.17078 | 0.549737635 | 0.093500224 | 9.05E-10 | 2.45E-08 |
| SYPL1 | 926.9104901 | 0.549727493 | 0.108910608 | 9.46E-08 | 1.69E-06 |
| FHDC1 | 116.4587399 | 0.549532168 | 0.21584069 | 0.00152938 | 0.007509597 |
| CAPN13 | 87.09646789 | 0.549409758 | 0.201647042 | 0.00096427 | 0.005102745 |
| SRPRB | 2824.72484 | 0.549262826 | 0.085002763 | 2.33E-11 | 8.69E-10 |
| TFE3 | 1194.519214 | 0.548789739 | 0.187851766 | 0.00055691 | 0.003200541 |
| TMEM176A | 26.95164891 | 0.548550683 | 0.323834139 | 0.00787385 | 0.029169116 |
| RUNDC1 | 1912.494312 | 0.54834694 | 0.147485599 | 3.78E-05 | 0.00032225 |
| GOLGA2 | 2746.49312 | 0.547880257 | 0.130147116 | 5.13E-06 | 5.61E-05 |
| BIRC2 | 1636.324989 | 0.547607481 | 0.13219785 | 6.86E-06 | 7.24E-05 |
| HTT-AS | 3.812925091 | 0.54757544 | 0.727342101 | 0.00963471 | 0.034464342 |
| MAILR | 174.2104188 | 0.547410169 | 0.186107377 | 0.00053105 | 0.003078815 |
| TXNRD3 | 241.29909 | 0.54739356 | 0.12970261 | 4.92E-06 | 5.40E-05 |
| MED26 | 233.8932682 | 0.547116382 | 0.183259096 | 0.00046669 | 0.002761741 |
| PLOD3 | 1297.199324 | 0.546974971 | 0.161302688 | 0.00012568 | 0.000901146 |
| NMNAT1 | 123.8450416 | 0.546251916 | 0.188156739 | 0.00059969 | 0.003406608 |
| H2AC11 | 35.63973299 | 0.546221075 | 0.294218351 | 0.00636059 | 0.024512572 |
| GORAB | 891.225197 | 0.545733022 | 0.108035656 | 9.53E-08 | 1.70E-06 |
| TNRC6B | 3117.430353 | 0.545148989 | 0.090490622 | 3.87E-10 | 1.14E-08 |
| CDC37L1 | 914.4688453 | 0.54491137 | 0.095224968 | 2.37E-09 | 5.90E-08 |
| ZHX2 | 206.9290935 | 0.544880896 | 0.221787885 | 0.00197538 | 0.009351224 |
| PDC-AS1 | 86.75560989 | 0.544857592 | 0.199453643 | 0.00098241 | 0.005184077 |

| | | | | | |
|---|---|---|---|---|---|
| RMND5A | 5582.197556 | 0.544853086 | 0.081936378 | 6.83E-12 | 2.80E-10 |
| TMEM128 | 1527.416467 | 0.544814612 | 0.089295555 | 2.41E-10 | 7.34E-09 |
| CFAP58 | 10.53032805 | 0.544202249 | 0.443541564 | 0.01254658 | 0.042660046 |
| MTHFD2 | 15631.97293 | 0.544094926 | 0.087841777 | 1.35E-10 | 4.30E-09 |
| MAP3K2 | 5258.599689 | 0.543709524 | 0.091760376 | 7.14E-10 | 1.97E-08 |
| EPHA1-AS1 | 4.519638855 | 0.54367886 | 0.685620531 | 0.01079181 | 0.037812692 |
| IL12RB2 | 94.52356088 | 0.543615952 | 0.209420448 | 0.00141889 | 0.007058045 |
| ATP2B1-AS1 | 29.52650413 | 0.5435123 | 0.299966467 | 0.00696677 | 0.026351243 |
| SRPRA | 3651.658825 | 0.543493629 | 0.116313376 | 6.40E-07 | 9.02E-06 |
| TYW1 | 1665.714854 | 0.543186596 | 0.087333401 | 1.16E-10 | 3.75E-09 |
| GEMIN2 | 842.4160974 | 0.543163024 | 0.122178024 | 1.86E-06 | 2.29E-05 |
| SETDB2 | 1106.244902 | 0.54299837 | 0.1259866 | 3.42E-06 | 3.92E-05 |
| MIR2116 | 6.720493771 | 0.542993482 | 0.56070886 | 0.01292254 | 0.043649665 |
| HOXD4 | 287.3211356 | 0.542622909 | 0.144424488 | 3.40E-05 | 0.000294536 |
| PRKG1 | 85.25580055 | 0.542561574 | 0.198480869 | 0.00099632 | 0.0052413 |
| ID2 | 5877.48175 | 0.542139948 | 0.090348676 | 4.56E-10 | 1.31E-08 |
| DNAJB4 | 1990.567195 | 0.542024366 | 0.09220727 | 9.59E-10 | 2.59E-08 |
| COL2A1 | 67.07799478 | 0.541289816 | 0.27899948 | 0.0058015 | 0.0226767 |
| RBM11 | 44.82517492 | 0.540994064 | 0.268056575 | 0.00508346 | 0.020377447 |
| B4GALT4 | 642.5205574 | 0.54046194 | 0.116514929 | 7.69E-07 | 1.05E-05 |
| GIHCG | 410.8555568 | 0.540391986 | 0.135008982 | 1.30E-05 | 0.00012781 |
| KLF4 | 147.6411771 | 0.540377356 | 0.198373828 | 0.00104096 | 0.00544272 |
| TBX4 | 97.07240295 | 0.540354997 | 0.227063397 | 0.00245478 | 0.011175566 |
| FOXC1 | 455.4889161 | 0.540320373 | 0.187298684 | 0.00066372 | 0.003724084 |
| AZI2 | 1091.049937 | 0.539734375 | 0.091078058 | 7.29E-10 | 2.00E-08 |
| SV2C | 4171.831209 | 0.539733901 | 0.164743922 | 0.0001964 | 0.001320742 |
| GGNBP2 | 3527.623232 | 0.539554338 | 0.078540281 | 1.55E-12 | 7.15E-11 |
| INSIG2 | 727.8068498 | 0.539273562 | 0.120972056 | 1.80E-06 | 2.24E-05 |
| SLC22A4 | 55.2395892 | 0.539090016 | 0.228575366 | 0.00260354 | 0.011723069 |
| RFX3 | 3100.567958 | 0.538851718 | 0.124144359 | 3.08E-06 | 3.58E-05 |
| TTLL7 | 2540.510739 | 0.538846959 | 0.102244656 | 3.14E-08 | 6.21E-07 |
| RHOXF1-AS1 | 35.32289217 | 0.538770365 | 0.264917359 | 0.00504045 | 0.020228674 |
| SLC35B1 | 5231.29977 | 0.5382657 | 0.088558838 | 2.90E-10 | 8.72E-09 |
| NCAL1 | 16.01452762 | 0.537774634 | 0.375829648 | 0.01149516 | 0.039768919 |
| EDEM2 | 324.6820126 | 0.537472001 | 0.136896098 | 1.81E-05 | 0.00017169 |
| PTGER4P2 | 22.99728686 | 0.537165605 | 0.321835628 | 0.00895144 | 0.032507428 |
| MVD | 963.5498778 | 0.537074108 | 0.139927735 | 2.58E-05 | 0.000232462 |
| SERINC1 | 11946.50323 | 0.53688359 | 0.092008921 | 1.28E-09 | 3.37E-08 |
| FOXO3 | 6064.356912 | 0.536462476 | 0.111389538 | 3.34E-07 | 5.09E-06 |
| SLU7 | 2875.296376 | 0.536436411 | 0.083396814 | 3.06E-11 | 1.12E-09 |
| NOL4L | 1876.310111 | 0.536291721 | 0.156949461 | 0.00012452 | 0.000895616 |
| CERS4 | 222.0638309 | 0.536150617 | 0.164033074 | 0.00020707 | 0.001378056 |
| LCLAT1 | 1714.715361 | 0.535987929 | 0.086836549 | 1.63E-10 | 5.14E-09 |
| OTUD7B | 1892.641151 | 0.535984996 | 0.111201711 | 3.28E-07 | 5.03E-06 |
| RNF111 | 3088.769103 | 0.535830163 | 0.103016912 | 4.63E-08 | 8.85E-07 |
| TRIM56 | 7.074659842 | 0.535827252 | 0.554434626 | 0.01358709 | 0.045402172 |
| DGKE | 4675.882573 | 0.535745937 | 0.097199059 | 8.43E-09 | 1.88E-07 |
| TRAPPC13 | 1233.502791 | 0.535586103 | 0.109176478 | 2.15E-07 | 3.47E-06 |
| CRYAB | 13.16189945 | 0.53552468 | 0.421106058 | 0.01311056 | 0.044118089 |
| FAM131A | 661.331399 | 0.535310036 | 0.139741003 | 2.69E-05 | 0.000240679 |

| | | | | | |
|---|---|---|---|---|---|
| MYO1D | 181.9345438 | 0.535197008 | 0.220664592 | 0.00230865 | 0.010637495 |
| ZCCHC10 | 1192.573735 | 0.534947482 | 0.103697162 | 5.84E-08 | 1.09E-06 |
| APCDD1L-DT | 34.99211684 | 0.534627916 | 0.280170707 | 0.00647717 | 0.024873286 |
| ZNF222 | 317.8027637 | 0.534617848 | 0.138701539 | 2.46E-05 | 0.000223526 |
| ZNF256 | 439.4947435 | 0.534505503 | 0.132199062 | 1.14E-05 | 0.000114231 |
| GLRA3 | 9.532330788 | 0.534308975 | 0.476541153 | 0.01397499 | 0.04641926 |
| MIR99AHG | 1820.670525 | 0.533954134 | 0.116455888 | 1.04E-06 | 1.38E-05 |
| ACVR1 | 900.949546 | 0.533669718 | 0.142043873 | 3.63E-05 | 0.000310551 |
| ZNF20 | 55.58698216 | 0.533551831 | 0.217292036 | 0.00218069 | 0.010160746 |
| LMAN2L | 1264.54176 | 0.533496673 | 0.085537543 | 1.10E-10 | 3.58E-09 |
| UBL3 | 1099.446215 | 0.533199958 | 0.09945993 | 1.98E-08 | 4.09E-07 |
| MYSM1 | 7298.06361 | 0.532661146 | 0.105918425 | 1.17E-07 | 2.03E-06 |
| STOX1 | 224.2635707 | 0.532551191 | 0.170479077 | 0.0003411 | 0.002112748 |
| BCDIN3D | 278.8213668 | 0.532512697 | 0.135918129 | 1.94E-05 | 0.00018197 |
| SMPD1 | 355.4569448 | 0.532504728 | 0.166147713 | 0.00026262 | 0.001692417 |
| SEMA5B | 219.6186181 | 0.532440103 | 0.215765916 | 0.00213786 | 0.010000689 |
| GMPPB | 1034.113543 | 0.532392682 | 0.150329551 | 8.22E-05 | 0.000624131 |
| SLC38A6 | 221.4739586 | 0.532150719 | 0.154833886 | 0.00011971 | 0.000864656 |
| LINC00487 | 11.23456825 | 0.532120922 | 0.499948915 | 0.01425816 | 0.047169384 |
| TARS1 | 9983.812205 | 0.531901692 | 0.092650882 | 2.31E-09 | 5.76E-08 |
| CCR10 | 74.31487082 | 0.531858614 | 0.200807346 | 0.00136449 | 0.006842957 |
| PRKG1-AS1 | 53.2397019 | 0.531670712 | 0.25351175 | 0.00476671 | 0.019318606 |
| NAMPT | 2134.376446 | 0.531195066 | 0.090421509 | 1.05E-09 | 2.80E-08 |
| ALDH1L1 | 32.80347409 | 0.531034806 | 0.272604869 | 0.0062574 | 0.024196537 |
| ALPK1 | 213.4264525 | 0.530806206 | 0.206562604 | 0.00168609 | 0.008165996 |
| BLOC1S2 | 1893.200512 | 0.530721082 | 0.088398574 | 4.81E-10 | 1.38E-08 |
| TYW1B | 711.5625596 | 0.530474372 | 0.098770785 | 1.91E-08 | 3.96E-07 |
| TBRG1 | 1066.484269 | 0.530283048 | 0.09984314 | 2.65E-08 | 5.34E-07 |
| ZNF329 | 641.9044496 | 0.530154955 | 0.127342888 | 7.09E-06 | 7.46E-05 |
| PYGM | 65.91872166 | 0.530042708 | 0.23870898 | 0.0037873 | 0.015992223 |
| ADCK5 | 238.371029 | 0.529838987 | 0.155856413 | 0.00013881 | 0.000987026 |
| FAM53C | 3639.621962 | 0.529828178 | 0.112792426 | 6.23E-07 | 8.81E-06 |
| FBXL17 | 848.7197642 | 0.529187295 | 0.120724293 | 2.71E-06 | 3.21E-05 |
| FBXO3 | 2243.919087 | 0.528933294 | 0.100831793 | 3.81E-08 | 7.44E-07 |
| SNAI1 | 926.2257887 | 0.528924067 | 0.129438959 | 9.92E-06 | 0.000100387 |
| RIOK2 | 1688.008526 | 0.528861088 | 0.103484341 | 7.82E-08 | 1.43E-06 |
| SYTL5 | 20.55395242 | 0.528583667 | 0.34884553 | 0.01145687 | 0.039675473 |
| DOCK9-DT | 5.376584671 | 0.527888253 | 0.570992541 | 0.01406304 | 0.046674046 |
| ZFAND3 | 2804.497696 | 0.527804925 | 0.160899629 | 0.0002119 | 0.00140793 |
| PLEKHM3 | 2077.21683 | 0.527435443 | 0.114720486 | 1.02E-06 | 1.36E-05 |
| UAP1 | 3023.049555 | 0.527150804 | 0.097209725 | 1.46E-08 | 3.11E-07 |
| PEX5L | 962.3520145 | 0.527032798 | 0.15112667 | 0.00010385 | 0.00076406 |
| IPMK | 1203.859016 | 0.526860535 | 0.120100225 | 2.71E-06 | 3.21E-05 |
| GAS2 | 117.0119217 | 0.526785746 | 0.166787665 | 0.00031896 | 0.001994793 |
| ZBTB8A | 1032.479701 | 0.526726228 | 0.136352631 | 2.51E-05 | 0.000227749 |
| CLCA4-AS1 | 23.80999493 | 0.526641789 | 0.329105285 | 0.01063176 | 0.037366519 |
| VPREB3 | 7.771215266 | 0.526522566 | 0.5078374 | 0.01482762 | 0.048739623 |
| ANKRD13A | 2497.606765 | 0.526397809 | 0.090066091 | 1.29E-09 | 3.39E-08 |
| C11orf87 | 33.93822528 | 0.526029664 | 0.272654759 | 0.00674127 | 0.025648005 |
| ZNF287 | 778.680499 | 0.525954391 | 0.117820071 | 1.92E-06 | 2.36E-05 |

| | | | | | |
|---|---|---|---|---|---|
| CLEC18B | 23.15051606 | 0.525936046 | 0.318250353 | 0.0100531 | 0.035674197 |
| UBE2A | 2288.25687 | 0.525934178 | 0.082430228 | 4.59E-11 | 1.61E-09 |
| DRAIC | 16.62322539 | 0.525932139 | 0.373766336 | 0.01291312 | 0.043627806 |
| CYB5AP2 | 14.69879962 | 0.525910503 | 0.416444495 | 0.01420447 | 0.04701445 |
| DRAM1 | 198.3394345 | 0.525858111 | 0.162925504 | 0.00025626 | 0.001656096 |
| SH3BP5-AS1 | 908.1282001 | 0.525017802 | 0.130274821 | 1.29E-05 | 0.00012708 |
| PRDM6 | 10.21449798 | 0.525009078 | 0.484925568 | 0.01503766 | 0.04927253 |
| NADK2 | 1139.296833 | 0.524484716 | 0.109926759 | 4.50E-07 | 6.67E-06 |
| SLC48A1 | 403.2819827 | 0.524299243 | 0.178290849 | 0.00063879 | 0.003599882 |
| RRAD | 16.61917138 | 0.524299224 | 0.392587832 | 0.01379051 | 0.045940082 |
| RYR3 | 2108.851215 | 0.52417528 | 0.092275652 | 3.45E-09 | 8.32E-08 |
| CEP126 | 391.4716654 | 0.524139747 | 0.122712999 | 4.62E-06 | 5.11E-05 |
| ZNF674 | 275.1456237 | 0.524071383 | 0.126750008 | 8.35E-06 | 8.62E-05 |
| FBXO28 | 3271.848272 | 0.523941298 | 0.080759588 | 2.30E-11 | 8.62E-10 |
| BEND6 | 315.5300061 | 0.523685349 | 0.148768856 | 9.45E-05 | 0.000705224 |
| BCL2L10 | 5.020524787 | 0.523530895 | 0.617649987 | 0.01352409 | 0.045235821 |
| ZNF846 | 188.2443858 | 0.523484832 | 0.175043702 | 0.00055314 | 0.003181766 |
| OXNAD1 | 747.7239772 | 0.52342633 | 0.107551643 | 2.82E-07 | 4.40E-06 |
| TECPR2 | 797.9680034 | 0.523241862 | 0.14211547 | 5.21E-05 | 0.000422492 |
| COLGALT2 | 309.8023363 | 0.522999798 | 0.15766608 | 0.00019381 | 0.001305657 |
| RLF | 2584.259097 | 0.522841177 | 0.101692811 | 6.91E-08 | 1.27E-06 |
| GCNT3 | 10.21902756 | 0.522590059 | 0.51473461 | 0.0152069 | 0.049747816 |
| GCNA | 202.3509259 | 0.522023229 | 0.165989935 | 0.00034516 | 0.002135354 |
| BMP8A | 141.5657966 | 0.521166275 | 0.180545093 | 0.00076746 | 0.004199966 |
| NRSN2-AS1 | 478.5304371 | 0.521121133 | 0.139258702 | 4.19E-05 | 0.000352616 |
| RPS20P13 | 2.522811114 | 0.521042049 | 0.799237339 | 0.00816392 | 0.03006161 |
| ARF4 | 7051.237156 | 0.520548776 | 0.082157896 | 6.32E-11 | 2.17E-09 |
| HSCB | 381.3391122 | 0.520546854 | 0.144745715 | 7.31E-05 | 0.000564672 |
| UNC5B-AS1 | 24.56757882 | 0.520488745 | 0.350711551 | 0.01263475 | 0.042874765 |
| BRD2 | 16937.36211 | 0.520172548 | 0.141142328 | 5.24E-05 | 0.000424427 |
| STAB1 | 22.54164245 | 0.52000552 | 0.327881396 | 0.01143061 | 0.039615467 |
| KLHL36 | 114.1726863 | 0.519952566 | 0.230433183 | 0.00382579 | 0.016127576 |
| ADGRF5P1 | 93.41377101 | 0.519938709 | 0.209727292 | 0.00230544 | 0.010625064 |
| STOX2 | 3447.353368 | 0.519908361 | 0.096146394 | 1.67E-08 | 3.49E-07 |
| PRKAB1 | 2009.767465 | 0.519851149 | 0.131824476 | 1.90E-05 | 0.000179135 |
| SPEF2 | 352.2897103 | 0.519602436 | 0.15001922 | 0.0001192 | 0.000861304 |
| BBX | 2755.133992 | 0.519393928 | 0.108109898 | 3.95E-07 | 5.92E-06 |
| SIDT1 | 6.0061159 | 0.519117715 | 0.567685334 | 0.01488806 | 0.048907046 |
| DBX2 | 42.96656836 | 0.518779932 | 0.276815456 | 0.00784465 | 0.029085511 |
| NKIRAS1 | 968.3542548 | 0.518040665 | 0.101763608 | 9.29E-08 | 1.66E-06 |
| NOCT | 461.656703 | 0.517415859 | 0.171652564 | 0.00053845 | 0.003114656 |
| MTHFD1L | 3937.997447 | 0.517382051 | 0.114466103 | 1.56E-06 | 1.98E-05 |
| BCL2L13 | 2319.231154 | 0.51734891 | 0.101524256 | 9.08E-08 | 1.63E-06 |
| TMEM260 | 983.4956546 | 0.516797787 | 0.11756753 | 2.78E-06 | 3.28E-05 |
| RHBDF2 | 149.5909049 | 0.516736555 | 0.214489178 | 0.0027948 | 0.012477686 |
| HAVCR2 | 38.74708493 | 0.516677566 | 0.292685638 | 0.00938965 | 0.033765053 |
| DOCK9 | 1240.005352 | 0.516002003 | 0.127536044 | 1.28E-05 | 0.000126434 |
| ANKRD42 | 413.3728016 | 0.515609665 | 0.131658299 | 2.19E-05 | 0.00020136 |
| SERP1 | 5943.206351 | 0.515608067 | 0.080852706 | 4.98E-11 | 1.74E-09 |
| TM7SF2 | 809.8842641 | 0.515605616 | 0.128596778 | 1.49E-05 | 0.000144857 |

| | | | | | |
|---|---|---|---|---|---|
| DUSP4 | 2390.402207 | 0.515348961 | 0.156034523 | 0.00021496 | 0.001425019 |
| CD53 | 18.2751401 | 0.515057113 | 0.343057122 | 0.01301167 | 0.043849864 |
| ERBB3 | 20.2809635 | 0.514703263 | 0.369311342 | 0.01431615 | 0.047341285 |
| ANKRD13C-DT | 49.43278944 | 0.514691786 | 0.235658244 | 0.00464701 | 0.01892078 |
| ASB7 | 1511.586648 | 0.514503837 | 0.089334166 | 2.31E-09 | 5.76E-08 |
| ZNF550 | 777.0880084 | 0.514225143 | 0.119433964 | 4.23E-06 | 4.73E-05 |
| RPL13AP20 | 46.45660614 | 0.514047717 | 0.256754445 | 0.00660051 | 0.025229166 |
| DDX19A-DT | 14.02395005 | 0.513761288 | 0.36902696 | 0.01444484 | 0.047679562 |
| SSBP2 | 1095.699905 | 0.513752681 | 0.13797567 | 4.74E-05 | 0.000391303 |
| PPP1R26-AS1 | 93.64779666 | 0.513500771 | 0.176750797 | 0.00077004 | 0.00421032 |
| ESRRB | 58.08611493 | 0.513405963 | 0.315961124 | 0.01156958 | 0.039952444 |
| DNAJC3-DT | 200.7102625 | 0.513100247 | 0.169037591 | 0.00052079 | 0.003027001 |
| PODNL1 | 30.89105425 | 0.513089268 | 0.285356018 | 0.00924344 | 0.033365309 |
| ADAMTS9-AS2 | 113.980305 | 0.513058942 | 0.20047893 | 0.00199788 | 0.009438161 |
| DMXL1 | 2358.720982 | 0.513034122 | 0.097644925 | 4.02E-08 | 7.82E-07 |
| SYT4 | 4324.454701 | 0.512704773 | 0.104551245 | 2.50E-07 | 3.96E-06 |
| BTG1-DT | 31.81297382 | 0.51202615 | 0.284148391 | 0.00927602 | 0.033447687 |
| AURKC | 24.4173623 | 0.511674384 | 0.294383141 | 0.01017873 | 0.036032951 |
| TCIRG1 | 99.75199653 | 0.511659723 | 0.190975084 | 0.00147672 | 0.007289312 |
| MAPT-AS1 | 83.35501998 | 0.51156299 | 0.248333287 | 0.00607153 | 0.02355755 |
| CACNA1C | 971.3898277 | 0.511547193 | 0.138064161 | 5.16E-05 | 0.00041954 |
| SLC43A2 | 1015.285332 | 0.511434371 | 0.163769569 | 0.00040012 | 0.002421445 |
| JSRP1 | 38.15856508 | 0.511111451 | 0.271557102 | 0.00827592 | 0.030428403 |
| TAF7L | 42.46939204 | 0.510973193 | 0.250330038 | 0.00631665 | 0.024379817 |
| ARFGEF3 | 165.1752041 | 0.510925308 | 0.19850259 | 0.00195697 | 0.009283659 |
| ZNF510 | 2871.175955 | 0.510550904 | 0.131624428 | 2.63E-05 | 0.000236395 |
| PFN4 | 21.85647902 | 0.510456252 | 0.355938295 | 0.01436553 | 0.047455859 |
| ZFP37 | 759.4310487 | 0.510112766 | 0.100960278 | 1.19E-07 | 2.06E-06 |
| INSYN1 | 554.2353435 | 0.510106621 | 0.155809705 | 0.00024592 | 0.001599345 |
| HAP1 | 36.22169895 | 0.509045731 | 0.284368901 | 0.00968338 | 0.034602955 |
| GOLGA1 | 1139.629157 | 0.509035005 | 0.098917772 | 7.33E-08 | 1.34E-06 |
| BBOX1-AS1 | 37.90233346 | 0.508938896 | 0.25042655 | 0.00653197 | 0.025032334 |
| ZNF136 | 1028.813946 | 0.508855244 | 0.126147283 | 1.41E-05 | 0.000137821 |
| GRM1 | 96.56757569 | 0.508716747 | 0.216028264 | 0.0033805 | 0.014531061 |
| CCDC28A | 670.0157976 | 0.508542653 | 0.102738594 | 2.03E-07 | 3.31E-06 |
| RUNDC3B | 567.1357637 | 0.508402009 | 0.116044818 | 3.13E-06 | 3.64E-05 |
| PREPL | 5635.048426 | 0.508142269 | 0.094155105 | 1.90E-08 | 3.94E-07 |
| C8orf89 | 4.480951092 | 0.50797736 | 0.612768396 | 0.01481107 | 0.048708594 |
| MIR23AHG | 212.2878551 | 0.507908027 | 0.169788109 | 0.0006193 | 0.003501563 |
| NR2F1-AS1 | 82.3876873 | 0.507789443 | 0.20001041 | 0.00219213 | 0.010203464 |
| SERAC1 | 947.6847125 | 0.507759187 | 0.09547093 | 2.93E-08 | 5.86E-07 |
| TAL2 | 30.41563107 | 0.507343879 | 0.314087221 | 0.01232589 | 0.042041736 |
| UBOX5 | 407.864121 | 0.507312766 | 0.162802963 | 0.00042108 | 0.002532656 |
| CEP57 | 3397.775615 | 0.507241027 | 0.129708154 | 2.36E-05 | 0.000215664 |
| ABHD17AP4 | 4.282272361 | 0.506847228 | 0.662923049 | 0.0133708 | 0.044839704 |
| SLC35B3 | 1253.885971 | 0.506756933 | 0.107375461 | 6.47E-07 | 9.10E-06 |
| AMOTL2 | 1013.909157 | 0.506745145 | 0.193860987 | 0.00182284 | 0.008733643 |
| RGS3 | 1221.325598 | 0.506594759 | 0.155313882 | 0.00026253 | 0.00169234 |
| CLMN | 1709.779737 | 0.50612357 | 0.128851014 | 2.22E-05 | 0.000203525 |
| POLR3C | 3097.90327 | 0.50607767 | 0.08215341 | 2.11E-10 | 6.49E-09 |

| | | | | | |
|---|---|---|---|---|---|
| CCDC112 | 1771.421291 | 0.505652331 | 0.096970554 | 5.20E-08 | 9.85E-07 |
| LINC03048 | 66.97988038 | 0.505487428 | 0.302877185 | 0.01174716 | 0.040464174 |
| HERC4 | 2553.452257 | 0.505249794 | 0.108443305 | 8.74E-07 | 1.18E-05 |
| ZNF440 | 1147.203736 | 0.505170255 | 0.130541879 | 2.82E-05 | 0.000250728 |
| VPS37D | 204.4144241 | 0.505069655 | 0.168962962 | 0.00063634 | 0.003590052 |
| RNF103 | 860.5460983 | 0.504992467 | 0.106306688 | 5.62E-07 | 8.10E-06 |
| PDSS2 | 589.5795714 | 0.504984562 | 0.105239907 | 4.44E-07 | 6.61E-06 |
| FADS3 | 701.4848275 | 0.504426453 | 0.163306897 | 0.00046878 | 0.002773301 |
| DENND2A | 1000.503144 | 0.504220342 | 0.145953765 | 0.00013676 | 0.000973779 |
| BAMBI | 1824.780673 | 0.504205439 | 0.130304688 | 2.84E-05 | 0.000252535 |
| GAB2 | 3472.117971 | 0.504031962 | 0.180184564 | 0.00112899 | 0.005840638 |
| TTC7A | 534.7834759 | 0.504009591 | 0.142172248 | 9.88E-05 | 0.000730771 |
| ECM1 | 329.7127286 | 0.504001314 | 0.165986294 | 0.00055437 | 0.003187938 |
| LINC00680-GUSBP4 | 5.515312772 | 0.503795741 | 0.617534217 | 0.01500056 | 0.049182304 |
| LPAR1 | 1590.454765 | 0.503354449 | 0.123605803 | 1.24E-05 | 0.000122878 |
| MAP9 | 1554.204584 | 0.503269017 | 0.098278289 | 8.66E-08 | 1.56E-06 |
| LINC01140 | 17.83313997 | 0.503142871 | 0.336322674 | 0.01447397 | 0.04776039 |
| ATP6AP1-DT | 42.92491311 | 0.502896295 | 0.254374684 | 0.00758582 | 0.028290661 |
| MIR2052HG | 20.07257752 | 0.502790487 | 0.325247679 | 0.013825 | 0.046017684 |
| ZBTB34 | 1958.337942 | 0.502709599 | 0.099055205 | 1.10E-07 | 1.93E-06 |
| CDH3 | 122.0870903 | 0.502479739 | 0.182734246 | 0.00130517 | 0.0065961 |
| HDLBP | 19880.50287 | 0.502170402 | 0.140214867 | 8.74E-05 | 0.000657085 |
| SYNM | 1947.548745 | 0.501520346 | 0.112388722 | 2.25E-06 | 2.71E-05 |
| UHMK1 | 6440.358844 | 0.501491596 | 0.098012365 | 8.95E-08 | 1.61E-06 |
| PHLDA3 | 1022.832913 | 0.501254043 | 0.123808466 | 1.39E-05 | 0.000136237 |
| SLC22A17 | 3615.673055 | 0.500955301 | 0.16484791 | 0.00056205 | 0.003226671 |
| JHY | 125.9706328 | 0.500887299 | 0.171278373 | 0.00079769 | 0.004332698 |
| CCBE1 | 70.16021391 | 0.500821105 | 0.206017226 | 0.00301537 | 0.013275114 |
| ARHGAP23 | 1707.126452 | 0.500408722 | 0.172135836 | 0.00084288 | 0.004543368 |
| CTBS | 280.3715273 | 0.500234595 | 0.128744076 | 2.73E-05 | 0.00024459 |
| NXF1 | 4567.262199 | 0.500059431 | 0.147852048 | 0.00018164 | 0.001240129 |
| NGF | 170.2058215 | 0.499954211 | 0.158006146 | 0.0003796 | 0.002317696 |
| TEKT2 | 86.73241873 | 0.499842353 | 0.20669777 | 0.00313207 | 0.0136805 |
| GPR68 | 49.32928053 | 0.49964944 | 0.265325869 | 0.00911835 | 0.033006453 |
| PAQR9 | 358.9872209 | 0.49941273 | 0.122396131 | 1.23E-05 | 0.000121957 |
| SNHG11 | 558.6394948 | 0.499110562 | 0.112133285 | 2.41E-06 | 2.88E-05 |
| DGKA | 208.4445013 | 0.498974241 | 0.151527952 | 0.00024885 | 0.001614826 |
| TSPO2 | 25.16145082 | 0.498956879 | 0.294335508 | 0.01202545 | 0.041229707 |
| ZFP3 | 1311.412751 | 0.498901345 | 0.105563432 | 6.56E-07 | 9.21E-06 |
| TRA2A | 5762.120673 | 0.498818784 | 0.113110684 | 2.91E-06 | 3.42E-05 |
| CYP51A1P2 | 25.07831549 | 0.498689905 | 0.328823637 | 0.01474648 | 0.048527223 |
| CFAP418 | 156.892099 | 0.498064682 | 0.162858278 | 0.00053975 | 0.003121295 |
| AAK1 | 2115.841157 | 0.498010238 | 0.133206668 | 4.95E-05 | 0.000404965 |
| EIF5 | 14342.43395 | 0.4980033 | 0.084193821 | 1.00E-09 | 2.68E-08 |
| RICTOR | 2188.803343 | 0.497947024 | 0.112232277 | 2.59E-06 | 3.07E-05 |
| UBE2J1 | 4166.821294 | 0.497898195 | 0.07900239 | 8.93E-11 | 2.95E-09 |
| CDH20 | 32.55380551 | 0.497779784 | 0.320998012 | 0.01436363 | 0.047455859 |
| USP30 | 1596.159395 | 0.497715121 | 0.082527189 | 4.94E-10 | 1.41E-08 |
| ATXN2 | 3453.580021 | 0.49761167 | 0.166501149 | 0.00067213 | 0.003761017 |
| TCTE1 | 117.717734 | 0.497483902 | 0.218553007 | 0.00440897 | 0.018111494 |

| | | | | | |
|---|---|---|---|---|---|
| DEPTOR | 63.55467219 | 0.497454069 | 0.229139637 | 0.00550605 | 0.021710289 |
| RCBTB2 | 780.222368 | 0.497241125 | 0.113672101 | 3.45E-06 | 3.95E-05 |
| ATE1 | 2370.988175 | 0.497228642 | 0.108711427 | 1.37E-06 | 1.77E-05 |
| FAM184A | 2706.211386 | 0.496196442 | 0.108855185 | 1.49E-06 | 1.90E-05 |
| DUSP15 | 87.95743805 | 0.496137884 | 0.213049636 | 0.00396659 | 0.016611889 |
| ZNF503 | 1614.849071 | 0.496084554 | 0.179828416 | 0.00133446 | 0.006721006 |
| PER2 | 1017.952352 | 0.49587863 | 0.155725466 | 0.00036536 | 0.002243402 |
| RBM5 | 6965.632309 | 0.495844824 | 0.102369688 | 3.74E-07 | 5.63E-06 |
| FBXO38 | 2209.638995 | 0.495756314 | 0.100382177 | 2.32E-07 | 3.72E-06 |
| HCRTR1 | 47.22822094 | 0.495741882 | 0.296642793 | 0.01274647 | 0.043182593 |
| S1PR2 | 156.7331253 | 0.495177779 | 0.205903515 | 0.00335229 | 0.014438576 |
| UFSP2 | 834.0586985 | 0.495167217 | 0.108566664 | 1.48E-06 | 1.89E-05 |
| RUNX1 | 138.0894693 | 0.495012062 | 0.222138365 | 0.00498509 | 0.020061331 |
| SLC25A47 | 3.259213403 | 0.494930005 | 0.751478003 | 0.01052692 | 0.037074091 |
| AKR1C3 | 23.9037966 | 0.494909141 | 0.313757715 | 0.01431703 | 0.047341285 |
| FAM169A | 1072.510626 | 0.494865975 | 0.112310536 | 3.03E-06 | 3.54E-05 |
| HSPB1 | 1306.718375 | 0.494668818 | 0.287950835 | 0.01211641 | 0.041457862 |
| TXNL4B | 526.5082482 | 0.494568102 | 0.106821133 | 1.07E-06 | 1.42E-05 |
| GPX3 | 239.5567067 | 0.494465279 | 0.19166714 | 0.00218662 | 0.010182433 |
| MAGED2 | 3871.452479 | 0.494234611 | 0.080284651 | 2.31E-10 | 7.09E-09 |
| RNF10 | 5080.49913 | 0.494176075 | 0.179786171 | 0.00139131 | 0.006942684 |
| CCDC191 | 58.40817657 | 0.494172542 | 0.235760914 | 0.00656117 | 0.025120795 |
| INTS12 | 867.6990646 | 0.493706913 | 0.101616437 | 3.52E-07 | 5.33E-06 |
| CDC14B | 1558.830016 | 0.493422762 | 0.137148529 | 8.71E-05 | 0.000655027 |
| TMTC1 | 473.0366315 | 0.493257746 | 0.154074231 | 0.00035145 | 0.002169006 |
| GNB5 | 920.870113 | 0.492789631 | 0.112575596 | 3.50E-06 | 3.99E-05 |
| C2CD6 | 19.57233646 | 0.492545098 | 0.314812961 | 0.01481432 | 0.04871085 |
| ANKHD1-EIF4EBP3 | 163.3838263 | 0.492479767 | 0.154927783 | 0.00038066 | 0.002322128 |
| SNHG32 | 2003.525926 | 0.492015828 | 0.101410529 | 3.67E-07 | 5.55E-06 |
| PPIB | 4098.269644 | 0.491602078 | 0.098832339 | 1.98E-07 | 3.24E-06 |
| MGARP | 60.65782765 | 0.490874641 | 0.222047742 | 0.00534997 | 0.021214185 |
| CEP95 | 9674.301455 | 0.490581425 | 0.09331087 | 4.49E-08 | 8.65E-07 |
| ALG5 | 1320.171262 | 0.490525375 | 0.088597952 | 9.58E-09 | 2.11E-07 |
| ARRDC4 | 1439.943778 | 0.490400342 | 0.092919203 | 4.03E-08 | 7.84E-07 |
| MUC20P1 | 34.4811288 | 0.490367468 | 0.309632606 | 0.01479043 | 0.048648499 |
| PBX4 | 143.8193779 | 0.490302101 | 0.188707805 | 0.00214996 | 0.010045843 |
| ST20-AS1 | 183.4493394 | 0.490124419 | 0.174060108 | 0.00118396 | 0.006082155 |
| ADAM32 | 57.8719578 | 0.48974593 | 0.236128972 | 0.00709783 | 0.026761977 |
| TIGD4 | 69.45658044 | 0.48958164 | 0.21159363 | 0.00431961 | 0.017819733 |
| PPFIBP1 | 2066.644798 | 0.489386061 | 0.117311467 | 8.86E-06 | 9.05E-05 |
| ENC1 | 2371.280716 | 0.489342439 | 0.102651187 | 5.67E-07 | 8.16E-06 |
| ZCCHC2 | 591.7419177 | 0.489328085 | 0.125549703 | 2.79E-05 | 0.000248423 |
| UBR3 | 4972.387701 | 0.489289819 | 0.09424025 | 6.43E-08 | 1.19E-06 |
| MORC3 | 1717.289217 | 0.489106476 | 0.111162653 | 3.23E-06 | 3.73E-05 |
| LRRC46 | 80.93126441 | 0.488801786 | 0.212492357 | 0.00447891 | 0.018354421 |
| TCERG1L | 100.8401299 | 0.48874635 | 0.201884364 | 0.00339091 | 0.014566677 |
| RSRC2 | 7343.229101 | 0.488593618 | 0.085861452 | 3.99E-09 | 9.48E-08 |
| NFKBIL1 | 550.6217981 | 0.488519342 | 0.140889677 | 0.00014472 | 0.001021998 |
| PRDX1P1 | 4.303531717 | 0.488501579 | 0.654008991 | 0.01483761 | 0.048749083 |
| LINC02820 | 113.152877 | 0.488407442 | 0.18123637 | 0.00168305 | 0.008155074 |

| | | | | | |
|---|---|---|---|---|---|
| PDCL3P4 | 309.0635889 | 0.488362092 | 0.16639204 | 0.00084361 | 0.004544928 |
| WDR31 | 156.9863901 | 0.488053735 | 0.167181746 | 0.0008863 | 0.004750008 |
| ADPRM | 288.1713614 | 0.488034795 | 0.155506767 | 0.00044736 | 0.002663431 |
| COPG1 | 5931.567504 | 0.487288606 | 0.118594287 | 1.17E-05 | 0.000116799 |
| CEP76 | 937.3276186 | 0.487103547 | 0.108091756 | 2.00E-06 | 2.44E-05 |
| KIN | 960.0599837 | 0.486905359 | 0.099808855 | 3.31E-07 | 5.07E-06 |
| TAMM41 | 517.6505355 | 0.486616334 | 0.122799883 | 2.17E-05 | 0.000200384 |
| DCAF4L1 | 123.9523524 | 0.486511862 | 0.202564084 | 0.00361002 | 0.015357908 |
| SUPT20H | 2746.18811 | 0.486135453 | 0.098892675 | 2.75E-07 | 4.30E-06 |
| NFX1 | 3950.71959 | 0.485975582 | 0.087565292 | 9.11E-09 | 2.01E-07 |
| GOLGA6A | 3.604880978 | 0.48589254 | 0.677307018 | 0.01404116 | 0.046616475 |
| ZNF571 | 352.0824876 | 0.485545513 | 0.16090739 | 0.00066771 | 0.003741354 |
| SAXO2 | 77.38643876 | 0.48548659 | 0.233832971 | 0.00731971 | 0.027437568 |
| TTBK2 | 3844.036275 | 0.485464433 | 0.121111654 | 1.81E-05 | 0.000171729 |
| BCAS2 | 3169.9181 | 0.485413273 | 0.086026407 | 5.37E-09 | 1.24E-07 |
| PEA15 | 2316.834597 | 0.485347213 | 0.120020392 | 1.56E-05 | 0.00015057 |
| AGPAT4 | 3567.821539 | 0.484829337 | 0.093301732 | 6.44E-08 | 1.19E-06 |
| ARHGEF37 | 168.904377 | 0.484663954 | 0.167283651 | 0.00096888 | 0.005123158 |
| MCPH1-AS1 | 93.81150158 | 0.484635154 | 0.21064756 | 0.00462281 | 0.0188431 |
| PRNP | 2159.904986 | 0.484596538 | 0.102102365 | 6.46E-07 | 9.09E-06 |
| INTS13 | 1840.377907 | 0.484496465 | 0.105467866 | 1.35E-06 | 1.74E-05 |
| NCOA7 | 1647.656612 | 0.484213165 | 0.137236715 | 0.00011919 | 0.000861304 |
| LIN7B | 236.439643 | 0.483974702 | 0.158149944 | 0.00059018 | 0.003360945 |
| EPG5 | 2232.340937 | 0.483941933 | 0.102228888 | 6.88E-07 | 9.58E-06 |
| PSD2 | 187.9781531 | 0.483916431 | 0.168361691 | 0.00104254 | 0.005448239 |
| AIFM2 | 124.5659066 | 0.483840937 | 0.19425731 | 0.00296702 | 0.013087452 |
| LINC02803 | 83.62068899 | 0.483680243 | 0.189339231 | 0.0025273 | 0.011439899 |
| NFU1 | 708.8608656 | 0.483543733 | 0.103843901 | 1.00E-06 | 1.34E-05 |
| FAM135A | 2218.854135 | 0.483079665 | 0.108707236 | 2.73E-06 | 3.23E-05 |
| TMLHE | 294.8717212 | 0.483004713 | 0.132859757 | 8.06E-05 | 0.000613865 |
| FGFR1OP2 | 2250.521744 | 0.482647646 | 0.103732762 | 1.03E-06 | 1.37E-05 |
| PCYT1A | 2667.624767 | 0.482607248 | 0.082770972 | 1.80E-09 | 4.59E-08 |
| SLC7A11-AS1 | 3.176278271 | 0.482599329 | 0.734762563 | 0.01162021 | 0.040086889 |
| TRAPPC8 | 2579.068446 | 0.482533899 | 0.092166029 | 5.29E-08 | 1.00E-06 |
| HECTD2 | 2257.992668 | 0.482388954 | 0.123108718 | 2.67E-05 | 0.000239235 |
| RBM48 | 1055.189662 | 0.482378052 | 0.106893525 | 1.99E-06 | 2.44E-05 |
| MIGA1 | 1846.61847 | 0.482160425 | 0.092971567 | 6.90E-08 | 1.27E-06 |
| BEND4 | 3631.130217 | 0.481852224 | 0.105330596 | 1.50E-06 | 1.91E-05 |
| CCDC17 | 54.1582484 | 0.481728893 | 0.250784319 | 0.00997943 | 0.035449488 |
| GADD45A | 3356.874174 | 0.481375939 | 0.166047393 | 0.00098553 | 0.00519917 |
| NEK4 | 1781.151766 | 0.481276657 | 0.101440817 | 6.63E-07 | 9.28E-06 |
| APTR | 529.1577627 | 0.481093748 | 0.125290248 | 3.69E-05 | 0.00031534 |
| MFSD8 | 1438.306199 | 0.48064943 | 0.12751376 | 4.89E-05 | 0.000400958 |
| UBALD2 | 2953.374555 | 0.480505172 | 0.157708969 | 0.00062998 | 0.003556116 |
| CDC25C | 883.7548198 | 0.480359767 | 0.120838568 | 2.14E-05 | 0.000198061 |
| PAK3 | 1609.295933 | 0.480259339 | 0.106788771 | 2.17E-06 | 2.62E-05 |
| SHROOM3-AS1 | 3.042382087 | 0.479266055 | 0.727729727 | 0.01205896 | 0.041309419 |
| RAD52 | 1230.339273 | 0.479235575 | 0.102278535 | 8.93E-07 | 1.21E-05 |
| MAP2K4 | 2294.295339 | 0.478920555 | 0.084954438 | 5.72E-09 | 1.31E-07 |
| GOLGA6L5P | 69.55732064 | 0.478810243 | 0.193600762 | 0.00321599 | 0.013934589 |

| | | | | | |
|---|---|---|---|---|---|
| TNFRSF10B | 6230.318763 | 0.478596531 | 0.101138722 | 7.15E-07 | 9.90E-06 |
| RP9 | 786.3099542 | 0.478160911 | 0.117801766 | 1.53E-05 | 0.000147646 |
| RAB3GAP2 | 7247.251728 | 0.477761189 | 0.086570267 | 1.13E-08 | 2.47E-07 |
| TAF1A | 1583.170417 | 0.477652575 | 0.100943032 | 7.19E-07 | 9.93E-06 |
| MCCC1-AS1 | 58.26889977 | 0.477619364 | 0.234438996 | 0.00839885 | 0.030792285 |
| CRTAC1 | 383.5101281 | 0.477507459 | 0.150142417 | 0.00041745 | 0.002513016 |
| LINC01778 | 2.395604283 | 0.477270458 | 0.768124028 | 0.01015244 | 0.035946079 |
| LRRTM2 | 1303.770643 | 0.477038561 | 0.142730334 | 0.00024199 | 0.001577295 |
| TWF2-DT | 50.38046674 | 0.477003532 | 0.253959558 | 0.01113614 | 0.038749215 |
| PJA1 | 2338.770263 | 0.476803479 | 0.08345231 | 3.72E-09 | 8.91E-08 |
| DEF6 | 59.54460758 | 0.476692591 | 0.232703432 | 0.00828583 | 0.030459387 |
| SOS2 | 2453.111423 | 0.476640934 | 0.092708636 | 9.02E-08 | 1.62E-06 |
| KCNK12 | 356.0495348 | 0.476607185 | 0.151530088 | 0.00047104 | 0.002785011 |
| STARD8 | 52.28027323 | 0.47605115 | 0.273873341 | 0.01384244 | 0.046045942 |
| PNRC1 | 1098.327767 | 0.47572598 | 0.152473209 | 0.00051416 | 0.002995257 |
| SGCD | 607.9275206 | 0.475387593 | 0.164740249 | 0.00106742 | 0.005562089 |
| OSBPL2 | 1287.689562 | 0.474810223 | 0.111002886 | 6.06E-06 | 6.49E-05 |
| ZNF699 | 335.612937 | 0.474625476 | 0.13266137 | 0.00010526 | 0.000773318 |
| KRTAP5-8 | 3.500489985 | 0.474168403 | 0.73797644 | 0.0117163 | 0.040391264 |
| ZNF805 | 571.8381923 | 0.473376636 | 0.119380072 | 2.32E-05 | 0.000212107 |
| POLD4 | 34.92344041 | 0.473192322 | 0.26731156 | 0.01356159 | 0.045324315 |
| ZNF165 | 39.554871 | 0.473057173 | 0.231889632 | 0.00866928 | 0.031614646 |
| NCK1 | 871.6462837 | 0.472676902 | 0.103562955 | 1.65E-06 | 2.08E-05 |
| ZNF577 | 718.2525155 | 0.472569669 | 0.12218417 | 3.47E-05 | 0.000299894 |
| MED8 | 844.6726091 | 0.471707424 | 0.107208593 | 3.56E-06 | 4.05E-05 |
| TRANK1 | 167.8920322 | 0.471628809 | 0.201088416 | 0.00460717 | 0.018786801 |
| MUC20 | 453.710704 | 0.471265501 | 0.193143991 | 0.00368661 | 0.015615113 |
| TAF2 | 3287.135601 | 0.471235809 | 0.092091317 | 1.05E-07 | 1.86E-06 |
| GALNT6 | 756.7238589 | 0.471122953 | 0.145973 | 0.00037141 | 0.002276505 |
| REV3L | 3143.381951 | 0.471032844 | 0.107753392 | 4.06E-06 | 4.57E-05 |
| AKAP6 | 826.6959924 | 0.470760891 | 0.138006236 | 0.00019737 | 0.001326365 |
| SIK2 | 1560.782676 | 0.470494718 | 0.126734414 | 6.47E-05 | 0.000508797 |
| SAP130 | 2487.84715 | 0.470375343 | 0.180670892 | 0.00244462 | 0.011139192 |
| ATG4A | 260.976643 | 0.470236641 | 0.128918499 | 8.31E-05 | 0.000629694 |
| GABRE | 296.9413896 | 0.469969317 | 0.150313182 | 0.00052196 | 0.003032951 |
| USP6 | 116.512888 | 0.469823805 | 0.215726984 | 0.00676118 | 0.025709497 |
| ZIM2-AS1 | 3.414630722 | 0.469371527 | 0.727506989 | 0.01239598 | 0.042223215 |
| KPNA4 | 4054.445131 | 0.469259793 | 0.095880259 | 3.36E-07 | 5.11E-06 |
| IDE | 3407.906559 | 0.468746496 | 0.094411083 | 2.35E-07 | 3.76E-06 |
| RETSAT | 1064.79835 | 0.468462792 | 0.114499315 | 1.41E-05 | 0.000137596 |
| EIF2S2 | 5215.999886 | 0.468295937 | 0.091065041 | 9.36E-08 | 1.67E-06 |
| SLC39A7 | 3621.644365 | 0.468033096 | 0.106915013 | 4.02E-06 | 4.53E-05 |
| SYF2P2 | 3.052891455 | 0.467998862 | 0.716062504 | 0.01304113 | 0.04393477 |
| PKD1P6 | 700.845088 | 0.467747811 | 0.191559501 | 0.00376396 | 0.015907283 |
| SPRYD7 | 786.4363848 | 0.467507274 | 0.10679928 | 4.03E-06 | 4.54E-05 |
| ELMO1 | 269.6396437 | 0.467314257 | 0.162098462 | 0.00113462 | 0.005862413 |
| HSP90B2P | 80.67100799 | 0.467089787 | 0.190092032 | 0.00364081 | 0.015454609 |
| COX19 | 880.4073411 | 0.467076847 | 0.104162683 | 2.48E-06 | 2.96E-05 |
| POLR3GL | 188.3068726 | 0.466925632 | 0.167551794 | 0.00150753 | 0.007413594 |
| THUMPD2 | 1651.098719 | 0.466887693 | 0.097096151 | 5.22E-07 | 7.58E-06 |

| | | | | | |
|---|---|---|---|---|---|
| CDNF | 73.15304852 | 0.466448064 | 0.23286052 | 0.00978892 | 0.034923533 |
| SDHAP3 | 58.40024274 | 0.466134528 | 0.223769937 | 0.00843064 | 0.030897816 |
| OSBPL6 | 1218.24189 | 0.466078697 | 0.094064531 | 2.51E-07 | 3.97E-06 |
| ZEB1 | 7058.113448 | 0.465864841 | 0.093804418 | 2.37E-07 | 3.78E-06 |
| SLMAP | 2238.443487 | 0.465801114 | 0.094086594 | 2.57E-07 | 4.04E-06 |
| COL11A2 | 181.2293206 | 0.465780487 | 0.182966372 | 0.00293609 | 0.012970516 |
| TENM1 | 293.2983665 | 0.465649458 | 0.148200781 | 0.0005098 | 0.002975725 |
| ZC2HC1A | 1373.539985 | 0.465344639 | 0.1306443 | 0.00011785 | 0.00085305 |
| ARID3B | 1562.573558 | 0.465268542 | 0.175041166 | 0.0021857 | 0.010180479 |
| TRIM2 | 2903.241531 | 0.465237551 | 0.093652212 | 2.36E-07 | 3.77E-06 |
| PDE9A | 139.4766215 | 0.465055459 | 0.200814164 | 0.0051763 | 0.020644977 |
| KMT2C | 6290.357468 | 0.464960578 | 0.116441542 | 2.17E-05 | 0.000200239 |
| PARGP1 | 180.7495376 | 0.464800542 | 0.178147087 | 0.00250224 | 0.011346425 |
| FOCAD | 2045.264801 | 0.464296805 | 0.101066867 | 1.50E-06 | 1.91E-05 |
| ZNF707 | 540.9756176 | 0.463289459 | 0.1452121 | 0.00044085 | 0.002633181 |
| PPARGC1A | 1456.843023 | 0.462792819 | 0.096147088 | 5.21E-07 | 7.56E-06 |
| REPS1 | 2694.684276 | 0.462789132 | 0.088498215 | 6.06E-08 | 1.13E-06 |
| LINC01569 | 70.21531962 | 0.462516809 | 0.197265894 | 0.00493233 | 0.019887926 |
| PCNX2 | 3097.596351 | 0.462505859 | 0.136423297 | 0.00022319 | 0.001470491 |
| P4HB | 16849.11479 | 0.462309943 | 0.127722369 | 9.66E-05 | 0.00071701 |
| CCDC168 | 39.95405984 | 0.46218619 | 0.233699691 | 0.01060748 | 0.037313061 |
| SHB | 754.4788682 | 0.46216069 | 0.16525321 | 0.00151109 | 0.007426892 |
| CPB2-AS1 | 159.0933354 | 0.461969454 | 0.193019604 | 0.00441121 | 0.018117049 |
| TGFBI | 3.737502609 | 0.46182691 | 0.744975434 | 0.0116633 | 0.040222053 |
| TVP23B | 1712.968413 | 0.461823423 | 0.100224838 | 1.42E-06 | 1.83E-05 |
| C1orf52 | 2192.457472 | 0.461766731 | 0.09635562 | 5.81E-07 | 8.32E-06 |
| SCN8A | 3339.576911 | 0.461750195 | 0.130163763 | 0.00012693 | 0.000909136 |
| UBQLN1-AS1 | 55.15822277 | 0.461710195 | 0.2061567 | 0.00629601 | 0.024322974 |
| NAGS | 207.5795864 | 0.461442222 | 0.181098269 | 0.00300665 | 0.013242396 |
| TSC22D1 | 6318.438298 | 0.461321423 | 0.088050706 | 5.78E-08 | 1.08E-06 |
| ABHD18 | 1025.180402 | 0.460762806 | 0.12478234 | 7.38E-05 | 0.000569259 |
| EIF2S2P4 | 160.1670247 | 0.460682222 | 0.171256604 | 0.00206375 | 0.009711835 |
| OSTM1 | 1489.408117 | 0.460587112 | 0.095058181 | 4.49E-07 | 6.67E-06 |
| CHM | 1226.299501 | 0.4604251 | 0.105528035 | 4.47E-06 | 4.96E-05 |
| NSUN6 | 854.3941338 | 0.4602035 | 0.120706378 | 4.63E-05 | 0.000384076 |
| NUDT4 | 2984.471673 | 0.459934634 | 0.128315952 | 0.00011178 | 0.000814565 |
| TBC1D22B | 896.9097649 | 0.458957591 | 0.120725876 | 4.87E-05 | 0.000400297 |
| XPOT | 14813.60062 | 0.458908831 | 0.093558602 | 3.36E-07 | 5.11E-06 |
| LMF1 | 341.3261472 | 0.458852978 | 0.159950225 | 0.00124906 | 0.006357837 |
| DNAJB1 | 3853.818448 | 0.458708485 | 0.132635392 | 0.0001788 | 0.001223595 |
| SIK3 | 2228.336076 | 0.458666074 | 0.133348507 | 0.00019138 | 0.00129201 |
| ALKBH1 | 861.3091702 | 0.458599646 | 0.095707922 | 5.93E-07 | 8.44E-06 |
| TMEM269 | 160.0967334 | 0.458464892 | 0.168715018 | 0.00193986 | 0.009221283 |
| TLCD4 | 411.1413986 | 0.458285429 | 0.14791482 | 0.00061445 | 0.003480838 |
| ITGB5 | 94.30116722 | 0.458115353 | 0.234710325 | 0.01147582 | 0.039715373 |
| BAAT | 82.30158993 | 0.457208011 | 0.216077013 | 0.00845679 | 0.030960563 |
| CSRNP2 | 3827.649345 | 0.457149064 | 0.097688402 | 1.03E-06 | 1.38E-05 |
| MEIOC | 293.5262097 | 0.457068296 | 0.135009718 | 0.00023423 | 0.001534943 |
| PPP1R12A | 4448.315855 | 0.456965248 | 0.093021414 | 3.26E-07 | 5.01E-06 |
| CABP1 | 50.35801871 | 0.456756403 | 0.235325788 | 0.01182558 | 0.04066575 |

| | | | | | |
|---|---|---|---|---|---|
| TRPC3 | 118.945121 | 0.456548954 | 0.17568513 | 0.00271875 | 0.012169871 |
| LMAN1 | 8022.846389 | 0.456287771 | 0.099721556 | 1.71E-06 | 2.14E-05 |
| FXYD2 | 79.83177254 | 0.456116299 | 0.249398354 | 0.01432638 | 0.047364577 |
| ERAP2 | 49.93251596 | 0.455947676 | 0.234005923 | 0.01174543 | 0.040464174 |
| BRICD5 | 332.8062825 | 0.45591852 | 0.163947025 | 0.00164588 | 0.007999506 |
| IRF5 | 119.6517683 | 0.455709908 | 0.196085004 | 0.00543253 | 0.021487705 |
| CYP27A1 | 61.55255307 | 0.455597588 | 0.220973367 | 0.00953587 | 0.034174898 |
| CLASP2 | 12694.5786 | 0.455447702 | 0.107047327 | 7.44E-06 | 7.79E-05 |
| RAB11FIP1 | 754.5028234 | 0.455330566 | 0.131107694 | 0.00017292 | 0.001189653 |
| ZNF264 | 1125.48423 | 0.455011026 | 0.115203002 | 2.74E-05 | 0.000244707 |
| EIF1 | 16929.11489 | 0.454891935 | 0.126008249 | 0.00010443 | 0.000768024 |
| FOXL1 | 3.358720202 | 0.454451372 | 0.727974065 | 0.01282853 | 0.043398236 |
| SUGP1 | 1819.314102 | 0.453573911 | 0.121042654 | 6.21E-05 | 0.000490798 |
| SPINT2 | 66.77177279 | 0.4535712 | 0.216881116 | 0.00913428 | 0.033042831 |
| ARHGEF40 | 1613.923424 | 0.45305283 | 0.172571796 | 0.00258957 | 0.011676942 |
| NECTIN3 | 733.1464845 | 0.453009842 | 0.104505037 | 5.27E-06 | 5.75E-05 |
| TBC1D12 | 746.9968472 | 0.452907325 | 0.115933945 | 3.30E-05 | 0.000287111 |
| LAMB2P1 | 53.86837139 | 0.452525933 | 0.232335347 | 0.01208251 | 0.041376294 |
| EEFSEC | 790.5370878 | 0.452358994 | 0.120462353 | 6.06E-05 | 0.000480343 |
| FAM227B | 155.2123308 | 0.452178174 | 0.153605807 | 0.00103724 | 0.005426066 |
| RC3H1 | 4045.246722 | 0.452159518 | 0.085934104 | 5.36E-08 | 1.01E-06 |
| RGMB | 1182.935117 | 0.452112556 | 0.136697691 | 0.00031654 | 0.001983311 |
| EIF2AK3-DT | 73.38585727 | 0.452102747 | 0.209732589 | 0.00809452 | 0.029849125 |
| ARHGAP24 | 110.1169085 | 0.452043978 | 0.175197145 | 0.00294154 | 0.012990775 |
| PDIA2 | 603.0179225 | 0.452011049 | 0.15298899 | 0.00100444 | 0.005278651 |
| LATS1 | 5417.173273 | 0.45200364 | 0.087487284 | 8.94E-08 | 1.61E-06 |
| MLH3 | 66.06701769 | 0.451666762 | 0.21181849 | 0.00852412 | 0.031162653 |
| SLC2A11 | 507.3950465 | 0.451534354 | 0.140665834 | 0.00044257 | 0.002640405 |
| TENT2 | 2485.896638 | 0.450886784 | 0.098128139 | 1.60E-06 | 2.02E-05 |
| SLC30A5 | 2414.96537 | 0.450872667 | 0.093668588 | 5.53E-07 | 7.98E-06 |
| SCOC | 3639.565266 | 0.450751746 | 0.10566396 | 7.25E-06 | 7.61E-05 |
| CBLB | 2005.42441 | 0.450704207 | 0.114793816 | 3.08E-05 | 0.000271009 |
| RRAS2 | 990.5921602 | 0.450657503 | 0.103375622 | 4.78E-06 | 5.26E-05 |
| TUT7 | 1056.150497 | 0.450537942 | 0.117342824 | 4.38E-05 | 0.000366286 |
| ZXDB | 863.2124825 | 0.450332455 | 0.104257288 | 5.73E-06 | 6.18E-05 |
| MRPS31 | 1051.645123 | 0.450331545 | 0.118114786 | 4.88E-05 | 0.000400958 |
| METTL15 | 943.0380866 | 0.45030809 | 0.100990408 | 3.04E-06 | 3.54E-05 |
| KIDINS220 | 10590.01155 | 0.450259147 | 0.086594874 | 7.56E-08 | 1.38E-06 |
| INAFM2 | 2074.287929 | 0.450108724 | 0.129861264 | 0.00018251 | 0.001244006 |
| GDAP1 | 5863.258126 | 0.449853354 | 0.082670297 | 2.02E-08 | 4.15E-07 |
| DZIP3 | 2171.390362 | 0.449842342 | 0.118366624 | 5.14E-05 | 0.000418347 |
| ANAPC10 | 252.8684861 | 0.449713414 | 0.12818739 | 0.00015684 | 0.001095565 |
| SLC39A14 | 4530.330596 | 0.449640205 | 0.087519837 | 1.05E-07 | 1.86E-06 |
| SAFB2 | 3579.492805 | 0.449490279 | 0.127736234 | 0.00015102 | 0.001060256 |
| FBXO48 | 293.2744481 | 0.449490159 | 0.160000064 | 0.00157745 | 0.007716151 |
| ZYX | 1928.414007 | 0.449139913 | 0.194681601 | 0.0059178 | 0.023065514 |
| FGGY-DT | 55.68888715 | 0.449072461 | 0.244534215 | 0.01499135 | 0.049159961 |
| SLIT3-AS2 | 70.71469034 | 0.448949419 | 0.195100894 | 0.0060075 | 0.023362005 |
| NUMB | 2185.416656 | 0.448785574 | 0.100525522 | 2.98E-06 | 3.49E-05 |
| NCOA1 | 2019.706939 | 0.44858673 | 0.107228997 | 1.05E-05 | 0.000105868 |

| | | | | | |
|---|---|---|---|---|---|
| SLC10A5 | 41.51517096 | 0.44813118 | 0.233520753 | 0.01315583 | 0.044248695 |
| COG5 | 1334.782114 | 0.448126074 | 0.109547292 | 1.57E-05 | 0.00015147 |
| SUCLA2 | 2188.812779 | 0.447734589 | 0.106983867 | 1.05E-05 | 0.000105634 |
| KDM4D | 143.6092691 | 0.447655657 | 0.162542223 | 0.00187178 | 0.008945232 |
| LIAS | 978.6755949 | 0.447525777 | 0.108481939 | 1.36E-05 | 0.000133536 |
| SYTL1 | 80.22388347 | 0.447387245 | 0.220052759 | 0.01073076 | 0.037656545 |
| NUFIP2 | 9177.303308 | 0.447376075 | 0.089356163 | 2.11E-07 | 3.42E-06 |
| CD101 | 366.7983157 | 0.447268924 | 0.125490057 | 0.00012926 | 0.000924873 |
| PDCD4-AS1 | 193.1282486 | 0.447191076 | 0.17832156 | 0.00366923 | 0.01555113 |
| SNTB2 | 1505.713043 | 0.446702484 | 0.110495324 | 1.94E-05 | 0.000182278 |
| PLIN5 | 91.54833166 | 0.446690662 | 0.237374128 | 0.01418274 | 0.046981561 |
| WDR20 | 1256.88258 | 0.446684647 | 0.095144575 | 1.01E-06 | 1.35E-05 |
| POFUT2 | 1033.455557 | 0.446444972 | 0.137286523 | 0.00039533 | 0.002398109 |
| FSIP2 | 36.87159453 | 0.44640543 | 0.241688399 | 0.01505711 | 0.049328416 |
| UBR5 | 10294.38897 | 0.446177168 | 0.087075922 | 1.15E-07 | 2.01E-06 |
| GOSR2 | 4226.804024 | 0.445845897 | 0.091697474 | 4.44E-07 | 6.61E-06 |
| LCTL | 86.28194986 | 0.445812175 | 0.220533707 | 0.01110324 | 0.038666723 |
| ANKZF1 | 1893.537448 | 0.445394176 | 0.093479679 | 7.23E-07 | 9.96E-06 |
| STRC | 76.00082892 | 0.445365825 | 0.191211279 | 0.00576811 | 0.022576223 |
| MAP3K2-DT | 66.52223287 | 0.445172528 | 0.220588839 | 0.01122981 | 0.039015011 |
| RAB21 | 3467.764627 | 0.444753893 | 0.105294655 | 9.00E-06 | 9.18E-05 |
| KLC4 | 1218.002559 | 0.44436562 | 0.104791304 | 8.38E-06 | 8.63E-05 |
| NEBL | 116.4970386 | 0.444012535 | 0.170225406 | 0.00287201 | 0.012736663 |
| NPIPB9 | 71.01906025 | 0.443951652 | 0.211903766 | 0.00973798 | 0.034778002 |
| STX12 | 2146.691202 | 0.443940319 | 0.087832457 | 1.68E-07 | 2.79E-06 |
| ZRSR2 | 717.6002324 | 0.443795097 | 0.117457717 | 5.79E-05 | 0.000461559 |
| ATF2 | 4453.921549 | 0.44364346 | 0.102761103 | 5.98E-06 | 6.41E-05 |
| STK17B | 722.0414462 | 0.443449321 | 0.113802151 | 3.61E-05 | 0.000309863 |
| GPR137C | 1479.534339 | 0.443396399 | 0.15858508 | 0.0017021 | 0.008226096 |
| NMD3 | 3163.03485 | 0.443114302 | 0.095682655 | 1.40E-06 | 1.80E-05 |
| PTGER3 | 65.30027199 | 0.442597082 | 0.221231388 | 0.011837 | 0.040698217 |
| PCNX1 | 3791.897743 | 0.442585898 | 0.09234866 | 6.39E-07 | 9.01E-06 |
| USP32 | 5085.880355 | 0.442580905 | 0.086985598 | 1.42E-07 | 2.41E-06 |
| TMEM263 | 2464.632722 | 0.442399784 | 0.09547917 | 1.39E-06 | 1.79E-05 |
| ADCY5 | 727.3268257 | 0.442311409 | 0.14793718 | 0.00095301 | 0.005053617 |
| EPB41L5 | 2378.629682 | 0.44222233 | 0.099246254 | 3.21E-06 | 3.71E-05 |
| MAGI3 | 5645.161653 | 0.442064582 | 0.097404077 | 2.18E-06 | 2.64E-05 |
| KIF1C | 1257.1648 | 0.441955927 | 0.168647365 | 0.00281458 | 0.012538802 |
| CYSTM1 | 212.9757494 | 0.441588482 | 0.146007134 | 0.00085826 | 0.004616578 |
| SPRTN | 1412.369312 | 0.441582877 | 0.087376096 | 1.70E-07 | 2.83E-06 |
| TNFRSF10D | 3.046163245 | 0.4412261 | 0.700441934 | 0.01501456 | 0.049212519 |
| PHB1P9 | 129.0568319 | 0.441204976 | 0.185541891 | 0.00528018 | 0.021006305 |
| CLDN1 | 2.698607193 | 0.441151361 | 0.752542589 | 0.01157568 | 0.039963706 |
| KLF5 | 49.86840876 | 0.441070048 | 0.22381251 | 0.01266188 | 0.042945558 |
| NAA16 | 4138.771909 | 0.440957922 | 0.078238993 | 6.95E-09 | 1.58E-07 |
| C17orf80 | 4109.736962 | 0.440785392 | 0.094881133 | 1.32E-06 | 1.71E-05 |
| HSPA7 | 3.030345516 | 0.440640441 | 0.699327636 | 0.01511292 | 0.049495463 |
| ADGRB3 | 1337.431128 | 0.440596718 | 0.112768948 | 3.51E-05 | 0.000303163 |
| ZFR | 10154.24988 | 0.44050268 | 0.087448183 | 1.86E-07 | 3.07E-06 |
| COPB2 | 4698.152596 | 0.440464591 | 0.096385142 | 1.90E-06 | 2.34E-05 |

| | | | | | |
|---|---|---|---|---|---|
| HOMEZ | 244.9196212 | 0.440415468 | 0.180351923 | 0.00453252 | 0.018526473 |
| IARS1 | 18371.55518 | 0.440266898 | 0.083628166 | 5.59E-08 | 1.05E-06 |
| PCGF5 | 711.2006138 | 0.440115221 | 0.12901134 | 0.00023457 | 0.001536666 |
| HBP1 | 1001.83128 | 0.440056023 | 0.12806996 | 0.00021481 | 0.001424506 |
| GORASP2 | 3840.523992 | 0.440035334 | 0.116357301 | 5.83E-05 | 0.000463941 |
| ZNF671 | 526.520448 | 0.439214168 | 0.146619882 | 0.00095279 | 0.005053617 |
| ASPHD2 | 213.1689413 | 0.439067069 | 0.176301658 | 0.00404327 | 0.016872777 |
| IGSF22 | 100.2933318 | 0.438940563 | 0.182700716 | 0.00504731 | 0.0202483 |
| ABHD5 | 1031.827019 | 0.438837157 | 0.104514491 | 1.03E-05 | 0.000104328 |
| MAPK9 | 2668.27048 | 0.438628346 | 0.081983928 | 3.53E-08 | 6.91E-07 |
| H1-2 | 104.2191421 | 0.438235916 | 0.173982413 | 0.00377669 | 0.01595093 |
| ZNF391 | 455.4069423 | 0.438171369 | 0.124920596 | 0.00016726 | 0.001156564 |
| SNX14 | 802.5369414 | 0.438007002 | 0.15368515 | 0.00149824 | 0.007381349 |
| TAF12 | 574.4168049 | 0.437753017 | 0.123072574 | 0.0001397 | 0.000993032 |
| CABYR | 296.8410776 | 0.437665239 | 0.138867228 | 0.00058109 | 0.003318398 |
| DYNC2I1 | 1143.679722 | 0.437572289 | 0.105886281 | 1.39E-05 | 0.000135949 |
| HSPA14 | 1392.102614 | 0.437534301 | 0.100500872 | 5.23E-06 | 5.71E-05 |
| CRCP | 2550.77396 | 0.437343577 | 0.107244099 | 1.75E-05 | 0.000166839 |
| DUSP26 | 3121.922565 | 0.437186648 | 0.110251154 | 2.81E-05 | 0.000250223 |
| CLIP3 | 2675.609245 | 0.437021542 | 0.199255337 | 0.00831674 | 0.030551184 |
| CLU | 2140.752289 | 0.436726977 | 0.143934311 | 0.00085522 | 0.00460147 |
| EPC1 | 2129.857363 | 0.436665961 | 0.089475775 | 4.24E-07 | 6.33E-06 |
| ELF2 | 2163.851988 | 0.436321652 | 0.097026804 | 2.73E-06 | 3.22E-05 |
| ATP8B2 | 5587.621762 | 0.436203889 | 0.127111368 | 0.00022274 | 0.001468927 |
| ECD | 1833.54191 | 0.436130485 | 0.093277396 | 1.17E-06 | 1.53E-05 |
| PPP1R3B | 311.0952095 | 0.436076704 | 0.125103568 | 0.00018322 | 0.001247587 |
| SUCO | 5090.84458 | 0.436017278 | 0.08529175 | 1.29E-07 | 2.21E-06 |
| TCEAL5 | 390.4103754 | 0.435518649 | 0.121875237 | 0.00013283 | 0.000949127 |
| CCDC9 | 489.160495 | 0.435445299 | 0.181744007 | 0.00524755 | 0.020900745 |
| YTHDC2 | 5687.03243 | 0.43520402 | 0.098070825 | 3.61E-06 | 4.10E-05 |
| NME9 | 64.95096245 | 0.434814195 | 0.190641681 | 0.00693402 | 0.0262599 |
| BLZF1 | 1029.467096 | 0.434671374 | 0.11004733 | 3.04E-05 | 0.000267587 |
| ACER3 | 1964.343478 | 0.434627885 | 0.091029619 | 7.25E-07 | 9.98E-06 |
| CTXN2-AS1 | 63.8190259 | 0.434305256 | 0.222674886 | 0.01383083 | 0.046025116 |
| ZNG1F | 417.2726509 | 0.433936027 | 0.132095935 | 0.00037794 | 0.00231102 |
| EEIG1 | 1155.747047 | 0.433691925 | 0.16154959 | 0.00248059 | 0.011263142 |
| ANO10 | 573.6268613 | 0.433532441 | 0.110609604 | 3.46E-05 | 0.000299327 |
| UNC80 | 2052.573304 | 0.433502225 | 0.095139722 | 2.09E-06 | 2.54E-05 |
| ZSWIM9 | 462.8520406 | 0.433446565 | 0.152612161 | 0.00158412 | 0.007739575 |
| UNC5D | 551.0734525 | 0.433368392 | 0.139681868 | 0.00069913 | 0.003884611 |
| PKNOX2 | 258.5517228 | 0.433287256 | 0.201349397 | 0.00933337 | 0.03359555 |
| TMEM178B | 3082.242412 | 0.433020238 | 0.097973111 | 3.96E-06 | 4.46E-05 |
| SYF2 | 1395.74773 | 0.432946448 | 0.113740538 | 5.48E-05 | 0.000439468 |
| SULT4A1 | 1855.646535 | 0.432896799 | 0.111059353 | 3.79E-05 | 0.000322634 |
| SPAG16 | 716.1033064 | 0.432847359 | 0.102720292 | 9.97E-06 | 0.000100878 |
| TCEA1 | 6220.447836 | 0.432802198 | 0.095172205 | 2.19E-06 | 2.64E-05 |
| GSKIP | 1062.547582 | 0.432791842 | 0.096826977 | 3.15E-06 | 3.65E-05 |
| TRIM32 | 1292.941828 | 0.432668027 | 0.096853352 | 3.18E-06 | 3.70E-05 |
| HBEGF | 99.98592599 | 0.432650095 | 0.167225252 | 0.00326196 | 0.014108592 |
| TANK | 1389.741501 | 0.43248428 | 0.105348419 | 1.60E-05 | 0.000153687 |

| | | | | | |
|---|---|---|---|---|---|
| TAF3 | 1410.628023 | 0.432374828 | 0.157952102 | 0.00215242 | 0.010055046 |
| ZNF596 | 609.9187657 | 0.432364841 | 0.104021999 | 1.28E-05 | 0.000126635 |
| EPM2A-DT | 257.3300605 | 0.432304672 | 0.141133695 | 0.00079907 | 0.004336435 |
| NEMF | 2906.601544 | 0.432203263 | 0.090919393 | 8.13E-07 | 1.11E-05 |
| MATCAP2 | 927.8802527 | 0.432151643 | 0.101545279 | 8.32E-06 | 8.59E-05 |
| TMEM59L | 611.3890112 | 0.431988299 | 0.155154573 | 0.00188387 | 0.008996733 |
| WDR91 | 1490.892755 | 0.431573801 | 0.107780115 | 2.46E-05 | 0.000223557 |
| ZNF132 | 459.338924 | 0.43143769 | 0.134219801 | 0.00048736 | 0.002867564 |
| MKLN1 | 2589.968506 | 0.431071151 | 0.099963468 | 6.50E-06 | 6.91E-05 |
| HBS1L | 3260.140617 | 0.431038602 | 0.085599159 | 1.97E-07 | 3.22E-06 |
| PRKACB | 2880.420375 | 0.431009922 | 0.087945362 | 3.92E-07 | 5.88E-06 |
| ABCD4 | 2254.550646 | 0.43099243 | 0.097815817 | 4.25E-06 | 4.75E-05 |
| MAST4 | 311.3107448 | 0.430892678 | 0.171621694 | 0.00403825 | 0.016861439 |
| COL3A1 | 56.80274605 | 0.430798717 | 0.206346912 | 0.01086387 | 0.038004877 |
| C5orf22 | 1731.60782 | 0.4306894 | 0.090093219 | 7.17E-07 | 9.91E-06 |
| CRY1 | 2687.587333 | 0.430687694 | 0.101930123 | 9.58E-06 | 9.72E-05 |
| MED10 | 1163.3481 | 0.430559501 | 0.109222352 | 3.20E-05 | 0.000279465 |
| FNIP1 | 1648.85772 | 0.429886492 | 0.105917592 | 1.97E-05 | 0.000184979 |
| SNAPC1 | 670.4838194 | 0.428656607 | 0.103225192 | 1.33E-05 | 0.000130566 |
| ELF1 | 253.480849 | 0.428501263 | 0.138982275 | 0.00076459 | 0.004188345 |
| PRSS27 | 349.247111 | 0.428140761 | 0.155434442 | 0.00209996 | 0.009853284 |
| OS9 | 4531.370943 | 0.428006988 | 0.116608196 | 9.55E-05 | 0.000710311 |
| PDIA3 | 16894.25692 | 0.427942639 | 0.098267563 | 5.45E-06 | 5.91E-05 |
| YTHDC1 | 5278.919026 | 0.427919546 | 0.079351894 | 2.93E-08 | 5.86E-07 |
| MBD5 | 1033.059128 | 0.42790261 | 0.091107611 | 1.10E-06 | 1.45E-05 |
| THAP5 | 1700.398338 | 0.427818893 | 0.111210722 | 4.78E-05 | 0.000393801 |
| DMWD | 2320.804782 | 0.427603554 | 0.173479429 | 0.00463848 | 0.018898287 |
| FDXR | 1109.269552 | 0.427530033 | 0.117059131 | 0.00010266 | 0.000756399 |
| SNX2 | 2602.179532 | 0.427306831 | 0.094899566 | 2.77E-06 | 3.27E-05 |
| YY1AP1 | 3367.04099 | 0.427268144 | 0.126870274 | 0.0002931 | 0.001852841 |
| WDR37 | 1773.508008 | 0.427053786 | 0.104788105 | 1.86E-05 | 0.000175723 |
| MAP3K14-AS1 | 168.6122792 | 0.427013412 | 0.161450204 | 0.00288222 | 0.01277646 |
| ZNG1DP | 90.30651638 | 0.426977875 | 0.191135395 | 0.00813144 | 0.029972219 |
| GKAP1 | 808.7242302 | 0.426826474 | 0.123159443 | 0.00020675 | 0.001377691 |
| HSPA13 | 5833.865514 | 0.42680396 | 0.114502845 | 7.70E-05 | 0.000590484 |
| PLAA | 2755.299792 | 0.426102258 | 0.095643997 | 3.47E-06 | 3.97E-05 |
| NRBF2 | 841.5758148 | 0.426082563 | 0.106097947 | 2.41E-05 | 0.000219148 |
| ATG2B | 3472.162652 | 0.426046721 | 0.097238461 | 4.87E-06 | 5.36E-05 |
| TOX4 | 2339.302368 | 0.425567531 | 0.091282309 | 1.31E-06 | 1.70E-05 |
| HIVEP2 | 2118.982096 | 0.425387135 | 0.137790834 | 0.00076727 | 0.004199966 |
| DEUP1 | 2.646436543 | 0.425229437 | 0.739377045 | 0.01269734 | 0.043044532 |
| MON2 | 4042.154735 | 0.425165947 | 0.090453433 | 1.09E-06 | 1.44E-05 |
| AGO4 | 2580.314582 | 0.424737539 | 0.093352056 | 2.25E-06 | 2.70E-05 |
| PDIK1L | 1085.101376 | 0.424577882 | 0.120999892 | 0.00017847 | 0.001222101 |
| MTRF1L | 1159.631223 | 0.424336112 | 0.096247547 | 4.33E-06 | 4.83E-05 |
| HLCS | 959.3568178 | 0.423800306 | 0.106455859 | 2.81E-05 | 0.000250223 |
| JUNB | 770.4559294 | 0.423431444 | 0.198881547 | 0.01048397 | 0.036954471 |
| FDFT1 | 9663.485481 | 0.422930199 | 0.110489154 | 5.28E-05 | 0.000426797 |
| CAPN7 | 1996.789353 | 0.422901429 | 0.105900759 | 2.68E-05 | 0.000240508 |
| SDAD1P1 | 91.84547382 | 0.422645717 | 0.168891728 | 0.00433764 | 0.017875584 |

| | | | | | |
|---|---|---|---|---|---|
| BTG1 | 2530.706938 | 0.422614386 | 0.11885364 | 0.00015148 | 0.001062792 |
| FBN1-DT | 2.232259837 | 0.422582846 | 0.712051381 | 0.01483615 | 0.048749083 |
| IQCC | 301.1665574 | 0.422255053 | 0.144776104 | 0.00133812 | 0.006734334 |
| PCM1 | 11026.36226 | 0.422193114 | 0.090821041 | 1.42E-06 | 1.82E-05 |
| MKRN7P | 91.65056676 | 0.422116531 | 0.188086301 | 0.00819766 | 0.030167637 |
| HAX1 | 3609.176556 | 0.421873197 | 0.095068849 | 3.84E-06 | 4.34E-05 |
| LEMD3 | 2530.026385 | 0.421750512 | 0.090599792 | 1.38E-06 | 1.78E-05 |
| GINM1 | 860.3333297 | 0.421630035 | 0.113503678 | 8.30E-05 | 0.000629694 |
| LINC01936 | 2.31451306 | 0.421358407 | 0.714812443 | 0.01465411 | 0.048254142 |
| IQCK | 208.1604101 | 0.421327046 | 0.136726122 | 0.00079844 | 0.004334484 |
| PIK3CA | 1915.434588 | 0.421003499 | 0.091138275 | 1.64E-06 | 2.07E-05 |
| XPNPEP3 | 1385.482603 | 0.420864178 | 0.119588636 | 0.00017497 | 0.001201924 |
| PRR14 | 1233.066905 | 0.420433414 | 0.173051894 | 0.00530132 | 0.021070026 |
| FAM24B | 118.8270376 | 0.420317863 | 0.160917121 | 0.00328017 | 0.01417385 |
| MMP13 | 2.480853055 | 0.420168233 | 0.725570043 | 0.0138357 | 0.046030948 |
| TMEM242 | 960.8249635 | 0.419574992 | 0.103623529 | 2.16E-05 | 0.000199425 |
| SLC38A9 | 968.0104816 | 0.418738439 | 0.105586437 | 3.07E-05 | 0.000270096 |
| RASSF8 | 1254.278894 | 0.41871726 | 0.103091249 | 2.05E-05 | 0.000191298 |
| PLCB4 | 3116.800074 | 0.418448585 | 0.110956091 | 6.75E-05 | 0.00052787 |
| BRF2 | 400.9343968 | 0.418273918 | 0.132125889 | 0.00061477 | 0.003481725 |
| SRP68 | 8264.732896 | 0.417722005 | 0.117625781 | 0.00015776 | 0.001101214 |
| ZC3H7A | 4934.496118 | 0.417558925 | 0.080139177 | 8.28E-08 | 1.50E-06 |
| KDM4A | 1490.599261 | 0.417424633 | 0.163115005 | 0.00385017 | 0.016197808 |
| NBR1 | 5928.458965 | 0.417404105 | 0.083022834 | 2.18E-07 | 3.51E-06 |
| ZNF419 | 961.2592839 | 0.417110194 | 0.141366232 | 0.0012383 | 0.006315596 |
| CACNB1 | 1757.691993 | 0.417091976 | 0.162406295 | 0.0037641 | 0.015907283 |
| NPC1 | 1319.327723 | 0.416573501 | 0.127748587 | 0.00044921 | 0.002672909 |
| CHKA | 1877.47489 | 0.416314657 | 0.102470825 | 2.07E-05 | 0.000192232 |
| STX18 | 1185.686585 | 0.416289165 | 0.096539153 | 6.97E-06 | 7.34E-05 |
| NSMCE2 | 679.4900196 | 0.416075063 | 0.110676528 | 7.16E-05 | 0.00055447 |
| PPFIA3 | 2576.726297 | 0.41586952 | 0.13748625 | 0.00098617 | 0.005201236 |
| STXBP5 | 1372.710652 | 0.415807224 | 0.11053658 | 7.10E-05 | 0.000550527 |
| FAM227A | 945.1915231 | 0.415691155 | 0.138584859 | 0.00106968 | 0.005568681 |
| FRG1 | 1212.418496 | 0.415628568 | 0.094715372 | 4.95E-06 | 5.44E-05 |
| THSD1 | 74.65218078 | 0.415531292 | 0.186839145 | 0.00895537 | 0.032507428 |
| MIRLET7IHG | 574.3142311 | 0.415404521 | 0.114761263 | 0.00012329 | 0.000887995 |
| WDR7 | 1367.981894 | 0.415047428 | 0.106948925 | 4.43E-05 | 0.000369209 |
| PGAP1 | 8102.593894 | 0.414979848 | 0.087747744 | 9.90E-07 | 1.33E-05 |
| ZNF547 | 106.4070797 | 0.414836044 | 0.181926312 | 0.00789659 | 0.029227043 |
| IFRD1 | 626.9099631 | 0.414787049 | 0.099690971 | 1.37E-05 | 0.000134146 |
| MFF-DT | 100.8420315 | 0.414653831 | 0.182089844 | 0.00796236 | 0.029443978 |
| WWC2 | 788.4835663 | 0.414609678 | 0.108040964 | 5.28E-05 | 0.000426797 |
| RHOQ | 2585.682993 | 0.413947784 | 0.095122254 | 5.89E-06 | 6.33E-05 |
| FAM171B | 2292.620208 | 0.41368808 | 0.090298491 | 2.03E-06 | 2.47E-05 |
| RBM39 | 15764.91578 | 0.413272196 | 0.10206568 | 2.22E-05 | 0.000203693 |
| PGM2L1 | 983.4898649 | 0.413103669 | 0.113434144 | 0.00011458 | 0.000831783 |
| STAMBPL1 | 1001.072622 | 0.413100603 | 0.116580561 | 0.00016606 | 0.001149811 |
| GATAD1 | 2468.250559 | 0.413041661 | 0.096986779 | 8.97E-06 | 9.16E-05 |
| SENP7 | 1753.664005 | 0.412942178 | 0.131477141 | 0.00068625 | 0.003829583 |
| MFSD9 | 1112.22984 | 0.412886566 | 0.090111878 | 2.03E-06 | 2.48E-05 |

| | | | | | |
|---|---|---|---|---|---|
| DPM1 | 1891.07268 | 0.412633793 | 0.110850284 | 8.41E-05 | 0.000635992 |
| APOLD1 | 721.555084 | 0.412434971 | 0.162584343 | 0.0042057 | 0.017431369 |
| ZNF578 | 891.9811167 | 0.412364981 | 0.118930824 | 0.00022071 | 0.001459862 |
| ZFYVE16 | 2827.785506 | 0.412349122 | 0.100274288 | 1.71E-05 | 0.000162632 |
| LGMN | 1070.318772 | 0.41200185 | 0.151002521 | 0.00247691 | 0.011248916 |
| NUB1 | 2332.720061 | 0.411672586 | 0.083509877 | 3.69E-07 | 5.57E-06 |
| DCAF6 | 1463.636923 | 0.411505954 | 0.088580894 | 1.51E-06 | 1.92E-05 |
| LONP1 | 6532.562821 | 0.411448959 | 0.162728272 | 0.00432384 | 0.017831155 |
| PDE4D | 616.9725187 | 0.411302326 | 0.163607963 | 0.00449683 | 0.018406212 |
| UBLCP1 | 1910.472752 | 0.411237601 | 0.095086267 | 6.73E-06 | 7.12E-05 |
| ERCC4 | 1394.458975 | 0.411136774 | 0.105855458 | 4.45E-05 | 0.000370722 |
| DCP1A | 3831.486745 | 0.410797461 | 0.0853998 | 6.76E-07 | 9.44E-06 |
| ZNF621 | 4625.236864 | 0.410750144 | 0.101067132 | 2.11E-05 | 0.000195398 |
| CUL3 | 4368.795694 | 0.410089108 | 0.083976951 | 4.70E-07 | 6.93E-06 |
| TMEM255A | 277.8366964 | 0.409624888 | 0.162944608 | 0.00454059 | 0.018552076 |
| CBY1 | 472.9187306 | 0.409567565 | 0.110818325 | 9.46E-05 | 0.000705607 |
| NCAM2 | 309.9445969 | 0.409091706 | 0.157174558 | 0.00358396 | 0.015277881 |
| HHIP-AS1 | 110.7315997 | 0.408739812 | 0.167983117 | 0.00562748 | 0.022105716 |
| TSNAX | 2721.871772 | 0.40871671 | 0.093952789 | 6.07E-06 | 6.50E-05 |
| CNOT10 | 2376.735387 | 0.408509568 | 0.084251983 | 5.63E-07 | 8.11E-06 |
| RTN3P1 | 67.87614259 | 0.408370789 | 0.190975551 | 0.01134347 | 0.039363338 |
| SHISA3 | 459.4630366 | 0.40795115 | 0.118555114 | 0.00024798 | 0.001610687 |
| GORASP1 | 1574.911076 | 0.40794041 | 0.110981703 | 0.00010302 | 0.000758532 |
| ZSCAN12 | 2706.703797 | 0.407817881 | 0.103740289 | 3.72E-05 | 0.00031765 |
| RNF169 | 2367.588128 | 0.40725879 | 0.107347266 | 6.51E-05 | 0.000511233 |
| GDPD3 | 132.9796258 | 0.407214543 | 0.155341748 | 0.00343977 | 0.014758074 |
| CHMP2B | 1188.34861 | 0.407181374 | 0.090902557 | 3.38E-06 | 3.88E-05 |
| STARD10 | 497.0761408 | 0.407179603 | 0.167518598 | 0.00571551 | 0.022404369 |
| MEF2D | 2233.540907 | 0.407166618 | 0.183303535 | 0.00949436 | 0.034061759 |
| MCFD2 | 4586.196262 | 0.406931058 | 0.083349779 | 4.79E-07 | 7.03E-06 |
| MTA3 | 2292.007244 | 0.40689683 | 0.106471728 | 5.83E-05 | 0.000464332 |
| ZNF112 | 476.3856798 | 0.406815756 | 0.136464512 | 0.00118663 | 0.006092821 |
| SEC61A2 | 1710.075586 | 0.406758445 | 0.108119766 | 7.39E-05 | 0.000569933 |
| MED13L | 4050.41445 | 0.40635214 | 0.117875406 | 0.00024423 | 0.001589854 |
| PPM1B | 1380.029462 | 0.406009267 | 0.083969638 | 6.09E-07 | 8.63E-06 |
| TP53INP1 | 1755.324686 | 0.405717694 | 0.1413224 | 0.00167919 | 0.008140593 |
| TLCD2 | 77.00488066 | 0.405156529 | 0.199590631 | 0.01455531 | 0.047990288 |
| RNF19B | 609.4104005 | 0.405077906 | 0.179415563 | 0.00883563 | 0.032175662 |
| STAT3 | 6691.547612 | 0.404228834 | 0.151879899 | 0.00312525 | 0.013665243 |
| TAX1BP1 | 3715.608534 | 0.404031107 | 0.097003235 | 1.41E-05 | 0.000137596 |
| NINJ1 | 232.3164737 | 0.403568959 | 0.151999873 | 0.00319364 | 0.013855268 |
| ZNNT1 | 283.431021 | 0.403513219 | 0.183145996 | 0.01011732 | 0.035852587 |
| TBC1D10A | 143.4959331 | 0.403392598 | 0.177489753 | 0.00861528 | 0.031465427 |
| SNX13 | 1858.911051 | 0.403372525 | 0.113232317 | 0.00016196 | 0.001127498 |
| RHOC | 395.3362163 | 0.403073748 | 0.147739499 | 0.00259871 | 0.01170388 |
| RNF181 | 921.0031963 | 0.402761845 | 0.097627575 | 1.68E-05 | 0.000160574 |
| MED13 | 13157.07245 | 0.402651713 | 0.084817722 | 9.56E-07 | 1.29E-05 |
| NOVA1 | 499.1213496 | 0.401717521 | 0.114154361 | 0.00019176 | 0.001294204 |
| PRKAA2 | 2372.089681 | 0.401604729 | 0.09671313 | 1.50E-05 | 0.000145413 |
| WASHC3 | 413.5041504 | 0.4014404 | 0.122149759 | 0.00044259 | 0.002640405 |

| | | | | | |
|---|---|---|---|---|---|
| PDCD4 | 2474.512005 | 0.400853328 | 0.096270734 | 1.44E-05 | 0.000139841 |
| TRIM67 | 2500.77566 | 0.400840081 | 0.150018345 | 0.00308934 | 0.013539875 |
| POR | 1346.754987 | 0.400830055 | 0.151828214 | 0.00337999 | 0.014531061 |
| SLC50A1 | 655.8226962 | 0.400473818 | 0.120142534 | 0.0003774 | 0.002308879 |
| LEPROTL1 | 4681.79632 | 0.400448505 | 0.090943795 | 4.94E-06 | 5.43E-05 |
| SSR1 | 7062.239499 | 0.400410537 | 0.087009519 | 1.95E-06 | 2.39E-05 |
| ZNF75D | 755.3686515 | 0.400379814 | 0.096364816 | 1.50E-05 | 0.000144872 |
| ZNF879 | 283.2605681 | 0.40036551 | 0.140165504 | 0.00180309 | 0.008643064 |
| MED27 | 1282.057804 | 0.400327888 | 0.128985836 | 0.00082548 | 0.004456613 |
| GDPGP1 | 111.7393846 | 0.400285696 | 0.168187269 | 0.0067562 | 0.025695317 |
| RBM26-AS1 | 243.0491108 | 0.400056613 | 0.186994214 | 0.01192121 | 0.040946661 |
| KIF5A | 5036.546953 | 0.400015111 | 0.128136971 | 0.00077878 | 0.004251332 |
| NSDHL | 1060.877673 | 0.400000902 | 0.088498353 | 2.88E-06 | 3.39E-05 |
| CSMD2 | 1568.414744 | 0.399667415 | 0.163260851 | 0.00570259 | 0.022366679 |
| SRP14-DT | 336.2773094 | 0.399271122 | 0.13535212 | 0.00135922 | 0.006820666 |
| ZNF229 | 1014.712102 | 0.399191499 | 0.111309202 | 0.00015083 | 0.001059679 |
| ZBTB7C | 92.57235428 | 0.399189176 | 0.191370425 | 0.01345683 | 0.04505474 |
| IQCE | 959.0033224 | 0.398759437 | 0.155131433 | 0.00414094 | 0.017206373 |
| AKIP1 | 602.3945102 | 0.398695949 | 0.10963419 | 0.00012488 | 0.000897267 |
| SGSM1 | 487.5647915 | 0.398691621 | 0.1429463 | 0.00222454 | 0.010333293 |
| FAM204A | 1566.279027 | 0.398428784 | 0.0856911 | 1.56E-06 | 1.98E-05 |
| PSMG3-AS1 | 328.8435702 | 0.398088696 | 0.165801225 | 0.00648803 | 0.024905692 |
| ZNF429 | 953.5894213 | 0.398010574 | 0.142689032 | 0.00223054 | 0.010354113 |
| SLC2A4RG | 1153.992562 | 0.397969251 | 0.154588324 | 0.0041142 | 0.017122938 |
| MKRN2 | 1132.566267 | 0.397968069 | 0.102862132 | 5.02E-05 | 0.000409264 |
| VPS13B | 2225.7132 | 0.397886579 | 0.084275489 | 1.11E-06 | 1.46E-05 |
| RDH10 | 449.1447823 | 0.397683545 | 0.139928656 | 0.00190848 | 0.009099451 |
| CCDC82 | 3825.428815 | 0.397510901 | 0.113209905 | 0.0002012 | 0.001347024 |
| SCART1 | 346.2217642 | 0.397079402 | 0.18769891 | 0.01279298 | 0.043318756 |
| ABHD13 | 1427.295285 | 0.397005879 | 0.114251434 | 0.00023061 | 0.001514627 |
| HYCC2 | 2027.462003 | 0.396915763 | 0.097160321 | 2.05E-05 | 0.000190881 |
| SSPN | 204.1935329 | 0.39661919 | 0.145032653 | 0.00263717 | 0.011851247 |
| STK19 | 566.8278265 | 0.396232949 | 0.131395163 | 0.0011204 | 0.00580349 |
| TAPBPL | 227.5008924 | 0.396063851 | 0.162435489 | 0.00596495 | 0.023227286 |
| ANO4 | 241.4292049 | 0.395733557 | 0.15741282 | 0.00490005 | 0.019777149 |
| KRAS | 5251.091121 | 0.395237204 | 0.076811026 | 1.29E-07 | 2.21E-06 |
| RPAP2 | 3221.052168 | 0.39503046 | 0.106771271 | 9.95E-05 | 0.000735554 |
| SLC4A1AP | 2595.747515 | 0.394901683 | 0.085187843 | 1.70E-06 | 2.13E-05 |
| KMT2A | 7643.997063 | 0.39446654 | 0.097105698 | 2.28E-05 | 0.000208966 |
| SFT2D1 | 1032.333598 | 0.39430956 | 0.124840928 | 0.00070807 | 0.003926852 |
| DPF2 | 3364.32571 | 0.394282757 | 0.128662314 | 0.00096582 | 0.005108314 |
| HELZ | 7233.261659 | 0.39404219 | 0.089567127 | 5.16E-06 | 5.64E-05 |
| AP1S3 | 159.5156052 | 0.393977358 | 0.143341992 | 0.00256705 | 0.011599407 |
| TRIM35 | 1172.496026 | 0.39393739 | 0.13572704 | 0.00161685 | 0.00787514 |
| COL7A1 | 707.2698131 | 0.393680857 | 0.160541633 | 0.00582889 | 0.022758118 |
| MBNL1 | 2865.542853 | 0.393585041 | 0.101812048 | 5.18E-05 | 0.000420693 |
| TAGLN | 187.2491968 | 0.393551896 | 0.186087573 | 0.01308893 | 0.044059698 |
| KSR1 | 1485.826468 | 0.393495402 | 0.14636622 | 0.00306156 | 0.013437057 |
| GRIP1 | 738.5586941 | 0.393471441 | 0.138360213 | 0.00193829 | 0.009215942 |
| DMPK | 2622.633905 | 0.393438036 | 0.160897072 | 0.00594233 | 0.023147966 |

| | | | | | |
|---|---|---|---|---|---|
| UBE2B | 2689.415158 | 0.393219708 | 0.089635193 | 5.49E-06 | 5.94E-05 |
| OTUD6B | 1814.348778 | 0.393097886 | 0.095988452 | 1.99E-05 | 0.000186734 |
| APPBP2 | 6802.532376 | 0.392982858 | 0.084103385 | 1.43E-06 | 1.84E-05 |
| INPP1 | 371.7592557 | 0.391850027 | 0.117207029 | 0.00037913 | 0.002315651 |
| BNIP2 | 3915.03788 | 0.391744354 | 0.094763195 | 1.70E-05 | 0.00016214 |
| LMLN | 439.9960245 | 0.391707779 | 0.103003103 | 6.72E-05 | 0.00052563 |
| FIP1L1 | 1864.326398 | 0.391488206 | 0.085310505 | 2.15E-06 | 2.61E-05 |
| PFKFB2 | 288.0992787 | 0.39147579 | 0.126237786 | 0.00086812 | 0.004662295 |
| ATP6V1H | 1461.004342 | 0.391459461 | 0.124575056 | 0.00075726 | 0.004157049 |
| GAPVD1 | 3604.913289 | 0.391382614 | 0.084833086 | 1.92E-06 | 2.36E-05 |
| TTC1 | 1581.467838 | 0.391380127 | 0.101335668 | 5.30E-05 | 0.000428124 |
| CHIC1 | 487.0217973 | 0.39130993 | 0.122812508 | 0.00065401 | 0.003672612 |
| GOLT1B | 2851.087062 | 0.391237646 | 0.099614492 | 4.06E-05 | 0.000343071 |
| LRRC17 | 131.5911305 | 0.390770163 | 0.163565231 | 0.00696153 | 0.02633981 |
| C12orf4 | 1783.84958 | 0.390748897 | 0.103427702 | 7.45E-05 | 0.0005738 |
| IFT57 | 1319.631557 | 0.390566484 | 0.088706313 | 5.16E-06 | 5.63E-05 |
| PHF21A | 3859.200316 | 0.390334974 | 0.109264738 | 0.00016535 | 0.001145906 |
| KDM6B | 5726.29194 | 0.390273716 | 0.186125992 | 0.01388473 | 0.046149293 |
| AGAP4 | 1663.022553 | 0.390064361 | 0.102896396 | 7.10E-05 | 0.000550527 |
| ZSCAN30 | 1643.191231 | 0.389883469 | 0.097751844 | 3.17E-05 | 0.000277765 |
| BORA | 846.0598964 | 0.389835146 | 0.151607989 | 0.00432876 | 0.017846132 |
| DYNC2H1 | 1074.644484 | 0.389740225 | 0.101600701 | 5.94E-05 | 0.000471858 |
| ZNF383 | 1011.980997 | 0.389519558 | 0.119973774 | 0.00053653 | 0.003106192 |
| DPF3 | 972.4152732 | 0.388978905 | 0.166981632 | 0.00815762 | 0.030057912 |
| ZNF225 | 400.9382334 | 0.388567746 | 0.118122197 | 0.00046456 | 0.002750434 |
| FAF1 | 2466.690681 | 0.388361929 | 0.090564459 | 8.74E-06 | 8.94E-05 |
| ARL1 | 2015.937612 | 0.388127486 | 0.089521526 | 7.07E-06 | 7.44E-05 |
| OGA | 12077.26717 | 0.387853315 | 0.091858901 | 1.17E-05 | 0.000116799 |
| FAM3C | 1770.17393 | 0.387790763 | 0.092832311 | 1.43E-05 | 0.000139373 |
| EDRF1-AS1 | 134.6101612 | 0.387681045 | 0.155819329 | 0.00547788 | 0.021621319 |
| YARS1 | 7416.431855 | 0.387567985 | 0.12350895 | 0.00078227 | 0.004264713 |
| SAR1B | 2070.487622 | 0.387473041 | 0.095621047 | 2.45E-05 | 0.000222985 |
| MSH2-OT1 | 93.13993307 | 0.387345937 | 0.184870939 | 0.01417881 | 0.046981561 |
| ZNF155 | 286.2667461 | 0.387074521 | 0.137878313 | 0.00223136 | 0.010355596 |
| TBC1D15 | 2763.827376 | 0.386846638 | 0.097498312 | 3.51E-05 | 0.000302701 |
| XPC-AS1 | 194.8559354 | 0.386636505 | 0.156611492 | 0.00579464 | 0.022654183 |
| ZNF160 | 1826.228576 | 0.386433082 | 0.098807666 | 4.44E-05 | 0.000369999 |
| AJM1 | 280.4679928 | 0.386385292 | 0.167828341 | 0.00884445 | 0.032202054 |
| PRRC1 | 4463.740253 | 0.385773339 | 0.08295804 | 1.64E-06 | 2.07E-05 |
| KIF27 | 586.0352749 | 0.385470529 | 0.141256911 | 0.00283526 | 0.012609076 |
| TASP1 | 341.6121696 | 0.385458495 | 0.127245036 | 0.00112799 | 0.005836922 |
| SMAP1 | 758.2956699 | 0.385360385 | 0.096340914 | 3.08E-05 | 0.000271009 |
| SYNE1 | 2024.746485 | 0.385284413 | 0.123100931 | 0.00081216 | 0.004396216 |
| SLC7A1 | 8524.144999 | 0.385275099 | 0.145170877 | 0.00351927 | 0.015055182 |
| TRIM39 | 488.9350731 | 0.385259572 | 0.130429537 | 0.00143585 | 0.007123497 |
| PCDH19 | 419.9802646 | 0.38512279 | 0.158090489 | 0.00636042 | 0.024512572 |
| LINC00632 | 1612.596451 | 0.384904743 | 0.099768227 | 5.54E-05 | 0.000443385 |
| DKK1 | 1359.726667 | 0.384795283 | 0.156926395 | 0.00611178 | 0.023686906 |
| CARF | 531.602647 | 0.384786459 | 0.145483265 | 0.00361929 | 0.015390226 |
| KYAT3 | 1667.119622 | 0.384700005 | 0.113689735 | 0.00033878 | 0.002100278 |

| | | | | | |
|---|---|---|---|---|---|
| PPP1R12B | 2291.07406 | 0.384367176 | 0.116455411 | 0.00045547 | 0.002704643 |
| ACVR2A | 1207.863594 | 0.384367076 | 0.101550189 | 7.45E-05 | 0.000573764 |
| ANKRD44 | 984.7078909 | 0.384286631 | 0.147623647 | 0.00407813 | 0.016990019 |
| NAA35 | 2783.493571 | 0.383853708 | 0.083208503 | 1.98E-06 | 2.42E-05 |
| COQ6 | 295.6697056 | 0.383541598 | 0.142216098 | 0.00313958 | 0.013701688 |
| ZSCAN26 | 942.9537978 | 0.38346056 | 0.129790919 | 0.00144482 | 0.007159353 |
| RHOU | 4038.48689 | 0.383364548 | 0.108334681 | 0.00019333 | 0.001303474 |
| RNF146 | 2130.359897 | 0.383264494 | 0.10749541 | 0.00017499 | 0.001201924 |
| PLGLB1 | 135.5345954 | 0.383208771 | 0.16615918 | 0.00891093 | 0.032398236 |
| DPP6 | 1910.095963 | 0.38286481 | 0.136880186 | 0.00234782 | 0.010769713 |
| ZBTB6 | 1474.787479 | 0.382386 | 0.130716577 | 0.00159087 | 0.007770685 |
| GPATCH8 | 10944.86249 | 0.382358578 | 0.101693752 | 8.30E-05 | 0.000629227 |
| EEA1 | 1994.313277 | 0.382316188 | 0.085744926 | 4.12E-06 | 4.62E-05 |
| TMEM231 | 323.8203519 | 0.38209601 | 0.136137404 | 0.00228958 | 0.01056146 |
| CHPT1 | 1178.806953 | 0.38178552 | 0.119163078 | 0.00064302 | 0.003619804 |
| FKBP14 | 1065.970741 | 0.381744682 | 0.133213214 | 0.00191836 | 0.009140229 |
| IL6ST | 2386.468155 | 0.381612854 | 0.105252006 | 0.0001402 | 0.000995555 |
| USP47 | 3813.568468 | 0.381291107 | 0.093948101 | 2.45E-05 | 0.000222713 |
| KLF3 | 2140.734289 | 0.381225822 | 0.090358329 | 1.22E-05 | 0.000121284 |
| SURF4 | 5979.109415 | 0.381063181 | 0.149005126 | 0.00470112 | 0.019109283 |
| RGL1 | 654.0740747 | 0.380942029 | 0.120999914 | 0.00077866 | 0.004251332 |
| RAB3GAP1 | 4048.720821 | 0.380777222 | 0.084032307 | 2.95E-06 | 3.46E-05 |
| TAB3 | 1726.924148 | 0.380726723 | 0.08821537 | 7.96E-06 | 8.27E-05 |
| DDX50 | 3867.535695 | 0.380683188 | 0.096891923 | 4.22E-05 | 0.000354609 |
| DNAJC27 | 528.0960215 | 0.380542354 | 0.140767403 | 0.00312668 | 0.013665661 |
| HERC2P9 | 207.9628963 | 0.380256621 | 0.166886845 | 0.0096856 | 0.034602955 |
| SNRNP48 | 1525.797788 | 0.380226167 | 0.105959789 | 0.00016252 | 0.001130222 |
| CSNK2A2 | 1858.777929 | 0.379871502 | 0.157727668 | 0.00702031 | 0.026508568 |
| PTPRN | 4325.850132 | 0.379172997 | 0.175766217 | 0.01294832 | 0.043693679 |
| PLPP5 | 1397.452657 | 0.378998432 | 0.110169869 | 0.00028328 | 0.001800718 |
| CEP120 | 4797.342046 | 0.378834029 | 0.104396589 | 0.00014017 | 0.000995555 |
| ZZEF1 | 4481.115238 | 0.378788447 | 0.159210251 | 0.00760666 | 0.028347824 |
| IKZF5 | 2029.186501 | 0.378709599 | 0.106434983 | 0.00018333 | 0.001247627 |
| MARK3 | 5091.501038 | 0.378592697 | 0.085242408 | 4.53E-06 | 5.02E-05 |
| SIPA1L2 | 4873.125658 | 0.378528212 | 0.100809948 | 8.60E-05 | 0.000648279 |
| CCNDBP1 | 1617.74625 | 0.378256132 | 0.094181854 | 2.96E-05 | 0.000262055 |
| FEM1C | 1468.439996 | 0.378246842 | 0.108482967 | 0.00023962 | 0.001564797 |
| WASHC4 | 3554.978387 | 0.378116902 | 0.098235412 | 5.91E-05 | 0.00047028 |
| ACAP2 | 3386.434749 | 0.378024251 | 0.096147245 | 4.22E-05 | 0.000354522 |
| SLC35G2 | 1570.366612 | 0.377831482 | 0.112873408 | 0.00039728 | 0.00240782 |
| USP33 | 7942.52039 | 0.377623838 | 0.093567202 | 2.74E-05 | 0.00024459 |
| FAM8A1 | 3853.023546 | 0.3776069 | 0.112550184 | 0.00038705 | 0.00235621 |
| GRIK2 | 1806.66129 | 0.377396236 | 0.104563484 | 0.00015193 | 0.00106523 |
| GPRASP2 | 2122.744453 | 0.377382163 | 0.07806555 | 6.87E-07 | 9.58E-06 |
| SRPK2 | 2751.593008 | 0.377201952 | 0.0817363 | 2.01E-06 | 2.45E-05 |
| CDC42BPA | 5773.405097 | 0.377175062 | 0.090549396 | 1.57E-05 | 0.00015143 |
| CSRNP3 | 1954.073277 | 0.376920294 | 0.105207838 | 0.00016842 | 0.001163352 |
| FURIN | 1073.402757 | 0.376705932 | 0.168017359 | 0.01077449 | 0.037764898 |
| ERCC6L2-AS1 | 230.5369773 | 0.376551691 | 0.127137767 | 0.001457 | 0.007212626 |
| SELENOW | 1226.18608 | 0.376404927 | 0.112367705 | 0.00039636 | 0.002403614 |

| | | | | | |
|---|---|---|---|---|---|
| EFEMP2 | 262.8333129 | 0.376400161 | 0.15822861 | 0.0077073 | 0.028681308 |
| MSMO1 | 3934.991368 | 0.376370276 | 0.125936011 | 0.00133915 | 0.006734685 |
| MADD | 5152.021731 | 0.376285137 | 0.117916417 | 0.00068789 | 0.003832519 |
| MAFG | 3822.514297 | 0.376270524 | 0.144055121 | 0.00413816 | 0.017201758 |
| KDM4B | 3345.450032 | 0.376251072 | 0.160802404 | 0.00850928 | 0.031113939 |
| COPB1 | 6321.291681 | 0.37595849 | 0.093661692 | 3.02E-05 | 0.000266597 |
| MSX1 | 101.9522581 | 0.375906767 | 0.170706598 | 0.01189092 | 0.040863112 |
| METTL6 | 1135.655095 | 0.375730181 | 0.086517988 | 7.19E-06 | 7.55E-05 |
| CDKL3 | 114.8933106 | 0.375565281 | 0.16321794 | 0.00940083 | 0.033796981 |
| SYNJ2 | 486.337737 | 0.375354474 | 0.125149898 | 0.0013002 | 0.00657261 |
| GDPD1 | 1092.07357 | 0.375341751 | 0.121209769 | 0.00094754 | 0.005029735 |
| UGDH-AS1 | 229.6614572 | 0.374695001 | 0.173974428 | 0.01338028 | 0.044864201 |
| PLPBP | 1754.813319 | 0.374450013 | 0.083344793 | 3.63E-06 | 4.12E-05 |
| LZTS3 | 913.1513352 | 0.374303438 | 0.136129734 | 0.00281338 | 0.012536367 |
| NLRX1 | 269.4156193 | 0.373364088 | 0.144700331 | 0.00458356 | 0.01870166 |
| UNC119B | 3604.980421 | 0.373352099 | 0.109307927 | 0.00031708 | 0.001985676 |
| GPR180 | 1323.564683 | 0.372783381 | 0.105568072 | 0.00020781 | 0.001382534 |
| AFF4 | 8031.230815 | 0.372640784 | 0.088188025 | 1.23E-05 | 0.000121804 |
| NR1H3 | 159.0006637 | 0.372475757 | 0.144133549 | 0.00455256 | 0.0185899 |
| STARD7-AS1 | 500.4037447 | 0.372265208 | 0.137139777 | 0.00314747 | 0.01372382 |
| CAMK2D | 1900.54332 | 0.372189824 | 0.111027436 | 0.00039992 | 0.002420951 |
| NVL | 3165.616994 | 0.372051927 | 0.092181918 | 2.79E-05 | 0.000248962 |
| PREB | 2219.938766 | 0.372001057 | 0.130236815 | 0.00206384 | 0.009711835 |
| BCL11B | 2271.878867 | 0.371928607 | 0.108471974 | 0.00030406 | 0.001912215 |
| DNAH14 | 1167.644328 | 0.371893521 | 0.101325017 | 0.00012293 | 0.000886044 |
| CCSER2 | 2172.463095 | 0.37165959 | 0.08891975 | 1.51E-05 | 0.000145921 |
| BET1L | 1050.764617 | 0.371483772 | 0.124016606 | 0.00133876 | 0.006734417 |
| TSEN15 | 2223.549339 | 0.370981174 | 0.086884871 | 1.02E-05 | 0.000102662 |
| C12orf57 | 1441.804771 | 0.370585279 | 0.110298533 | 0.00039184 | 0.002381173 |
| MLF1 | 603.6102801 | 0.370362562 | 0.113005746 | 0.00052443 | 0.003045544 |
| PVT1 | 859.8672797 | 0.370314191 | 0.108609293 | 0.00032813 | 0.002042904 |
| SERTAD4 | 1798.10633 | 0.370072108 | 0.104164576 | 0.00019384 | 0.001305657 |
| MTMR7 | 576.458644 | 0.369913849 | 0.132730348 | 0.00256936 | 0.01160266 |
| RUSF1 | 1179.959291 | 0.369879354 | 0.126583797 | 0.0017006 | 0.008220771 |
| C9orf85 | 588.1765668 | 0.369873671 | 0.104045306 | 0.00019242 | 0.001298199 |
| FAM76B | 2330.904834 | 0.369728012 | 0.121168925 | 0.00112625 | 0.005829417 |
| TIMM44 | 2735.06042 | 0.369652195 | 0.100785657 | 0.00012529 | 0.000899298 |
| TTC7B | 1204.639066 | 0.369237316 | 0.161729047 | 0.01014971 | 0.035942601 |
| CSPP1 | 2157.681657 | 0.368817947 | 0.090964721 | 2.62E-05 | 0.000235602 |
| ZDHHC20 | 3259.32395 | 0.368742831 | 0.113052773 | 0.00055757 | 0.003202738 |
| VIPAS39 | 996.0728164 | 0.368697408 | 0.104169241 | 0.00020492 | 0.001367734 |
| SHC1 | 3841.663186 | 0.368429832 | 0.161899243 | 0.01037697 | 0.036658986 |
| MSANTD7 | 2274.988401 | 0.368132562 | 0.094170903 | 4.81E-05 | 0.000396252 |
| PTK6 | 135.2260743 | 0.367946407 | 0.16629635 | 0.01210193 | 0.041415224 |
| LINC02035 | 828.8882164 | 0.367847591 | 0.129736409 | 0.00224137 | 0.0103927 |
| KLHL5 | 1751.611562 | 0.36767348 | 0.105361635 | 0.0002476 | 0.001608706 |
| ZKSCAN5 | 1455.273425 | 0.367637619 | 0.103047674 | 0.00018509 | 0.001258217 |
| HLX | 425.1235049 | 0.367060061 | 0.159461793 | 0.0097995 | 0.034943111 |
| PAN3 | 2138.602441 | 0.366765175 | 0.092621915 | 3.93E-05 | 0.000332541 |
| NCAM1-AS1 | 416.4578243 | 0.366668661 | 0.134146405 | 0.00305574 | 0.013418366 |

| | | | | | |
|---|---|---|---|---|---|
| IFT56 | 721.5920659 | 0.366060502 | 0.114354393 | 0.00069438 | 0.003862378 |
| MTMR6 | 3101.559134 | 0.365813717 | 0.093938723 | 5.17E-05 | 0.00042001 |
| ACOT8 | 655.0395995 | 0.365791353 | 0.111064068 | 0.00050516 | 0.002955389 |
| FBXO30 | 2857.719105 | 0.365588554 | 0.131591692 | 0.00268891 | 0.012057273 |
| DHX32 | 1071.681097 | 0.365266571 | 0.1054919 | 0.00027625 | 0.001762163 |
| PAPOLG | 2045.735411 | 0.364985811 | 0.095180295 | 6.60E-05 | 0.000517508 |
| UFM1 | 3245.561288 | 0.364899179 | 0.092394779 | 4.13E-05 | 0.000348345 |
| NCOA2 | 2550.415608 | 0.364893677 | 0.091103962 | 3.27E-05 | 0.000284904 |
| ZBTB48 | 724.8098661 | 0.364438025 | 0.128671438 | 0.00229631 | 0.010585362 |
| KIF18A | 1640.328104 | 0.364256568 | 0.095348293 | 7.01E-05 | 0.000545793 |
| ACOX1 | 3135.393431 | 0.363990496 | 0.100256329 | 0.00014782 | 0.001040979 |
| MAP3K1 | 1524.150048 | 0.363217044 | 0.136490212 | 0.00382642 | 0.016127576 |
| ZNF568 | 1782.592645 | 0.363200218 | 0.125157452 | 0.00186373 | 0.008908802 |
| NBAS | 91226.9607 | 0.362918888 | 0.095591332 | 7.75E-05 | 0.000593193 |
| NUDT6 | 154.3778466 | 0.362899126 | 0.152866295 | 0.0083461 | 0.030615264 |
| ATP10D | 2689.887693 | 0.362520683 | 0.097848234 | 0.0001115 | 0.000813095 |
| UBA6-DT | 410.6619471 | 0.362148403 | 0.122186401 | 0.00154094 | 0.007557325 |
| KBTBD8 | 618.1256866 | 0.362086412 | 0.131807878 | 0.00299511 | 0.0131944 |
| BCAS4 | 106.8929137 | 0.362039545 | 0.156704301 | 0.00983881 | 0.035046814 |
| RPN1 | 8092.822916 | 0.361651534 | 0.083824276 | 8.63E-06 | 8.85E-05 |
| SMC6 | 3031.607777 | 0.361290347 | 0.082301981 | 6.13E-06 | 6.56E-05 |
| PDIA3P1 | 915.546856 | 0.361263492 | 0.108407731 | 0.00044861 | 0.002670097 |
| PDIA6 | 7491.138533 | 0.361020507 | 0.092156263 | 4.78E-05 | 0.000394075 |
| H6PD | 598.7722874 | 0.360872646 | 0.14621745 | 0.00659301 | 0.025214546 |
| DUSP11 | 1061.828741 | 0.359963316 | 0.101611569 | 0.00020967 | 0.001393986 |
| CEP19 | 131.9603552 | 0.359924302 | 0.158216867 | 0.01085022 | 0.03796562 |
| SECISBP2L | 3144.705259 | 0.359089889 | 0.103335501 | 0.00027041 | 0.001733894 |
| TTLL5 | 1274.319354 | 0.358784394 | 0.088248383 | 2.59E-05 | 0.000233657 |
| SCD | 33266.39174 | 0.35866348 | 0.090430501 | 3.94E-05 | 0.000333808 |
| USP25 | 1539.271051 | 0.358634784 | 0.100036535 | 0.00017956 | 0.001227938 |
| SUPT7L | 3330.813329 | 0.358161822 | 0.085744621 | 1.61E-05 | 0.000154183 |
| H1-0 | 6314.191248 | 0.357773027 | 0.126404491 | 0.00237718 | 0.010884935 |
| SNRNP27 | 1382.72415 | 0.357601954 | 0.096590008 | 0.00011482 | 0.000832548 |
| TSSK5P | 225.3442708 | 0.357527015 | 0.163211596 | 0.01344407 | 0.045026691 |
| CLSTN3 | 572.5797977 | 0.357497036 | 0.160346063 | 0.01225666 | 0.041868084 |
| ACADSB | 1354.189345 | 0.357401411 | 0.105607555 | 0.00037868 | 0.00231488 |
| MAPK10 | 1654.947506 | 0.357017599 | 0.098330679 | 0.00015175 | 0.001064348 |
| ZNF34 | 290.223252 | 0.356942612 | 0.133790224 | 0.00385836 | 0.016225624 |
| ZNF584 | 390.221584 | 0.356700446 | 0.13459877 | 0.00406498 | 0.016942092 |
| TMEM9 | 3752.725739 | 0.356643082 | 0.086828652 | 2.19E-05 | 0.000201239 |
| STON2 | 1241.994161 | 0.356402923 | 0.118421089 | 0.00136354 | 0.006840652 |
| PHKB | 2932.737715 | 0.356311405 | 0.098593809 | 0.0001623 | 0.001129096 |
| AFTPH | 1812.036941 | 0.35625991 | 0.095808149 | 0.00010831 | 0.000793488 |
| MCCC1 | 1750.312722 | 0.356236345 | 0.081372205 | 6.60E-06 | 7.00E-05 |
| SMC5 | 3866.938399 | 0.356223888 | 0.102428782 | 0.00027065 | 0.001734919 |
| RNF152 | 641.2185156 | 0.356194968 | 0.112340796 | 0.00080163 | 0.004347248 |
| BAK1 | 838.029213 | 0.356095005 | 0.144596774 | 0.00683737 | 0.025956019 |
| TFB1M | 579.0183149 | 0.355574414 | 0.102153153 | 0.00026832 | 0.001723737 |
| EDRF1 | 2350.231729 | 0.355524156 | 0.105891778 | 0.00041992 | 0.002526415 |
| MRPL49 | 2247.819973 | 0.355409068 | 0.108680528 | 0.00057134 | 0.003264529 |

| | | | | | |
|---|---|---|---|---|---|
| BROX | 2376.725867 | 0.355403462 | 0.104598499 | 0.00036348 | 0.002233224 |
| BNIP1 | 376.6762793 | 0.355081659 | 0.124947466 | 0.00232176 | 0.010678755 |
| EFNB2 | 261.7140986 | 0.354815559 | 0.138989112 | 0.00538645 | 0.021334156 |
| TLK2P1 | 161.7803482 | 0.35471761 | 0.154374633 | 0.01051688 | 0.037051421 |
| ZNF585B | 1249.713436 | 0.354665066 | 0.114486754 | 0.00102918 | 0.005387984 |
| LONRF2 | 1765.900957 | 0.35418072 | 0.110661513 | 0.00072997 | 0.00402724 |
| NAA30 | 1675.353261 | 0.354111044 | 0.099096564 | 0.00019093 | 0.001289844 |
| GBF1 | 5952.976115 | 0.353447654 | 0.134414292 | 0.00437187 | 0.01798782 |
| TMED5 | 4387.69995 | 0.353060474 | 0.091901548 | 6.71E-05 | 0.00052502 |
| ITCH | 1225.458008 | 0.352776473 | 0.114978286 | 0.00114362 | 0.005897032 |
| TPH1 | 332.0581796 | 0.352665989 | 0.154920229 | 0.01119524 | 0.038927769 |
| THAP9-AS1 | 3518.463183 | 0.352625988 | 0.116758183 | 0.00133842 | 0.006734334 |
| ATP6V0A2 | 4166.871043 | 0.352396655 | 0.092051414 | 7.10E-05 | 0.000550527 |
| ZNF175 | 687.4794917 | 0.352393224 | 0.10725915 | 0.00054822 | 0.003161197 |
| LTN1 | 3984.15739 | 0.352343462 | 0.107060362 | 0.00053761 | 0.003111407 |
| SWAP70 | 2131.963213 | 0.352327471 | 0.086935922 | 2.80E-05 | 0.000249702 |
| ADD3 | 3068.879905 | 0.352317538 | 0.115487945 | 0.00121337 | 0.006205367 |
| CYP2R1 | 610.6866496 | 0.351864929 | 0.106603432 | 0.00052073 | 0.003027001 |
| FAM219A | 2683.475918 | 0.351651981 | 0.147383493 | 0.00853269 | 0.031188462 |
| FBF1 | 290.239013 | 0.351406109 | 0.163256853 | 0.01515062 | 0.049587368 |
| LINC02610 | 210.822694 | 0.351091057 | 0.157486855 | 0.01265103 | 0.042915847 |
| ANKRD12 | 1723.221839 | 0.35093113 | 0.12342457 | 0.00235184 | 0.010780906 |
| IRF2 | 242.0146078 | 0.350919892 | 0.128036827 | 0.0032015 | 0.013880611 |
| CDC6 | 4812.032339 | 0.350887766 | 0.088928326 | 4.42E-05 | 0.000368709 |
| ZMAT3 | 2192.291759 | 0.350606347 | 0.126001833 | 0.00283074 | 0.012594419 |
| RHPN2 | 420.3591781 | 0.350539682 | 0.108184641 | 0.00064625 | 0.00363695 |
| NHSL2 | 2726.990211 | 0.350233466 | 0.133979948 | 0.0046316 | 0.018875183 |
| TIGD2 | 901.5014546 | 0.349939159 | 0.103468796 | 0.00039282 | 0.002384982 |
| SNHG8 | 1615.905234 | 0.34984512 | 0.100101779 | 0.00026031 | 0.001679127 |
| GSPT2 | 549.5876258 | 0.349720181 | 0.107582185 | 0.00062511 | 0.003533455 |
| STRN3 | 1925.304329 | 0.349329339 | 0.090428301 | 6.24E-05 | 0.000492482 |
| SUSD6 | 654.8111003 | 0.349320532 | 0.131045607 | 0.0040134 | 0.01677472 |
| NT5C2 | 1662.30952 | 0.349140445 | 0.095751053 | 0.00014725 | 0.00103734 |
| C2orf49 | 1623.045142 | 0.34912904 | 0.092444125 | 8.84E-05 | 0.000662838 |
| PPIL3 | 898.3695053 | 0.348996183 | 0.106399585 | 0.00056603 | 0.003241365 |
| MALT1 | 1315.277345 | 0.348655841 | 0.1161285 | 0.00144037 | 0.007142462 |
| TATDN1 | 1003.413872 | 0.348558603 | 0.122947374 | 0.00243559 | 0.011102958 |
| ZNF829 | 966.6524736 | 0.348272241 | 0.117674242 | 0.00165376 | 0.008030212 |
| RIC8B | 1960.689457 | 0.348211192 | 0.084072178 | 1.94E-05 | 0.000182278 |
| ZER1 | 1432.625616 | 0.348187693 | 0.148501208 | 0.00963674 | 0.034464342 |
| ARHGEF9 | 1825.526166 | 0.348175313 | 0.086056585 | 2.93E-05 | 0.00025959 |
| TSPY26P | 436.0586339 | 0.348069667 | 0.161437921 | 0.01527226 | 0.049921559 |
| LRRC23 | 147.8209471 | 0.347864256 | 0.158999985 | 0.01418792 | 0.046982342 |
| VPS8 | 2535.804485 | 0.347825449 | 0.112904789 | 0.00111929 | 0.005800669 |
| CCDC59 | 1637.367912 | 0.347461299 | 0.099610111 | 0.00026923 | 0.001728502 |
| HACD2 | 3091.960417 | 0.347248694 | 0.096525357 | 0.00017879 | 0.001223595 |
| ARL14EP | 2073.317857 | 0.346148665 | 0.107183935 | 0.00068203 | 0.003808102 |
| DCAF10 | 2644.602716 | 0.345629103 | 0.081386747 | 1.24E-05 | 0.000122381 |
| PCGF6 | 1188.863066 | 0.345581287 | 0.091465965 | 8.90E-05 | 0.000666452 |
| SHCBP1 | 2008.030286 | 0.345536783 | 0.09220086 | 0.00010046 | 0.000741808 |

| | | | | | |
|---|---|---|---|---|---|
| KMT5A | 3349.666234 | 0.345326312 | 0.153310032 | 0.01228962 | 0.041950582 |
| PGBD1 | 1140.997002 | 0.344839402 | 0.096433562 | 0.00019577 | 0.001317323 |
| ZNF546 | 736.9606949 | 0.34469688 | 0.127222553 | 0.00360319 | 0.015334431 |
| KCTD9 | 1442.618934 | 0.344494977 | 0.103770323 | 0.00050072 | 0.002931895 |
| DTNB | 872.2078346 | 0.344486485 | 0.124209327 | 0.00298326 | 0.013145022 |
| PIK3C2A | 6447.80873 | 0.344401284 | 0.096907611 | 0.00021308 | 0.001413902 |
| KCTD5 | 2026.062538 | 0.344218251 | 0.151720537 | 0.01186812 | 0.040791584 |
| SAV1 | 980.5528331 | 0.344123461 | 0.110497543 | 0.00101571 | 0.005329497 |
| SSR2 | 6407.321737 | 0.343777203 | 0.073547131 | 1.71E-06 | 2.14E-05 |
| GTF2A1 | 3857.335722 | 0.343727959 | 0.092127407 | 0.00010799 | 0.000791389 |
| DAAM1 | 1824.443948 | 0.343106334 | 0.098332883 | 0.00027257 | 0.001743978 |
| CRBN | 1186.181253 | 0.343094439 | 0.121036382 | 0.00249288 | 0.011308982 |
| SLC35A1 | 314.9753971 | 0.34308734 | 0.142320173 | 0.00831426 | 0.030547525 |
| DNASE1 | 1215.250217 | 0.3429679 | 0.113164737 | 0.00134367 | 0.006752517 |
| PPFIA2 | 768.789325 | 0.3425017 | 0.117471022 | 0.00194432 | 0.009238215 |
| CNNM2 | 1131.983802 | 0.342496822 | 0.134337944 | 0.005734 | 0.022468363 |
| B3GALT2 | 751.0769585 | 0.342421896 | 0.125392406 | 0.00341498 | 0.014660894 |
| LIN7C | 2012.933299 | 0.341945475 | 0.114771433 | 0.00159208 | 0.007774743 |
| COPS4 | 2049.335721 | 0.341669574 | 0.110138898 | 0.00106757 | 0.005562089 |
| SDHA | 442.1683492 | 0.341570997 | 0.133831293 | 0.00571297 | 0.022398744 |
| RB1CC1 | 3730.54583 | 0.341420287 | 0.107385616 | 0.00082428 | 0.004452459 |
| UNC79 | 3314.839508 | 0.341362295 | 0.08892838 | 7.09E-05 | 0.000550245 |
| LRCH2 | 1680.200787 | 0.341329899 | 0.093387016 | 0.00014652 | 0.00103364 |
| MOB2 | 519.9061141 | 0.341042563 | 0.106005299 | 0.0007253 | 0.004006411 |
| PIGL | 732.090215 | 0.341038829 | 0.128207778 | 0.00422169 | 0.017485342 |
| ZNF205 | 500.9270744 | 0.34082614 | 0.151938626 | 0.01283433 | 0.043408734 |
| CEP83 | 1451.531835 | 0.340773536 | 0.102907857 | 0.0005229 | 0.003037536 |
| ZNF652 | 3800.858865 | 0.340541989 | 0.086123471 | 4.43E-05 | 0.000369209 |
| AKTIP | 742.6011171 | 0.340449761 | 0.113084251 | 0.00144822 | 0.007174476 |
| SPTLC1 | 3171.808755 | 0.340428572 | 0.097410069 | 0.00026963 | 0.001730535 |
| ARHGAP5 | 2724.652007 | 0.340410258 | 0.112761007 | 0.00141013 | 0.007022967 |
| SLC9A8 | 847.4443985 | 0.340304067 | 0.12796502 | 0.00424287 | 0.017555415 |
| LGALS8 | 2173.983955 | 0.340270232 | 0.112941086 | 0.00143882 | 0.00713649 |
| NFE2L2 | 4898.035685 | 0.340237809 | 0.088252644 | 6.65E-05 | 0.000521149 |
| SMAD1 | 1137.705647 | 0.339988109 | 0.095318627 | 0.0002062 | 0.001374924 |
| DCAF5 | 2621.980726 | 0.339812403 | 0.112285321 | 0.00137952 | 0.006893763 |
| RBM18 | 2968.750116 | 0.339714965 | 0.100551479 | 0.00041336 | 0.002490591 |
| FBXL3 | 1234.77891 | 0.33960372 | 0.128713488 | 0.00451828 | 0.018479292 |
| IMPA1 | 2618.508013 | 0.339404 | 0.107028763 | 0.00085411 | 0.004596697 |
| PLEKHM1 | 1825.386292 | 0.33922805 | 0.14878709 | 0.01180764 | 0.040617637 |
| MED12L | 875.5303935 | 0.33908692 | 0.116415379 | 0.00198839 | 0.009404134 |
| HINT3 | 1504.977725 | 0.339054332 | 0.105620532 | 0.00074875 | 0.004122085 |
| ASCC3 | 5774.488255 | 0.338943219 | 0.096052882 | 0.0002389 | 0.001561095 |
| RPAP3 | 3391.977084 | 0.338753021 | 0.090284796 | 0.00010115 | 0.000746096 |
| ZNF800 | 932.7596484 | 0.338552585 | 0.097592943 | 0.0002986 | 0.00188124 |
| CDADC1 | 363.7935787 | 0.338469726 | 0.125974951 | 0.0039478 | 0.016544221 |
| PPIG | 4845.602561 | 0.338258778 | 0.10051363 | 0.00043575 | 0.002607928 |
| TMEM182 | 205.3076299 | 0.337872976 | 0.132836653 | 0.00594358 | 0.023148439 |
| CNST | 2839.211185 | 0.337702333 | 0.105399995 | 0.00076845 | 0.004202767 |
| PLRG1 | 3079.041954 | 0.337616921 | 0.107293587 | 0.00093404 | 0.004973465 |

| | | | | | |
|---|---|---|---|---|---|
| ZNF721 | 1600.992939 | 0.337507877 | 0.084093359 | 3.49E-05 | 0.000301539 |
| CCNL1 | 5986.978397 | 0.337501652 | 0.098157468 | 0.00033542 | 0.002082616 |
| SESN1 | 1094.847072 | 0.337115124 | 0.149341458 | 0.01260514 | 0.042830806 |
| MSTO1 | 1815.820545 | 0.337075051 | 0.136274303 | 0.00721929 | 0.027150473 |
| FRS2 | 2577.894249 | 0.336472761 | 0.111616984 | 0.00145167 | 0.007188084 |
| RNF14 | 1406.335656 | 0.33645326 | 0.103408818 | 0.00065053 | 0.003656065 |
| ATAD1 | 4806.91834 | 0.336245816 | 0.083849911 | 3.56E-05 | 0.000305932 |
| PITHD1 | 1430.638029 | 0.336020801 | 0.096660734 | 0.00029336 | 0.001853917 |
| OSTC | 3271.991049 | 0.335901218 | 0.108050935 | 0.00106745 | 0.005562089 |
| ARFIP2 | 2113.569259 | 0.335718898 | 0.09624872 | 0.00028119 | 0.001789131 |
| KLHDC2 | 1981.772199 | 0.33506653 | 0.09636374 | 0.00029348 | 0.001854099 |
| SWI5 | 234.453854 | 0.33493098 | 0.133691099 | 0.00668301 | 0.025487638 |
| USP54 | 1178.995413 | 0.334791612 | 0.100643653 | 0.00050672 | 0.002962807 |
| POLR1H | 515.6173422 | 0.33467331 | 0.115334772 | 0.00209318 | 0.009827435 |
| PORCN | 554.9345352 | 0.334233282 | 0.123570711 | 0.00380929 | 0.01607515 |
| DDX19B | 433.4398526 | 0.334040256 | 0.133109208 | 0.00662991 | 0.025326867 |
| TBC1D31 | 1277.593431 | 0.333750153 | 0.112279969 | 0.00167927 | 0.008140593 |
| NBPF12 | 1591.377769 | 0.333578533 | 0.083630622 | 3.93E-05 | 0.000332541 |
| TTC17 | 4188.861277 | 0.333517021 | 0.095952002 | 0.00029644 | 0.001868772 |
| FLYWCH2 | 674.3422747 | 0.333407192 | 0.129357598 | 0.00550781 | 0.021710289 |
| WBP4 | 776.1472563 | 0.333201155 | 0.099992882 | 0.00049934 | 0.002925341 |
| EXOC1 | 2045.786529 | 0.333144984 | 0.093784344 | 0.00022325 | 0.001470491 |
| MPC2 | 603.459543 | 0.332919194 | 0.102084857 | 0.00064179 | 0.003614857 |
| ZNF12 | 2203.958404 | 0.332670295 | 0.107951435 | 0.00118186 | 0.006074433 |
| MBIP | 944.0567141 | 0.332367501 | 0.105394607 | 0.0009304 | 0.004955404 |
| SERGEF | 439.0934985 | 0.332137269 | 0.111963371 | 0.00172216 | 0.008311027 |
| OSBPL9 | 5269.444583 | 0.332117942 | 0.092080822 | 0.00018216 | 0.001242405 |
| FAM167A | 3083.529638 | 0.331584057 | 0.143812374 | 0.01144746 | 0.039657278 |
| DNAAF9 | 1921.422991 | 0.331539914 | 0.101725258 | 0.00064956 | 0.003651614 |
| SNRPA1P1 | 249.4026965 | 0.331484754 | 0.138776768 | 0.00925606 | 0.03340499 |
| ZNF583 | 673.0867987 | 0.331364868 | 0.105867117 | 0.00101086 | 0.005309648 |
| ZNG1B | 1273.430925 | 0.331247616 | 0.118326664 | 0.00290624 | 0.012863499 |
| RSF1 | 6896.417663 | 0.330717179 | 0.088280076 | 0.00010644 | 0.000781164 |
| YIPF5 | 2446.436971 | 0.330513214 | 0.093941045 | 0.0002559 | 0.001654301 |
| PDRG1 | 1011.684548 | 0.330442214 | 0.099904808 | 0.00055033 | 0.003170923 |
| IFT52 | 1904.43892 | 0.330370668 | 0.090835642 | 0.00016321 | 0.001133498 |
| PITRM1 | 2430.540595 | 0.330143327 | 0.113075622 | 0.00201279 | 0.009497698 |
| COMMD8 | 444.961775 | 0.329861183 | 0.113757795 | 0.00214542 | 0.010029197 |
| CLGN | 1656.518491 | 0.329544097 | 0.09801458 | 0.0004546 | 0.002700286 |
| KCTD18 | 442.9602125 | 0.329398957 | 0.117122398 | 0.00281343 | 0.012536367 |
| NUDT16L1 | 872.8814265 | 0.329337997 | 0.136640888 | 0.0088237 | 0.03214358 |
| TTC13 | 1535.654756 | 0.329221955 | 0.088662094 | 0.00012184 | 0.000878843 |
| ZNF438 | 179.3993912 | 0.32921865 | 0.135175575 | 0.00825698 | 0.030364176 |
| ATR | 2895.769027 | 0.32914222 | 0.106609449 | 0.00117482 | 0.00604275 |
| RAB12 | 1108.378256 | 0.328474813 | 0.101354885 | 0.00069989 | 0.00388718 |
| DNAJC2 | 2046.902819 | 0.328414105 | 0.098640785 | 0.00051307 | 0.002991421 |
| CENPC | 1280.03291 | 0.328127904 | 0.107629084 | 0.00133947 | 0.006734685 |
| MIS12 | 2527.039315 | 0.32776467 | 0.099840814 | 0.000606 | 0.003438707 |
| CAMK2G | 3003.157982 | 0.327669985 | 0.116029213 | 0.00273338 | 0.012227349 |
| TXNL1 | 3195.587977 | 0.327630297 | 0.072565457 | 3.86E-06 | 4.36E-05 |

| | | | | | |
|---|---|---|---|---|---|
| CEPT1 | 2424.735352 | 0.32730296 | 0.136851174 | 0.00934259 | 0.033611108 |
| TTC33 | 1309.602835 | 0.327215648 | 0.106930995 | 0.00129464 | 0.006550944 |
| DHRS12 | 181.080694 | 0.327211592 | 0.145565439 | 0.013472 | 0.045090844 |
| SLC25A25 | 1656.507406 | 0.32682514 | 0.145482908 | 0.01354021 | 0.045260232 |
| ASH1L | 7168.264723 | 0.326735478 | 0.091494783 | 0.00021257 | 0.001411033 |
| ZNF804A | 1856.886098 | 0.32670831 | 0.114279251 | 0.00246466 | 0.011205675 |
| EAF1 | 2075.72569 | 0.326490842 | 0.090241172 | 0.00017794 | 0.001218894 |
| COP1 | 2616.553745 | 0.326403877 | 0.090684381 | 0.00019114 | 0.001290849 |
| UBA5 | 3557.542514 | 0.326228645 | 0.0802004 | 2.88E-05 | 0.000255445 |
| ERCC5 | 343.2424126 | 0.326115832 | 0.119824586 | 0.00374068 | 0.015814849 |
| EDEM3 | 4202.865649 | 0.326078101 | 0.099194264 | 0.00060027 | 0.003408996 |
| KAT7 | 6409.262306 | 0.325863401 | 0.10935691 | 0.0016887 | 0.008170937 |
| EBAG9 | 840.8161018 | 0.325585753 | 0.107803949 | 0.00148377 | 0.007320606 |
| SNHG15 | 2543.156634 | 0.32477014 | 0.111452161 | 0.00209011 | 0.009817511 |
| GOPC | 5603.538853 | 0.324567482 | 0.093643895 | 0.00031717 | 0.001985676 |
| SEPHS2 | 2397.796581 | 0.324504861 | 0.097879014 | 0.00054664 | 0.003153167 |
| PABIR3 | 333.313027 | 0.324439242 | 0.135098572 | 0.00921107 | 0.033277658 |
| USPL1 | 2610.79595 | 0.324358864 | 0.135818897 | 0.00954212 | 0.034191342 |
| SNAP25 | 3661.055596 | 0.324252085 | 0.085509286 | 9.07E-05 | 0.000678218 |
| RAD17 | 2479.374643 | 0.324245 | 0.101551772 | 0.00083788 | 0.00451762 |
| RSBN1L | 2846.224019 | 0.323857668 | 0.092702578 | 0.00028714 | 0.001821917 |
| ABI1 | 1243.05828 | 0.323681948 | 0.093885306 | 0.00034033 | 0.002108633 |
| RAB33B | 679.3795824 | 0.323528696 | 0.112302208 | 0.00232895 | 0.010704642 |
| NT5DC1 | 1583.944087 | 0.323216276 | 0.087502306 | 0.00013416 | 0.000957576 |
| LRRFIP2 | 2054.55795 | 0.323036703 | 0.091247456 | 0.00024183 | 0.001576749 |
| PTPRE | 263.3277736 | 0.322792981 | 0.13511109 | 0.0095793 | 0.034294686 |
| MCL1 | 18004.99489 | 0.322693561 | 0.088386329 | 0.00015875 | 0.00110736 |
| PHYH | 498.0654766 | 0.322084849 | 0.117418245 | 0.00356343 | 0.015208469 |
| LAMB2 | 791.5886461 | 0.321716029 | 0.142058855 | 0.01322387 | 0.044448466 |
| DLG1 | 3054.948475 | 0.321313713 | 0.093077767 | 0.00033749 | 0.00209357 |
| LINGO2 | 305.4042318 | 0.321239117 | 0.129724704 | 0.00763833 | 0.028460713 |
| ZUP1 | 968.7009325 | 0.321107551 | 0.101420396 | 0.00092812 | 0.004944498 |
| ZNF567 | 942.3155806 | 0.320853958 | 0.136038696 | 0.01045378 | 0.036879339 |
| TDRKH | 1170.959067 | 0.320796653 | 0.101192843 | 0.00091643 | 0.004888562 |
| MFAP3 | 4458.952992 | 0.3207882 | 0.105911129 | 0.00146731 | 0.00725491 |
| MBD2 | 1164.114102 | 0.320314122 | 0.103649477 | 0.00120031 | 0.006155427 |
| ARIH2 | 5070.295799 | 0.319983566 | 0.082945796 | 7.05E-05 | 0.000548312 |
| TIGD6 | 255.1960953 | 0.318969587 | 0.128540226 | 0.0075985 | 0.028327686 |
| TRMT44 | 890.6902785 | 0.318620668 | 0.12379034 | 0.00589256 | 0.022984576 |
| BECN1 | 2485.08895 | 0.318610248 | 0.08073855 | 4.92E-05 | 0.000403272 |
| RNF11 | 4281.245364 | 0.318516666 | 0.093291513 | 0.00039129 | 0.002378511 |
| DCAF17 | 2055.027466 | 0.318421553 | 0.106717024 | 0.00171213 | 0.008266771 |
| MSANTD4 | 2060.93852 | 0.318231282 | 0.106491479 | 0.00168844 | 0.008170937 |
| GPATCH1 | 952.7432392 | 0.317985992 | 0.090558645 | 0.00027401 | 0.001750974 |
| SLC22A5 | 887.8539257 | 0.317958429 | 0.10296649 | 0.00121978 | 0.006230401 |
| RABGAP1 | 3787.759754 | 0.317944085 | 0.079163715 | 3.68E-05 | 0.000314675 |
| BBS9 | 409.798136 | 0.317812549 | 0.131217027 | 0.00895563 | 0.032507428 |
| SIRT1 | 3391.788071 | 0.317793914 | 0.091577216 | 0.00031943 | 0.001995993 |
| RNF2 | 2271.302753 | 0.317671375 | 0.106957801 | 0.00179405 | 0.008603754 |
| UPF2 | 2899.70278 | 0.317601979 | 0.092908599 | 0.0003865 | 0.002354118 |

| | | | | | |
|---|---|---|---|---|---|
| PDZD8 | 2723.101818 | 0.317488234 | 0.086775435 | 0.0001567 | 0.001094934 |
| PTPN5 | 706.6587281 | 0.317314303 | 0.138565634 | 0.01263472 | 0.042874765 |
| SYT14 | 3847.583472 | 0.317309125 | 0.09984378 | 0.00090286 | 0.004824327 |
| AACS | 3310.671996 | 0.31719596 | 0.115716113 | 0.00364846 | 0.015472679 |
| ASXL2 | 2608.022799 | 0.316977675 | 0.102063678 | 0.00115384 | 0.005945282 |
| PEAK1 | 3437.235575 | 0.316958246 | 0.133960739 | 0.0104116 | 0.036749751 |
| PPFIA1 | 5191.975334 | 0.316877164 | 0.110137427 | 0.00241346 | 0.011014279 |
| ZNF330 | 2919.570817 | 0.316740205 | 0.088224596 | 0.00020443 | 0.001365387 |
| RALGPS2 | 2514.275363 | 0.316599124 | 0.107111595 | 0.00188495 | 0.008999783 |
| RANBP10 | 1633.406944 | 0.316580408 | 0.121072346 | 0.00528727 | 0.021026366 |
| FBXO42 | 1305.745071 | 0.31655817 | 0.115302222 | 0.0036102 | 0.015357908 |
| SETD2 | 6546.327901 | 0.316431953 | 0.077257382 | 2.64E-05 | 0.000236752 |
| CREM | 494.4515973 | 0.316366664 | 0.098449342 | 0.00080218 | 0.004349028 |
| GSTCD | 1547.602973 | 0.316109035 | 0.095903995 | 0.00060211 | 0.003418486 |
| TP53BP1 | 9777.762358 | 0.316076345 | 0.100405662 | 0.00100406 | 0.005278011 |
| MTMR14 | 1382.512449 | 0.316063625 | 0.124436794 | 0.00654157 | 0.025064424 |
| CLCN6 | 1711.729199 | 0.315462664 | 0.137228001 | 0.0124474 | 0.042357817 |
| RAB27A | 563.8718472 | 0.315448192 | 0.113234433 | 0.00321079 | 0.013915011 |
| DDRGK1 | 1233.547735 | 0.315010081 | 0.094174293 | 0.00050812 | 0.002967625 |
| PEX1 | 1463.118961 | 0.314891153 | 0.106663785 | 0.00191813 | 0.009140229 |
| ATRNL1 | 998.4717372 | 0.314768763 | 0.13886729 | 0.0135337 | 0.045253159 |
| TXNRD2 | 652.5569436 | 0.314642363 | 0.136408329 | 0.01224298 | 0.041835268 |
| NHLRC2 | 3918.591455 | 0.31454145 | 0.11650957 | 0.00416475 | 0.017287848 |
| ANKRD33B | 780.1693414 | 0.314486207 | 0.11890479 | 0.00488759 | 0.019734633 |
| C3orf38 | 1248.740146 | 0.314391588 | 0.097078595 | 0.00074061 | 0.004080912 |
| DYM | 2920.709538 | 0.313979912 | 0.088191495 | 0.00023111 | 0.00151738 |
| NDUFAF1 | 595.7469259 | 0.313649331 | 0.10102018 | 0.0011704 | 0.00602155 |
| IFT46 | 705.48876 | 0.3134013 | 0.116409939 | 0.00427487 | 0.017677143 |
| UBXN4 | 9083.051135 | 0.31334367 | 0.085888589 | 0.00016539 | 0.001145906 |
| PRPF38B | 9712.779577 | 0.31334362 | 0.074081361 | 1.48E-05 | 0.000143895 |
| SEC31A | 7715.465947 | 0.31313519 | 0.08209195 | 8.59E-05 | 0.000647988 |
| ZSWIM5 | 669.3301215 | 0.312947679 | 0.130560205 | 0.00975178 | 0.034815217 |
| RNF25 | 1015.886965 | 0.312850865 | 0.111390585 | 0.00302366 | 0.013305927 |
| CCDC93 | 1891.32788 | 0.312745766 | 0.08452528 | 0.00013548 | 0.000965651 |
| LRRFIP1 | 3250.907989 | 0.312172091 | 0.079581392 | 5.54E-05 | 0.000443351 |
| DENND4C | 2543.305297 | 0.31184941 | 0.107254372 | 0.00223279 | 0.010359913 |
| JAKMIP2 | 1634.316648 | 0.311353475 | 0.086646654 | 0.00020549 | 0.001370663 |
| ZNF189 | 1665.506223 | 0.31092544 | 0.104174355 | 0.00175214 | 0.008432227 |
| ELOVL1 | 721.5125063 | 0.310889828 | 0.139420021 | 0.01505999 | 0.049329981 |
| JAK1 | 4026.203799 | 0.310534104 | 0.090814308 | 0.00039412 | 0.002392181 |
| NSMCE3 | 963.602196 | 0.310489428 | 0.118506762 | 0.00532613 | 0.021131878 |
| MOSPD1 | 465.3421605 | 0.310189982 | 0.130147763 | 0.0102136 | 0.036150172 |
| CERT1 | 1974.797659 | 0.310158756 | 0.085661763 | 0.00018577 | 0.001261182 |
| ZSCAN32 | 996.7349511 | 0.309822861 | 0.092968304 | 0.00054023 | 0.003122578 |
| GTF2H1 | 2047.630159 | 0.309435511 | 0.100370306 | 0.0012768 | 0.006474985 |
| ACAT2 | 5510.499797 | 0.309406267 | 0.102077427 | 0.00151474 | 0.007441277 |
| DYNC2LI1 | 445.6710452 | 0.309394023 | 0.123122561 | 0.00722704 | 0.027169459 |
| TMEM67 | 797.0292508 | 0.309173842 | 0.12497887 | 0.00805031 | 0.029731766 |
| VPS54 | 2014.276173 | 0.309043583 | 0.085338189 | 0.00018595 | 0.00126202 |
| CTNND1 | 343.3221825 | 0.308914763 | 0.122415346 | 0.00703058 | 0.0265376 |

| | | | | | |
|---|---|---|---|---|---|
| RBCK1 | 2285.958777 | 0.308739637 | 0.132140808 | 0.01160458 | 0.040039708 |
| GIT2 | 1359.536296 | 0.30835487 | 0.111403429 | 0.00347465 | 0.014889083 |
| SLC16A14 | 1914.253004 | 0.308304786 | 0.10807109 | 0.0026816 | 0.012029768 |
| DTWD1 | 1799.150246 | 0.307908935 | 0.124392613 | 0.00805507 | 0.029744014 |
| FAM193B | 3638.371212 | 0.307887357 | 0.133180974 | 0.01240306 | 0.042227912 |
| MOCS3 | 633.921806 | 0.307417165 | 0.123373263 | 0.00771473 | 0.028698583 |
| SUPT3H | 1059.122888 | 0.307390202 | 0.09262185 | 0.00057188 | 0.003266712 |
| RGS2 | 857.5381141 | 0.307336475 | 0.127034896 | 0.00938648 | 0.033763082 |
| TBC1D3L | 313.2478216 | 0.307192491 | 0.132632695 | 0.01229926 | 0.041971788 |
| CCDC90B | 1822.776832 | 0.307115895 | 0.101810764 | 0.00160063 | 0.007809092 |
| PEX7 | 252.1008818 | 0.307035981 | 0.127861327 | 0.009857 | 0.03508729 |
| SENP2 | 3311.819677 | 0.306982303 | 0.080676008 | 9.09E-05 | 0.000679842 |
| TSC1 | 3571.469565 | 0.306937495 | 0.078261664 | 5.65E-05 | 0.000451282 |
| RNF139 | 1783.659168 | 0.306864636 | 0.093736206 | 0.00067153 | 0.003758667 |
| TMEM59 | 7349.427192 | 0.306628448 | 0.096178742 | 0.00090399 | 0.004828482 |
| MTDH | 8432.003184 | 0.306564212 | 0.07738842 | 4.80E-05 | 0.000395415 |
| LRIG2 | 2924.37814 | 0.30642496 | 0.094469409 | 0.00074689 | 0.004113317 |
| MAP3K9 | 1650.978021 | 0.306161249 | 0.135670979 | 0.01435335 | 0.047438498 |
| PDPK1 | 2533.910609 | 0.305838669 | 0.123482321 | 0.00808337 | 0.029821728 |
| PXK | 1085.113085 | 0.30545559 | 0.109609494 | 0.00331513 | 0.014306902 |
| XRRA1 | 539.7984911 | 0.305455522 | 0.13464036 | 0.01396444 | 0.046391731 |
| GPN2 | 1541.736456 | 0.305250578 | 0.122360866 | 0.0077137 | 0.028698583 |
| ANKRD36C | 299.4158343 | 0.304337908 | 0.12979463 | 0.01154688 | 0.039887451 |
| TCAF1P1 | 469.2237467 | 0.304228845 | 0.133553673 | 0.01370558 | 0.045716377 |
| CRNKL1 | 2090.106844 | 0.303888343 | 0.105929121 | 0.00259015 | 0.011676942 |
| STK33 | 698.7846773 | 0.303574776 | 0.099162919 | 0.00139803 | 0.00697456 |
| ZKSCAN7 | 987.04906 | 0.303499087 | 0.09542968 | 0.00093777 | 0.004990337 |
| MSRB2 | 599.4683397 | 0.303482869 | 0.121067889 | 0.00750979 | 0.028052924 |
| CNEP1R1 | 652.6791779 | 0.303428786 | 0.114438364 | 0.00498723 | 0.020065108 |
| EXOSC1 | 1278.59366 | 0.303304911 | 0.108291412 | 0.0031997 | 0.013875711 |
| TBCC | 1597.037519 | 0.303206328 | 0.111548465 | 0.00410404 | 0.017087544 |
| POC1B | 1226.730103 | 0.303080114 | 0.125753377 | 0.0097704 | 0.03486501 |
| CAMKMT | 346.4410061 | 0.302962658 | 0.122055652 | 0.00804825 | 0.029729504 |
| PPP3CC | 302.7813998 | 0.30288921 | 0.117170666 | 0.00604547 | 0.023475859 |
| ANKRD13C | 2160.61842 | 0.302885087 | 0.090520334 | 0.00052642 | 0.003055422 |
| RB1 | 4894.794055 | 0.302678749 | 0.08877873 | 0.00041921 | 0.002522849 |
| CDK17 | 2248.739745 | 0.30247113 | 0.102461996 | 0.00200137 | 0.009448161 |
| UBALD1 | 1157.692216 | 0.302451224 | 0.132514981 | 0.01364344 | 0.045566219 |
| IQCB1 | 1994.245305 | 0.301768055 | 0.116643375 | 0.00603355 | 0.023441124 |
| ANKRD26 | 1008.323823 | 0.301706764 | 0.109318554 | 0.00364151 | 0.015454609 |
| SLC31A1 | 2340.705922 | 0.301612526 | 0.085037514 | 0.00025278 | 0.001637729 |
| IFT70A | 366.1264161 | 0.301554435 | 0.126195161 | 0.01037407 | 0.036655059 |
| ZNF529 | 2294.493631 | 0.301535044 | 0.114457822 | 0.00527295 | 0.020989732 |
| UMAD1 | 617.940423 | 0.301352705 | 0.114916572 | 0.00546339 | 0.021580702 |
| ARMT1 | 1393.287168 | 0.300659896 | 0.102669936 | 0.00217056 | 0.010128314 |
| PRDM5 | 602.3709424 | 0.300611812 | 0.114361567 | 0.00537976 | 0.02131586 |
| CCNT2 | 6063.168525 | 0.300601271 | 0.102120487 | 0.00206864 | 0.009730001 |
| DIAPH3 | 1637.006282 | 0.300422077 | 0.098187628 | 0.00142033 | 0.007060091 |
| ARMCX6 | 410.9321612 | 0.300416533 | 0.119396741 | 0.00739688 | 0.027686446 |
| PIGG | 3190.696486 | 0.300007546 | 0.098600024 | 0.0015041 | 0.007401651 |

| | | | | | |
|---|---|---|---|---|---|
| ZNF619 | 342.0403614 | 0.299334883 | 0.130919849 | 0.01366267 | 0.045610246 |
| APH1B | 1030.765114 | 0.299201673 | 0.106448176 | 0.00314678 | 0.01372382 |
| CDK5RAP1 | 1740.076158 | 0.299043103 | 0.083119501 | 0.00020999 | 0.001395688 |
| RBM33 | 7029.508128 | 0.298852816 | 0.100889962 | 0.00196049 | 0.009296755 |
| MTMR3 | 363.3563003 | 0.298830144 | 0.108291556 | 0.00368109 | 0.015598153 |
| ZNF227 | 1024.988158 | 0.298748462 | 0.107839351 | 0.00356398 | 0.015208469 |
| ZNF570 | 1079.41067 | 0.298596275 | 0.123153669 | 0.00955264 | 0.034223078 |
| KTN1 | 8759.659394 | 0.298488306 | 0.110232571 | 0.0043 | 0.017770318 |
| ING2 | 359.3372494 | 0.297944115 | 0.120405888 | 0.00836564 | 0.030681475 |
| BAG4 | 1497.124938 | 0.297180015 | 0.112727143 | 0.00532383 | 0.021126841 |
| ISY1 | 676.6813151 | 0.297071203 | 0.099795726 | 0.00188121 | 0.008988184 |
| MTRR | 2244.973211 | 0.296998324 | 0.103563385 | 0.00265792 | 0.011933935 |
| PSMD12 | 6542.494902 | 0.29676438 | 0.089309934 | 0.00058281 | 0.003326349 |
| PTPN21 | 1519.612144 | 0.296587207 | 0.125707011 | 0.01143434 | 0.03961848 |
| CCDC91 | 1562.12132 | 0.296329984 | 0.096608246 | 0.00140298 | 0.006997533 |
| KANSL1L | 1649.275252 | 0.295945743 | 0.108213822 | 0.00400177 | 0.016732897 |
| WDR26 | 8827.797569 | 0.295917246 | 0.085723913 | 0.00036623 | 0.002248081 |
| ZNF506 | 1959.190698 | 0.295892694 | 0.116209124 | 0.0069094 | 0.02617627 |
| BTAF1 | 5909.960271 | 0.295602605 | 0.101240788 | 0.00226749 | 0.010478368 |
| GFPT2 | 298.4364974 | 0.29527635 | 0.120296083 | 0.00891378 | 0.032402856 |
| ARHGAP29 | 2378.180995 | 0.295217281 | 0.123589206 | 0.01063637 | 0.037376332 |
| VMP1 | 3747.440274 | 0.295052456 | 0.081204754 | 0.00018532 | 0.001259401 |
| KDM5A | 4075.83581 | 0.294579463 | 0.100750379 | 0.00224586 | 0.01040879 |
| ZNF200 | 854.7417477 | 0.29456347 | 0.090731117 | 0.00076753 | 0.004199966 |
| SMARCA2 | 2155.587762 | 0.293660171 | 0.098425505 | 0.00186053 | 0.008898154 |
| LSM12 | 2013.879756 | 0.293612275 | 0.117914529 | 0.00813774 | 0.029990051 |
| ZFP28 | 982.5564726 | 0.293508369 | 0.107358631 | 0.00404496 | 0.016872777 |
| SNIP1 | 1160.075324 | 0.293386507 | 0.089207664 | 0.00066431 | 0.003726374 |
| RAB22A | 1571.370621 | 0.293169536 | 0.097000316 | 0.00164285 | 0.007988556 |
| COPS5 | 1835.582038 | 0.293133423 | 0.101206302 | 0.00246063 | 0.011197241 |
| SPOPL | 1450.765021 | 0.292973065 | 0.103008878 | 0.00289772 | 0.012831347 |
| SRP54 | 2189.291081 | 0.292968663 | 0.082875637 | 0.00027125 | 0.001736607 |
| YTHDF1 | 4540.960121 | 0.292812366 | 0.126384407 | 0.01294034 | 0.043681084 |
| DAGLB | 1262.008603 | 0.292371005 | 0.127614563 | 0.01384616 | 0.046050855 |
| SCMH1 | 3231.221175 | 0.292183582 | 0.102798836 | 0.00292188 | 0.012911857 |
| TRIP11 | 1777.344544 | 0.292172627 | 0.113137611 | 0.00631649 | 0.024379817 |
| SNX25 | 756.5184651 | 0.291895608 | 0.094368498 | 0.00130561 | 0.006596704 |
| EFL1 | 1369.347757 | 0.291892815 | 0.105169392 | 0.00358847 | 0.015293962 |
| PYCR1 | 6849.041322 | 0.291675741 | 0.088095164 | 0.00061755 | 0.003494946 |
| ADAM9 | 1718.523341 | 0.291209771 | 0.08908857 | 0.0007175 | 0.003970578 |
| FAM118B | 738.0548494 | 0.291053335 | 0.120915412 | 0.01027687 | 0.036349088 |
| GDAP2 | 909.994079 | 0.291017911 | 0.118077523 | 0.0088009 | 0.032071909 |
| SBNO1 | 5589.043073 | 0.290898944 | 0.086889386 | 0.00054251 | 0.003132023 |
| RRAGC | 551.031863 | 0.290896845 | 0.108957951 | 0.00493081 | 0.019885697 |
| BRWD1 | 5551.875816 | 0.290709929 | 0.112459717 | 0.00630117 | 0.024338326 |
| ABCF3 | 2101.40994 | 0.290513886 | 0.122919477 | 0.01155595 | 0.039912084 |
| CLDN12 | 1909.031481 | 0.290337193 | 0.112790735 | 0.00650699 | 0.024959849 |
| RPRD2 | 5257.386313 | 0.289869463 | 0.124934644 | 0.01296283 | 0.043713937 |
| PARN | 1404.654841 | 0.289605667 | 0.092485049 | 0.00115653 | 0.005956154 |
| ITGAE | 843.2691816 | 0.289506802 | 0.105185201 | 0.00387711 | 0.016291135 |

| | | | | | |
|---|---|---|---|---|---|
| GON4L | 4819.526176 | 0.289461121 | 0.112134561 | 0.00639295 | 0.024609612 |
| ACBD5 | 2274.617246 | 0.289280462 | 0.093852879 | 0.00136467 | 0.006842957 |
| MRPL1 | 1150.703219 | 0.289056873 | 0.117903367 | 0.00917788 | 0.033163565 |
| FEM1B | 8286.23984 | 0.289022515 | 0.092821401 | 0.0012294 | 0.006274845 |
| NEK1 | 2043.879634 | 0.2889036 | 0.115039507 | 0.00779568 | 0.028952562 |
| CBL | 3057.007372 | 0.288804592 | 0.107490257 | 0.00472282 | 0.019186107 |
| YIPF1 | 992.7286934 | 0.288423723 | 0.099251584 | 0.00242297 | 0.011055255 |
| SPG11 | 4333.82457 | 0.288168445 | 0.084804148 | 0.00045649 | 0.002709164 |
| MAP4K3 | 2369.415467 | 0.287988758 | 0.086307527 | 0.00056949 | 0.003258509 |
| WDFY1 | 1890.325206 | 0.287660386 | 0.088305666 | 0.00075421 | 0.004148042 |
| MYO9A | 2258.664187 | 0.287321102 | 0.097955034 | 0.00223035 | 0.010354113 |
| FBXW7 | 1857.176149 | 0.287057519 | 0.102659408 | 0.00342115 | 0.014684327 |
| XRCC1 | 1450.071465 | 0.287014593 | 0.098340441 | 0.00233852 | 0.010736607 |
| ZNF131 | 3119.17317 | 0.286997037 | 0.099205411 | 0.00253569 | 0.011470288 |
| DDB2 | 1288.976614 | 0.286802279 | 0.11238428 | 0.00700759 | 0.026465401 |
| BNIP3L | 3940.197763 | 0.286446431 | 0.095312203 | 0.0017729 | 0.008514241 |
| EPRS1 | 14362.90111 | 0.286444858 | 0.094484376 | 0.00162691 | 0.007920388 |
| AHCYL1 | 8580.017446 | 0.286433942 | 0.069533225 | 2.60E-05 | 0.000234424 |
| SMNDC1 | 2470.876361 | 0.286432127 | 0.078694694 | 0.00018545 | 0.001259846 |
| MARCHF7 | 5157.235267 | 0.28591147 | 0.102713566 | 0.00356809 | 0.015222877 |
| NT5C3A | 1710.606144 | 0.28533489 | 0.089040715 | 0.00091282 | 0.004870576 |
| STK38L | 2332.334982 | 0.285051362 | 0.099740566 | 0.00284637 | 0.012642065 |
| MYT1L | 2506.018843 | 0.284532709 | 0.093407371 | 0.00156026 | 0.007644789 |
| BOD1L1 | 4662.565072 | 0.284321527 | 0.087565403 | 0.00079038 | 0.004302092 |
| UBE2W | 1520.252717 | 0.284265304 | 0.115350323 | 0.00900773 | 0.032669299 |
| CTNNAL1 | 2490.54518 | 0.284152369 | 0.098283334 | 0.00257249 | 0.01161119 |
| ATP2B1 | 5198.820062 | 0.284038407 | 0.096873807 | 0.0022611 | 0.010465069 |
| CDK7 | 781.0535709 | 0.28358154 | 0.102853547 | 0.00389356 | 0.016346911 |
| SNX27 | 3988.941413 | 0.283176559 | 0.100267381 | 0.00317775 | 0.013818411 |
| SLC12A2 | 1678.367988 | 0.283115619 | 0.087338878 | 0.0008082 | 0.004375919 |
| FNBP4 | 7722.411833 | 0.282879063 | 0.085542649 | 0.00064297 | 0.003619804 |
| THAP1 | 373.6392775 | 0.28269843 | 0.118936155 | 0.01145279 | 0.03966905 |
| RNF141 | 1740.682984 | 0.282647847 | 0.12027036 | 0.01229101 | 0.041950582 |
| RYBP | 3675.954132 | 0.282217054 | 0.08214817 | 0.00040516 | 0.002445488 |
| ZFP69 | 546.0249195 | 0.282049907 | 0.104685183 | 0.00471965 | 0.019177019 |
| REV1 | 3353.243265 | 0.281633221 | 0.079957337 | 0.00029407 | 0.001857286 |
| LARP4B | 3801.945296 | 0.28132478 | 0.105409056 | 0.00509712 | 0.020400762 |
| ASCC1 | 1063.00606 | 0.281154052 | 0.112898774 | 0.00847386 | 0.031000957 |
| RP2 | 850.509661 | 0.281124391 | 0.097229671 | 0.00259467 | 0.011690783 |
| LCORL | 2018.992363 | 0.280853345 | 0.11436623 | 0.00932501 | 0.033577212 |
| ZMYM2 | 9168.521064 | 0.280799489 | 0.103031303 | 0.00431975 | 0.017819733 |
| SEPTIN6 | 4388.300301 | 0.280379163 | 0.110115445 | 0.00727031 | 0.027292177 |
| JMY | 2574.117115 | 0.279756074 | 0.119656292 | 0.01280807 | 0.043353208 |
| ZZZ3 | 4547.072356 | 0.279215179 | 0.099239704 | 0.00332356 | 0.014331259 |
| CLUAP1 | 945.5534215 | 0.279115885 | 0.097721274 | 0.00291337 | 0.012883556 |
| ATF1 | 1847.691072 | 0.2781375 | 0.101937017 | 0.00431435 | 0.017811726 |
| WDFY2 | 2440.300307 | 0.277869556 | 0.117133355 | 0.01177902 | 0.040539554 |
| DSTYK | 2155.958236 | 0.277155486 | 0.094172016 | 0.00222823 | 0.010348072 |
| AGO3 | 3581.969022 | 0.276917148 | 0.089182122 | 0.00131185 | 0.006620076 |
| RAB28 | 937.0005555 | 0.276853041 | 0.100732959 | 0.00408062 | 0.016996943 |

| | | | | | |
|---|---|---|---|---|---|
| AFF2 | 1316.978452 | 0.276714736 | 0.092117597 | 0.00183342 | 0.008780284 |
| GPR158 | 420.9875253 | 0.276484417 | 0.112819395 | 0.00958709 | 0.03431659 |
| ZNF502 | 885.624504 | 0.276307651 | 0.118600676 | 0.01323516 | 0.044471867 |
| MAP3K12 | 2592.949764 | 0.276225729 | 0.107862414 | 0.00706673 | 0.026659369 |
| MARS1 | 11692.16981 | 0.276015877 | 0.112910745 | 0.00976265 | 0.034847981 |
| ZNF26 | 1999.271102 | 0.275717978 | 0.108046174 | 0.00726115 | 0.027262795 |
| RHOT1 | 2892.327046 | 0.274806334 | 0.089552536 | 0.00149025 | 0.007349027 |
| RWDD1 | 1780.522844 | 0.27399799 | 0.101501243 | 0.00476331 | 0.019308648 |
| PTCD2 | 1531.77778 | 0.273886022 | 0.102341244 | 0.00510348 | 0.020418186 |
| SLC25A46 | 4875.916817 | 0.273797504 | 0.103967935 | 0.00578323 | 0.022622454 |
| CGGBP1 | 9871.945254 | 0.273752956 | 0.111084987 | 0.00931548 | 0.033554639 |
| DEDD | 1652.433943 | 0.273738361 | 0.111373923 | 0.00948347 | 0.034040518 |
| LSM1 | 604.1339725 | 0.273645938 | 0.108199223 | 0.00779015 | 0.028937267 |
| CDKAL1 | 1309.854695 | 0.27309252 | 0.081165278 | 0.00053773 | 0.003111407 |
| PIGK | 1946.387908 | 0.27263062 | 0.115998875 | 0.01269164 | 0.04303228 |
| CCNC | 2304.242403 | 0.272624015 | 0.11011325 | 0.0090581 | 0.032823039 |
| VWA8 | 1604.275294 | 0.27259049 | 0.091411184 | 0.00199206 | 0.009417178 |
| DDHD2 | 2305.069861 | 0.272504658 | 0.087363073 | 0.00126678 | 0.006433697 |
| GMPPA | 1086.823798 | 0.272497561 | 0.11368458 | 0.01122091 | 0.039003858 |
| KIFAP3 | 1423.26824 | 0.272435666 | 0.096144416 | 0.00318763 | 0.01384417 |
| STAC | 2950.117993 | 0.272053486 | 0.092916423 | 0.00237339 | 0.010872447 |
| RNF138 | 3272.39303 | 0.271908141 | 0.093830316 | 0.00261179 | 0.011755054 |
| ZSCAN20 | 621.2896474 | 0.271877524 | 0.112943043 | 0.01093858 | 0.038187183 |
| MOB4 | 1841.476061 | 0.271863622 | 0.100120007 | 0.00457208 | 0.018662204 |
| PPP3CA | 3571.622226 | 0.271829475 | 0.087553757 | 0.00133258 | 0.006713188 |
| STK16 | 882.4604452 | 0.271487667 | 0.101274969 | 0.00507253 | 0.020337577 |
| PKN2 | 5050.783514 | 0.271378398 | 0.10845677 | 0.0084553 | 0.030960563 |
| KIF5B | 13820.28419 | 0.271139573 | 0.088791616 | 0.00158253 | 0.007733627 |
| USP38 | 1999.960499 | 0.271099205 | 0.082939959 | 0.00076081 | 0.004173162 |
| NCOA4 | 5501.847413 | 0.270833059 | 0.109006283 | 0.00889223 | 0.032341654 |
| TIMM23B | 601.5721744 | 0.270152642 | 0.10579627 | 0.00735075 | 0.027543882 |
| JMJD6 | 2123.109607 | 0.270096257 | 0.111586078 | 0.01061254 | 0.037320945 |
| DISC1 | 293.9806855 | 0.270017912 | 0.116274691 | 0.01376932 | 0.045876922 |
| OGFOD1 | 2531.820302 | 0.269829928 | 0.086652431 | 0.00129921 | 0.006569205 |
| ZNF606 | 769.3729528 | 0.26956904 | 0.097878024 | 0.00409785 | 0.017065248 |
| CNKSR2 | 567.5730505 | 0.269214344 | 0.100438869 | 0.00511141 | 0.020437693 |
| CLPX | 3413.852942 | 0.268492176 | 0.086866414 | 0.0014089 | 0.007018545 |
| WDR74 | 2292.675966 | 0.268349393 | 0.109999192 | 0.01013388 | 0.035905068 |
| WASHC2A | 1804.81746 | 0.267748978 | 0.082150124 | 0.00079343 | 0.004314119 |
| DIS3 | 4704.976226 | 0.267621663 | 0.091083033 | 0.00232593 | 0.01069372 |
| RTCA | 1479.387408 | 0.26707753 | 0.090956011 | 0.00234351 | 0.010754727 |
| RRAGB | 730.5258328 | 0.267076338 | 0.111359657 | 0.01136909 | 0.039432602 |
| FCHO2 | 873.4820923 | 0.267021421 | 0.11604767 | 0.01468642 | 0.048337308 |
| AGAP13P | 501.3646939 | 0.267007579 | 0.115585459 | 0.01434631 | 0.047422838 |
| TCEAL7 | 1502.0666 | 0.266969438 | 0.099459353 | 0.00508721 | 0.020384518 |
| ELOVL6 | 3217.860807 | 0.26687038 | 0.093047195 | 0.00290919 | 0.012873825 |
| ABRAXAS2 | 1591.602654 | 0.266701513 | 0.101190207 | 0.00586924 | 0.022897953 |
| APAF1 | 1757.351485 | 0.266526456 | 0.099803685 | 0.00530348 | 0.021074534 |
| SEC61B | 1474.782731 | 0.265904902 | 0.098699662 | 0.00495602 | 0.019963887 |
| STRIP1 | 4270.255685 | 0.265460909 | 0.11139918 | 0.01190284 | 0.040897231 |

| | | | | | |
|---|---|---|---|---|---|
| ARFGEF1 | 3243.994643 | 0.265319856 | 0.09471408 | 0.00359398 | 0.015311068 |
| POLR2C | 3310.108759 | 0.265274796 | 0.074650114 | 0.00027331 | 0.001748157 |
| AEBP2 | 2430.67488 | 0.265230572 | 0.087184843 | 0.00167097 | 0.00810498 |
| CTNNA2 | 1564.910512 | 0.265204384 | 0.084120649 | 0.00115428 | 0.005946026 |
| RAE1 | 2670.80634 | 0.264723247 | 0.112994322 | 0.01326532 | 0.044565935 |
| NCAM1 | 18285.60592 | 0.264695043 | 0.1054985 | 0.00846472 | 0.030973043 |
| CYB5R1 | 1305.432337 | 0.26435594 | 0.107936127 | 0.00999112 | 0.03547263 |
| OTULINL | 1011.984684 | 0.264279321 | 0.098568089 | 0.00517266 | 0.020638445 |
| DR1 | 6149.44576 | 0.264022037 | 0.078385371 | 0.00054396 | 0.003138631 |
| PIKFYVE | 4198.964848 | 0.263961689 | 0.092912354 | 0.00319191 | 0.013853623 |
| NDUFAF7 | 1281.77269 | 0.263820109 | 0.114795843 | 0.01493966 | 0.049013918 |
| ACTR8 | 1936.184136 | 0.263746625 | 0.081644513 | 0.00088694 | 0.004752219 |
| SEC24B | 2609.839257 | 0.263685479 | 0.099130595 | 0.00551477 | 0.021726804 |
| RC3H2 | 5523.877554 | 0.263577656 | 0.09827317 | 0.00516919 | 0.020628595 |
| PHLPP2 | 638.0417776 | 0.263467206 | 0.110855796 | 0.01217712 | 0.041630964 |
| CNOT6L | 1273.220601 | 0.263387932 | 0.103230814 | 0.00754322 | 0.028157338 |
| COQ5 | 1570.036605 | 0.263172454 | 0.104125928 | 0.00807545 | 0.029805278 |
| RASAL2 | 2380.888303 | 0.263090842 | 0.094568424 | 0.00383581 | 0.016150615 |
| RECK | 1060.13031 | 0.263048524 | 0.101171481 | 0.00657459 | 0.025167502 |
| CDK20 | 793.3907456 | 0.261887227 | 0.109266714 | 0.01159601 | 0.040016839 |
| TMX3 | 4217.868013 | 0.261481539 | 0.110435636 | 0.01254364 | 0.042657081 |
| CWC27 | 1557.651905 | 0.260740295 | 0.083994208 | 0.00137578 | 0.00687857 |
| TXNDC9 | 1268.927266 | 0.260303767 | 0.109119918 | 0.01200722 | 0.041187082 |
| ZSCAN29 | 4899.47139 | 0.259738177 | 0.079653355 | 0.00080606 | 0.004367451 |
| MRS2 | 2397.036332 | 0.259703074 | 0.097129154 | 0.00535553 | 0.021232125 |
| RALGAPA1 | 2292.539218 | 0.259064464 | 0.096947084 | 0.00538988 | 0.021343647 |
| TEX10 | 3974.256671 | 0.258874948 | 0.096697155 | 0.00531471 | 0.021102836 |
| SGCE | 1494.970174 | 0.258810143 | 0.094518443 | 0.00443237 | 0.018193072 |
| PRPF18 | 1915.263547 | 0.258677727 | 0.093247354 | 0.00397774 | 0.01664595 |
| PIGA | 1051.044631 | 0.258264319 | 0.109992698 | 0.01334044 | 0.044755225 |
| TRIM26 | 2146.893671 | 0.258245728 | 0.092674685 | 0.00383382 | 0.016145556 |
| PARD3 | 2622.226322 | 0.258019889 | 0.087447622 | 0.0022954 | 0.010583562 |
| SNRPA1 | 3565.835105 | 0.257182329 | 0.083534942 | 0.00151258 | 0.007432445 |
| GGPS1 | 1208.393291 | 0.256699295 | 0.099742067 | 0.00721938 | 0.027150473 |
| N4BP2 | 3178.286524 | 0.256497548 | 0.095565777 | 0.00524319 | 0.020887452 |
| NDC80 | 3259.753079 | 0.256181996 | 0.10200133 | 0.00861341 | 0.031465427 |
| SNX19 | 3535.131197 | 0.255694372 | 0.098260842 | 0.00667081 | 0.025445824 |
| TMEM170B | 5215.586708 | 0.255576718 | 0.09438495 | 0.00489765 | 0.019771347 |
| ZNF226 | 710.6224195 | 0.255527045 | 0.102802866 | 0.00927466 | 0.033447687 |
| FAM114A2 | 1045.592771 | 0.255425696 | 0.107262718 | 0.01231553 | 0.042020322 |
| SPCS2 | 4850.946794 | 0.255382258 | 0.09582535 | 0.0055607 | 0.021868479 |
| LYSMD1 | 656.3777588 | 0.254840271 | 0.096259801 | 0.00586471 | 0.022884616 |
| MKS1 | 2210.507457 | 0.254100704 | 0.076569204 | 0.00066695 | 0.003738104 |
| NDUFAF5 | 1110.860153 | 0.254040369 | 0.092253511 | 0.00428503 | 0.017715572 |
| ZNF830 | 1055.068488 | 0.253977597 | 0.086407499 | 0.00240546 | 0.010982668 |
| PPP2R5A | 625.5784966 | 0.253926855 | 0.098391888 | 0.00712846 | 0.026843009 |
| PAAF1 | 1492.538746 | 0.253918142 | 0.088512784 | 0.00300848 | 0.013247617 |
| CAP2 | 1805.003057 | 0.253748004 | 0.096238928 | 0.00607009 | 0.023556376 |
| IDH3A | 1266.017398 | 0.253705518 | 0.089807988 | 0.0034492 | 0.014792343 |
| MSL2 | 2838.953233 | 0.253639431 | 0.085399147 | 0.00218097 | 0.010160746 |

| | | | | | |
|---|---|---|---|---|---|
| CALR | 15705.57573 | 0.253563757 | 0.107709382 | 0.01330689 | 0.044676394 |
| TNKS2 | 4931.785974 | 0.253459834 | 0.08885356 | 0.00316819 | 0.013797273 |
| ERP29 | 2719.549105 | 0.253393882 | 0.099032752 | 0.00760234 | 0.028336878 |
| CTTNBP2NL | 2461.030498 | 0.253230457 | 0.077987551 | 0.00086026 | 0.004625739 |
| PRKCI | 3951.646098 | 0.25305077 | 0.083911849 | 0.00188313 | 0.008995257 |
| RALB | 725.0752001 | 0.252767367 | 0.108005001 | 0.01383171 | 0.046025116 |
| NAPB | 2152.329086 | 0.252601642 | 0.088081209 | 0.00302739 | 0.013319357 |
| SCN3B | 5772.134682 | 0.252598457 | 0.093298198 | 0.00494291 | 0.019918886 |
| TTI2 | 618.3659146 | 0.252580574 | 0.100571707 | 0.00870319 | 0.031721462 |
| ZNF182 | 700.9660201 | 0.252427725 | 0.105337933 | 0.01193127 | 0.040974396 |
| SNRK | 1653.627965 | 0.252184495 | 0.08730244 | 0.00283889 | 0.012622479 |
| MAIP1 | 560.7804581 | 0.251523535 | 0.10535304 | 0.01225115 | 0.041856224 |
| UPF3B | 1894.750832 | 0.251261761 | 0.096434236 | 0.006688 | 0.025497211 |
| ZNF398 | 1555.517928 | 0.251210594 | 0.103150191 | 0.01077426 | 0.037764898 |
| RFK | 1403.42996 | 0.2509074 | 0.086445931 | 0.00272644 | 0.012201613 |
| VPS11 | 1620.001533 | 0.25088709 | 0.100528294 | 0.00913643 | 0.033042831 |
| SEC23A | 4255.969972 | 0.250588683 | 0.097769232 | 0.00756637 | 0.028228368 |
| DDX42 | 17515.38337 | 0.25006208 | 0.0714979 | 0.00035078 | 0.002166216 |
| FAM120B | 2925.68608 | 0.249737267 | 0.087635088 | 0.00322752 | 0.013980108 |
| TPRA1 | 1155.985519 | 0.249735377 | 0.099686426 | 0.0089256 | 0.032440097 |
| SLC39A9 | 3791.495984 | 0.249384304 | 0.088381958 | 0.00352456 | 0.015066293 |
| SASS6 | 2185.947922 | 0.24914677 | 0.104390352 | 0.01235775 | 0.042143426 |
| ZBTB11 | 2556.351678 | 0.249021296 | 0.107845092 | 0.01517037 | 0.04963618 |
| SENP1 | 2427.97467 | 0.248694762 | 0.096489959 | 0.00730034 | 0.027379922 |
| TATDN3 | 1391.079764 | 0.248132384 | 0.091624048 | 0.0049911 | 0.020069707 |
| CDC42EP3 | 8645.546277 | 0.248125725 | 0.087689409 | 0.00345037 | 0.014794299 |
| SLC38A2 | 17878.47123 | 0.247850291 | 0.08661579 | 0.00312647 | 0.013665661 |
| WRN | 3120.615632 | 0.246969301 | 0.096791821 | 0.0078943 | 0.029223827 |
| MAP3K7 | 4242.124944 | 0.246755483 | 0.100948964 | 0.0106437 | 0.037395691 |
| SELENOF | 8171.540289 | 0.246592251 | 0.104578887 | 0.01343453 | 0.045002071 |
| KLHL20 | 1951.048674 | 0.246545394 | 0.093908872 | 0.00639489 | 0.024612475 |
| DYRK1A | 3310.645678 | 0.245619037 | 0.079229725 | 0.00145061 | 0.007184572 |
| ATP6AP2 | 2977.429657 | 0.24515738 | 0.09433093 | 0.00692869 | 0.026244523 |
| TERF2IP | 2524.247316 | 0.245038308 | 0.102314624 | 0.01222442 | 0.041778775 |
| RNF185 | 1498.36728 | 0.244314633 | 0.082840151 | 0.00238857 | 0.010922513 |
| ZNF274 | 986.2130979 | 0.243453841 | 0.100347913 | 0.01128652 | 0.039198772 |
| RBM27 | 1744.856129 | 0.243421784 | 0.10121669 | 0.01195184 | 0.041031333 |
| SLC25A4 | 1818.473682 | 0.243147712 | 0.091321918 | 0.00578774 | 0.02263152 |
| ILKAP | 1956.139737 | 0.243118234 | 0.095104193 | 0.00786762 | 0.029151292 |
| LTO1 | 1537.692025 | 0.242996786 | 0.10039173 | 0.01147368 | 0.03971465 |
| CNTLN | 1848.924025 | 0.242995903 | 0.092072119 | 0.00619983 | 0.02399646 |
| RLIM | 7750.072227 | 0.242906928 | 0.096304806 | 0.0086667 | 0.031610859 |
| CLCN5 | 1574.555651 | 0.242460394 | 0.099408726 | 0.01092524 | 0.038164442 |
| BLTP1 | 3642.2623 | 0.242100884 | 0.103760369 | 0.01451557 | 0.047882301 |
| JADE1 | 3441.005632 | 0.241755627 | 0.076777491 | 0.00124221 | 0.006329253 |
| WDFY3 | 3522.06144 | 0.241577973 | 0.091924072 | 0.00642843 | 0.024713827 |
| MLX | 3470.378931 | 0.241511706 | 0.087115578 | 0.00418423 | 0.017358188 |
| ATP6V1B2 | 6765.374037 | 0.241480406 | 0.074446657 | 0.00089616 | 0.004792903 |
| EXOC2 | 2053.680106 | 0.240643352 | 0.089348922 | 0.00531858 | 0.021110077 |
| TCTN1 | 1571.145657 | 0.240642227 | 0.103182608 | 0.01461388 | 0.048160228 |

| | | | | | |
|---|---|---|---|---|---|
| BMPR2 | 5083.57055 | 0.240338049 | 0.099573963 | 0.0117705 | 0.040517006 |
| DDX27 | 2265.677465 | 0.239939082 | 0.098153838 | 0.01083436 | 0.037939583 |
| TRMT11 | 1689.77499 | 0.239790652 | 0.092429654 | 0.00711947 | 0.026816371 |
| KDSR | 1559.928259 | 0.23932448 | 0.091403463 | 0.00665001 | 0.025375931 |
| PIAS2 | 2414.463931 | 0.239021053 | 0.094986874 | 0.00890125 | 0.032368746 |
| EXOC4 | 3301.861045 | 0.238950918 | 0.076060988 | 0.00128168 | 0.006494923 |
| RAP1GDS1 | 2555.917386 | 0.238803877 | 0.088201849 | 0.00512248 | 0.020473994 |
| NKRF | 1166.545479 | 0.23851524 | 0.086824068 | 0.00455076 | 0.01858624 |
| INTS6L | 1652.823998 | 0.238350705 | 0.10157455 | 0.01415753 | 0.046919488 |
| TERF1 | 2525.712432 | 0.238212395 | 0.093234453 | 0.00799866 | 0.029572899 |
| CSNK1A1 | 5889.832445 | 0.237412692 | 0.074841103 | 0.00115865 | 0.005964069 |
| MCPH1 | 1752.335271 | 0.237086353 | 0.100590336 | 0.01381938 | 0.046013007 |
| GHITM | 7020.434814 | 0.234245904 | 0.08989863 | 0.00698684 | 0.02640641 |
| SETX | 4853.441326 | 0.234075823 | 0.091737782 | 0.00816118 | 0.030060257 |
| PKP4 | 2915.270116 | 0.233856861 | 0.100682167 | 0.0152525 | 0.049873171 |
| ANXA5 | 2123.944472 | 0.233220896 | 0.093777555 | 0.00980593 | 0.034959965 |
| USP9X | 8501.24462 | 0.233195706 | 0.088619847 | 0.00650055 | 0.024944455 |
| NOL4 | 1358.719854 | 0.2315533 | 0.087121497 | 0.00604118 | 0.023466324 |
| GOLGA7 | 1009.763143 | 0.231444041 | 0.087947478 | 0.00652463 | 0.02500886 |
| CLCN3 | 4393.844044 | 0.231331627 | 0.086039956 | 0.00551807 | 0.021726905 |
| BTG3 | 2334.565084 | 0.230347123 | 0.097062327 | 0.01346605 | 0.045078269 |
| VIRMA | 4188.551681 | 0.230339746 | 0.09058549 | 0.00844469 | 0.030938292 |
| GPHN | 2147.570892 | 0.230320442 | 0.087973648 | 0.00680561 | 0.025849733 |
| INVS | 1224.599453 | 0.230054644 | 0.095524595 | 0.01226136 | 0.041877181 |
| C16orf87 | 897.4968237 | 0.229542453 | 0.095448126 | 0.01239162 | 0.042216918 |
| TUBGCP5 | 1858.848545 | 0.228948419 | 0.094756789 | 0.01203684 | 0.041254255 |
| GYG1 | 834.3912082 | 0.228482209 | 0.096403944 | 0.01364294 | 0.045566219 |
| AGFG1 | 4064.387651 | 0.227869279 | 0.076619166 | 0.00229341 | 0.010576725 |
| BTN2A1 | 2423.071517 | 0.227811957 | 0.084654364 | 0.00552465 | 0.021747499 |
| DYNC1LI1 | 2746.067639 | 0.227442396 | 0.086880472 | 0.00685653 | 0.026009523 |
| LPIN1 | 4270.815652 | 0.226961825 | 0.093744027 | 0.01193571 | 0.040982805 |
| ZNF106 | 3487.511962 | 0.226885432 | 0.091630289 | 0.01026519 | 0.036320277 |
| SNX9 | 1916.796345 | 0.225863268 | 0.083285621 | 0.00521569 | 0.020789979 |
| HAGH | 1586.497149 | 0.225754401 | 0.088648641 | 0.00844662 | 0.030939844 |
| MMAB | 1331.153465 | 0.225272642 | 0.089935875 | 0.00951341 | 0.034112257 |
| SGPL1 | 3433.390503 | 0.225161554 | 0.091275776 | 0.01057578 | 0.03723931 |
| UTP6 | 3137.671133 | 0.22496526 | 0.093610188 | 0.01258904 | 0.04279025 |
| SEC23IP | 3523.411478 | 0.224113926 | 0.09072082 | 0.01049912 | 0.036995441 |
| PPIE | 1458.035124 | 0.223506684 | 0.0941251 | 0.01364505 | 0.045566219 |
| ASF1A | 2667.869663 | 0.223341879 | 0.091038392 | 0.01102637 | 0.038431557 |
| LCMT1 | 1070.828201 | 0.223311146 | 0.091644645 | 0.01153957 | 0.039882309 |
| TMX4 | 4166.820818 | 0.223059816 | 0.081301808 | 0.00477226 | 0.019333494 |
| RALGAPB | 5579.599313 | 0.22213629 | 0.08439953 | 0.0066636 | 0.025423068 |
| CAB39 | 4226.932938 | 0.221807265 | 0.087998883 | 0.00917709 | 0.033163565 |
| ARIH1 | 6147.46951 | 0.2203447 | 0.082986474 | 0.00625088 | 0.024177519 |
| MAPK1 | 6553.666504 | 0.219670756 | 0.080406838 | 0.00498071 | 0.020047635 |
| WASHC2C | 1769.832431 | 0.219584238 | 0.078828014 | 0.00423258 | 0.0175199 |
| ALS2 | 2662.323929 | 0.218997792 | 0.086670501 | 0.00907686 | 0.03288521 |
| ALDH9A1 | 2333.460817 | 0.218358977 | 0.085775131 | 0.00861612 | 0.031465427 |
| SAAL1 | 1736.025012 | 0.217919553 | 0.09300059 | 0.01503262 | 0.049263875 |

| | | | | | |
|---|---|---|---|---|---|
| CHD9 | 5004.969749 | 0.217021683 | 0.08949534 | 0.01209379 | 0.041401147 |
| KCMF1 | 2358.768597 | 0.216772719 | 0.092668956 | 0.01523325 | 0.049826072 |
| TM9SF2 | 5490.27112 | 0.216556685 | 0.074874343 | 0.00305648 | 0.013418767 |
| NRDC | 9711.533915 | 0.212910003 | 0.086163706 | 0.01075685 | 0.037726011 |
| MMUT | 3949.353033 | 0.20992291 | 0.085847396 | 0.01162407 | 0.040093484 |
| CRK | 4457.327768 | 0.209709216 | 0.076089541 | 0.00473252 | 0.019203628 |
| NCBP1 | 5187.534245 | 0.209235569 | 0.085982409 | 0.01202567 | 0.041229707 |
| CHD2 | 8235.246907 | 0.209067839 | 0.074842236 | 0.00422751 | 0.01750243 |
| SMURF2 | 4411.19852 | 0.207618674 | 0.08491496 | 0.01169229 | 0.040315257 |
| TRAK2 | 2372.580963 | 0.207072201 | 0.085653946 | 0.01261975 | 0.042845066 |
| PTPN12 | 3886.627197 | 0.206835923 | 0.071069308 | 0.00294602 | 0.013006016 |
| AEN | 5565.1344 | 0.202486482 | 0.084292818 | 0.01328847 | 0.044636393 |
| RTN3 | 8680.22984 | 0.200369847 | 0.083037416 | 0.01296157 | 0.043713937 |
| C6orf62 | 7087.995061 | 0.199888435 | 0.071911043 | 0.00449417 | 0.018404254 |
| WTAP | 5345.074746 | 0.199519592 | 0.083818677 | 0.01418311 | 0.046981561 |
| EPC2 | 3011.929576 | 0.198816792 | 0.071813406 | 0.00465993 | 0.018964332 |
| USP39 | 4328.6371 | 0.198638446 | 0.080838342 | 0.01152345 | 0.03983327 |
| CCDC117 | 2580.694018 | 0.197769435 | 0.079839089 | 0.0109256 | 0.038164442 |
| AVL9 | 3584.678674 | 0.197058656 | 0.078297307 | 0.00979066 | 0.034923681 |
| ARFGEF2 | 3382.206813 | 0.193884658 | 0.080345366 | 0.01313128 | 0.044173337 |
| SOX11 | 20217.5024 | -0.199827205 | 0.083515151 | 0.01371044 | 0.045725167 |
| TMED3 | 2654.988616 | -0.203306942 | 0.07749351 | 0.00711765 | 0.026816371 |
| ATRAID | 1503.121445 | -0.205205647 | 0.086569576 | 0.01439107 | 0.047509714 |
| TRMU | 3200.133371 | -0.207794046 | 0.08478697 | 0.01150413 | 0.039793273 |
| DCLRE1B | 2021.499003 | -0.211609087 | 0.089985164 | 0.01491792 | 0.048956506 |
| DGCR8 | 3864.856196 | -0.212672576 | 0.077478421 | 0.00486443 | 0.019664264 |
| ACAD9 | 2109.414116 | -0.214422179 | 0.082436804 | 0.00741843 | 0.027751955 |
| CDV3 | 7059.639035 | -0.214829262 | 0.081821211 | 0.00690191 | 0.026157546 |
| PRMT2 | 3279.3985 | -0.215042628 | 0.086730294 | 0.01045728 | 0.036879339 |
| TOR3A | 1469.723351 | -0.215405544 | 0.08242681 | 0.00714415 | 0.026887302 |
| NDUFA3 | 817.828011 | -0.215447739 | 0.089667839 | 0.01289263 | 0.043591592 |
| TPM4 | 10692.38898 | -0.215665776 | 0.082532026 | 0.00714369 | 0.026887302 |
| CARS2 | 2443.325175 | -0.216957636 | 0.080158273 | 0.00540773 | 0.021406198 |
| AMOTL1 | 5084.679476 | -0.217994916 | 0.088724966 | 0.01105275 | 0.038516695 |
| PIGX | 1284.165878 | -0.219299868 | 0.091837433 | 0.01329841 | 0.0446552 |
| RBM28 | 3339.698896 | -0.219828479 | 0.08049028 | 0.00499205 | 0.020069707 |
| SLC41A3 | 1236.036896 | -0.221434309 | 0.093657879 | 0.0140952 | 0.046752752 |
| ASXL1 | 6772.094316 | -0.221757689 | 0.076101955 | 0.00282056 | 0.01255727 |
| BRD9 | 3766.836612 | -0.222447857 | 0.093374495 | 0.01339889 | 0.044919273 |
| DEGS1 | 3163.431606 | -0.222832459 | 0.092409101 | 0.01237783 | 0.042196432 |
| ITPA | 1348.062629 | -0.223141904 | 0.089785337 | 0.01009719 | 0.035795118 |
| GOLIM4 | 5529.27841 | -0.223840522 | 0.089075668 | 0.00933145 | 0.033594538 |
| DCTD | 2599.740239 | -0.224198919 | 0.094489568 | 0.01368908 | 0.045676169 |
| RHOBTB3 | 8126.118314 | -0.224410002 | 0.088765277 | 0.00892872 | 0.032445702 |
| DNAAF2 | 1513.284458 | -0.226002162 | 0.092676533 | 0.01140529 | 0.039544495 |
| AHSA2P | 3483.692426 | -0.229605865 | 0.097482955 | 0.0141477 | 0.046900231 |
| BAX | 2184.824414 | -0.230288522 | 0.089607814 | 0.00781753 | 0.029018004 |
| KIF22 | 5685.575372 | -0.23047136 | 0.093380181 | 0.01040287 | 0.036737857 |
| DCAF11 | 2011.193978 | -0.231222535 | 0.095940211 | 0.0121681 | 0.041607037 |
| TDP1 | 1271.427655 | -0.232527898 | 0.095172731 | 0.01108203 | 0.038605949 |

| | | | | | |
|---|---|---|---|---|---|
| CD151 | 1895.333885 | -0.232928126 | 0.093930245 | 0.01001001 | 0.035533546 |
| PGRMC2 | 2618.957753 | -0.232975744 | 0.082129322 | 0.00350486 | 0.015002324 |
| SOD2 | 3175.432097 | -0.235018344 | 0.094323242 | 0.00963574 | 0.034464342 |
| SENP3 | 1380.310705 | -0.235165514 | 0.092049267 | 0.00806523 | 0.029776178 |
| PDCD2 | 3168.123124 | -0.236153275 | 0.089705637 | 0.00643212 | 0.024723392 |
| TSR1 | 3892.659224 | -0.23653802 | 0.09829732 | 0.0121253 | 0.041481382 |
| SLC9B2 | 938.7585351 | -0.237303009 | 0.097141368 | 0.01095853 | 0.038246985 |
| POLR3E | 2237.45032 | -0.237377301 | 0.085412494 | 0.00413985 | 0.017205317 |
| C22orf39 | 1683.710552 | -0.238129533 | 0.090311078 | 0.00632178 | 0.024385884 |
| BABAM1 | 929.6543649 | -0.238187882 | 0.100930036 | 0.01367085 | 0.045630139 |
| GET3 | 1786.42465 | -0.239811291 | 0.087609862 | 0.00467285 | 0.019013146 |
| EXOSC2 | 4648.29483 | -0.241229858 | 0.082435485 | 0.00259046 | 0.011676942 |
| TRAF5 | 2876.729185 | -0.241472276 | 0.102774491 | 0.01393015 | 0.046285293 |
| ADAM23 | 1801.715086 | -0.241628345 | 0.103987761 | 0.01490655 | 0.04893347 |
| CCAR2 | 4944.415755 | -0.241789555 | 0.102209915 | 0.01333742 | 0.044755225 |
| RHOA | 7735.502744 | -0.241904707 | 0.101370575 | 0.01261607 | 0.042841043 |
| LDB1 | 9862.845238 | -0.242047868 | 0.086089972 | 0.00370504 | 0.015680258 |
| COX8A | 2836.591488 | -0.242153732 | 0.104253483 | 0.01491978 | 0.048956506 |
| SAP30 | 1337.112658 | -0.242227211 | 0.088688008 | 0.00472957 | 0.019203628 |
| ENOSF1 | 2412.714078 | -0.242545888 | 0.089229978 | 0.00491394 | 0.019825453 |
| TRUB2 | 1897.744684 | -0.242961381 | 0.094409422 | 0.00749564 | 0.028005132 |
| NAF1 | 1084.195302 | -0.243146984 | 0.102013578 | 0.01267078 | 0.042968649 |
| ANKRD27 | 1421.749112 | -0.243223574 | 0.099456311 | 0.01071102 | 0.037593679 |
| MCF2L | 2966.17444 | -0.243230989 | 0.105179566 | 0.01527806 | 0.04992499 |
| PPM1L | 1428.771557 | -0.24371634 | 0.096622677 | 0.00864632 | 0.031553309 |
| LLPH | 1303.214449 | -0.244199266 | 0.093966597 | 0.00694851 | 0.026302838 |
| PLEKHJ1 | 1736.103304 | -0.24458703 | 0.093739145 | 0.00673549 | 0.025635501 |
| FTL | 13986.67965 | -0.244704762 | 0.090798459 | 0.00523655 | 0.020865022 |
| LPCAT4 | 1452.702111 | -0.244880165 | 0.091772807 | 0.0056644 | 0.022229516 |
| ATF7IP | 5086.994684 | -0.24494981 | 0.101696489 | 0.01178515 | 0.040553836 |
| APMAP | 2922.027363 | -0.245130665 | 0.090663331 | 0.00509691 | 0.020400762 |
| ARNT | 2679.714226 | -0.2451358 | 0.100985662 | 0.01119491 | 0.038927769 |
| SUMF2 | 3029.343077 | -0.245173914 | 0.08206088 | 0.00210481 | 0.009870756 |
| DDAH1 | 1668.933323 | -0.245660671 | 0.100634852 | 0.01077046 | 0.037763627 |
| RAMP2 | 726.5090289 | -0.245873688 | 0.089786587 | 0.00458521 | 0.018704658 |
| GSS | 1204.970101 | -0.246533375 | 0.092956041 | 0.00591454 | 0.023057161 |
| SFXN1 | 5007.948393 | -0.24707488 | 0.104376739 | 0.0130935 | 0.044067862 |
| RAB40C | 880.706292 | -0.247097628 | 0.096752158 | 0.0078387 | 0.029075585 |
| ITFG2 | 1660.506534 | -0.247712008 | 0.101288785 | 0.01058001 | 0.03723931 |
| CBX5 | 29562.38042 | -0.247738755 | 0.099675466 | 0.00948073 | 0.034036612 |
| C2orf68 | 2469.59666 | -0.248149048 | 0.098146747 | 0.00840191 | 0.030798014 |
| SH3GLB2 | 2023.929632 | -0.248715884 | 0.10036481 | 0.00964938 | 0.034503534 |
| C6orf120 | 1468.514031 | -0.248718429 | 0.106759258 | 0.0143867 | 0.047502914 |
| WAC-AS1 | 1044.546858 | -0.248870405 | 0.097837846 | 0.00803043 | 0.029669016 |
| GIGYF1 | 7139.57136 | -0.251883538 | 0.103536989 | 0.01083028 | 0.037934572 |
| THRAP3 | 6545.211702 | -0.251903625 | 0.099368479 | 0.00816116 | 0.030060257 |
| GATC | 1612.118304 | -0.251970837 | 0.092957423 | 0.00490461 | 0.019791694 |
| VHL | 3532.963245 | -0.252542162 | 0.095834068 | 0.00611571 | 0.023697638 |
| GPANK1 | 973.4819431 | -0.252909675 | 0.098487592 | 0.00741504 | 0.027744332 |
| SLC25A13 | 3011.106692 | -0.253804947 | 0.101738993 | 0.00908845 | 0.032915629 |

| | | | | | |
|---|---|---|---|---|---|
| PCGF1 | 2680.6511 | -0.25384539 | 0.096847885 | 0.00634926 | 0.024487278 |
| UBQLN4 | 4752.372834 | -0.253970207 | 0.096959041 | 0.00637874 | 0.024564118 |
| FN3KRP | 2462.578154 | -0.254370122 | 0.089059159 | 0.0031249 | 0.013665243 |
| ARRB2 | 2523.021542 | -0.254951135 | 0.106745973 | 0.01210138 | 0.041415224 |
| PSAP | 10017.49773 | -0.25506325 | 0.099089797 | 0.0072435 | 0.027211404 |
| SLC35E3 | 2644.84227 | -0.255477939 | 0.086677484 | 0.00233417 | 0.010723705 |
| NAGLU | 1196.121243 | -0.255681396 | 0.102484897 | 0.00903684 | 0.032755653 |
| TMEM138 | 1469.380759 | -0.255854993 | 0.103920641 | 0.00988797 | 0.035184489 |
| MMD | 10282.07207 | -0.256078439 | 0.094186991 | 0.00473274 | 0.019203628 |
| SLCO5A1 | 1158.161851 | -0.256148161 | 0.095450215 | 0.00525461 | 0.020924817 |
| AADAT | 2780.666283 | -0.256260506 | 0.106321154 | 0.01136918 | 0.039432602 |
| CILK1 | 2745.691294 | -0.25663123 | 0.105582961 | 0.01074884 | 0.037711269 |
| RBM23 | 3311.868056 | -0.25666448 | 0.097153819 | 0.00593121 | 0.023113405 |
| LINC02987 | 507.3429862 | -0.258190584 | 0.11030314 | 0.01360102 | 0.045441334 |
| LAS1L | 3600.715709 | -0.258333657 | 0.094012409 | 0.0043108 | 0.017806121 |
| ENO2 | 4873.613043 | -0.258829816 | 0.109498738 | 0.01277572 | 0.043267444 |
| HAND1 | 2463.036892 | -0.258852365 | 0.094367268 | 0.00436785 | 0.017974883 |
| LAGE3 | 563.0721839 | -0.259054468 | 0.107555382 | 0.01132868 | 0.039331933 |
| CLNS1A | 5406.508631 | -0.259990204 | 0.108840002 | 0.01191441 | 0.040930147 |
| ZMYND19 | 1863.219999 | -0.260092803 | 0.109691383 | 0.01248227 | 0.042469462 |
| PUM1 | 7923.34234 | -0.26026278 | 0.098891695 | 0.00604591 | 0.023475859 |
| KNSTRN | 2623.758779 | -0.26030819 | 0.108172462 | 0.0113526 | 0.039388362 |
| ATF6B | 2459.172048 | -0.260343814 | 0.103860056 | 0.00862824 | 0.031498506 |
| CMBL | 1502.105781 | -0.260635887 | 0.108557839 | 0.01151111 | 0.039810704 |
| SCG3 | 9410.371419 | -0.260960398 | 0.090440622 | 0.00280099 | 0.012499923 |
| ATP5IF1 | 1172.949671 | -0.260970714 | 0.087578905 | 0.00207219 | 0.00973999 |
| TMEM131L | 2305.006663 | -0.261161869 | 0.106129166 | 0.00977082 | 0.03486501 |
| USF1 | 2072.068908 | -0.261679329 | 0.085455476 | 0.00157921 | 0.007719233 |
| ANKRD40 | 5051.094634 | -0.26224689 | 0.094488676 | 0.00392272 | 0.016452546 |
| ARHGEF25 | 952.7581624 | -0.263003118 | 0.108647677 | 0.0108376 | 0.03794081 |
| ATP6V1F | 1478.692337 | -0.263035047 | 0.105732836 | 0.00902298 | 0.032713071 |
| STK25 | 6391.668839 | -0.263112898 | 0.093442258 | 0.00345866 | 0.014823635 |
| TMEM97 | 5086.428579 | -0.263554794 | 0.109170116 | 0.01101203 | 0.038388105 |
| CENPA | 964.1904144 | -0.263614705 | 0.097361319 | 0.00479243 | 0.019407558 |
| CENPH | 1833.518125 | -0.263766284 | 0.097541274 | 0.00483965 | 0.019575639 |
| NCBP2 | 6377.712463 | -0.264177496 | 0.105650047 | 0.00868031 | 0.031649277 |
| NARF | 5454.095907 | -0.264252557 | 0.081077502 | 0.00080096 | 0.004344745 |
| UBTD2 | 3114.242572 | -0.264644181 | 0.102929522 | 0.00710798 | 0.026790397 |
| METTL9 | 4409.61203 | -0.264728119 | 0.115493471 | 0.01513092 | 0.049546549 |
| BRD7 | 3801.045361 | -0.265241404 | 0.093709961 | 0.0032859 | 0.014195632 |
| FAM136A | 2332.299895 | -0.26562314 | 0.113572013 | 0.01336807 | 0.044837856 |
| C19orf48P | 4256.936889 | -0.265637208 | 0.084897706 | 0.00124961 | 0.006359052 |
| MRPS6 | 1248.386605 | -0.26595201 | 0.094100086 | 0.00332199 | 0.014327494 |
| MT-CO1 | 402312.2104 | -0.26615827 | 0.114424028 | 0.01379763 | 0.045956325 |
| DCXR | 3225.542118 | -0.266623417 | 0.090425551 | 0.00225663 | 0.01044696 |
| TADA3 | 2251.498654 | -0.26684733 | 0.095710184 | 0.00372578 | 0.015758944 |
| LINC02210 | 4254.435252 | -0.26687329 | 0.101673885 | 0.0060538 | 0.023502053 |
| GID8 | 4980.096345 | -0.267099584 | 0.087349463 | 0.00157858 | 0.007717988 |
| LIG3 | 3461.507347 | -0.267401032 | 0.109161471 | 0.00988944 | 0.035184489 |
| KLHL23 | 6072.56772 | -0.267583103 | 0.116722701 | 0.01498505 | 0.049154547 |

| | | | | | |
|---|---|---|---|---|---|
| PIN1 | 2685.324666 | -0.267636278 | 0.089298917 | 0.00192384 | 0.009162081 |
| RNF130 | 2396.044525 | -0.267887167 | 0.086977212 | 0.00146349 | 0.007239644 |
| GGA2 | 5457.507964 | -0.267993104 | 0.091325624 | 0.0023514 | 0.010780906 |
| E2F3 | 5719.220471 | -0.268414166 | 0.08564402 | 0.00121869 | 0.006229029 |
| TTPAL | 1465.076162 | -0.268475765 | 0.101186567 | 0.00554656 | 0.021825379 |
| UNC119 | 566.7633012 | -0.268986258 | 0.108431388 | 0.00903791 | 0.032755653 |
| MT-TP | 987.0774349 | -0.269244525 | 0.11472917 | 0.01294585 | 0.043692513 |
| SUMF1 | 437.6189009 | -0.269361591 | 0.117311199 | 0.01476192 | 0.048565269 |
| MRPL24 | 997.2512373 | -0.269387116 | 0.094798513 | 0.00313641 | 0.01369366 |
| AP2B1 | 9530.202612 | -0.269585889 | 0.116656426 | 0.01419823 | 0.047001347 |
| UBE3C | 4100.356179 | -0.269901779 | 0.095778155 | 0.0033697 | 0.014499799 |
| MTA2 | 5019.356135 | -0.270004454 | 0.112332699 | 0.01110887 | 0.03867979 |
| STMN3 | 8075.319529 | -0.270485067 | 0.103059437 | 0.00599389 | 0.023313479 |
| MAPKAPK5 | 4797.497905 | -0.270546086 | 0.098113368 | 0.00404507 | 0.016872777 |
| GSTK1 | 1183.99059 | -0.270674869 | 0.117178929 | 0.01418658 | 0.046982342 |
| DENND11 | 2355.948148 | -0.270801352 | 0.10588807 | 0.00725574 | 0.02724981 |
| ZNF532 | 9443.932243 | -0.271626652 | 0.085681059 | 0.00106824 | 0.005562723 |
| MPV17 | 1486.214444 | -0.272155307 | 0.097528861 | 0.00364195 | 0.015454609 |
| DHFR2 | 410.2616839 | -0.272290183 | 0.118629812 | 0.01465662 | 0.048254671 |
| CCDC144A | 545.8362998 | -0.272633682 | 0.118808869 | 0.01466188 | 0.04826427 |
| FAHD1 | 1026.478105 | -0.273108372 | 0.100971952 | 0.00470092 | 0.019109283 |
| TIPIN | 1093.039699 | -0.273225671 | 0.118364729 | 0.01412972 | 0.04685753 |
| PRUNE1 | 1775.676204 | -0.273791703 | 0.094190375 | 0.00252543 | 0.011433932 |
| TMEM64 | 1779.861277 | -0.273981035 | 0.105652619 | 0.0064906 | 0.024910898 |
| UXS1 | 1310.349903 | -0.273994675 | 0.107887474 | 0.0075552 | 0.028196937 |
| YWHAH | 5108.232456 | -0.274603092 | 0.10009936 | 0.00417146 | 0.017309018 |
| MED22 | 2063.747024 | -0.274665585 | 0.096783989 | 0.00312488 | 0.013665243 |
| USF2 | 2417.800706 | -0.275114704 | 0.111406792 | 0.00914724 | 0.033076127 |
| CLIC1 | 2400.651472 | -0.275710712 | 0.104731234 | 0.00576521 | 0.022569163 |
| PSME1 | 817.1240455 | -0.275935566 | 0.098656544 | 0.00353162 | 0.01509232 |
| PEX26 | 3358.092568 | -0.276006508 | 0.089264005 | 0.00137399 | 0.006871557 |
| TENT4A | 4706.25375 | -0.27614445 | 0.09475561 | 0.00244937 | 0.011153427 |
| CXADR | 3454.696248 | -0.276480423 | 0.114197322 | 0.01038692 | 0.036687831 |
| SKA2 | 8795.015382 | -0.276501217 | 0.114335304 | 0.01046307 | 0.036893453 |
| STK32C | 396.849641 | -0.276752787 | 0.117723757 | 0.01250624 | 0.042543956 |
| DHX35 | 1263.257744 | -0.276796253 | 0.117810812 | 0.01254941 | 0.042662602 |
| INA | 12356.79611 | -0.277034351 | 0.074996732 | 0.00015423 | 0.00107948 |
| COPS7B | 2363.453919 | -0.277364673 | 0.10236082 | 0.00457437 | 0.018667853 |
| PPT1 | 3660.854006 | -0.277892869 | 0.119830953 | 0.01353745 | 0.045258345 |
| PHB2 | 11947.23015 | -0.278503174 | 0.111743431 | 0.00849819 | 0.031078924 |
| ACTR1A | 3955.449642 | -0.278629625 | 0.109598356 | 0.00739211 | 0.027673647 |
| RPS9 | 18038.87445 | -0.278644964 | 0.116231868 | 0.01099297 | 0.038341161 |
| TMEM147-AS1 | 431.428128 | -0.278893348 | 0.115735813 | 0.01062546 | 0.037350775 |
| PRIM1 | 1704.130079 | -0.279295653 | 0.10771485 | 0.0063908 | 0.02460594 |
| NDUFA6 | 2204.807744 | -0.279302171 | 0.114735684 | 0.00993264 | 0.035301547 |
| ATP11C | 3927.278187 | -0.279829067 | 0.114735023 | 0.00979302 | 0.03492606 |
| GYG2 | 836.0729114 | -0.27997123 | 0.104980461 | 0.00514861 | 0.020554952 |
| SNX8 | 736.728794 | -0.27997224 | 0.113606355 | 0.00913296 | 0.033042831 |
| ZBED4 | 4592.752259 | -0.280167148 | 0.120078582 | 0.01295161 | 0.043697592 |
| ORC1 | 2872.384657 | -0.280399895 | 0.107224747 | 0.00597676 | 0.023260063 |

| | | | | | |
|---|---|---|---|---|---|
| GPS1 | 7115.085386 | -0.280514152 | 0.116134358 | 0.0104099 | 0.036749751 |
| ZNF765 | 520.5815698 | -0.280616867 | 0.121030767 | 0.01343307 | 0.045002071 |
| FAM53B | 1564.610299 | -0.281241664 | 0.11559137 | 0.00990371 | 0.035223072 |
| SERF2 | 6266.472204 | -0.281277245 | 0.09865871 | 0.00294194 | 0.012990775 |
| ASRGL1 | 1501.385085 | -0.281317258 | 0.099599087 | 0.00319307 | 0.013855268 |
| ERLIN1 | 1959.68596 | -0.281566273 | 0.118870177 | 0.01174749 | 0.040464174 |
| AKAP12 | 16816.80456 | -0.282331614 | 0.104101141 | 0.00447296 | 0.018337695 |
| MPHOSPH9 | 5097.902812 | -0.282489467 | 0.117108755 | 0.0104329 | 0.036812321 |
| PLA2G12A | 1052.401157 | -0.282503186 | 0.119980699 | 0.01215385 | 0.041565224 |
| PIP4K2B | 5817.947764 | -0.282863039 | 0.098391735 | 0.00271799 | 0.012169096 |
| CDC45 | 3055.125779 | -0.282890631 | 0.095543781 | 0.00206932 | 0.009730961 |
| EXOSC3 | 1245.347485 | -0.282958322 | 0.113923175 | 0.00857807 | 0.031348761 |
| MDM4 | 10599.00196 | -0.28299489 | 0.111643351 | 0.0074446 | 0.027834662 |
| CHCHD2 | 8512.152308 | -0.283000507 | 0.093585446 | 0.00168576 | 0.008165996 |
| AOPEP | 2313.450976 | -0.283006611 | 0.089717628 | 0.00109109 | 0.005663096 |
| SMC1A | 12965.81661 | -0.283068752 | 0.119336424 | 0.01158678 | 0.039991713 |
| PDZD4 | 5716.698653 | -0.283085275 | 0.123335158 | 0.01414626 | 0.046900231 |
| SNHG20 | 799.6982545 | -0.283173236 | 0.112783312 | 0.0079559 | 0.029430688 |
| CDK6 | 26179.72936 | -0.283408242 | 0.115796205 | 0.00945789 | 0.033966486 |
| PPP4C | 2956.430862 | -0.283472347 | 0.101090898 | 0.00337675 | 0.014521024 |
| BMAL2 | 845.3678844 | -0.28354407 | 0.114416995 | 0.00869445 | 0.031695196 |
| SLC25A24 | 4223.222861 | -0.283564699 | 0.111783338 | 0.00739107 | 0.027673647 |
| SPECC1 | 688.8462774 | -0.283803816 | 0.105204669 | 0.00464739 | 0.01892078 |
| PARD6G | 828.0238904 | -0.28401365 | 0.116610303 | 0.00974798 | 0.034807688 |
| SRI | 1634.924698 | -0.284522945 | 0.122822833 | 0.0133277 | 0.04473166 |
| IDH2 | 8698.953175 | -0.284620691 | 0.119485241 | 0.01122533 | 0.039012636 |
| HAUS8 | 657.8177407 | -0.284977437 | 0.104573709 | 0.00426716 | 0.017648816 |
| NTAQ1 | 670.6600372 | -0.284980993 | 0.114990217 | 0.00865034 | 0.031562383 |
| GTF2I | 6715.085234 | -0.285008033 | 0.092072826 | 0.00132294 | 0.006672794 |
| FANCC | 2080.651538 | -0.28512795 | 0.10101546 | 0.00317427 | 0.013809133 |
| PGLS | 712.0887022 | -0.285291595 | 0.118604695 | 0.01052627 | 0.037074091 |
| NAP1L4 | 8457.719989 | -0.285310172 | 0.092856899 | 0.00142639 | 0.007085073 |
| IGSF3 | 11115.76918 | -0.285392729 | 0.121772755 | 0.01238534 | 0.042202501 |
| BICD1 | 6733.032694 | -0.285416021 | 0.104339826 | 0.00413199 | 0.017179603 |
| PCCB | 1328.670113 | -0.286104681 | 0.093561771 | 0.00149374 | 0.007361659 |
| NLGN2 | 6271.822878 | -0.286129728 | 0.123792679 | 0.01343283 | 0.045002071 |
| EIF4B | 28383.91384 | -0.2861795 | 0.122997449 | 0.01290842 | 0.043627806 |
| KBTBD4 | 486.9707359 | -0.286233971 | 0.103109541 | 0.00364727 | 0.015470787 |
| DHODH | 369.8848043 | -0.286769648 | 0.109567641 | 0.00581989 | 0.022731297 |
| PABPC1 | 35959.91337 | -0.286843106 | 0.098804563 | 0.00245686 | 0.011182559 |
| ZNF589 | 711.815424 | -0.287099052 | 0.11908264 | 0.01030947 | 0.036458108 |
| MACROH2A1 | 12415.36226 | -0.287316544 | 0.11164431 | 0.00658338 | 0.025196483 |
| PSME3 | 11683.47128 | -0.287403863 | 0.120174622 | 0.0108444 | 0.037951698 |
| LRRC42 | 1892.612231 | -0.287525248 | 0.102484715 | 0.00332013 | 0.014322445 |
| RPL3P4 | 1813.64763 | -0.287609565 | 0.123979477 | 0.01307651 | 0.044032324 |
| PISD | 1945.334931 | -0.287789976 | 0.091690309 | 0.00113443 | 0.005862413 |
| PHOX2B-AS1 | 412.1555954 | -0.288010653 | 0.126449246 | 0.01454679 | 0.047977561 |
| SLC25A15 | 1415.310986 | -0.288537159 | 0.092546388 | 0.00121475 | 0.006210846 |
| NFATC2IP-AS1 | 630.1534001 | -0.288617072 | 0.123738234 | 0.01261087 | 0.042841043 |
| C18orf32 | 301.9173524 | -0.28866674 | 0.127864433 | 0.01526964 | 0.049921267 |

| | | | | | |
|---|---|---|---|---|---|
| RPL36 | 5708.228414 | -0.288686724 | 0.109405519 | 0.0054374 | 0.021498672 |
| PCBD2 | 949.9505637 | -0.288745905 | 0.103760692 | 0.00354476 | 0.015139015 |
| DIXDC1 | 2101.83749 | -0.288867545 | 0.1022918 | 0.00312453 | 0.013665243 |
| WDR82 | 11908.6807 | -0.288889334 | 0.107761021 | 0.00480509 | 0.019451146 |
| PAN2 | 2742.111188 | -0.288925092 | 0.092435607 | 0.00118136 | 0.006073369 |
| NUP43 | 4028.030706 | -0.289435042 | 0.099453541 | 0.00238302 | 0.010909259 |
| MECR | 656.2687651 | -0.289566331 | 0.103474612 | 0.00337132 | 0.014503744 |
| CENPM | 1144.288822 | -0.289702431 | 0.105070555 | 0.00381837 | 0.016100241 |
| HPS4 | 3045.204646 | -0.289984206 | 0.111726881 | 0.00613081 | 0.023751694 |
| NIPSNAP1 | 4424.936188 | -0.290017933 | 0.083845738 | 0.0003633 | 0.002232799 |
| C17orf58 | 407.3972608 | -0.290095567 | 0.114701843 | 0.00739021 | 0.027673647 |
| LAMTOR4 | 758.0947574 | -0.290102841 | 0.123787935 | 0.01219048 | 0.041669709 |
| NDRG3 | 1785.888803 | -0.290103741 | 0.103588276 | 0.00334389 | 0.014406833 |
| PES1 | 4079.517735 | -0.290132213 | 0.105236186 | 0.00381606 | 0.016093826 |
| SLC25A25-AS1 | 1487.387813 | -0.290145431 | 0.107376915 | 0.00449456 | 0.018404254 |
| RRP7A | 1913.267752 | -0.290188088 | 0.098448969 | 0.00211135 | 0.009892583 |
| DCBLD2 | 4010.79079 | -0.290424169 | 0.114683734 | 0.00731519 | 0.027425635 |
| FRZB | 1327.677824 | -0.290435023 | 0.10043696 | 0.00251848 | 0.01141254 |
| GTPBP2 | 4783.782074 | -0.290728612 | 0.105629597 | 0.00386212 | 0.016234794 |
| EIF4E3 | 1487.702611 | -0.290895644 | 0.119721654 | 0.00967621 | 0.034587431 |
| MORF4L1 | 13688.16043 | -0.290915563 | 0.128878181 | 0.01516051 | 0.049611828 |
| FAM83D | 2585.406682 | -0.291197602 | 0.110661221 | 0.0055074 | 0.021710289 |
| STIP1 | 11001.97174 | -0.291253007 | 0.100439911 | 0.00244913 | 0.011153427 |
| ERCC1 | 1473.641629 | -0.29126174 | 0.089506484 | 0.0007552 | 0.004151254 |
| SPAG5 | 6501.439396 | -0.291569624 | 0.108750998 | 0.00476016 | 0.019303482 |
| LYRM2 | 2339.194993 | -0.291737055 | 0.125772674 | 0.01289876 | 0.043605135 |
| LZTS2 | 3562.591135 | -0.291755124 | 0.106282522 | 0.0039341 | 0.016493553 |
| HP1BP3 | 8431.151951 | -0.292184888 | 0.081680604 | 0.00023184 | 0.001521746 |
| LEPROT | 2572.357815 | -0.292209025 | 0.113395004 | 0.0064158 | 0.024669864 |
| RBMX | 21618.66138 | -0.292391524 | 0.111320322 | 0.00556319 | 0.021869914 |
| RPS2P5 | 14631.16476 | -0.292581014 | 0.1059697 | 0.00374049 | 0.015814849 |
| TAF11 | 2341.125259 | -0.293226611 | 0.111202334 | 0.00538531 | 0.02133377 |
| ZNF512B | 5491.780063 | -0.293234416 | 0.123591323 | 0.01117486 | 0.038870077 |
| CTTN | 7192.37084 | -0.293280793 | 0.098032982 | 0.00181473 | 0.008696846 |
| DAZAP1 | 7897.08371 | -0.293293972 | 0.110191743 | 0.00500996 | 0.020125982 |
| CASKIN1 | 1123.284419 | -0.293398159 | 0.127958715 | 0.01373018 | 0.04578359 |
| CDH2 | 9141.72307 | -0.293439024 | 0.087819796 | 0.00055114 | 0.003172025 |
| FKRP | 1316.868443 | -0.293456689 | 0.118874787 | 0.00863382 | 0.031513288 |
| SYNCRIP | 12021.20266 | -0.293527011 | 0.125151927 | 0.01198863 | 0.041137033 |
| STX10 | 1201.991012 | -0.293551131 | 0.093176801 | 0.00107117 | 0.005573761 |
| ZFHX3 | 1860.05581 | -0.293593536 | 0.105343522 | 0.00344588 | 0.014781213 |
| EIF3F | 6387.451596 | -0.293805003 | 0.097860455 | 0.00175036 | 0.008425641 |
| UBE2G1 | 3329.172392 | -0.294265072 | 0.099875364 | 0.00209298 | 0.009827435 |
| ABCB8 | 1493.865331 | -0.29478206 | 0.109602152 | 0.00459413 | 0.018737344 |
| KLHL42 | 5031.235361 | -0.295206916 | 0.104932064 | 0.00316353 | 0.013782795 |
| M6PR | 2356.18604 | -0.295221046 | 0.105019855 | 0.00318494 | 0.013841369 |
| MACROD1 | 1006.317418 | -0.295314483 | 0.127736028 | 0.01298919 | 0.043788482 |
| LIN28B | 2540.832591 | -0.295528779 | 0.118666858 | 0.0080764 | 0.029805278 |
| ZNF691 | 561.2462966 | -0.295737407 | 0.128108614 | 0.01308391 | 0.044050027 |
| C15orf61 | 365.0303459 | -0.295962726 | 0.121720476 | 0.00946437 | 0.033983809 |

| | | | | | |
|---|---|---|---|---|---|
| XPO5 | 15050.81813 | -0.296035941 | 0.124657491 | 0.01100197 | 0.038359516 |
| GAP43 | 1069.276945 | -0.296283375 | 0.125022407 | 0.01113635 | 0.038749215 |
| CENPO | 2076.8476 | -0.296355308 | 0.116313185 | 0.00686572 | 0.026034793 |
| CINP | 755.7021176 | -0.296388375 | 0.100172006 | 0.00199756 | 0.009438161 |
| CCDC12 | 822.9930471 | -0.296649023 | 0.111192739 | 0.00486542 | 0.01966441 |
| TUSC2 | 762.6016363 | -0.296788157 | 0.094820463 | 0.00113641 | 0.00587018 |
| HSP90AB1 | 166927.421 | -0.296811678 | 0.123848079 | 0.01035824 | 0.036605407 |
| QTRT2 | 2020.351667 | -0.296852741 | 0.10098838 | 0.00212159 | 0.009931342 |
| SPTLC2 | 3244.596505 | -0.296883632 | 0.104393882 | 0.0028638 | 0.012705758 |
| CENPP | 802.4357565 | -0.296933213 | 0.122616762 | 0.00968501 | 0.034602955 |
| FAM217B | 1639.631398 | -0.297313847 | 0.126321898 | 0.01157673 | 0.039963706 |
| BEX3 | 7474.880041 | -0.297441019 | 0.112197959 | 0.0050965 | 0.020400762 |
| MAGEF1 | 2570.917077 | -0.297524536 | 0.102894401 | 0.00246337 | 0.011204783 |
| MTG1 | 481.868158 | -0.297547253 | 0.108687969 | 0.00394741 | 0.016544221 |
| ANKRD39 | 332.9523756 | -0.297675335 | 0.113938938 | 0.00568994 | 0.022325495 |
| WARS2 | 467.3683911 | -0.297827003 | 0.125950252 | 0.01122738 | 0.039013159 |
| SFI1 | 1963.766368 | -0.297908799 | 0.087011194 | 0.00040364 | 0.002439137 |
| PANK4 | 583.1434835 | -0.297957889 | 0.107554515 | 0.00357348 | 0.015239534 |
| RFPL1S | 328.3415459 | -0.29856367 | 0.116908839 | 0.00670189 | 0.025540696 |
| PGK1 | 11456.38471 | -0.298632068 | 0.12014846 | 0.00809764 | 0.029852965 |
| ORC6 | 4954.112534 | -0.298792493 | 0.105844694 | 0.00303574 | 0.013341932 |
| SEM1 | 1991.470075 | -0.2988127 | 0.124666913 | 0.01027642 | 0.036349088 |
| HEATR3 | 1494.87874 | -0.29883733 | 0.114515728 | 0.00571658 | 0.022404369 |
| SCRN2 | 1384.505958 | -0.298846412 | 0.097065088 | 0.00133982 | 0.006734816 |
| SMIM12 | 1105.72314 | -0.298868203 | 0.099830085 | 0.00177061 | 0.008505192 |
| LINC00641 | 3511.608455 | -0.299332877 | 0.114919452 | 0.0057867 | 0.02263152 |
| HADH | 3025.479708 | -0.299494663 | 0.117351732 | 0.0067124 | 0.025571294 |
| CHD4 | 1618.131606 | -0.299760569 | 0.126199896 | 0.01084259 | 0.037951698 |
| GNAI2 | 2954.017027 | -0.299955521 | 0.100054504 | 0.00174083 | 0.008387646 |
| RBM3 | 11105.34625 | -0.300061708 | 0.114402811 | 0.00547993 | 0.02162527 |
| TBX2-AS1 | 903.4081402 | -0.300464511 | 0.130784295 | 0.01323029 | 0.044462774 |
| TTN | 1871.636701 | -0.300742868 | 0.116028113 | 0.00597161 | 0.023248797 |
| COMMD9 | 1371.050725 | -0.300813782 | 0.129538443 | 0.01240042 | 0.042225896 |
| STX2 | 3578.779983 | -0.301071416 | 0.098312387 | 0.00140468 | 0.007001384 |
| APH1A | 6366.171618 | -0.301104398 | 0.085779653 | 0.00029046 | 0.00183901 |
| NAGA | 824.0375994 | -0.301330483 | 0.122202918 | 0.00846407 | 0.030973043 |
| DGCR2 | 2833.287676 | -0.301384625 | 0.130575067 | 0.01282911 | 0.043398236 |
| NUTF2 | 4117.899486 | -0.301521348 | 0.104548433 | 0.0024893 | 0.011295221 |
| GLRX | 449.4459192 | -0.301538161 | 0.10471005 | 0.00252279 | 0.011427012 |
| MYH10 | 13046.51804 | -0.30160583 | 0.103153711 | 0.00219538 | 0.010213958 |
| ZNF771 | 649.1083527 | -0.301831427 | 0.130459446 | 0.01262974 | 0.0428719 |
| DDX23 | 6916.038873 | -0.301948636 | 0.101348381 | 0.00183677 | 0.008794271 |
| MRPL42 | 6473.905629 | -0.302613477 | 0.127702815 | 0.01089009 | 0.038059783 |
| RPS2 | 70062.35156 | -0.302655432 | 0.102228701 | 0.00194625 | 0.009243117 |
| SMARCA5-AS1 | 189.580053 | -0.302658581 | 0.1289387 | 0.01154243 | 0.039885473 |
| DBF4 | 2398.806303 | -0.302810785 | 0.123951231 | 0.00895607 | 0.032507428 |
| FAM72B | 709.8251694 | -0.302989945 | 0.106768528 | 0.00285962 | 0.012689935 |
| STK11IP | 794.3787571 | -0.303115521 | 0.092600702 | 0.00068046 | 0.003802463 |
| TMEM250 | 835.6970721 | -0.303283037 | 0.121722213 | 0.00783412 | 0.029063843 |
| ZFAND5 | 8406.250822 | -0.30328844 | 0.08871228 | 0.00040429 | 0.00244234 |

| | | | | | |
|---|---|---|---|---|---|
| PNKP | 1690.226442 | -0.303474656 | 0.130860224 | 0.01237798 | 0.042196432 |
| ELP6 | 1780.261954 | -0.303602429 | 0.104485358 | 0.00230966 | 0.010639737 |
| RPL8 | 29168.59907 | -0.303673074 | 0.102834084 | 0.00198705 | 0.009399976 |
| SNX33 | 823.8029387 | -0.303700125 | 0.128417988 | 0.01098199 | 0.038322345 |
| RTN4 | 11498.69591 | -0.303789347 | 0.116326834 | 0.0055885 | 0.021956795 |
| RABGGTA | 490.968281 | -0.303853899 | 0.13203098 | 0.01293281 | 0.043675587 |
| BAZ1A | 6650.106151 | -0.303901981 | 0.122108467 | 0.00787729 | 0.029176613 |
| DHRS11 | 552.9223693 | -0.303912174 | 0.115495287 | 0.00527642 | 0.020996103 |
| C20orf96 | 1048.611468 | -0.303949567 | 0.100950291 | 0.00164751 | 0.008005517 |
| SLC35B4 | 3497.403711 | -0.304062643 | 0.097886408 | 0.00120234 | 0.006162757 |
| RAB5C | 868.1324294 | -0.304161205 | 0.114797411 | 0.00500225 | 0.02009891 |
| HIP1R | 1455.683297 | -0.30416383 | 0.129581869 | 0.01147944 | 0.039721223 |
| WARS2-AS1 | 1130.512461 | -0.30418554 | 0.121921776 | 0.00773798 | 0.028774657 |
| MACROH2A2 | 4682.278937 | -0.304399233 | 0.106066508 | 0.00257687 | 0.011625866 |
| TADA2A | 931.7993661 | -0.304408403 | 0.12926214 | 0.01124179 | 0.039050027 |
| NCOA5 | 2833.348755 | -0.304929778 | 0.113162377 | 0.00437359 | 0.017991287 |
| CNPY4 | 481.0835036 | -0.304968509 | 0.128077644 | 0.0104765 | 0.036934479 |
| AMD1 | 7082.043511 | -0.304994157 | 0.135811813 | 0.01481649 | 0.04871085 |
| VPS28 | 1706.822997 | -0.305078124 | 0.090859283 | 0.00050141 | 0.002935081 |
| MYCBP | 657.165013 | -0.305211368 | 0.13497606 | 0.01424134 | 0.047121306 |
| NFATC2IP | 3117.549303 | -0.305560965 | 0.088550009 | 0.00035711 | 0.002200682 |
| DPYSL2 | 7726.629117 | -0.305772782 | 0.124006297 | 0.00833079 | 0.030586399 |
| FAM72A | 420.1855272 | -0.305787294 | 0.131356619 | 0.01199556 | 0.041153928 |
| NDUFB11 | 3350.833947 | -0.305838287 | 0.117026241 | 0.00551836 | 0.021726905 |
| SF3B5 | 2630.639041 | -0.306018296 | 0.120464115 | 0.00677979 | 0.025766601 |
| RBBP4 | 12751.36613 | -0.306126306 | 0.119685856 | 0.0064548 | 0.024801274 |
| MTA1 | 8357.596319 | -0.306189068 | 0.123673781 | 0.00809515 | 0.029849125 |
| VGLL4 | 1191.31339 | -0.306389431 | 0.136245162 | 0.01462405 | 0.048186007 |
| RALY | 4848.362903 | -0.306392045 | 0.118832343 | 0.00608507 | 0.02359671 |
| WDR3 | 6050.806394 | -0.30655621 | 0.137195592 | 0.0151456 | 0.049578829 |
| CAPN2 | 4077.370008 | -0.306778217 | 0.090777681 | 0.00046117 | 0.002733008 |
| TP53RK | 1671.444971 | -0.307085957 | 0.100536011 | 0.00141373 | 0.007037504 |
| DUS1L | 6083.353082 | -0.307215485 | 0.110262234 | 0.00330055 | 0.014249957 |
| NT5C3B | 2759.284003 | -0.307267337 | 0.085310857 | 0.00020163 | 0.001348373 |
| PMF1 | 1062.165224 | -0.307366171 | 0.089937356 | 0.00040089 | 0.002425414 |
| TMEM147 | 1370.945269 | -0.307384682 | 0.105105039 | 0.00214875 | 0.010042459 |
| PNKD | 806.0351261 | -0.307434164 | 0.114034631 | 0.00431846 | 0.017819733 |
| INKA2 | 3461.272479 | -0.307853952 | 0.110065588 | 0.00318727 | 0.01384417 |
| NMT1 | 8888.061117 | -0.308045552 | 0.117549229 | 0.0053644 | 0.021263197 |
| KIAA1614 | 417.4067132 | -0.308147271 | 0.12365535 | 0.00768959 | 0.028625746 |
| MCUR1 | 2874.325871 | -0.308219822 | 0.079394635 | 6.63E-05 | 0.000519578 |
| TRIR | 3954.091532 | -0.308488458 | 0.10812161 | 0.00267688 | 0.012011217 |
| NR2F6 | 1034.684585 | -0.30863487 | 0.137220665 | 0.01448388 | 0.047785411 |
| TSPAN11 | 3414.415382 | -0.308667435 | 0.134947075 | 0.01316012 | 0.044255882 |
| CTCF | 4903.233805 | -0.308734279 | 0.119272025 | 0.00586371 | 0.022884616 |
| ZNF738 | 3759.091541 | -0.308873349 | 0.113238411 | 0.00391263 | 0.016413589 |
| MDN1 | 11566.12846 | -0.309093431 | 0.11518139 | 0.00445297 | 0.018259413 |
| QSER1 | 6948.076269 | -0.309417473 | 0.116960564 | 0.00496867 | 0.020009572 |
| TBCB | 2623.106769 | -0.309541577 | 0.101790856 | 0.00146595 | 0.007250065 |
| TALDO1 | 2733.072808 | -0.309691992 | 0.109927716 | 0.00297755 | 0.013122655 |

| | | | | | |
|---|---|---|---|---|---|
| TAF5L | 2198.406209 | -0.309698854 | 0.112069166 | 0.00350506 | 0.015002324 |
| SUV39H1 | 562.5921663 | -0.309706451 | 0.129443124 | 0.00998644 | 0.035462122 |
| ZNF43 | 3034.012532 | -0.309760715 | 0.127023301 | 0.00883417 | 0.032175662 |
| SLC39A6 | 8793.094382 | -0.30989315 | 0.120934926 | 0.0062839 | 0.024285292 |
| WDR12 | 3743.369939 | -0.310022197 | 0.104507434 | 0.00186314 | 0.008908091 |
| BVES | 1837.932282 | -0.310255492 | 0.10746325 | 0.00239412 | 0.01094033 |
| REC8 | 6274.717422 | -0.310286324 | 0.129076092 | 0.00967161 | 0.034577017 |
| KCNC1 | 2407.178303 | -0.310311086 | 0.12388708 | 0.00736538 | 0.027588659 |
| BCCIP | 3517.628808 | -0.310312617 | 0.136428744 | 0.01350499 | 0.045186545 |
| POLR1B | 4030.693773 | -0.310633876 | 0.100968247 | 0.00129892 | 0.006569205 |
| ZNF587B | 1015.468782 | -0.310635453 | 0.123938758 | 0.00732484 | 0.027451777 |
| HNRNPD | 17026.91826 | -0.310820418 | 0.128803698 | 0.00941873 | 0.033843569 |
| ACTN4 | 5342.413321 | -0.310895475 | 0.109373233 | 0.00274353 | 0.012264751 |
| ARHGEF17 | 3672.886831 | -0.31123957 | 0.131431028 | 0.01058842 | 0.037258767 |
| ZNF8-DT | 269.0037821 | -0.311335308 | 0.122534872 | 0.00664184 | 0.025352939 |
| IFT25 | 1102.352779 | -0.31137721 | 0.123151272 | 0.00687323 | 0.026053641 |
| KRBA1 | 724.4494853 | -0.311407327 | 0.135764449 | 0.01281159 | 0.043353208 |
| RAMP2-AS1 | 908.9105853 | -0.311565649 | 0.119224242 | 0.0054088 | 0.021406198 |
| ADGRL2 | 9788.774668 | -0.311623581 | 0.117820047 | 0.00493777 | 0.019902048 |
| MTCL3 | 961.7431523 | -0.311637837 | 0.123776504 | 0.00707234 | 0.026675648 |
| LBX2 | 480.7283491 | -0.311753298 | 0.124253338 | 0.00724163 | 0.02720938 |
| TMED4 | 3607.476492 | -0.312007952 | 0.08468266 | 0.00014433 | 0.001019545 |
| TAF10 | 321.5068461 | -0.31219892 | 0.118636324 | 0.00512081 | 0.020471278 |
| ITPK1 | 1497.83485 | -0.312210185 | 0.138963078 | 0.01437 | 0.047463012 |
| POGK | 6270.06674 | -0.312423854 | 0.102401149 | 0.00140477 | 0.007001384 |
| ALG3 | 1585.909246 | -0.312545472 | 0.112808458 | 0.00339656 | 0.014587885 |
| TBCA | 2913.298392 | -0.312623643 | 0.113564032 | 0.00358154 | 0.015270725 |
| DHX30 | 6388.819303 | -0.312693229 | 0.122191024 | 0.00628002 | 0.024274878 |
| DNAJC14 | 1040.795176 | -0.312787088 | 0.134695693 | 0.01185224 | 0.040743811 |
| PLCG1 | 10385.27303 | -0.312831539 | 0.112383521 | 0.00326232 | 0.014108592 |
| ZNF22-AS1 | 185.1539483 | -0.3128366 | 0.135060226 | 0.01203022 | 0.041238451 |
| NAT10 | 3839.970391 | -0.313017635 | 0.118900559 | 0.00509043 | 0.020393439 |
| FZD2 | 1400.560258 | -0.313241356 | 0.13834515 | 0.01370107 | 0.045708755 |
| TMEM94 | 6025.105638 | -0.313262112 | 0.135942998 | 0.01238047 | 0.042196432 |
| MTCH2 | 3861.508567 | -0.313307439 | 0.124924776 | 0.00722067 | 0.027150473 |
| TAF8 | 1387.59277 | -0.313311031 | 0.085652024 | 0.00015931 | 0.001110554 |
| TPM3 | 11254.17449 | -0.313334623 | 0.112046166 | 0.00313191 | 0.0136805 |
| ZNF367 | 1538.78263 | -0.313685658 | 0.123497596 | 0.00659873 | 0.025227027 |
| SRSF9 | 4445.993214 | -0.313771726 | 0.121881287 | 0.00599103 | 0.023306778 |
| C14orf93 | 485.70494 | -0.313773054 | 0.125610136 | 0.00740831 | 0.027724203 |
| MICAL2 | 216.8726365 | -0.313801762 | 0.141159629 | 0.01513922 | 0.04956581 |
| FBXO41 | 2483.170456 | -0.31382318 | 0.140321633 | 0.01464532 | 0.048240648 |
| PCDHGA6 | 432.4670005 | -0.313969715 | 0.120465518 | 0.00546936 | 0.021595996 |
| CCT7 | 15861.54934 | -0.313985392 | 0.120496629 | 0.00547751 | 0.021621319 |
| UBA52 | 14301.34905 | -0.314012915 | 0.131575179 | 0.00998146 | 0.035450561 |
| GPR137B | 348.050768 | -0.314020958 | 0.133643477 | 0.01098863 | 0.038332501 |
| NUP85 | 6005.40388 | -0.314262853 | 0.091012615 | 0.0003444 | 0.002131303 |
| DEPDC5 | 658.9640991 | -0.314401555 | 0.11938678 | 0.00505218 | 0.020263899 |
| STMN2 | 6007.051145 | -0.314620725 | 0.118782848 | 0.0048312 | 0.019552985 |
| GAMT | 1100.219037 | -0.314665404 | 0.123798869 | 0.0065431 | 0.025065647 |

| | | | | | |
|---|---|---|---|---|---|
| POGLUT3 | 392.1360374 | -0.314673391 | 0.128932926 | 0.00862612 | 0.031496354 |
| HNRNPCP2 | 664.4469905 | -0.31477877 | 0.135563596 | 0.01176557 | 0.040506812 |
| GTSE1 | 4710.65306 | -0.314814983 | 0.103824295 | 0.00148186 | 0.007312912 |
| MGST3 | 1395.06488 | -0.31486425 | 0.132083815 | 0.01002067 | 0.035565249 |
| PCDHB14 | 968.9484268 | -0.315058013 | 0.112877175 | 0.00316348 | 0.013782795 |
| DOCK10 | 1383.31722 | -0.315188165 | 0.137956508 | 0.01291395 | 0.043627806 |
| DMAC2 | 1478.767586 | -0.315189806 | 0.090552097 | 0.00030989 | 0.001947594 |
| MSRB1 | 493.4815737 | -0.315199487 | 0.120991442 | 0.00546279 | 0.021580702 |
| RPUSD4 | 1538.599069 | -0.315289212 | 0.10535986 | 0.00168303 | 0.008155074 |
| NENF | 1002.020979 | -0.315399386 | 0.105087024 | 0.0016351 | 0.007956495 |
| MOB3A | 681.181743 | -0.315401538 | 0.141647672 | 0.014908 | 0.04893347 |
| VMA21 | 3340.61725 | -0.315440904 | 0.142251977 | 0.01524523 | 0.049857339 |
| MAP4K4 | 21285.77877 | -0.315763237 | 0.108185603 | 0.00212655 | 0.009952304 |
| MCM10 | 3876.503441 | -0.315879086 | 0.124862039 | 0.00673275 | 0.025629821 |
| ATP5MF | 1856.345997 | -0.315879501 | 0.104876637 | 0.00157724 | 0.007716151 |
| CTDSPL | 1039.76739 | -0.315987479 | 0.107401822 | 0.00197314 | 0.009342787 |
| TMEM141 | 220.9805713 | -0.316520386 | 0.132713258 | 0.00992587 | 0.035289689 |
| CNP | 2026.021258 | -0.316558116 | 0.120576195 | 0.00513013 | 0.020491451 |
| ARHGAP11A | 3544.884591 | -0.317085747 | 0.109571281 | 0.00228862 | 0.010559361 |
| DCAKD | 2219.33371 | -0.317156978 | 0.09773598 | 0.00071735 | 0.003970578 |
| POLD3 | 2234.920916 | -0.317184696 | 0.112930963 | 0.00297687 | 0.013122492 |
| QDPR | 2007.647709 | -0.317281111 | 0.118826006 | 0.00449609 | 0.018406212 |
| NOSTRIN | 419.9034081 | -0.317465359 | 0.137535053 | 0.01205652 | 0.041307941 |
| CATSPER2 | 557.0042031 | -0.317546464 | 0.141564764 | 0.01419104 | 0.046985097 |
| SDHC | 2924.572084 | -0.317689479 | 0.129992156 | 0.00845142 | 0.030951904 |
| RAD51 | 1269.788243 | -0.317793558 | 0.113878596 | 0.00313759 | 0.013695902 |
| STEEP1 | 848.8696542 | -0.317965502 | 0.112887359 | 0.00289694 | 0.012830653 |
| EBNA1BP2 | 2252.076193 | -0.317968465 | 0.139533347 | 0.01296154 | 0.043713937 |
| KIAA1143 | 3549.51821 | -0.318490646 | 0.118976446 | 0.00438843 | 0.018037923 |
| TRIM52-AS1 | 287.6195918 | -0.318698224 | 0.126428504 | 0.00683005 | 0.025932993 |
| SHD | 4493.844669 | -0.318764386 | 0.139674961 | 0.01281041 | 0.043353208 |
| RUVBL1 | 4231.622187 | -0.318849882 | 0.132138175 | 0.00913596 | 0.033042831 |
| ARNT2-DT | 645.1394437 | -0.318861155 | 0.121149107 | 0.00499115 | 0.020069707 |
| ARHGAP28 | 480.1149383 | -0.319088417 | 0.115914509 | 0.00349971 | 0.014990209 |
| FAM120A | 4125.600962 | -0.319322992 | 0.085045321 | 0.0001069 | 0.000784219 |
| RUSC1 | 1923.155322 | -0.319340094 | 0.105624117 | 0.00150138 | 0.007391518 |
| RBMXL1 | 3154.814784 | -0.31954138 | 0.132422029 | 0.00911004 | 0.03298219 |
| NSMCE4A | 1936.806952 | -0.319641929 | 0.094505508 | 0.00043764 | 0.002618463 |
| CUL4A | 5032.957355 | -0.31976728 | 0.091364901 | 0.0002843 | 0.001806573 |
| PRDX3 | 8606.990707 | -0.319888464 | 0.139002093 | 0.01215133 | 0.041563534 |
| PCBD1 | 703.9940037 | -0.31999827 | 0.101140773 | 0.00093872 | 0.004993223 |
| OSGEPL1 | 683.8474204 | -0.320084066 | 0.144434624 | 0.01500695 | 0.04919542 |
| GALNT16 | 1327.65236 | -0.320118217 | 0.098700869 | 0.00071441 | 0.003955624 |
| GDI1 | 9889.313949 | -0.320145913 | 0.115890804 | 0.00338648 | 0.014553735 |
| NDUFAF3 | 1297.678265 | -0.320386679 | 0.098877764 | 0.00072144 | 0.00398703 |
| DRC3 | 334.0324027 | -0.320454013 | 0.134705866 | 0.0099301 | 0.035298619 |
| VTI1B | 1504.184828 | -0.320670301 | 0.124480672 | 0.00581109 | 0.022709254 |
| MRPL34 | 684.7094335 | -0.320781031 | 0.10749613 | 0.00169616 | 0.008203155 |
| DCPS | 565.9022295 | -0.320789804 | 0.103295692 | 0.00113923 | 0.005880415 |
| TRMO | 390.5959608 | -0.320823436 | 0.127474382 | 0.00684995 | 0.025989354 |

| | | | | | |
|---|---|---|---|---|---|
| MPP1 | 613.1151526 | -0.321032802 | 0.13705073 | 0.01089006 | 0.038059783 |
| TBC1D24 | 1575.898891 | -0.321128858 | 0.100809581 | 0.00086874 | 0.004664411 |
| RNF4 | 2250.926971 | -0.321370782 | 0.103010171 | 0.00108403 | 0.005632114 |
| DCAF7 | 20725.57328 | -0.321432567 | 0.115955398 | 0.00327513 | 0.01415803 |
| UROD | 1136.117287 | -0.321468508 | 0.130386134 | 0.0078579 | 0.02912053 |
| PSMB3 | 2921.750928 | -0.321537107 | 0.126500652 | 0.00637451 | 0.024552446 |
| CBX6 | 3234.881775 | -0.321611091 | 0.1155912 | 0.00317315 | 0.01380897 |
| HNRNPK | 37544.89613 | -0.321704518 | 0.119031859 | 0.00402148 | 0.016798218 |
| UBAC1 | 1604.932041 | -0.321796336 | 0.095430197 | 0.00045071 | 0.002679496 |
| C19orf53 | 1542.289992 | -0.321927655 | 0.10103788 | 0.00086424 | 0.004642692 |
| AKR1A1 | 2235.998131 | -0.321995336 | 0.107463007 | 0.00162362 | 0.007906256 |
| IDH3G | 1496.191779 | -0.322109384 | 0.105367986 | 0.00133184 | 0.0067111 |
| HNRNPDL | 20435.30944 | -0.322268788 | 0.121641295 | 0.00468752 | 0.019068229 |
| LARP7 | 1425.326634 | -0.322298105 | 0.143369163 | 0.01374811 | 0.045828531 |
| FMNL2 | 6007.514264 | -0.322298425 | 0.105831668 | 0.00138265 | 0.006906195 |
| MCM9 | 595.0045362 | -0.322333796 | 0.10661453 | 0.00148563 | 0.007328011 |
| SCARB2 | 10347.46414 | -0.322415538 | 0.110238134 | 0.00203789 | 0.00960062 |
| UBQLN2 | 2788.104962 | -0.322773192 | 0.107626915 | 0.00160525 | 0.007826059 |
| TMEM209 | 1034.218724 | -0.322786748 | 0.11638776 | 0.00324573 | 0.014048234 |
| FOXO3B | 1505.283885 | -0.322858099 | 0.134370859 | 0.0092395 | 0.033356919 |
| ANP32B | 5935.348107 | -0.322990094 | 0.126868276 | 0.00626729 | 0.024230223 |
| GINS1 | 3097.136679 | -0.323022957 | 0.102593313 | 0.00097832 | 0.00516583 |
| NFS1 | 272.278735 | -0.323057441 | 0.120975912 | 0.00439665 | 0.018068092 |
| MBTPS1 | 2413.715953 | -0.323061648 | 0.105161598 | 0.00126311 | 0.006419813 |
| NUP210 | 6418.264247 | -0.323092705 | 0.12686994 | 0.00625131 | 0.024177519 |
| ATPSCKMT | 680.9472995 | -0.323183117 | 0.118014241 | 0.00359728 | 0.015318795 |
| SPHK2 | 1169.329697 | -0.323343032 | 0.129836875 | 0.00729162 | 0.027355778 |
| HNRNPUL1 | 9619.488461 | -0.323410342 | 0.123602426 | 0.00512781 | 0.020487309 |
| NDNF | 442.3841184 | -0.323437575 | 0.142443629 | 0.01293597 | 0.043675587 |
| SYNE2 | 5197.047383 | -0.323454637 | 0.097079963 | 0.00051748 | 0.003011182 |
| PDE7A | 4081.696336 | -0.323528234 | 0.09461466 | 0.00037748 | 0.002308879 |
| CNOT9 | 5105.88449 | -0.32358127 | 0.100686431 | 0.00078186 | 0.004264137 |
| UQCRC1 | 5174.502718 | -0.323746207 | 0.101025992 | 0.00080642 | 0.004367451 |
| MLLT6 | 8845.873818 | -0.323758267 | 0.104741624 | 0.00118327 | 0.006080153 |
| ZNF107 | 2168.122086 | -0.323824151 | 0.10232427 | 0.00092379 | 0.004925294 |
| CEP41 | 1369.396653 | -0.323921525 | 0.108916411 | 0.00173242 | 0.008351026 |
| DNMT1 | 14087.58369 | -0.324153655 | 0.135642723 | 0.00951037 | 0.034107293 |
| ALKBH2 | 1781.737698 | -0.324304262 | 0.114454549 | 0.00268885 | 0.012057273 |
| NRAS | 7797.507747 | -0.32439599 | 0.132016219 | 0.0079428 | 0.029392791 |
| ZSWIM7 | 590.236689 | -0.324463565 | 0.113987933 | 0.00258202 | 0.011646549 |
| SLAIN1 | 3768.57635 | -0.324566788 | 0.114281211 | 0.00263239 | 0.011837404 |
| DNAJC9 | 1440.29263 | -0.324606176 | 0.119625604 | 0.00385303 | 0.016206523 |
| ZBTB18 | 11846.28324 | -0.324677068 | 0.119950568 | 0.00392946 | 0.016477455 |
| MRPS17 | 352.5316171 | -0.324704186 | 0.119800185 | 0.00388737 | 0.016330919 |
| ARPC5 | 3995.991107 | -0.324772697 | 0.12640205 | 0.00582897 | 0.022758118 |
| JAG2 | 1042.089181 | -0.324833178 | 0.143476088 | 0.01307442 | 0.044032324 |
| MTAP | 2572.176871 | -0.324901926 | 0.131782016 | 0.00775357 | 0.028817001 |
| BLTP2 | 7219.590829 | -0.324973802 | 0.113641967 | 0.00247461 | 0.011240971 |
| GLRX5 | 1735.076972 | -0.325063443 | 0.127885451 | 0.00628711 | 0.024293137 |
| NIP7 | 1737.075406 | -0.325437672 | 0.111409885 | 0.00203892 | 0.00960062 |

| | | | | | |
|---|---|---|---|---|---|
| MED24 | 5079.02309 | -0.325468813 | 0.118675257 | 0.00352436 | 0.015066293 |
| SEH1L | 3930.443716 | -0.325510499 | 0.133242895 | 0.00821575 | 0.030228795 |
| POMK | 3376.905915 | -0.325563093 | 0.145742777 | 0.01404367 | 0.046617269 |
| FAM72D | 377.4851696 | -0.325658517 | 0.129141033 | 0.006631 | 0.025326867 |
| NABP2 | 4036.78099 | -0.3256635 | 0.106946048 | 0.00136735 | 0.006853091 |
| CHRNA3 | 15368.07963 | -0.325695422 | 0.140809448 | 0.01151871 | 0.039823603 |
| CORO1C | 6884.232324 | -0.325993742 | 0.122447853 | 0.00445155 | 0.018257206 |
| LRRC7 | 260.7300824 | -0.326035359 | 0.142540532 | 0.01227334 | 0.04190421 |
| TGIF2 | 2321.36265 | -0.326059921 | 0.12774604 | 0.00608121 | 0.023589104 |
| COX7B | 1584.457566 | -0.326206668 | 0.116920934 | 0.00304816 | 0.013390812 |
| ZNF611 | 584.6624626 | -0.326218138 | 0.135703398 | 0.00908735 | 0.032915629 |
| DECR2 | 428.1602396 | -0.326542985 | 0.125648661 | 0.00532739 | 0.021132786 |
| GUCY1A1 | 874.6097398 | -0.326589005 | 0.129484672 | 0.00659774 | 0.025227027 |
| NUDT3 | 578.3822876 | -0.326868579 | 0.128536242 | 0.00622244 | 0.024079467 |
| C2orf49-DT | 550.7581484 | -0.326878976 | 0.126527866 | 0.00555696 | 0.021857929 |
| ZBTB7A | 1100.082819 | -0.326903436 | 0.123919452 | 0.00475767 | 0.019297186 |
| GNPTAB | 1517.860954 | -0.327053876 | 0.122450384 | 0.00432425 | 0.017831155 |
| ARRDC1-AS1 | 716.2859675 | -0.327068254 | 0.116167885 | 0.00281198 | 0.012536367 |
| RBBP7 | 5090.144439 | -0.327127914 | 0.114149764 | 0.00240869 | 0.010994977 |
| CEP15 | 1188.753322 | -0.327586714 | 0.140189161 | 0.01075789 | 0.037726011 |
| ELF3-AS1 | 229.7162975 | -0.327627332 | 0.146543251 | 0.01385636 | 0.04606986 |
| TRIM14 | 725.1164859 | -0.327949104 | 0.108362601 | 0.00144157 | 0.007146678 |
| ITGB1P1 | 374.332478 | -0.328091929 | 0.126691376 | 0.00543423 | 0.021490284 |
| PLXNB2 | 8723.028119 | -0.328167571 | 0.133611418 | 0.00784703 | 0.029085511 |
| PLCE1 | 489.7033363 | -0.328418983 | 0.133638268 | 0.0078099 | 0.02900015 |
| TUBB2A | 4552.093695 | -0.328520162 | 0.132924392 | 0.00751869 | 0.028081056 |
| BORCS8 | 423.3733616 | -0.328602497 | 0.129736645 | 0.0063563 | 0.024505222 |
| TMEM219 | 860.1998374 | -0.328614972 | 0.097757098 | 0.00045715 | 0.0027115 |
| CNNM3 | 1885.550956 | -0.32865034 | 0.125648495 | 0.00503803 | 0.020224257 |
| PTGES3 | 18949.10547 | -0.328676253 | 0.116122609 | 0.00267033 | 0.011984415 |
| TMEM121B | 701.5132951 | -0.328943207 | 0.145568678 | 0.01298694 | 0.043788073 |
| CBX3 | 15249.08962 | -0.32899266 | 0.130957822 | 0.00671736 | 0.025581128 |
| PLOD2 | 779.8740385 | -0.329024146 | 0.126428305 | 0.0052225 | 0.020813086 |
| TRIM71 | 458.8181834 | -0.329161061 | 0.122009647 | 0.00396544 | 0.016611388 |
| FHOD1 | 755.9357161 | -0.32916468 | 0.146309281 | 0.01329673 | 0.0446552 |
| GNA12 | 2648.874891 | -0.32921384 | 0.135134813 | 0.00824179 | 0.030319153 |
| NID1 | 1117.973103 | -0.329793819 | 0.141436648 | 0.0107963 | 0.037821995 |
| NOP56 | 13017.78123 | -0.330119735 | 0.108231845 | 0.00132359 | 0.00667442 |
| PSD | 1791.484394 | -0.330140069 | 0.115647337 | 0.00246461 | 0.011205675 |
| NAA38 | 750.3538889 | -0.330429393 | 0.105237138 | 0.00098127 | 0.005179391 |
| DNAAF10 | 440.1813284 | -0.330735348 | 0.118710867 | 0.00303209 | 0.013331602 |
| ATCAY | 4464.461148 | -0.330884142 | 0.132948181 | 0.00710552 | 0.02678606 |
| FUT11 | 649.9404574 | -0.3309519 | 0.106931514 | 0.00113795 | 0.005876629 |
| CCDC120 | 715.0150246 | -0.331114928 | 0.114795827 | 0.00223983 | 0.010387881 |
| GSTZ1 | 494.7399046 | -0.331376791 | 0.118107295 | 0.00284989 | 0.012654942 |
| ZNF71 | 1547.480046 | -0.331633309 | 0.109852146 | 0.00145733 | 0.007212626 |
| RPL6P10 | 597.3882532 | -0.331781944 | 0.143668787 | 0.01133197 | 0.039336696 |
| MGA | 4253.30603 | -0.331974437 | 0.128044873 | 0.00530715 | 0.02108507 |
| ABCB6 | 534.3248581 | -0.332038572 | 0.110553653 | 0.00153001 | 0.007510903 |
| EIF3B | 11670.21896 | -0.332129382 | 0.116238354 | 0.0024266 | 0.011068544 |

| | | | | | |
|---|---|---|---|---|---|
| COCH | 1017.376635 | -0.3321484 | 0.111163004 | 0.00160762 | 0.007835767 |
| TIAM1 | 5073.999409 | -0.332187273 | 0.118774972 | 0.00291918 | 0.012904113 |
| PRRT2 | 1186.913609 | -0.332267629 | 0.138785665 | 0.00909446 | 0.032931584 |
| CDC7 | 5495.485194 | -0.332397793 | 0.139619491 | 0.00941199 | 0.033825264 |
| TNRC18 | 5494.70875 | -0.332414905 | 0.149964139 | 0.01421699 | 0.047048294 |
| NUBP2 | 2683.542267 | -0.332430234 | 0.105376143 | 0.00092659 | 0.004938929 |
| PRRC2A | 20230.85512 | -0.332786108 | 0.144427227 | 0.01143154 | 0.039615467 |
| ARNT2 | 4624.642099 | -0.332897608 | 0.132086466 | 0.00646703 | 0.024843653 |
| RALBP1 | 2872.422482 | -0.333191113 | 0.099907778 | 0.00049452 | 0.002900556 |
| ADAT2 | 1125.608161 | -0.333224439 | 0.138574982 | 0.00881121 | 0.032103771 |
| CSNK1E | 5167.231515 | -0.333233909 | 0.105295203 | 0.00089296 | 0.004777002 |
| THAP7 | 971.0147177 | -0.333268662 | 0.095606329 | 0.00028596 | 0.001815519 |
| TWNK | 2148.281845 | -0.333275711 | 0.147593971 | 0.01279799 | 0.043328604 |
| SLC1A2 | 2933.703459 | -0.333394798 | 0.119411639 | 0.00294812 | 0.013012461 |
| DBI | 1583.364482 | -0.333471288 | 0.126769538 | 0.00473743 | 0.019218876 |
| ZNF143 | 790.4825677 | -0.333700217 | 0.143058579 | 0.01059515 | 0.037276089 |
| DNA2 | 2912.925621 | -0.333831071 | 0.140549527 | 0.00949124 | 0.034056524 |
| SRSF1 | 12096.33923 | -0.334031181 | 0.123697542 | 0.00386188 | 0.016234794 |
| PPP1R3E | 508.923271 | -0.334132199 | 0.152463193 | 0.01498731 | 0.049154547 |
| GGH | 1337.145336 | -0.334142056 | 0.124551618 | 0.00406405 | 0.016941651 |
| CHTF18 | 4924.322348 | -0.334193289 | 0.116700413 | 0.00236065 | 0.010818899 |
| RABL6 | 7107.703287 | -0.334406128 | 0.134242824 | 0.00696151 | 0.02633981 |
| ALDH7A1 | 5895.809044 | -0.334477001 | 0.117403532 | 0.00246772 | 0.011214621 |
| FRAT2 | 345.3461592 | -0.334514817 | 0.148132676 | 0.01272383 | 0.043113019 |
| EEF1A1P5 | 12877.17616 | -0.334552867 | 0.147191507 | 0.01226718 | 0.041890123 |
| MPPED2 | 714.4684496 | -0.334882626 | 0.103367535 | 0.00068658 | 0.003830385 |
| NOP14 | 2787.161075 | -0.335042058 | 0.119005861 | 0.00272908 | 0.012210812 |
| DERA | 1277.412197 | -0.335096428 | 0.151803387 | 0.01436414 | 0.047455859 |
| SH3PXD2A | 6866.434302 | -0.335266905 | 0.121251548 | 0.00317357 | 0.01380897 |
| GGCT | 1274.276675 | -0.335369177 | 0.128535331 | 0.00499647 | 0.020080714 |
| MT-ND4 | 198506.5569 | -0.335441752 | 0.143424372 | 0.0103513 | 0.03659344 |
| GALNT13 | 1843.69198 | -0.335584668 | 0.111145735 | 0.00143408 | 0.007116422 |
| RPS15 | 15315.03064 | -0.335626225 | 0.122530819 | 0.0034237 | 0.014692188 |
| NIT2 | 1206.647185 | -0.335710574 | 0.141427166 | 0.00945108 | 0.03395388 |
| LZIC | 1251.037856 | -0.335821358 | 0.104834713 | 0.00077557 | 0.004239432 |
| DCAF4 | 1201.901825 | -0.335840429 | 0.109369828 | 0.00121117 | 0.006198721 |
| RABIF | 794.7074578 | -0.335852024 | 0.112305048 | 0.00157228 | 0.007694495 |
| CALM2 | 20646.92066 | -0.335986527 | 0.15062168 | 0.01352411 | 0.045235821 |
| RPL18AP3 | 2864.25083 | -0.336320651 | 0.096001636 | 0.00026516 | 0.001707165 |
| PRELID1P6 | 228.416284 | -0.336568257 | 0.136228831 | 0.00729123 | 0.027355778 |
| MRPL33 | 741.8809758 | -0.336825654 | 0.13612651 | 0.00721024 | 0.027126125 |
| OTUB1 | 3446.664023 | -0.336933796 | 0.095254024 | 0.00023308 | 0.001528426 |
| ARGLU1 | 10718.23626 | -0.336934304 | 0.134442775 | 0.00661169 | 0.025267213 |
| TFDP2 | 3557.18399 | -0.337147138 | 0.091874241 | 0.00014051 | 0.000996692 |
| GALNT18 | 445.6320346 | -0.337227675 | 0.151143942 | 0.01343263 | 0.045002071 |
| CLN8-AS1 | 183.5325094 | -0.337257601 | 0.152114719 | 0.01389623 | 0.04618004 |
| FUNDC2 | 2056.317799 | -0.337546467 | 0.154167958 | 0.01483139 | 0.048744221 |
| ATP5MG | 816.8819583 | -0.337702101 | 0.145252735 | 0.01061859 | 0.037332988 |
| TMC6 | 315.687801 | -0.337721143 | 0.142116651 | 0.0093062 | 0.03352711 |
| SNRNP25 | 1303.046104 | -0.337745172 | 0.110906524 | 0.00130697 | 0.006599768 |

| | | | | | |
|---|---|---|---|---|---|
| ENSA | 7649.243877 | -0.337767678 | 0.101772732 | 0.00051507 | 0.002999179 |
| PRKCSH | 5396.467876 | -0.337794408 | 0.148325076 | 0.01195918 | 0.04104968 |
| SOX4 | 8703.850922 | -0.33787062 | 0.11119894 | 0.00133604 | 0.006727315 |
| PDE3A | 1231.941446 | -0.338199399 | 0.131022349 | 0.00534385 | 0.021194015 |
| SLC4A3 | 760.2801318 | -0.338332721 | 0.117794185 | 0.00226258 | 0.010465069 |
| RPL24 | 13445.70585 | -0.338405703 | 0.148196625 | 0.01174724 | 0.040464174 |
| RPL17P7 | 521.0382187 | -0.338497289 | 0.148971055 | 0.01207532 | 0.041358562 |
| COA7 | 5612.943315 | -0.338613949 | 0.104660326 | 0.00068754 | 0.003832519 |
| ENY2 | 3977.427163 | -0.338633815 | 0.133892814 | 0.00616453 | 0.023868834 |
| TMUB1 | 782.1157507 | -0.338700455 | 0.132249348 | 0.00564176 | 0.022153348 |
| RABEPK | 2262.138908 | -0.338721441 | 0.108577508 | 0.00101853 | 0.005340383 |
| AMOT | 717.5319254 | -0.338747151 | 0.142900233 | 0.00940629 | 0.033810671 |
| FLAD1 | 3618.532263 | -0.338882271 | 0.081186755 | 1.74E-05 | 0.000166197 |
| DBN1 | 12362.4481 | -0.339069766 | 0.123523705 | 0.00331907 | 0.014320891 |
| SCARF2 | 1516.070591 | -0.33907672 | 0.152209103 | 0.01343361 | 0.045002071 |
| C4orf50 | 390.4736627 | -0.339106996 | 0.131826073 | 0.00545747 | 0.021569753 |
| ZNF891 | 2172.669708 | -0.339160571 | 0.110549885 | 0.00120699 | 0.006181474 |
| XRCC5 | 25875.40444 | -0.33917612 | 0.136621556 | 0.00698241 | 0.026399402 |
| HCG18 | 3999.291049 | -0.339215608 | 0.095743223 | 0.00022624 | 0.001487843 |
| GSTA4 | 2914.100627 | -0.339221674 | 0.128040453 | 0.00438628 | 0.018032666 |
| ZDHHC16 | 1468.068579 | -0.339245474 | 0.086847027 | 5.42E-05 | 0.000436449 |
| WLS | 2572.579022 | -0.339295683 | 0.127923029 | 0.00434718 | 0.017911318 |
| ELOVL2 | 3223.467607 | -0.339325773 | 0.123011765 | 0.00318505 | 0.013841369 |
| WDR46 | 2653.154155 | -0.339567776 | 0.100779022 | 0.00042731 | 0.002564839 |
| CELF5 | 945.1371712 | -0.339703938 | 0.129146305 | 0.00462173 | 0.01884242 |
| GCA | 656.1083587 | -0.339814892 | 0.149593829 | 0.0120237 | 0.041229707 |
| KAT8 | 1957.929151 | -0.339964756 | 0.089016827 | 7.71E-05 | 0.000590484 |
| CEP85 | 2486.07618 | -0.339969097 | 0.108742895 | 0.00099119 | 0.005221768 |
| DLEU2 | 854.2741327 | -0.339986463 | 0.146203989 | 0.01049919 | 0.036995441 |
| GIT1 | 2684.074002 | -0.340140896 | 0.147318792 | 0.01093954 | 0.038187183 |
| STARD7 | 10249.60188 | -0.340189337 | 0.084984452 | 3.61E-05 | 0.000309863 |
| MMP24OS | 284.028897 | -0.340498197 | 0.153061966 | 0.01344843 | 0.045033964 |
| PHACTR4 | 3710.660892 | -0.340519829 | 0.102567735 | 0.00050789 | 0.002967107 |
| SPAG7 | 1144.075089 | -0.34055084 | 0.107025142 | 0.00081974 | 0.004433772 |
| SAMD14 | 1483.25476 | -0.340644806 | 0.124793969 | 0.00345126 | 0.014795005 |
| SNX12 | 1421.832252 | -0.340717136 | 0.101644665 | 0.00045282 | 0.00269049 |
| CLCN4 | 1164.872247 | -0.340724922 | 0.1038426 | 0.00058172 | 0.003321038 |
| NOP10 | 1717.540528 | -0.340990502 | 0.146410059 | 0.01035808 | 0.036605407 |
| CELF2 | 1647.767946 | -0.34116209 | 0.120286813 | 0.00249985 | 0.011338111 |
| EXOSC5 | 1209.111206 | -0.341239963 | 0.114596458 | 0.00160467 | 0.007825114 |
| GRM2 | 831.8395034 | -0.341263673 | 0.114322589 | 0.00156722 | 0.007671595 |
| AMER3 | 2423.098935 | -0.341405818 | 0.130873054 | 0.00488133 | 0.019713215 |
| TPM2 | 373.2473853 | -0.341414973 | 0.144138443 | 0.00934194 | 0.033611108 |
| ATP9A | 5322.995842 | -0.341420231 | 0.108928573 | 0.00095971 | 0.005083813 |
| NFE2L3 | 602.2761361 | -0.34159839 | 0.117652725 | 0.00202712 | 0.009556554 |
| THBS3 | 1976.359735 | -0.341641186 | 0.102321424 | 0.0004729 | 0.002791276 |
| MFSD12 | 1565.306336 | -0.341690538 | 0.130448675 | 0.00473034 | 0.019203628 |
| RHOF | 152.544911 | -0.341793436 | 0.148079611 | 0.01087835 | 0.03803169 |
| TMEM14B | 2663.075761 | -0.341878861 | 0.139237303 | 0.007426 | 0.027775253 |
| TMEFF1 | 744.5236813 | -0.341885513 | 0.126907219 | 0.00381296 | 0.016087345 |

| | | | | | |
|---|---|---|---|---|---|
| URI1 | 4041.47263 | -0.342271296 | 0.112446868 | 0.00129076 | 0.006532899 |
| GRAMD4 | 1662.415263 | -0.342410193 | 0.110027227 | 0.00103012 | 0.005391568 |
| HNRNPU | 43607.12971 | -0.342584267 | 0.095219134 | 0.00018161 | 0.001240129 |
| MFSD5 | 702.7270915 | -0.342657863 | 0.111950343 | 0.00121925 | 0.006229241 |
| ZNF232 | 1328.80516 | -0.342712606 | 0.132039424 | 0.00503836 | 0.020224257 |
| DCLK1 | 2223.736967 | -0.342730548 | 0.109441758 | 0.0009637 | 0.005101039 |
| EEF1AKMT1 | 187.1675631 | -0.342790594 | 0.150681595 | 0.01176517 | 0.040506812 |
| ZC3HAV1L | 648.4805534 | -0.342908164 | 0.122297001 | 0.00273927 | 0.012248373 |
| PKN3 | 1229.05466 | -0.342921379 | 0.100704188 | 0.00037098 | 0.00227452 |
| NDUFB4 | 1910.254974 | -0.342924567 | 0.132685695 | 0.0051925 | 0.020705565 |
| TRDMT1 | 440.0661741 | -0.343066669 | 0.156044825 | 0.01414944 | 0.046900231 |
| LETM1 | 2631.880523 | -0.343152647 | 0.136831928 | 0.00641117 | 0.024661284 |
| NDUFV1 | 6253.598062 | -0.343179184 | 0.108584844 | 0.00087303 | 0.004685026 |
| ULK1 | 4921.846701 | -0.343324799 | 0.137233299 | 0.00651328 | 0.024976023 |
| NUCKS1 | 41767.20589 | -0.343566208 | 0.132530278 | 0.00506538 | 0.020312877 |
| GNB1 | 12664.38371 | -0.343617799 | 0.129154718 | 0.00417154 | 0.017309018 |
| SDHAF2 | 325.5495612 | -0.343624159 | 0.149053592 | 0.01086514 | 0.038004877 |
| UBE2G2 | 7532.69337 | -0.343644761 | 0.093464823 | 0.00013357 | 0.000954051 |
| CDK2AP1 | 7021.354454 | -0.343854731 | 0.11457992 | 0.00147416 | 0.007281909 |
| ZNF573 | 267.8830536 | -0.34398437 | 0.150779654 | 0.01151625 | 0.03982178 |
| CD99L2 | 1214.628599 | -0.344165967 | 0.110817014 | 0.00104503 | 0.005459834 |
| MAP2K7 | 2361.948452 | -0.34423815 | 0.118101016 | 0.0019362 | 0.009210279 |
| RRM1 | 9583.433907 | -0.34445181 | 0.146321089 | 0.00956858 | 0.034268245 |
| UBAP2L | 11448.12065 | -0.344636501 | 0.123002543 | 0.00273703 | 0.012241044 |
| TAGLN2 | 269.2785458 | -0.344762124 | 0.134840401 | 0.00556557 | 0.021875085 |
| PRPF31 | 1627.209661 | -0.344926127 | 0.120934195 | 0.00234513 | 0.010759749 |
| ITGB1BP1 | 1602.722147 | -0.344934277 | 0.107358277 | 0.00072534 | 0.004006411 |
| PEMT | 526.3666937 | -0.34494104 | 0.123915112 | 0.00288758 | 0.012794685 |
| HNRNPAB | 12955.25457 | -0.345200365 | 0.110669621 | 0.00099514 | 0.005236445 |
| CCDC85C | 2213.037078 | -0.345232244 | 0.138988241 | 0.00677996 | 0.025766601 |
| USB1 | 1823.728674 | -0.345253727 | 0.120954559 | 0.00232606 | 0.01069372 |
| GATA3 | 8688.925063 | -0.345339539 | 0.14006344 | 0.00711828 | 0.026816371 |
| SLC25A22 | 1826.576149 | -0.345735844 | 0.137899676 | 0.00635224 | 0.024494178 |
| SLC27A4 | 1045.334707 | -0.345780688 | 0.149849732 | 0.01070156 | 0.037566878 |
| LYRM7 | 1751.597817 | -0.345863937 | 0.125874245 | 0.00320311 | 0.013884628 |
| PPP4R2P4 | 135.0948959 | -0.346104218 | 0.151835119 | 0.0114585 | 0.039675473 |
| CDK4 | 6736.801998 | -0.3461504 | 0.096079177 | 0.00017594 | 0.001208042 |
| EBPL | 1342.042954 | -0.346159904 | 0.111797857 | 0.00106992 | 0.005568681 |
| CNBP | 16356.03081 | -0.346426585 | 0.134682457 | 0.0052953 | 0.021050173 |
| NFIB | 9257.730729 | -0.346524506 | 0.110967358 | 0.00097834 | 0.00516583 |
| SMUG1 | 1085.775929 | -0.346693593 | 0.121071395 | 0.00224844 | 0.010418417 |
| CACFD1 | 576.8358738 | -0.346722971 | 0.134898766 | 0.00531707 | 0.02110815 |
| SELENOH | 291.0491514 | -0.346727093 | 0.1282483 | 0.00363463 | 0.015449045 |
| NOL3 | 1154.866838 | -0.346869546 | 0.112550113 | 0.00111899 | 0.005800571 |
| CLSPN | 3226.858467 | -0.346927234 | 0.119846982 | 0.00203927 | 0.00960062 |
| SLC4A8 | 7163.975783 | -0.347059908 | 0.097061696 | 0.00019437 | 0.001308767 |
| UTP25 | 3440.396887 | -0.347060946 | 0.123245207 | 0.00259647 | 0.011696362 |
| ZNF559 | 1482.123092 | -0.347223527 | 0.156294966 | 0.01312881 | 0.044172274 |
| ARPC4 | 971.6486886 | -0.347345885 | 0.131568151 | 0.00435537 | 0.017937885 |
| GBA1 | 697.756097 | -0.347605713 | 0.101647459 | 0.0003459 | 0.002138611 |

| | | | | | |
|---|---|---|---|---|---|
| NUDT13 | 203.6594155 | -0.347690703 | 0.151633454 | 0.01100037 | 0.038359516 |
| AGRN | 2123.088958 | -0.347722773 | 0.159332032 | 0.01438165 | 0.047493854 |
| ZNF335 | 920.4349608 | -0.348037145 | 0.146444642 | 0.00888695 | 0.032333905 |
| ISOC1 | 567.2575541 | -0.348109912 | 0.123458395 | 0.002557 | 0.011556532 |
| ATP5ME | 336.7190701 | -0.348332312 | 0.146945016 | 0.0090152 | 0.032690616 |
| PPCDC | 265.889491 | -0.348380901 | 0.12463284 | 0.00274985 | 0.012290332 |
| CACNA1H | 4642.663848 | -0.348465408 | 0.161733557 | 0.01529607 | 0.049975898 |
| ROR1 | 569.5416625 | -0.348692096 | 0.15699874 | 0.0130567 | 0.043980006 |
| SATB1 | 14178.67484 | -0.348771043 | 0.1072991 | 0.00062799 | 0.003546806 |
| RPL18A | 26607.33807 | -0.348939443 | 0.100784287 | 0.00029465 | 0.00186033 |
| ALKBH7 | 460.847095 | -0.348942987 | 0.135252425 | 0.00512619 | 0.020484827 |
| YDJC | 1020.383674 | -0.34930749 | 0.139503692 | 0.00630915 | 0.024359988 |
| SMPD4 | 6721.233466 | -0.349367119 | 0.130284695 | 0.00383218 | 0.016141957 |
| BAHCC1 | 5549.393293 | -0.349886904 | 0.144330126 | 0.00778555 | 0.02892538 |
| SHANK3 | 1908.402085 | -0.349998013 | 0.143444606 | 0.00746547 | 0.027902552 |
| ANP32E | 14287.18498 | -0.350228268 | 0.142103814 | 0.00698516 | 0.026404936 |
| IMPDH1 | 1879.741684 | -0.350472703 | 0.150402713 | 0.00989164 | 0.035186227 |
| INO80C | 679.5041249 | -0.350588443 | 0.142173148 | 0.0069492 | 0.026302838 |
| SLC39A3 | 862.7921423 | -0.350683711 | 0.139239137 | 0.00602626 | 0.02342497 |
| NDUFB6 | 1243.167509 | -0.350751812 | 0.148040893 | 0.00894392 | 0.03248943 |
| HNRNPA2B1 | 57947.85563 | -0.350759985 | 0.120531952 | 0.00191398 | 0.009123563 |
| NDC1 | 7305.56781 | -0.35091065 | 0.136027083 | 0.00508595 | 0.020383431 |
| ZNF271P | 1706.99833 | -0.351221644 | 0.135386013 | 0.00487699 | 0.019699568 |
| DIMT1 | 838.0823696 | -0.351324443 | 0.117050834 | 0.00142902 | 0.007094721 |
| USP22 | 13779.07735 | -0.351366273 | 0.149327547 | 0.00929828 | 0.033504426 |
| SNRPG | 1941.250091 | -0.351395207 | 0.138681411 | 0.00576193 | 0.022569163 |
| HNRNPL | 10209.2896 | -0.351406001 | 0.123436592 | 0.00232172 | 0.010678755 |
| RABEP2 | 301.2912017 | -0.351418095 | 0.151559362 | 0.01013866 | 0.035915825 |
| GOT2 | 5612.917571 | -0.351556767 | 0.121300442 | 0.00197975 | 0.009369774 |
| FGFBP3 | 275.267915 | -0.351583166 | 0.134187196 | 0.00452696 | 0.018507432 |
| CDKN2AIPNL | 1350.215041 | -0.35182457 | 0.119188798 | 0.00167113 | 0.00810498 |
| TGFBRAP1 | 1966.112291 | -0.351898557 | 0.099258562 | 0.00021406 | 0.001419952 |
| PTTG1 | 2984.321281 | -0.352132903 | 0.142254048 | 0.00672901 | 0.025620311 |
| UBE2L3 | 3553.669029 | -0.352514355 | 0.111219906 | 0.0008166 | 0.00441908 |
| SUMO2 | 18775.99664 | -0.352530955 | 0.146212424 | 0.00796177 | 0.029443978 |
| SNAPIN | 1015.413257 | -0.352642137 | 0.128454351 | 0.00313634 | 0.01369366 |
| RPA1 | 6093.659061 | -0.352745845 | 0.121903943 | 0.00199848 | 0.009438852 |
| XYLT2 | 1333.088592 | -0.352780493 | 0.089400369 | 4.38E-05 | 0.00036625 |
| TRIM69 | 874.979139 | -0.352914053 | 0.117562476 | 0.00141778 | 0.007054266 |
| C12orf75 | 1578.027833 | -0.352937304 | 0.125812284 | 0.00261806 | 0.011775555 |
| GMNN | 3516.357017 | -0.352942725 | 0.154054953 | 0.01078513 | 0.037795728 |
| GNB4 | 2117.04236 | -0.353051069 | 0.120545954 | 0.00178833 | 0.00858029 |
| TNFAIP8L1 | 321.6112532 | -0.353302366 | 0.132988871 | 0.00404915 | 0.016886376 |
| SMIM27 | 141.8406685 | -0.353518488 | 0.146369282 | 0.00784018 | 0.029075825 |
| MYL6 | 6556.961954 | -0.354148097 | 0.132466557 | 0.00384343 | 0.01617607 |
| ST3GAL3 | 169.5160288 | -0.354304692 | 0.159123697 | 0.01253823 | 0.042645745 |
| CDC25A | 3361.741067 | -0.354719159 | 0.105233873 | 0.00040162 | 0.00242911 |
| MFGE8 | 1096.258397 | -0.355312649 | 0.095138498 | 0.00010199 | 0.000751715 |
| SIX3 | 2165.534189 | -0.355428132 | 0.135791877 | 0.0044852 | 0.018373249 |
| CRACD | 6018.450108 | -0.355650101 | 0.119693435 | 0.00154628 | 0.00757991 |

| | | | | | |
|---|---|---|---|---|---|
| CNPY3 | 1592.965297 | -0.355757953 | 0.102367787 | 0.00027363 | 0.001749636 |
| CCNA2 | 5431.546019 | -0.355783228 | 0.12128952 | 0.00174306 | 0.008394423 |
| FAM86B1 | 129.1894737 | -0.356231467 | 0.144631882 | 0.00682682 | 0.025925529 |
| PKM | 54084.53091 | -0.356541264 | 0.148073842 | 0.00788186 | 0.029188305 |
| KIFC1 | 5365.895481 | -0.35657179 | 0.103100713 | 0.00029015 | 0.001838163 |
| SERBP1P5 | 351.7104762 | -0.356592875 | 0.151220725 | 0.00895677 | 0.032507428 |
| DRD2 | 1073.33223 | -0.356650221 | 0.1414867 | 0.00583346 | 0.022771324 |
| C12orf43 | 1061.416605 | -0.356984518 | 0.123404113 | 0.00196866 | 0.009329595 |
| HSPD1P1 | 388.3661989 | -0.357067793 | 0.160957012 | 0.01261434 | 0.042841043 |
| TXN | 4945.428085 | -0.357075732 | 0.158130903 | 0.01146271 | 0.039683347 |
| NTN4 | 269.0831806 | -0.357130696 | 0.161936328 | 0.01300753 | 0.043843128 |
| TMCO6 | 805.8930159 | -0.357195181 | 0.106622837 | 0.00042825 | 0.002569003 |
| CPOX | 909.2235118 | -0.357243246 | 0.091094326 | 4.77E-05 | 0.000393334 |
| EEF1AKMT4 | 129.7244959 | -0.357332242 | 0.16133802 | 0.01270646 | 0.043061238 |
| DIAPH1 | 5553.29955 | -0.357385127 | 0.123484392 | 0.00195704 | 0.009283659 |
| GABPB1-AS1 | 13102.42165 | -0.357400812 | 0.123516419 | 0.00196071 | 0.009296755 |
| MTARC1 | 1904.500039 | -0.357593329 | 0.093358206 | 6.91E-05 | 0.000538447 |
| KMT5C | 1920.503303 | -0.35766191 | 0.12462022 | 0.0021065 | 0.009876416 |
| ARPC1A | 705.1255483 | -0.357795915 | 0.11931919 | 0.00140448 | 0.007001384 |
| TMEM18 | 1542.918196 | -0.357811126 | 0.117194124 | 0.00117713 | 0.006053144 |
| FYCO1 | 527.0696552 | -0.357880211 | 0.139394574 | 0.00510355 | 0.020418186 |
| CADM4 | 915.1176672 | -0.357909925 | 0.117980173 | 0.00125381 | 0.00637762 |
| POLRMT | 2725.114293 | -0.358037752 | 0.123358294 | 0.00190224 | 0.009076038 |
| PIP5K1C | 2633.888439 | -0.358169951 | 0.135174633 | 0.00404361 | 0.016872777 |
| CKAP4 | 6457.045406 | -0.358231136 | 0.121157368 | 0.00160248 | 0.007816248 |
| UBE2I | 5271.107264 | -0.35826274 | 0.10135913 | 0.00021744 | 0.001439613 |
| H2AZ1 | 16517.13194 | -0.358366476 | 0.147165828 | 0.00727754 | 0.027314337 |
| SUSD5 | 1964.486522 | -0.358415769 | 0.111544844 | 0.00068741 | 0.003832519 |
| MRTO4 | 1618.33885 | -0.358608093 | 0.111083279 | 0.00065223 | 0.003663609 |
| ARF5 | 2206.189412 | -0.358644816 | 0.098748798 | 0.00015017 | 0.001055413 |
| FANCD2 | 4028.004872 | -0.358761259 | 0.127989815 | 0.00257142 | 0.011608923 |
| HPRT1 | 1638.388803 | -0.358926178 | 0.129671014 | 0.00285421 | 0.012668664 |
| MAPRE1 | 9399.679158 | -0.359045713 | 0.127915126 | 0.00253856 | 0.011480741 |
| PRR36 | 1062.038102 | -0.359144621 | 0.164711743 | 0.01365581 | 0.045594752 |
| ZNF514 | 1745.312475 | -0.359666532 | 0.103543646 | 0.00027121 | 0.001736607 |
| ASTN1 | 954.1825415 | -0.359706321 | 0.122980599 | 0.00176076 | 0.008467785 |
| NOLC1 | 14458.91783 | -0.359798479 | 0.110971212 | 0.00061857 | 0.003498582 |
| GRIN3A | 437.4395329 | -0.359825084 | 0.161204872 | 0.01204149 | 0.041263318 |
| DAXX | 3605.714336 | -0.359876736 | 0.110905308 | 0.00061278 | 0.003472345 |
| ADI1 | 2392.642425 | -0.359962795 | 0.117052156 | 0.00108532 | 0.00563598 |
| PCOLCE | 3362.500008 | -0.359980195 | 0.139795235 | 0.00494914 | 0.019940082 |
| TLE5 | 4254.239146 | -0.360103961 | 0.116453607 | 0.00102553 | 0.005370273 |
| PWWP2B | 928.654649 | -0.360206294 | 0.143927355 | 0.00602711 | 0.02342497 |
| KRBA2 | 930.0078811 | -0.360226513 | 0.106846247 | 0.00039207 | 0.0023813 |
| SNUPN | 546.7717566 | -0.360257901 | 0.12654364 | 0.00223652 | 0.010374853 |
| LZTR1 | 654.6472953 | -0.360321454 | 0.128637235 | 0.00256947 | 0.01160266 |
| MRPS7 | 4706.107724 | -0.360510169 | 0.110039774 | 0.00054834 | 0.003161197 |
| ECI1 | 1465.633042 | -0.360563642 | 0.113636436 | 0.00078151 | 0.004263996 |
| CTBP1 | 4236.19762 | -0.360743174 | 0.116346553 | 0.00099501 | 0.005236445 |
| EVL | 8756.102693 | -0.360825359 | 0.106287983 | 0.00035968 | 0.002214528 |

| | | | | | |
|---|---|---|---|---|---|
| SELENBP1 | 297.5263112 | -0.361355458 | 0.166325881 | 0.01375287 | 0.045836958 |
| QTRT1 | 2987.40393 | -0.361417629 | 0.081492029 | 4.98E-06 | 5.46E-05 |
| PRR7 | 447.6409586 | -0.361631597 | 0.138659049 | 0.00447971 | 0.018354421 |
| TMEM98 | 2893.64249 | -0.361790493 | 0.117724189 | 0.00108435 | 0.005632352 |
| NALF2 | 482.8775056 | -0.361846253 | 0.165997303 | 0.01348828 | 0.045137986 |
| GLIPR2 | 756.5017258 | -0.361898093 | 0.114338866 | 0.00079808 | 0.00433368 |
| NDUFS6 | 1157.671261 | -0.36190324 | 0.123726492 | 0.00174378 | 0.008395929 |
| FGD5-AS1 | 8297.856397 | -0.361927286 | 0.094148954 | 6.42E-05 | 0.000505625 |
| TMA7 | 1085.516327 | -0.362059649 | 0.140032506 | 0.00476174 | 0.019306092 |
| BAP1 | 5076.494754 | -0.362077922 | 0.103264951 | 0.00023804 | 0.001556428 |
| PLEKHG3 | 768.4469108 | -0.362439168 | 0.116923971 | 0.0009907 | 0.005221095 |
| PAXX | 1222.088429 | -0.362441228 | 0.104892133 | 0.00028679 | 0.001820238 |
| CYB5RL | 1319.785984 | -0.362470309 | 0.112982816 | 0.00068789 | 0.003832519 |
| GLB1L2 | 1121.307155 | -0.362546385 | 0.132705212 | 0.00312506 | 0.013665243 |
| SF3A2 | 4900.706534 | -0.362975291 | 0.126830601 | 0.00211135 | 0.009892583 |
| AGPAT5 | 5281.048382 | -0.363006574 | 0.130294312 | 0.00265669 | 0.011931004 |
| PCGF3 | 3856.201753 | -0.363069537 | 0.088307506 | 2.10E-05 | 0.000194529 |
| PBXIP1 | 410.3914774 | -0.363282839 | 0.1343976 | 0.00338953 | 0.01456377 |
| RPS10 | 110.9809378 | -0.363304307 | 0.16813986 | 0.01399405 | 0.046475075 |
| ENTR1 | 3304.979924 | -0.363311232 | 0.089185296 | 2.46E-05 | 0.00022364 |
| AK4 | 1917.319441 | -0.36351721 | 0.157906833 | 0.00996883 | 0.035417928 |
| MED18 | 486.8701925 | -0.363536337 | 0.109021458 | 0.00044125 | 0.00263391 |
| TFDP1 | 7395.288774 | -0.363625586 | 0.094750647 | 6.55E-05 | 0.000514497 |
| SIPA1L3 | 928.7358278 | -0.363667801 | 0.156015095 | 0.00927037 | 0.033439026 |
| PRPF8 | 24292.05585 | -0.363740754 | 0.159375974 | 0.01045562 | 0.036879339 |
| MYCN | 74542.638 | -0.363841382 | 0.127814574 | 0.00220402 | 0.01024955 |
| FPGS | 1978.219781 | -0.364001883 | 0.116287571 | 0.00088967 | 0.004763104 |
| TOP2A | 31098.71646 | -0.364096047 | 0.153480453 | 0.00833391 | 0.030592393 |
| TBC1D25 | 557.1050741 | -0.364130154 | 0.138265109 | 0.00412178 | 0.017140596 |
| ALDH18A1 | 6290.119334 | -0.364201935 | 0.123845698 | 0.0016434 | 0.007989347 |
| FOXK1 | 3211.725842 | -0.364377174 | 0.128918203 | 0.00233838 | 0.010736607 |
| THTPA | 251.1355951 | -0.364467585 | 0.135677484 | 0.00353941 | 0.015122458 |
| TBC1D8 | 987.0524901 | -0.364475522 | 0.099949623 | 0.00013874 | 0.000986893 |
| ARVCF | 4457.220254 | -0.364552387 | 0.104995971 | 0.00026749 | 0.001719522 |
| RPSAP47 | 196.1747701 | -0.36462232 | 0.159514591 | 0.01031959 | 0.036487636 |
| WSB2 | 2686.374784 | -0.364628951 | 0.120075493 | 0.00120728 | 0.006181474 |
| ENTPD1-AS1 | 136.597024 | -0.364729532 | 0.170416626 | 0.01455898 | 0.047994684 |
| DGLUCY | 833.9085819 | -0.364846969 | 0.095916646 | 7.47E-05 | 0.000574582 |
| WIZ | 4740.359949 | -0.364916406 | 0.160399424 | 0.01057964 | 0.03723931 |
| PRPS2 | 1035.082616 | -0.365053357 | 0.12536707 | 0.0017926 | 0.008598799 |
| SH3BP2 | 2883.390452 | -0.365227796 | 0.102421741 | 0.00018809 | 0.001272767 |
| CARM1 | 3119.586369 | -0.365230282 | 0.144426086 | 0.0054853 | 0.021642311 |
| FAM200B | 1150.425267 | -0.365327042 | 0.15028477 | 0.00712008 | 0.026816371 |
| CCDC77 | 2284.535058 | -0.36589769 | 0.126561918 | 0.00190545 | 0.009089231 |
| MIR124-1HG | 2690.472547 | -0.366021772 | 0.142813672 | 0.00497786 | 0.020040077 |
| DPF1 | 2055.769669 | -0.366089379 | 0.10612695 | 0.00028873 | 0.001830826 |
| ARPP19 | 20249.2214 | -0.366193816 | 0.129204564 | 0.00226556 | 0.010471778 |
| COQ7 | 946.7409058 | -0.366314718 | 0.146801778 | 0.00597502 | 0.023257685 |
| ZWINT | 4875.760594 | -0.366486608 | 0.118771046 | 0.00102056 | 0.005346943 |
| RAD23A | 2270.370542 | -0.366704556 | 0.138750942 | 0.00396718 | 0.016611889 |

| | | | | | |
|---|---|---|---|---|---|
| LDLRAD3 | 2494.119239 | -0.366718362 | 0.120921147 | 0.00121207 | 0.006201793 |
| THG1L | 798.8624062 | -0.366892205 | 0.135269243 | 0.00324631 | 0.014048234 |
| NUBP1 | 255.118159 | -0.366933831 | 0.132343478 | 0.00271794 | 0.012169096 |
| MZT2B | 1507.910989 | -0.366949349 | 0.099941346 | 0.00012473 | 0.000896513 |
| ATXN7L2 | 1423.04287 | -0.367161967 | 0.096707465 | 7.62E-05 | 0.000585318 |
| APOO | 447.2467249 | -0.367182823 | 0.127577978 | 0.00197216 | 0.009340286 |
| THEM6 | 727.3027897 | -0.367421429 | 0.133584819 | 0.0028957 | 0.012827925 |
| NEIL1 | 998.0822961 | -0.367451494 | 0.134572795 | 0.00307048 | 0.013468713 |
| MMP24 | 1797.49726 | -0.367502329 | 0.166589227 | 0.01231894 | 0.042024984 |
| NCK2 | 2477.347175 | -0.367634158 | 0.129209805 | 0.00217686 | 0.010148583 |
| UBE2E3 | 4500.105806 | -0.367726477 | 0.099582294 | 0.00011459 | 0.000831783 |
| ATAD3A | 1148.322352 | -0.367886819 | 0.146332649 | 0.00563868 | 0.022145507 |
| SCD5 | 2107.190175 | -0.368032273 | 0.106839193 | 0.00029154 | 0.001845267 |
| TMEM106C | 2359.192078 | -0.36814775 | 0.137424373 | 0.00355549 | 0.015178545 |
| LYSET | 295.6517996 | -0.368197848 | 0.138793266 | 0.00383004 | 0.016136239 |
| RAB11A | 6975.606433 | -0.368263446 | 0.144525407 | 0.00513084 | 0.020491451 |
| DNAJC12 | 1134.242281 | -0.368333616 | 0.154807291 | 0.00801714 | 0.029630564 |
| GTF3C6 | 1832.503833 | -0.36836552 | 0.123778754 | 0.00144387 | 0.007156353 |
| SNHG16 | 3167.096355 | -0.368390016 | 0.122037351 | 0.0012593 | 0.006402012 |
| SNRPE | 3744.15692 | -0.3685656 | 0.151756908 | 0.0070504 | 0.026607518 |
| TOMM22 | 2485.644499 | -0.36864735 | 0.112550052 | 0.00053182 | 0.003082415 |
| DUXAP9 | 171.4939573 | -0.368764016 | 0.152742322 | 0.00731165 | 0.027417354 |
| CDCA3 | 1294.603417 | -0.368906411 | 0.080309316 | 2.30E-06 | 2.76E-05 |
| POLR3G | 329.0925161 | -0.369304906 | 0.154331042 | 0.007705 | 0.028677939 |
| MTSS1 | 1722.454231 | -0.369377135 | 0.14017254 | 0.00400543 | 0.01674481 |
| LINC01560 | 106.1050195 | -0.369401469 | 0.164545371 | 0.01111992 | 0.038711701 |
| HMGN1P38 | 87.98957892 | -0.369603822 | 0.175388982 | 0.01527458 | 0.049921559 |
| MT-ND2 | 75737.56135 | -0.369685898 | 0.128811491 | 0.00199993 | 0.00944354 |
| PLPP1 | 288.3895192 | -0.369994764 | 0.142744922 | 0.00450815 | 0.018448866 |
| COX7C | 3113.743889 | -0.370025985 | 0.154862181 | 0.00774775 | 0.028805756 |
| CRMP1 | 19170.96201 | -0.370053828 | 0.125517209 | 0.00156504 | 0.007662724 |
| LYRM4 | 1438.445354 | -0.370139877 | 0.122518311 | 0.00124005 | 0.006321369 |
| EPN2 | 1099.07465 | -0.370327551 | 0.122020887 | 0.00118478 | 0.006084842 |
| CCNE1 | 605.5348834 | -0.370452403 | 0.107945212 | 0.00030253 | 0.001903695 |
| RAPGEF1 | 4267.220608 | -0.370511891 | 0.138928401 | 0.00364002 | 0.015454609 |
| ZNF217 | 3083.89969 | -0.370552481 | 0.126599088 | 0.00166944 | 0.008100622 |
| COQ10A | 1004.578143 | -0.370782696 | 0.102680393 | 0.00015504 | 0.001084069 |
| FUS | 21784.45601 | -0.370941732 | 0.095282747 | 5.09E-05 | 0.000414541 |
| C14orf132 | 18722.75351 | -0.371081211 | 0.118285026 | 0.00084358 | 0.004544928 |
| RASSF2 | 287.7835293 | -0.371242871 | 0.142012957 | 0.0042128 | 0.017452052 |
| SPSB4 | 1378.42193 | -0.371363789 | 0.112997238 | 0.00050591 | 0.002958907 |
| APC2 | 6427.847876 | -0.371511075 | 0.155428891 | 0.00767405 | 0.028578246 |
| GTDC1 | 549.2769086 | -0.371564316 | 0.137060192 | 0.00319033 | 0.013849666 |
| C4orf46 | 4227.690717 | -0.371699422 | 0.130891846 | 0.00217391 | 0.0101393 |
| SYN2 | 2391.742816 | -0.371873875 | 0.13710015 | 0.00317221 | 0.01380897 |
| CENPN | 807.0778742 | -0.371905983 | 0.107910133 | 0.00028517 | 0.001811064 |
| ATP1A1-AS1 | 750.0934095 | -0.372134796 | 0.113393541 | 0.00051259 | 0.002989447 |
| CDCA7L | 1235.062152 | -0.372204569 | 0.123097953 | 0.00121891 | 0.006229029 |
| AARS2 | 5042.177605 | -0.372211114 | 0.107114308 | 0.00025662 | 0.001657921 |
| SACS | 7519.340768 | -0.372241349 | 0.139370939 | 0.00357035 | 0.015229339 |

| | | | | | |
|---|---|---|---|---|---|
| FUBP1 | 20726.97401 | -0.372282407 | 0.131042504 | 0.00216041 | 0.01008553 |
| RAD51D | 716.6059725 | -0.372420376 | 0.104386706 | 0.00018151 | 0.001240021 |
| MRPS21 | 3324.51442 | -0.372498168 | 0.124319533 | 0.00132916 | 0.006699224 |
| ABCC8 | 2488.247626 | -0.372558797 | 0.173804348 | 0.01385457 | 0.04606986 |
| TACC3 | 5684.693673 | -0.372668242 | 0.112881263 | 0.00047754 | 0.002815404 |
| XKR5 | 555.381882 | -0.37284902 | 0.122293841 | 0.00112038 | 0.00580349 |
| HMGB2 | 24008.76435 | -0.372849351 | 0.138773856 | 0.00340157 | 0.014606379 |
| WNK2 | 4233.775614 | -0.372875685 | 0.137280035 | 0.00312334 | 0.013665243 |
| RMI2 | 1074.362616 | -0.372880892 | 0.096040317 | 5.27E-05 | 0.000426042 |
| RBM12B | 2005.142476 | -0.37293569 | 0.12612887 | 0.00150416 | 0.007401651 |
| BBLN | 584.4229853 | -0.372985091 | 0.145358278 | 0.00477652 | 0.019346918 |
| TAFA5 | 4609.724601 | -0.372985471 | 0.123704573 | 0.00124854 | 0.006356775 |
| ADCY3 | 1773.242132 | -0.373146223 | 0.118788665 | 0.0008248 | 0.004454093 |
| SMARCC2 | 7018.529189 | -0.373287626 | 0.114630155 | 0.00055703 | 0.003200541 |
| DLGAP3 | 975.8559141 | -0.373364434 | 0.1675686 | 0.01133747 | 0.039349159 |
| ECE1 | 1380.175287 | -0.373527747 | 0.172639304 | 0.01316502 | 0.044265105 |
| PPP5C | 4086.869465 | -0.373566124 | 0.099097861 | 8.27E-05 | 0.00062799 |
| DHX33 | 5347.400254 | -0.373784976 | 0.096239091 | 5.22E-05 | 0.000422802 |
| GNPTG | 1238.595211 | -0.373799296 | 0.08885202 | 1.33E-05 | 0.000130566 |
| CTSL | 2076.654476 | -0.37387767 | 0.13908739 | 0.00337227 | 0.014504759 |
| MT-TY | 607.0502665 | -0.373984912 | 0.163833663 | 0.00991203 | 0.03524658 |
| CARHSP1 | 2441.720488 | -0.374026266 | 0.093313796 | 3.12E-05 | 0.00027347 |
| DOK3 | 876.514414 | -0.374038854 | 0.113955001 | 0.00050752 | 0.002966661 |
| SMS | 4775.842192 | -0.374045018 | 0.116803577 | 0.00066856 | 0.003745101 |
| CLPP | 1327.357816 | -0.374140986 | 0.138707105 | 0.00327924 | 0.014172817 |
| SHMT1 | 1303.776672 | -0.374161088 | 0.104568734 | 0.00017326 | 0.001191202 |
| PAGR1 | 881.2478437 | -0.374410655 | 0.118896672 | 0.00079922 | 0.004336435 |
| PSD3 | 3435.98844 | -0.374419175 | 0.118999555 | 0.00080632 | 0.004367451 |
| FHOD3 | 993.157898 | -0.374521154 | 0.131934648 | 0.00215324 | 0.010056598 |
| KBTBD11 | 1470.817515 | -0.374568874 | 0.106804706 | 0.00022569 | 0.001484688 |
| RBX1 | 2275.351079 | -0.374584857 | 0.151234106 | 0.00603006 | 0.023431988 |
| REXO2 | 662.3362476 | -0.374597602 | 0.12539723 | 0.00135535 | 0.00680286 |
| TIFA | 507.389223 | -0.374797611 | 0.128806919 | 0.00172857 | 0.008336379 |
| STAT5B | 4359.356209 | -0.375270112 | 0.090996644 | 1.90E-05 | 0.000178542 |
| SRRD | 727.142432 | -0.37527806 | 0.114835649 | 0.00053075 | 0.003078529 |
| CDT1 | 1366.275474 | -0.375364586 | 0.166174859 | 0.01042475 | 0.03678985 |
| ZNF706 | 1862.110679 | -0.375411036 | 0.105936803 | 0.00019618 | 0.001319674 |
| CRTAP | 3551.76889 | -0.375427655 | 0.10566355 | 0.00018943 | 0.001280982 |
| PRPS1 | 2947.550978 | -0.375590148 | 0.132559095 | 0.00217754 | 0.010149362 |
| HSPA8 | 66333.01421 | -0.375627119 | 0.154897216 | 0.00687275 | 0.026053641 |
| ALS2CL | 199.8244998 | -0.375630424 | 0.17260324 | 0.0126427 | 0.042894639 |
| TBC1D4 | 1252.966243 | -0.375632578 | 0.110831934 | 0.00034541 | 0.002136215 |
| RBM19 | 2358.857729 | -0.375651354 | 0.105475802 | 0.00018335 | 0.001247627 |
| CSDE1 | 46418.56015 | -0.375940182 | 0.136598447 | 0.00277121 | 0.012380457 |
| DLX5 | 375.0752087 | -0.376309108 | 0.175998053 | 0.01373819 | 0.045802891 |
| EYA1 | 5306.860267 | -0.376460379 | 0.137733202 | 0.00292218 | 0.012911857 |
| ZNF362 | 1841.968919 | -0.376552958 | 0.133205358 | 0.00221114 | 0.010275685 |
| SF3B3 | 6904.335015 | -0.376839184 | 0.110802303 | 0.00032938 | 0.002050047 |
| HNRNPUL2 | 2036.349386 | -0.37700489 | 0.094840517 | 3.54E-05 | 0.000304684 |
| GTF3A | 3636.0003 | -0.377217597 | 0.105315936 | 0.00016875 | 0.001164895 |

| | | | | | |
|---|---|---|---|---|---|
| PEX2 | 926.8258093 | -0.377348148 | 0.141303059 | 0.00348918 | 0.014948207 |
| PUS7 | 3115.570093 | -0.37746618 | 0.116223937 | 0.00056386 | 0.003234354 |
| IGFBP2 | 2874.152491 | -0.377503089 | 0.174113472 | 0.0127487 | 0.043183029 |
| HSBP1 | 3144.834945 | -0.377505414 | 0.157954498 | 0.00744973 | 0.02784878 |
| CSK | 2806.091778 | -0.378150905 | 0.166119378 | 0.00984648 | 0.035068075 |
| COX5A | 3927.63782 | -0.378653169 | 0.109908735 | 0.00027854 | 0.001773896 |
| IGF1R | 6063.240452 | -0.378677182 | 0.092215789 | 2.02E-05 | 0.000188602 |
| DRAXIN | 2454.567774 | -0.378714732 | 0.169875382 | 0.01098403 | 0.038322958 |
| PSRC1 | 2339.959692 | -0.37889446 | 0.096807223 | 4.52E-05 | 0.000375695 |
| ZBTB3 | 213.3864945 | -0.379020402 | 0.133576098 | 0.00211689 | 0.009913841 |
| DTYMK | 2992.477872 | -0.379211693 | 0.08393381 | 3.16E-06 | 3.67E-05 |
| MMP17 | 1015.975767 | -0.379330972 | 0.140477395 | 0.00317261 | 0.01380897 |
| EIF4G1 | 21813.42062 | -0.379337412 | 0.152852212 | 0.00581315 | 0.022709254 |
| HDGF | 16961.0886 | -0.379346797 | 0.109184558 | 0.00024958 | 0.001618534 |
| ARHGAP42 | 632.4689715 | -0.379593005 | 0.15993997 | 0.00768756 | 0.028623395 |
| CPSF6 | 13943.85117 | -0.379677762 | 0.115590616 | 0.00049138 | 0.002886265 |
| SNRPC | 3935.151085 | -0.379795596 | 0.112118563 | 0.00034151 | 0.002114643 |
| MFSD13A | 759.3031974 | -0.380039618 | 0.103836927 | 0.00012374 | 0.000890622 |
| PXMP4 | 880.7796368 | -0.380043165 | 0.112432059 | 0.00035012 | 0.002162741 |
| UBE2L6 | 800.1410772 | -0.380279213 | 0.119380877 | 0.00068916 | 0.003837511 |
| CEP44 | 14023.2034 | -0.380299885 | 0.170734216 | 0.01093915 | 0.038187183 |
| NCDN | 2000.487955 | -0.380622699 | 0.123588366 | 0.00097845 | 0.00516583 |
| DISP2 | 1167.892004 | -0.380648385 | 0.163091635 | 0.00843822 | 0.030920097 |
| SEPTIN5 | 345.3056693 | -0.380851537 | 0.141875863 | 0.00329705 | 0.014237821 |
| RING1 | 1613.716973 | -0.381166842 | 0.100916415 | 7.79E-05 | 0.000595631 |
| POLE2 | 990.6030739 | -0.381197364 | 0.135724059 | 0.00228606 | 0.010549938 |
| WDR18 | 1686.744982 | -0.381200839 | 0.10237464 | 9.62E-05 | 0.000714394 |
| MFSD4B | 1533.480016 | -0.381305484 | 0.123599157 | 0.00095879 | 0.005080287 |
| SRD5A3 | 163.6658103 | -0.381378545 | 0.135646089 | 0.00226398 | 0.010469182 |
| STS | 861.7922357 | -0.381559001 | 0.111281001 | 0.00029199 | 0.001846935 |
| FKBP9 | 2444.022919 | -0.381591401 | 0.146631419 | 0.00414179 | 0.017206436 |
| TEN1-CDK3 | 155.8509705 | -0.381631759 | 0.169499926 | 0.01025762 | 0.036299717 |
| CHAC2 | 429.5962136 | -0.381875961 | 0.136725015 | 0.00238759 | 0.010920475 |
| ATXN7L3 | 5896.482421 | -0.381974423 | 0.123630702 | 0.00094151 | 0.005004178 |
| TMPO-AS1 | 358.4838737 | -0.382000336 | 0.122567407 | 0.00086109 | 0.004628165 |
| AGO1 | 4489.055001 | -0.382044841 | 0.116766463 | 0.00050849 | 0.002968919 |
| BOK | 113.5410491 | -0.382156354 | 0.161829289 | 0.00781529 | 0.029014924 |
| ENSAP2 | 113.0082549 | -0.382216726 | 0.166257298 | 0.00912046 | 0.033008296 |
| TUFM | 4967.67992 | -0.3822661 | 0.134584192 | 0.00206705 | 0.009724753 |
| MIR4453HG | 279.7786898 | -0.382412431 | 0.172412264 | 0.01105454 | 0.038516695 |
| TBL1X | 2167.186007 | -0.382455969 | 0.11002056 | 0.00024462 | 0.001591908 |
| KIF3C | 3297.005388 | -0.382536605 | 0.15231988 | 0.00528487 | 0.021020902 |
| NSD2 | 15495.27733 | -0.38269845 | 0.10888378 | 0.00021197 | 0.001407936 |
| LINC01291 | 190.7224138 | -0.382750598 | 0.153032416 | 0.0054265 | 0.02146798 |
| FAM136DP | 109.7775833 | -0.382794666 | 0.163825085 | 0.00828782 | 0.030461272 |
| ZDHHC8 | 2924.077481 | -0.38316318 | 0.121991363 | 0.00079024 | 0.004302092 |
| ENTREP3 | 3257.075763 | -0.383315972 | 0.148614277 | 0.00437779 | 0.018004954 |
| CHRNB2 | 2918.565255 | -0.38336663 | 0.152782955 | 0.00529346 | 0.021046927 |
| SLC27A2 | 131.4173211 | -0.38349913 | 0.158299062 | 0.00664186 | 0.025352939 |
| B3GAT3 | 1230.149619 | -0.383546656 | 0.091719548 | 1.43E-05 | 0.00013917 |

| | | | | | |
|---|---|---|---|---|---|
| EIF4E2 | 2382.046284 | -0.383581768 | 0.133395132 | 0.00184536 | 0.008833316 |
| THSD7B | 4546.091184 | -0.38359639 | 0.130447563 | 0.00150803 | 0.007413594 |
| E2F7 | 2035.726234 | -0.383673068 | 0.111431575 | 0.00027465 | 0.001754572 |
| RABL3 | 971.2597111 | -0.383826426 | 0.172760598 | 0.0108731 | 0.038026245 |
| PREP | 4099.803651 | -0.383830581 | 0.106346583 | 0.0001479 | 0.00104119 |
| BCL9 | 2987.820811 | -0.383862886 | 0.142878672 | 0.00322846 | 0.013980108 |
| EBF3 | 5403.724275 | -0.384065711 | 0.116800669 | 0.00047615 | 0.002807985 |
| MCM8 | 2555.710489 | -0.384066804 | 0.12687608 | 0.00114314 | 0.005896051 |
| SPEN-AS1 | 90.17894453 | -0.384132337 | 0.177504405 | 0.01239758 | 0.042223215 |
| IQSEC1 | 3897.041579 | -0.384231424 | 0.162107277 | 0.00756163 | 0.028215813 |
| CLVS2 | 8166.059446 | -0.384598223 | 0.136522947 | 0.00219303 | 0.010205374 |
| MRPL11 | 2560.915934 | -0.384955468 | 0.140776986 | 0.00279599 | 0.012480285 |
| GSTM1 | 987.1972747 | -0.385036746 | 0.132658749 | 0.00168691 | 0.00816802 |
| GVQW3 | 3236.1228 | -0.385045568 | 0.113044107 | 0.00031214 | 0.001958752 |
| NT5C | 685.3040697 | -0.385069385 | 0.10889851 | 0.00019384 | 0.001305657 |
| NUMA1 | 10804.01994 | -0.385188539 | 0.140687416 | 0.00276526 | 0.012356533 |
| PROS1 | 330.4392233 | -0.385215318 | 0.147142084 | 0.003895 | 0.01634961 |
| NOP14-AS1 | 453.4303934 | -0.38530941 | 0.145925718 | 0.00365537 | 0.01549877 |
| MMP16 | 2288.437562 | -0.385398797 | 0.130794294 | 0.00146763 | 0.00725491 |
| ERMAP | 242.1040929 | -0.38554236 | 0.137027108 | 0.00220607 | 0.010256772 |
| TRAF7 | 3258.917583 | -0.385628139 | 0.143191784 | 0.00314194 | 0.013709068 |
| CUTA | 4599.527319 | -0.385961323 | 0.111394986 | 0.00025126 | 0.001628403 |
| RFLNA | 379.0885172 | -0.385999045 | 0.137274106 | 0.00221275 | 0.010280816 |
| FBXL15 | 584.3042481 | -0.386037005 | 0.11102046 | 0.00024004 | 0.001567043 |
| CYB561A3 | 734.021469 | -0.386785319 | 0.116181628 | 0.00040759 | 0.00245794 |
| BSG | 9866.32962 | -0.386921874 | 0.135822183 | 0.00197116 | 0.009337677 |
| URM1 | 1883.461246 | -0.387356749 | 0.091019527 | 1.01E-05 | 0.000102466 |
| NXT2 | 256.2987826 | -0.3873874 | 0.15374683 | 0.00504276 | 0.020234009 |
| RELT | 626.9618021 | -0.387425458 | 0.104579406 | 0.00010089 | 0.000744661 |
| SLC25A5 | 6614.24141 | -0.387431047 | 0.126931342 | 0.00103756 | 0.005426328 |
| UBXN8 | 543.7079219 | -0.387518044 | 0.14985007 | 0.00420696 | 0.017431369 |
| APP | 23290.93161 | -0.387684927 | 0.12597102 | 0.00095439 | 0.005059595 |
| BCL11A | 2186.735544 | -0.387812228 | 0.119878121 | 0.00056276 | 0.003229862 |
| SCAMP5 | 8554.153097 | -0.387824934 | 0.078366993 | 3.69E-07 | 5.57E-06 |
| KIAA0930 | 2459.65934 | -0.387911087 | 0.11728737 | 0.00043782 | 0.002618747 |
| CCDC61 | 233.3570988 | -0.387996014 | 0.149624557 | 0.00411544 | 0.017124622 |
| CDK1 | 6316.184849 | -0.388052599 | 0.155732936 | 0.00541301 | 0.021418746 |
| MREG | 871.5992625 | -0.388119402 | 0.125880852 | 0.00093501 | 0.004977379 |
| SLC6A6 | 2470.215849 | -0.388195545 | 0.11714573 | 0.00042752 | 0.002565366 |
| HECTD1 | 6269.390896 | -0.388388407 | 0.125602504 | 0.00090676 | 0.004840766 |
| POLE4 | 554.2453346 | -0.388394611 | 0.129650433 | 0.00123907 | 0.006317911 |
| PCDHB10 | 448.4833329 | -0.388559556 | 0.133298557 | 0.00159607 | 0.007790544 |
| PGAP4 | 2109.440633 | -0.38858104 | 0.118434594 | 0.00047859 | 0.002819972 |
| ZCRB1 | 1666.058516 | -0.388685915 | 0.130926604 | 0.00134794 | 0.006767317 |
| MILIP | 325.3548437 | -0.38868675 | 0.136955829 | 0.00201889 | 0.009524317 |
| EEF1A1P6 | 855.6164103 | -0.38871138 | 0.145135358 | 0.00322853 | 0.013980108 |
| TMEM80 | 513.6783575 | -0.388990391 | 0.15160997 | 0.00441472 | 0.018127857 |
| IFRD2 | 1716.14934 | -0.389100394 | 0.12795183 | 0.00106792 | 0.005562506 |
| SSU72 | 1788.815318 | -0.389205926 | 0.128177892 | 0.00108317 | 0.005629071 |
| SKP1 | 579.6129047 | -0.389258696 | 0.119412889 | 0.00051351 | 0.002993121 |

| | | | | | |
|---|---|---|---|---|---|
| PIK3R3 | 3484.545101 | -0.389406543 | 0.117463205 | 0.0004232 | 0.002544623 |
| DKC1 | 7774.742973 | -0.389609323 | 0.124183011 | 0.00077652 | 0.004242724 |
| GAPDH | 84866.80184 | -0.389644955 | 0.109150901 | 0.00016747 | 0.001157608 |
| ALYREF | 6683.637854 | -0.389682099 | 0.133472696 | 0.00156502 | 0.007662724 |
| WDR13 | 1751.134218 | -0.389811482 | 0.088995141 | 5.74E-06 | 6.19E-05 |
| CHERP | 2669.704777 | -0.390229537 | 0.171701034 | 0.0092974 | 0.033504426 |
| PXMP2 | 259.0454146 | -0.390415799 | 0.131890253 | 0.00137361 | 0.006871557 |
| ANTKMT | 602.7317945 | -0.390628268 | 0.142791735 | 0.00271017 | 0.012142033 |
| GNL3L | 2792.042084 | -0.390715539 | 0.101861859 | 5.92E-05 | 0.00047076 |
| GPAT4 | 4543.966986 | -0.39076407 | 0.100772952 | 4.99E-05 | 0.000407796 |
| ANP32A | 18301.1759 | -0.390781131 | 0.124232655 | 0.00075157 | 0.004134642 |
| YJU2 | 478.0757168 | -0.390802722 | 0.14135716 | 0.00248737 | 0.011291435 |
| CLPTM1 | 2130.169347 | -0.39098901 | 0.149932862 | 0.00389222 | 0.016344975 |
| CPNE2 | 1061.078997 | -0.390991172 | 0.11319177 | 0.00025545 | 0.001652409 |
| ZNF692 | 2681.267271 | -0.391134062 | 0.083496771 | 1.36E-06 | 1.76E-05 |
| EPOP | 769.4434503 | -0.39170509 | 0.165928466 | 0.00743805 | 0.02781525 |
| CST3 | 1191.672887 | -0.391802357 | 0.113574693 | 0.00025874 | 0.001671091 |
| POLR1E | 2367.947827 | -0.391828435 | 0.125096666 | 0.00078195 | 0.004264137 |
| BRAT1 | 2197.164338 | -0.391890588 | 0.112005528 | 0.00021601 | 0.001431069 |
| ZNF576 | 475.7882514 | -0.391960718 | 0.116672565 | 0.00035772 | 0.002203758 |
| MT-ND4L | 16851.70773 | -0.392244022 | 0.140315358 | 0.00225458 | 0.010444499 |
| TRMT2B | 394.3327825 | -0.392307324 | 0.132651873 | 0.00137305 | 0.006871557 |
| TOP1MT | 812.0988937 | -0.392450489 | 0.116950202 | 0.00036176 | 0.002225283 |
| MTCO1P12 | 884.1247378 | -0.392465696 | 0.163487359 | 0.00669932 | 0.025535635 |
| LAT2 | 145.1077764 | -0.392478004 | 0.176828943 | 0.0104049 | 0.036738734 |
| CYB561D1 | 662.5305237 | -0.392829939 | 0.095423725 | 1.82E-05 | 0.00017241 |
| RNF26 | 1455.900952 | -0.392838164 | 0.110977811 | 0.00018478 | 0.001256581 |
| ST3GAL4 | 683.9163399 | -0.392931883 | 0.127253314 | 0.00090075 | 0.004815077 |
| MZT2A | 2110.404693 | -0.393165493 | 0.104316596 | 7.64E-05 | 0.000586567 |
| HMGN1 | 13897.15385 | -0.393350516 | 0.129417189 | 0.00105221 | 0.005491787 |
| FAR2 | 981.6910385 | -0.393457129 | 0.136002234 | 0.00166827 | 0.008098752 |
| SETMAR | 1625.091251 | -0.393474305 | 0.146765712 | 0.00312783 | 0.013667809 |
| GUCD1 | 1646.980183 | -0.393516141 | 0.099892971 | 3.83E-05 | 0.000325829 |
| TMEM203 | 1385.086683 | -0.39362318 | 0.110611539 | 0.00017163 | 0.001181958 |
| CALHM2 | 475.4174282 | -0.393631595 | 0.129816417 | 0.00107542 | 0.005591646 |
| RRP1 | 1974.238335 | -0.3937978 | 0.121600579 | 0.00054128 | 0.003125788 |
| SLC25A1 | 1940.716299 | -0.393877799 | 0.131016606 | 0.00116717 | 0.006006437 |
| CRYBG3 | 1099.045252 | -0.393885042 | 0.134945554 | 0.00153714 | 0.007542301 |
| RBM38 | 1097.188291 | -0.393886259 | 0.143199054 | 0.00255245 | 0.011541014 |
| NDE1 | 934.5519705 | -0.39401206 | 0.136432054 | 0.00168977 | 0.008174198 |
| SPATA31H1 | 310.5037329 | -0.394026741 | 0.115898841 | 0.00030694 | 0.00192967 |
| ADORA2B | 377.9245692 | -0.394066823 | 0.147337537 | 0.00317561 | 0.01381204 |
| SLC39A10 | 5654.19437 | -0.39414676 | 0.160261675 | 0.00570651 | 0.022377696 |
| TMEM201 | 1081.665133 | -0.394182026 | 0.18253528 | 0.011814 | 0.040632713 |
| IGSF8 | 639.5223172 | -0.394413651 | 0.11875154 | 0.00040481 | 0.002444089 |
| LRATD2 | 489.0599923 | -0.394871902 | 0.166022308 | 0.00699179 | 0.026420297 |
| EFTUD2 | 11061.53302 | -0.394877556 | 0.150293355 | 0.00361234 | 0.015363856 |
| HNRNPA1P10 | 867.6867346 | -0.394877764 | 0.128771731 | 0.0009568 | 0.005071047 |
| CDK18 | 226.7439634 | -0.395103595 | 0.158762554 | 0.00525823 | 0.020935181 |
| TSPAN17 | 1320.628663 | -0.395215532 | 0.117756081 | 0.0003566 | 0.00219819 |

| | | | | | |
|---|---|---|---|---|---|
| RFX2 | 177.6803317 | -0.395500249 | 0.170365837 | 0.00802215 | 0.029643739 |
| PSMB5 | 4269.80547 | -0.39552491 | 0.128765576 | 0.00093793 | 0.004990337 |
| WDR4 | 1267.627455 | -0.395556835 | 0.139175232 | 0.00192816 | 0.009176294 |
| DFFA | 1882.473931 | -0.395884426 | 0.096016275 | 1.75E-05 | 0.000166567 |
| H3P6 | 2336.462684 | -0.395911815 | 0.139670165 | 0.001969 | 0.009329595 |
| MYO1B | 7699.097139 | -0.396095261 | 0.12329497 | 0.00058503 | 0.003335377 |
| TCF4 | 18854.41581 | -0.396110233 | 0.129475201 | 0.00097354 | 0.005145189 |
| PARK7 | 3147.754609 | -0.396187395 | 0.139433636 | 0.00192647 | 0.009170368 |
| RHOT2 | 4780.601942 | -0.39652703 | 0.10569094 | 8.06E-05 | 0.000613865 |
| IKBKB | 1674.454456 | -0.39672818 | 0.091402046 | 6.68E-06 | 7.08E-05 |
| DCTN6 | 856.5717797 | -0.397010735 | 0.147928302 | 0.00304699 | 0.013388525 |
| GPR75 | 323.6241006 | -0.397076743 | 0.142576218 | 0.00227008 | 0.010486419 |
| INSM1 | 3720.570396 | -0.397183658 | 0.193014509 | 0.01446282 | 0.047731262 |
| ASAP2 | 1570.848116 | -0.397212555 | 0.123006302 | 0.00054992 | 0.003169417 |
| TMEM276 | 189.632154 | -0.397249982 | 0.1668706 | 0.00686254 | 0.026027524 |
| NPM3 | 1055.727516 | -0.397729563 | 0.143802671 | 0.00239447 | 0.01094033 |
| DTD2 | 347.717142 | -0.398014827 | 0.120002773 | 0.00040468 | 0.002443975 |
| SKP2 | 8115.282824 | -0.39824413 | 0.131819073 | 0.00108999 | 0.00565881 |
| RAB11FIP1P1 | 174.5489038 | -0.398264711 | 0.183248548 | 0.01117486 | 0.038870077 |
| LINC00294 | 724.8606544 | -0.39828326 | 0.101038841 | 3.72E-05 | 0.000317367 |
| SNCB | 90.36201474 | -0.398754064 | 0.175442428 | 0.00885192 | 0.032223541 |
| SEPHS1 | 6384.741505 | -0.398848482 | 0.098414528 | 2.33E-05 | 0.00021304 |
| MYO10 | 989.6899013 | -0.3988721 | 0.169051334 | 0.0071679 | 0.026971756 |
| MTERF3 | 953.8329652 | -0.398892283 | 0.145777571 | 0.00259449 | 0.011690783 |
| PPIAP22 | 306.7750868 | -0.399031371 | 0.178386424 | 0.0096176 | 0.034419832 |
| MCM6 | 8955.555655 | -0.399299153 | 0.138657713 | 0.00168827 | 0.008170937 |
| PTGFRN | 3937.07223 | -0.399349852 | 0.138871906 | 0.001709 | 0.008253592 |
| GLI4 | 344.0455293 | -0.399421027 | 0.123118141 | 0.00051699 | 0.003009173 |
| TRIL | 309.9175417 | -0.399432144 | 0.148031382 | 0.00288707 | 0.012794685 |
| CAD | 6729.117851 | -0.399583964 | 0.171475208 | 0.00767106 | 0.028572291 |
| ATP5MC3 | 5204.354447 | -0.399932174 | 0.127796368 | 0.00075955 | 0.004168499 |
| SLC16A1 | 13911.90957 | -0.39996304 | 0.140467609 | 0.00185669 | 0.008883417 |
| UBL7-DT | 230.1912846 | -0.399983437 | 0.149653749 | 0.00309633 | 0.013567643 |
| UBE2D4 | 383.702299 | -0.400050226 | 0.124954809 | 0.00059615 | 0.003387643 |
| GALNT7 | 2359.435204 | -0.400081167 | 0.127367871 | 0.00073009 | 0.00402724 |
| NFYC-AS1 | 152.3419743 | -0.400121721 | 0.195820322 | 0.01463069 | 0.048200185 |
| TEDC2-AS1 | 87.2942769 | -0.40036606 | 0.189796991 | 0.01270171 | 0.043052248 |
| HNRNPA3 | 14552.54445 | -0.400415949 | 0.128575886 | 0.00079688 | 0.004329446 |
| RPL7A | 36765.29617 | -0.400499564 | 0.142021987 | 0.00201062 | 0.009489627 |
| MRPS26 | 1599.114511 | -0.400620459 | 0.08926114 | 3.33E-06 | 3.84E-05 |
| SPAG5-AS1 | 227.6873486 | -0.400813638 | 0.139466858 | 0.0017057 | 0.008239617 |
| CYFIP1 | 4752.147225 | -0.40115283 | 0.104663208 | 5.72E-05 | 0.000456471 |
| TAGLN3 | 3069.960085 | -0.401153577 | 0.113738983 | 0.00018671 | 0.001265911 |
| NDUFS2 | 4960.940074 | -0.401167577 | 0.115402498 | 0.00022512 | 0.001481373 |
| SLC35F2 | 879.383289 | -0.4011885 | 0.123238089 | 0.00049337 | 0.002896182 |
| TIMM8B | 957.7918236 | -0.401206798 | 0.113111297 | 0.00017324 | 0.001191202 |
| GALNT1 | 2344.138567 | -0.401252602 | 0.129246864 | 0.00081942 | 0.004433162 |
| NAXE | 1536.090274 | -0.401369921 | 0.118937721 | 0.00032469 | 0.002023898 |
| SDC3 | 1458.3284 | -0.40154977 | 0.134379059 | 0.00119126 | 0.006113583 |
| SERBP1P1 | 405.2897058 | -0.401778546 | 0.135126168 | 0.001247 | 0.00635052 |

| | | | | | |
|---|---|---|---|---|---|
| LPL | 139.4387514 | -0.401804521 | 0.187615902 | 0.01173568 | 0.040451323 |
| SLC9A3 | 311.8957905 | -0.401813572 | 0.161639818 | 0.00510677 | 0.02042311 |
| SERBP1 | 37556.9773 | -0.401876519 | 0.126038498 | 0.00061762 | 0.003494946 |
| ELAVL2 | 5676.764916 | -0.401958124 | 0.135549592 | 0.00127752 | 0.00647707 |
| H1-10-AS1 | 444.556664 | -0.402105112 | 0.124633138 | 0.00054271 | 0.003132262 |
| ABCC1 | 5365.943537 | -0.402243244 | 0.153382617 | 0.0035228 | 0.015066293 |
| PLPP3 | 341.9103518 | -0.402305136 | 0.156575517 | 0.00407453 | 0.016978461 |
| PYM1 | 772.0547652 | -0.402732722 | 0.118218644 | 0.00028767 | 0.001824714 |
| MIATNB | 147.0645571 | -0.402752487 | 0.168600766 | 0.00651693 | 0.024983982 |
| ARL10 | 6429.057174 | -0.402841515 | 0.092207952 | 5.72E-06 | 6.18E-05 |
| TPGS2 | 4740.265456 | -0.402981069 | 0.092401592 | 5.92E-06 | 6.35E-05 |
| PGD | 4271.05167 | -0.403113707 | 0.129501425 | 0.00079025 | 0.004302092 |
| SKI | 2005.113114 | -0.403168941 | 0.174300148 | 0.00783401 | 0.029063843 |
| PRELID1 | 2281.542012 | -0.403326506 | 0.130449003 | 0.00084574 | 0.004555234 |
| SMO | 2591.763991 | -0.403669374 | 0.098307271 | 1.82E-05 | 0.000172241 |
| ARPIN | 1454.866964 | -0.403676933 | 0.103255783 | 4.15E-05 | 0.000349435 |
| PHF7 | 187.8976736 | -0.403720408 | 0.157960977 | 0.00419476 | 0.017398365 |
| PTCHD4 | 509.9812402 | -0.403809985 | 0.147508097 | 0.00252014 | 0.011417535 |
| UQCC2 | 1342.855847 | -0.403841196 | 0.140260718 | 0.00165276 | 0.008027259 |
| CD2BP2 | 3155.055164 | -0.404153075 | 0.087822087 | 1.92E-06 | 2.36E-05 |
| SMARCD2 | 4875.013998 | -0.404182822 | 0.108463335 | 8.61E-05 | 0.000648775 |
| RPH3A | 786.0760841 | -0.404248462 | 0.171793099 | 0.00706035 | 0.026640181 |
| CDKL1 | 631.6074303 | -0.404266463 | 0.14811707 | 0.00257434 | 0.011617003 |
| DNPEP | 1154.92461 | -0.404277346 | 0.113435978 | 0.00016034 | 0.001116833 |
| TMEM240 | 102.0525515 | -0.404425171 | 0.182640248 | 0.00980964 | 0.03496714 |
| HPS6 | 602.4195074 | -0.40447643 | 0.117410899 | 0.00024849 | 0.001613475 |
| CANT1 | 3260.602861 | -0.404743903 | 0.106289785 | 6.22E-05 | 0.000491302 |
| KEAP1 | 3151.229064 | -0.404788735 | 0.145302288 | 0.00217688 | 0.010148583 |
| EBF1 | 1336.200634 | -0.404980595 | 0.140379545 | 0.00161495 | 0.007867745 |
| PAICS | 21587.89276 | -0.405117252 | 0.121379935 | 0.00036373 | 0.00223407 |
| CTPS1 | 7528.500319 | -0.4051541 | 0.108873913 | 8.74E-05 | 0.000657085 |
| ZNF180 | 868.995599 | -0.405204855 | 0.129902566 | 0.00076567 | 0.00419314 |
| DDX39A | 5536.771822 | -0.405365921 | 0.107505556 | 7.19E-05 | 0.000556814 |
| ZNRF1 | 755.1246085 | -0.405375549 | 0.142569738 | 0.00182825 | 0.00875755 |
| CASP6 | 203.0741467 | -0.40544218 | 0.141654004 | 0.00172641 | 0.008327896 |
| RNPEPL1 | 738.0231011 | -0.40551516 | 0.150807568 | 0.00287174 | 0.012736663 |
| MAP3K6 | 292.4818137 | -0.405540975 | 0.201042083 | 0.01489823 | 0.048921315 |
| ITGA9-AS1 | 104.7110253 | -0.405550425 | 0.160635299 | 0.00451455 | 0.0184677 |
| ENDOD1 | 358.1110284 | -0.405692446 | 0.182433382 | 0.00952526 | 0.034142836 |
| CDPF1 | 210.559693 | -0.405762133 | 0.141832824 | 0.00173066 | 0.008344475 |
| CYB5R3 | 1884.104841 | -0.405774298 | 0.153671775 | 0.00328728 | 0.014198638 |
| COBLL1 | 308.7089339 | -0.405790867 | 0.135233182 | 0.0011211 | 0.005805654 |
| PRXL2B | 580.8493132 | -0.406018908 | 0.138268777 | 0.00137036 | 0.006864826 |
| CHMP7 | 1835.168364 | -0.406085525 | 0.103036747 | 3.60E-05 | 0.000309081 |
| NDRG4 | 4732.10817 | -0.406419538 | 0.104282346 | 4.30E-05 | 0.000360613 |
| MRM1 | 206.7279253 | -0.406540989 | 0.160797593 | 0.00444676 | 0.018241222 |
| TMED1 | 223.8892167 | -0.406633343 | 0.126819012 | 0.00056819 | 0.003251951 |
| LDB2 | 257.8539393 | -0.406927827 | 0.177967055 | 0.00816447 | 0.03006161 |
| BLACAT1 | 144.6515765 | -0.407093937 | 0.187555445 | 0.01065251 | 0.037420251 |
| USP13 | 2000.10628 | -0.407158305 | 0.114147837 | 0.00015644 | 0.001093467 |

| | | | | | |
|---|---|---|---|---|---|
| CNTFR-AS1 | 200.6811888 | -0.407493841 | 0.128717044 | 0.00064869 | 0.003648667 |
| TIMM50 | 3775.02665 | -0.407947977 | 0.094265939 | 6.74E-06 | 7.13E-05 |
| ADAMTS16 | 106.566791 | -0.408018286 | 0.188258269 | 0.01066641 | 0.037462677 |
| FLNB | 5435.513398 | -0.408225661 | 0.176444594 | 0.00759315 | 0.028312862 |
| ASIC1 | 689.2306375 | -0.408259635 | 0.151942699 | 0.00284618 | 0.012642065 |
| FCSK | 200.4201974 | -0.408288525 | 0.157229008 | 0.00365969 | 0.015513889 |
| MAN2C1 | 4978.615623 | -0.408524305 | 0.089377058 | 2.19E-06 | 2.64E-05 |
| HNRNPM | 17408.68497 | -0.408919392 | 0.124774498 | 0.00044087 | 0.002633181 |
| VOPP1 | 1396.963871 | -0.408964829 | 0.119978849 | 0.00027738 | 0.001767893 |
| TRIB1 | 570.6610273 | -0.409054551 | 0.12654699 | 0.00051396 | 0.002994929 |
| SLC39A8 | 302.093886 | -0.409125757 | 0.156677042 | 0.00350066 | 0.014991148 |
| BOLA3-DT | 1195.877764 | -0.409187941 | 0.104319183 | 3.84E-05 | 0.000325919 |
| TSC22D2 | 4118.877994 | -0.409222512 | 0.118530065 | 0.00023645 | 0.001547533 |
| HNRNPF | 7884.397086 | -0.409493874 | 0.106024343 | 4.89E-05 | 0.000401492 |
| UBE2R2 | 4083.040581 | -0.409496037 | 0.129473361 | 0.00064886 | 0.003648667 |
| P2RX3 | 799.6155454 | -0.409610389 | 0.174746242 | 0.00699543 | 0.026429163 |
| PMP22 | 288.1440479 | -0.40961091 | 0.167408392 | 0.00540841 | 0.021406198 |
| NOPCHAP1 | 2413.012574 | -0.409652882 | 0.119571164 | 0.0002597 | 0.001676246 |
| RANP1 | 132.9313758 | -0.409730929 | 0.19911637 | 0.01334125 | 0.044755225 |
| GPR153 | 199.6852179 | -0.409794142 | 0.180135671 | 0.0082478 | 0.030335854 |
| RNASEH2C | 1501.47104 | -0.41003312 | 0.109839115 | 8.17E-05 | 0.000621266 |
| DUXAP8 | 1580.079287 | -0.410049236 | 0.102002443 | 2.55E-05 | 0.00023035 |
| PLP2 | 639.7303031 | -0.410199529 | 0.140415922 | 0.00140575 | 0.007004587 |
| DDA1 | 1763.060862 | -0.410207526 | 0.103415816 | 3.18E-05 | 0.000278271 |
| ZNF84 | 6520.939931 | -0.410275656 | 0.149212889 | 0.00235158 | 0.010780906 |
| TMEM119 | 622.265167 | -0.410307365 | 0.116483539 | 0.00018196 | 0.001241449 |
| PTRHD1 | 396.296363 | -0.410324438 | 0.149593364 | 0.00239691 | 0.01094847 |
| RXRA | 1948.330913 | -0.410412312 | 0.131733396 | 0.00075565 | 0.004152045 |
| STAP2 | 173.4857929 | -0.410429882 | 0.169605285 | 0.00576429 | 0.022569163 |
| MON1A | 630.4479005 | -0.410430193 | 0.130936264 | 0.00070939 | 0.003933097 |
| ZNF169 | 466.3385338 | -0.410465126 | 0.150642879 | 0.00252422 | 0.01143097 |
| TMEM145 | 2257.851967 | -0.410498678 | 0.121668017 | 0.00031172 | 0.001956699 |
| HMGB1P5 | 2191.263936 | -0.410729455 | 0.156614417 | 0.00336342 | 0.014475784 |
| PERPP2 | 344.7842955 | -0.410791016 | 0.126715651 | 0.00049349 | 0.002896182 |
| SBNO1-AS1 | 52.6642792 | -0.410792488 | 0.202557091 | 0.01407542 | 0.046707591 |
| KCNJ11 | 261.1147381 | -0.410820429 | 0.135527232 | 0.00099133 | 0.005221768 |
| YBX1P10 | 408.9330246 | -0.410975148 | 0.153130614 | 0.00282946 | 0.012591428 |
| WASF3 | 1759.927099 | -0.410992873 | 0.109759348 | 7.77E-05 | 0.000594642 |
| NCL | 49060.78257 | -0.411043991 | 0.122600561 | 0.00033518 | 0.00208174 |
| DHX37 | 2267.46795 | -0.411045413 | 0.169661354 | 0.00570261 | 0.022366679 |
| KHDRBS1 | 13095.53651 | -0.411201397 | 0.129038197 | 0.00059389 | 0.003379268 |
| METTL8 | 1230.344731 | -0.411205939 | 0.104201451 | 3.45E-05 | 0.000298493 |
| ASB16-AS1 | 908.2741835 | -0.411571608 | 0.143010394 | 0.00159445 | 0.007784488 |
| SHISA6 | 377.502172 | -0.411714116 | 0.18716506 | 0.00969259 | 0.034621923 |
| TRAF4 | 2503.731191 | -0.41202931 | 0.117829608 | 0.00019838 | 0.001331667 |
| DARS2 | 2829.808058 | -0.412087954 | 0.113682067 | 0.00012279 | 0.000885362 |
| CTSA | 1605.67418 | -0.41216102 | 0.085174079 | 5.82E-07 | 8.32E-06 |
| GALNT17 | 1017.239953 | -0.412207672 | 0.160508122 | 0.00387079 | 0.01626793 |
| LAMTOR1 | 1318.023096 | -0.412266351 | 0.117332841 | 0.00018624 | 0.001263539 |
| AAAS | 3045.159963 | -0.412375406 | 0.112847996 | 0.00010962 | 0.000801631 |

| | | | | | |
|---|---|---|---|---|---|
| U2AF2 | 10724.12937 | -0.412478333 | 0.105391584 | 3.92E-05 | 0.000332028 |
| NGRN | 3425.855448 | -0.412572652 | 0.092073736 | 3.28E-06 | 3.78E-05 |
| EIF3CL | 557.6118496 | -0.412573214 | 0.20332501 | 0.01387703 | 0.046131128 |
| POLR2D | 3731.712375 | -0.412620655 | 0.09472689 | 5.81E-06 | 6.25E-05 |
| DENND6A | 2347.842283 | -0.412715541 | 0.085017181 | 5.37E-07 | 7.78E-06 |
| PAG1 | 1015.812053 | -0.412783985 | 0.149594772 | 0.0022552 | 0.010445018 |
| B4GALNT4 | 8005.270913 | -0.412807107 | 0.114088104 | 0.00012548 | 0.000900367 |
| CD9 | 829.7798471 | -0.413088599 | 0.119938203 | 0.00023921 | 0.001562644 |
| IPP | 312.1314764 | -0.413172684 | 0.153273863 | 0.00270336 | 0.012114176 |
| ATP6V0E1 | 1388.283617 | -0.413401614 | 0.137821454 | 0.00108243 | 0.005626682 |
| ZNF497 | 115.0965351 | -0.413462441 | 0.167996333 | 0.00509721 | 0.020400762 |
| NONO | 29435.08667 | -0.413464552 | 0.10711069 | 4.85E-05 | 0.000398655 |
| CTBP1-DT | 869.5714772 | -0.413522602 | 0.122947132 | 0.00031885 | 0.001994774 |
| ZNF784 | 274.3906307 | -0.413644907 | 0.141999148 | 0.00141659 | 0.007050045 |
| FUOM | 239.665469 | -0.413660618 | 0.144108039 | 0.00161298 | 0.007860019 |
| DSCC1 | 1530.308863 | -0.413688545 | 0.123722737 | 0.00034161 | 0.002114643 |
| AFG3L1P | 1132.418101 | -0.413795909 | 0.092116509 | 3.09E-06 | 3.60E-05 |
| HIPK2 | 8275.089036 | -0.413815425 | 0.166129056 | 0.00470862 | 0.019135971 |
| BRD3 | 2935.000392 | -0.414042331 | 0.133668896 | 0.00078697 | 0.004288072 |
| CCDC3 | 3759.34504 | -0.414183138 | 0.128455437 | 0.00051513 | 0.002999179 |
| TSPAN14 | 1792.245273 | -0.414253696 | 0.118174002 | 0.00019014 | 0.001285328 |
| MGAT1 | 1426.442974 | -0.414490452 | 0.113821446 | 0.00011393 | 0.000827535 |
| CCDC183-AS1 | 88.79987948 | -0.414643262 | 0.208745941 | 0.01489106 | 0.048909094 |
| POMGNT1 | 1520.050283 | -0.414694973 | 0.097830992 | 9.72E-06 | 9.85E-05 |
| ATP5MC2 | 7045.620606 | -0.414930373 | 0.121708726 | 0.00026896 | 0.001727322 |
| LDHB | 2414.722231 | -0.41501352 | 0.140989627 | 0.00127957 | 0.006485851 |
| EIF4A1P10 | 446.6981618 | -0.415244436 | 0.13725723 | 0.00098781 | 0.005208556 |
| NME2 | 491.306297 | -0.415318019 | 0.117840331 | 0.00017637 | 0.001210143 |
| XYLB | 941.1074195 | -0.41563353 | 0.117546182 | 0.00016874 | 0.001164895 |
| DCAF1 | 3041.713873 | -0.415831417 | 0.115090359 | 0.00012618 | 0.000904381 |
| SAMD1 | 2236.292473 | -0.41590829 | 0.156527682 | 0.00296863 | 0.013091778 |
| MRPL28 | 1359.377956 | -0.416126468 | 0.097273076 | 8.12E-06 | 8.42E-05 |
| BLOC1S1 | 189.8604178 | -0.416207729 | 0.170660173 | 0.00531069 | 0.021095026 |
| IDUA | 490.2322638 | -0.416291773 | 0.108741627 | 5.44E-05 | 0.000437129 |
| NDUFS8 | 2122.637104 | -0.41650148 | 0.13895371 | 0.00107237 | 0.005578623 |
| WDR6 | 9885.707047 | -0.416660279 | 0.094696629 | 4.67E-06 | 5.16E-05 |
| EHMT2 | 5549.994521 | -0.416756017 | 0.114279508 | 0.00011047 | 0.0008073 |
| HCFC1R1 | 289.2986317 | -0.416793093 | 0.161476419 | 0.00364099 | 0.015454609 |
| VCF2 | 557.1486206 | -0.41687159 | 0.163577549 | 0.00397388 | 0.016636566 |
| SRRT | 9469.20814 | -0.416911617 | 0.121209911 | 0.00023875 | 0.001560608 |
| HMGN2 | 22722.28025 | -0.417006436 | 0.126687356 | 0.00040312 | 0.002436704 |
| PRR11 | 9551.355562 | -0.417013663 | 0.092478023 | 2.81E-06 | 3.31E-05 |
| GRIN1 | 1656.323569 | -0.417074381 | 0.133506717 | 0.00071037 | 0.00393537 |
| NRIP1 | 6237.627528 | -0.417185296 | 0.12817155 | 0.00045714 | 0.0027115 |
| CIART | 202.0495336 | -0.417306819 | 0.143078287 | 0.00137404 | 0.006871557 |
| ILVBL | 2183.008166 | -0.41733037 | 0.098455086 | 9.61E-06 | 9.74E-05 |
| ETAA1 | 2951.448976 | -0.417439324 | 0.129227669 | 0.00049666 | 0.002912239 |
| HNRNPA3P6 | 106.0264661 | -0.417611246 | 0.190073439 | 0.00939031 | 0.033765053 |
| NEK6 | 352.6523352 | -0.417615913 | 0.168921379 | 0.00483391 | 0.019560124 |
| ICMT | 2077.797366 | -0.417749874 | 0.086366786 | 5.74E-07 | 8.23E-06 |

| | | | | | |
|---|---|---|---|---|---|
| COQ8A | 148.9205328 | -0.417914869 | 0.156428498 | 0.00281838 | 0.012550281 |
| MYO1B-AS1 | 96.22625339 | -0.417944722 | 0.189761724 | 0.00925816 | 0.033406694 |
| GAR1 | 840.036512 | -0.418531375 | 0.113613549 | 9.50E-05 | 0.000707859 |
| RAN | 29493.24707 | -0.418541671 | 0.142162489 | 0.00125388 | 0.00637762 |
| PHLDA1 | 2660.633243 | -0.418661946 | 0.129466667 | 0.00048778 | 0.002869176 |
| APEX1 | 9016.743872 | -0.418878967 | 0.142669241 | 0.00128344 | 0.006502265 |
| FOXN4 | 541.9200689 | -0.419209849 | 0.144336004 | 0.00141219 | 0.007031541 |
| KCNB2 | 897.5739126 | -0.419222727 | 0.170061198 | 0.00487671 | 0.019699568 |
| GHDC | 425.6358836 | -0.419310376 | 0.134229463 | 0.00070262 | 0.003899307 |
| SAYSD1 | 735.2511838 | -0.419325453 | 0.159081217 | 0.00308863 | 0.013539674 |
| MLLT1 | 4343.109255 | -0.419806039 | 0.146016305 | 0.0015393 | 0.007551072 |
| AGAP3 | 4690.758708 | -0.419937584 | 0.118087857 | 0.00015326 | 0.001073812 |
| PABPN1 | 2601.842177 | -0.419971176 | 0.112757659 | 8.06E-05 | 0.000613773 |
| CYB561D2 | 296.2429694 | -0.420132564 | 0.180201756 | 0.00676448 | 0.025717279 |
| NAA40 | 3374.861243 | -0.420329853 | 0.081723588 | 1.17E-07 | 2.04E-06 |
| NDUFB7 | 1332.411854 | -0.420336039 | 0.135262254 | 0.00073726 | 0.004065717 |
| FBL | 6464.141732 | -0.420473513 | 0.118481082 | 0.00015707 | 0.001096743 |
| GNG4 | 9252.89355 | -0.42055754 | 0.091670462 | 1.91E-06 | 2.36E-05 |
| DHRS13 | 365.1586335 | -0.420767716 | 0.13475734 | 0.00070061 | 0.003889667 |
| LINC01829 | 4608.736593 | -0.420780442 | 0.134078344 | 0.00066474 | 0.003727353 |
| CENPI | 1763.18606 | -0.420928294 | 0.127604674 | 0.00038566 | 0.00235054 |
| IGLVIVOR22-1 | 56.70674294 | -0.421188701 | 0.21047007 | 0.01382136 | 0.046013007 |
| FBXO43 | 521.6278111 | -0.421192299 | 0.142887057 | 0.00122242 | 0.006242324 |
| ZMYND8 | 2170.94308 | -0.421247845 | 0.108556468 | 4.31E-05 | 0.000360613 |
| CBX2 | 13785.99925 | -0.421281221 | 0.114595841 | 9.65E-05 | 0.000716733 |
| MEX3A | 13349.72219 | -0.421450496 | 0.09339196 | 2.71E-06 | 3.21E-05 |
| MDC1 | 4164.891748 | -0.421604306 | 0.151723572 | 0.00203039 | 0.009568279 |
| C2orf72 | 238.7483618 | -0.421875267 | 0.159957506 | 0.00303126 | 0.013330795 |
| ZBTB47 | 425.9728974 | -0.421923097 | 0.13738885 | 0.00082357 | 0.004449793 |
| ASIC4 | 323.8477371 | -0.422036397 | 0.19425719 | 0.0096324 | 0.034464342 |
| LARP1 | 15331.91248 | -0.422120071 | 0.168018405 | 0.00424025 | 0.017548082 |
| SLC37A4 | 1319.084634 | -0.422333842 | 0.112262417 | 6.88E-05 | 0.000536234 |
| EIF5AP4 | 175.3347075 | -0.422360803 | 0.191308435 | 0.00889058 | 0.032341396 |
| TRIB2 | 4300.561393 | -0.422582823 | 0.106201332 | 2.85E-05 | 0.000253252 |
| CHST2 | 3235.71164 | -0.422674808 | 0.147917833 | 0.00159796 | 0.00779792 |
| RBFA | 1600.830872 | -0.422894686 | 0.113561021 | 7.96E-05 | 0.000607285 |
| SYBU | 924.540988 | -0.422895981 | 0.125405316 | 0.00029499 | 0.001861326 |
| ZNF74 | 2216.540427 | -0.42303247 | 0.09189574 | 1.76E-06 | 2.19E-05 |
| FANCB | 583.7344755 | -0.423166709 | 0.155693941 | 0.00239946 | 0.010957689 |
| ESYT3 | 250.5761931 | -0.423264273 | 0.161327214 | 0.00312353 | 0.013665243 |
| TET1 | 1116.913968 | -0.423706127 | 0.108429044 | 3.81E-05 | 0.000324036 |
| RBBP9 | 1132.104431 | -0.4237954 | 0.115251439 | 9.50E-05 | 0.000707859 |
| HMGB1P10 | 388.8206236 | -0.423802698 | 0.18979398 | 0.00832029 | 0.030553293 |
| CHAF1A | 4360.859053 | -0.42391357 | 0.129356082 | 0.00040929 | 0.002467487 |
| MIR17HG | 337.969633 | -0.424398022 | 0.193924541 | 0.00915651 | 0.033103807 |
| FAM157C | 110.4830164 | -0.424420997 | 0.216691583 | 0.01464977 | 0.048247566 |
| C2orf69 | 1105.217295 | -0.424697007 | 0.125915395 | 0.00029164 | 0.001845313 |
| LINC02268 | 4724.743843 | -0.424754449 | 0.112223634 | 6.21E-05 | 0.000490732 |
| TOR4A | 143.1520729 | -0.424797904 | 0.168690087 | 0.00410918 | 0.017105508 |
| SFPQ | 34509.56263 | -0.424849277 | 0.083552127 | 1.56E-07 | 2.62E-06 |

| | | | | | |
|---|---|---|---|---|---|
| S100A6 | 185.5196047 | -0.424887211 | 0.181492071 | 0.00641537 | 0.024669864 |
| ZNF710-AS1 | 714.9095444 | -0.424910231 | 0.16210103 | 0.00311395 | 0.013639013 |
| KIAA2013 | 1458.450822 | -0.425304983 | 0.13757813 | 0.00075725 | 0.004157049 |
| SUPT16H | 14999.2077 | -0.425316305 | 0.136664306 | 0.00070759 | 0.003925222 |
| RBM15B | 5581.845588 | -0.425413915 | 0.13811531 | 0.00078494 | 0.004278176 |
| C1QL3 | 185.5678412 | -0.425478977 | 0.177927759 | 0.00565452 | 0.022194979 |
| NBEAL2 | 3535.349723 | -0.425669845 | 0.120495621 | 0.00016258 | 0.001130274 |
| MDH2 | 6556.940107 | -0.425698199 | 0.145878717 | 0.00130695 | 0.006599768 |
| PRMT1 | 10901.68799 | -0.425704237 | 0.10311352 | 1.49E-05 | 0.000144857 |
| CHAF1B | 1759.757025 | -0.425728908 | 0.127604078 | 0.00033002 | 0.002053384 |
| MKI67 | 19189.42477 | -0.425850591 | 0.116714026 | 0.00010495 | 0.00077133 |
| SCAMP1-AS1 | 238.63959 | -0.426127787 | 0.137364059 | 0.00072784 | 0.004019151 |
| GAL | 7831.855495 | -0.42613223 | 0.14217321 | 0.00101924 | 0.005341417 |
| CDH10 | 176.8176128 | -0.42615285 | 0.162951682 | 0.00314308 | 0.013711139 |
| FNBP1P1 | 344.5563486 | -0.426221572 | 0.118967762 | 0.00013453 | 0.000959936 |
| SIVA1 | 2421.070781 | -0.426247254 | 0.105458502 | 2.16E-05 | 0.000199294 |
| SINHCAF | 14129.88735 | -0.426272028 | 0.130646927 | 0.00042476 | 0.002552522 |
| RPS3P4 | 230.2789818 | -0.426587229 | 0.163255292 | 0.00315399 | 0.013747063 |
| MTND4P12 | 98.19470689 | -0.426606241 | 0.183725994 | 0.00664276 | 0.025352939 |
| POLR2F | 1461.08481 | -0.426921936 | 0.126966437 | 0.00029915 | 0.00188415 |
| URB2 | 3529.698976 | -0.426996682 | 0.130060735 | 0.00039464 | 0.002394605 |
| SLC25A29 | 4777.993327 | -0.427006009 | 0.113806254 | 6.99E-05 | 0.000544548 |
| DCLK2 | 1205.16442 | -0.427014337 | 0.090791493 | 1.07E-06 | 1.42E-05 |
| ZNRF2 | 412.2815133 | -0.427152985 | 0.134397727 | 0.00056295 | 0.003230032 |
| GPR162 | 689.5073018 | -0.427339693 | 0.124064231 | 0.00022264 | 0.001468861 |
| C19orf12 | 1330.106727 | -0.427498145 | 0.083171821 | 1.15E-07 | 2.01E-06 |
| NIN | 3599.27238 | -0.428033437 | 0.132508279 | 0.00047032 | 0.002781624 |
| CENPF | 30746.3214 | -0.428066348 | 0.143366955 | 0.00104563 | 0.005460074 |
| GPI | 8725.302293 | -0.428157981 | 0.13184957 | 0.00044345 | 0.002644763 |
| ERVK13-1 | 1399.548177 | -0.428165348 | 0.1196215 | 0.00013491 | 0.000962276 |
| MYO19 | 3366.063155 | -0.42826114 | 0.095421236 | 2.95E-06 | 3.46E-05 |
| AP2S1 | 1152.646895 | -0.428401121 | 0.123462571 | 0.00020196 | 0.001350137 |
| ZC3H10 | 657.5454079 | -0.428489665 | 0.100643286 | 8.40E-06 | 8.65E-05 |
| DACH1 | 5009.008733 | -0.42850554 | 0.114686544 | 7.38E-05 | 0.000569259 |
| MMP11 | 235.3190095 | -0.428550132 | 0.146630721 | 0.00126894 | 0.006439894 |
| TINF2 | 1055.883869 | -0.428717853 | 0.115120742 | 7.73E-05 | 0.00059232 |
| CD248 | 806.9279201 | -0.428850192 | 0.197687636 | 0.0092675 | 0.03343455 |
| POLH-AS1 | 147.3971463 | -0.429050255 | 0.150553967 | 0.00157824 | 0.007717988 |
| ATG9B | 379.2846018 | -0.429050874 | 0.130367562 | 0.00037975 | 0.002317963 |
| FAM163A | 3066.95184 | -0.429124381 | 0.121768551 | 0.00016492 | 0.001144215 |
| ST8SIA4 | 215.5399389 | -0.42919701 | 0.142981935 | 0.00098829 | 0.005209716 |
| SPOCK1 | 6455.91795 | -0.429281012 | 0.169349205 | 0.00382782 | 0.01613016 |
| COPS7A | 1424.408949 | -0.429310452 | 0.089486823 | 6.63E-07 | 9.28E-06 |
| TNPO2 | 8698.579291 | -0.429365707 | 0.121445186 | 0.00015793 | 0.001102031 |
| SORD | 4450.521477 | -0.429604734 | 0.115986936 | 8.33E-05 | 0.000630757 |
| ARK2C | 4182.238715 | -0.429730635 | 0.094010963 | 1.98E-06 | 2.43E-05 |
| DAPK1 | 2813.888435 | -0.429760568 | 0.12230908 | 0.00017081 | 0.00117709 |
| VPS51 | 2397.423382 | -0.429771773 | 0.137042491 | 0.00063872 | 0.003599882 |
| CCDC86 | 1022.743838 | -0.429804408 | 0.151472416 | 0.00163033 | 0.007935162 |
| PRDM10 | 825.5771865 | -0.42984321 | 0.130105064 | 0.00036186 | 0.002225283 |

| | | | | | |
|---|---|---|---|---|---|
| ESRRA | 895.0500069 | -0.429991872 | 0.114739822 | 7.01E-05 | 0.000545793 |
| B4GAT1 | 1446.594345 | -0.430169561 | 0.114598671 | 6.84E-05 | 0.000533434 |
| FAM215B | 73.7163711 | -0.430184926 | 0.202423166 | 0.01009328 | 0.03579206 |
| SLC25A23 | 1722.301764 | -0.430218257 | 0.140249151 | 0.00079674 | 0.004329446 |
| KAT5 | 547.9378126 | -0.430229699 | 0.112800135 | 5.38E-05 | 0.00043359 |
| SIGMAR1 | 3854.188158 | -0.430525758 | 0.101178885 | 8.41E-06 | 8.65E-05 |
| EMC10 | 2432.523818 | -0.430566769 | 0.09441131 | 2.08E-06 | 2.53E-05 |
| MARK4 | 2529.502278 | -0.431072731 | 0.171318464 | 0.00397637 | 0.016643599 |
| FAAP20 | 1210.629178 | -0.431230397 | 0.155519151 | 0.00195601 | 0.009283039 |
| MCM7 | 17885.96083 | -0.431239403 | 0.100182391 | 6.73E-06 | 7.12E-05 |
| G3BP1 | 13379.46816 | -0.431487393 | 0.134646479 | 0.00050364 | 0.002947284 |
| NCKIPSD | 1297.323595 | -0.431515435 | 0.099885971 | 6.26E-06 | 6.68E-05 |
| NCLN | 3267.684494 | -0.431584457 | 0.137711854 | 0.00063687 | 0.003592038 |
| MRPS12 | 685.3055764 | -0.43189843 | 0.125679815 | 0.00022387 | 0.001474136 |
| ANXA2 | 6000.925249 | -0.431996821 | 0.179417133 | 0.00521023 | 0.02077224 |
| COX6B1 | 2275.650293 | -0.432323959 | 0.151323129 | 0.0015164 | 0.00744763 |
| ZNF496-DT | 514.6559398 | -0.432400761 | 0.130248517 | 0.00033771 | 0.002094291 |
| ZNF135 | 308.8352432 | -0.432451931 | 0.146476877 | 0.00113286 | 0.005857708 |
| NASP | 13612.89148 | -0.432488556 | 0.133482909 | 0.00044432 | 0.002649177 |
| ZNF837 | 99.8658058 | -0.432581463 | 0.2187469 | 0.01332097 | 0.044716349 |
| ADAMTS7 | 4096.698439 | -0.433019351 | 0.114021699 | 5.67E-05 | 0.000452198 |
| MRPS34 | 1626.385365 | -0.433043414 | 0.093190042 | 1.36E-06 | 1.76E-05 |
| MELTF-AS1 | 107.0090377 | -0.433092765 | 0.200396669 | 0.00916217 | 0.033118436 |
| DCLK3 | 477.4658576 | -0.433108973 | 0.203524592 | 0.00983041 | 0.035022973 |
| EMID1 | 502.8835806 | -0.433159008 | 0.136605851 | 0.00055882 | 0.003209048 |
| NDUFAF8 | 1428.554543 | -0.433561366 | 0.130535681 | 0.00033379 | 0.002075388 |
| BICD2 | 2571.552503 | -0.433588235 | 0.117897503 | 9.03E-05 | 0.000676234 |
| XXYLT1 | 2238.461907 | -0.433673463 | 0.099657376 | 5.38E-06 | 5.85E-05 |
| ANGPTL2 | 1145.816701 | -0.433951745 | 0.173870918 | 0.00411837 | 0.017129882 |
| MIR503HG | 467.3963205 | -0.434169031 | 0.121523788 | 0.00013415 | 0.000957576 |
| PUS1 | 2019.326894 | -0.434201033 | 0.100122797 | 5.74E-06 | 6.19E-05 |
| SAP30L | 1414.758646 | -0.434229795 | 0.095576599 | 2.22E-06 | 2.67E-05 |
| PROSER3 | 861.9695097 | -0.434793409 | 0.137436937 | 0.00056747 | 0.003248734 |
| MTND2P28 | 3370.835476 | -0.434871057 | 0.141375778 | 0.00075574 | 0.004152045 |
| GAPDHP1 | 458.4742018 | -0.435031005 | 0.151490409 | 0.00142854 | 0.007094032 |
| SPARC | 3293.752881 | -0.435059695 | 0.123907698 | 0.00016779 | 0.001159444 |
| MPDU1 | 1850.485159 | -0.435077827 | 0.115086044 | 6.00E-05 | 0.000476211 |
| FEN1 | 5707.799963 | -0.435093139 | 0.117888481 | 8.52E-05 | 0.00064356 |
| C1orf53 | 94.10858404 | -0.435105367 | 0.208613264 | 0.01058108 | 0.03723931 |
| H3-3A | 21352.06271 | -0.435107706 | 0.142332212 | 0.00080238 | 0.004349028 |
| ATP2B3 | 2672.64128 | -0.435112811 | 0.159627407 | 0.00218921 | 0.010192196 |
| SPG21 | 3379.589388 | -0.43530394 | 0.090827836 | 6.61E-07 | 9.26E-06 |
| COPRS | 927.8651828 | -0.435605283 | 0.110245149 | 3.01E-05 | 0.000265518 |
| CTBP1-AS | 218.3554453 | -0.435635522 | 0.14674116 | 0.00105754 | 0.005518211 |
| SYTL4 | 1823.214962 | -0.435700167 | 0.132493158 | 0.00036994 | 0.002268869 |
| PFKM | 5546.712218 | -0.43579302 | 0.118952662 | 9.43E-05 | 0.000703816 |
| ABCB10 | 2173.468763 | -0.435849631 | 0.098046725 | 3.47E-06 | 3.97E-05 |
| PSMB2 | 6070.935983 | -0.435863255 | 0.092085002 | 8.83E-07 | 1.19E-05 |
| ADGRA3 | 6964.26808 | -0.43611085 | 0.111895462 | 3.74E-05 | 0.000318566 |
| PCDHB9 | 635.9690211 | -0.436146385 | 0.158207919 | 0.00199099 | 0.0094143 |

| | | | | | |
|---|---|---|---|---|---|
| RRP1B | 6139.906192 | -0.436216984 | 0.120630679 | 0.00011273 | 0.000821137 |
| MTNR1A | 96.97218743 | -0.436346514 | 0.204834727 | 0.00956506 | 0.034261601 |
| HELLPAR | 665.0747934 | -0.436435228 | 0.146117582 | 0.00099477 | 0.005236445 |
| STMN1 | 29251.10159 | -0.436502048 | 0.156167281 | 0.00177776 | 0.008535553 |
| PA2G4 | 8174.439658 | -0.436527374 | 0.109402956 | 2.55E-05 | 0.00023035 |
| ARSB | 747.3753733 | -0.436683148 | 0.112256526 | 3.84E-05 | 0.000326184 |
| KCNV2 | 50.86376355 | -0.436756008 | 0.215699609 | 0.01180188 | 0.040604629 |
| POLR3K | 1071.219935 | -0.43677433 | 0.146228416 | 0.00099269 | 0.005227577 |
| LAPTM4B | 8811.751796 | -0.436849169 | 0.114929207 | 5.49E-05 | 0.00043999 |
| ADD2 | 5460.270123 | -0.437300423 | 0.133649517 | 0.00038817 | 0.002360962 |
| STX1A | 2227.046521 | -0.437331872 | 0.123337672 | 0.00014589 | 0.001029879 |
| BAIAP2-DT | 854.6457426 | -0.4374326 | 0.156162245 | 0.00173704 | 0.008371322 |
| CHD7 | 6734.108269 | -0.43743428 | 0.135092882 | 0.0004356 | 0.002607777 |
| TSKU | 1016.725034 | -0.437693443 | 0.173077605 | 0.00369167 | 0.01563335 |
| GPR161 | 4079.122974 | -0.437778505 | 0.112112224 | 3.60E-05 | 0.00030879 |
| TOMM40L | 896.1083064 | -0.437811571 | 0.108806625 | 2.20E-05 | 0.000202075 |
| MAPK12 | 2364.048166 | -0.438187083 | 0.101782732 | 6.49E-06 | 6.90E-05 |
| TSR3 | 1134.664906 | -0.438207769 | 0.12873487 | 0.00024339 | 0.001584892 |
| FBLN1 | 7606.973609 | -0.43830597 | 0.169709164 | 0.0031893 | 0.013848116 |
| TTC31 | 1794.129546 | -0.438690659 | 0.090557658 | 5.03E-07 | 7.33E-06 |
| RNU6-529P | 2.294851948 | -0.43875474 | 0.708742347 | 0.01455314 | 0.047990288 |
| USP31 | 1158.645339 | -0.439062034 | 0.11233679 | 3.52E-05 | 0.0003035 |
| UCK2 | 5334.382888 | -0.439203066 | 0.114589522 | 4.77E-05 | 0.000393801 |
| ELAC2 | 3250.015784 | -0.439207795 | 0.121423311 | 0.00011056 | 0.000807676 |
| CLMP | 1627.542362 | -0.439335282 | 0.12200385 | 0.00011744 | 0.000850696 |
| NELL2 | 13354.45343 | -0.439740979 | 0.137546442 | 0.00049395 | 0.002898037 |
| SERPING1 | 830.5103301 | -0.439770147 | 0.208843736 | 0.00982174 | 0.034998155 |
| LYN | 1600.599838 | -0.440478997 | 0.162964171 | 0.00226173 | 0.010465069 |
| SCRIB | 6591.068017 | -0.440603017 | 0.157854797 | 0.00175502 | 0.008444146 |
| POU3F1 | 48.81704227 | -0.440716943 | 0.21805982 | 0.01154492 | 0.039887394 |
| RAB11FIP4 | 2627.763441 | -0.440737184 | 0.123913422 | 0.00013756 | 0.000978826 |
| CDCA7 | 5140.869641 | -0.440745982 | 0.124815504 | 0.00015128 | 0.001061724 |
| BAG1 | 2800.496554 | -0.440850282 | 0.120862563 | 9.77E-05 | 0.000723544 |
| DPP7 | 3710.600965 | -0.441230657 | 0.090847906 | 4.67E-07 | 6.88E-06 |
| FZD8 | 330.6035119 | -0.441254321 | 0.182099825 | 0.00472447 | 0.019189029 |
| MICAL1 | 5852.208427 | -0.441309902 | 0.084268065 | 6.45E-08 | 1.19E-06 |
| CDK5 | 735.7695579 | -0.441360314 | 0.109830287 | 2.21E-05 | 0.000202616 |
| GPX4 | 3521.9276 | -0.441386098 | 0.124730561 | 0.00014662 | 0.001033977 |
| ATP6V0E2 | 920.3820301 | -0.441501966 | 0.146713203 | 0.00090078 | 0.004815077 |
| TONSL | 4279.839491 | -0.441599495 | 0.112951837 | 3.46E-05 | 0.000299327 |
| TLCD3A | 976.9402713 | -0.441662356 | 0.104481543 | 9.01E-06 | 9.18E-05 |
| C1orf220 | 102.321081 | -0.441863714 | 0.211956468 | 0.01009762 | 0.035795118 |
| TTI1 | 2106.377976 | -0.441873509 | 0.112862171 | 3.38E-05 | 0.00029315 |
| YBX1 | 55706.00885 | -0.441889304 | 0.120985213 | 9.53E-05 | 0.000709386 |
| GATD1 | 2785.808844 | -0.44207756 | 0.126521179 | 0.00017215 | 0.001184779 |
| KHSRP | 11134.67028 | -0.442102415 | 0.149338582 | 0.00104585 | 0.005460074 |
| EARS2 | 1895.911013 | -0.44226343 | 0.109159307 | 1.91E-05 | 0.000179932 |
| PYCR3 | 447.7135337 | -0.44253008 | 0.137282057 | 0.00044493 | 0.002652028 |
| PIGO | 1236.464325 | -0.442565357 | 0.116707527 | 5.52E-05 | 0.000442175 |
| NME3 | 773.4572186 | -0.442692871 | 0.111701011 | 2.76E-05 | 0.000246431 |

| | | | | | |
|---|---|---|---|---|---|
| FAM78A | 51.65647991 | -0.442923189 | 0.229942578 | 0.0135285 | 0.045243145 |
| COL9A3 | 274.4134048 | -0.443080745 | 0.160833896 | 0.00192046 | 0.009148118 |
| SIX2 | 848.0510339 | -0.443152435 | 0.164968528 | 0.00233125 | 0.01071281 |
| NELFB | 3068.33305 | -0.443233477 | 0.126405355 | 0.00016353 | 0.00113529 |
| SLC16A1-AS1 | 1429.483972 | -0.443557818 | 0.115750396 | 4.68E-05 | 0.000387831 |
| NXN | 2312.245924 | -0.443670556 | 0.126503878 | 0.00016271 | 0.001130775 |
| ANKDD1A | 865.7713837 | -0.443809209 | 0.103161929 | 6.40E-06 | 6.81E-05 |
| TUBA1A | 59146.4277 | -0.443850249 | 0.113099751 | 3.22E-05 | 0.000280974 |
| HNRNPKP4 | 232.1421736 | -0.443919585 | 0.136108841 | 0.00038805 | 0.002360962 |
| AIP | 1374.850725 | -0.444077125 | 0.121984457 | 9.87E-05 | 0.000730346 |
| YWHAZP4 | 161.9377644 | -0.444378521 | 0.234285196 | 0.01408132 | 0.046719623 |
| MAPK11 | 579.7191964 | -0.444449341 | 0.114826333 | 3.99E-05 | 0.000337582 |
| PPEF1 | 225.6135979 | -0.444621894 | 0.180319196 | 0.00415377 | 0.017249202 |
| HNRNPH2 | 2717.917205 | -0.444818699 | 0.142498875 | 0.00061639 | 0.003489906 |
| L3MBTL3 | 571.9635926 | -0.444825695 | 0.14491153 | 0.00072932 | 0.004026243 |
| GALK1 | 603.5784233 | -0.444949769 | 0.138014204 | 0.00043833 | 0.002620269 |
| FBXL19-AS1 | 614.3994741 | -0.444975105 | 0.112366581 | 2.76E-05 | 0.000246668 |
| C21orf58 | 1007.872981 | -0.445056249 | 0.093970919 | 8.33E-07 | 1.13E-05 |
| ST3GAL1-DT | 50.16468204 | -0.445075922 | 0.224869963 | 0.01208936 | 0.041392876 |
| ZCCHC18 | 298.9528068 | -0.445339986 | 0.127080548 | 0.00016291 | 0.001131763 |
| CACNG4 | 3431.366905 | -0.445560653 | 0.183421223 | 0.00452409 | 0.018499397 |
| MIR600HG | 1345.054891 | -0.445725372 | 0.153303444 | 0.00120439 | 0.006171701 |
| UQCR10 | 888.0130323 | -0.44603176 | 0.139542585 | 0.00047795 | 0.002817017 |
| GEMIN6 | 770.6129923 | -0.446067677 | 0.132457918 | 0.00026533 | 0.001707208 |
| POLD2 | 4721.214229 | -0.446121212 | 0.117416044 | 5.26E-05 | 0.000425671 |
| TMSB15B-AS1 | 141.9878549 | -0.446240419 | 0.166382083 | 0.00231437 | 0.010656658 |
| POLA2 | 778.5796628 | -0.446308083 | 0.092115113 | 4.83E-07 | 7.08E-06 |
| KLHDC3 | 4924.53951 | -0.446354906 | 0.087862634 | 1.45E-07 | 2.46E-06 |
| KAT2A | 2822.663576 | -0.446529279 | 0.091369133 | 3.91E-07 | 5.87E-06 |
| ZNF124 | 2026.098655 | -0.446626947 | 0.134701443 | 0.0003172 | 0.001985676 |
| PHC2 | 4273.327043 | -0.446636493 | 0.149941974 | 0.00096218 | 0.005095542 |
| EXOSC6 | 1413.446317 | -0.446653637 | 0.09876401 | 2.30E-06 | 2.76E-05 |
| CNTFR | 3697.872392 | -0.446683851 | 0.139700685 | 0.00047452 | 0.002800012 |
| SNRNP70 | 9899.581185 | -0.446890581 | 0.093014852 | 5.90E-07 | 8.41E-06 |
| TONSL-AS1 | 58.68716363 | -0.446927808 | 0.231617161 | 0.01303636 | 0.043925879 |
| POLR2L | 877.1332819 | -0.447471654 | 0.136048148 | 0.00034613 | 0.002139397 |
| QSOX1 | 2728.270704 | -0.447633689 | 0.129466123 | 0.00019082 | 0.001289512 |
| ALDH4A1 | 543.1230335 | -0.447706762 | 0.134862448 | 0.0003109 | 0.001952759 |
| SLC35B2 | 2436.56881 | -0.447793056 | 0.104050797 | 6.23E-06 | 6.64E-05 |
| AFAP1-AS1 | 40.16509218 | -0.44782789 | 0.229117961 | 0.01238151 | 0.042196432 |
| STMP1 | 3487.881006 | -0.447985576 | 0.123171895 | 9.79E-05 | 0.000724775 |
| HMGB1P6 | 2580.682225 | -0.448125323 | 0.155971563 | 0.00131685 | 0.006643668 |
| RNH1 | 2353.139281 | -0.448486509 | 0.10439758 | 6.42E-06 | 6.83E-05 |
| PTGES2 | 1904.247472 | -0.448561356 | 0.109786333 | 1.60E-05 | 0.000154053 |
| MPV17L | 255.2733657 | -0.448838055 | 0.142587529 | 0.00055268 | 0.00317997 |
| NOP16 | 1258.005479 | -0.44917568 | 0.099278814 | 2.25E-06 | 2.71E-05 |
| KCNN2 | 367.6816263 | -0.449263681 | 0.181978927 | 0.00399778 | 0.016723021 |
| C3orf70 | 171.8445542 | -0.449385979 | 0.194824696 | 0.00591357 | 0.023057161 |
| BYSL | 1961.150694 | -0.449449287 | 0.095399967 | 9.21E-07 | 1.24E-05 |
| ALDH5A1 | 1027.548498 | -0.449477196 | 0.124959938 | 0.00011298 | 0.000822396 |

| | | | | | |
|---|---|---|---|---|---|
| BOP1 | 4759.241802 | -0.449556217 | 0.135825929 | 0.00031848 | 0.001993058 |
| TSEN54 | 2594.202946 | -0.449630262 | 0.079400612 | 5.72E-09 | 1.31E-07 |
| HILPDA | 1070.893174 | -0.449661576 | 0.132473228 | 0.00023663 | 0.001548222 |
| TTLL12 | 2987.202398 | -0.449749955 | 0.151922218 | 0.00099989 | 0.005258738 |
| ZNF428 | 2131.247755 | -0.449837186 | 0.105935752 | 7.95E-06 | 8.25E-05 |
| CD276 | 4286.594251 | -0.449866686 | 0.146301995 | 0.00069641 | 0.003870546 |
| DPH6 | 160.7598736 | -0.450176212 | 0.155762048 | 0.00123575 | 0.006305697 |
| SEPTIN3 | 6691.753906 | -0.450222731 | 0.11025772 | 1.61E-05 | 0.00015416 |
| PTMAP4 | 39.5678861 | -0.450283591 | 0.244730222 | 0.01476277 | 0.048565269 |
| FLT4 | 88.18360923 | -0.450351947 | 0.185235725 | 0.00435886 | 0.017948647 |
| ADORA1 | 68.93342777 | -0.450364552 | 0.234432004 | 0.01288004 | 0.043556195 |
| TEDC1 | 736.312384 | -0.450384666 | 0.116699415 | 4.05E-05 | 0.000342126 |
| SLC27A5 | 84.933945 | -0.450564074 | 0.171638866 | 0.00263314 | 0.011838208 |
| SLC25A19 | 496.8612653 | -0.450648126 | 0.117974644 | 4.74E-05 | 0.000391303 |
| PLXND1 | 1910.313308 | -0.450751817 | 0.1595343 | 0.00149306 | 0.007361151 |
| RNFT2 | 833.3092812 | -0.450856897 | 0.100005317 | 2.41E-06 | 2.88E-05 |
| IKZF4 | 1300.024475 | -0.451228667 | 0.120301605 | 6.20E-05 | 0.000490486 |
| CDCA4 | 1854.899949 | -0.451250542 | 0.101483025 | 3.20E-06 | 3.71E-05 |
| DOC2A | 765.5045149 | -0.451300388 | 0.107046512 | 9.02E-06 | 9.18E-05 |
| UNK | 4228.131162 | -0.451760429 | 0.081855917 | 1.29E-08 | 2.78E-07 |
| GRAMD1A | 2406.938767 | -0.452080099 | 0.142817647 | 0.00051176 | 0.002986344 |
| FBXO46 | 704.2852065 | -0.452164343 | 0.111692468 | 1.84E-05 | 0.000174083 |
| TXNDC17 | 594.5334859 | -0.452166933 | 0.157132868 | 0.00126777 | 0.006435544 |
| CDHR2 | 242.2180922 | -0.452662406 | 0.169950514 | 0.00233912 | 0.010736962 |
| MAGEE1 | 179.2823938 | -0.452677953 | 0.201534314 | 0.00662734 | 0.025322298 |
| MRPL20 | 1301.981418 | -0.45299392 | 0.126924147 | 0.00012305 | 0.000886631 |
| TXNRD1 | 12900.55714 | -0.453011147 | 0.143947716 | 0.00054081 | 0.003123932 |
| RPUSD1 | 1018.95257 | -0.453357121 | 0.105510963 | 6.24E-06 | 6.65E-05 |
| GMEB2 | 1230.600237 | -0.453412396 | 0.127091082 | 0.00012341 | 0.000888577 |
| BACE2 | 211.4267151 | -0.453578722 | 0.159968015 | 0.00142463 | 0.00707806 |
| CHDH | 347.9065259 | -0.453644046 | 0.12288674 | 7.71E-05 | 0.000590484 |
| COL16A1 | 538.0415733 | -0.453663708 | 0.149288746 | 0.00076413 | 0.004187393 |
| PPIA | 36950.41264 | -0.453916928 | 0.151857345 | 0.0008921 | 0.004773672 |
| TMEM60 | 363.9650214 | -0.454033672 | 0.15855774 | 0.00130718 | 0.006599768 |
| CGREF1 | 562.3042897 | -0.454091784 | 0.107231531 | 8.18E-06 | 8.47E-05 |
| C16orf92 | 191.2267809 | -0.454180286 | 0.134569059 | 0.00024699 | 0.001605814 |
| RBM8A | 7162.997976 | -0.454400701 | 0.09313382 | 3.91E-07 | 5.88E-06 |
| SMIM10L1 | 1678.096227 | -0.455113439 | 0.112483053 | 1.83E-05 | 0.000173265 |
| IMPDH1P8 | 177.1923494 | -0.455125129 | 0.202857336 | 0.00654903 | 0.025083657 |
| TMC4 | 61.33530281 | -0.455184504 | 0.22041652 | 0.00950388 | 0.034089965 |
| KPTN | 541.1582391 | -0.455305364 | 0.10248812 | 3.19E-06 | 3.70E-05 |
| PWP2 | 432.4606485 | -0.455345818 | 0.155364728 | 0.00105856 | 0.005522142 |
| CENPS | 55.33374232 | -0.455837949 | 0.204983444 | 0.00680543 | 0.025849733 |
| TSACC | 82.18343056 | -0.455885789 | 0.179147015 | 0.00314799 | 0.01372382 |
| KLF13 | 4659.731939 | -0.455917522 | 0.143209391 | 0.00047122 | 0.002785011 |
| CNNM1 | 1115.510478 | -0.455935351 | 0.124805866 | 8.82E-05 | 0.000661986 |
| REPIN1 | 4852.405431 | -0.456162278 | 0.137264111 | 0.00029276 | 0.001851292 |
| SLC36A1 | 660.5923531 | -0.456312667 | 0.135729182 | 0.00025556 | 0.001652602 |
| ACVR2B | 6360.237798 | -0.45638198 | 0.135121216 | 0.00024178 | 0.001576749 |
| SPNS2 | 679.6355057 | -0.456870424 | 0.125956425 | 9.68E-05 | 0.000717852 |

| | | | | | |
|---|---|---|---|---|---|
| PKIB | 723.9930549 | -0.457032168 | 0.168507136 | 0.00198637 | 0.009398906 |
| PRMT6 | 3131.195147 | -0.457038551 | 0.112270391 | 1.63E-05 | 0.000156604 |
| RAB35-AS1 | 105.4477956 | -0.457114606 | 0.171451334 | 0.00225624 | 0.01044696 |
| PDGFRB | 635.9153736 | -0.457257107 | 0.133242351 | 0.00019842 | 0.001331667 |
| PTBP1 | 17851.66382 | -0.457347455 | 0.133801542 | 0.00020825 | 0.001384982 |
| FASN | 53610.77286 | -0.457396842 | 0.180359126 | 0.00318771 | 0.01384417 |
| LSM7 | 2576.3098 | -0.457441878 | 0.124651472 | 8.21E-05 | 0.000623881 |
| OAF | 391.1808064 | -0.457504616 | 0.124956742 | 8.47E-05 | 0.000640401 |
| CPLX2 | 10073.56853 | -0.457542214 | 0.172339424 | 0.00232047 | 0.010677597 |
| XKR8 | 354.9495799 | -0.457747129 | 0.125863408 | 9.29E-05 | 0.00069399 |
| SNHG19 | 200.8571154 | -0.457965636 | 0.179247248 | 0.00302309 | 0.013305927 |
| RNF220 | 1684.788151 | -0.458066937 | 0.123745248 | 7.23E-05 | 0.000559479 |
| LLGL1 | 3423.549832 | -0.458167525 | 0.119427858 | 4.26E-05 | 0.000357776 |
| MT-ATP6 | 83811.19218 | -0.458178525 | 0.127533542 | 0.00010947 | 0.000800839 |
| KIF20A | 6040.345269 | -0.458249621 | 0.143770822 | 0.00045897 | 0.002721534 |
| PCDHGA8 | 71.69079399 | -0.458434537 | 0.197550426 | 0.00537101 | 0.021285288 |
| CCDC88C | 1196.79394 | -0.458499313 | 0.157550234 | 0.00110536 | 0.005734259 |
| SLC9A3-AS1 | 1727.170802 | -0.458638727 | 0.14302617 | 0.00042934 | 0.002573304 |
| PYGL | 68.54565705 | -0.458668531 | 0.19522734 | 0.00501344 | 0.02013601 |
| PATZ1 | 3760.755293 | -0.458910671 | 0.114611592 | 2.14E-05 | 0.000198065 |
| INPP5E | 1514.768825 | -0.459020505 | 0.126734741 | 9.76E-05 | 0.000723173 |
| LINC01003 | 183.4251661 | -0.4592057 | 0.16313175 | 0.0014604 | 0.007226087 |
| SNRPD1 | 4745.535195 | -0.459205798 | 0.151330635 | 0.00074935 | 0.004123521 |
| TMEM249 | 40.32610285 | -0.459321571 | 0.257198928 | 0.01489954 | 0.048921315 |
| MTCH1P2 | 57.9573218 | -0.459391508 | 0.231305714 | 0.01068153 | 0.0375094 |
| HNRNPA1L3 | 676.8997432 | -0.459470803 | 0.17034437 | 0.00203847 | 0.00960062 |
| GPC3 | 122.059352 | -0.459531076 | 0.181673945 | 0.00319621 | 0.013863482 |
| COL25A1 | 1785.824196 | -0.45961797 | 0.135345946 | 0.00022246 | 0.001468199 |
| EIF4A1 | 978.9950557 | -0.460044738 | 0.112180337 | 1.42E-05 | 0.000138107 |
| PCED1A | 2345.544586 | -0.460240951 | 0.100982476 | 1.82E-06 | 2.26E-05 |
| PAK4 | 1550.152863 | -0.460252128 | 0.188737129 | 0.0039995 | 0.016726814 |
| LIFR | 2878.904366 | -0.460730412 | 0.144896634 | 0.00046388 | 0.002747461 |
| PRDX2 | 8046.986583 | -0.46088107 | 0.128763 | 0.00011332 | 0.000824037 |
| MRPL14 | 2404.182461 | -0.461190292 | 0.102700673 | 2.48E-06 | 2.96E-05 |
| RASL10B | 1357.675004 | -0.461379546 | 0.122053484 | 5.23E-05 | 0.000423563 |
| ITPRID2 | 7159.261734 | -0.461382659 | 0.137118737 | 0.00024567 | 0.001598187 |
| MCRS1 | 2965.95744 | -0.461499077 | 0.122750629 | 5.67E-05 | 0.000452198 |
| AP5S1 | 299.7022536 | -0.461625815 | 0.120305336 | 4.17E-05 | 0.000350849 |
| RFC3 | 4420.681685 | -0.461672333 | 0.142818344 | 0.00038656 | 0.002354118 |
| KIRREL3 | 55.98827196 | -0.461791163 | 0.204683545 | 0.00606691 | 0.023548483 |
| ZNF618 | 3901.946204 | -0.462092627 | 0.15758018 | 0.00101026 | 0.005307877 |
| KDM1A | 9637.035692 | -0.462825438 | 0.115749171 | 2.15E-05 | 0.000198336 |
| ZFP64 | 1209.070462 | -0.462890764 | 0.153509983 | 0.00077792 | 0.004248926 |
| CDC25B | 3930.490109 | -0.462987748 | 0.138873608 | 0.00027116 | 0.001736607 |
| EPHB6 | 172.7674534 | -0.463099423 | 0.1412441 | 0.00032774 | 0.00204106 |
| NDUFB10 | 1916.978572 | -0.46319437 | 0.124153002 | 6.28E-05 | 0.000495412 |
| CBX1 | 22469.74751 | -0.463298035 | 0.113846749 | 1.59E-05 | 0.000152906 |
| RELL1 | 297.9870921 | -0.4634388 | 0.164169907 | 0.00138866 | 0.006932836 |
| SRM | 2382.731828 | -0.463540967 | 0.199727534 | 0.0051737 | 0.020638591 |
| ABHD17A | 3151.62161 | -0.463568179 | 0.11495276 | 1.85E-05 | 0.000174745 |

| | | | | | |
|---|---|---|---|---|---|
| PDK2 | 644.3732814 | -0.463585978 | 0.144901463 | 0.00042724 | 0.002564839 |
| RIIAD1 | 141.2449359 | -0.463801393 | 0.159435724 | 0.00107294 | 0.005580158 |
| MIR4458HG | 496.8455295 | -0.464078399 | 0.153820742 | 0.00076828 | 0.004202767 |
| USP46 | 2970.129102 | -0.464121101 | 0.140121952 | 0.0002903 | 0.001838514 |
| SNRPD2 | 6256.126721 | -0.464245277 | 0.142005513 | 0.00033625 | 0.002086479 |
| HDHD5 | 2918.960655 | -0.464492645 | 0.081758317 | 4.78E-09 | 1.12E-07 |
| ANAPC15 | 235.923878 | -0.464538893 | 0.148603051 | 0.00054074 | 0.003123932 |
| GATA3-AS1 | 581.5599181 | -0.464542635 | 0.132126964 | 0.00014031 | 0.000995694 |
| HSPBP1 | 1784.677601 | -0.464965264 | 0.106626061 | 4.41E-06 | 4.90E-05 |
| GALNTL6 | 112.7870141 | -0.465074073 | 0.183412442 | 0.00302801 | 0.013319357 |
| SMAD3 | 1666.578809 | -0.465220962 | 0.132437212 | 0.00014136 | 0.0010024 |
| RPL7L1 | 11618.74308 | -0.465229127 | 0.11165769 | 1.04E-05 | 0.000104613 |
| AURKAIP1 | 1118.688807 | -0.465378067 | 0.154092071 | 0.00075466 | 0.004149394 |
| BLOC1S4 | 503.6926338 | -0.465399313 | 0.105111743 | 3.24E-06 | 3.74E-05 |
| LINC01963 | 3427.377391 | -0.465557087 | 0.150685226 | 0.00060457 | 0.003431505 |
| RTN1 | 5829.833044 | -0.465577824 | 0.111956026 | 1.07E-05 | 0.000107607 |
| DOCK7-DT | 51.11303205 | -0.465764212 | 0.252709062 | 0.01293659 | 0.043675587 |
| GOLM1 | 5723.518279 | -0.465890689 | 0.11492492 | 1.67E-05 | 0.000160052 |
| HIGD2A | 1189.039347 | -0.466017662 | 0.121878242 | 4.29E-05 | 0.000359819 |
| HNRNPA1 | 98602.31413 | -0.466051024 | 0.164670489 | 0.00133666 | 0.006728772 |
| CCDC78 | 387.9478815 | -0.466632625 | 0.150604394 | 0.0005838 | 0.003330155 |
| RANBP1 | 7531.457089 | -0.466954585 | 0.130620414 | 0.00011126 | 0.000811644 |
| RND3 | 1962.344321 | -0.467028055 | 0.149903941 | 0.00055104 | 0.003172025 |
| NDST1 | 5350.717956 | -0.467080874 | 0.158320892 | 0.00092721 | 0.004940931 |
| ENTPD6 | 1771.570454 | -0.467336276 | 0.089919732 | 7.03E-08 | 1.29E-06 |
| NACC1 | 2536.377994 | -0.467593855 | 0.156661302 | 0.00083101 | 0.004481767 |
| TIMM13 | 2558.442597 | -0.467699093 | 0.12015337 | 3.23E-05 | 0.000281381 |
| ASCL1 | 2227.751535 | -0.467762555 | 0.199882852 | 0.00479763 | 0.019424767 |
| HMGB1 | 33220.30393 | -0.467953578 | 0.14815239 | 0.00047606 | 0.002807985 |
| VWCE | 1987.927238 | -0.468091507 | 0.086499435 | 2.18E-08 | 4.45E-07 |
| SMARCC1 | 15429.38526 | -0.468210339 | 0.110782493 | 7.87E-06 | 8.19E-05 |
| UQCRH | 4190.867927 | -0.468610982 | 0.148820784 | 0.00048957 | 0.00287806 |
| HNRNPR | 11239.43778 | -0.468690723 | 0.12267241 | 4.28E-05 | 0.000358786 |
| TMEM44 | 669.8623939 | -0.4688323 | 0.109066945 | 5.70E-06 | 6.16E-05 |
| ITGB1BP2 | 120.9194944 | -0.468970588 | 0.191848251 | 0.00370659 | 0.015683614 |
| SLC26A10P | 294.8513195 | -0.46901341 | 0.158230021 | 0.00087763 | 0.004706001 |
| FAM110A | 559.1638829 | -0.469573595 | 0.123698791 | 4.69E-05 | 0.000388243 |
| GRIK3 | 6880.63316 | -0.469641198 | 0.13406543 | 0.00014272 | 0.001010314 |
| LCMT2 | 1627.815871 | -0.469726024 | 0.128019767 | 7.67E-05 | 0.000588605 |
| MTHFD1 | 7033.041249 | -0.469787318 | 0.125019178 | 5.45E-05 | 0.000437495 |
| CMTM1 | 151.3809579 | -0.469863702 | 0.181695311 | 0.00256937 | 0.01160266 |
| SYT7 | 1356.635247 | -0.470180281 | 0.184623167 | 0.00284037 | 0.012623586 |
| LMNB2 | 11837.91342 | -0.470201125 | 0.126886613 | 6.65E-05 | 0.000521149 |
| PHOSPHO2 | 153.7756615 | -0.470228681 | 0.161719143 | 0.00103378 | 0.005409299 |
| LRP3 | 3515.590563 | -0.470672339 | 0.129271389 | 8.50E-05 | 0.000641624 |
| TBC1D9B | 3915.559652 | -0.470795967 | 0.097943743 | 5.16E-07 | 7.50E-06 |
| TTF2 | 6545.094188 | -0.470877569 | 0.128997674 | 8.19E-05 | 0.000622952 |
| DUX4L50 | 208.5195706 | -0.470912505 | 0.141429504 | 0.00026244 | 0.001692329 |
| TBC1D1 | 1684.323096 | -0.4710627 | 0.139744069 | 0.00022709 | 0.001492464 |
| ARSA | 437.195026 | -0.472101337 | 0.136854793 | 0.00017065 | 0.00117641 |

| | | | | | |
|---|---|---|---|---|---|
| TEDC2 | 1542.828094 | -0.472256305 | 0.102405275 | 1.32E-06 | 1.72E-05 |
| MCM5 | 5504.942299 | -0.472365362 | 0.127169861 | 6.35E-05 | 0.000500408 |
| DELE1 | 3109.381093 | -0.4724141 | 0.097975825 | 4.75E-07 | 6.99E-06 |
| RPL7AP70 | 86.85541588 | -0.472567266 | 0.210509723 | 0.00573724 | 0.022476791 |
| PHF19 | 2835.159251 | -0.47268991 | 0.113728883 | 1.04E-05 | 0.000104915 |
| ENO1P2 | 42.05040462 | -0.472731047 | 0.268532058 | 0.01380599 | 0.045976745 |
| EMILIN3 | 324.2726672 | -0.472796695 | 0.132544236 | 0.00011069 | 0.000808352 |
| PHB1 | 5964.183073 | -0.472945435 | 0.12343448 | 4.00E-05 | 0.000337831 |
| TESC | 2346.160465 | -0.473284133 | 0.179274264 | 0.00217318 | 0.01013822 |
| PGAM1 | 7761.303071 | -0.473358045 | 0.12321414 | 3.83E-05 | 0.00032574 |
| CHST10 | 1262.084272 | -0.473421225 | 0.107093146 | 3.21E-06 | 3.71E-05 |
| RNF44 | 5542.243576 | -0.473480353 | 0.114352867 | 1.11E-05 | 0.000111131 |
| EIF5AL1 | 283.369088 | -0.473482892 | 0.159758022 | 0.00085371 | 0.00459574 |
| SNORD83A | 47.93858579 | -0.473915804 | 0.220499005 | 0.00694868 | 0.026302838 |
| IGDCC4 | 1117.329528 | -0.473973001 | 0.125389851 | 4.87E-05 | 0.000400448 |
| CISD3 | 347.002818 | -0.474030199 | 0.117949828 | 1.85E-05 | 0.000174727 |
| SSBP4 | 2674.317876 | -0.474045866 | 0.141145592 | 0.00023286 | 0.001527455 |
| ENO1 | 32132.27852 | -0.474174607 | 0.120800713 | 2.72E-05 | 0.000243573 |
| PLEKHO1 | 3552.283929 | -0.474365404 | 0.109171413 | 4.49E-06 | 4.98E-05 |
| NDUFB3 | 856.9724176 | -0.47498015 | 0.165853052 | 0.00114182 | 0.005890735 |
| FAM27E3 | 118.518553 | -0.475147519 | 0.202795409 | 0.00451316 | 0.01846569 |
| TMOD2 | 9991.296106 | -0.475195413 | 0.13137691 | 9.06E-05 | 0.000677639 |
| NOL6 | 3495.600459 | -0.475288923 | 0.142345859 | 0.00024741 | 0.001607983 |
| H3P16 | 348.7536111 | -0.475331441 | 0.183772544 | 0.00246959 | 0.011220642 |
| KCNH1 | 1000.417011 | -0.47546667 | 0.159398759 | 0.00079462 | 0.004319496 |
| MT-ND3 | 13228.80433 | -0.475787866 | 0.116589298 | 1.41E-05 | 0.000137956 |
| ANAPC11 | 2446.345099 | -0.476023459 | 0.120812057 | 2.53E-05 | 0.000229342 |
| CD81 | 4466.159905 | -0.476108777 | 0.148162172 | 0.00037739 | 0.002308879 |
| AP1S1 | 1118.422234 | -0.476256733 | 0.122072145 | 2.96E-05 | 0.000262055 |
| DHFR | 9294.4492 | -0.476280697 | 0.122546948 | 3.15E-05 | 0.000275716 |
| IGF2BP1 | 19829.70899 | -0.476334136 | 0.15433996 | 0.000571 | 0.0032635 |
| SPNS1 | 106.2341691 | -0.476347568 | 0.180509514 | 0.00213361 | 0.009983052 |
| LINC00526 | 103.7745713 | -0.476508274 | 0.18904928 | 0.00290381 | 0.012855549 |
| PCCA-DT | 215.2230594 | -0.476579661 | 0.143198672 | 0.00025487 | 0.001650211 |
| HNRNPLP2 | 107.9001776 | -0.47663972 | 0.177147462 | 0.00185004 | 0.008853683 |
| DLC1 | 2819.402807 | -0.477010484 | 0.151713283 | 0.00047158 | 0.002785862 |
| NR1H2 | 607.7671541 | -0.477085772 | 0.09905056 | 4.75E-07 | 6.99E-06 |
| AP4M1 | 610.3997543 | -0.477201778 | 0.101023207 | 7.50E-07 | 1.03E-05 |
| GNPDA1 | 2893.286907 | -0.477234646 | 0.110351 | 4.85E-06 | 5.34E-05 |
| ADSL | 803.8619639 | -0.477361735 | 0.143254564 | 0.00025001 | 0.001620808 |
| BMP7 | 373.0656891 | -0.477446445 | 0.164773785 | 0.00101619 | 0.005329497 |
| MRPL30 | 2654.551198 | -0.47752636 | 0.117400222 | 1.48E-05 | 0.000143657 |
| HMGA1 | 16037.88364 | -0.477797654 | 0.121399683 | 2.55E-05 | 0.000230777 |
| HMGB3 | 6091.241649 | -0.47780145 | 0.134615119 | 0.00011471 | 0.000832206 |
| PRXL2A | 5572.067652 | -0.477870599 | 0.102642127 | 1.04E-06 | 1.38E-05 |
| EEF1D | 5669.566006 | -0.47794363 | 0.085411427 | 7.31E-09 | 1.64E-07 |
| BID | 2174.878997 | -0.478377451 | 0.132946046 | 9.53E-05 | 0.000709386 |
| PTPRVP | 111.3466352 | -0.478398223 | 0.187099856 | 0.0026099 | 0.011749139 |
| FERMT1 | 85.34657447 | -0.478651781 | 0.227680585 | 0.00735391 | 0.027550703 |
| LINC02827 | 32.93840557 | -0.478962881 | 0.282424794 | 0.01428495 | 0.047250392 |

| | | | | | |
|---|---|---|---|---|---|
| ESRRG | 2054.851321 | -0.479060353 | 0.15260139 | 0.00047288 | 0.002791276 |
| TAS2R4 | 88.79971042 | -0.479061847 | 0.20293224 | 0.00420619 | 0.017431369 |
| TUB | 11770.00056 | -0.479181999 | 0.098998451 | 4.18E-07 | 6.24E-06 |
| RAI14 | 11603.18531 | -0.479190508 | 0.13196337 | 8.38E-05 | 0.000634141 |
| PIP4K2A | 1064.892757 | -0.479424086 | 0.121111451 | 2.30E-05 | 0.000210826 |
| RPL22L1 | 3038.18738 | -0.479459414 | 0.205301493 | 0.00444617 | 0.018241222 |
| MEIS1 | 11619.72516 | -0.479510294 | 0.118495047 | 1.60E-05 | 0.000153669 |
| ELAVL3 | 9987.927428 | -0.479514321 | 0.136069601 | 0.00012461 | 0.000895953 |
| SEMA6A-AS1 | 65.86720003 | -0.479515136 | 0.206119941 | 0.00453678 | 0.018540192 |
| WDR86 | 671.7533351 | -0.479522496 | 0.144508053 | 0.00025885 | 0.001671238 |
| GEMIN4 | 2317.540017 | -0.479684238 | 0.137895648 | 0.00014685 | 0.001034892 |
| GTF2H5 | 772.1105338 | -0.479733947 | 0.15144523 | 0.00042908 | 0.002572465 |
| RECQL4 | 4891.247548 | -0.479939329 | 0.096961607 | 2.39E-07 | 3.81E-06 |
| RTN4RL1 | 70.74439625 | -0.480025051 | 0.224235667 | 0.00673805 | 0.025640501 |
| TBC1D14 | 2101.306446 | -0.48003016 | 0.122107836 | 2.57E-05 | 0.000231863 |
| MIER2 | 724.892687 | -0.480091782 | 0.162717459 | 0.00085129 | 0.004583928 |
| BIRC5 | 9608.685315 | -0.480201531 | 0.11758143 | 1.36E-05 | 0.000133464 |
| MT-CO3 | 129481.7638 | -0.480388433 | 0.125210837 | 3.75E-05 | 0.000319784 |
| SNRPB | 5976.688284 | -0.480560065 | 0.100621701 | 5.70E-07 | 8.19E-06 |
| FAAP100 | 3457.290972 | -0.480650826 | 0.127767452 | 5.03E-05 | 0.000410357 |
| PROCA1 | 404.0926541 | -0.481019824 | 0.10912706 | 3.26E-06 | 3.76E-05 |
| NUDT15 | 2672.682444 | -0.481129821 | 0.146539916 | 0.00028893 | 0.001831527 |
| GRWD1 | 2018.369701 | -0.481148545 | 0.146974796 | 0.00029847 | 0.001880997 |
| TBCD | 11329.74399 | -0.481261679 | 0.13104125 | 7.08E-05 | 0.000549509 |
| RAB3D | 308.246272 | -0.481365126 | 0.165192518 | 0.00094251 | 0.005006932 |
| ZC3H4 | 2428.012941 | -0.481414174 | 0.150069299 | 0.00037163 | 0.002277181 |
| SUMO3 | 4903.192338 | -0.48143174 | 0.130611823 | 6.72E-05 | 0.000525365 |
| DDX54 | 2937.446676 | -0.481939094 | 0.137521629 | 0.00013187 | 0.000942871 |
| OPA3 | 1445.751251 | -0.48197211 | 0.097872428 | 2.69E-07 | 4.22E-06 |
| SLC10A4 | 1609.77492 | -0.482125459 | 0.116423139 | 1.05E-05 | 0.000105882 |
| MFSD4A | 42.15080644 | -0.482703149 | 0.248764068 | 0.00957721 | 0.034293192 |
| FGD6 | 836.1931066 | -0.48282716 | 0.131225505 | 6.83E-05 | 0.000533272 |
| ROR2 | 454.5800772 | -0.482943697 | 0.185801037 | 0.00226221 | 0.010465069 |
| ING2-DT | 26.22698722 | -0.48323816 | 0.274277076 | 0.0125932 | 0.042797304 |
| NPR1 | 134.3775793 | -0.483383392 | 0.22453003 | 0.00640073 | 0.024630368 |
| GSR | 2450.359655 | -0.484072042 | 0.114516925 | 7.18E-06 | 7.54E-05 |
| GLB1L | 145.7588305 | -0.484086581 | 0.148913282 | 0.00031651 | 0.001983311 |
| TRMT2A | 1207.126745 | -0.484248845 | 0.088295248 | 1.33E-08 | 2.85E-07 |
| JPT2 | 5995.252389 | -0.484254565 | 0.124672545 | 3.03E-05 | 0.000267173 |
| GABPB1-IT1 | 1590.030195 | -0.484672596 | 0.126473925 | 3.72E-05 | 0.00031765 |
| MOSPD3 | 422.3906245 | -0.484770922 | 0.143250448 | 0.00019924 | 0.001335434 |
| LYPLA2 | 1151.108975 | -0.484852928 | 0.09834385 | 2.58E-07 | 4.05E-06 |
| METTL13 | 4044.484019 | -0.485496983 | 0.128600453 | 4.64E-05 | 0.000384326 |
| AVPR1A | 61.53678005 | -0.485605555 | 0.233298053 | 0.00723913 | 0.027204933 |
| CLSTN2 | 2149.986985 | -0.485694488 | 0.121909738 | 2.00E-05 | 0.000187194 |
| DAZAP2 | 4501.101196 | -0.485866679 | 0.126296035 | 3.48E-05 | 0.000301193 |
| MTATP6P27 | 32.24681405 | -0.485913191 | 0.265170055 | 0.01109744 | 0.038653065 |
| SEMA6C | 992.8968776 | -0.48598723 | 0.124632705 | 2.82E-05 | 0.000250728 |
| PTOV1-AS1 | 110.9685725 | -0.486264542 | 0.163587797 | 0.00076362 | 0.004186345 |
| VDAC1 | 7153.489481 | -0.486276678 | 0.128268026 | 4.34E-05 | 0.000363088 |

| | | | | | |
|---|---|---|---|---|---|
| FAM171A2 | 3309.72849 | -0.486298042 | 0.131151073 | 5.99E-05 | 0.000475594 |
| GUSBP5 | 36.29386897 | -0.486326744 | 0.266726021 | 0.01121272 | 0.038981958 |
| CHRNA5 | 1926.316763 | -0.486344371 | 0.145211816 | 0.0002228 | 0.001468927 |
| UCP2 | 1717.637614 | -0.486346546 | 0.155612579 | 0.0004718 | 0.002786336 |
| RPL13P8 | 340.5007015 | -0.486412369 | 0.153744182 | 0.00041622 | 0.002507088 |
| MAD2L1 | 8341.308139 | -0.486421335 | 0.152868923 | 0.0003921 | 0.0023813 |
| TXNL4A | 3249.098683 | -0.486452999 | 0.10222642 | 6.01E-07 | 8.54E-06 |
| EIF4EBP2 | 9519.062152 | -0.486718297 | 0.092063813 | 3.92E-08 | 7.64E-07 |
| MHENCR | 327.6581562 | -0.486906336 | 0.186250194 | 0.00211305 | 0.009898101 |
| SPTBN2 | 16271.76359 | -0.487059018 | 0.173205946 | 0.00122306 | 0.006244022 |
| GEMIN5 | 3316.704633 | -0.48707056 | 0.134578064 | 8.36E-05 | 0.000632574 |
| HSPE1 | 2989.97488 | -0.487295876 | 0.158868907 | 0.00056454 | 0.003235569 |
| MRPL57 | 1157.663559 | -0.487321904 | 0.124645782 | 2.69E-05 | 0.000240508 |
| PVALEF | 77.232816 | -0.487432975 | 0.216033704 | 0.00498798 | 0.020065108 |
| PHLDB2 | 2159.946073 | -0.487515846 | 0.145065541 | 0.0002125 | 0.001410999 |
| BAZ1A-AS1 | 109.390563 | -0.48757565 | 0.178156742 | 0.00150802 | 0.007413594 |
| HSPA12A | 1284.511065 | -0.488102619 | 0.097889926 | 1.90E-07 | 3.13E-06 |
| OR7E12P | 293.063054 | -0.488160044 | 0.152637476 | 0.00036745 | 0.002254273 |
| CYBC1 | 3880.760745 | -0.488374258 | 0.083944241 | 1.89E-09 | 4.77E-08 |
| TRIM45 | 771.480111 | -0.488475724 | 0.101487815 | 4.54E-07 | 6.73E-06 |
| PFN1 | 7454.745637 | -0.488720266 | 0.142619081 | 0.00016713 | 0.001156184 |
| TMEM271 | 842.2893318 | -0.48883098 | 0.14336688 | 0.00017744 | 0.001215905 |
| ACAP3 | 5513.660038 | -0.488857402 | 0.11664659 | 8.18E-06 | 8.46E-05 |
| EMC3-AS1 | 98.25213922 | -0.488922764 | 0.187400391 | 0.00211139 | 0.009892583 |
| PTPRO | 1800.294501 | -0.488935563 | 0.127383379 | 3.54E-05 | 0.000304867 |
| PRRT3 | 227.3842907 | -0.488956568 | 0.163579791 | 0.00071254 | 0.003946313 |
| PCNA | 11105.0555 | -0.489160298 | 0.157404397 | 0.00049022 | 0.002881075 |
| PLSCR4 | 72.78589701 | -0.489241568 | 0.210261726 | 0.00420555 | 0.017431369 |
| LIG1 | 3587.170898 | -0.489598489 | 0.099672622 | 2.75E-07 | 4.30E-06 |
| ASF1B | 1270.512494 | -0.489669631 | 0.130602331 | 5.00E-05 | 0.000408463 |
| NAGPA | 387.1632759 | -0.48975403 | 0.149570122 | 0.0002817 | 0.00179181 |
| MIR3176 | 94.17501272 | -0.489793343 | 0.189075879 | 0.00220214 | 0.010243093 |
| ECI1-AS1 | 144.959686 | -0.489813208 | 0.15754159 | 0.00048584 | 0.002859401 |
| ZNF887P | 188.887219 | -0.489983827 | 0.156622746 | 0.000456 | 0.002707054 |
| DIRAS1 | 1656.864473 | -0.4900306 | 0.125280106 | 2.62E-05 | 0.000235898 |
| ANKS6 | 1636.519173 | -0.490099676 | 0.09195599 | 3.04E-08 | 6.06E-07 |
| YBX1P1 | 604.6703163 | -0.490249302 | 0.165067895 | 0.00074889 | 0.004122085 |
| ESPL1 | 5589.476461 | -0.490479207 | 0.133019611 | 6.32E-05 | 0.00049864 |
| LENG8 | 12588.80748 | -0.490666646 | 0.109685525 | 2.28E-06 | 2.74E-05 |
| TMEM143 | 307.6684182 | -0.490924637 | 0.13474266 | 7.45E-05 | 0.000573771 |
| TUBG1 | 3306.693467 | -0.491007817 | 0.126827524 | 3.06E-05 | 0.000269671 |
| MXD3 | 1497.943162 | -0.491752822 | 0.154568359 | 0.00037915 | 0.002315651 |
| POLR1A | 8051.062365 | -0.49191237 | 0.156677913 | 0.00043412 | 0.002599684 |
| CDK10 | 1317.674054 | -0.492382701 | 0.108331445 | 1.62E-06 | 2.04E-05 |
| FAM162B | 209.2216332 | -0.49257913 | 0.184803776 | 0.0017688 | 0.008498517 |
| CENPX | 1498.926945 | -0.492695481 | 0.113405472 | 4.06E-06 | 4.56E-05 |
| NEK8 | 195.3205604 | -0.492850391 | 0.142359073 | 0.00014352 | 0.001014882 |
| SLC4A11 | 150.2278328 | -0.492969348 | 0.173845394 | 0.00109578 | 0.005685977 |
| DNAAF5 | 2492.443364 | -0.49338155 | 0.128316007 | 3.36E-05 | 0.000291378 |
| TSPAN15 | 216.9533607 | -0.49345805 | 0.144604951 | 0.00017054 | 0.00117641 |

| | | | | | |
|---|---|---|---|---|---|
| RN7SKP87 | 34.66570744 | -0.493603223 | 0.304232922 | 0.0137679 | 0.045876922 |
| SLC35E2B | 3431.571723 | -0.493723148 | 0.113105066 | 3.68E-06 | 4.18E-05 |
| LRP8 | 6286.454706 | -0.49382174 | 0.097847585 | 1.35E-07 | 2.30E-06 |
| CBLN1 | 2030.023086 | -0.493982076 | 0.111066738 | 2.52E-06 | 3.00E-05 |
| PGBD2 | 396.0016813 | -0.494177602 | 0.188227552 | 0.00194497 | 0.009239156 |
| DNPH1 | 730.2708991 | -0.494381017 | 0.145645533 | 0.00018064 | 0.001234551 |
| PPP1R14BP3 | 282.8128042 | -0.494632852 | 0.133824521 | 5.96E-05 | 0.000473117 |
| LHX2 | 29.47863271 | -0.494885218 | 0.317353129 | 0.01459557 | 0.048107585 |
| MBNL3 | 1739.003528 | -0.494996842 | 0.120560274 | 1.14E-05 | 0.000113558 |
| TMTC4 | 2481.652729 | -0.495197446 | 0.121064539 | 1.21E-05 | 0.000120278 |
| DAG1 | 2647.185243 | -0.495243489 | 0.126905185 | 2.63E-05 | 0.000236752 |
| PABIR2 | 2994.200634 | -0.495312527 | 0.115341347 | 5.00E-06 | 5.48E-05 |
| E2F2 | 1320.882847 | -0.495389734 | 0.154060204 | 0.00033062 | 0.002056519 |
| AHCY | 6720.482559 | -0.495689448 | 0.130938087 | 4.18E-05 | 0.000351757 |
| EMX2OS | 34.60565631 | -0.495768245 | 0.298570276 | 0.01291301 | 0.043627806 |
| CCDC124 | 1618.425872 | -0.495894468 | 0.149184618 | 0.00022841 | 0.001500654 |
| CEBPZOS | 1639.615562 | -0.496256758 | 0.101759952 | 3.16E-07 | 4.87E-06 |
| OGG1 | 1040.168833 | -0.496526128 | 0.122244541 | 1.36E-05 | 0.000133134 |
| BEX1 | 5788.346868 | -0.496757237 | 0.151554843 | 0.00026596 | 0.001710162 |
| EIF5A | 12723.55916 | -0.49688902 | 0.136814578 | 7.49E-05 | 0.000575915 |
| PSMG4 | 1233.422705 | -0.497013857 | 0.118712466 | 7.93E-06 | 8.24E-05 |
| DYNC2I2 | 2797.208805 | -0.497021567 | 0.119495892 | 8.93E-06 | 9.12E-05 |
| ARHGEF39 | 860.4750404 | -0.497072367 | 0.096582819 | 7.84E-08 | 1.43E-06 |
| MRFAP1L2 | 983.4700879 | -0.497344824 | 0.108778363 | 1.39E-06 | 1.78E-05 |
| DYDC2 | 48.4570833 | -0.497593273 | 0.238643154 | 0.00651364 | 0.024976023 |
| PAFAH1B3 | 2322.679293 | -0.497640296 | 0.130915971 | 3.89E-05 | 0.000329734 |
| ERCC6L | 947.7210031 | -0.49802687 | 0.160248131 | 0.00046134 | 0.002733211 |
| SMYD5 | 1198.631392 | -0.49825022 | 0.107763916 | 1.08E-06 | 1.43E-05 |
| UBE2C | 4846.573881 | -0.498449908 | 0.129726517 | 3.29E-05 | 0.000286031 |
| PTPN3 | 579.0118141 | -0.498755572 | 0.160679439 | 0.00046465 | 0.002750434 |
| CROCC | 1054.781496 | -0.498902757 | 0.138580684 | 8.33E-05 | 0.000630846 |
| TSPAN11-AS1 | 121.6880401 | -0.499539396 | 0.188039736 | 0.00172037 | 0.008304643 |
| LSM2 | 1198.310206 | -0.499564173 | 0.11219372 | 2.38E-06 | 2.86E-05 |
| TLNRD1 | 2588.898039 | -0.499631082 | 0.142796339 | 0.00012025 | 0.000867954 |
| DPM2 | 967.027522 | -0.499749414 | 0.108595753 | 1.19E-06 | 1.55E-05 |
| PARP1 | 21234.57629 | -0.500020736 | 0.125217257 | 1.77E-05 | 0.000167898 |
| PLEKHG4B | 2368.37132 | -0.500047342 | 0.093405481 | 2.53E-08 | 5.09E-07 |
| B9D2 | 160.7349487 | -0.500119118 | 0.146691255 | 0.00016506 | 0.001144756 |
| CCER2 | 49.06162253 | -0.50017955 | 0.231186636 | 0.00546589 | 0.021586437 |
| ARMC5 | 637.2488449 | -0.50031363 | 0.162829648 | 0.00050788 | 0.002967107 |
| DUT | 6202.533206 | -0.500324431 | 0.11906032 | 7.27E-06 | 7.62E-05 |
| WDR77 | 3737.937513 | -0.500512983 | 0.113555714 | 2.91E-06 | 3.42E-05 |
| SRSF2 | 17107.39957 | -0.500551732 | 0.115301145 | 3.93E-06 | 4.43E-05 |
| VASH1 | 1262.205745 | -0.500680469 | 0.102233713 | 2.78E-07 | 4.34E-06 |
| ZNF880 | 620.799498 | -0.500838185 | 0.143842895 | 0.00012688 | 0.000909107 |
| MAGI1 | 3819.441784 | -0.500959047 | 0.149408267 | 0.00019986 | 0.001338754 |
| SPON2 | 58.02326849 | -0.501425347 | 0.247169389 | 0.00700682 | 0.026465401 |
| MAPK7 | 1918.121435 | -0.501635936 | 0.151009318 | 0.00022144 | 0.001462766 |
| PABPC4 | 7018.764024 | -0.50167851 | 0.126956017 | 2.07E-05 | 0.000192627 |
| FSD1 | 2811.799819 | -0.501901215 | 0.082972476 | 4.31E-10 | 1.25E-08 |

| | | | | | |
|---|---|---|---|---|---|
| HMGN2P5 | 118.6160632 | -0.501947786 | 0.197442205 | 0.00227403 | 0.010501504 |
| MMP2 | 1536.506275 | -0.50222895 | 0.120299688 | 8.09E-06 | 8.39E-05 |
| GPM6B | 800.438361 | -0.502274335 | 0.149869674 | 0.00019917 | 0.001335397 |
| TLX2 | 13563.47777 | -0.502308277 | 0.124491854 | 1.46E-05 | 0.000142338 |
| FKBP10 | 4267.494651 | -0.502332374 | 0.078137332 | 3.83E-11 | 1.38E-09 |
| TELO2 | 2702.622704 | -0.502636496 | 0.113523228 | 2.62E-06 | 3.11E-05 |
| HECW1 | 304.2396119 | -0.502659343 | 0.166326276 | 0.00058504 | 0.003335377 |
| GREB1L | 2608.494849 | -0.502734486 | 0.127102597 | 2.03E-05 | 0.000189466 |
| PIGCP1 | 214.4344246 | -0.502865224 | 0.127803463 | 2.20E-05 | 0.00020238 |
| ARAF | 1332.555121 | -0.502916921 | 0.106972845 | 7.23E-07 | 9.96E-06 |
| MT-ND6 | 13566.27636 | -0.502985418 | 0.298318386 | 0.01174925 | 0.040464174 |
| RSAD1 | 2199.018266 | -0.503154957 | 0.107157846 | 7.42E-07 | 1.02E-05 |
| PTPRF | 7985.53357 | -0.503453556 | 0.184719354 | 0.00138578 | 0.006920125 |
| PAQR4 | 1052.300487 | -0.503622425 | 0.127403429 | 2.04E-05 | 0.000189983 |
| NDUFB2 | 2448.670958 | -0.503762437 | 0.155830597 | 0.00029497 | 0.001861326 |
| IGDCC3 | 705.8913338 | -0.504033108 | 0.124910404 | 1.45E-05 | 0.000140838 |
| NLN | 3348.834807 | -0.50409246 | 0.143343014 | 0.00010947 | 0.000800839 |
| MYLK | 408.6095499 | -0.504119602 | 0.154821626 | 0.00027213 | 0.001741684 |
| CCNT2-AS1 | 216.8803519 | -0.504128814 | 0.184348288 | 0.00134446 | 0.006754813 |
| ILF3 | 33847.8492 | -0.504379151 | 0.115318565 | 3.32E-06 | 3.82E-05 |
| MCM3 | 15475.18713 | -0.504663297 | 0.117597315 | 4.78E-06 | 5.27E-05 |
| HEATR5A-DT | 63.33049593 | -0.504699288 | 0.199799744 | 0.00231884 | 0.010672461 |
| STARD4-AS1 | 1663.797771 | -0.504869771 | 0.167646138 | 0.0005951 | 0.003383374 |
| LINC01089 | 1429.497582 | -0.504971832 | 0.134535564 | 4.47E-05 | 0.000371743 |
| SLC35A4 | 2723.992251 | -0.505037193 | 0.137543106 | 6.11E-05 | 0.000483823 |
| GPT | 158.1305988 | -0.505072299 | 0.205046636 | 0.00269882 | 0.012099093 |
| SLC25A48 | 59.01944704 | -0.505100498 | 0.216271772 | 0.0036367 | 0.015454609 |
| SIX5 | 36.33300043 | -0.50520246 | 0.244839826 | 0.00636682 | 0.02453198 |
| PNMA3 | 301.1491252 | -0.505394655 | 0.135011369 | 4.64E-05 | 0.000384276 |
| CPPED1 | 740.4131173 | -0.505820651 | 0.118686988 | 5.41E-06 | 5.88E-05 |
| BTD | 657.4372119 | -0.505834595 | 0.131488308 | 3.08E-05 | 0.000270745 |
| UTP20 | 6474.409114 | -0.505864724 | 0.143171042 | 0.00010184 | 0.000750887 |
| DDX12B | 1224.462951 | -0.505870767 | 0.104170721 | 3.31E-07 | 5.07E-06 |
| TYMP | 61.74495906 | -0.506242603 | 0.209000862 | 0.00294968 | 0.013016554 |
| IMP3 | 1530.888807 | -0.506248154 | 0.104904981 | 3.85E-07 | 5.78E-06 |
| NDUFS5 | 4498.291287 | -0.506366704 | 0.155821365 | 0.00027381 | 0.001750295 |
| SCAND1 | 880.9398924 | -0.506953497 | 0.144024236 | 0.00010615 | 0.000779302 |
| GCDH | 1427.475151 | -0.507203379 | 0.113543367 | 2.13E-06 | 2.58E-05 |
| CYCSP49 | 38.07919252 | -0.507479487 | 0.263490134 | 0.00795344 | 0.029426885 |
| TMPO | 21862.73754 | -0.507493119 | 0.116555367 | 3.56E-06 | 4.05E-05 |
| MRPL27 | 2393.483787 | -0.507718821 | 0.168886735 | 0.00059255 | 0.003372613 |
| LINC02666 | 52.70123145 | -0.507874678 | 0.211772765 | 0.00308028 | 0.01350881 |
| SMARCA4 | 17744.40443 | -0.508034377 | 0.184829924 | 0.00125597 | 0.006386684 |
| PACSIN3 | 308.4321214 | -0.508149412 | 0.137445358 | 5.44E-05 | 0.000437129 |
| KCTD12 | 7007.387754 | -0.508253909 | 0.177626048 | 0.00091041 | 0.004858959 |
| DCHS1 | 8371.033449 | -0.508262524 | 0.170304547 | 0.00063103 | 0.003561054 |
| SH2D7 | 39.50712187 | -0.508382114 | 0.242033491 | 0.00577904 | 0.022610397 |
| VPS9D1-AS1 | 279.915177 | -0.508398592 | 0.149834755 | 0.00016528 | 0.001145899 |
| CUEDC2 | 1637.159984 | -0.50871179 | 0.1093206 | 8.79E-07 | 1.19E-05 |
| HAGHL | 412.3887886 | -0.509065257 | 0.138126086 | 5.65E-05 | 0.000451308 |

| | | | | | |
|---|---|---|---|---|---|
| METTL26 | 1214.802589 | -0.509150931 | 0.093097001 | 1.26E-08 | 2.73E-07 |
| KCNQ2 | 7501.929242 | -0.509244678 | 0.111578669 | 1.34E-06 | 1.73E-05 |
| SSTR2 | 12634.96129 | -0.509252751 | 0.11368054 | 1.98E-06 | 2.43E-05 |
| BASP1 | 9560.234996 | -0.509275829 | 0.110528909 | 1.09E-06 | 1.44E-05 |
| PIK3AP1 | 186.5881049 | -0.509306088 | 0.160771866 | 0.00035133 | 0.002168952 |
| NETO2 | 6494.074585 | -0.509499255 | 0.119291773 | 5.08E-06 | 5.56E-05 |
| FBXL19 | 2100.455224 | -0.509660889 | 0.137395411 | 5.14E-05 | 0.000418212 |
| ANO8 | 1672.290919 | -0.510299473 | 0.128490541 | 1.81E-05 | 0.00017169 |
| N6AMT1 | 688.787186 | -0.510326443 | 0.218744277 | 0.00350546 | 0.015002324 |
| MCM4 | 14186.30697 | -0.510526769 | 0.115253338 | 2.48E-06 | 2.96E-05 |
| NDUFC2 | 879.9885637 | -0.511462361 | 0.118811899 | 4.32E-06 | 4.82E-05 |
| COMMD4 | 3252.54734 | -0.511502793 | 0.1178478 | 3.69E-06 | 4.19E-05 |
| PANX1 | 1862.73843 | -0.511572723 | 0.135533402 | 3.94E-05 | 0.000333808 |
| MAMSTR | 86.63558371 | -0.511626177 | 0.211412553 | 0.00283966 | 0.012623162 |
| KREMEN1 | 6728.847048 | -0.511704008 | 0.15515259 | 0.00022411 | 0.001475224 |
| FANCL | 3201.563186 | -0.512073228 | 0.141997079 | 7.47E-05 | 0.00057447 |
| RRP9 | 2018.684736 | -0.512096291 | 0.102499557 | 1.57E-07 | 2.63E-06 |
| LINC02985 | 38.63743386 | -0.512246866 | 0.243674596 | 0.0055752 | 0.021908734 |
| CHCHD5 | 443.5525603 | -0.512338892 | 0.132589926 | 2.76E-05 | 0.000246128 |
| SNORA66 | 66.30767475 | -0.512421344 | 0.230468252 | 0.0043678 | 0.017974883 |
| BRINP1 | 258.9871383 | -0.512535196 | 0.177583311 | 0.00082032 | 0.004435704 |
| DANCR | 4251.381107 | -0.512562384 | 0.143730303 | 8.63E-05 | 0.000650368 |
| NPR2 | 1347.174289 | -0.512645951 | 0.143782697 | 8.65E-05 | 0.000651242 |
| FOXM1 | 5099.454093 | -0.513586687 | 0.190126128 | 0.00137076 | 0.006865147 |
| DBNDD1 | 514.7564254 | -0.513712761 | 0.179529033 | 0.00087387 | 0.004688296 |
| VSTM2A | 1708.780004 | -0.514247871 | 0.132267459 | 2.47E-05 | 0.000224429 |
| ZNF497-AS1 | 54.72374563 | -0.514450641 | 0.245148616 | 0.00551034 | 0.021716097 |
| GDF11 | 1305.968068 | -0.514618029 | 0.112304167 | 1.19E-06 | 1.56E-05 |
| KCNQ4 | 276.8141974 | -0.514664134 | 0.138581434 | 4.89E-05 | 0.000400958 |
| SIRPA | 76.81869007 | -0.514675942 | 0.265165374 | 0.00729257 | 0.027355778 |
| TUBB4B | 7942.182255 | -0.51469719 | 0.12498718 | 9.53E-06 | 9.68E-05 |
| MIR6835 | 19.17322032 | -0.514734363 | 0.32497989 | 0.01198385 | 0.041127468 |
| ZNF407 | 776.6125133 | -0.514764284 | 0.131743181 | 2.28E-05 | 0.000208903 |
| SECTM1 | 90.37165114 | -0.514830355 | 0.184464179 | 0.00105989 | 0.005527673 |
| DIPK1B | 1342.939723 | -0.514844388 | 0.113586599 | 1.50E-06 | 1.91E-05 |
| POP1 | 2795.634828 | -0.514883843 | 0.14201363 | 6.81E-05 | 0.00053218 |
| PFDN6 | 1109.061481 | -0.515031213 | 0.135130876 | 3.34E-05 | 0.000290038 |
| LIMD1 | 517.2507855 | -0.515157747 | 0.139385653 | 5.21E-05 | 0.000422518 |
| PUF60 | 3934.692578 | -0.515180193 | 0.139372082 | 5.20E-05 | 0.000422492 |
| MSH5 | 522.4528451 | -0.51542192 | 0.148376962 | 0.00011836 | 0.000856119 |
| NOL12 | 182.5135395 | -0.515425307 | 0.174351145 | 0.000652 | 0.003663323 |
| ASPSCR1 | 2706.382969 | -0.515464854 | 0.105150752 | 2.48E-07 | 3.93E-06 |
| P2RX5-TAX1BP3 | 47.47632154 | -0.515535206 | 0.271696964 | 0.00777384 | 0.02888711 |
| REX1BD | 1199.795572 | -0.515619955 | 0.114534929 | 1.72E-06 | 2.15E-05 |
| IMP4 | 2868.386532 | -0.515884695 | 0.117721455 | 2.97E-06 | 3.48E-05 |
| TNFSF13 | 25.95191948 | -0.516142373 | 0.300798947 | 0.01008885 | 0.035784026 |
| ZNF750 | 44.56421735 | -0.516257706 | 0.246423421 | 0.00546288 | 0.021580702 |
| PDXP | 392.2658597 | -0.516285555 | 0.159387905 | 0.00026442 | 0.001702924 |
| NOS3 | 49.4257454 | -0.516634632 | 0.261985366 | 0.00680307 | 0.025849638 |
| TK1 | 3575.697868 | -0.516724558 | 0.128688597 | 1.45E-05 | 0.0001409 |

| | | | | | |
|---|---|---|---|---|---|
| RAD54L | 1106.459693 | -0.51673756 | 0.126832946 | 1.13E-05 | 0.00011326 |
| LRCH1 | 2616.802736 | -0.516843515 | 0.135094012 | 3.11E-05 | 0.000273334 |
| ZFTA | 3317.971903 | -0.516868526 | 0.119167137 | 3.61E-06 | 4.11E-05 |
| LINC00624 | 107.8790272 | -0.516971337 | 0.182837229 | 0.00094007 | 0.004999094 |
| MDK | 12614.30725 | -0.517183222 | 0.108393652 | 4.71E-07 | 6.94E-06 |
| PABPC4-AS1 | 64.20108408 | -0.517244979 | 0.205667818 | 0.0021584 | 0.010078409 |
| HMGXB4 | 2546.122455 | -0.517411736 | 0.131321353 | 1.96E-05 | 0.000184068 |
| MPLKIP | 947.1071946 | -0.517509384 | 0.115628109 | 1.92E-06 | 2.36E-05 |
| CCNE2 | 531.9226174 | -0.517666932 | 0.145440972 | 8.55E-05 | 0.000644958 |
| TRIP13 | 2814.268509 | -0.517680208 | 0.117466791 | 2.63E-06 | 3.11E-05 |
| GAREM2 | 1647.300942 | -0.517718346 | 0.124484702 | 7.85E-06 | 8.17E-05 |
| HMBS | 954.4639004 | -0.518315305 | 0.110522597 | 6.96E-07 | 9.66E-06 |
| MLST8 | 1846.735645 | -0.518374344 | 0.13850824 | 4.27E-05 | 0.000358087 |
| BVES-AS1 | 114.8716793 | -0.518569069 | 0.192525992 | 0.00134671 | 0.006762795 |
| GATM | 598.0922231 | -0.518611109 | 0.124155903 | 7.22E-06 | 7.57E-05 |
| CENPW | 669.5358638 | -0.518642998 | 0.147998813 | 0.00010367 | 0.000762995 |
| SLC25A3 | 17157.36593 | -0.519037585 | 0.137839872 | 3.89E-05 | 0.000329734 |
| SMIM7 | 1389.757163 | -0.519105976 | 0.138491487 | 4.15E-05 | 0.000349643 |
| OLFM1 | 11527.5126 | -0.51949541 | 0.153591344 | 0.00015881 | 0.001107406 |
| ARRB1 | 1146.925974 | -0.520061003 | 0.156583222 | 0.00019516 | 0.001313683 |
| EMG1 | 443.4067856 | -0.520096847 | 0.164610829 | 0.00033395 | 0.002075388 |
| PPIL2 | 2402.173819 | -0.520399288 | 0.121349789 | 4.39E-06 | 4.88E-05 |
| DDX12P | 2333.357122 | -0.520590363 | 0.100636115 | 5.93E-08 | 1.11E-06 |
| TRAPPC1 | 985.1483444 | -0.520654857 | 0.13293836 | 2.11E-05 | 0.000195733 |
| HNRNPA3P5 | 53.24900708 | -0.520821662 | 0.228157128 | 0.00358981 | 0.015296505 |
| ZBTB22 | 538.5617136 | -0.520824402 | 0.142390302 | 5.80E-05 | 0.000461995 |
| PEG10 | 8710.770192 | -0.521234833 | 0.100321952 | 5.23E-08 | 9.91E-07 |
| H2AX | 3736.681293 | -0.521286952 | 0.129765424 | 1.39E-05 | 0.000136401 |
| ZNF511 | 285.3772027 | -0.521555271 | 0.146460895 | 8.26E-05 | 0.000627612 |
| MTX1LP | 181.3931985 | -0.521671676 | 0.245376193 | 0.0049151 | 0.019826242 |
| PGAP2 | 725.0020586 | -0.522048503 | 0.121633389 | 4.27E-06 | 4.77E-05 |
| SNRPD3 | 3427.911455 | -0.522275621 | 0.099149574 | 3.53E-08 | 6.92E-07 |
| H2AZ2P1 | 42.84994111 | -0.522303573 | 0.272297204 | 0.00709079 | 0.026740317 |
| SALL1 | 91.13293593 | -0.52234693 | 0.231704997 | 0.00376591 | 0.015911704 |
| RAD51-AS1 | 505.5056163 | -0.522459669 | 0.168181821 | 0.00038892 | 0.00236478 |
| COA3 | 1773.306693 | -0.522465608 | 0.145060563 | 7.07E-05 | 0.00054916 |
| CCNB1 | 6529.393919 | -0.522709513 | 0.159400454 | 0.00022079 | 0.001459862 |
| LDHA | 24055.35409 | -0.522746168 | 0.150083235 | 0.00010875 | 0.000796108 |
| PRTG | 873.1284371 | -0.522790471 | 0.177232083 | 0.00062895 | 0.003551286 |
| DEAF1 | 1471.982598 | -0.522902494 | 0.131751971 | 1.68E-05 | 0.000160693 |
| GALNT2 | 5972.743376 | -0.522939171 | 0.147752159 | 8.86E-05 | 0.000664151 |
| SPHK1 | 30.96311163 | -0.523161063 | 0.297961114 | 0.00899667 | 0.032634948 |
| LOXL1-AS1 | 807.0058827 | -0.523265 | 0.100197498 | 4.48E-08 | 8.64E-07 |
| PLK1 | 3769.594388 | -0.523296251 | 0.148523119 | 9.37E-05 | 0.000699767 |
| USP5 | 6428.159256 | -0.523460998 | 0.156791959 | 0.00017952 | 0.001227938 |
| ATP5F1D | 1582.454155 | -0.523857954 | 0.145713173 | 7.17E-05 | 0.000555352 |
| KIF18B-DT | 11.83799308 | -0.524252159 | 0.400834129 | 0.01403805 | 0.046613675 |
| TBC1D22A-DT | 78.77265081 | -0.524417945 | 0.231791443 | 0.00363766 | 0.015454609 |
| ATAT1 | 2916.270614 | -0.524483728 | 0.082894176 | 6.53E-11 | 2.22E-09 |
| NMU | 169.5730676 | -0.524720056 | 0.158680906 | 0.00019826 | 0.001331653 |

| | | | | | |
|---|---|---|---|---|---|
| RSL24D1P3 | 51.05883278 | -0.524910043 | 0.237368965 | 0.00403135 | 0.016836032 |
| SLC29A2 | 1934.917035 | -0.524971159 | 0.129096868 | 1.11E-05 | 0.000110904 |
| CORO2B | 887.5426717 | -0.525136054 | 0.220931882 | 0.00280913 | 0.012528088 |
| KPNA2 | 25490.10442 | -0.52515305 | 0.145002402 | 6.44E-05 | 0.000506629 |
| PRAG1 | 1015.583556 | -0.525202957 | 0.173587407 | 0.00049059 | 0.002882402 |
| MAST3 | 635.0396202 | -0.525629038 | 0.17736672 | 0.00059057 | 0.003362265 |
| RNF227 | 187.0243568 | -0.52588649 | 0.158401888 | 0.00018797 | 0.001272325 |
| MMP15 | 718.03362 | -0.525964271 | 0.141965002 | 4.68E-05 | 0.000387471 |
| ADCYAP1 | 3141.585975 | -0.525964376 | 0.165454244 | 0.00030033 | 0.001891004 |
| ALPL | 185.7574886 | -0.526611967 | 0.208532027 | 0.00194979 | 0.009255639 |
| CALML4 | 113.9491798 | -0.526705455 | 0.174635906 | 0.00049946 | 0.002925341 |
| DUS3L | 2137.365883 | -0.526922765 | 0.121169389 | 3.22E-06 | 3.72E-05 |
| HAND2-AS1 | 35456.46733 | -0.527253608 | 0.13409841 | 1.90E-05 | 0.000178793 |
| TBXAS1 | 182.0254255 | -0.527866832 | 0.138709379 | 3.13E-05 | 0.000274283 |
| PTPRU | 1534.300018 | -0.528012673 | 0.156469554 | 0.00015344 | 0.001074344 |
| RGS12 | 2210.196589 | -0.528040981 | 0.115613154 | 1.17E-06 | 1.53E-05 |
| GTPBP3 | 1932.821731 | -0.528303441 | 0.097316972 | 1.41E-08 | 3.01E-07 |
| PTER | 201.2561083 | -0.528773209 | 0.173169554 | 0.00043796 | 0.002618864 |
| ECE2 | 76.78229976 | -0.529039331 | 0.185320549 | 0.00079191 | 0.004308139 |
| GSTP1 | 8908.343975 | -0.529852563 | 0.144565569 | 5.29E-05 | 0.000427495 |
| JPT1 | 11239.97025 | -0.530021342 | 0.141081281 | 3.72E-05 | 0.00031765 |
| PTPN18 | 1011.339062 | -0.530368899 | 0.099825264 | 2.63E-08 | 5.28E-07 |
| LINC00896 | 84.08960061 | -0.530416175 | 0.202700863 | 0.00149999 | 0.007388226 |
| TMEM25 | 555.1904898 | -0.530537916 | 0.122561599 | 3.43E-06 | 3.93E-05 |
| CDKN2C | 4343.840866 | -0.530594316 | 0.133346157 | 1.53E-05 | 0.000147787 |
| TMSB15A | 2538.51868 | -0.530656988 | 0.168508499 | 0.00031919 | 0.001995062 |
| PASK | 4344.243218 | -0.530699548 | 0.090638528 | 1.18E-09 | 3.13E-08 |
| SOX12 | 6049.352982 | -0.530753762 | 0.136466555 | 2.20E-05 | 0.000202075 |
| ANTXR2 | 2566.900392 | -0.531068569 | 0.127255724 | 6.75E-06 | 7.14E-05 |
| KCTD17 | 1116.967647 | -0.532006432 | 0.096464927 | 8.48E-09 | 1.88E-07 |
| MMACHC | 441.0876595 | -0.532105026 | 0.13080893 | 1.05E-05 | 0.000105522 |
| PITX1 | 2070.738793 | -0.532222326 | 0.128953016 | 8.16E-06 | 8.45E-05 |
| VASH2 | 3377.60568 | -0.532345494 | 0.112307275 | 4.98E-07 | 7.27E-06 |
| MIR27B | 8.289681785 | -0.53262289 | 0.466085671 | 0.01409585 | 0.046752752 |
| LINC00938 | 1356.347904 | -0.532743675 | 0.145071448 | 5.05E-05 | 0.000411486 |
| CTNS-AS1 | 47.26628321 | -0.532810001 | 0.228484946 | 0.0029139 | 0.012883556 |
| COMTD1 | 244.9548283 | -0.533020216 | 0.140290023 | 3.09E-05 | 0.000271425 |
| GJC1 | 10816.48473 | -0.533038986 | 0.103379746 | 5.99E-08 | 1.12E-06 |
| RPIA | 1104.191897 | -0.533167811 | 0.106004889 | 1.16E-07 | 2.02E-06 |
| MAB21L1 | 7143.202939 | -0.533344609 | 0.145414237 | 5.11E-05 | 0.000416273 |
| NSG1 | 1930.298052 | -0.533447253 | 0.14564429 | 5.21E-05 | 0.000422492 |
| RN7SL481P | 31.12116693 | -0.53356108 | 0.265807593 | 0.00552754 | 0.021754702 |
| RNU6-388P | 9.744968519 | -0.533980954 | 0.442990438 | 0.01368714 | 0.045676169 |
| LGALS3BP | 3507.088522 | -0.533994179 | 0.124222155 | 3.83E-06 | 4.33E-05 |
| MIR5587 | 70.46712454 | -0.534134292 | 0.204517937 | 0.00147416 | 0.007281909 |
| KRT8P12 | 212.4227483 | -0.534394225 | 0.147994166 | 6.27E-05 | 0.000495132 |
| KDM2B | 3210.639519 | -0.534433993 | 0.151466481 | 8.49E-05 | 0.000641307 |
| RNASEH2A | 35.21200891 | -0.534580412 | 0.250976985 | 0.00436565 | 0.017973028 |
| GATD1-DT | 182.0405501 | -0.534587174 | 0.171035448 | 0.00033396 | 0.002075388 |
| PFN2 | 13000.01213 | -0.534896135 | 0.129657152 | 8.07E-06 | 8.37E-05 |

| | | | | | |
|---|---|---|---|---|---|
| KLF16 | 1058.097107 | -0.535308824 | 0.191396547 | 0.00088777 | 0.00475517 |
| CHST12 | 1253.272066 | -0.535355701 | 0.112643086 | 4.59E-07 | 6.79E-06 |
| SLC12A9 | 797.2376751 | -0.535492723 | 0.109017757 | 2.08E-07 | 3.37E-06 |
| LINC02367 | 339.1808938 | -0.535632454 | 0.140958141 | 3.03E-05 | 0.000267016 |
| NTAN1 | 386.9924728 | -0.535662636 | 0.166691896 | 0.00024937 | 0.001617663 |
| FSCN1 | 14057.5917 | -0.535711388 | 0.165706728 | 0.00023387 | 0.001533081 |
| TPI1 | 19336.57097 | -0.536112388 | 0.130183878 | 8.25E-06 | 8.53E-05 |
| NRDE2 | 1318.584768 | -0.536775173 | 0.111072739 | 3.07E-07 | 4.74E-06 |
| TAS2R19 | 76.14342358 | -0.536850796 | 0.249689917 | 0.00411647 | 0.017125463 |
| KIT | 1676.177354 | -0.537496291 | 0.169105681 | 0.0002756 | 0.001758992 |
| NCAPD2 | 11284.01595 | -0.537843741 | 0.139437002 | 2.38E-05 | 0.000216793 |
| NCBP2AS2 | 772.346544 | -0.537858544 | 0.110266597 | 2.43E-07 | 3.86E-06 |
| HTR7 | 28.99882114 | -0.538039866 | 0.28777454 | 0.00668736 | 0.025497211 |
| KCNH7 | 471.0169909 | -0.538354214 | 0.143516285 | 3.59E-05 | 0.000308207 |
| LSM4 | 4568.984325 | -0.538665364 | 0.119883589 | 1.54E-06 | 1.96E-05 |
| NUDT11 | 1799.362385 | -0.538735236 | 0.143567347 | 3.56E-05 | 0.000306052 |
| MYB | 551.7360058 | -0.538779236 | 0.153513983 | 8.80E-05 | 0.00066087 |
| RCAN3 | 120.3571701 | -0.538889944 | 0.247606779 | 0.00383893 | 0.016160436 |
| GFOD2 | 1533.22352 | -0.539224425 | 0.09601569 | 4.56E-09 | 1.07E-07 |
| ROR1-AS1 | 4.992276867 | -0.539230035 | 0.563285291 | 0.01320706 | 0.044399225 |
| TPM3P6 | 22.76969808 | -0.539462101 | 0.318322528 | 0.00849429 | 0.031070188 |
| SLC5A6 | 3499.831069 | -0.539836124 | 0.111605724 | 2.94E-07 | 4.57E-06 |
| CENPB | 2833.25216 | -0.540398634 | 0.142178032 | 2.91E-05 | 0.000258179 |
| PTMA | 48788.47134 | -0.540556768 | 0.138989491 | 2.05E-05 | 0.000191196 |
| LOXL1 | 1726.498228 | -0.540720878 | 0.204061917 | 0.00126466 | 0.006424526 |
| YWHAQ | 20852.68942 | -0.540854223 | 0.112458921 | 3.35E-07 | 5.10E-06 |
| MRPL54 | 486.2751347 | -0.540928942 | 0.139025234 | 2.03E-05 | 0.000189924 |
| SMPD3 | 1526.962314 | -0.540963953 | 0.1389272 | 2.01E-05 | 0.000187954 |
| AKAP1 | 5665.442852 | -0.541001049 | 0.121375673 | 1.79E-06 | 2.22E-05 |
| RPL23AP20 | 70.57634705 | -0.541017351 | 0.210278141 | 0.00153688 | 0.007542301 |
| DOT1L | 7156.653253 | -0.541062227 | 0.109075347 | 1.57E-07 | 2.63E-06 |
| IL17RD | 755.102286 | -0.541477404 | 0.179767225 | 0.0004505 | 0.002679018 |
| MRPL12 | 1166.621899 | -0.541842372 | 0.148911995 | 5.36E-05 | 0.000432223 |
| RNASEH1-DT | 446.7154042 | -0.541865295 | 0.12505165 | 3.11E-06 | 3.62E-05 |
| ZNF783 | 2196.906526 | -0.542500788 | 0.0933549 | 1.43E-09 | 3.70E-08 |
| PALM2AKAP2 | 2237.593038 | -0.542503838 | 0.13219491 | 8.38E-06 | 8.63E-05 |
| HPCAL1 | 626.0165802 | -0.542592705 | 0.152980301 | 7.47E-05 | 0.000574582 |
| LMO3 | 442.5956993 | -0.542611265 | 0.337911282 | 0.00923556 | 0.033354432 |
| PDE1A | 330.0309648 | -0.542682563 | 0.18914811 | 0.00068139 | 0.003805581 |
| HOMER3 | 283.2700012 | -0.543117891 | 0.168152209 | 0.00022308 | 0.001470295 |
| LSAMP | 4729.758987 | -0.543216086 | 0.138562288 | 1.78E-05 | 0.000168853 |
| PRR19 | 72.75636216 | -0.543454122 | 0.197525935 | 0.00094235 | 0.005006932 |
| SYNM-AS1 | 21.77169188 | -0.543562222 | 0.372701381 | 0.01070143 | 0.037566878 |
| CFAP157 | 108.3279473 | -0.543801443 | 0.192955003 | 0.00078025 | 0.004258264 |
| PUDP | 659.7527486 | -0.543981539 | 0.151233944 | 6.16E-05 | 0.000486873 |
| TEX15 | 782.6033215 | -0.544427435 | 0.162061713 | 0.00014293 | 0.001011382 |
| RASL11B | 202.5375395 | -0.544496299 | 0.163659137 | 0.00015954 | 0.001111739 |
| IMPDH1P6 | 4.172882521 | -0.544928592 | 0.66723032 | 0.01113245 | 0.038748765 |
| MAZ | 2791.910849 | -0.544946122 | 0.159095629 | 0.00011329 | 0.000824037 |
| BACH1-AS1 | 19.02017723 | -0.545018097 | 0.348344259 | 0.00948852 | 0.034052673 |

| | | | | | |
|---|---|---|---|---|---|
| MCRIP2 | 524.4164532 | -0.545049821 | 0.117293471 | 7.16E-07 | 9.90E-06 |
| UNC5A | 2130.282431 | -0.545339268 | 0.140499565 | 2.05E-05 | 0.000190647 |
| RN7SL1 | 620.1544367 | -0.545592043 | 0.45898894 | 0.01261648 | 0.042841043 |
| HNRNPA1P7 | 2404.761724 | -0.545902417 | 0.157016197 | 9.38E-05 | 0.000700281 |
| CKB | 14021.72258 | -0.545952743 | 0.144471273 | 3.05E-05 | 0.000268292 |
| PRKX | 3197.989355 | -0.546243446 | 0.112305112 | 2.46E-07 | 3.90E-06 |
| GINS3 | 1832.907985 | -0.546578531 | 0.14515043 | 3.19E-05 | 0.000278968 |
| TYMS | 2716.315815 | -0.546640003 | 0.102188036 | 1.94E-08 | 4.01E-07 |
| IL15 | 15.87015664 | -0.546819655 | 0.389373008 | 0.01093728 | 0.038187183 |
| KCNS2 | 150.9552219 | -0.546866384 | 0.178097117 | 0.00036047 | 0.002218694 |
| ROBO1 | 6745.679661 | -0.547267711 | 0.125129059 | 2.50E-06 | 2.98E-05 |
| DLK1 | 210.5991539 | -0.547393574 | 0.151380815 | 5.59E-05 | 0.000446894 |
| RNVU1-25 | 4.942062456 | -0.547424648 | 0.575788676 | 0.01242797 | 0.042298716 |
| HMX1 | 1953.950352 | -0.547738289 | 0.144500719 | 2.87E-05 | 0.000254959 |
| DCX | 9076.212152 | -0.547833745 | 0.133302862 | 7.87E-06 | 8.18E-05 |
| PEBP1 | 13412.78411 | -0.547836601 | 0.116688992 | 5.58E-07 | 8.06E-06 |
| TNNT2 | 8.458090552 | -0.547910152 | 0.46310224 | 0.01242624 | 0.042298716 |
| COL18A1 | 715.1632504 | -0.548252661 | 0.14904646 | 4.40E-05 | 0.000367324 |
| PCDHGA12 | 123.7206566 | -0.548376237 | 0.208878739 | 0.00126387 | 0.006422095 |
| SERF1A | 86.03675001 | -0.548413077 | 0.193230106 | 0.00071024 | 0.00393537 |
| PPP2R1A | 8817.701572 | -0.548484966 | 0.140273139 | 1.78E-05 | 0.000169227 |
| RAVER1 | 1424.640588 | -0.548523998 | 0.174161353 | 0.00027743 | 0.001767893 |
| NES | 4074.827537 | -0.548701364 | 0.145580007 | 3.10E-05 | 0.000272313 |
| RNF207-AS1 | 40.97212398 | -0.549106341 | 0.262610952 | 0.00414662 | 0.017223006 |
| LY6E | 3196.453613 | -0.549214745 | 0.146998665 | 3.51E-05 | 0.000302701 |
| FAM171A1 | 5132.645493 | -0.549222157 | 0.169560014 | 0.00020631 | 0.001375205 |
| NHP2 | 1621.166521 | -0.5493972 | 0.123275048 | 1.69E-06 | 2.12E-05 |
| VCAN | 3994.433445 | -0.549771414 | 0.180685268 | 0.00038383 | 0.002340781 |
| SEMA4C | 2853.876802 | -0.549921396 | 0.088846924 | 1.34E-10 | 4.27E-09 |
| MYCNOS | 4741.63524 | -0.549930062 | 0.124824202 | 2.12E-06 | 2.58E-05 |
| RPSAP54 | 55.23792215 | -0.549958936 | 0.231176362 | 0.00226453 | 0.010469387 |
| WDR86-AS1 | 65.05777231 | -0.550212828 | 0.24171573 | 0.00283148 | 0.012594957 |
| FAM210B | 351.2302134 | -0.550286856 | 0.146338063 | 3.17E-05 | 0.000277445 |
| MZF1 | 1452.748811 | -0.550317037 | 0.113876573 | 2.80E-07 | 4.37E-06 |
| PCDHGA10 | 305.1934912 | -0.550454878 | 0.245772231 | 0.00305182 | 0.013404022 |
| CHML | 6631.535968 | -0.550710373 | 0.152115039 | 5.37E-05 | 0.000432662 |
| MTATP6P1 | 4928.399455 | -0.551026369 | 0.144596095 | 2.59E-05 | 0.000233613 |
| AKT2 | 4804.277039 | -0.551107042 | 0.090452008 | 2.44E-10 | 7.40E-09 |
| MGAT4B | 1750.126674 | -0.55151783 | 0.115915336 | 4.01E-07 | 6.00E-06 |
| ATP8B1 | 65.84586839 | -0.551567322 | 0.195352434 | 0.00072121 | 0.003986791 |
| MRPL23 | 651.5121771 | -0.551852802 | 0.144417014 | 2.47E-05 | 0.000224337 |
| FXYD6 | 4901.911385 | -0.5521336 | 0.117323643 | 5.13E-07 | 7.47E-06 |
| MEIS1-AS3 | 117.5813365 | -0.552158797 | 0.171783024 | 0.00021863 | 0.001446577 |
| YPEL1 | 621.2941658 | -0.552398327 | 0.163897799 | 0.00012935 | 0.000925175 |
| DPH2 | 1527.228987 | -0.552689966 | 0.160845493 | 0.0001027 | 0.000756436 |
| CTU1 | 317.4109606 | -0.552824935 | 0.150959626 | 4.52E-05 | 0.000375424 |
| PRDX1 | 5995.824555 | -0.553497288 | 0.152161047 | 4.93E-05 | 0.000403272 |
| ZNF726 | 393.6143094 | -0.553628155 | 0.148572403 | 3.52E-05 | 0.0003035 |
| ADAMTS9 | 3123.734481 | -0.553673725 | 0.155413297 | 6.48E-05 | 0.000509758 |
| LRRC8A | 2596.706355 | -0.554366735 | 0.142427169 | 1.83E-05 | 0.00017323 |

| | | | | | |
|---|---|---|---|---|---|
| MICOS13 | 455.5369224 | -0.554500827 | 0.128971573 | 3.30E-06 | 3.81E-05 |
| SEPTIN9 | 16457.85815 | -0.554899815 | 0.16706275 | 0.0001498 | 0.001053491 |
| LYRM4-AS1 | 70.76132631 | -0.555375952 | 0.212117217 | 0.00121273 | 0.0062036 |
| MCRIP1 | 3878.378507 | -0.555620999 | 0.100748253 | 7.26E-09 | 1.64E-07 |
| TMOD4 | 98.96065062 | -0.555644801 | 0.188186451 | 0.00048003 | 0.002827625 |
| UGT8 | 752.0066232 | -0.555819451 | 0.172067308 | 0.0002015 | 0.00134794 |
| NREP | 1841.298454 | -0.555829069 | 0.149678556 | 3.64E-05 | 0.000311295 |
| FGF10-AS1 | 20.76798862 | -0.556524727 | 0.387589028 | 0.0098545 | 0.035084492 |
| TEAD2 | 805.8468262 | -0.556678558 | 0.173524305 | 0.00021513 | 0.001425685 |
| AKT1 | 5161.072835 | -0.556805375 | 0.112375063 | 1.45E-07 | 2.45E-06 |
| ACMSD | 26.73628026 | -0.556821387 | 0.317159045 | 0.00674894 | 0.025672427 |
| NRIP2 | 321.6000062 | -0.556917412 | 0.135797778 | 7.63E-06 | 7.97E-05 |
| SMARCD1 | 8652.255872 | -0.557101996 | 0.14496822 | 2.18E-05 | 0.000200995 |
| NUDT1 | 1072.903 | -0.557293429 | 0.130163543 | 3.50E-06 | 3.99E-05 |
| CALM3 | 8926.299716 | -0.557472172 | 0.099079208 | 3.80E-09 | 9.06E-08 |
| ZNF668 | 264.8225865 | -0.557505516 | 0.159698683 | 8.15E-05 | 0.000619965 |
| C15orf32 | 18.97923346 | -0.557924468 | 0.335617424 | 0.00758516 | 0.028290661 |
| CDHR3 | 290.589398 | -0.558039291 | 0.151839487 | 4.13E-05 | 0.000348345 |
| RNF167 | 2146.768951 | -0.55807264 | 0.103140176 | 1.28E-08 | 2.75E-07 |
| PGA5 | 57.14014459 | -0.558407549 | 0.219396112 | 0.00141999 | 0.007060091 |
| ADAT3 | 163.7711942 | -0.558832811 | 0.154753461 | 5.21E-05 | 0.000422518 |
| MIR23B | 4.81195432 | -0.558916825 | 0.641502641 | 0.0108354 | 0.037939583 |
| PPRC1 | 6032.534343 | -0.558998138 | 0.126438093 | 1.85E-06 | 2.29E-05 |
| LINC01465 | 12.28525671 | -0.559043436 | 0.429318909 | 0.01075002 | 0.037711269 |
| MYBBP1A | 3667.394899 | -0.5590529 | 0.160012836 | 7.97E-05 | 0.000607427 |
| AACSP1 | 101.0641189 | -0.559433221 | 0.193416262 | 0.00055473 | 0.003189108 |
| ELF2P1 | 75.94109566 | -0.559465623 | 0.202303734 | 0.00079264 | 0.004310981 |
| MCUB | 130.1925423 | -0.559545071 | 0.1720898 | 0.00018233 | 0.00124315 |
| TIMM8A | 723.3282732 | -0.559649182 | 0.132005612 | 4.13E-06 | 4.63E-05 |
| APEX2 | 893.0128295 | -0.559707534 | 0.111854135 | 1.10E-07 | 1.93E-06 |
| ZBED10P | 1354.087448 | -0.55973573 | 0.108300949 | 4.69E-08 | 8.95E-07 |
| SLC38A5 | 1476.95552 | -0.560268031 | 0.114175372 | 1.80E-07 | 2.98E-06 |
| LINC00470 | 118.2058734 | -0.560405815 | 0.217381218 | 0.00128624 | 0.006514208 |
| KDM8 | 287.5043033 | -0.560510441 | 0.124707484 | 1.31E-06 | 1.70E-05 |
| BANF1 | 4154.120359 | -0.561288368 | 0.132587734 | 4.20E-06 | 4.69E-05 |
| UQCR11 | 602.4336891 | -0.561388529 | 0.124237689 | 1.16E-06 | 1.53E-05 |
| THOC6 | 1617.954375 | -0.561634769 | 0.121993098 | 7.81E-07 | 1.07E-05 |
| LINC01561 | 22.57293679 | -0.561866957 | 0.322401309 | 0.00658734 | 0.02520162 |
| PPP1R14B | 3413.038033 | -0.561933455 | 0.107924949 | 3.77E-08 | 7.35E-07 |
| FAM89B | 178.5722698 | -0.562095596 | 0.187454488 | 0.00039656 | 0.002404163 |
| RPL36A | 577.6130844 | -0.562143021 | 0.125178064 | 1.32E-06 | 1.71E-05 |
| PRDM15 | 755.7657502 | -0.563017494 | 0.12926258 | 2.42E-06 | 2.89E-05 |
| GPC2 | 3281.506509 | -0.563273526 | 0.098990941 | 2.53E-09 | 6.26E-08 |
| ACBD7 | 1170.428495 | -0.563371516 | 0.112301657 | 1.01E-07 | 1.79E-06 |
| TH | 922.4278589 | -0.56361321 | 0.146235177 | 1.98E-05 | 0.000185954 |
| UBE2FP1 | 31.3139087 | -0.563924401 | 0.28053152 | 0.00422504 | 0.017495706 |
| RAB29 | 323.3135278 | -0.564209412 | 0.133387242 | 4.17E-06 | 4.66E-05 |
| SLC25A10 | 337.0338118 | -0.565184634 | 0.16909375 | 0.00012842 | 0.000919201 |
| C1QBP | 5772.525337 | -0.565196019 | 0.123743605 | 9.02E-07 | 1.22E-05 |
| PA2G4P2 | 48.23267597 | -0.565210865 | 0.25193698 | 0.00266155 | 0.011947626 |

| | | | | | |
|---|---|---|---|---|---|
| SEMA7A | 459.6207801 | -0.565926613 | 0.131424141 | 2.95E-06 | 3.45E-05 |
| PPIAP11 | 154.5942614 | -0.566020098 | 0.167551866 | 0.00011289 | 0.000822057 |
| PDCL3 | 1090.125919 | -0.56621452 | 0.171063442 | 0.0001421 | 0.001006927 |
| ARL2 | 456.7595104 | -0.566273484 | 0.138794911 | 7.77E-06 | 8.09E-05 |
| ARHGDIA | 8733.221859 | -0.566276924 | 0.184092312 | 0.00030222 | 0.001902322 |
| WEE1 | 3388.577875 | -0.56627948 | 0.106988491 | 2.30E-08 | 4.68E-07 |
| LINC01424 | 102.9399954 | -0.566356515 | 0.17413745 | 0.00017192 | 0.001183582 |
| GRN | 2775.71186 | -0.566627269 | 0.150586508 | 2.77E-05 | 0.000246796 |
| TMEM35A | 617.8163518 | -0.566660904 | 0.156955217 | 4.91E-05 | 0.000402744 |
| ZNF778-DT | 47.76466249 | -0.56699319 | 0.250004121 | 0.00248885 | 0.011295221 |
| KRI1 | 2031.224973 | -0.567510432 | 0.093511921 | 2.53E-10 | 7.65E-09 |
| TMEM204 | 48.22524944 | -0.567570378 | 0.25369846 | 0.00264126 | 0.011866903 |
| B3GNTL1 | 581.5573415 | -0.567899733 | 0.127466823 | 1.49E-06 | 1.90E-05 |
| CHST14 | 1019.737702 | -0.568302823 | 0.129176667 | 1.91E-06 | 2.35E-05 |
| DMAC2L | 467.0116152 | -0.568388807 | 0.143141982 | 1.20E-05 | 0.000119045 |
| KCNK2 | 1128.978111 | -0.568576566 | 0.132601897 | 3.13E-06 | 3.64E-05 |
| ACTBP11 | 25.23502055 | -0.568610261 | 0.392908547 | 0.00887359 | 0.032291023 |
| RPL30-AS1 | 138.1791899 | -0.56949244 | 0.172644821 | 0.00014365 | 0.001015497 |
| NCAPH | 5173.57383 | -0.570198617 | 0.121325404 | 4.64E-07 | 6.85E-06 |
| ATXN7L3B | 11050.30923 | -0.571604726 | 0.128346511 | 1.46E-06 | 1.86E-05 |
| MEX3D | 1817.271746 | -0.571813653 | 0.139783722 | 7.11E-06 | 7.47E-05 |
| EID1 | 10917.96072 | -0.5718315 | 0.15459812 | 3.39E-05 | 0.000293937 |
| STUM | 343.440417 | -0.571981218 | 0.156288832 | 3.92E-05 | 0.000332156 |
| LINC01798 | 3806.547765 | -0.572056032 | 0.139408292 | 6.73E-06 | 7.12E-05 |
| PIMREG | 3235.755605 | -0.572189785 | 0.101928941 | 3.69E-09 | 8.87E-08 |
| B4GAT1-DT | 261.332675 | -0.572463369 | 0.133138677 | 2.88E-06 | 3.39E-05 |
| MICOS10 | 233.578556 | -0.572686576 | 0.13783446 | 5.39E-06 | 5.85E-05 |
| HSPD1P11 | 4.388715277 | -0.573067713 | 0.637688535 | 0.01010826 | 0.035826669 |
| NME1 | 5873.904583 | -0.573513255 | 0.146967236 | 1.52E-05 | 0.000146528 |
| TRAM1L1 | 178.7161973 | -0.573611124 | 0.206393373 | 0.00067572 | 0.003780043 |
| CFL1 | 12077.94025 | -0.573687902 | 0.12883102 | 1.44E-06 | 1.84E-05 |
| SLX1B | 7.291135423 | -0.573830505 | 0.540198864 | 0.01061335 | 0.037320945 |
| KIAA1549L | 2181.673213 | -0.573930381 | 0.120993982 | 3.65E-07 | 5.51E-06 |
| RCCD1 | 2215.520005 | -0.573936139 | 0.086247901 | 5.44E-12 | 2.29E-10 |
| SMTNL2 | 61.82514684 | -0.574152049 | 0.251091875 | 0.00224566 | 0.01040879 |
| DMAC1 | 1269.001398 | -0.574291677 | 0.15571841 | 3.46E-05 | 0.000299327 |
| TPM3P9 | 572.8538777 | -0.574353502 | 0.138563435 | 5.54E-06 | 6.00E-05 |
| CFAP298 | 442.3175587 | -0.574700258 | 0.137921442 | 5.04E-06 | 5.52E-05 |
| PRKAG2-AS1 | 213.7938722 | -0.574802513 | 0.130125643 | 1.67E-06 | 2.10E-05 |
| TUBA1C | 1491.018752 | -0.575022308 | 0.151101908 | 2.19E-05 | 0.00020173 |
| TBX1 | 595.4161301 | -0.575448271 | 0.223946197 | 0.00115173 | 0.005935879 |
| NPPA | 20.94016173 | -0.575632661 | 0.322004232 | 0.00550106 | 0.021700338 |
| PARM1 | 457.032296 | -0.575794353 | 0.173067697 | 0.00012391 | 0.000891523 |
| CCNF | 1912.737198 | -0.575817109 | 0.16917369 | 9.53E-05 | 0.000709386 |
| AQP11 | 168.2592765 | -0.576374657 | 0.174350185 | 0.00013238 | 0.000946186 |
| NUP210P3 | 16.57733584 | -0.576422097 | 0.397789023 | 0.00834163 | 0.030604314 |
| SPIB | 37.84475563 | -0.576923771 | 0.246912926 | 0.00196593 | 0.009319368 |
| LINGO1 | 742.1477258 | -0.57698527 | 0.146413822 | 1.26E-05 | 0.00012483 |
| SZRD1 | 1706.228726 | -0.577073913 | 0.169626599 | 9.49E-05 | 0.000707269 |
| CAMKV | 7405.816163 | -0.577835413 | 0.116589859 | 1.23E-07 | 2.14E-06 |

| | | | | | |
|---|---|---|---|---|---|
| LGSN | 11.82436943 | -0.577883084 | 0.425703045 | 0.00896421 | 0.03252293 |
| INCENP | 4243.625365 | -0.578714999 | 0.14371637 | 8.78E-06 | 8.98E-05 |
| CCIN | 16.79472792 | -0.579603334 | 0.347825319 | 0.00631896 | 0.024381904 |
| NR4A2 | 133.1176672 | -0.580149169 | 0.160329665 | 4.27E-05 | 0.000358324 |
| MIR1244-2 | 435.5705658 | -0.580169238 | 0.166319607 | 6.84E-05 | 0.000533434 |
| NFYA | 4564.771473 | -0.58024079 | 0.142797782 | 7.45E-06 | 7.79E-05 |
| CHRDL1 | 281.6064147 | -0.580278703 | 0.186736901 | 0.00024325 | 0.001584495 |
| TIMELESS | 9342.023957 | -0.58090752 | 0.133116733 | 2.03E-06 | 2.47E-05 |
| PRODHLP | 35.85505344 | -0.580913255 | 0.280649429 | 0.00326944 | 0.014136388 |
| PGM5 | 138.0522477 | -0.58162938 | 0.214051358 | 0.0007448 | 0.004102891 |
| SDC1 | 793.4373101 | -0.581708779 | 0.116201993 | 9.28E-08 | 1.66E-06 |
| ADAMTS5 | 279.0176769 | -0.581756244 | 0.181902627 | 0.0001801 | 0.001231221 |
| ITPRIPL1 | 1094.968109 | -0.581800964 | 0.118363082 | 1.47E-07 | 2.48E-06 |
| MIR573 | 44.23178795 | -0.582215169 | 0.269167746 | 0.00270046 | 0.012103782 |
| TRHDE | 107.8583327 | -0.582276206 | 0.216270342 | 0.00079113 | 0.004305049 |
| B3GNT5 | 757.4938869 | -0.58237767 | 0.192991445 | 0.00031474 | 0.001973858 |
| ARMC6 | 2517.024475 | -0.583016603 | 0.122635199 | 3.23E-07 | 4.96E-06 |
| CELSR2 | 8840.845424 | -0.583057482 | 0.180486006 | 0.00016038 | 0.001116833 |
| NRBP2 | 1143.853821 | -0.584116781 | 0.109323625 | 1.53E-08 | 3.24E-07 |
| TUBA1B | 21600.13761 | -0.584387591 | 0.145477321 | 8.73E-06 | 8.93E-05 |
| NECAB2 | 27.70408783 | -0.584388865 | 0.274069801 | 0.00281525 | 0.012539033 |
| EFS | 1748.686211 | -0.584790539 | 0.12744911 | 7.04E-07 | 9.76E-06 |
| FGFR2 | 863.3383101 | -0.585112703 | 0.144679038 | 7.76E-06 | 8.08E-05 |
| MEGF6 | 140.8036688 | -0.585114333 | 0.207726625 | 0.00055049 | 0.00317093 |
| FGD4 | 1640.805521 | -0.585301325 | 0.140140789 | 4.44E-06 | 4.93E-05 |
| MIR4477B | 98.86519253 | -0.585418447 | 0.190892696 | 0.00026389 | 0.001700095 |
| SPATA33 | 175.0539583 | -0.585421762 | 0.146071899 | 8.99E-06 | 9.17E-05 |
| RN7SKP271 | 13.47229588 | -0.585662691 | 0.395521152 | 0.0075275 | 0.028108845 |
| MIR3164 | 21.13892836 | -0.585828313 | 0.338995574 | 0.00551718 | 0.021726905 |
| KCNIP2 | 216.4232951 | -0.585920753 | 0.140884553 | 4.76E-06 | 5.25E-05 |
| MGAT5 | 3355.506678 | -0.585936317 | 0.128739387 | 8.29E-07 | 1.13E-05 |
| SYNGR4 | 182.6794054 | -0.586025287 | 0.163828008 | 4.73E-05 | 0.000390982 |
| PREX2 | 152.0628674 | -0.586134267 | 0.214832781 | 0.00069469 | 0.003863086 |
| AMH | 3965.325932 | -0.586194722 | 0.114490187 | 4.97E-08 | 9.44E-07 |
| NFIA-AS2 | 12.53062876 | -0.586619499 | 0.410851086 | 0.00788703 | 0.029202159 |
| MCM2 | 6910.697976 | -0.586702814 | 0.13929215 | 3.77E-06 | 4.28E-05 |
| USP46-DT | 83.21794353 | -0.586999231 | 0.193234973 | 0.00028438 | 0.001806573 |
| AIF1L | 668.8687528 | -0.587250621 | 0.14124587 | 4.74E-06 | 5.23E-05 |
| CYYR1 | 1447.089315 | -0.587608691 | 0.123647579 | 3.14E-07 | 4.85E-06 |
| CMTM3 | 607.9703654 | -0.587739432 | 0.115724322 | 6.09E-08 | 1.13E-06 |
| CCDC188 | 522.7389675 | -0.588083121 | 0.114943824 | 4.99E-08 | 9.48E-07 |
| TPI1P1 | 149.2629889 | -0.588156639 | 0.199401499 | 0.00036634 | 0.002248088 |
| SF1-DT | 36.68106256 | -0.588206724 | 0.290351438 | 0.00332972 | 0.01435178 |
| PCDHB8 | 118.6630741 | -0.588278674 | 0.236187172 | 0.00124619 | 0.006347957 |
| LINC03015 | 468.5121023 | -0.588920895 | 0.186175283 | 0.00018924 | 0.001280081 |
| APCDD1 | 209.6984888 | -0.589182128 | 0.154495901 | 1.89E-05 | 0.000178248 |
| TMEM132C | 695.4423151 | -0.589263306 | 0.203483074 | 0.00042409 | 0.002549238 |
| RNU6-415P | 27.95574755 | -0.589746309 | 0.339520741 | 0.00527659 | 0.020996103 |
| PKMYT1 | 3073.528573 | -0.590656405 | 0.144087794 | 5.89E-06 | 6.33E-05 |
| FBRSL1 | 6119.412898 | -0.590788791 | 0.148964943 | 1.02E-05 | 0.000102844 |

| | | | | | |
|---|---|---|---|---|---|
| NT5DC3 | 1731.226657 | -0.591201895 | 0.145125799 | 6.53E-06 | 6.93E-05 |
| LINC01797 | 2362.418028 | -0.591360806 | 0.141641795 | 4.25E-06 | 4.75E-05 |
| RRS1 | 891.6056234 | -0.591826129 | 0.138667562 | 2.84E-06 | 3.34E-05 |
| TUSC1 | 284.0443395 | -0.592053015 | 0.143612164 | 5.28E-06 | 5.75E-05 |
| MPPED1 | 95.59012136 | -0.592240356 | 0.176841711 | 9.99E-05 | 0.000738123 |
| CFAP410 | 438.9708495 | -0.592288084 | 0.124983778 | 3.23E-07 | 4.96E-06 |
| COX16 | 225.1055367 | -0.592359243 | 0.166250623 | 4.71E-05 | 0.000390008 |
| LDOC1 | 781.5343762 | -0.592781298 | 0.135493266 | 1.75E-06 | 2.19E-05 |
| RUNX1T1 | 4536.308576 | -0.59303572 | 0.114283464 | 3.27E-08 | 6.46E-07 |
| RPS6P25 | 70.44090548 | -0.593150031 | 0.233806259 | 0.00106656 | 0.005561011 |
| C1orf216 | 2012.545603 | -0.593237256 | 0.109034575 | 8.34E-09 | 1.86E-07 |
| METTL1 | 736.1018614 | -0.594028501 | 0.127218277 | 4.45E-07 | 6.62E-06 |
| NEUROG1 | 7.867632432 | -0.594069414 | 0.562875428 | 0.00928448 | 0.033472305 |
| MANEAL | 1830.92327 | -0.594129018 | 0.13809054 | 2.39E-06 | 2.87E-05 |
| UBAC2-AS1 | 51.64797111 | -0.594302588 | 0.242118107 | 0.00128674 | 0.006514208 |
| COPG2IT1 | 627.3719279 | -0.594488094 | 0.157958468 | 2.19E-05 | 0.00020173 |
| ZIC5 | 1387.765872 | -0.595098432 | 0.155310456 | 1.68E-05 | 0.00016026 |
| SHF | 2584.352429 | -0.595530073 | 0.158958321 | 2.32E-05 | 0.000212107 |
| ADAMTS14 | 296.8245788 | -0.596013031 | 0.137791658 | 2.13E-06 | 2.58E-05 |
| TLCD1 | 66.66497065 | -0.596299599 | 0.216646806 | 0.00059441 | 0.003381321 |
| RPL39P16 | 70.57125295 | -0.597267624 | 0.232984608 | 0.00096241 | 0.005095542 |
| NCS1 | 7721.520564 | -0.597283862 | 0.16780429 | 4.58E-05 | 0.000379861 |
| MVB12B | 2151.656342 | -0.59972597 | 0.165746242 | 3.61E-05 | 0.000309863 |
| ARHGAP33 | 2775.795072 | -0.599891274 | 0.126218199 | 2.84E-07 | 4.42E-06 |
| CENPV | 6947.15916 | -0.600374341 | 0.098646502 | 1.78E-10 | 5.54E-09 |
| ZNF414 | 439.8064743 | -0.600402261 | 0.171382049 | 5.44E-05 | 0.000437384 |
| LRFN3 | 621.5609895 | -0.600491619 | 0.203298524 | 0.00032411 | 0.002021545 |
| TAF6L | 899.8043868 | -0.600814753 | 0.091737787 | 9.03E-12 | 3.64E-10 |
| TP73 | 52.64309199 | -0.600909702 | 0.349142761 | 0.00493648 | 0.019900754 |
| ENTPD1 | 1162.350957 | -0.601000909 | 0.161287401 | 2.39E-05 | 0.000217806 |
| CNKSR1 | 62.12179241 | -0.601454777 | 0.218436861 | 0.00056577 | 0.003240801 |
| PGAP3 | 373.4245193 | -0.601723866 | 0.13927727 | 2.08E-06 | 2.52E-05 |
| ADAMTS8 | 201.6243041 | -0.601944819 | 0.172136683 | 5.49E-05 | 0.000440325 |
| RUNX3 | 14.88040858 | -0.602083683 | 0.413942651 | 0.0068491 | 0.025989354 |
| FGF5 | 56.90085052 | -0.60221025 | 0.242648453 | 0.00112363 | 0.005817277 |
| SACS-AS1 | 45.99352101 | -0.602284742 | 0.247802762 | 0.00126754 | 0.006435544 |
| NCAN | 2573.88713 | -0.602304519 | 0.131226644 | 6.06E-07 | 8.59E-06 |
| LZTS1 | 1537.743768 | -0.602405412 | 0.300423038 | 0.00306196 | 0.013437057 |
| MAPK6P3 | 8.129442908 | -0.602426049 | 0.494510727 | 0.00831997 | 0.030553293 |
| TRHDE-AS1 | 132.7284789 | -0.602559607 | 0.238407499 | 0.00100309 | 0.005274252 |
| LINC01315 | 145.1124203 | -0.603258775 | 0.170954586 | 4.85E-05 | 0.000398655 |
| RBM14 | 4534.671275 | -0.603481317 | 0.105351714 | 1.50E-09 | 3.85E-08 |
| SEMA3G | 92.63820731 | -0.603651513 | 0.225900298 | 0.00068856 | 0.003835179 |
| CDH6 | 263.9332122 | -0.603683736 | 0.225710553 | 0.00068407 | 0.003818437 |
| RPL5P18 | 12.04126843 | -0.60386946 | 0.460067919 | 0.00771885 | 0.028708705 |
| SPC24 | 1479.971413 | -0.603925585 | 0.105256461 | 1.41E-09 | 3.66E-08 |
| ATP5MC1 | 4070.244396 | -0.604169484 | 0.151844684 | 8.63E-06 | 8.85E-05 |
| CTSF | 1219.549969 | -0.604192023 | 0.153310533 | 1.01E-05 | 0.000101591 |
| GAS1 | 341.5209 | -0.606289117 | 0.141870612 | 2.45E-06 | 2.93E-05 |
| C4orf3 | 1520.497746 | -0.606518122 | 0.167132071 | 3.27E-05 | 0.000285028 |

| | | | | | |
|---|---|---|---|---|---|
| LIMD2 | 5038.084249 | -0.606617276 | 0.120451744 | 6.51E-08 | 1.20E-06 |
| C8orf82 | 878.7000661 | -0.606687835 | 0.167068071 | 3.24E-05 | 0.000282565 |
| CLN6 | 3397.16162 | -0.606725521 | 0.083820926 | 6.93E-14 | 3.87E-12 |
| SH2B2 | 380.5759022 | -0.606752571 | 0.155634362 | 1.16E-05 | 0.000116122 |
| VEGFD | 3362.716614 | -0.606838028 | 0.210499297 | 0.00037579 | 0.002300599 |
| CEROX1 | 84.60422434 | -0.606930794 | 0.290463423 | 0.0025153 | 0.011400623 |
| PLSCR1 | 1094.837448 | -0.607479067 | 0.157977456 | 1.43E-05 | 0.00013917 |
| ADAM11 | 1470.022991 | -0.607751142 | 0.110271748 | 5.00E-09 | 1.16E-07 |
| NPTX1 | 795.5105058 | -0.607836283 | 0.216459224 | 0.00045963 | 0.002724678 |
| ISOC2 | 642.530405 | -0.608133882 | 0.106314506 | 1.51E-09 | 3.88E-08 |
| TUBB3 | 263.4011568 | -0.608299569 | 0.149862579 | 5.99E-06 | 6.42E-05 |
| RCN3 | 165.9937422 | -0.609135818 | 0.172266256 | 4.48E-05 | 0.000372609 |
| PCLAF | 3109.116258 | -0.610085588 | 0.15604625 | 1.08E-05 | 0.000108483 |
| SLC40A1 | 113.5205489 | -0.610202553 | 0.211517211 | 0.00036231 | 0.002227369 |
| B4GALT2 | 2928.582515 | -0.610549114 | 0.146791752 | 3.86E-06 | 4.36E-05 |
| RHBDL3 | 824.2395191 | -0.612163817 | 0.154229811 | 8.37E-06 | 8.63E-05 |
| TFAP4 | 1298.121211 | -0.612600309 | 0.116887005 | 2.12E-08 | 4.34E-07 |
| ACYP1 | 738.9036394 | -0.612753916 | 0.130439276 | 3.33E-07 | 5.08E-06 |
| EDDM13 | 46.50204921 | -0.61310795 | 0.242195121 | 0.00090298 | 0.004824327 |
| GTF3C2 | 3358.312673 | -0.613539763 | 0.132804065 | 4.78E-07 | 7.02E-06 |
| PLD6 | 1231.173419 | -0.613635257 | 0.113156285 | 7.80E-09 | 1.74E-07 |
| RAB4A-AS1 | 719.7005162 | -0.613869773 | 0.175015551 | 4.75E-05 | 0.000392488 |
| CERK | 12376.55287 | -0.61427663 | 0.11187733 | 5.33E-09 | 1.23E-07 |
| KIF19 | 45.84836673 | -0.614362791 | 0.343786051 | 0.00401501 | 0.016777511 |
| GINS4 | 922.6229575 | -0.614727408 | 0.093570665 | 7.06E-12 | 2.88E-10 |
| TKFC | 980.8345834 | -0.614954763 | 0.117922076 | 2.38E-08 | 4.84E-07 |
| MTA1-DT | 232.6780341 | -0.614973979 | 0.151216938 | 5.47E-06 | 5.93E-05 |
| B4GALNT1 | 2399.270788 | -0.614974834 | 0.128198064 | 2.02E-07 | 3.29E-06 |
| CDK5R1 | 2223.391135 | -0.61501616 | 0.13888705 | 1.14E-06 | 1.50E-05 |
| SNRNP35 | 631.161491 | -0.615522957 | 0.140427562 | 1.39E-06 | 1.79E-05 |
| RAC3 | 2302.242146 | -0.616285342 | 0.098030219 | 4.43E-11 | 1.57E-09 |
| FAM78B | 2294.439489 | -0.616358927 | 0.147182126 | 3.25E-06 | 3.75E-05 |
| PRSS12 | 3744.036105 | -0.616452206 | 0.110074715 | 2.81E-09 | 6.89E-08 |
| CLUH | 5129.935029 | -0.616553202 | 0.169769869 | 2.96E-05 | 0.000262032 |
| PCBP1 | 671.7160957 | -0.616684245 | 0.183338406 | 7.63E-05 | 0.000586062 |
| C1GALT1C1 | 240.8544824 | -0.616686344 | 0.194163832 | 0.00014149 | 0.001002934 |
| BHLHE22 | 295.8413464 | -0.616795522 | 0.182511892 | 7.23E-05 | 0.000559138 |
| CDK2 | 2707.320372 | -0.616853387 | 0.133454336 | 4.59E-07 | 6.79E-06 |
| COL4A2 | 5557.812256 | -0.617210748 | 0.125570985 | 1.10E-07 | 1.93E-06 |
| CNIH2 | 1012.86101 | -0.617388861 | 0.182371941 | 7.05E-05 | 0.000548208 |
| H2AC20 | 56.88729167 | -0.617401981 | 0.264487748 | 0.00137909 | 0.006893432 |
| HBQ1 | 28.86919009 | -0.617453916 | 0.376516057 | 0.0048702 | 0.019679837 |
| USP2-AS1 | 117.7692981 | -0.618011447 | 0.194978748 | 0.00014268 | 0.001010314 |
| ANKRD9 | 438.486319 | -0.618297428 | 0.129897296 | 2.35E-07 | 3.76E-06 |
| SLC25A39 | 9032.028319 | -0.619271079 | 0.099509089 | 6.46E-11 | 2.21E-09 |
| SLC39A4 | 184.3087868 | -0.619481642 | 0.153181894 | 5.77E-06 | 6.21E-05 |
| MXRA8 | 139.1945045 | -0.619553784 | 0.19100695 | 0.00011085 | 0.000809164 |
| WASL-DT | 161.1943682 | -0.619845189 | 0.182318085 | 6.56E-05 | 0.000514522 |
| CPXM1 | 4824.952527 | -0.620087455 | 0.090375578 | 9.24E-13 | 4.33E-11 |
| TMSB4X | 13621.60936 | -0.620107271 | 0.189611932 | 0.00010102 | 0.000745363 |

| | | | | | |
|---|---|---|---|---|---|
| HNRNPA0 | 18656.14103 | -0.620574362 | 0.127578762 | 1.38E-07 | 2.35E-06 |
| RSKR | 1696.145995 | -0.620928876 | 0.099903033 | 6.70E-11 | 2.27E-09 |
| SEMA6A | 1075.355492 | -0.621200554 | 0.149132695 | 3.42E-06 | 3.92E-05 |
| HUNK | 1315.603663 | -0.621354706 | 0.137491417 | 7.14E-07 | 9.89E-06 |
| FAM81A | 60.79795569 | -0.621372027 | 0.289855145 | 0.00199657 | 0.009436308 |
| ZBTB12 | 1139.150807 | -0.622559285 | 0.09987166 | 5.89E-11 | 2.04E-09 |
| BRI3BP | 2891.662753 | -0.622704392 | 0.099079362 | 4.24E-11 | 1.51E-09 |
| COL4A1 | 1570.565914 | -0.622964539 | 0.108367632 | 1.13E-09 | 3.00E-08 |
| GABRA5 | 844.7506549 | -0.622995368 | 0.137971617 | 7.16E-07 | 9.90E-06 |
| HSF4 | 1266.20891 | -0.623174571 | 0.135392087 | 4.76E-07 | 6.99E-06 |
| GFRA2 | 10897.52382 | -0.623240625 | 0.122840797 | 4.66E-08 | 8.91E-07 |
| XRCC2 | 4592.074038 | -0.623337197 | 0.129727249 | 1.80E-07 | 2.98E-06 |
| COA4 | 1444.437134 | -0.623959925 | 0.156640338 | 7.10E-06 | 7.47E-05 |
| CCND1 | 32809.56216 | -0.624280564 | 0.132608433 | 2.86E-07 | 4.46E-06 |
| RPP25 | 192.4357094 | -0.624312421 | 0.135307541 | 4.47E-07 | 6.64E-06 |
| ZBTB46 | 594.2585571 | -0.624788626 | 0.184129017 | 6.40E-05 | 0.000504254 |
| FAM20B | 5263.428677 | -0.624824818 | 0.103463631 | 1.95E-10 | 6.02E-09 |
| SLC24A2 | 84.38669283 | -0.625100986 | 0.2175593 | 0.00032651 | 0.002034045 |
| DUSP5P1 | 317.4579388 | -0.62542883 | 0.14239482 | 1.23E-06 | 1.60E-05 |
| RNASET2 | 271.5214819 | -0.625649822 | 0.127768939 | 1.12E-07 | 1.96E-06 |
| CTSC | 2709.833159 | -0.62578542 | 0.125332525 | 6.89E-08 | 1.27E-06 |
| HK3 | 49.02268869 | -0.625936942 | 0.275633282 | 0.00147063 | 0.007267955 |
| RAB11B | 2254.69092 | -0.626044575 | 0.139061227 | 7.41E-07 | 1.02E-05 |
| PIF1 | 2758.067092 | -0.626297939 | 0.09674632 | 1.21E-11 | 4.77E-10 |
| APRT | 536.7800201 | -0.6267576 | 0.119509601 | 1.84E-08 | 3.83E-07 |
| RGS19 | 370.9526557 | -0.627056023 | 0.128843148 | 1.28E-07 | 2.21E-06 |
| JRK | 4231.940846 | -0.627659103 | 0.093752855 | 2.73E-12 | 1.22E-10 |
| MATCAP1 | 3858.467663 | -0.628164519 | 0.077219884 | 5.41E-17 | 4.80E-15 |
| CHRNB4 | 3287.643606 | -0.628518143 | 0.166149848 | 1.50E-05 | 0.000145413 |
| TMEM177 | 683.3230977 | -0.628860767 | 0.127875138 | 9.81E-08 | 1.74E-06 |
| LAMA1 | 1657.64426 | -0.629341571 | 0.138956957 | 6.35E-07 | 8.97E-06 |
| APBA2 | 4104.135487 | -0.629794041 | 0.108087834 | 6.72E-10 | 1.86E-08 |
| EMB | 406.3588456 | -0.630594691 | 0.173220185 | 2.52E-05 | 0.000228089 |
| LINC01285 | 24.7289306 | -0.630840025 | 0.321548684 | 0.00261367 | 0.01175835 |
| GCNT2 | 636.1172343 | -0.630852101 | 0.150841031 | 2.92E-06 | 3.42E-05 |
| HNRNPA1P16 | 82.68809058 | -0.631208099 | 0.216931866 | 0.00027781 | 0.001769797 |
| C1orf159 | 529.4286554 | -0.631322365 | 0.118321187 | 1.08E-08 | 2.35E-07 |
| TBC1D30 | 654.3314891 | -0.631335757 | 0.166919671 | 1.47E-05 | 0.000142548 |
| UBTF | 11482.44795 | -0.631381475 | 0.147311618 | 1.86E-06 | 2.29E-05 |
| HSPD1P6 | 29.83271323 | -0.631574801 | 0.323279717 | 0.0026372 | 0.011851247 |
| FIZ1 | 932.6042441 | -0.632431278 | 0.135236452 | 3.09E-07 | 4.77E-06 |
| B3GALT4 | 29.40222567 | -0.632853593 | 0.367474689 | 0.00384539 | 0.016180994 |
| B3GALT6 | 610.4199397 | -0.633115818 | 0.139414933 | 5.80E-07 | 8.30E-06 |
| SRPK3 | 238.5908056 | -0.633340565 | 0.190248456 | 7.35E-05 | 0.000567185 |
| PFAS | 7221.986144 | -0.633908098 | 0.140815848 | 6.90E-07 | 9.60E-06 |
| PIGW | 1201.211829 | -0.634200202 | 0.120575798 | 1.59E-08 | 3.33E-07 |
| CHODL | 190.394787 | -0.634414322 | 0.220427951 | 0.00029528 | 0.001862607 |
| EXOSC4 | 26.79748027 | -0.634515972 | 0.319885757 | 0.00243729 | 0.011108258 |
| TCF3 | 8304.746588 | -0.634907586 | 0.18615315 | 5.49E-05 | 0.00043999 |
| SH2D5 | 219.641649 | -0.63492769 | 0.147984276 | 1.76E-06 | 2.20E-05 |

| | | | | | |
|---|---|---|---|---|---|
| CHRM1 | 272.8480268 | -0.635050436 | 0.157623745 | 5.32E-06 | 5.79E-05 |
| NIBAN2 | 1518.885996 | -0.635444862 | 0.140832978 | 6.49E-07 | 9.12E-06 |
| MIR222HG | 28.31356922 | -0.63604088 | 0.29688887 | 0.00176651 | 0.00849087 |
| HS3ST4 | 614.8474541 | -0.636157684 | 0.13354827 | 1.96E-07 | 3.22E-06 |
| GM2A | 459.3873681 | -0.636602636 | 0.117906415 | 7.28E-09 | 1.64E-07 |
| PTENP1-AS | 119.1545867 | -0.638186811 | 0.162999073 | 8.18E-06 | 8.46E-05 |
| POLD1 | 4462.746638 | -0.638544969 | 0.126445341 | 4.59E-08 | 8.79E-07 |
| CORO1A | 685.9689202 | -0.638615628 | 0.109018053 | 5.16E-10 | 1.47E-08 |
| EME2 | 1207.267418 | -0.63865017 | 0.116607915 | 4.65E-09 | 1.09E-07 |
| NEPNP | 19.13114292 | -0.639299879 | 0.373621057 | 0.00372593 | 0.015758944 |
| NXPH2 | 12.73111439 | -0.63977577 | 0.513322957 | 0.00640659 | 0.024648297 |
| SBK1 | 4268.597522 | -0.640494682 | 0.172672628 | 1.78E-05 | 0.000168753 |
| CIB2 | 598.7851677 | -0.64067148 | 0.115889018 | 3.42E-09 | 8.25E-08 |
| ZNF813 | 358.3980544 | -0.640693371 | 0.185212048 | 4.39E-05 | 0.000366491 |
| CYP1B1 | 60.06648176 | -0.640749788 | 0.215056282 | 0.00020702 | 0.001378056 |
| HMG20B | 1123.756746 | -0.64082553 | 0.088304514 | 4.55E-14 | 2.60E-12 |
| CAVIN4 | 1061.491953 | -0.640968142 | 0.13977242 | 4.38E-07 | 6.53E-06 |
| H2AZ2 | 11086.86964 | -0.641226887 | 0.140123124 | 4.57E-07 | 6.77E-06 |
| RN7SKP16 | 19.65193692 | -0.641895265 | 0.4343991 | 0.00503619 | 0.02022342 |
| KNOP1 | 1835.462846 | -0.642036819 | 0.102443598 | 4.01E-11 | 1.44E-09 |
| KIF18B | 7995.626681 | -0.642119535 | 0.090776826 | 1.70E-13 | 8.91E-12 |
| SLC29A1 | 5670.90059 | -0.642288022 | 0.103523423 | 5.97E-11 | 2.06E-09 |
| CAMSAP1-DT | 137.4967834 | -0.643344191 | 0.183452433 | 3.62E-05 | 0.000309984 |
| DEXI | 254.8603243 | -0.643737594 | 0.170219632 | 1.31E-05 | 0.000128622 |
| HSPA12B | 51.36642557 | -0.643843741 | 0.262191273 | 0.00082264 | 0.004447087 |
| NKAIN1 | 2713.274903 | -0.643915634 | 0.190414398 | 5.56E-05 | 0.000444647 |
| POLR3H | 1878.60919 | -0.644443405 | 0.135995115 | 2.05E-07 | 3.34E-06 |
| FDXACB1 | 75.92303665 | -0.645890683 | 0.226817572 | 0.00028925 | 0.001833036 |
| MT-TK | 22.06351893 | -0.646128235 | 0.362733447 | 0.00316699 | 0.013794971 |
| POU2F2 | 900.0668482 | -0.646397884 | 0.12703746 | 3.50E-08 | 6.87E-07 |
| GPRIN1 | 4268.162186 | -0.647238611 | 0.149289895 | 1.29E-06 | 1.68E-05 |
| LRRN1 | 3392.998003 | -0.648919146 | 0.134062547 | 1.20E-07 | 2.08E-06 |
| GINS2 | 1221.87546 | -0.649165968 | 0.158020823 | 3.35E-06 | 3.86E-05 |
| SRSF9P1 | 8.03374391 | -0.649175083 | 0.524612219 | 0.00608196 | 0.023589104 |
| MARCKS | 20364.46442 | -0.649597647 | 0.105760443 | 8.24E-11 | 2.74E-09 |
| STAB2 | 32.23667061 | -0.649900059 | 0.290010503 | 0.00127517 | 0.006468332 |
| MAP2K6 | 2138.95115 | -0.650228164 | 0.184839477 | 3.26E-05 | 0.000283904 |
| TMEM108 | 3230.460072 | -0.651492523 | 0.105885106 | 7.57E-11 | 2.53E-09 |
| LINC01224 | 1560.923617 | -0.652210583 | 0.147025726 | 7.83E-07 | 1.07E-05 |
| EFNA4 | 26.08531362 | -0.652341271 | 0.343288497 | 0.00246117 | 0.011197241 |
| TUBB6 | 1625.936503 | -0.653015428 | 0.125424137 | 1.77E-08 | 3.69E-07 |
| HS3ST5 | 309.5606477 | -0.65332381 | 0.156842119 | 2.53E-06 | 3.01E-05 |
| MFAP2 | 192.0472746 | -0.653989306 | 0.18269872 | 2.52E-05 | 0.000227852 |
| ADM5 | 106.0488145 | -0.654663437 | 0.230308756 | 0.00027027 | 0.001733894 |
| ARX | 2.939283477 | -0.655420945 | 0.87646391 | 0.00463987 | 0.018898287 |
| LYPD6 | 262.4953676 | -0.65547181 | 0.17840392 | 1.75E-05 | 0.000166815 |
| COL26A1 | 56.73067495 | -0.655865664 | 0.232353613 | 0.00028265 | 0.001797272 |
| TPBGL | 290.8820227 | -0.65593604 | 0.436497879 | 0.00443943 | 0.018218415 |
| CCDC85B | 796.304088 | -0.656483442 | 0.155863051 | 2.01E-06 | 2.45E-05 |
| MGST1 | 188.072034 | -0.656787278 | 0.20455599 | 8.66E-05 | 0.000652206 |

| | | | | | |
|---|---|---|---|---|---|
| SLC15A3 | 63.48546369 | -0.657181336 | 0.244675689 | 0.00040631 | 0.002451697 |
| ABCF1 | 7314.755927 | -0.65724714 | 0.146513068 | 5.94E-07 | 8.45E-06 |
| MARVELD1 | 2026.010712 | -0.657542478 | 0.150610751 | 1.02E-06 | 1.36E-05 |
| PDE5A | 1662.139036 | -0.65788233 | 0.168077016 | 6.79E-06 | 7.17E-05 |
| PCDHB16 | 436.9943745 | -0.658102695 | 0.195084601 | 4.99E-05 | 0.000407425 |
| IGSF1 | 69.73779182 | -0.658247469 | 0.270122341 | 0.00075704 | 0.004157049 |
| MPDU1-AS1 | 125.5941675 | -0.658914785 | 0.192497173 | 4.17E-05 | 0.000351174 |
| ALDH1B1 | 1762.35885 | -0.659189008 | 0.141259661 | 2.51E-07 | 3.97E-06 |
| PLPPR1 | 738.5548767 | -0.659471138 | 0.155337316 | 1.69E-06 | 2.12E-05 |
| CDK3 | 103.9181648 | -0.660194102 | 0.17496918 | 1.15E-05 | 0.000114866 |
| NICOL1 | 462.1652906 | -0.660439755 | 0.147329663 | 5.85E-07 | 8.35E-06 |
| H1-10 | 5131.690015 | -0.660973703 | 0.159606566 | 2.60E-06 | 3.08E-05 |
| RIPPLY2 | 256.555584 | -0.661034106 | 0.146161379 | 4.84E-07 | 7.09E-06 |
| RCC1 | 3710.112399 | -0.661113592 | 0.119690479 | 2.90E-09 | 7.08E-08 |
| BOLA2P2 | 48.59894777 | -0.661587713 | 0.241649437 | 0.0003391 | 0.002101663 |
| RAB1B | 1492.444762 | -0.661898415 | 0.13220206 | 4.59E-08 | 8.79E-07 |
| ST6GAL1 | 3355.046468 | -0.661949785 | 0.147735993 | 5.82E-07 | 8.32E-06 |
| ASB13 | 1276.576734 | -0.662199449 | 0.131937964 | 4.29E-08 | 8.31E-07 |
| TSPEAR-AS2 | 26.46673275 | -0.662253123 | 0.333229782 | 0.00193674 | 0.009210731 |
| BDH1 | 2226.32979 | -0.662341571 | 0.112797007 | 3.80E-10 | 1.12E-08 |
| TENT5B | 40.81459865 | -0.662842377 | 0.344249061 | 0.00216679 | 0.010112979 |
| GTF3C2-AS2 | 536.7758418 | -0.663792177 | 0.105697574 | 3.02E-11 | 1.10E-09 |
| SLC19A3 | 108.0147905 | -0.664784673 | 0.201362989 | 5.95E-05 | 0.000473117 |
| MPST | 2449.207691 | -0.664953115 | 0.104262733 | 1.59E-11 | 6.14E-10 |
| ANKFN1 | 149.3531291 | -0.665514421 | 0.226697057 | 0.00018637 | 0.00126399 |
| TRABD | 1626.347244 | -0.665664222 | 0.089551411 | 9.74E-15 | 6.01E-13 |
| TMEM160 | 417.9148438 | -0.666040409 | 0.127341405 | 1.37E-08 | 2.94E-07 |
| DSTNP1 | 84.22304126 | -0.666536493 | 0.222343218 | 0.0001534 | 0.001074344 |
| TMEM158 | 98.14673523 | -0.666566819 | 0.252146899 | 0.00041713 | 0.002511839 |
| LINC01625 | 7.28567749 | -0.666739921 | 0.610597944 | 0.00593321 | 0.023116818 |
| DND1P1 | 65.26163929 | -0.667159177 | 0.251956826 | 0.00041015 | 0.002471956 |
| MUC19 | 153.0239356 | -0.667676709 | 0.178590998 | 1.22E-05 | 0.00012093 |
| GLI2 | 2663.293574 | -0.667768884 | 0.132014362 | 3.32E-08 | 6.54E-07 |
| ING5 | 3443.009562 | -0.668024874 | 0.094090434 | 1.11E-13 | 5.99E-12 |
| NRXN1 | 863.52672 | -0.668128232 | 0.180487353 | 1.39E-05 | 0.000136401 |
| COLEC12 | 1456.795785 | -0.668154424 | 0.173196653 | 7.66E-06 | 8.00E-05 |
| ATP8B1-AS1 | 94.4481584 | -0.669747232 | 0.221404636 | 0.00013699 | 0.000975094 |
| AURKB | 2934.426609 | -0.669974546 | 0.114234229 | 3.68E-10 | 1.09E-08 |
| KCNK5 | 139.0591541 | -0.670126994 | 0.227982949 | 0.00017646 | 0.001210349 |
| DDC | 1463.369423 | -0.67104313 | 0.138926857 | 1.01E-07 | 1.79E-06 |
| PIANP | 1769.363613 | -0.673115854 | 0.134941818 | 4.51E-08 | 8.66E-07 |
| CPNE7 | 113.9989514 | -0.673512878 | 0.188843252 | 2.18E-05 | 0.000200583 |
| TTPA | 21.25758522 | -0.675162085 | 0.378994958 | 0.00255537 | 0.011551668 |
| CKLF | 38.46740022 | -0.675255481 | 0.278982054 | 0.00067839 | 0.00379294 |
| PCDHB3 | 188.6808232 | -0.67544425 | 0.174532012 | 6.79E-06 | 7.17E-05 |
| TMEM19 | 1289.774378 | -0.676202371 | 0.160491892 | 1.64E-06 | 2.07E-05 |
| DPYSL5 | 7091.172982 | -0.676544743 | 0.143311514 | 1.63E-07 | 2.72E-06 |
| JUP | 735.7829354 | -0.677277254 | 0.179931422 | 1.00E-05 | 0.000101577 |
| HNRNPA1P35 | 35.08309331 | -0.677928215 | 0.26733795 | 0.00049765 | 0.002917055 |
| LNMICC | 52.78825974 | -0.678378628 | 0.234596823 | 0.0001877 | 0.001271352 |

| | | | | | |
|---|---|---|---|---|---|
| MLLT11 | 14619.38624 | -0.678776015 | 0.152917555 | 5.93E-07 | 8.44E-06 |
| FGFRL1 | 571.1580968 | -0.679142086 | 0.375914306 | 0.00237706 | 0.010884935 |
| ILF3-DT | 607.0814755 | -0.679202162 | 0.129997987 | 1.24E-08 | 2.68E-07 |
| RNF145 | 7719.119303 | -0.679288165 | 0.154263572 | 6.92E-07 | 9.62E-06 |
| ARHGAP22 | 595.5706435 | -0.679783042 | 0.222477808 | 0.00011329 | 0.000824037 |
| ALDH16A1 | 983.9937865 | -0.679824292 | 0.113923499 | 1.80E-10 | 5.60E-09 |
| IZUMO4 | 61.69638725 | -0.679927272 | 0.237289268 | 0.00019964 | 0.001337685 |
| ZIC2 | 3414.094162 | -0.68140481 | 0.107792777 | 1.94E-11 | 7.38E-10 |
| PROKR1 | 106.2236256 | -0.681467008 | 0.273584743 | 0.00054027 | 0.003122578 |
| DCTPP1 | 2198.281291 | -0.681589928 | 0.131985502 | 1.67E-08 | 3.49E-07 |
| LMNB1 | 11477.71709 | -0.682270266 | 0.129875797 | 1.03E-08 | 2.25E-07 |
| FBLN2 | 555.6903731 | -0.682990425 | 0.189664049 | 1.74E-05 | 0.000166244 |
| TMEM109 | 925.2157945 | -0.68306601 | 0.159758816 | 1.17E-06 | 1.53E-05 |
| IPO5P1 | 1161.270689 | -0.684365075 | 0.125918508 | 3.76E-09 | 8.97E-08 |
| LRRTM1 | 280.3868689 | -0.685179579 | 0.193073918 | 2.06E-05 | 0.000191864 |
| LINC01918 | 69.19361427 | -0.685213737 | 0.225605561 | 0.00011375 | 0.000826532 |
| SMOC1 | 188.8038817 | -0.685237202 | 0.201918625 | 3.56E-05 | 0.000306146 |
| DUBR | 342.5950779 | -0.686003334 | 0.147384881 | 2.03E-07 | 3.31E-06 |
| VWA1 | 52.64796047 | -0.686419593 | 0.256741143 | 0.00032131 | 0.002005894 |
| TMEM120B | 2548.646048 | -0.686696803 | 0.111896266 | 5.91E-11 | 2.05E-09 |
| GLDC | 921.4686222 | -0.687228245 | 0.18520394 | 1.11E-05 | 0.000111325 |
| PPIF | 4042.938057 | -0.687757722 | 0.162742666 | 1.38E-06 | 1.78E-05 |
| RNA5SP18 | 100.5781385 | -0.688396302 | 0.191089144 | 1.64E-05 | 0.000157385 |
| C10orf53 | 189.1549119 | -0.688848267 | 0.198066344 | 2.56E-05 | 0.000231245 |
| SETD6 | 2171.162165 | -0.688933717 | 0.11006298 | 2.67E-11 | 9.89E-10 |
| F2R | 3279.375881 | -0.690275222 | 0.132395287 | 1.17E-08 | 2.54E-07 |
| PHETA1 | 373.7409279 | -0.691096703 | 0.155196007 | 4.90E-07 | 7.16E-06 |
| TMEM132A | 15123.50185 | -0.69115179 | 0.11372991 | 8.17E-11 | 2.72E-09 |
| MTND2P40 | 56.48784209 | -0.69180329 | 0.248060111 | 0.00022239 | 0.001468199 |
| BCL7A | 4093.433336 | -0.692359885 | 0.153708431 | 3.82E-07 | 5.75E-06 |
| SCARA3 | 913.2176354 | -0.692381453 | 0.126542563 | 2.82E-09 | 6.91E-08 |
| MIR7156 | 2.739557286 | -0.693321975 | 0.916356433 | 0.00379298 | 0.016012901 |
| RENO1 | 427.5100214 | -0.694029642 | 0.17968984 | 5.78E-06 | 6.22E-05 |
| TMSB4XP8 | 204.045214 | -0.695216968 | 0.180834329 | 6.13E-06 | 6.55E-05 |
| WDR81 | 1613.224454 | -0.696682261 | 0.153991657 | 3.34E-07 | 5.09E-06 |
| LINC00997 | 162.4271692 | -0.697846993 | 0.180052108 | 5.27E-06 | 5.75E-05 |
| FGF22 | 40.81373868 | -0.698177385 | 0.2540188 | 0.00023492 | 0.001538522 |
| NUDT14 | 290.3203255 | -0.698367902 | 0.130378512 | 4.99E-09 | 1.16E-07 |
| DPP3 | 1179.38802 | -0.699020277 | 0.161263435 | 7.65E-07 | 1.05E-05 |
| SCRT2 | 862.7585708 | -0.699816845 | 0.178012307 | 4.14E-06 | 4.64E-05 |
| SLC7A2 | 1450.381303 | -0.700318196 | 0.167784879 | 1.51E-06 | 1.93E-05 |
| KCNAB3 | 595.6512452 | -0.700462804 | 0.099071281 | 9.84E-14 | 5.37E-12 |
| VAV3 | 271.3488183 | -0.700777127 | 0.168993662 | 1.69E-06 | 2.12E-05 |
| LINC01116 | 46.05529351 | -0.700884027 | 0.28401552 | 0.00048377 | 0.002848846 |
| SFXN2 | 1378.577596 | -0.700947646 | 0.150322568 | 1.67E-07 | 2.78E-06 |
| DCC | 236.4014491 | -0.701072082 | 0.242134468 | 0.00014996 | 0.001054284 |
| SMPD5 | 52.78462842 | -0.701749851 | 0.253797693 | 0.00021709 | 0.001437787 |
| SPRYD4 | 579.8671612 | -0.701825932 | 0.121321821 | 4.25E-10 | 1.23E-08 |
| B4GALT5 | 4135.317912 | -0.702028726 | 0.138553251 | 2.23E-08 | 4.54E-07 |
| COLGALT1 | 2943.874802 | -0.70206282 | 0.110874757 | 1.46E-11 | 5.66E-10 |

| | | | | | |
|---|---|---|---|---|---|
| MBD3 | 4165.074816 | -0.702231072 | 0.113398831 | 3.54E-11 | 1.28E-09 |
| NEUROG2-AS1 | 56.40477156 | -0.702598604 | 0.228169641 | 8.44E-05 | 0.000638302 |
| SLC19A1 | 1980.795028 | -0.703436052 | 0.127804265 | 2.09E-09 | 5.25E-08 |
| LINC01237 | 294.7620505 | -0.703935284 | 0.142955585 | 4.51E-08 | 8.66E-07 |
| RTL8B | 956.1510583 | -0.704135208 | 0.15280699 | 2.09E-07 | 3.38E-06 |
| TTYH3 | 7895.600336 | -0.704213408 | 0.143429499 | 4.83E-08 | 9.20E-07 |
| TTC39C-AS1 | 15.76681076 | -0.704472299 | 0.439338513 | 0.00280307 | 0.01250645 |
| PHOX2A | 12626.15367 | -0.705440311 | 0.146889299 | 8.10E-08 | 1.47E-06 |
| ZBTB45 | 445.6075081 | -0.705595951 | 0.167992188 | 1.28E-06 | 1.67E-05 |
| FXN | 442.2643935 | -0.705652908 | 0.134679 | 8.66E-09 | 1.92E-07 |
| H2BC9 | 96.40719454 | -0.705877331 | 0.247713061 | 0.000164 | 0.001138212 |
| MRPS18AP1 | 6.080471794 | -0.706098179 | 0.683255289 | 0.00485293 | 0.01962296 |
| ANKRD13B | 4198.190015 | -0.706625059 | 0.107992246 | 3.49E-12 | 1.52E-10 |
| PDLIM1 | 1138.391633 | -0.706878893 | 0.199634974 | 1.70E-05 | 0.000162578 |
| MT-ND5 | 51568.9061 | -0.707428186 | 0.208987468 | 2.94E-05 | 0.000260105 |
| MGAT5B | 1864.261817 | -0.708598959 | 0.127008601 | 1.29E-09 | 3.39E-08 |
| EPHA8 | 84.24356345 | -0.7090106 | 0.232740297 | 8.80E-05 | 0.00066087 |
| ENG | 163.750539 | -0.709174797 | 0.195003137 | 1.17E-05 | 0.000116662 |
| SNX18P12 | 9.428486627 | -0.709684794 | 0.538576759 | 0.00396323 | 0.016605507 |
| FAM111B | 42.16128077 | -0.71053565 | 0.324666797 | 0.00087104 | 0.004675573 |
| DDX11 | 5262.60523 | -0.710981335 | 0.09636322 | 9.22E-15 | 5.71E-13 |
| HPGD | 6.448193839 | -0.71138401 | 0.671060789 | 0.00469936 | 0.019109283 |
| HSBP1L1 | 253.8412495 | -0.711608779 | 0.1438203 | 3.68E-08 | 7.20E-07 |
| BNIP3P27 | 43.23226492 | -0.711766563 | 0.311528483 | 0.00069072 | 0.003845123 |
| RRM2 | 9972.596307 | -0.713452928 | 0.13285681 | 3.93E-09 | 9.34E-08 |
| HIF3A | 124.3566362 | -0.713738982 | 0.240199777 | 0.00010597 | 0.000778294 |
| VDAC1P1 | 121.1735739 | -0.714133694 | 0.168967882 | 1.04E-06 | 1.39E-05 |
| RPL22P12 | 44.93702878 | -0.715094879 | 0.280660535 | 0.00034841 | 0.002152837 |
| ZDHHC12 | 269.274349 | -0.716869683 | 0.207025907 | 2.04E-05 | 0.000189983 |
| RTL8C | 1758.228702 | -0.717455014 | 0.155366686 | 1.72E-07 | 2.86E-06 |
| JPH1 | 1030.491824 | -0.718178291 | 0.124837008 | 4.27E-10 | 1.24E-08 |
| DUXAP10 | 2003.891257 | -0.71852082 | 0.115860634 | 2.79E-11 | 1.03E-09 |
| RBM12 | 7007.43015 | -0.719420645 | 0.169897597 | 9.53E-07 | 1.28E-05 |
| MIF4GD-DT | 147.601287 | -0.720105375 | 0.169462859 | 8.87E-07 | 1.20E-05 |
| SARS2 | 186.2542727 | -0.722183339 | 0.169490558 | 8.24E-07 | 1.12E-05 |
| PCDHGA9 | 192.4501052 | -0.722386329 | 0.182660381 | 2.97E-06 | 3.48E-05 |
| CD24 | 27206.83618 | -0.722526143 | 0.120114682 | 8.47E-11 | 2.81E-09 |
| DPYSL4 | 5214.152241 | -0.723468614 | 0.151247649 | 7.30E-08 | 1.34E-06 |
| SLC26A1 | 224.9995464 | -0.723545816 | 0.192852992 | 6.54E-06 | 6.94E-05 |
| CERKL | 261.2006491 | -0.724177524 | 0.253481269 | 0.00013561 | 0.000966294 |
| COL5A1 | 4233.304517 | -0.724378741 | 0.14088961 | 1.18E-08 | 2.56E-07 |
| DCAF16 | 4930.357926 | -0.725478657 | 0.121340052 | 1.02E-10 | 3.35E-09 |
| TMSB10 | 14829.73813 | -0.725683515 | 0.179136149 | 1.94E-06 | 2.38E-05 |
| SNRPA | 2682.652652 | -0.726519141 | 0.142946453 | 1.57E-08 | 3.31E-07 |
| NUP62 | 6795.890734 | -0.726530188 | 0.123281866 | 1.69E-10 | 5.33E-09 |
| TM4SF4 | 6.203611125 | -0.726717639 | 0.637829238 | 0.00420006 | 0.01741683 |
| NR2C2AP | 616.5980412 | -0.726764696 | 0.169551432 | 7.02E-07 | 9.74E-06 |
| SLC25A53 | 458.5223426 | -0.72734711 | 0.150759117 | 5.71E-08 | 1.07E-06 |
| COX5BP6 | 47.27782076 | -0.727560977 | 0.268511604 | 0.00020123 | 0.001347024 |
| KIF12 | 14.56811222 | -0.729407738 | 0.452263367 | 0.00236535 | 0.01083803 |

| | | | | | |
|---|---|---|---|---|---|
| GLYCTK | 365.0871361 | -0.729453416 | 0.151372997 | 5.74E-08 | 1.08E-06 |
| ST6GALNAC5 | 2090.537085 | -0.730404715 | 0.168946432 | 5.73E-07 | 8.23E-06 |
| LRRC3 | 89.19028058 | -0.730463485 | 0.261612178 | 0.00015487 | 0.001083593 |
| MARCKSL1 | 17109.16143 | -0.730719471 | 0.126739112 | 3.44E-10 | 1.02E-08 |
| MTCL2 | 8218.47543 | -0.730989202 | 0.138440586 | 5.25E-09 | 1.22E-07 |
| SEZ6L | 5932.50837 | -0.731363223 | 0.158528419 | 1.51E-07 | 2.55E-06 |
| SRPX | 72.33746405 | -0.731398183 | 0.316068279 | 0.00055104 | 0.003172025 |
| TCF19 | 989.7908872 | -0.731676511 | 0.109882296 | 1.21E-12 | 5.63E-11 |
| RTL8A | 1371.732977 | -0.7318782 | 0.1572664 | 1.24E-07 | 2.14E-06 |
| MIR1915HG | 604.1045557 | -0.731997984 | 0.135696658 | 2.79E-09 | 6.85E-08 |
| PRSS35 | 24.58835821 | -0.732498154 | 0.393038269 | 0.00147644 | 0.007289312 |
| H3P18 | 15.3752797 | -0.73250517 | 0.47970576 | 0.00265597 | 0.011930395 |
| TEX14BP | 168.7026468 | -0.732594802 | 0.18817517 | 3.42E-06 | 3.92E-05 |
| TNFRSF21 | 2451.72934 | -0.734304081 | 0.170567145 | 5.95E-07 | 8.46E-06 |
| FABP5P7 | 1046.359922 | -0.734978912 | 0.199942636 | 7.75E-06 | 8.08E-05 |
| FGF14-AS2 | 71.01479582 | -0.73510979 | 0.234520499 | 5.18E-05 | 0.000420693 |
| MEST | 1078.034014 | -0.735460456 | 0.152768317 | 5.49E-08 | 1.03E-06 |
| FOXRED2 | 2422.849881 | -0.735584866 | 0.170473606 | 5.62E-07 | 8.10E-06 |
| DNAH11 | 58.05404339 | -0.737139696 | 0.248703731 | 8.72E-05 | 0.000655544 |
| SLC29A4 | 4460.929847 | -0.737316405 | 0.125016984 | 1.46E-10 | 4.60E-09 |
| LINC00682 | 3202.601055 | -0.738704963 | 0.192798747 | 4.09E-06 | 4.59E-05 |
| CEND1 | 504.53096 | -0.738875645 | 0.147874651 | 2.12E-08 | 4.34E-07 |
| PHOX2B | 14296.54302 | -0.739063793 | 0.15423729 | 5.88E-08 | 1.10E-06 |
| SNHG26 | 237.8638328 | -0.740786477 | 0.162817973 | 1.83E-07 | 3.02E-06 |
| PON2 | 125.1656009 | -0.741257841 | 0.253488991 | 9.48E-05 | 0.000706811 |
| AMDHD2 | 641.4373538 | -0.741490277 | 0.123253199 | 6.77E-11 | 2.29E-09 |
| TWIST1 | 4554.418347 | -0.742224915 | 0.12250609 | 5.17E-11 | 1.81E-09 |
| ARRDC2 | 118.9310352 | -0.74289949 | 0.247353748 | 7.31E-05 | 0.000564672 |
| GPR12 | 120.3806843 | -0.744327244 | 0.210053933 | 1.15E-05 | 0.000114912 |
| FABP5 | 5051.629052 | -0.745208235 | 0.203945371 | 7.57E-06 | 7.91E-05 |
| TUBAP2 | 140.7439801 | -0.745972405 | 0.241857995 | 5.49E-05 | 0.000440325 |
| TUBB | 62423.02386 | -0.746158016 | 0.147776413 | 1.49E-08 | 3.15E-07 |
| USH1C | 36.36262669 | -0.748335238 | 0.333143485 | 0.00057064 | 0.003263107 |
| ADRA2A | 2721.247602 | -0.74924292 | 0.176418935 | 6.52E-07 | 9.15E-06 |
| SSBL2P | 9.68123804 | -0.750601675 | 0.529615729 | 0.00280633 | 0.01251828 |
| RNU6-418P | 15.8130114 | -0.752059645 | 0.41798137 | 0.00147529 | 0.007285734 |
| ADISSP | 1553.366892 | -0.752591807 | 0.166365418 | 1.81E-07 | 2.99E-06 |
| SRRM2-AS1 | 127.3839835 | -0.753397573 | 0.175607042 | 5.15E-07 | 7.49E-06 |
| ARSI | 33.3875276 | -0.753999809 | 0.359483659 | 0.00077658 | 0.004242724 |
| CCKAR | 510.4528712 | -0.755838574 | 0.209019146 | 7.83E-06 | 8.15E-05 |
| RBMS2P1 | 7.556274464 | -0.756304194 | 0.586297815 | 0.00318011 | 0.013825762 |
| INF2 | 2988.3541 | -0.756727987 | 0.11356211 | 8.70E-13 | 4.10E-11 |
| EHMT2-AS1 | 59.81095794 | -0.757024791 | 0.309942324 | 0.00032568 | 0.002029503 |
| CD320 | 1886.217765 | -0.757916652 | 0.110351117 | 2.10E-13 | 1.10E-11 |
| MED11 | 148.323449 | -0.759614491 | 0.199043904 | 3.48E-06 | 3.98E-05 |
| CFD | 48.9254733 | -0.759629625 | 0.283663721 | 0.00016715 | 0.001156184 |
| BEND3 | 1174.510515 | -0.760010884 | 0.138338537 | 1.16E-09 | 3.06E-08 |
| BRME1 | 148.4322144 | -0.762450106 | 0.193056844 | 1.98E-06 | 2.43E-05 |
| LINC02078 | 761.702997 | -0.762482229 | 0.185399201 | 1.00E-06 | 1.34E-05 |
| CHRNA1 | 103.8508743 | -0.762852792 | 0.237098731 | 3.02E-05 | 0.000266597 |

| | | | | | |
|---|---|---|---|---|---|
| NTHL1 | 835.0413843 | -0.763885182 | 0.128583157 | 8.17E-11 | 2.72E-09 |
| MXRA5 | 426.5330028 | -0.76438456 | 0.235119954 | 2.66E-05 | 0.000238318 |
| FRMPD1 | 275.1635053 | -0.76440674 | 0.260528793 | 7.47E-05 | 0.00057447 |
| TOMM5 | 384.9958741 | -0.769248925 | 0.204968595 | 4.04E-06 | 4.54E-05 |
| RTN4R | 919.520029 | -0.771238196 | 0.173358566 | 2.07E-07 | 3.36E-06 |
| P3H4 | 1592.236243 | -0.771670535 | 0.099736517 | 2.85E-16 | 2.23E-14 |
| ACTG1 | 111075.8414 | -0.773187355 | 0.170087174 | 1.29E-07 | 2.21E-06 |
| EFNA2 | 382.1296063 | -0.77340131 | 0.197406889 | 2.01E-06 | 2.45E-05 |
| DMRTA2 | 1441.110293 | -0.775249947 | 0.151395771 | 7.29E-09 | 1.64E-07 |
| PPP1R13L | 59.57933567 | -0.77551246 | 0.262573056 | 6.37E-05 | 0.000501707 |
| HAND2 | 15087.95685 | -0.775549668 | 0.110616289 | 6.16E-14 | 3.47E-12 |
| LINC01521 | 208.2619321 | -0.775822479 | 0.155585643 | 1.45E-08 | 3.08E-07 |
| MFSD10 | 2938.425268 | -0.776928316 | 0.098408975 | 7.64E-17 | 6.60E-15 |
| TMSB15B | 57.09880406 | -0.777326083 | 0.237662724 | 2.20E-05 | 0.000201896 |
| GFRA1 | 2369.932293 | -0.778367704 | 0.177040997 | 2.42E-07 | 3.85E-06 |
| C15orf39 | 984.0806729 | -0.779205951 | 0.186910199 | 6.58E-07 | 9.23E-06 |
| RAB3A | 328.2058794 | -0.779393409 | 0.177284355 | 2.40E-07 | 3.82E-06 |
| UNG | 5925.564015 | -0.779565587 | 0.146723107 | 2.48E-09 | 6.16E-08 |
| LRFN1 | 235.8443686 | -0.78018954 | 0.216021695 | 6.22E-06 | 6.64E-05 |
| PPP1R14A | 682.9267411 | -0.780876683 | 0.176474408 | 2.07E-07 | 3.36E-06 |
| EOLA2-DT | 150.4152042 | -0.781273044 | 0.193508818 | 1.13E-06 | 1.48E-05 |
| CD163L1 | 15.17817787 | -0.783432423 | 0.450315557 | 0.00137211 | 0.006869699 |
| UHRF1 | 5978.236999 | -0.783877445 | 0.154101643 | 7.86E-09 | 1.75E-07 |
| PRR34-AS1 | 61.04598157 | -0.785259242 | 0.245736823 | 2.64E-05 | 0.000236977 |
| GMEB1 | 745.83926 | -0.785414678 | 0.132405057 | 6.60E-11 | 2.24E-09 |
| CAMTA1-DT | 13.45927384 | -0.786456408 | 0.467739859 | 0.00150653 | 0.007411538 |
| GLI3 | 1235.856348 | -0.789220662 | 0.216984127 | 5.15E-06 | 5.63E-05 |
| NPW | 356.4475875 | -0.792046091 | 0.194577963 | 8.72E-07 | 1.18E-05 |
| TNS3 | 543.5166412 | -0.792436925 | 0.217750928 | 4.95E-06 | 5.44E-05 |
| FBXL8 | 121.5213554 | -0.793087209 | 0.177211084 | 1.43E-07 | 2.43E-06 |
| BASP1-AS1 | 59.54324112 | -0.793878147 | 0.2713255 | 5.93E-05 | 0.000471858 |
| CMTM7 | 385.1066429 | -0.794704049 | 0.182452622 | 2.43E-07 | 3.86E-06 |
| TLE1 | 1236.487659 | -0.794819364 | 0.146755332 | 1.17E-09 | 3.10E-08 |
| TPSB2 | 10.13252926 | -0.795584057 | 0.545824776 | 0.00209674 | 0.009841914 |
| NRM | 1138.527755 | -0.796190236 | 0.146900242 | 1.13E-09 | 3.00E-08 |
| PODXL2 | 5158.09415 | -0.79665952 | 0.086047299 | 4.30E-22 | 7.23E-20 |
| ZIC3 | 3614.309655 | -0.801799097 | 0.156944273 | 5.65E-09 | 1.30E-07 |
| EXOC3L1 | 129.2646778 | -0.802563875 | 0.174400328 | 7.10E-08 | 1.30E-06 |
| PLSCR5 | 1506.557568 | -0.803248069 | 0.248037509 | 1.93E-05 | 0.000181836 |
| E2F1 | 5493.171714 | -0.804093108 | 0.121737026 | 7.05E-13 | 3.36E-11 |
| MSI1 | 3156.617424 | -0.804755619 | 0.196539871 | 6.88E-07 | 9.58E-06 |
| DHPS | 298.7152076 | -0.80503574 | 0.161011396 | 9.59E-09 | 2.11E-07 |
| ZNF321P | 40.8529568 | -0.805109206 | 0.304212596 | 0.00012735 | 0.000911858 |
| HCFC1 | 9872.356255 | -0.805147599 | 0.209482938 | 1.95E-06 | 2.39E-05 |
| KCNJ12 | 418.867715 | -0.806218997 | 0.135297813 | 4.31E-11 | 1.53E-09 |
| CAMKK2 | 4399.591228 | -0.806293105 | 0.13359641 | 2.70E-11 | 9.98E-10 |
| LRRC45 | 1723.260672 | -0.808243427 | 0.105786378 | 3.73E-16 | 2.85E-14 |
| MAB21L2 | 5010.877518 | -0.808869259 | 0.128851058 | 5.69E-12 | 2.36E-10 |
| CACNA1A | 98.89318652 | -0.81043127 | 0.219731976 | 3.43E-06 | 3.93E-05 |
| NME4 | 4897.837912 | -0.811364524 | 0.09863502 | 3.22E-18 | 3.35E-16 |

| | | | | | |
|---|---|---|---|---|---|
| TERT | 221.991039 | -0.816560014 | 0.141802734 | 1.26E-10 | 4.06E-09 |
| CHGA | 4992.834856 | -0.817850649 | 0.131010885 | 6.34E-12 | 2.62E-10 |
| METRN | 462.3506782 | -0.818349542 | 0.13327562 | 1.20E-11 | 4.75E-10 |
| SAPCD2 | 2003.866235 | -0.821341678 | 0.085332229 | 9.20E-24 | 1.73E-21 |
| SNX22 | 204.7344427 | -0.824802864 | 0.174589941 | 3.05E-08 | 6.07E-07 |
| LY86-AS1 | 82.06349577 | -0.827765995 | 0.277875604 | 3.78E-05 | 0.00032225 |
| TCF15 | 56.17617957 | -0.828351085 | 0.299814942 | 7.52E-05 | 0.000577747 |
| TSPEAR-AS1 | 28.71024931 | -0.829021479 | 0.380066742 | 0.00038811 | 0.002360962 |
| PTX3 | 685.890415 | -0.829805506 | 0.195925598 | 2.86E-07 | 4.45E-06 |
| NEUROG2 | 426.9339084 | -0.830006607 | 0.146230577 | 1.73E-10 | 5.41E-09 |
| LRFN4 | 1748.37872 | -0.830538858 | 0.165247293 | 6.21E-09 | 1.42E-07 |
| PITPNM2 | 1119.613734 | -0.833634201 | 0.18359047 | 6.72E-08 | 1.24E-06 |
| CLPB | 3125.808133 | -0.839925468 | 0.115595279 | 4.11E-15 | 2.73E-13 |
| CD24P4 | 68.64118507 | -0.840279196 | 0.222427037 | 1.80E-06 | 2.23E-05 |
| PRR22 | 256.6704714 | -0.841825412 | 0.162260256 | 2.31E-09 | 5.76E-08 |
| CHST6 | 49.84742685 | -0.843110593 | 0.299780582 | 5.77E-05 | 0.000460084 |
| SLCO4A1-AS1 | 18.41074087 | -0.845430571 | 0.48983776 | 0.00101265 | 0.00531632 |
| SNORD104 | 29.55540292 | -0.845794849 | 0.33844938 | 0.00014685 | 0.001034892 |
| ZBTB42 | 184.3773655 | -0.846091679 | 0.1934109 | 1.28E-07 | 2.20E-06 |
| TUBBP1 | 236.7544228 | -0.846255924 | 0.184921211 | 4.94E-08 | 9.40E-07 |
| RAB27B | 48.59544254 | -0.847179222 | 0.356725729 | 0.00020705 | 0.001378056 |
| ZNRF3 | 3433.189716 | -0.847199874 | 0.181308107 | 3.06E-08 | 6.09E-07 |
| CAPN15 | 4458.937895 | -0.847591932 | 0.11037871 | 1.60E-16 | 1.31E-14 |
| MBOAT1 | 118.3388211 | -0.847979758 | 0.206790726 | 4.29E-07 | 6.41E-06 |
| MIR6071 | 11.45820675 | -0.853391074 | 0.532236426 | 0.00123767 | 0.006313938 |
| ACTL8 | 12.50100106 | -0.855337915 | 0.622560248 | 0.00179449 | 0.008603864 |
| KLHL35 | 140.3945788 | -0.856014952 | 0.16464163 | 1.84E-09 | 4.66E-08 |
| SLC1A1 | 423.5589868 | -0.856342243 | 0.161179568 | 9.86E-10 | 2.65E-08 |
| LHX9 | 1806.994846 | -0.859500475 | 0.175488052 | 8.66E-09 | 1.92E-07 |
| MFNG | 262.4551437 | -0.860400859 | 0.157901054 | 4.40E-10 | 1.27E-08 |
| P2RX5 | 386.2794886 | -0.860697876 | 0.123255815 | 2.43E-14 | 1.43E-12 |
| CCNJL | 461.3564476 | -0.862464751 | 0.160471162 | 6.52E-10 | 1.82E-08 |
| MIR6783 | 13.45681548 | -0.864533982 | 0.542942492 | 0.00119992 | 0.006154923 |
| GSTM5P1 | 92.01404832 | -0.866450476 | 0.223908123 | 9.67E-07 | 1.30E-05 |
| NEURL1B | 1301.569142 | -0.86705408 | 0.139692551 | 4.24E-12 | 1.80E-10 |
| CA12 | 647.9546642 | -0.869349715 | 0.206227696 | 2.09E-07 | 3.39E-06 |
| FNDC10 | 147.6300151 | -0.873051784 | 0.210289649 | 2.69E-07 | 4.22E-06 |
| PDE2A | 5058.000941 | -0.875724747 | 0.160103111 | 3.27E-10 | 9.75E-09 |
| ZNF346-IT1 | 37.58432586 | -0.876232234 | 0.307054077 | 3.99E-05 | 0.000337582 |
| IFI44 | 1762.69753 | -0.87666596 | 0.20404126 | 1.35E-07 | 2.30E-06 |
| CENATAC-DT | 84.04160507 | -0.876797661 | 0.219886641 | 5.33E-07 | 7.72E-06 |
| ADAMTS9-AS1 | 53.68284549 | -0.877609614 | 0.42722276 | 0.00040678 | 0.002453785 |
| CDH7 | 1503.224216 | -0.877816771 | 0.220214823 | 5.31E-07 | 7.70E-06 |
| HK2-DT | 337.4196889 | -0.879312941 | 0.177163154 | 4.98E-09 | 1.16E-07 |
| RNF207 | 987.8561014 | -0.880915738 | 0.133393208 | 2.61E-13 | 1.34E-11 |
| OGFOD3 | 2075.563645 | -0.88286597 | 0.125242554 | 1.13E-14 | 6.93E-13 |
| ITGA4 | 1640.026696 | -0.88303083 | 0.230050399 | 9.51E-07 | 1.28E-05 |
| SDC2 | 1580.594249 | -0.883346016 | 0.209801136 | 1.87E-07 | 3.08E-06 |
| LINC01088 | 136.5925501 | -0.883371976 | 0.304869309 | 3.29E-05 | 0.000286145 |
| PLAUR | 404.7003824 | -0.884379258 | 0.141664075 | 2.72E-12 | 1.21E-10 |

| | | | | | |
|---|---|---|---|---|---|
| ASIP | 61.68557956 | -0.884620104 | 0.230747776 | 9.57E-07 | 1.29E-05 |
| IGFBPL1 | 23762.41856 | -0.884640782 | 0.168336878 | 9.83E-10 | 2.64E-08 |
| C1QTNF4 | 96.53506797 | -0.884977389 | 0.260505746 | 5.45E-06 | 5.91E-05 |
| COL6A6 | 96.28238303 | -0.887844065 | 0.293675163 | 2.08E-05 | 0.000192991 |
| FNDC5 | 5725.563632 | -0.889051361 | 0.113932558 | 3.40E-17 | 3.14E-15 |
| SNHG3 | 3211.753436 | -0.890937166 | 0.167485577 | 6.43E-10 | 1.80E-08 |
| NTN3 | 325.0289199 | -0.891885563 | 0.19061273 | 1.86E-08 | 3.87E-07 |
| ATOH8 | 2498.300459 | -0.894389616 | 0.165757921 | 4.03E-10 | 1.18E-08 |
| ST8SIA2 | 5157.837299 | -0.896397465 | 0.119771958 | 3.72E-16 | 2.85E-14 |
| RALGDS | 1410.38675 | -0.89789224 | 0.126232907 | 5.84E-15 | 3.78E-13 |
| ZBTB8B | 167.9876029 | -0.899333826 | 0.244800049 | 1.64E-06 | 2.07E-05 |
| PPP1R1B | 15.86398286 | -0.901009941 | 0.457430626 | 0.00044631 | 0.00265871 |
| AFP | 3.818824457 | -0.905668654 | 0.945478732 | 0.00194053 | 0.009222353 |
| MVP-DT | 291.1293932 | -0.906521679 | 0.13633027 | 1.39E-13 | 7.36E-12 |
| ACTB | 51459.28137 | -0.911657107 | 0.215025038 | 1.27E-07 | 2.19E-06 |
| CNTN2 | 4732.179743 | -0.912093461 | 0.214940086 | 1.24E-07 | 2.15E-06 |
| SNORA2B | 13.41356536 | -0.912558447 | 0.569231133 | 0.00095302 | 0.005053617 |
| GLYCTK-AS1 | 45.60655742 | -0.916547895 | 0.296455572 | 1.35E-05 | 0.000132333 |
| LPCAT1 | 3330.38188 | -0.918495473 | 0.169057178 | 2.53E-10 | 7.65E-09 |
| PHB1P4 | 11.84198907 | -0.91920429 | 0.629345925 | 0.00120231 | 0.006162757 |
| HYAL2 | 1361.618061 | -0.920884474 | 0.142313511 | 3.94E-13 | 1.96E-11 |
| CCDC68 | 17.5304373 | -0.939678893 | 0.533031535 | 0.00061019 | 0.003458633 |
| CTXN1 | 1545.003863 | -0.94035972 | 0.204469219 | 1.78E-08 | 3.70E-07 |
| TSPEAR | 56.85527935 | -0.942426217 | 0.292236287 | 7.02E-06 | 7.39E-05 |
| KCNH4 | 767.6189167 | -0.942855764 | 0.123525391 | 6.60E-17 | 5.73E-15 |
| EPAS1 | 50.39191787 | -0.943574344 | 0.303552515 | 1.07E-05 | 0.000107607 |
| LAMC3 | 141.8096794 | -0.943692914 | 0.230859171 | 1.96E-07 | 3.22E-06 |
| ZNF488 | 269.9881998 | -0.951478717 | 0.183449263 | 7.38E-10 | 2.02E-08 |
| PCDHB2 | 412.6752973 | -0.952548875 | 0.216024571 | 4.08E-08 | 7.93E-07 |
| ITPRID1 | 22.18337296 | -0.953266377 | 0.413536711 | 0.00014219 | 0.001007231 |
| LFNG | 350.3187193 | -0.954408975 | 0.156769012 | 3.37E-12 | 1.47E-10 |
| FAM86GP | 364.4464604 | -0.954859946 | 0.14975407 | 5.15E-13 | 2.53E-11 |
| LINC01537 | 129.1746234 | -0.957560283 | 0.233759108 | 1.71E-07 | 2.83E-06 |
| SNHG4 | 1525.973592 | -0.958743634 | 0.170905239 | 6.13E-11 | 2.11E-09 |
| SEC13P1 | 6.942094985 | -0.959090318 | 0.739388764 | 0.00132408 | 0.006675259 |
| RNU6-1223P | 25.81775016 | -0.96081841 | 0.516057157 | 0.00044566 | 0.0026556 |
| SPRY1 | 134.1573089 | -0.960821693 | 0.213360183 | 2.43E-08 | 4.92E-07 |
| CDH24 | 4383.445849 | -0.963581562 | 0.107444593 | 6.24E-22 | 1.00E-19 |
| LRRC20 | 886.9454746 | -0.974563432 | 0.178509096 | 1.29E-10 | 4.15E-09 |
| NAPRT | 379.9291711 | -0.978960591 | 0.143866108 | 2.17E-14 | 1.29E-12 |
| SNORA73B | 132.7080087 | -0.980026729 | 0.224757832 | 4.25E-08 | 8.24E-07 |
| INHBA | 47.56482144 | -0.98234531 | 0.405614368 | 8.82E-05 | 0.000662028 |
| ACTG2 | 9.38438933 | -0.98698276 | 0.614599445 | 0.00071008 | 0.00393537 |
| ZNF341 | 189.012101 | -0.996926769 | 0.167063847 | 5.02E-12 | 2.12E-10 |
| ENOX1-AS1 | 4.945250135 | -0.999026169 | 1.02254835 | 0.00136704 | 0.006853091 |
| SLC44A3 | 86.18878876 | -1.005146294 | 0.298037826 | 2.81E-06 | 3.31E-05 |
| NNAT | 12616.40701 | -1.008541174 | 0.1519229 | 5.42E-14 | 3.07E-12 |
| DDN | 815.9017546 | -1.017668666 | 0.171634176 | 5.51E-12 | 2.31E-10 |
| DDX11L2 | 205.0031099 | -1.019463102 | 0.167200809 | 1.87E-12 | 8.49E-11 |
| POLR1G | 256.7286665 | -1.023687529 | 0.204005293 | 1.11E-09 | 2.96E-08 |

| | | | | | |
|---|---|---|---|---|---|
| PRDM8 | 2435.717213 | -1.028606195 | 0.149207741 | 7.79E-15 | 4.91E-13 |
| RAPGEF5 | 1725.277871 | -1.034779678 | 0.227680165 | 1.25E-08 | 2.70E-07 |
| EGFLAM | 696.5259623 | -1.039675537 | 0.205672897 | 8.34E-10 | 2.27E-08 |
| C1QL1 | 3039.03845 | -1.044479137 | 0.183254255 | 1.96E-11 | 7.42E-10 |
| STC1 | 4512.061339 | -1.044736558 | 0.205829912 | 7.26E-10 | 2.00E-08 |
| SLCO4A1 | 809.01366 | -1.045190273 | 0.140604074 | 1.27E-16 | 1.07E-14 |
| VGLL3 | 112.7025975 | -1.053922153 | 0.276877499 | 3.70E-07 | 5.57E-06 |
| H3P21 | 31.46123875 | -1.059532867 | 0.520065503 | 0.0001988 | 0.001333339 |
| IGSF9 | 1712.177003 | -1.061345809 | 0.134336105 | 2.89E-18 | 3.02E-16 |
| PPP1R17 | 52.77097615 | -1.062544162 | 0.35847284 | 1.04E-05 | 0.000105012 |
| SNN | 2206.130883 | -1.063644538 | 0.183232081 | 9.40E-12 | 3.78E-10 |
| NWD2 | 418.0500562 | -1.092510999 | 0.43536485 | 4.46E-05 | 0.000371091 |
| THSD4-AS1 | 10.679281 | -1.096267385 | 0.719065242 | 0.00058324 | 0.003327911 |
| CCDC177 | 849.8951975 | -1.117571667 | 0.202053688 | 4.23E-11 | 1.51E-09 |
| LRRC55 | 596.5871461 | -1.119136325 | 0.227244437 | 1.29E-09 | 3.39E-08 |
| LINC02012 | 63.01767019 | -1.13593041 | 0.284802747 | 1.29E-07 | 2.21E-06 |
| LRRC4 | 632.8343767 | -1.145488674 | 0.148383397 | 1.09E-17 | 1.07E-15 |
| RNVU1-30 | 12.14479799 | -1.146464435 | 0.660077967 | 0.00032343 | 0.002017931 |
| HRH3 | 1126.208301 | -1.147471911 | 0.126333253 | 8.77E-23 | 1.54E-20 |
| HPCA | 1637.729711 | -1.150436477 | 0.244404688 | 3.88E-09 | 9.24E-08 |
| MIRLET7BHG | 796.5836029 | -1.154124472 | 0.172131098 | 2.14E-14 | 1.28E-12 |
| DLL3 | 892.718283 | -1.15706037 | 0.150084916 | 1.20E-17 | 1.18E-15 |
| NEUROD1 | 5360.658256 | -1.161036545 | 0.21677173 | 1.13E-10 | 3.67E-09 |
| PMEL | 1057.446307 | -1.16930777 | 0.176723941 | 4.01E-14 | 2.31E-12 |
| PDE2A-AS2 | 13.16944242 | -1.171681347 | 0.583532782 | 0.00015493 | 0.001083637 |
| TPSG1 | 46.18563981 | -1.194834344 | 0.318372385 | 3.31E-07 | 5.07E-06 |
| SLC2A4 | 41.98940553 | -1.198054667 | 0.394367043 | 5.59E-06 | 6.04E-05 |
| IGFBP5 | 9356.454017 | -1.204336085 | 0.231055849 | 2.57E-10 | 7.77E-09 |
| HSD11B2 | 232.2894691 | -1.234467161 | 0.147888665 | 8.59E-20 | 1.09E-17 |
| SUSD2 | 237.476548 | -1.239578147 | 0.231967841 | 1.29E-10 | 4.15E-09 |
| FJX1 | 2271.851506 | -1.252722107 | 0.184846518 | 1.68E-14 | 1.02E-12 |
| RAB26 | 377.8225 | -1.293668069 | 0.176390957 | 3.80E-16 | 2.89E-14 |
| SLCO4A1-AS2 | 31.87826065 | -1.300636416 | 0.505772548 | 2.28E-05 | 0.000209098 |
| HPDL | 749.0391661 | -1.32429625 | 0.191363247 | 8.83E-15 | 5.48E-13 |
| EBF2 | 321.9601032 | -1.325945578 | 0.251719289 | 2.41E-10 | 7.34E-09 |
| MT-TR | 100.2154373 | -1.333967022 | 0.232956746 | 1.89E-11 | 7.20E-10 |
| CDH5 | 44.63593284 | -1.385540849 | 0.341223277 | 8.92E-08 | 1.61E-06 |
| MT-TG | 90.40892736 | -1.420712319 | 0.255815983 | 6.80E-11 | 2.29E-09 |
| NEUROD6 | 75.07586765 | -1.422547138 | 0.315624508 | 1.31E-08 | 2.81E-07 |
| MT-TH | 70.62168347 | -1.428044887 | 0.29383832 | 2.52E-09 | 6.26E-08 |
| CDH15 | 277.8272142 | -1.46814053 | 0.219861082 | 9.47E-14 | 5.18E-12 |
| NAA11 | 7.178212134 | -1.484846113 | 0.87857581 | 0.00017133 | 0.00118033 |
| PLCH2 | 396.8735962 | -1.514701701 | 0.198332524 | 1.37E-16 | 1.14E-14 |
| ATP6V1G3 | 9.670491287 | -1.643213073 | 0.725915376 | 3.54E-05 | 0.000304681 |
| BTBD17 | 88.32564413 | -1.695945786 | 0.284260473 | 1.38E-11 | 5.37E-10 |
| LINC00989 | 8.881800139 | -1.954539161 | 0.939562761 | 3.88E-05 | 0.000329081 |

**Table S3. List of interactomes in SH-SY5Y cells**

| BAIT | PROTID | GENE | preyFreq | preyAvg_AvgCounts | preySD_AvgCounts | NS1_AVG_SPC | NS1_FC_A | NS1_FC_B | NS1_SP | NS1_iREF | IP_SH-SY5Y_NS1_R4 | IP_SH-SY5Y_NS1_R3 | IP_SH-SY5Y_NS1_R2 | IP_SH-SY5Y_NS1_R1 | IP_SH-SY5Y_GFP_R4 | IP_SH-SY5Y_GFP_R3 | IP_SH-SY5Y_GFP_R2 | IP_SH-SY5Y_GFP_R1 |
|---|---|---|---|---|---|---|---|---|---|---|---|---|---|---|---|---|---|---|
| NS1 | Q14697-2 | GANAB | 1 | 4.5 | 0 | 4.5 | 2.26 | 2.25 | 1 | 0 | 6 | 4 | 4 | 4 | 0 | 0 | 0 | 0 |
| NS1 | P11310-2 | ACADM | 1 | 5.75 | 0 | 5.75 | 2.62 | 2.59 | 1 | 0 | 6 | 7 | 6 | 4 | 0 | 0 | 0 | 0 |
| NS1 | O43707 | ACTN4 | 1 | 6 | 0 | 6 | 2.68 | 2.66 | 1 | 0 | 8 | 5 | 6 | 5 | 0 | 0 | 0 | 0 |
| NS1 | P33991 | MCM4 | 1 | 6 | 0 | 6 | 2.68 | 2.65 | 1 | 0 | 6 | 8 | 4 | 6 | 0 | 0 | 0 | 0 |
| NS1 | Q13642-5 | FHL1 | 1 | 6.5 | 0 | 6.5 | 2.81 | 2.8 | 1 | 0 | 6 | 5 | 7 | 8 | 0 | 0 | 0 | 0 |
| NS1 | P21796 | VDAC1 | 1 | 7 | 0 | 7 | 2.95 | 2.9 | 1 | 0 | 5 | 8 | 5 | 10 | 0 | 0 | 0 | 0 |
| NS1 | P61604 | HSPE1 | 1 | 7 | 0 | 7 | 2.96 | 2.95 | 1 | 0 | 7 | 7 | 8 | 6 | 0 | 0 | 0 | 0 |
| NS1 | P43304 | GPD2 | 1 | 7.25 | 0 | 7.25 | 3.03 | 2.92 | 1 | 0 | 6 | 12 | 4 | 7 | 0 | 0 | 0 | 0 |
| NS1 | Q14324 | MYBPC2 | 1 | 7.5 | 0 | 7.5 | 3.09 | 3.05 | 1 | 0 | 6 | 6 | 7 | 11 | 0 | 0 | 0 | 0 |
| NS1 | A0A0B4J2D5 | LOC102724023 | 1 | 7.5 | 0 | 7.5 | 3.1 | 3.07 | 1 | 0 | 7 | 7 | 10 | 6 | 0 | 0 | 0 | 0 |
| NS1 | Q8N142 | ADSSL1 | 1 | 7.75 | 0 | 7.75 | 3.17 | 3.16 | 1 | 0 | 9 | 7 | 8 | 7 | 0 | 0 | 0 | 0 |
| NS1 | P36957 | DLST | 1 | 7.75 | 0 | 7.75 | 3.17 | 3.15 | 1 | 0 | 8 | 9 | 6 | 8 | 0 | 0 | 0 | 0 |
| NS1 | P62328 | TMSB4X | 1 | 8 | 0 | 8 | 3.24 | 3.22 | 1 | 0 | 9 | 9 | 6 | 8 | 0 | 0 | 0 | 0 |
| NS1 | P99999 | CYCS | 1 | 8.25 | 0 | 8.25 | 3.3 | 3.29 | 1 | 0 | 7 | 8 | 8 | 10 | 0 | 0 | 0 | 0 |
| NS1 | P09104 | ENO2 | 1 | 8.25 | 0 | 8.25 | 3.31 | 3.3 | 1 | 0 | 9 | 9 | 8 | 7 | 0 | 0 | 0 | 0 |
| NS1 | P00167 | CYB5A | 1 | 8.25 | 0 | 8.25 | 3.31 | 3.3 | 1 | 0 | 8 | 7 | 9 | 9 | 0 | 0 | 0 | 0 |
| NS1 | P10515 | DLAT | 1 | 8.5 | 0 | 8.5 | 3.38 | 3.37 | 1 | 0 | 9 | 9 | 7 | 9 | 0 | 0 | 0 | 0 |
| NS1 | P31946 | YWHAB | 1 | 8.75 | 0 | 8.75 | 3.45 | 3.37 | 1 | 0 | 11 | 11 | 5 | 8 | 0 | 0 | 0 | 0 |
| NS1 | P11217-2 | PYGM | 1 | 8.75 | 0 | 8.75 | 3.45 | 3.45 | 1 | 0 | 8 | 9 | 9 | 9 | 0 | 0 | 0 | 0 |
| NS1 | P21695-2 | GPD1 | 1 | 8.75 | 0 | 8.75 | 3.46 | 3.44 | 1 | 0 | 9 | 9 | 10 | 7 | 0 | 0 | 0 | 0 |
| NS1 | Q16891-2 | IMMT | 1 | 8.75 | 0 | 8.75 | 3.46 | 3.34 | 1 | 0 | 9 | 14 | 6 | 6 | 0 | 0 | 0 | 0 |
| NS1 | Q15942 | ZYX | 1 | 9 | 0 | 9 | 3.53 | 3.48 | 1 | 0 | 11 | 11 | 7 | 7 | 0 | 0 | 0 | 0 |
| NS1 | A0A2R8YDQ9 | SUCLA2 | 1 | 9.25 | 0 | 9.25 | 3.59 | 3.56 | 1 | 0 | 8 | 12 | 8 | 9 | 0 | 0 | 0 | 0 |
| NS1 | P17540 | CKMT2 | 1 | 9.5 | 0 | 9.5 | 3.64 | 3.53 | 1 | 0 | 5 | 8 | 11 | 14 | 0 | 0 | 0 | 0 |
| NS1 | P07954-2 | FH | 1 | 18.75 | 0 | 18.75 | 4 | 3.55 | 1 | 0 | 16 | 22 | 18 | 19 | 0 | 0 | 1 | 1 |
| NS1 | P15090 | FABP4 | 1 | 10.75 | 0 | 10.75 | 4.01 | 4.01 | 1 | 0 | 11 | 11 | 11 | 10 | 0 | 0 | 0 | 0 |
| NS1 | P84243 | H3F3A;H3F3B | 1 | 10.75 | 0 | 10.75 | 4.02 | 3.82 | 1 | 0 | 19 | 8 | 7 | 9 | 0 | 0 | 0 | 0 |
| NS1 | P60174-3 | TPI1 | 1 | 11 | 0 | 11 | 4.08 | 4.06 | 1 | 0 | 12 | 9 | 12 | 11 | 0 | 0 | 0 | 0 |
| NS1 | P15259 | PGAM2 | 1 | 12.5 | 0 | 12.5 | 4.49 | 4.48 | 1 | 0 | 11 | 12 | 12 | 15 | 0 | 0 | 0 | 0 |
| NS1 | Q00872-10 | MYBPC1 | 1 | 12.75 | 0 | 12.75 | 4.55 | 4.49 | 1 | 0 | 9 | 12 | 13 | 17 | 0 | 0 | 0 | 0 |
| NS1 | Q6NZI2 | CAVIN1 | 1 | 12.75 | 0 | 12.75 | 4.57 | 4.55 | 1 | 0 | 12 | 15 | 12 | 12 | 0 | 0 | 0 | 0 |
| NS1 | P10916 | MYL2 | 1 | 13 | 0 | 13 | 4.63 | 4.58 | 1 | 0 | 9 | 13 | 15 | 15 | 0 | 0 | 0 | 0 |
| NS1 | O00217 | NDUFS8 | 1 | 14.25 | 0 | 14.25 | 4.96 | 4.86 | 1 | 0 | 11 | 11 | 14 | 21 | 0 | 0 | 0 | 0 |
| NS1 | A0A2R8YEA7 | ACTB | 1 | 14.75 | 0 | 14.75 | 5.12 | 5.11 | 1 | 0 | 15 | 14 | 13 | 17 | 0 | 0 | 0 | 0 |
| NS1 | P07205 | PGK2 | 1 | 14.75 | 0 | 14.75 | 5.13 | 5.12 | 1 | 0 | 14 | 15 | 16 | 14 | 0 | 0 | 0 | 0 |
| NS1 | P53396 | ACLY | 1 | 16.5 | 0 | 16.5 | 5.63 | 5.51 | 1 | 0 | 20 | 21 | 13 | 12 | 0 | 0 | 0 | 0 |
| NS1 | P40926 | MDH2 | 1 | 17.75 | 0 | 17.75 | 5.97 | 5.9 | 1 | 0 | 21 | 19 | 13 | 18 | 0 | 0 | 0 | 0 |
| NS1 | P27708 | CAD | 1 | 29.5 | 0 | 29.5 | 6.06 | 5.4 | 1 | 0 | 32 | 32 | 28 | 26 | 3 | 0 | 0 | 0 |
| NS1 | P08559-2 | PDHA1 | 1 | 18.25 | 0 | 18.25 | 6.12 | 6.01 | 1 | 0 | 17 | 25 | 17 | 14 | 0 | 0 | 0 | 0 |
| NS1 | P35609 | ACTN2 | 1 | 19 | 0 | 19 | 6.28 | 5.86 | 1 | 0 | 9 | 13 | 24 | 30 | 0 | 0 | 0 | 0 |
| NS1 | P48643 | CCT5 | 1 | 128.5 | 0 | 128.5 | 6.54 | 5.86 | 1 | 0 | 141 | 144 | 128 | 101 | 6 | 3 | 3 | 6 |
| NS1 | A0A0C4DG89 | DDX46 | 1 | 20.75 | 0 | 20.75 | 6.8 | 6.8 | 1 | 0 | 20 | 20 | 21 | 22 | 0 | 0 | 0 | 0 |
| NS1 | P06753-2 | TPM3 | 1 | 33.25 | 0 | 33.25 | 6.87 | 6.15 | 1 | 0 | 30 | 30 | 31 | 42 | 1 | 0 | 0 | 1 |
| NS1 | P08590 | MYL3 | 1 | 21.25 | 0 | 21.25 | 6.89 | 6.6 | 1 | 0 | 15 | 17 | 17 | 36 | 0 | 0 | 0 | 0 |
| NS1 | U3KQK2 | TPM2 | 1 | 21.25 | 0 | 21.25 | 6.94 | 6.9 | 1 | 0 | 17 | 21 | 23 | 24 | 0 | 0 | 0 | 0 |
| NS1 | P13804 | ETFA | 1 | 44.75 | 0 | 44.75 | 7.14 | 6.05 | 1 | 0 | 35 | 36 | 45 | 63 | 0 | 1 | 1 | 1 |
| NS1 | P48788-2 | TNNI2 | 1 | 22.25 | 0 | 22.25 | 7.19 | 7.03 | 1 | 0 | 16 | 18 | 23 | 32 | 0 | 0 | 0 | 0 |
| NS1 | Q13813 | SPTAN1 | 1 | 24.5 | 0 | 24.5 | 7.84 | 7.63 | 1 | 0 | 17 | 33 | 19 | 29 | 0 | 0 | 0 | 0 |
| NS1 | P00558-2 | PGK1 | 1 | 29.25 | 0 | 29.25 | 9.16 | 9.12 | 1 | 0 | 25 | 30 | 27 | 35 | 0 | 0 | 0 | 0 |
| NS1 | P40227 | CCT6A | 1 | 117 | 0 | 117 | 9.38 | 8.06 | 1 | 0 | 139 | 141 | 115 | 73 | 3 | 1 | 2 | 4 |
| NS1 | P18669 | PGAM1 | 1 | 31 | 0 | 31 | 9.65 | 9.57 | 1 | 0 | 26 | 27 | 32 | 39 | 0 | 0 | 0 | 0 |
| NS1 | P00558 | PGK1 | 1 | 31.75 | 0 | 31.75 | 9.87 | 9.84 | 1 | 0 | 28 | 34 | 29 | 36 | 0 | 0 | 0 | 0 |
| NS1 | P06733-2 | ENO1 | 1 | 32 | 0 | 32 | 9.99 | 9.73 | 1 | 0 | 45 | 31 | 28 | 24 | 0 | 0 | 0 | 0 |
| NS1 | P04075 | ALDOA | 1 | 32.75 | 0 | 32.75 | 10.13 | 10.05 | 1 | 0 | 28 | 29 | 32 | 42 | 0 | 0 | 0 | 0 |
| NS1 | P05976 | MYL1 | 1 | 33 | 0 | 33 | 10.18 | 9.98 | 1 | 0 | 24 | 30 | 31 | 47 | 0 | 0 | 0 | 0 |
| NS1 | O14983-2 | ATP2A1 | 1 | 34.5 | 0 | 34.5 | 10.63 | 10.53 | 1 | 0 | 26 | 34 | 37 | 41 | 0 | 0 | 0 | 0 |
| NS1 | F8WA37 | TNNT3 | 1 | 34.75 | 0 | 34.75 | 10.7 | 10.66 | 1 | 0 | 31 | 31 | 36 | 41 | 0 | 0 | 0 | 0 |
| NS1 | P45378-7 | TNNT3 | 1 | 34.75 | 0 | 34.75 | 10.7 | 10.66 | 1 | 0 | 31 | 31 | 36 | 41 | 0 | 0 | 0 | 0 |
| NS1 | P06753-5 | TPM3 | 1 | 35.25 | 0 | 35.25 | 10.83 | 10.75 | 1 | 0 | 31 | 32 | 33 | 45 | 0 | 0 | 0 | 0 |
| NS1 | Q96A32 | MYLPF | 1 | 36 | 0 | 36 | 11.03 | 10.76 | 1 | 0 | 23 | 33 | 41 | 47 | 0 | 0 | 0 | 0 |
| NS1 | A0A7I2V5A1 | ACO2 | 1 | 36.25 | 0 | 36.25 | 11.11 | 11.03 | 1 | 0 | 31 | 34 | 34 | 46 | 0 | 0 | 0 | 0 |
| NS1 | P13929 | ENO3 | 1 | 36.5 | 0 | 36.5 | 11.18 | 11.08 | 1 | 0 | 31 | 32 | 36 | 47 | 0 | 0 | 0 | 0 |
| NS1 | P06753-6 | TPM3 | 1 | 38.5 | 0 | 38.5 | 11.74 | 11.65 | 1 | 0 | 34 | 34 | 37 | 49 | 0 | 0 | 0 | 0 |
| NS1 | A7E2Y1 | MYH7B | 1 | 38.5 | 0 | 38.5 | 11.75 | 11.7 | 1 | 0 | 34 | 36 | 38 | 46 | 0 | 0 | 0 | 0 |
| NS1 | Q99832 | CCT7 | 1 | 85.5 | 0 | 85.5 | 12.12 | 10.16 | 1 | 0 | 94 | 101 | 84 | 63 | 1 | 2 | 1 | 0 |
| NS1 | P67936 | TPM4 | 1 | 40.25 | 0 | 40.25 | 12.22 | 12.09 | 1 | 0 | 33 | 35 | 41 | 52 | 0 | 0 | 0 | 0 |
| NS1 | P49368 | CCT3 | 1 | 137.75 | 0 | 137.75 | 12.72 | 11.58 | 1 | 0 | 163 | 152 | 138 | 98 | 2 | 2 | 2 | 2 |
| NS1 | Q08043 | ACTN3 | 1 | 42.25 | 0 | 42.25 | 12.77 | 12.3 | 1 | 0 | 26 | 34 | 52 | 57 | 0 | 0 | 0 | 0 |
| NS1 | P16615-2 | ATP2A2 | 1 | 45.25 | 0 | 45.25 | 13.65 | 13.59 | 1 | 0 | 37 | 48 | 49 | 47 | 0 | 0 | 0 | 0 |
| NS1 | P50991 | CCT4 | 1 | 112 | 0 | 112 | 14.25 | 11.8 | 1 | 0 | 127 | 126 | 113 | 82 | 0 | 1 | 0 | 3 |
| NS1 | H0YK48 | TPM1 | 1 | 55.75 | 0 | 55.75 | 16.53 | 16.35 | 1 | 0 | 47 | 50 | 52 | 74 | 0 | 0 | 0 | 0 |
| NS1 | P06753 | TPM3 | 1 | 57.75 | 0 | 57.75 | 17.1 | 16.99 | 1 | 0 | 50 | 53 | 55 | 73 | 0 | 0 | 0 | 0 |
| NS1 | Q99798 | ACO2 | 1 | 58.5 | 0 | 58.5 | 17.31 | 17.17 | 1 | 0 | 48 | 55 | 57 | 74 | 0 | 0 | 0 | 0 |
| NS1 | P20929-3 | NEB | 1 | 59 | 0 | 59 | 17.39 | 16.09 | 1 | 0 | 25 | 62 | 59 | 90 | 0 | 0 | 0 | 0 |
| NS1 | P09493-6 | TPM1 | 1 | 65.5 | 0 | 65.5 | 19.23 | 18.9 | 1 | 0 | 53 | 59 | 58 | 92 | 0 | 0 | 0 | 0 |
| NS1 | P06732 | CKM | 1 | 68.75 | 0 | 68.75 | 20.17 | 19.95 | 1 | 0 | 52 | 68 | 69 | 86 | 0 | 0 | 0 | 0 |
| NS1 | A7XZE4 | A7XZE4 | 1 | 69.5 | 0 | 69.5 | 20.36 | 20.06 | 1 | 0 | 55 | 63 | 66 | 94 | 0 | 0 | 0 | 0 |
| NS1 | Q6ZN40 | TPM1 | 1 | 78.25 | 0 | 78.25 | 22.8 | 22.52 | 1 | 0 | 66 | 73 | 69 | 105 | 0 | 0 | 0 | 0 |
| NS1 | P09493 | TPM1 | 1 | 91 | 0 | 91 | 26.35 | 26.06 | 1 | 0 | 78 | 84 | 81 | 121 | 0 | 0 | 0 | 0 |
| NS1 | P50990 | CCT8 | 1 | 120 | 0 | 120 | 34.7 | 34.12 | 1 | 0 | 134 | 138 | 119 | 89 | 0 | 0 | 0 | 0 |
| NS1 | Q8WZ42-11 | TTN | 1 | 129 | 0 | 129 | 36.78 | 32.13 | 1 | 0 | 42 | 110 | 154 | 210 | 0 | 0 | 0 | 0 |
| NS1 | Q8WZ42-12 | TTN | 1 | 130 | 0 | 130 | 37.06 | 32.34 | 1 | 0 | 42 | 111 | 156 | 211 | 0 | 0 | 0 | 0 |
| NS1 | P69905 | HBA2;HBA1 | 1 | 136.25 | 0 | 136.25 | 38.93 | 37.47 | 1 | 0 | 83 | 118 | 156 | 188 | 0 | 0 | 0 | 0 |
| NS1 | P78371 | CCT2 | 1 | 161.5 | 0 | 161.5 | 46.39 | 45.14 | 1 | 0 | 184 | 193 | 161 | 108 | 0 | 0 | 0 | 0 |
| NS1 | P13533 | MYH6 | 1 | 204.75 | 0 | 204.75 | 58.16 | 57.9 | 1 | 0 | 175 | 194 | 209 | 241 | 0 | 0 | 0 | 0 |
| NS1 | P12883 | MYH7 | 1 | 229.75 | 0 | 229.75 | 65.18 | 64.91 | 1 | 0 | 193 | 226 | 240 | 260 | 0 | 0 | 0 | 0 |
| NS1 | P11055 | MYH3 | 1 | 366.25 | 0 | 366.25 | 103.32 | 103.13 | 1 | 0 | 332 | 348 | 374 | 411 | 0 | 0 | 0 | 0 |
| NS1 | Q9UKX2 | MYH2 | 1 | 572.5 | 0 | 572.5 | 160.76 | 159.88 | 1 | 0 | 497 | 526 | 574 | 693 | 0 | 0 | 0 | 0 |
| NS1 | P13535 | MYH8 | 1 | 592 | 0 | 592 | 166.33 | 165.84 | 1 | 0 | 538 | 544 | 602 | 684 | 0 | 0 | 0 | 0 |
| NS1 | P12882 | MYH1 | 1 | 602.5 | 0 | 602.5 | 169.16 | 168.31 | 1 | 0 | 522 | 558 | 609 | 721 | 0 | 0 | 0 | 0 |
| NS1 | Q9Y623 | MYH4 | 1 | 687.5 | 0 | 687.5 | 192.79 | 191.32 | 1 | 0 | 574 | 629 | 701 | 846 | 0 | 0 | 0 | 0 |

| BAIT | PROTID | GENE | preyFreq | preyAvg_AvgCounts | preySD_AvgCounts | NS2B_AVG_SPC | NS2B_2-4_FC_A | NS2B_FC_B | NS2B_SP | NS2B_iREF | IP_SH-SY5Y_NS2B_R4 | IP_SH-SY5Y_NS2B_R3 | IP_SH-SY5Y_NS2B_R2 | IP_SH-SY5Y_GFP_R4 | IP_SH-SY5Y_GFP_R3 | IP_SH-SY5Y_GFP_R2 | IP_SH-SY5Y_GFP_R1 |
|---|---|---|---|---|---|---|---|---|---|---|---|---|---|---|---|---|---|
| NS2B | P62873-2 | GNB1 | 1 | 4 | 0 | 4 | 2.13 | 2.13 | 1 | 0 | 4 | 4 | 4 | 0 | 0 | 0 | 0 |
| NS2B | P15259 | PGAM2 | 1 | 4 | 0 | 4 | 2.14 | 2.1 | 1 | 0 | 3 | 3 | 6 | 0 | 0 | 0 | 0 |
| NS2B | P48059-2 | LIMS1 | 1 | 4.333333333 | 0 | 4.333333333 | 2.21 | 2.2 | 1 | 0 | 5 | 5 | 3 | 0 | 0 | 0 | 0 |
| NS2B | P18859-2 | ATP5J | 1 | 4.333333333 | 0 | 4.333333333 | 2.21 | 2.2 | 1 | 0 | 5 | 5 | 3 | 0 | 0 | 0 | 0 |
| NS2B | Q9UMR2-2 | DDX19B | 1 | 4.333333333 | 0 | 4.333333333 | 2.21 | 2.2 | 1 | 0 | 5 | 5 | 3 | 0 | 0 | 0 | 0 |
| NS2B | Q9Y2R0 | COA3 | 1 | 4.333333333 | 0 | 4.333333333 | 2.22 | 2.21 | 1 | 0 | 4 | 5 | 4 | 0 | 0 | 0 | 0 |
| NS2B | O95299 | NDUFA10 | 1 | 4.666666667 | 0 | 4.666666667 | 2.31 | 2.3 | 1 | 0 | 4 | 6 | 4 | 0 | 0 | 0 | 0 |
| NS2B | P56134-2 | ATP5J2 | 1 | 4.666666667 | 0 | 4.666666667 | 2.31 | 2.3 | 1 | 0 | 4 | 6 | 4 | 0 | 0 | 0 | 0 |
| NS2B | Q32P28-3 | P3H1 | 1 | 4.666666667 | 0 | 4.666666667 | 2.32 | 2.31 | 1 | 0 | 4 | 5 | 5 | 0 | 0 | 0 | 0 |
| NS2B | P13674-2 | P4HA1 | 1 | 5 | 0 | 5 | 2.4 | 2.29 | 1 | 0 | 9 | 3 | 3 | 0 | 0 | 0 | 0 |
| NS2B | O75477 | ERLIN1 | 1 | 5 | 0 | 5 | 2.4 | 2.39 | 1 | 0 | 6 | 5 | 4 | 0 | 0 | 0 | 0 |
| NS2B | Q9UNL2-2 | SSR3 | 1 | 5 | 0 | 5 | 2.41 | 2.4 | 1 | 0 | 4 | 6 | 5 | 0 | 0 | 0 | 0 |
| NS2B | P45880-1 | VDAC2 | 1 | 5 | 0 | 5 | 2.41 | 2.4 | 1 | 0 | 4 | 5 | 6 | 0 | 0 | 0 | 0 |
| NS2B | P63241 | EIF5A | 1 | 5 | 0 | 5 | 2.41 | 2.41 | 1 | 0 | 5 | 5 | 5 | 0 | 0 | 0 | 0 |
| NS2B | Q9H4M9 | EHD1 | 1 | 5 | 0 | 5 | 2.41 | 2.41 | 1 | 0 | 5 | 5 | 5 | 0 | 0 | 0 | 0 |
| NS2B | Q93045-2 | STMN2 | 1 | 5 | 0 | 5 | 2.41 | 2.41 | 1 | 0 | 5 | 5 | 5 | 0 | 0 | 0 | 0 |
| NS2B | Q9HCN8 | SDF2L1 | 1 | 5.333333333 | 0 | 5.333333333 | 2.49 | 2.48 | 1 | 0 | 6 | 6 | 4 | 0 | 0 | 0 | 0 |
| NS2B | O60313-10 | OPA1 | 1 | 5.333333333 | 0 | 5.333333333 | 2.49 | 2.47 | 1 | 0 | 7 | 5 | 4 | 0 | 0 | 0 | 0 |
| NS2B | P07197 | NEFM | 1 | 5.333333333 | 0 | 5.333333333 | 2.5 | 2.5 | 1 | 0 | 5 | 6 | 5 | 0 | 0 | 0 | 0 |
| NS2B | O14980 | XPO1 | 1 | 5.333333333 | 0 | 5.333333333 | 2.5 | 2.5 | 1 | 0 | 5 | 6 | 5 | 0 | 0 | 0 | 0 |
| NS2B | Q02338 | BDH1 | 1 | 5.333333333 | 0 | 5.333333333 | 2.51 | 2.49 | 1 | 0 | 6 | 4 | 6 | 0 | 0 | 0 | 0 |
| NS2B | P38606 | ATP6V1A | 1 | 5.666666667 | 0 | 5.666666667 | 2.59 | 2.59 | 1 | 0 | 6 | 6 | 5 | 0 | 0 | 0 | 0 |
| NS2B | P25786-2 | PSMA1 | 1 | 5.666666667 | 0 | 5.666666667 | 2.59 | 2.55 | 1 | 0 | 8 | 4 | 5 | 0 | 0 | 0 | 0 |
| NS2B | Q15070-2 | OXA1L | 1 | 5.666666667 | 0 | 5.666666667 | 2.59 | 2.59 | 1 | 0 | 6 | 6 | 5 | 0 | 0 | 0 | 0 |
| NS2B | P07919 | UQCRH | 1 | 5.666666667 | 0 | 5.666666667 | 2.59 | 2.55 | 1 | 0 | 8 | 5 | 4 | 0 | 0 | 0 | 0 |
| NS2B | Q9UNH7 | SNX6 | 1 | 5.666666667 | 0 | 5.666666667 | 2.59 | 2.59 | 1 | 0 | 6 | 6 | 5 | 0 | 0 | 0 | 0 |
| NS2B | P62820 | RAB1A | 1 | 5.666666667 | 0 | 5.666666667 | 2.6 | 2.59 | 1 | 0 | 6 | 5 | 6 | 0 | 0 | 0 | 0 |
| NS2B | Q9Y281 | CFL2 | 1 | 6 | 0 | 6 | 2.68 | 2.65 | 1 | 0 | 7 | 7 | 4 | 0 | 0 | 0 | 0 |
| NS2B | Q9BSJ8-2 | ESYT1 | 1 | 6 | 0 | 6 | 2.69 | 2.69 | 1 | 0 | 6 | 6 | 6 | 0 | 0 | 0 | 0 |
| NS2B | Q8NBJ5 | COLGALT1 | 1 | 6.333333333 | 0 | 6.333333333 | 2.78 | 2.78 | 1 | 0 | 7 | 6 | 6 | 0 | 0 | 0 | 0 |
| NS2B | Q9H0U4 | RAB1B | 1 | 6.333333333 | 0 | 6.333333333 | 2.78 | 2.77 | 1 | 0 | 7 | 7 | 5 | 0 | 0 | 0 | 0 |
| NS2B | P61604 | HSPE1 | 1 | 6.333333333 | 0 | 6.333333333 | 2.79 | 2.77 | 1 | 0 | 5 | 7 | 7 | 0 | 0 | 0 | 0 |
| NS2B | P27824-2 | CANX | 1 | 6.333333333 | 0 | 6.333333333 | 2.79 | 2.77 | 1 | 0 | 7 | 5 | 7 | 0 | 0 | 0 | 0 |
| NS2B | P27797 | CALR | 1 | 6.333333333 | 0 | 6.333333333 | 2.79 | 2.78 | 1 | 0 | 6 | 6 | 7 | 0 | 0 | 0 | 0 |
| NS2B | P30153 | PPP2R1A | 1 | 6.666666667 | 0 | 6.666666667 | 2.87 | 2.85 | 1 | 0 | 8 | 7 | 5 | 0 | 0 | 0 | 0 |
| NS2B | O14579-3 | COPE | 1 | 6.666666667 | 0 | 6.666666667 | 2.87 | 2.86 | 1 | 0 | 6 | 8 | 6 | 0 | 0 | 0 | 0 |
| NS2B | P11172 | UMPS | 1 | 6.666666667 | 0 | 6.666666667 | 2.87 | 2.86 | 1 | 0 | 8 | 6 | 6 | 0 | 0 | 0 | 0 |
| NS2B | P26006 | ITGA3 | 1 | 6.666666667 | 0 | 6.666666667 | 2.87 | 2.86 | 1 | 0 | 8 | 6 | 6 | 0 | 0 | 0 | 0 |
| NS2B | P62847 | RPS24 | 1 | 6.666666667 | 0 | 6.666666667 | 2.88 | 2.86 | 1 | 0 | 8 | 5 | 7 | 0 | 0 | 0 | 0 |
| NS2B | P18669 | PGAM1 | 1 | 6.666666667 | 0 | 6.666666667 | 2.89 | 2.84 | 1 | 0 | 5 | 6 | 9 | 0 | 0 | 0 | 0 |
| NS2B | G5E9W2 | CHPF2 | 1 | 7 | 0 | 7 | 2.96 | 2.93 | 1 | 0 | 9 | 7 | 5 | 0 | 0 | 0 | 0 |
| NS2B | Q92616 | GCN1 | 1 | 7 | 0 | 7 | 2.96 | 2.93 | 1 | 0 | 9 | 7 | 5 | 0 | 0 | 0 | 0 |
| NS2B | P24534 | EEF1B2 | 1 | 7 | 0 | 7 | 2.97 | 2.96 | 1 | 0 | 6 | 8 | 7 | 0 | 0 | 0 | 0 |
| NS2B | Q14203-3 | DCTN1 | 1 | 7 | 0 | 7 | 2.97 | 2.96 | 1 | 0 | 8 | 7 | 6 | 0 | 0 | 0 | 0 |
| NS2B | P42704 | LRPPRC | 1 | 7.333333333 | 0 | 7.333333333 | 3.04 | 2.97 | 1 | 0 | 9 | 9 | 4 | 0 | 0 | 0 | 0 |
| NS2B | P09172 | DBH | 1 | 7.333333333 | 0 | 7.333333333 | 3.06 | 3.05 | 1 | 0 | 8 | 8 | 6 | 0 | 0 | 0 | 0 |
| NS2B | Q13015 | MLLT11 | 1 | 7.333333333 | 0 | 7.333333333 | 3.06 | 3.06 | 1 | 0 | 7 | 8 | 7 | 0 | 0 | 0 | 0 |
| NS2B | Q58FF3 | HSP90B2P | 1 | 7.333333333 | 0 | 7.333333333 | 3.06 | 3.06 | 1 | 0 | 8 | 7 | 7 | 0 | 0 | 0 | 0 |
| NS2B | P52292 | KPNA2 | 1 | 7.333333333 | 0 | 7.333333333 | 3.07 | 3.06 | 1 | 0 | 7 | 7 | 8 | 0 | 0 | 0 | 0 |
| NS2B | Q15773 | MLF2 | 1 | 7.666666667 | 0 | 7.666666667 | 3.16 | 3.15 | 1 | 0 | 9 | 7 | 7 | 0 | 0 | 0 | 0 |
| NS2B | O75746 | SLC25A12 | 1 | 7.666666667 | 0 | 7.666666667 | 3.16 | 3.12 | 1 | 0 | 10 | 6 | 7 | 0 | 0 | 0 | 0 |
| NS2B | P54136-2 | RARS | 1 | 7.666666667 | 0 | 7.666666667 | 3.16 | 3.14 | 1 | 0 | 9 | 6 | 8 | 0 | 0 | 0 | 0 |
| NS2B | Q9UBS4 | DNAJB11 | 1 | 7.666666667 | 0 | 7.666666667 | 3.16 | 3.16 | 1 | 0 | 7 | 8 | 8 | 0 | 0 | 0 | 0 |
| NS2B | Q9Y3Z3 | SAMHD1 | 1 | 7.666666667 | 0 | 7.666666667 | 3.16 | 3.16 | 1 | 0 | 7 | 8 | 8 | 0 | 0 | 0 | 0 |
| NS2B | P42025 | ACTR1B | 1 | 7.666666667 | 0 | 7.666666667 | 3.16 | 3.16 | 1 | 0 | 7 | 8 | 8 | 0 | 0 | 0 | 0 |
| NS2B | P50993 | ATP1A2 | 1 | 7.666666667 | 0 | 7.666666667 | 3.17 | 3.14 | 1 | 0 | 6 | 8 | 9 | 0 | 0 | 0 | 0 |
| NS2B | Q8WYP5-2 | AHCTF1 | 1 | 8 | 0 | 8 | 3.24 | 3.16 | 1 | 0 | 12 | 6 | 6 | 0 | 0 | 0 | 0 |
| NS2B | P13995-2 | MTHFD2 | 1 | 8 | 0 | 8 | 3.25 | 3.25 | 1 | 0 | 8 | 8 | 8 | 0 | 0 | 0 | 0 |
| NS2B | O14949 | UQCRQ | 1 | 8 | 0 | 8 | 3.25 | 3.2 | 1 | 0 | 11 | 6 | 7 | 0 | 0 | 0 | 0 |
| NS2B | Q9UBE0-2 | SAE1 | 1 | 8.333333333 | 0 | 8.333333333 | 3.34 | 3.3 | 1 | 0 | 11 | 7 | 7 | 0 | 0 | 0 | 0 |
| NS2B | A0A8V8TQ18 | CLTC | 1 | 8.333333333 | 0 | 8.333333333 | 3.35 | 3.34 | 1 | 0 | 7 | 9 | 9 | 0 | 0 | 0 | 0 |
| NS2B | Q9NSE4 | IARS2 | 1 | 8.333333333 | 0 | 8.333333333 | 3.35 | 3.33 | 1 | 0 | 10 | 7 | 8 | 0 | 0 | 0 | 0 |
| NS2B | O43251-6 | RBFOX2 | 1 | 8.666666667 | 0 | 8.666666667 | 3.43 | 3.42 | 1 | 0 | 10 | 9 | 7 | 0 | 0 | 0 | 0 |
| NS2B | Q9BSD7 | NTPCR | 1 | 8.666666667 | 0 | 8.666666667 | 3.43 | 3.42 | 1 | 0 | 9 | 10 | 7 | 0 | 0 | 0 | 0 |
| NS2B | O43615 | TIMM44 | 1 | 8.666666667 | 0 | 8.666666667 | 3.43 | 3.38 | 1 | 0 | 11 | 9 | 6 | 0 | 0 | 0 | 0 |
| NS2B | Q5JPH6 | EARS2 | 1 | 8.666666667 | 0 | 8.666666667 | 3.43 | 3.38 | 1 | 0 | 12 | 7 | 7 | 0 | 0 | 0 | 0 |
| NS2B | Q9UNX3 | RPL26L1 | 1 | 8.666666667 | 0 | 8.666666667 | 3.44 | 3.44 | 1 | 0 | 8 | 9 | 9 | 0 | 0 | 0 | 0 |
| NS2B | Q53H12 | AGK | 1 | 8.666666667 | 0 | 8.666666667 | 3.44 | 3.42 | 1 | 0 | 10 | 7 | 9 | 0 | 0 | 0 | 0 |
| NS2B | A1L0T0 | ILVBL | 1 | 8.666666667 | 0 | 8.666666667 | 3.44 | 3.32 | 1 | 0 | 13 | 5 | 8 | 0 | 0 | 0 | 0 |
| NS2B | P16949 | STMN1 | 1 | 8.666666667 | 0 | 8.666666667 | 3.45 | 3.42 | 1 | 0 | 7 | 9 | 10 | 0 | 0 | 0 | 0 |
| NS2B | Q99798 | ACO2 | 1 | 8.666666667 | 0 | 8.666666667 | 3.46 | 3.37 | 1 | 0 | 6 | 8 | 12 | 0 | 0 | 0 | 0 |

| | | | | | | | | | | | | | | | | |
|---|---|---|---|---|---|---|---|---|---|---|---|---|---|---|---|---|
| NS2B | F8W543 | NAP1L1 | 1 | 9 | 0 | 9 | 3.53 | 3.52 | 1 | 0 | 10 | 9 | 8 | 0 | 0 | 0 | 0 |
| NS2B | O00165-2 | HAX1 | 1 | 9 | 0 | 9 | 3.53 | 3.48 | 1 | 0 | 12 | 7 | 8 | 0 | 0 | 0 | 0 |
| NS2B | P17812 | CTPS1 | 1 | 9 | 0 | 9 | 3.53 | 3.52 | 1 | 0 | 9 | 10 | 8 | 0 | 0 | 0 | 0 |
| NS2B | P56962 | STX17 | 1 | 9 | 0 | 9 | 3.53 | 3.53 | 1 | 0 | 9 | 9 | 9 | 0 | 0 | 0 | 0 |
| NS2B | Q99470 | SDF2 | 1 | 9 | 0 | 9 | 3.53 | 3.52 | 1 | 0 | 10 | 9 | 8 | 0 | 0 | 0 | 0 |
| NS2B | Q53GQ0 | HSD17B12 | 1 | 9 | 0 | 9 | 3.54 | 3.51 | 1 | 0 | 10 | 7 | 10 | 0 | 0 | 0 | 0 |
| NS2B | P14314-2 | PRKCSH | 1 | 9 | 0 | 9 | 3.55 | 3.49 | 1 | 0 | 7 | 8 | 12 | 0 | 0 | 0 | 0 |
| NS2B | P69905 | HBA2;HBA1 | 1 | 9 | 0 | 9 | 3.56 | 3.45 | 1 | 0 | 6 | 8 | 13 | 0 | 0 | 0 | 0 |
| NS2B | Q9Y679 | AUP1 | 1 | 9.333333333 | 0 | 9.333333333 | 3.63 | 3.58 | 1 | 0 | 12 | 7 | 9 | 0 | 0 | 0 | 0 |
| NS2B | Q9ULX6-2 | AKAP8L | 1 | 9.333333333 | 0 | 9.333333333 | 3.63 | 3.63 | 1 | 0 | 9 | 9 | 10 | 0 | 0 | 0 | 0 |
| NS2B | P47897-2 | QARS | 1 | 9.666666667 | 0 | 9.666666667 | 3.69 | 3.57 | 1 | 0 | 11 | 13 | 5 | 0 | 0 | 0 | 0 |
| NS2B | P20674 | COX5A | 1 | 9.666666667 | 0 | 9.666666667 | 3.72 | 3.72 | 1 | 0 | 10 | 9 | 10 | 0 | 0 | 0 | 0 |
| NS2B | P06753 | TPM3 | 1 | 9.666666667 | 0 | 9.666666667 | 3.75 | 3.63 | 1 | 0 | 7 | 8 | 14 | 0 | 0 | 0 | 0 |
| NS2B | Q03252 | LMNB2 | 1 | 10 | 0 | 10 | 3.79 | 3.67 | 1 | 0 | 9 | 15 | 6 | 0 | 0 | 0 | 0 |
| NS2B | Q96A33 | CCDC47 | 1 | 10 | 0 | 10 | 3.8 | 3.72 | 1 | 0 | 14 | 9 | 7 | 0 | 0 | 0 | 0 |
| NS2B | O95573 | ACSL3 | 1 | 10 | 0 | 10 | 3.8 | 3.65 | 1 | 0 | 16 | 7 | 7 | 0 | 0 | 0 | 0 |
| NS2B | P50990 | CCT8 | 1 | 10 | 0 | 10 | 3.81 | 3.81 | 1 | 0 | 10 | 11 | 9 | 0 | 0 | 0 | 0 |
| NS2B | P54136 | RARS | 1 | 10 | 0 | 10 | 3.82 | 3.79 | 1 | 0 | 12 | 8 | 10 | 0 | 0 | 0 | 0 |
| NS2B | P67936 | TPM4 | 1 | 10 | 0 | 10 | 3.83 | 3.77 | 1 | 0 | 8 | 9 | 13 | 0 | 0 | 0 | 0 |
| NS2B | P56192 | MARS | 1 | 10.33333333 | 0 | 10.33333333 | 3.89 | 3.84 | 1 | 0 | 12 | 12 | 7 | 0 | 0 | 0 | 0 |
| NS2B | Q9UN37 | VPS4A | 1 | 10.33333333 | 0 | 10.33333333 | 3.9 | 3.89 | 1 | 0 | 10 | 12 | 9 | 0 | 0 | 0 | 0 |
| NS2B | Q9NYU2-2 | UGGT1 | 1 | 10.33333333 | 0 | 10.33333333 | 3.9 | 3.9 | 1 | 0 | 11 | 11 | 9 | 0 | 0 | 0 | 0 |
| NS2B | Q9BRK5 | SDF4 | 1 | 10.33333333 | 0 | 10.33333333 | 3.9 | 3.89 | 1 | 0 | 12 | 10 | 9 | 0 | 0 | 0 | 0 |
| NS2B | P08865 | RPSA | 1 | 10.33333333 | 0 | 10.33333333 | 3.91 | 3.91 | 1 | 0 | 11 | 10 | 10 | 0 | 0 | 0 | 0 |
| NS2B | P40926 | MDH2 | 1 | 10.33333333 | 0 | 10.33333333 | 3.92 | 3.84 | 1 | 0 | 7 | 11 | 13 | 0 | 0 | 0 | 0 |
| NS2B | Q5JTZ9 | AARS2 | 1 | 10.66666667 | 0 | 10.66666667 | 3.99 | 3.94 | 1 | 0 | 14 | 10 | 8 | 0 | 0 | 0 | 0 |
| NS2B | O75600-2 | GCAT | 1 | 10.66666667 | 0 | 10.66666667 | 3.99 | 3.97 | 1 | 0 | 13 | 10 | 9 | 0 | 0 | 0 | 0 |
| NS2B | Q8WTW3 | COG1 | 1 | 10.66666667 | 0 | 10.66666667 | 3.99 | 3.95 | 1 | 0 | 11 | 13 | 8 | 0 | 0 | 0 | 0 |
| NS2B | O95864 | FADS2 | 1 | 10.66666667 | 0 | 10.66666667 | 3.99 | 3.86 | 1 | 0 | 16 | 9 | 7 | 0 | 0 | 0 | 0 |
| NS2B | P11310-2 | ACADM | 1 | 10.66666667 | 0 | 10.66666667 | 4 | 3.99 | 1 | 0 | 10 | 12 | 10 | 0 | 0 | 0 | 0 |
| NS2B | Q6KCM7-2 | SLC25A25 | 1 | 11 | 0 | 11 | 4.09 | 4.05 | 1 | 0 | 14 | 10 | 9 | 0 | 0 | 0 | 0 |
| NS2B | P33991 | MCM4 | 1 | 11 | 0 | 11 | 4.09 | 4.07 | 1 | 0 | 13 | 11 | 9 | 0 | 0 | 0 | 0 |
| NS2B | P53396 | ACLY | 1 | 11 | 0 | 11 | 4.11 | 4.06 | 1 | 0 | 9 | 10 | 14 | 0 | 0 | 0 | 0 |
| NS2B | A7XZE4 | A7XZE4 | 1 | 11.33333333 | 0 | 11.33333333 | 4.24 | 3.95 | 1 | 0 | 8 | 7 | 19 | 0 | 0 | 0 | 0 |
| NS2B | Q16891-2 | IMMT | 1 | 11.66666667 | 0 | 11.66666667 | 4.27 | 4.24 | 1 | 0 | 12 | 14 | 9 | 0 | 0 | 0 | 0 |
| NS2B | O75947-2 | ATP5H | 1 | 11.66666667 | 0 | 11.66666667 | 4.28 | 4.27 | 1 | 0 | 11 | 13 | 11 | 0 | 0 | 0 | 0 |
| NS2B | Q15942 | ZYX | 1 | 11.66666667 | 0 | 11.66666667 | 4.28 | 4.27 | 1 | 0 | 13 | 12 | 10 | 0 | 0 | 0 | 0 |
| NS2B | P52701-4 | MSH6 | 1 | 11.66666667 | 0 | 11.66666667 | 4.29 | 4.24 | 1 | 0 | 14 | 9 | 12 | 0 | 0 | 0 | 0 |
| NS2B | P00558-2 | PGK1 | 1 | 12 | 0 | 12 | 4.41 | 4.28 | 1 | 0 | 9 | 10 | 17 | 0 | 0 | 0 | 0 |
| NS2B | O14983-2 | ATP2A1 | 1 | 12.33333333 | 0 | 12.33333333 | 4.46 | 4.46 | 1 | 0 | 13 | 13 | 11 | 0 | 0 | 0 | 0 |
| NS2B | Q9UKV5 | AMFR | 1 | 12.33333333 | 0 | 12.33333333 | 4.47 | 4.41 | 1 | 0 | 16 | 10 | 11 | 0 | 0 | 0 | 0 |
| NS2B | H0YK48 | TPM1 | 1 | 12.33333333 | 0 | 12.33333333 | 4.48 | 4.45 | 1 | 0 | 13 | 10 | 14 | 0 | 0 | 0 | 0 |
| NS2B | P78371 | CCT2 | 1 | 12.33333333 | 0 | 12.33333333 | 4.48 | 4.46 | 1 | 0 | 13 | 11 | 13 | 0 | 0 | 0 | 0 |
| NS2B | P04843 | RPN1 | 1 | 20.33333333 | 0 | 20.33333333 | 4.49 | 4.04 | 1 | 0 | 21 | 22 | 18 | 1 | 0 | 0 | 1 |
| NS2B | Q8NBM4-2 | UBAC2 | 1 | 12.66666667 | 0 | 12.66666667 | 4.56 | 4.56 | 1 | 0 | 14 | 12 | 12 | 0 | 0 | 0 | 0 |
| NS2B | Q8IXI1 | RHOT2 | 1 | 12.66666667 | 0 | 12.66666667 | 4.56 | 4.56 | 1 | 0 | 12 | 14 | 12 | 0 | 0 | 0 | 0 |
| NS2B | Q14318-2 | FKBP8 | 1 | 12.66666667 | 0 | 12.66666667 | 4.57 | 4.56 | 1 | 0 | 13 | 12 | 13 | 0 | 0 | 0 | 0 |
| NS2B | P00558 | PGK1 | 1 | 13 | 0 | 13 | 4.69 | 4.56 | 1 | 0 | 10 | 11 | 18 | 0 | 0 | 0 | 0 |
| NS2B | P09493-6 | TPM1 | 1 | 13.66666667 | 0 | 13.66666667 | 4.86 | 4.81 | 1 | 0 | 14 | 11 | 16 | 0 | 0 | 0 | 0 |
| NS2B | Q6NUK1-2 | SLC25A24 | 1 | 14.66666667 | 0 | 14.66666667 | 5.11 | 5.04 | 1 | 0 | 19 | 14 | 11 | 0 | 0 | 0 | 0 |
| NS2B | P04075 | ALDOA | 1 | 14.66666667 | 0 | 14.66666667 | 5.17 | 4.98 | 1 | 0 | 10 | 13 | 21 | 0 | 0 | 0 | 0 |
| NS2B | Q6ZN40 | TPM1 | 1 | 15 | 0 | 15 | 5.25 | 5.12 | 1 | 0 | 14 | 11 | 20 | 0 | 0 | 0 | 0 |
| NS2B | P00367 | GLUD1 | 1 | 15.33333333 | 0 | 15.33333333 | 5.31 | 5.29 | 1 | 0 | 14 | 18 | 14 | 0 | 0 | 0 | 0 |
| NS2B | P05023-3 | ATP1A1 | 1 | 15.33333333 | 0 | 15.33333333 | 5.32 | 5.29 | 1 | 0 | 17 | 13 | 16 | 0 | 0 | 0 | 0 |
| NS2B | Q9H5Q4 | TFB2M | 1 | 15.33333333 | 0 | 15.33333333 | 5.32 | 5.31 | 1 | 0 | 16 | 14 | 16 | 0 | 0 | 0 | 0 |
| NS2B | P06753-6 | TPM3 | 1 | 15.33333333 | 0 | 15.33333333 | 5.34 | 5.26 | 1 | 0 | 12 | 15 | 19 | 0 | 0 | 0 | 0 |
| NS2B | O95394-3 | PGM3 | 1 | 15.66666667 | 0 | 15.66666667 | 5.4 | 5.39 | 1 | 0 | 18 | 15 | 14 | 0 | 0 | 0 | 0 |
| NS2B | Q9UHD8-2 | SEPTIN9 | 1 | 15.66666667 | 0 | 15.66666667 | 5.41 | 5.4 | 1 | 0 | 17 | 15 | 15 | 0 | 0 | 0 | 0 |
| NS2B | P51570 | GALK1 | 1 | 16 | 0 | 16 | 5.49 | 5.44 | 1 | 0 | 20 | 15 | 13 | 0 | 0 | 0 | 0 |
| NS2B | P13533 | MYH6 | 1 | 16.33333333 | 0 | 16.33333333 | 5.65 | 5.22 | 1 | 0 | 8 | 15 | 26 | 0 | 0 | 0 | 0 |
| NS2B | Q13418 | ILK | 1 | 16.66666667 | 0 | 16.66666667 | 5.69 | 5.69 | 1 | 0 | 17 | 16 | 17 | 0 | 0 | 0 | 0 |
| NS2B | P12883 | MYH7 | 1 | 16.66666667 | 0 | 16.66666667 | 5.75 | 5.27 | 1 | 0 | 8 | 15 | 27 | 0 | 0 | 0 | 0 |
| NS2B | P35998 | PSMC2 | 1 | 27.33333333 | 0 | 27.33333333 | 5.79 | 5.19 | 1 | 0 | 31 | 28 | 23 | 1 | 1 | 0 | 0 |
| NS2B | P09493 | TPM1 | 1 | 17.66666667 | 0 | 17.66666667 | 6.01 | 5.87 | 1 | 0 | 17 | 13 | 23 | 0 | 0 | 0 | 0 |
| NS2B | P06753-5 | TPM3 | 1 | 18 | 0 | 18 | 6.1 | 5.99 | 1 | 0 | 15 | 16 | 23 | 0 | 0 | 0 | 0 |
| NS2B | Q9NZ01 | TECR | 1 | 22 | 0 | 22 | 6.1 | 5.81 | 1 | 0 | 24 | 22 | 20 | 1 | 0 | 0 | 0 |
| NS2B | R4GNH3 | PSMC3 | 1 | 18.33333333 | 0 | 18.33333333 | 6.15 | 6.15 | 1 | 0 | 19 | 19 | 17 | 0 | 0 | 0 | 0 |
| NS2B | Q14204 | DYNC1H1 | 1 | 18.66666667 | 0 | 18.66666667 | 6.24 | 6.23 | 1 | 0 | 20 | 20 | 16 | 0 | 0 | 0 | 0 |
| NS2B | P62191-2 | PSMC1 | 1 | 18.66666667 | 0 | 18.66666667 | 6.26 | 6.25 | 1 | 0 | 19 | 18 | 19 | 0 | 0 | 0 | 0 |
| NS2B | Q7Z6Z7-2 | HUWE1 | 1 | 26 | 0 | 26 | 6.68 | 6.26 | 1 | 0 | 25 | 29 | 24 | 0 | 0 | 1 | 0 |
| NS2B | A0A2R8YDQ9 | SUCLA2 | 1 | 20.66666667 | 0 | 20.66666667 | 6.82 | 6.72 | 1 | 0 | 26 | 16 | 20 | 0 | 0 | 0 | 0 |
| NS2B | Q9UJS0 | SLC25A13 | 1 | 22 | 0 | 22 | 7.18 | 7.17 | 1 | 0 | 25 | 21 | 20 | 0 | 0 | 0 | 0 |
| NS2B | A0A2R8YEA7 | ACTB | 1 | 23.33333333 | 0 | 23.33333333 | 7.57 | 7.55 | 1 | 0 | 21 | 25 | 24 | 0 | 0 | 0 | 0 |
| NS2B | A0A0C4DG89 | DDX46 | 1 | 23.66666667 | 0 | 23.66666667 | 7.64 | 7.61 | 1 | 0 | 24 | 27 | 20 | 0 | 0 | 0 | 0 |
| NS2B | Q14697-2 | GANAB | 1 | 24 | 0 | 24 | 7.75 | 7.74 | 1 | 0 | 26 | 23 | 23 | 0 | 0 | 0 | 0 |
| NS2B | Q9Y4W6 | AFG3L2 | 1 | 25.66666667 | 0 | 25.66666667 | 8.22 | 8.22 | 1 | 0 | 27 | 25 | 25 | 0 | 0 | 0 | 0 |
| NS2B | Q5T9A4 | ATAD3B | 1 | 26 | 0 | 26 | 8.3 | 8.29 | 1 | 0 | 27 | 28 | 23 | 0 | 0 | 0 | 0 |
| NS2B | P06733-2 | ENO1 | 1 | 33 | 0 | 33 | 10.32 | 10.21 | 1 | 0 | 28 | 32 | 39 | 0 | 0 | 0 | 0 |
| NS2B | Q9Y285 | FARSA | 1 | 35 | 0 | 35 | 10.83 | 10.73 | 1 | 0 | 43 | 32 | 30 | 0 | 0 | 0 | 0 |
| NS2B | P16615-2 | ATP2A2 | 1 | 42.33333333 | 0 | 42.33333333 | 12.88 | 12.78 | 1 | 0 | 51 | 41 | 35 | 0 | 0 | 0 | 0 |
| NS2B | P11055 | MYH3 | 1 | 92.66666667 | 0 | 92.66666667 | 27.22 | 25.26 | 1 | 0 | 48 | 109 | 121 | 0 | 0 | 0 | 0 |
| NS2B | Q9UKX2 | MYH2 | 1 | 112.6666667 | 0 | 112.6666667 | 32.93 | 30.19 | 1 | 0 | 56 | 128 | 154 | 0 | 0 | 0 | 0 |
| NS2B | P12882 | MYH1 | 1 | 117.3333333 | 0 | 117.3333333 | 34.26 | 31.53 | 1 | 0 | 60 | 130 | 162 | 0 | 0 | 0 | 0 |
| NS2B | Q9Y623 | MYH4 | 1 | 135 | 0 | 135 | 39.27 | 36.36 | 1 | 0 | 72 | 147 | 186 | 0 | 0 | 0 | 0 |
| NS2B | P13535 | MYH8 | 1 | 180 | 0 | 180 | 51.83 | 51.17 | 1 | 0 | 148 | 183 | 209 | 0 | 0 | 0 | 0 |

| BAIT | PROTID | GENE | preyFreq | preyAvg_AvgCounts | preySD_AvgCounts | NS3_AVG_SPC | NS3_FC_A | NS3_FC_B | NS3_SP | NS3_iREF | IP_SH-SY5Y_NS3_R4 | IP_SH-SY5Y_NS3_R3 | IP_SH-SY5Y_NS3_R2 | IP_SH-SY5Y_NS3_R1 | IP_SH-SY5Y_GFP_R4 | IP_SH-SY5Y_GFP_R3 | IP_SH-SY5Y_GFP_R2 | IP_SH-SY5Y_GFP_R1 |
|---|---|---|---|---|---|---|---|---|---|---|---|---|---|---|---|---|---|---|
| NS3 | Q13185 | CBX3 | 1 | 5 | 0 | 5 | 2.62 | 2.61 | 1 | 0 | 6 | 6 | 5 | 3 | 0 | 0 | 0 | 0 |
| NS3 | Q03252 | LMNB2 | 1 | 5 | 0 | 5 | 2.69 | 2.67 | 1 | 0 | 6 | 5 | 4 | 5 | 0 | 0 | 0 | 0 |
| NS3 | P99999 | CYCS | 1 | 6 | 0 | 6 | 2.97 | 2.95 | 1 | 0 | 8 | 6 | 6 | 4 | 0 | 0 | 0 | 0 |
| NS3 | P00167 | CYB5A | 1 | 6 | 0 | 6 | 2.97 | 2.95 | 1 | 0 | 6 | 6 | 8 | 4 | 0 | 0 | 0 | 0 |
| NS3 | O00217 | NDUFS8 | 1 | 7 | 0 | 7 | 3.44 | 3.34 | 1 | 0 | 6 | 7 | 6 | 9 | 0 | 0 | 0 | 0 |
| NS3 | P09104 | ENO2 | 1 | 8.5 | 0 | 8.5 | 3.7 | 3.64 | 1 | 0 | 9 | 13 | 8 | 4 | 0 | 0 | 0 | 0 |
| NS3 | P36871 | PGM1 | 1 | 9.5 | 0 | 9.5 | 4.02 | 3.94 | 1 | 0 | 10 | 13 | 11 | 4 | 0 | 0 | 0 | 0 |
| NS3 | P50990 | CCT8 | 1 | 9.5 | 0 | 9.5 | 4.45 | 3.95 | 1 | 0 | 5 | 5 | 13 | 15 | 0 | 0 | 0 | 0 |
| NS3 | P07205 | PGK2 | 1 | 10.25 | 0 | 10.25 | 4.47 | 4.43 | 1 | 0 | 10 | 10 | 11 | 10 | 0 | 0 | 0 | 0 |
| NS3 | P15090 | FABP4 | 1 | 12 | 0 | 12 | 4.82 | 4.71 | 1 | 0 | 13 | 15 | 15 | 5 | 0 | 0 | 0 | 0 |
| NS3 | Q6NZI2 | CAVIN1 | 1 | 12.75 | 0 | 12.75 | 5 | 4.79 | 1 | 0 | 16 | 17 | 14 | 4 | 0 | 0 | 0 | 0 |
| NS3 | P34932 | HSPA4 | 1 | 12 | 0 | 12 | 5.02 | 5 | 1 | 0 | 13 | 13 | 11 | 11 | 0 | 0 | 0 | 0 |
| NS3 | Q02218-2 | OGDH | 1 | 13.5 | 0 | 13.5 | 5.32 | 5.24 | 1 | 0 | 16 | 19 | 12 | 7 | 0 | 0 | 0 | 0 |
| NS3 | P08590 | MYL3 | 1 | 15 | 0 | 15 | 5.74 | 5.41 | 1 | 0 | 16 | 28 | 8 | 8 | 0 | 0 | 0 | 0 |
| NS3 | A0A2R8YEA7 | ACTB | 1 | 14.5 | 0 | 14.5 | 5.81 | 5.77 | 1 | 0 | 14 | 15 | 18 | 11 | 0 | 0 | 0 | 0 |
| NS3 | P47929 | LGALS7;LGALS7B | 1 | 22.25 | 0 | 22.25 | 5.96 | 5.02 | 1 | 0 | 18 | 20 | 19 | 32 | 0 | 0 | 2 | 0 |
| NS3 | P08559-2 | PDHA1 | 1 | 18 | 0 | 18 | 6.64 | 6.24 | 1 | 0 | 23 | 23 | 21 | 5 | 0 | 0 | 0 | 0 |
| NS3 | P06753-2 | TPM3 | 1 | 32.25 | 0 | 32.25 | 7.58 | 6.74 | 1 | 0 | 38 | 43 | 31 | 17 | 1 | 0 | 0 | 1 |
| NS3 | A0A0C4DG89 | DDX46 | 1 | 19 | 0 | 19 | 7.69 | 7.21 | 1 | 0 | 20 | 12 | 19 | 25 | 0 | 0 | 0 | 0 |
| NS3 | F8WA37 | TNNT3 | 1 | 20 | 0 | 20 | 7.71 | 7.64 | 1 | 0 | 21 | 23 | 17 | 19 | 0 | 0 | 0 | 0 |
| NS3 | P45378-7 | TNNT3 | 1 | 20 | 0 | 20 | 7.71 | 7.64 | 1 | 0 | 21 | 23 | 17 | 19 | 0 | 0 | 0 | 0 |
| NS3 | P00558-2 | PGK1 | 1 | 21.5 | 0 | 21.5 | 7.95 | 7.88 | 1 | 0 | 26 | 25 | 23 | 12 | 0 | 0 | 0 | 0 |
| NS3 | Q13813 | SPTAN1 | 1 | 22 | 0 | 22 | 7.98 | 7.74 | 1 | 0 | 27 | 29 | 23 | 9 | 0 | 0 | 0 | 0 |
| NS3 | A7E2Y1 | MYH7B | 1 | 23 | 0 | 23 | 8.17 | 7.1 | 1 | 0 | 33 | 42 | 7 | 10 | 0 | 0 | 0 | 0 |
| NS3 | P13929 | ENO3 | 1 | 23.5 | 0 | 23.5 | 8.44 | 8.06 | 1 | 0 | 27 | 40 | 15 | 12 | 0 | 0 | 0 | 0 |
| NS3 | O14983-2 | ATP2A1 | 1 | 22.75 | 0 | 22.75 | 8.58 | 8.22 | 1 | 0 | 25 | 32 | 12 | 22 | 0 | 0 | 0 | 0 |
| NS3 | Q08043 | ACTN3 | 1 | 24.5 | 0 | 24.5 | 8.95 | 8.63 | 1 | 0 | 27 | 39 | 14 | 18 | 0 | 0 | 0 | 0 |
| NS3 | P05976 | MYL1 | 1 | 25 | 0 | 25 | 9.02 | 8.62 | 1 | 0 | 28 | 42 | 14 | 16 | 0 | 0 | 0 | 0 |
| NS3 | P18669 | PGAM1 | 1 | 25.25 | 0 | 25.25 | 9.19 | 9.14 | 1 | 0 | 28 | 30 | 28 | 15 | 0 | 0 | 0 | 0 |
| NS3 | P67936 | TPM4 | 1 | 26.25 | 0 | 26.25 | 9.28 | 8.76 | 1 | 0 | 35 | 42 | 16 | 12 | 0 | 0 | 0 | 0 |
| NS3 | A0A7I2V5A1 | ACO2 | 1 | 25.75 | 0 | 25.75 | 9.32 | 9.13 | 1 | 0 | 31 | 27 | 32 | 13 | 0 | 0 | 0 | 0 |
| NS3 | P00558 | PGK1 | 1 | 26 | 0 | 26 | 9.34 | 9.2 | 1 | 0 | 30 | 33 | 28 | 13 | 0 | 0 | 0 | 0 |
| NS3 | P78371 | CCT2 | 1 | 22.75 | 0 | 22.75 | 9.35 | 7.96 | 1 | 0 | 12 | 15 | 24 | 40 | 0 | 0 | 0 | 0 |
| NS3 | P06753-6 | TPM3 | 1 | 31.75 | 0 | 31.75 | 11.12 | 10.87 | 1 | 0 | 38 | 46 | 27 | 16 | 0 | 0 | 0 | 0 |
| NS3 | P04075 | ALDOA | 1 | 32.25 | 0 | 32.25 | 11.19 | 10.73 | 1 | 0 | 36 | 44 | 37 | 12 | 0 | 0 | 0 | 0 |
| NS3 | P06753-5 | TPM3 | 1 | 33.25 | 0 | 33.25 | 11.63 | 11.43 | 1 | 0 | 39 | 46 | 31 | 17 | 0 | 0 | 0 | 0 |
| NS3 | P16615-2 | ATP2A2 | 1 | 31.75 | 0 | 31.75 | 12.03 | 11.52 | 1 | 0 | 27 | 31 | 29 | 40 | 0 | 0 | 0 | 0 |
| NS3 | H0YK48 | TPM1 | 1 | 35.25 | 0 | 35.25 | 12.61 | 12.33 | 1 | 0 | 41 | 48 | 23 | 29 | 0 | 0 | 0 | 0 |
| NS3 | P06733-2 | ENO1 | 1 | 35.75 | 0 | 35.75 | 12.84 | 12.76 | 1 | 0 | 44 | 34 | 38 | 27 | 0 | 0 | 0 | 0 |
| NS3 | O95831 | AIFM1 | 1 | 35.25 | 0 | 35.25 | 13.22 | 12.7 | 1 | 0 | 36 | 31 | 31 | 43 | 0 | 0 | 0 | 0 |
| NS3 | P06753 | TPM3 | 1 | 39 | 0 | 39 | 13.27 | 12.43 | 1 | 0 | 53 | 62 | 24 | 17 | 0 | 0 | 0 | 0 |
| NS3 | A7XZE4 | A7XZE4 | 1 | 39.75 | 0 | 39.75 | 13.45 | 11.88 | 1 | 0 | 57 | 69 | 15 | 18 | 0 | 0 | 0 | 0 |
| NS3 | Q99798 | ACO2 | 1 | 39.25 | 0 | 39.25 | 13.9 | 13.85 | 1 | 0 | 42 | 42 | 46 | 27 | 0 | 0 | 0 | 0 |
| NS3 | P09493-6 | TPM1 | 1 | 40 | 0 | 40 | 14.1 | 13.67 | 1 | 0 | 51 | 54 | 24 | 31 | 0 | 0 | 0 | 0 |
| NS3 | P06732 | CKM | 1 | 48 | 0 | 48 | 16.67 | 15.58 | 1 | 0 | 53 | 78 | 22 | 39 | 0 | 0 | 0 | 0 |
| NS3 | Q6ZN40 | TPM1 | 1 | 49.25 | 0 | 49.25 | 16.97 | 16.29 | 1 | 0 | 66 | 69 | 28 | 34 | 0 | 0 | 0 | 0 |
| NS3 | P69905 | HBA2;HBA1 | 1 | 45 | 0 | 45 | 17.23 | 15.34 | 1 | 0 | 38 | 31 | 39 | 72 | 0 | 0 | 0 | 0 |
| NS3 | P09493 | TPM1 | 1 | 55 | 0 | 55 | 18.7 | 17.71 | 1 | 0 | 76 | 80 | 29 | 35 | 0 | 0 | 0 | 0 |
| NS3 | P13533 | MYH6 | 1 | 121.25 | 0 | 121.25 | 39.93 | 36.27 | 1 | 0 | 153 | 202 | 49 | 81 | 0 | 0 | 0 | 0 |
| NS3 | P12883 | MYH7 | 1 | 137.5 | 0 | 137.5 | 44.77 | 39.65 | 1 | 0 | 184 | 234 | 50 | 82 | 0 | 0 | 0 | 0 |
| NS3 | P11055 | MYH3 | 1 | 231.5 | 0 | 231.5 | 81.14 | 77.63 | 1 | 0 | 219 | 225 | 198 | 284 | 0 | 0 | 0 | 0 |
| NS3 | Q9UKX2 | MYH2 | 1 | 358 | 0 | 358 | 121.85 | 118.67 | 1 | 0 | 370 | 431 | 262 | 369 | 0 | 0 | 0 | 0 |
| NS3 | P12882 | MYH1 | 1 | 375.5 | 0 | 375.5 | 127.18 | 124.04 | 1 | 0 | 395 | 465 | 269 | 373 | 0 | 0 | 0 | 0 |
| NS3 | P13535 | MYH8 | 1 | 406.25 | 0 | 406.25 | 139.54 | 135.6 | 1 | 0 | 409 | 442 | 326 | 448 | 0 | 0 | 0 | 0 |
| NS3 | Q9Y623 | MYH4 | 1 | 422.25 | 0 | 422.25 | 144.17 | 139.92 | 1 | 0 | 427 | 499 | 311 | 452 | 0 | 0 | 0 | 0 |

| BAIT | PROTID | GENE | preyFreq | preyAvg_AvgCounts | preySD_AvgCounts | NS4A_AVG_SPC | NS4A_FC_A | NS4A_FC_B | NS4A_SP | NS4A_iREF | IP_SH-SY5Y_NS4A_R4 | IP_SH-SY5Y_NS4A_R3 | IP_SH-SY5Y_NS4A_R2 | IP_SH-SY5Y_NS4A_R1 | IP_SH-SY5Y_GFP_R4 | IP_SH-SY5Y_GFP_R3 | IP_SH-SY5Y_GFP_R2 | IP_SH-SY5Y_GFP_R1 |
|---|---|---|---|---|---|---|---|---|---|---|---|---|---|---|---|---|---|---|
| NS4A | Q8TED1 | GPX8 | 1 | 4 | 0 | 4 | 2.66 | 2.64 | 1 | 0 | 3 | 3 | 6 | 4 | 0 | 0 | 0 | 0 |
| NS4A | O14683 | TP53I11 | 1 | 4.25 | 0 | 4.25 | 2.83 | 2.81 | 1 | 0 | 5 | 4 | 4 | 4 | 0 | 0 | 0 | 0 |
| NS4A | Q8WUY8 | NAT14 | 1 | 4.5 | 0 | 4.5 | 2.88 | 2.86 | 1 | 0 | 5 | 3 | 6 | 4 | 0 | 0 | 0 | 0 |
| NS4A | P09172 | DBH | 1 | 4.5 | 0 | 4.5 | 2.91 | 2.9 | 1 | 0 | 5 | 4 | 5 | 4 | 0 | 0 | 0 | 0 |
| NS4A | Q9GZP9 | DERL2 | 1 | 4.5 | 0 | 4.5 | 2.92 | 2.87 | 1 | 0 | 5 | 6 | 4 | 3 | 0 | 0 | 0 | 0 |
| NS4A | Q9UKV5 | AMFR | 1 | 4.5 | 0 | 4.5 | 2.95 | 2.92 | 1 | 0 | 5 | 4 | 4 | 5 | 0 | 0 | 0 | 0 |
| NS4A | P14406 | COX7A2 | 1 | 4.5 | 0 | 4.5 | 2.95 | 2.92 | 1 | 0 | 5 | 4 | 4 | 5 | 0 | 0 | 0 | 0 |
| NS4A | P27105 | STOM | 1 | 4.5 | 0 | 4.5 | 2.97 | 2.91 | 1 | 0 | 4 | 4 | 4 | 6 | 0 | 0 | 0 | 0 |
| NS4A | A0A087WYV9 | SYNJ2BP-COX16 | 1 | 4.75 | 0 | 4.75 | 3.04 | 3.02 | 1 | 0 | 5 | 4 | 5 | 5 | 0 | 0 | 0 | 0 |
| NS4A | Q9Y5Y0 | FLVCR1 | 1 | 4.75 | 0 | 4.75 | 3.08 | 3.02 | 1 | 0 | 5 | 4 | 4 | 6 | 0 | 0 | 0 | 0 |
| NS4A | Q8N0U8 | VKORC1L1 | 1 | 5 | 0 | 5 | 3.09 | 3.06 | 1 | 0 | 5 | 6 | 6 | 3 | 0 | 0 | 0 | 0 |
| NS4A | Q6P4E1-5 | CASC4 | 1 | 4.75 | 0 | 4.75 | 3.1 | 3.01 | 1 | 0 | 5 | 5 | 3 | 6 | 0 | 0 | 0 | 0 |
| NS4A | O76024 | WFS1 | 1 | 5 | 0 | 5 | 3.12 | 3.11 | 1 | 0 | 4 | 5 | 6 | 5 | 0 | 0 | 0 | 0 |
| NS4A | Q5H8A4-2 | PIGG | 1 | 5 | 0 | 5 | 3.12 | 3.11 | 1 | 0 | 5 | 4 | 6 | 5 | 0 | 0 | 0 | 0 |
| NS4A | Q9P0J0-2 | NDUFA13 | 1 | 5.25 | 0 | 5.25 | 3.12 | 3.07 | 1 | 0 | 5 | 4 | 9 | 3 | 0 | 0 | 0 | 0 |
| NS4A | O96008-2 | TOMM40 | 1 | 5 | 0 | 5 | 3.13 | 3.11 | 1 | 0 | 5 | 6 | 5 | 4 | 0 | 0 | 0 | 0 |
| NS4A | Q9NRK6 | ABCB10 | 1 | 5 | 0 | 5 | 3.17 | 3.1 | 1 | 0 | 7 | 4 | 4 | 5 | 0 | 0 | 0 | 0 |
| NS4A | P33527-2 | ABCC1 | 1 | 5.25 | 0 | 5.25 | 3.23 | 3.13 | 1 | 0 | 4 | 3 | 7 | 7 | 0 | 0 | 0 | 0 |
| NS4A | Q8WVX9 | FAR1 | 1 | 5.25 | 0 | 5.25 | 3.24 | 3.21 | 1 | 0 | 6 | 6 | 5 | 4 | 0 | 0 | 0 | 0 |
| NS4A | B7ZBM8 | STX16 | 1 | 5.5 | 0 | 5.5 | 3.33 | 3.3 | 1 | 0 | 5 | 4 | 7 | 6 | 0 | 0 | 0 | 0 |
| NS4A | Q8IXI2-2 | RHOT1 | 1 | 5.5 | 0 | 5.5 | 3.33 | 3.26 | 1 | 0 | 8 | 4 | 6 | 4 | 0 | 0 | 0 | 0 |
| NS4A | Q96G23 | CERS2 | 1 | 5.75 | 0 | 5.75 | 3.45 | 3.43 | 1 | 0 | 7 | 5 | 6 | 5 | 0 | 0 | 0 | 0 |

| | | | | | | | | | | | | | | | | | | |
|---|---|---|---|---|---|---|---|---|---|---|---|---|---|---|---|---|---|---|
| NS4A | Q5JTV8 | TOR1AIP1 | 1 | 5.75 | 0 | 5.75 | 3.45 | 3.44 | 1 | 0 | 6 | 6 | 6 | 5 | 0 | 0 | 0 | 0 |
| NS4A | O15400-2 | STX7 | 1 | 6 | 0 | 6 | 3.5 | 3.5 | 1 | 0 | 5 | 6 | 8 | 5 | 0 | 0 | 0 | 0 |
| NS4A | Q8NBI6 | XXYLT1 | 1 | 5.75 | 0 | 5.75 | 3.51 | 3.44 | 1 | 0 | 6 | 5 | 5 | 7 | 0 | 0 | 0 | 0 |
| NS4A | Q9P0T7 | TMEM9 | 1 | 6 | 0 | 6 | 3.52 | 3.49 | 1 | 0 | 6 | 4 | 8 | 6 | 0 | 0 | 0 | 0 |
| NS4A | Q9BX79-3 | STRA6 | 1 | 5.75 | 0 | 5.75 | 3.53 | 3.43 | 1 | 0 | 5 | 7 | 4 | 7 | 0 | 0 | 0 | 0 |
| NS4A | O14980 | XPO1 | 1 | 6 | 0 | 6 | 3.55 | 3.49 | 1 | 0 | 6 | 8 | 6 | 4 | 0 | 0 | 0 | 0 |
| NS4A | Q68DH5 | LMBRD2 | 1 | 6 | 0 | 6 | 3.56 | 3.52 | 1 | 0 | 5 | 5 | 7 | 7 | 0 | 0 | 0 | 0 |
| NS4A | P98194-2 | ATP2C1 | 1 | 6 | 0 | 6 | 3.59 | 3.55 | 1 | 0 | 5 | 6 | 6 | 7 | 0 | 0 | 0 | 0 |
| NS4A | P13073 | COX4I1 | 1 | 6.25 | 0 | 6.25 | 3.62 | 3.56 | 1 | 0 | 8 | 4 | 8 | 5 | 0 | 0 | 0 | 0 |
| NS4A | P19404 | NDUFV2 | 1 | 6.5 | 0 | 6.5 | 3.68 | 3.66 | 1 | 0 | 6 | 7 | 9 | 4 | 0 | 0 | 0 | 0 |
| NS4A | Q9BWL3-2 | C1ORF43 | 1 | 6.5 | 0 | 6.5 | 3.71 | 3.58 | 1 | 0 | 4 | 10 | 8 | 4 | 0 | 0 | 0 | 0 |
| NS4A | P39656-3 | DDOST | 1 | 10.75 | 0 | 10.75 | 3.73 | 3.34 | 1 | 0 | 12 | 11 | 11 | 9 | 1 | 0 | 0 | 1 |
| NS4A | Q9H4M9 | EHD1 | 1 | 6.5 | 0 | 6.5 | 3.75 | 3.73 | 1 | 0 | 6 | 5 | 8 | 7 | 0 | 0 | 0 | 0 |
| NS4A | Q8N6S5 | ARL6IP6 | 1 | 6.5 | 0 | 6.5 | 3.8 | 3.75 | 1 | 0 | 6 | 8 | 6 | 6 | 0 | 0 | 0 | 0 |
| NS4A | Q93050-1 | ATP6V0A1 | 1 | 6.75 | 0 | 6.75 | 3.8 | 3.76 | 1 | 0 | 7 | 7 | 9 | 4 | 0 | 0 | 0 | 0 |
| NS4A | Q9BVK6 | TMED9 | 1 | 6.5 | 0 | 6.5 | 3.82 | 3.76 | 1 | 0 | 6 | 6 | 6 | 8 | 0 | 0 | 0 | 0 |
| NS4A | Q5TF21 | SOGA3 | 1 | 6.75 | 0 | 6.75 | 3.85 | 3.77 | 1 | 0 | 9 | 4 | 8 | 6 | 0 | 0 | 0 | 0 |
| NS4A | A6NKG5 | RTL1 | 1 | 6.75 | 0 | 6.75 | 3.88 | 3.85 | 1 | 0 | 8 | 6 | 7 | 6 | 0 | 0 | 0 | 0 |
| NS4A | P07919 | UQCRH | 1 | 7 | 0 | 7 | 3.94 | 3.87 | 1 | 0 | 9 | 7 | 8 | 4 | 0 | 0 | 0 | 0 |
| NS4A | A0A0G2JJD3 | ABHD16A | 1 | 6.75 | 0 | 6.75 | 3.95 | 3.84 | 1 | 0 | 6 | 9 | 5 | 7 | 0 | 0 | 0 | 0 |
| NS4A | O95182 | NDUFA7 | 1 | 7 | 0 | 7 | 3.96 | 3.91 | 1 | 0 | 9 | 5 | 8 | 6 | 0 | 0 | 0 | 0 |
| NS4A | O94906-2 | PRPF6 | 1 | 21.5 | 0 | 21.5 | 3.98 | 3.31 | 1 | 0 | 21 | 21 | 26 | 18 | 0 | 2 | 1 | 2 |
| NS4A | P20020-1 | ATP2B1 | 1 | 7 | 0 | 7 | 4.02 | 3.88 | 1 | 0 | 11 | 5 | 6 | 6 | 0 | 0 | 0 | 0 |
| NS4A | Q15070-2 | OXA1L | 1 | 7.25 | 0 | 7.25 | 4.03 | 4.01 | 1 | 0 | 7 | 8 | 9 | 5 | 0 | 0 | 0 | 0 |
| NS4A | O14662-2 | STX16 | 1 | 7.25 | 0 | 7.25 | 4.1 | 4.07 | 1 | 0 | 7 | 6 | 8 | 8 | 0 | 0 | 0 | 0 |
| NS4A | Q8WVQ1-3 | CANT1 | 1 | 7.5 | 0 | 7.5 | 4.11 | 4.09 | 1 | 0 | 8 | 7 | 10 | 5 | 0 | 0 | 0 | 0 |
| NS4A | O60704 | TPST2 | 1 | 7.25 | 0 | 7.25 | 4.17 | 4.06 | 1 | 0 | 8 | 6 | 6 | 9 | 0 | 0 | 0 | 0 |
| NS4A | Q13190-4 | STX5 | 1 | 7.5 | 0 | 7.5 | 4.2 | 4.12 | 1 | 0 | 9 | 9 | 7 | 5 | 0 | 0 | 0 | 0 |
| NS4A | O60507 | TPST1 | 1 | 7.5 | 0 | 7.5 | 4.2 | 4.18 | 1 | 0 | 7 | 8 | 8 | 7 | 0 | 0 | 0 | 0 |
| NS4A | A0A590UJA9 | TMUB1 | 1 | 7.5 | 0 | 7.5 | 4.2 | 4.18 | 1 | 0 | 8 | 7 | 8 | 7 | 0 | 0 | 0 | 0 |
| NS4A | Q9UH62 | ARMCX3 | 1 | 7.75 | 0 | 7.75 | 4.29 | 4.26 | 1 | 0 | 6 | 8 | 9 | 8 | 0 | 0 | 0 | 0 |
| NS4A | O14828 | SCAMP3 | 1 | 7.75 | 0 | 7.75 | 4.34 | 4.27 | 1 | 0 | 7 | 10 | 7 | 7 | 0 | 0 | 0 | 0 |
| NS4A | Q5BJH7-3 | YIF1B | 1 | 7.75 | 0 | 7.75 | 4.35 | 4.29 | 1 | 0 | 7 | 9 | 7 | 8 | 0 | 0 | 0 | 0 |
| NS4A | O96011 | PEX11B | 1 | 8 | 0 | 8 | 4.36 | 4.36 | 1 | 0 | 8 | 7 | 10 | 7 | 0 | 0 | 0 | 0 |
| NS4A | Q9GZM5 | YIPF3 | 1 | 8 | 0 | 8 | 4.38 | 4.3 | 1 | 0 | 11 | 6 | 9 | 6 | 0 | 0 | 0 | 0 |
| NS4A | Q9UJS0-2 | SLC25A13 | 1 | 21.75 | 0 | 21.75 | 4.39 | 3.69 | 1 | 0 | 19 | 22 | 28 | 18 | 1 | 0 | 2 | 2 |
| NS4A | Q9Y262-2 | EIF3L | 1 | 8 | 0 | 8 | 4.4 | 4.36 | 1 | 0 | 6 | 9 | 9 | 8 | 0 | 0 | 0 | 0 |
| NS4A | P23634-2 | ATP2B4 | 1 | 8 | 0 | 8 | 4.41 | 4.24 | 1 | 0 | 11 | 4 | 9 | 8 | 0 | 0 | 0 | 0 |
| NS4A | O00461 | GOLIM4 | 1 | 8 | 0 | 8 | 4.43 | 4.36 | 1 | 0 | 10 | 6 | 8 | 8 | 0 | 0 | 0 | 0 |
| NS4A | Q8N699 | MYCT1 | 1 | 8.25 | 0 | 8.25 | 4.49 | 4.48 | 1 | 0 | 8 | 7 | 10 | 8 | 0 | 0 | 0 | 0 |
| NS4A | Q8N2K0-2 | ABHD12 | 1 | 8.25 | 0 | 8.25 | 4.5 | 4.49 | 1 | 0 | 9 | 8 | 9 | 7 | 0 | 0 | 0 | 0 |
| NS4A | Q9NP58-4 | ABCB6 | 1 | 8.5 | 0 | 8.5 | 4.61 | 4.58 | 1 | 0 | 8 | 10 | 9 | 7 | 0 | 0 | 0 | 0 |
| NS4A | Q96N66-2 | MBOAT7 | 1 | 8.5 | 0 | 8.5 | 4.67 | 4.6 | 1 | 0 | 10 | 7 | 8 | 9 | 0 | 0 | 0 | 0 |
| NS4A | P04844 | RPN2 | 1 | 8.75 | 0 | 8.75 | 4.71 | 4.65 | 1 | 0 | 7 | 7 | 11 | 10 | 0 | 0 | 0 | 0 |
| NS4A | B7ZAQ6-2 | GPR89A | 1 | 8.75 | 0 | 8.75 | 4.74 | 4.72 | 1 | 0 | 9 | 9 | 9 | 8 | 0 | 0 | 0 | 0 |
| NS4A | Q9NXF1-2 | TEX10 | 1 | 8.75 | 0 | 8.75 | 4.77 | 4.71 | 1 | 0 | 9 | 10 | 8 | 8 | 0 | 0 | 0 | 0 |
| NS4A | P03928 | MT-ATP8 | 1 | 9 | 0 | 9 | 4.82 | 4.8 | 1 | 0 | 10 | 8 | 10 | 8 | 0 | 0 | 0 | 0 |
| NS4A | O60427 | FADS1 | 1 | 9 | 0 | 9 | 4.82 | 4.8 | 1 | 0 | 10 | 8 | 10 | 8 | 0 | 0 | 0 | 0 |
| NS4A | Q6ZPD9 | DPY19L3 | 1 | 9.25 | 0 | 9.25 | 4.92 | 4.84 | 1 | 0 | 11 | 6 | 11 | 9 | 0 | 0 | 0 | 0 |
| NS4A | Q9BQG0-2 | MYBBP1A | 1 | 9.5 | 0 | 9.5 | 5 | 4.95 | 1 | 0 | 8 | 12 | 11 | 7 | 0 | 0 | 0 | 0 |
| NS4A | Q15021 | NCAPD2 | 1 | 9.25 | 0 | 9.25 | 5.02 | 4.89 | 1 | 0 | 11 | 11 | 7 | 8 | 0 | 0 | 0 | 0 |
| NS4A | P11166 | SLC2A1 | 1 | 9.5 | 0 | 9.5 | 5.02 | 4.87 | 1 | 0 | 14 | 8 | 10 | 6 | 0 | 0 | 0 | 0 |
| NS4A | P46977 | STT3A | 1 | 9.5 | 0 | 9.5 | 5.03 | 4.93 | 1 | 0 | 9 | 6 | 12 | 11 | 0 | 0 | 0 | 0 |
| NS4A | Q9BV81 | EMC6 | 1 | 9.5 | 0 | 9.5 | 5.08 | 5.03 | 1 | 0 | 11 | 9 | 9 | 9 | 0 | 0 | 0 | 0 |
| NS4A | P18084 | ITGB5 | 1 | 9.5 | 0 | 9.5 | 5.11 | 5.01 | 1 | 0 | 9 | 12 | 8 | 9 | 0 | 0 | 0 | 0 |
| NS4A | Q9H8M9 | EVA1A | 1 | 9.5 | 0 | 9.5 | 5.13 | 5.04 | 1 | 0 | 10 | 10 | 8 | 10 | 0 | 0 | 0 | 0 |
| NS4A | Q9UM00-1 | TMCO1 | 1 | 9.75 | 0 | 9.75 | 5.15 | 5.13 | 1 | 0 | 9 | 9 | 11 | 10 | 0 | 0 | 0 | 0 |
| NS4A | O00116 | AGPS | 1 | 10 | 0 | 10 | 5.18 | 5.13 | 1 | 0 | 7 | 9 | 14 | 10 | 0 | 0 | 0 | 0 |
| NS4A | A0A0A0MR51 | FADS1 | 1 | 10 | 0 | 10 | 5.28 | 5.24 | 1 | 0 | 11 | 10 | 10 | 9 | 0 | 0 | 0 | 0 |
| NS4A | P08240-2 | SRPRA | 1 | 12.25 | 0 | 12.25 | 5.28 | 5 | 1 | 0 | 11 | 14 | 13 | 11 | 1 | 0 | 0 | 0 |
| NS4A | Q9Y679 | AUP1 | 1 | 10.5 | 0 | 10.5 | 5.33 | 5.3 | 1 | 0 | 10 | 10 | 15 | 7 | 0 | 0 | 0 | 0 |
| NS4A | P08574 | CYC1 | 1 | 10.25 | 0 | 10.25 | 5.33 | 5.26 | 1 | 0 | 10 | 7 | 13 | 11 | 0 | 0 | 0 | 0 |
| NS4A | O14681 | EI24 | 1 | 10.25 | 0 | 10.25 | 5.39 | 5.35 | 1 | 0 | 10 | 9 | 11 | 11 | 0 | 0 | 0 | 0 |
| NS4A | Q9BU23-2 | LMF2 | 1 | 10.5 | 0 | 10.5 | 5.44 | 5.37 | 1 | 0 | 13 | 11 | 11 | 7 | 0 | 0 | 0 | 0 |
| NS4A | Q5T2N8 | ATAD3C | 1 | 10.75 | 0 | 10.75 | 5.49 | 5.47 | 1 | 0 | 11 | 10 | 14 | 8 | 0 | 0 | 0 | 0 |
| NS4A | O43292 | GPAA1 | 1 | 10.5 | 0 | 10.5 | 5.49 | 5.44 | 1 | 0 | 12 | 9 | 11 | 10 | 0 | 0 | 0 | 0 |
| NS4A | P28288 | ABCD3 | 1 | 11 | 0 | 11 | 5.54 | 5.51 | 1 | 0 | 9 | 11 | 16 | 8 | 0 | 0 | 0 | 0 |
| NS4A | Q12981 | BNIP1 | 1 | 10.75 | 0 | 10.75 | 5.58 | 5.5 | 1 | 0 | 14 | 9 | 11 | 9 | 0 | 0 | 0 | 0 |
| NS4A | O75063 | FAM20B | 1 | 11 | 0 | 11 | 5.65 | 5.64 | 1 | 0 | 11 | 10 | 13 | 10 | 0 | 0 | 0 | 0 |
| NS4A | P10606 | COX5B | 1 | 10.75 | 0 | 10.75 | 5.67 | 5.56 | 1 | 0 | 11 | 12 | 9 | 11 | 0 | 0 | 0 | 0 |
| NS4A | Q9Y2R0 | COA3 | 1 | 11 | 0 | 11 | 5.71 | 5.66 | 1 | 0 | 10 | 10 | 12 | 12 | 0 | 0 | 0 | 0 |
| NS4A | P49755 | TMED10 | 1 | 11.25 | 0 | 11.25 | 5.73 | 5.73 | 1 | 0 | 11 | 10 | 14 | 10 | 0 | 0 | 0 | 0 |
| NS4A | P42167 | TMPO | 1 | 25.25 | 0 | 25.25 | 5.92 | 5.03 | 1 | 0 | 24 | 25 | 26 | 26 | 2 | 2 | 0 | 0 |
| NS4A | O75396 | SEC22B | 1 | 14.25 | 0 | 14.25 | 5.94 | 5.65 | 1 | 0 | 15 | 14 | 17 | 11 | 1 | 0 | 0 | 0 |
| NS4A | Q9UEU0 | VTI1B | 1 | 11.75 | 0 | 11.75 | 6.01 | 5.96 | 1 | 0 | 14 | 11 | 12 | 10 | 0 | 0 | 0 | 0 |
| NS4A | Q9NVI7-2 | ATAD3A | 1 | 48.25 | 0 | 48.25 | 6.06 | 5.24 | 1 | 0 | 42 | 57 | 56 | 38 | 2 | 5 | 1 | 1 |
| NS4A | Q96BQ5 | CCDC127 | 1 | 12 | 0 | 12 | 6.15 | 6.07 | 1 | 0 | 13 | 14 | 11 | 10 | 0 | 0 | 0 | 0 |
| NS4A | Q6UWP7-3 | LCLAT1 | 1 | 12 | 0 | 12 | 6.17 | 6.05 | 1 | 0 | 14 | 9 | 12 | 13 | 0 | 0 | 0 | 0 |
| NS4A | P05023-4 | ATP1A1 | 1 | 29 | 0 | 29 | 6.28 | 5.29 | 1 | 0 | 30 | 28 | 30 | 28 | 1 | 1 | 0 | 2 |
| NS4A | P09669 | COX6C | 1 | 15.25 | 0 | 15.25 | 6.31 | 5.98 | 1 | 0 | 17 | 14 | 18 | 12 | 1 | 0 | 0 | 0 |
| NS4A | Q9HCU5 | PREB | 1 | 13 | 0 | 13 | 6.41 | 6.36 | 1 | 0 | 11 | 15 | 17 | 9 | 0 | 0 | 0 | 0 |
| NS4A | A0A669KB27 | DUS4L-BCAP29 | 1 | 12.75 | 0 | 12.75 | 6.45 | 6.42 | 1 | 0 | 13 | 13 | 13 | 12 | 0 | 0 | 0 | 0 |
| NS4A | H0YJH7 | TMPO | 1 | 20.75 | 0 | 20.75 | 6.58 | 5.89 | 1 | 0 | 20 | 21 | 22 | 20 | 1 | 1 | 0 | 0 |
| NS4A | Q07065 | CKAP4 | 1 | 42.75 | 0 | 42.75 | 6.58 | 5.38 | 1 | 0 | 39 | 47 | 53 | 32 | 7 | 0 | 0 | 2 |
| NS4A | O95470 | SGPL1 | 1 | 13 | 0 | 13 | 6.58 | 6.52 | 1 | 0 | 14 | 12 | 13 | 13 | 0 | 0 | 0 | 0 |
| NS4A | Q9HBL7 | PLGRKT | 1 | 13.25 | 0 | 13.25 | 6.67 | 6.57 | 1 | 0 | 11 | 17 | 13 | 12 | 0 | 0 | 0 | 0 |
| NS4A | Q8TB61-2 | SLC35B2 | 1 | 13.25 | 0 | 13.25 | 6.69 | 6.63 | 1 | 0 | 14 | 13 | 13 | 13 | 0 | 0 | 0 | 0 |
| NS4A | Q5T8D3-2 | ACBD5 | 1 | 13.5 | 0 | 13.5 | 6.8 | 6.68 | 1 | 0 | 17 | 14 | 12 | 11 | 0 | 0 | 0 | 0 |
| NS4A | Q8IXI1 | RHOT2 | 1 | 13.75 | 0 | 13.75 | 6.84 | 6.82 | 1 | 0 | 13 | 15 | 15 | 12 | 0 | 0 | 0 | 0 |
| NS4A | Q96F07-2 | CYFIP2 | 1 | 14.5 | 0 | 14.5 | 7.15 | 7.12 | 1 | 0 | 14 | 16 | 16 | 12 | 0 | 0 | 0 | 0 |
| NS4A | Q9BUN8-2 | DERL1 | 1 | 14.75 | 0 | 14.75 | 7.37 | 7.25 | 1 | 0 | 15 | 17 | 13 | 14 | 0 | 0 | 0 | 0 |
| NS4A | Q9HD20-2 | ATP13A1 | 1 | 14.75 | 0 | 14.75 | 7.37 | 7.27 | 1 | 0 | 14 | 15 | 14 | 16 | 0 | 0 | 0 | 0 |
| NS4A | Q9Y4W6 | AFG3L2 | 1 | 15 | 0 | 15 | 7.38 | 7.35 | 1 | 0 | 16 | 15 | 16 | 13 | 0 | 0 | 0 | 0 |
| NS4A | P20674 | COX5A | 1 | 15.25 | 0 | 15.25 | 7.4 | 7.38 | 1 | 0 | 16 | 14 | 19 | 12 | 0 | 0 | 0 | 0 |
| NS4A | Q10471 | GALNT2 | 1 | 15.75 | 0 | 15.75 | 7.62 | 7.55 | 1 | 0 | 19 | 13 | 19 | 12 | 0 | 0 | 0 | 0 |
| NS4A | C9JRZ6 | CHCHD3 | 1 | 22 | 0 | 22 | 7.78 | 7.02 | 1 | 0 | 22 | 24 | 19 | 23 | 2 | 0 | 0 | 0 |
| NS4A | Q9BTX1-2 | NDC1 | 1 | 15.75 | 0 | 15.75 | 7.88 | 7.66 | 1 | 0 | 15 | 16 | 13 | 19 | 0 | 0 | 0 | 0 |
| NS4A | O43242 | PSMD3 | 1 | 28.75 | 0 | 28.75 | 8.03 | 7.09 | 1 | 0 | 28 | 29 | 33 | 25 | 0 | 1 | 0 | 1 |
| NS4A | Q6ZXV5-2 | TMTC3 | 1 | 16.75 | 0 | 16.75 | 8.13 | 8.1 | 1 | 0 | 17 | 17 | 18 | 15 | 0 | 0 | 0 | 0 |
| NS4A | Q56VL3 | OCIAD2 | 1 | 17.5 | 0 | 17.5 | 8.34 | 8.31 | 1 | 0 | 17 | 14 | 23 | 16 | 0 | 0 | 0 | 0 |
| NS4A | O95864 | FADS2 | 1 | 17.5 | 0 | 17.5 | 8.38 | 8.37 | 1 | 0 | 17 | 17 | 21 | 15 | 0 | 0 | 0 | 0 |
| NS4A | P35610 | SOAT1 | 1 | 18.5 | 0 | 18.5 | 8.73 | 8.51 | 1 | 0 | 25 | 14 | 23 | 12 | 0 | 0 | 0 | 0 |
| NS4A | Q8WVM8 | SCFD1 | 1 | 19 | 0 | 19 | 9.09 | 9.02 | 1 | 0 | 22 | 18 | 20 | 16 | 0 | 0 | 0 | 0 |
| NS4A | Q9NZ01 | TECR | 1 | 23.25 | 0 | 23.25 | 9.21 | 8.72 | 1 | 0 | 27 | 21 | 26 | 19 | 1 | 0 | 0 | 0 |
| NS4A | Q5JTH9-2 | RRP12 | 1 | 23 | 0 | 23 | 9.25 | 8.64 | 1 | 0 | 22 | 29 | 21 | 20 | 1 | 0 | 0 | 0 |
| NS4A | Q6P2E9-2 | EDC4 | 1 | 20.25 | 0 | 20.25 | 9.53 | 9.5 | 1 | 0 | 22 | 19 | 24 | 16 | 0 | 0 | 0 | 0 |
| NS4A | Q9BVC6 | TMEM109 | 1 | 20 | 0 | 20 | 9.61 | 9.5 | 1 | 0 | 21 | 20 | 19 | 20 | 0 | 0 | 0 | 0 |
| NS4A | Q7L576 | CYFIP1 | 1 | 20.5 | 0 | 20.5 | 9.72 | 9.68 | 1 | 0 | 21 | 21 | 22 | 18 | 0 | 0 | 0 | 0 |
| NS4A | P08183-2 | ABCB1 | 1 | 20.75 | 0 | 20.75 | 9.75 | 9.7 | 1 | 0 | 23 | 17 | 25 | 18 | 0 | 0 | 0 | 0 |
| NS4A | Q8NBM4 | UBAC2 | 1 | 22.25 | 0 | 22.25 | 10.47 | 10.42 | 1 | 0 | 21 | 24 | 24 | 20 | 0 | 0 | 0 | 0 |
| NS4A | E9PCW1 | GOSR1 | 1 | 22.25 | 0 | 22.25 | 10.58 | 10.45 | 1 | 0 | 22 | 21 | 22 | 24 | 0 | 0 | 0 | 0 |
| NS4A | Q86VP6 | CAND1 | 1 | 22.5 | 0 | 22.5 | 10.58 | 10.53 | 1 | 0 | 22 | 24 | 24 | 20 | 0 | 0 | 0 | 0 |
| NS4A | Q5HYI8 | RABL3 | 1 | 22.5 | 0 | 22.5 | 10.62 | 10.55 | 1 | 0 | 24 | 22 | 23 | 21 | 0 | 0 | 0 | 0 |
| NS4A | O95249 | GOSR1 | 1 | 23 | 0 | 23 | 10.9 | 10.76 | 1 | 0 | 22 | 22 | 23 | 25 | 0 | 0 | 0 | 0 |
| NS4A | Q9NVH1-3 | DNAJC11 | 1 | 23.5 | 0 | 23.5 | 10.94 | 10.87 | 1 | 0 | 26 | 24 | 26 | 18 | 0 | 0 | 0 | 0 |
| NS4A | Q9Y277-2 | VDAC3 | 1 | 23.5 | 0 | 23.5 | 11.02 | 10.96 | 1 | 0 | 25 | 22 | 25 | 22 | 0 | 0 | 0 | 0 |
| NS4A | Q9P0L0 | VAPA | 1 | 25.25 | 0 | 25.25 | 11.74 | 11.7 | 1 | 0 | 25 | 24 | 28 | 24 | 0 | 0 | 0 | 0 |
| NS4A | Q99623 | PHB2 | 1 | 146 | 0 | 146 | 11.81 | 10.71 | 1 | 0 | 151 | 171 | 146 | 116 | 6 | 4 | 3 | 4 |
| NS4A | E9PIE4 | MTCH2 | 1 | 25.25 | 0 | 25.25 | 11.87 | 11.67 | 1 | 0 | 30 | 25 | 23 | 23 | 0 | 0 | 0 | 0 |
| NS4A | P16615 | ATP2A2 | 1 | 51.75 | 0 | 51.75 | 12.09 | 10.34 | 1 | 0 | 51 | 54 | 52 | 50 | 2 | 0 | 1 | 1 |
| NS4A | Q14739 | LBR | 1 | 26 | 0 | 26 | 12.1 | 12.02 | 1 | 0 | 28 | 25 | 27 | 24 | 0 | 0 | 0 | 0 |
| NS4A | O94766 | B3GAT3 | 1 | 27.25 | 0 | 27.25 | 12.64 | 12.5 | 1 | 0 | 27 | 32 | 27 | 23 | 0 | 0 | 0 | 0 |
| NS4A | O95292 | VAPB | 1 | 27.75 | 0 | 27.75 | 13.02 | 12.77 | 1 | 0 | 29 | 27 | 25 | 30 | 0 | 0 | 0 | 0 |
| NS4A | Q14204 | DYNC1H1 | 1 | 30.25 | 0 | 30.25 | 13.98 | 13.73 | 1 | 0 | 29 | 38 | 28 | 26 | 0 | 0 | 0 | 0 |
| NS4A | P78527 | PRKDC | 1 | 80 | 0 | 80 | 14.07 | 11.66 | 1 | 0 | 77 | 89 | 85 | 69 | 3 | 1 | 0 | 2 |
| NS4A | Q9Y6C9 | MTCH2 | 1 | 30.75 | 0 | 30.75 | 14.22 | 13.99 | 1 | 0 | 36 | 32 | 28 | 27 | 0 | 0 | 0 | 0 |
| NS4A | Q8NC56 | LEMD2 | 1 | 33.5 | 0 | 33.5 | 15.37 | 15.18 | 1 | 0 | 29 | 37 | 34 | 34 | 0 | 0 | 0 | 0 |
| NS4A | P51572-2 | BCAP31 | 1 | 37.5 | 0 | 37.5 | 16.78 | 16.73 | 1 | 0 | 40 | 32 | 46 | 32 | 0 | 0 | 0 | 0 |
| NS4A | Q9C0E8-2 | LNP | 1 | 38 | 0 | 38 | 17.28 | 16.97 | 1 | 0 | 36 | 30 | 42 | 44 | 0 | 0 | 0 | 0 |
| NS4A | P35232 | PHB | 1 | 163.5 | 0 | 163.5 | 18.85 | 16.68 | 1 | 0 | 171 | 177 | 164 | 142 | 2 | 2 | 2 | 4 |
| NS4A | P21796 | VDAC1 | 1 | 43 | 0 | 43 | 19.15 | 19.11 | 1 | 0 | 42 | 45 | 50 | 35 | 0 | 0 | 0 | 0 |
| NS4A | Q92616 | GCN1 | 1 | 42 | 0 | 42 | 19.24 | 18.78 | 1 | 0 | 41 | 45 | 36 | 46 | 0 | 0 | 0 | 0 |
| NS4A | P45880 | VDAC2 | 1 | 44.25 | 0 | 44.25 | 19.78 | 19.73 | 1 | 0 | 44 | 45 | 49 | 39 | 0 | 0 | 0 | 0 |
| NS4A | A0A1B0GW05 | DPY19L1 | 1 | 50.75 | 0 | 50.75 | 22.64 | 22.4 | 1 | 0 | 60 | 45 | 53 | 45 | 0 | 0 | 0 | 0 |
| NS4A | Q53H12 | AGK | 1 | 51.75 | 0 | 51.75 | 23.04 | 22.85 | 1 | 0 | 54 | 43 | 58 | 52 | 0 | 0 | 0 | 0 |
| NS4A | Q16891-2 | IMMT | 1 | 93.5 | 0 | 93.5 | 41.23 | 40.34 | 1 | 0 | 98 | 113 | 83 | 80 | 0 | 0 | 0 | 0 |
| NS4A | Q9BSJ8-2 | ESYT1 | 1 | 300.75 | 0 | 300.75 | 129.66 | 128.49 | 1 | 0 | 323 | 281 | 310 | 289 | 0 | 0 | 0 | 0 |

| BAIT | PROTID | GENE | preyFreq | preyAvg_AvgCounts | preySD_AvgCounts | NS4B_AVG_SPC | NS4B_FC_A | NS4B_FC_B | NS4B_SP | NS4B_iREF | IP_SH-SY5Y_NS4B_R4 | IP_SH-SY5Y_NS4B_R3 | IP_SH-SY5Y_NS4B_R2 | IP_SH-SY5Y_NS4B_R1 | IP_SH-SY5Y_GFP_R4 | IP_SH-SY5Y_GFP_R3 | IP_SH-SY5Y_GFP_R2 | IP_SH-SY5Y_GFP_R1 |
|---|---|---|---|---|---|---|---|---|---|---|---|---|---|---|---|---|---|---|
| NS4B | P27544-2 | CERS1 | 1 | 4 | 0 | 4 | 2.77 | 2.76 | 1 | 0 | 5 | 4 | 4 | 3 | 0 | 0 | 0 | 0 |
| NS4B | Q96K37 | SLC35E1 | 1 | 4.25 | 0 | 4.25 | 2.9 | 2.89 | 1 | 0 | 4 | 4 | 5 | 4 | 0 | 0 | 0 | 0 |
| NS4B | Q969P0-3 | IGSF8 | 1 | 4.75 | 0 | 4.75 | 3.11 | 3.09 | 1 | 0 | 6 | 5 | 4 | 4 | 0 | 0 | 0 | 0 |
| NS4B | Q9BV81 | EMC6 | 1 | 4.75 | 0 | 4.75 | 3.12 | 3.11 | 1 | 0 | 5 | 5 | 5 | 4 | 0 | 0 | 0 | 0 |
| NS4B | P21397-2 | MAOA | 1 | 5 | 0 | 5 | 3.18 | 3.17 | 1 | 0 | 7 | 4 | 5 | 4 | 0 | 0 | 0 | 0 |
| NS4B | Q96GQ5 | C16ORF58 | 1 | 5 | 0 | 5 | 3.18 | 3.12 | 1 | 0 | 8 | 4 | 3 | 5 | 0 | 0 | 0 | 0 |
| NS4B | Q9P0T7 | TMEM9 | 1 | 5.25 | 0 | 5.25 | 3.35 | 3.32 | 1 | 0 | 6 | 5 | 4 | 6 | 0 | 0 | 0 | 0 |
| NS4B | Q32P28 | P3H1 | 1 | 5.25 | 0 | 5.25 | 3.39 | 3.33 | 1 | 0 | 5 | 6 | 4 | 6 | 0 | 0 | 0 | 0 |
| NS4B | P24534 | EEF1B2 | 1 | 5.75 | 0 | 5.75 | 3.53 | 3.51 | 1 | 0 | 6 | 4 | 7 | 6 | 0 | 0 | 0 | 0 |
| NS4B | Q9Y4W6 | AFG3L2 | 1 | 5.75 | 0 | 5.75 | 3.56 | 3.52 | 1 | 0 | 6 | 6 | 7 | 4 | 0 | 0 | 0 | 0 |
| NS4B | P98194-2 | ATP2C1 | 1 | 6 | 0 | 6 | 3.65 | 3.65 | 1 | 0 | 7 | 5 | 6 | 6 | 0 | 0 | 0 | 0 |
| NS4B | O00165-2 | HAX1 | 1 | 6 | 0 | 6 | 3.66 | 3.65 | 1 | 0 | 6 | 5 | 7 | 6 | 0 | 0 | 0 | 0 |
| NS4B | P42704 | LRPPRC | 1 | 6.25 | 0 | 6.25 | 3.67 | 3.61 | 1 | 0 | 9 | 4 | 8 | 4 | 0 | 0 | 0 | 0 |
| NS4B | Q96JJ7-2 | TMX3 | 1 | 6.25 | 0 | 6.25 | 3.69 | 3.64 | 1 | 0 | 10 | 5 | 6 | 4 | 0 | 0 | 0 | 0 |
| NS4B | Q9P0J0-2 | NDUFA13 | 1 | 6.25 | 0 | 6.25 | 3.72 | 3.7 | 1 | 0 | 9 | 5 | 6 | 5 | 0 | 0 | 0 | 0 |
| NS4B | Q9UM00-1 | TMCO1 | 1 | 6.25 | 0 | 6.25 | 3.76 | 3.71 | 1 | 0 | 7 | 4 | 6 | 8 | 0 | 0 | 0 | 0 |
| NS4B | Q9NRK6 | ABCB10 | 1 | 6.25 | 0 | 6.25 | 3.76 | 3.75 | 1 | 0 | 7 | 5 | 7 | 6 | 0 | 0 | 0 | 0 |
| NS4B | Q93045 | STMN2 | 1 | 6.25 | 0 | 6.25 | 3.78 | 3.76 | 1 | 0 | 6 | 5 | 7 | 7 | 0 | 0 | 0 | 0 |
| NS4B | Q9Y679 | AUP1 | 1 | 7.25 | 0 | 7.25 | 4.1 | 4.04 | 1 | 0 | 11 | 4 | 8 | 6 | 0 | 0 | 0 | 0 |
| NS4B | P03928 | MT-ATP8 | 1 | 7 | 0 | 7 | 4.11 | 4.1 | 1 | 0 | 7 | 6 | 8 | 7 | 0 | 0 | 0 | 0 |
| NS4B | Q14318-2 | FKBP8 | 1 | 7.25 | 0 | 7.25 | 4.2 | 4.2 | 1 | 0 | 8 | 6 | 8 | 7 | 0 | 0 | 0 | 0 |
| NS4B | A0A0A0MR51 | FADS1 | 1 | 7.75 | 0 | 7.75 | 4.46 | 4.44 | 1 | 0 | 8 | 7 | 8 | 8 | 0 | 0 | 0 | 0 |
| NS4B | P08183-2 | ABCB1 | 1 | 7.75 | 0 | 7.75 | 4.46 | 4.44 | 1 | 0 | 8 | 7 | 8 | 8 | 0 | 0 | 0 | 0 |
| NS4B | P30837 | ALDH1B1 | 1 | 11.5 | 0 | 11.5 | 4.47 | 4.01 | 1 | 0 | 20 | 8 | 11 | 7 | 0 | 0 | 0 | 1 |
| NS4B | O14949 | UQCRQ | 1 | 8 | 0 | 8 | 4.51 | 4.45 | 1 | 0 | 11 | 6 | 6 | 9 | 0 | 0 | 0 | 0 |
| NS4B | Q3ZCQ8-2 | TIMM50 | 1 | 13.75 | 0 | 13.75 | 4.52 | 4.03 | 1 | 0 | 16 | 11 | 14 | 14 | 0 | 0 | 1 | 1 |
| NS4B | O14828 | SCAMP3 | 1 | 8.25 | 0 | 8.25 | 4.64 | 4.59 | 1 | 0 | 10 | 6 | 7 | 10 | 0 | 0 | 0 | 0 |

| | | | | | | | | | | | | | | | | | | |
|---|---|---|---|---|---|---|---|---|---|---|---|---|---|---|---|---|---|---|
| NS4B | Q6ZXV5-2 | TMTC3 | 1 | 8.5 | 0 | 8.5 | 4.73 | 4.72 | 1 | 0 | 10 | 6 | 9 | 9 | 0 | 0 | 0 | 0 |
| NS4B | O60427 | FADS1 | 1 | 8.75 | 0 | 8.75 | 4.92 | 4.89 | 1 | 0 | 9 | 9 | 9 | 8 | 0 | 0 | 0 | 0 |
| NS4B | Q15070-2 | OXA1L | 1 | 9 | 0 | 9 | 4.95 | 4.9 | 1 | 0 | 13 | 8 | 7 | 8 | 0 | 0 | 0 | 0 |
| NS4B | O75746-2 | SLC25A12 | 1 | 9 | 0 | 9 | 5 | 4.95 | 1 | 0 | 11 | 8 | 7 | 10 | 0 | 0 | 0 | 0 |
| NS4B | Q9UNL2-2 | SSR3 | 1 | 9.5 | 0 | 9.5 | 5.15 | 5.1 | 1 | 0 | 14 | 8 | 8 | 8 | 0 | 0 | 0 | 0 |
| NS4B | O00116 | AGPS | 1 | 10.75 | 0 | 10.75 | 5.51 | 5.25 | 1 | 0 | 20 | 6 | 11 | 6 | 0 | 0 | 0 | 0 |
| NS4B | Q6IAA8 | LAMTOR1 | 1 | 10.25 | 0 | 10.25 | 5.59 | 5.54 | 1 | 0 | 10 | 9 | 10 | 12 | 0 | 0 | 0 | 0 |
| NS4B | Q8IXI1 | RHOT2 | 1 | 11 | 0 | 11 | 5.82 | 5.77 | 1 | 0 | 15 | 10 | 11 | 8 | 0 | 0 | 0 | 0 |
| NS4B | P09172 | DBH | 1 | 11 | 0 | 11 | 5.83 | 5.79 | 1 | 0 | 14 | 10 | 12 | 8 | 0 | 0 | 0 | 0 |
| NS4B | Q9BUN8-2 | DERL1 | 1 | 10.75 | 0 | 10.75 | 5.87 | 5.74 | 1 | 0 | 9 | 10 | 10 | 14 | 0 | 0 | 0 | 0 |
| NS4B | P13637-2 | ATP1A3 | 1 | 26.75 | 0 | 26.75 | 6.02 | 5.09 | 1 | 0 | 31 | 26 | 28 | 22 | 1 | 1 | 0 | 2 |
| NS4B | Q9Y277-2 | VDAC3 | 1 | 11.5 | 0 | 11.5 | 6.14 | 6.05 | 1 | 0 | 13 | 10 | 9 | 14 | 0 | 0 | 0 | 0 |
| NS4B | Q9H4A4 | RNPEP | 1 | 27.75 | 0 | 27.75 | 6.2 | 5.23 | 1 | 0 | 29 | 24 | 35 | 23 | 1 | 2 | 0 | 1 |
| NS4B | Q9HD20-2 | ATP13A1 | 1 | 12.25 | 0 | 12.25 | 6.38 | 6.32 | 1 | 0 | 17 | 10 | 10 | 12 | 0 | 0 | 0 | 0 |
| NS4B | Q16891-2 | IMMT | 1 | 12.5 | 0 | 12.5 | 6.47 | 6.38 | 1 | 0 | 15 | 11 | 16 | 8 | 0 | 0 | 0 | 0 |
| NS4B | P45880 | VDAC2 | 1 | 14.25 | 0 | 14.25 | 7.39 | 7.29 | 1 | 0 | 15 | 13 | 12 | 17 | 0 | 0 | 0 | 0 |
| NS4B | Q14739 | LBR | 1 | 14.75 | 0 | 14.75 | 7.48 | 7.44 | 1 | 0 | 16 | 12 | 19 | 12 | 0 | 0 | 0 | 0 |
| NS4B | Q9UJS0-2 | SLC25A13 | 1 | 38 | 0 | 38 | 7.8 | 6.53 | 1 | 0 | 42 | 37 | 36 | 37 | 1 | 0 | 2 | 2 |
| NS4B | P11166 | SLC2A1 | 1 | 15.5 | 0 | 15.5 | 7.93 | 7.88 | 1 | 0 | 17 | 16 | 15 | 14 | 0 | 0 | 0 | 0 |
| NS4B | Q8NBM4 | UBAC2 | 1 | 16.25 | 0 | 16.25 | 8.11 | 8.06 | 1 | 0 | 21 | 14 | 18 | 12 | 0 | 0 | 0 | 0 |
| NS4B | P05023-4 | ATP1A1 | 1 | 39.5 | 0 | 39.5 | 8.68 | 7.36 | 1 | 0 | 42 | 34 | 42 | 40 | 1 | 1 | 0 | 2 |
| NS4B | O95292 | VAPB | 1 | 18 | 0 | 18 | 9.03 | 8.99 | 1 | 0 | 18 | 17 | 20 | 17 | 0 | 0 | 0 | 0 |
| NS4B | O95864 | FADS2 | 1 | 21 | 0 | 21 | 10.4 | 10.33 | 1 | 0 | 20 | 20 | 23 | 21 | 0 | 0 | 0 | 0 |
| NS4B | P16615 | ATP2A2 | 1 | 52 | 0 | 52 | 12.49 | 10.74 | 1 | 0 | 57 | 40 | 58 | 53 | 2 | 0 | 1 | 1 |
| NS4B | Q9NZ01 | TECR | 1 | 31.75 | 0 | 31.75 | 12.7 | 12.01 | 1 | 0 | 34 | 22 | 38 | 33 | 1 | 0 | 0 | 0 |
| NS4B | P21796 | VDAC1 | 1 | 36.5 | 0 | 36.5 | 17.09 | 16.84 | 1 | 0 | 49 | 30 | 28 | 39 | 0 | 0 | 0 | 0 |

| BAIT | PROTID | GENE | preyFreq | preyAvg AvgCounts | preySD AvgCounts | NS5 AVG SPC | NS5 FC A | NS5 FC B | NS5 SP | NS5 iREF | IP SH-SY5Y NS5 R4 | IP SH-SY5Y NS5 R3 | IP SH-SY5Y NS5 R2 | IP SH-SY5Y NS5 R1 | IP SH-SY5Y GFP R4 | IP SH-SY5Y GFP R3 | IP SH-SY5Y GFP R2 | IP SH-SY5Y GFP R1 |
|---|---|---|---|---|---|---|---|---|---|---|---|---|---|---|---|---|---|---|
| NS5 | Q9NX63 | CHCHD3 | 1 | 6.5 | 0 | 6.5 | 3.25 | 3.23 | 1 | 0 | 5 | 6 | 9 | 6 | 0 | 0 | 0 | 0 |
| NS5 | P15090 | FABP4 | 1 | 8.5 | 0 | 8.5 | 3.79 | 3.61 | 1 | 0 | 3 | 6 | 13 | 12 | 0 | 0 | 0 | 0 |
| NS5 | P48788-2 | TNNI2 | 1 | 8.25 | 0 | 8.25 | 3.79 | 3.73 | 1 | 0 | 5 | 7 | 13 | 8 | 0 | 0 | 0 | 0 |
| NS5 | Q9UNX3 | RPL26L1 | 1 | 9.25 | 0 | 9.25 | 4.03 | 3.86 | 1 | 0 | 3 | 15 | 10 | 9 | 0 | 0 | 0 | 0 |
| NS5 | P07205 | PGK2 | 1 | 9 | 0 | 9 | 4.12 | 4.05 | 1 | 0 | 7 | 6 | 11 | 12 | 0 | 0 | 0 | 0 |
| NS5 | Q6NZI2 | CAVIN1 | 1 | 11 | 0 | 11 | 4.61 | 4.34 | 1 | 0 | 4 | 7 | 18 | 15 | 0 | 0 | 0 | 0 |
| NS5 | P09104 | ENO2 | 1 | 11.5 | 0 | 11.5 | 4.84 | 4.69 | 1 | 0 | 6 | 10 | 20 | 10 | 0 | 0 | 0 | 0 |
| NS5 | P08590 | MYL3 | 1 | 12.25 | 0 | 12.25 | 5.02 | 4.58 | 1 | 0 | 5 | 6 | 25 | 13 | 0 | 0 | 0 | 0 |
| NS5 | Q16891-2 | IMMT | 1 | 12.75 | 0 | 12.75 | 5.47 | 5.42 | 1 | 0 | 11 | 15 | 14 | 11 | 0 | 0 | 0 | 0 |
| NS5 | P09429 | HMGB1 | 1 | 14.25 | 0 | 14.25 | 5.83 | 5.45 | 1 | 0 | 9 | 6 | 23 | 19 | 0 | 0 | 0 | 0 |
| NS5 | F8WA37 | TNNT3 | 1 | 15.25 | 0 | 15.25 | 6.12 | 5.89 | 1 | 0 | 8 | 10 | 20 | 23 | 0 | 0 | 0 | 0 |
| NS5 | P45378-7 | TNNT3 | 1 | 15.25 | 0 | 15.25 | 6.12 | 5.89 | 1 | 0 | 8 | 10 | 21 | 22 | 0 | 0 | 0 | 0 |
| NS5 | Q03252 | LMNB2 | 1 | 15.25 | 0 | 15.25 | 6.4 | 5.66 | 1 | 0 | 14 | 31 | 8 | 8 | 0 | 0 | 0 | 0 |
| NS5 | P84243 | H3F3A;H3F3B | 1 | 16.75 | 0 | 16.75 | 6.5 | 6.2 | 1 | 0 | 6 | 14 | 24 | 23 | 0 | 0 | 0 | 0 |
| NS5 | A0A2R8YEA7 | ACTB | 1 | 15.75 | 0 | 15.75 | 6.66 | 6.51 | 1 | 0 | 17 | 14 | 17 | 15 | 0 | 0 | 0 | 0 |
| NS5 | Q13813 | SPTAN1 | 1 | 18.25 | 0 | 18.25 | 6.87 | 5.97 | 1 | 0 | 4 | 10 | 31 | 28 | 0 | 0 | 0 | 0 |
| NS5 | P05976 | MYL1 | 1 | 18 | 0 | 18 | 6.99 | 6.62 | 1 | 0 | 9 | 12 | 33 | 18 | 0 | 0 | 0 | 0 |
| NS5 | P00558-2 | PGK1 | 1 | 18.25 | 0 | 18.25 | 7.04 | 6.4 | 1 | 0 | 8 | 8 | 31 | 26 | 0 | 0 | 0 | 0 |
| NS5 | O14983-2 | ATP2A1 | 1 | 18 | 0 | 18 | 7.15 | 7.13 | 1 | 0 | 12 | 17 | 22 | 21 | 0 | 0 | 0 | 0 |
| NS5 | A0A7I2V5A1 | ACO2 | 1 | 19 | 0 | 19 | 7.23 | 6.81 | 1 | 0 | 7 | 15 | 33 | 21 | 0 | 0 | 0 | 0 |
| NS5 | P00558 | PGK1 | 1 | 22 | 0 | 22 | 8.21 | 7.22 | 1 | 0 | 8 | 9 | 39 | 32 | 0 | 0 | 0 | 0 |
| NS5 | P13929 | ENO3 | 1 | 22.5 | 0 | 22.5 | 8.5 | 7.99 | 1 | 0 | 11 | 13 | 37 | 29 | 0 | 0 | 0 | 0 |
| NS5 | P16615-2 | ATP2A2 | 1 | 23.75 | 0 | 23.75 | 9.4 | 9.24 | 1 | 0 | 22 | 27 | 19 | 27 | 0 | 0 | 0 | 0 |
| NS5 | P04075 | ALDOA | 1 | 26.25 | 0 | 26.25 | 9.63 | 9.15 | 1 | 0 | 10 | 22 | 42 | 31 | 0 | 0 | 0 | 0 |
| NS5 | H0YK48 | TPM1 | 1 | 27.75 | 0 | 27.75 | 10.06 | 9.13 | 1 | 0 | 9 | 17 | 45 | 40 | 0 | 0 | 0 | 0 |
| NS5 | Q99798 | ACO2 | 1 | 28.25 | 0 | 28.25 | 10.42 | 10.04 | 1 | 0 | 14 | 23 | 47 | 29 | 0 | 0 | 0 | 0 |
| NS5 | P69905 | HBA2;HBA1 | 1 | 29 | 0 | 29 | 10.95 | 10.94 | 1 | 0 | 20 | 30 | 33 | 33 | 0 | 0 | 0 | 0 |
| NS5 | P09493-6 | TPM1 | 1 | 32 | 0 | 32 | 11.38 | 10.08 | 1 | 0 | 9 | 19 | 54 | 46 | 0 | 0 | 0 | 0 |
| NS5 | P06732 | CKM | 1 | 38.25 | 0 | 38.25 | 13.72 | 13.28 | 1 | 0 | 18 | 33 | 59 | 43 | 0 | 0 | 0 | 0 |
| NS5 | P06733-2 | ENO1 | 1 | 37.5 | 0 | 37.5 | 13.98 | 13.85 | 1 | 0 | 29 | 31 | 46 | 44 | 0 | 0 | 0 | 0 |
| NS5 | Q6ZN40 | TPM1 | 1 | 40.25 | 0 | 40.25 | 14 | 12.07 | 1 | 0 | 10 | 23 | 67 | 61 | 0 | 0 | 0 | 0 |
| NS5 | Q92616 | GCN1 | 1 | 48.5 | 0 | 48.5 | 19.32 | 15.75 | 1 | 0 | 71 | 62 | 14 | 47 | 0 | 0 | 0 | 0 |
| NS5 | P13533 | MYH6 | 1 | 103.75 | 0 | 103.75 | 34.96 | 31.73 | 1 | 0 | 37 | 68 | 180 | 130 | 0 | 0 | 0 | 0 |
| NS5 | P12883 | MYH7 | 1 | 118.25 | 0 | 118.25 | 39.47 | 34.43 | 1 | 0 | 37 | 67 | 215 | 154 | 0 | 0 | 0 | 0 |
| NS5 | P11055 | MYH3 | 1 | 185.25 | 0 | 185.25 | 66.55 | 65 | 1 | 0 | 172 | 177 | 156 | 236 | 0 | 0 | 0 | 0 |
| NS5 | Q9UKX2 | MYH2 | 1 | 281.5 | 0 | 281.5 | 98.36 | 97.54 | 1 | 0 | 213 | 246 | 316 | 351 | 0 | 0 | 0 | 0 |
| NS5 | P12882 | MYH1 | 1 | 300.75 | 0 | 300.75 | 104.42 | 103.52 | 1 | 0 | 214 | 261 | 350 | 378 | 0 | 0 | 0 | 0 |
| NS5 | P13535 | MYH8 | 1 | 317.75 | 0 | 317.75 | 112.69 | 111.4 | 1 | 0 | 280 | 305 | 307 | 379 | 0 | 0 | 0 | 0 |
| NS5 | Q9Y623 | MYH4 | 1 | 331.75 | 0 | 331.75 | 115.76 | 115.16 | 1 | 0 | 251 | 307 | 366 | 403 | 0 | 0 | 0 | 0 |

**Table S4. List of interactomes in USP7 cells**

| BAIT | PROTID | GENE | preyFreq | preyAvg_AvgCounts | preySD_AvgCounts | NS1_AVG_SPC | NS1_FC_A | NS1_FC_B | NS1_SP | NS1_iREF | IP_USP7_NS1_R4 | IP_USP7_NS1_R3 | IP_USP7_NS1_R2 | IP_USP7_NS1_R1 | IP_USP7_GFP_R8 | IP_USP7_GFP_R7 | IP_USP7_GFP_R6 | IP_USP7_GFP_R5 | IP_USP7_GFP_R4 | IP_USP7_GFP_R3 | IP_USP7_GFP_R1 |
|---|---|---|---|---|---|---|---|---|---|---|---|---|---|---|---|---|---|---|---|---|---|
| NS1 | E7EQR6 | TCP1 | 1 | 137.25 | 0 | 137.25 | 24.75 | 24.71 | 1 | 0 | 118 | 132 | 130 | 169 | 0 | 0 | 0 | 0 | 0 | 0 | 0 |
| NS1 | F5GWF6 | CCT2 | 1 | 195.25 | 0 | 195.25 | 23.98 | 16.84 | 1 | 0 | 183 | 194 | 157 | 247 | 1 | 0 | 0 | 0 | 0 | 0 | 4 |
| NS1 | A0A7I2V599 | HSPD1 | 1 | 125.25 | 0 | 125.25 | 22.92 | 22.25 | 1 | 0 | 147 | 134 | 79 | 141 | 0 | 0 | 0 | 0 | 0 | 0 | 0 |
| NS1 | E9PCA1 | CCT5 | 1 | 119 | 0 | 119 | 21.64 | 21.55 | 1 | 0 | 116 | 108 | 104 | 148 | 0 | 0 | 0 | 0 | 0 | 0 | 0 |
| NS1 | E7ENZ3 | CCT5 | 1 | 127.75 | 0 | 127.75 | 19.84 | 16.62 | 1 | 0 | 118 | 117 | 109 | 167 | 0 | 0 | 0 | 1 | 0 | 0 | 0 |
| NS1 | H7C4C8 | CCT8 | 1 | 93 | 0 | 93 | 17.22 | 17.18 | 1 | 0 | 95 | 91 | 84 | 102 | 0 | 0 | 0 | 0 | 0 | 0 | 0 |
| NS1 | B4DUR8 | CCT3 | 1 | 146.5 | 0 | 146.5 | 15.74 | 10.22 | 1 | 0 | 131 | 138 | 138 | 179 | 0 | 1 | 0 | 3 | 0 | 0 | 0 |
| NS1 | E9PRC8 | CCT3 | 1 | 57.75 | 0 | 57.75 | 10.97 | 10.93 | 1 | 0 | 49 | 54 | 53 | 75 | 0 | 0 | 0 | 0 | 0 | 0 | 0 |
| NS1 | E7ERF2 | TCP1 | 1 | 134.25 | 0 | 134.25 | 10.75 | 7.1 | 1 | 0 | 124 | 129 | 122 | 162 | 2 | 2 | 1 | 1 | 0 | 0 | 1 |
| NS1 | F8WAM2 | CCT7 | 1 | 34 | 0 | 34 | 6.8 | 6.65 | 1 | 0 | 26 | 31 | 25 | 54 | 0 | 0 | 0 | 0 | 0 | 0 | 0 |
| NS1 | E7EMC7 | SQSTM1 | 1 | 16.5 | 0 | 16.5 | 3.83 | 3.79 | 1 | 0 | 12 | 13 | 18 | 23 | 0 | 0 | 0 | 0 | 0 | 0 | 0 |
| NS1 | F8WBP8 | CCT7 | 1 | 14.25 | 0 | 14.25 | 3.47 | 3.46 | 1 | 0 | 13 | 12 | 15 | 17 | 0 | 0 | 0 | 0 | 0 | 0 | 0 |
| NS1 | F8VPD4 | CAD | 1 | 11.75 | 0 | 11.75 | 3.04 | 3.03 | 1 | 0 | 11 | 10 | 12 | 14 | 0 | 0 | 0 | 0 | 0 | 0 | 0 |
| NS1 | S4R3D2 | ANKRD13A | 1 | 9 | 0 | 9 | 2.59 | 2.57 | 1 | 0 | 10 | 9 | 10 | 7 | 0 | 0 | 0 | 0 | 0 | 0 | 0 |
| NS1 | H7C2U0 | CCT8 | 1 | 9 | 0 | 9 | 2.56 | 2.56 | 1 | 0 | 8 | 8 | 10 | 10 | 0 | 0 | 0 | 0 | 0 | 0 | 0 |
| NS1 | H0YIN8 | ANKRD13A | 1 | 8.75 | 0 | 8.75 | 2.52 | 2.52 | 1 | 0 | 8 | 9 | 8 | 10 | 0 | 0 | 0 | 0 | 0 | 0 | 0 |
| NS1 | A0A2R8Y6Y7 | SUCLA2 | 1 | 7.25 | 0 | 7.25 | 2.24 | 2.22 | 1 | 0 | 4 | 7 | 9 | 9 | 0 | 0 | 0 | 0 | 0 | 0 | 0 |
| NS1 | A0A2R8Y706 | DYNC1H1 | 1 | 8.75 | 0 | 8.75 | 2.51 | 2.44 | 0.99 | 0 | 4 | 8 | 14 | 9 | 0 | 0 | 0 | 0 | 0 | 0 | 0 |
| NS1 | H0YKD8 | RPL28 | 1 | 7.25 | 0 | 7.25 | 2.26 | 2.23 | 0.99 | 0 | 7 | 4 | 10 | 8 | 0 | 0 | 0 | 0 | 0 | 0 | 0 |
| NS1 | A0A0D9SEI9 | SLC25A22 | 1 | 5 | 0 | 5 | 1.88 | 1.87 | 0.99 | 0 | 5 | 5 | 6 | 4 | 0 | 0 | 0 | 0 | 0 | 0 | 0 |
| NS1 | A0A2R8Y891 | PFKM | 1 | 5 | 0 | 5 | 1.87 | 1.86 | 0.99 | 0 | 4 | 4 | 7 | 5 | 0 | 0 | 0 | 0 | 0 | 0 | 0 |
| NS1 | A0A6I8PTT9 | GFPT1 | 1 | 5.5 | 0 | 5.5 | 1.97 | 1.95 | 0.98 | 0 | 6 | 4 | 7 | 5 | 0 | 0 | 0 | 0 | 0 | 0 | 0 |
| NS1 | M0R0Y3 | RUVBL2 | 1 | 4.75 | 0 | 4.75 | 1.83 | 1.83 | 0.98 | 0 | 4 | 5 | 5 | 5 | 0 | 0 | 0 | 0 | 0 | 0 | 0 |
| NS1 | C9JFK9 | BAG3 | 1 | 5.25 | 0 | 5.25 | 1.91 | 1.87 | 0.97 | 0 | 7 | 3 | 3 | 8 | 0 | 0 | 0 | 0 | 0 | 0 | 0 |
| NS1 | C9JAW5 | C9JAW5 | 1 | 3.5 | 0 | 3.5 | 1.61 | 1.6 | 0.97 | 0 | 3 | 2 | 5 | 4 | 0 | 0 | 0 | 0 | 0 | 0 | 0 |
| NS1 | H7C5W9 | ATP2A2 | 1 | 5 | 0 | 5 | 1.88 | 1.87 | 0.96 | 0 | 6 | 5 | 4 | 5 | 0 | 0 | 0 | 0 | 0 | 0 | 0 |
| NS1 | H7C4G5 | RUVBL1 | 1 | 3.75 | 0 | 3.75 | 1.64 | 1.63 | 0.96 | 0 | 3 | 4 | 2 | 6 | 0 | 0 | 0 | 0 | 0 | 0 | 0 |
| NS1 | C9JNW5 | RPL24 | 1 | 4.25 | 0 | 4.25 | 1.72 | 1.7 | 0.95 | 0 | 3 | 2 | 5 | 7 | 0 | 0 | 0 | 0 | 0 | 0 | 0 |
| NS1 | C9IZN5 | NDUFA5 | 1 | 3.75 | 0 | 3.75 | 1.64 | 1.63 | 0.95 | 0 | 2 | 3 | 5 | 5 | 0 | 0 | 0 | 0 | 0 | 0 | 0 |
| NS1 | G3V1B3 | RPL21 | 1 | 2.75 | 0 | 2.75 | 1.48 | 1.48 | 0.95 | 0 | 3 | 3 | 2 | 3 | 0 | 0 | 0 | 0 | 0 | 0 | 0 |
| NS1 | A0A0A0MTH3 | ILK | 1 | 4 | 0 | 4 | 1.71 | 1.69 | 0.94 | 0 | 4 | 5 | 5 | 2 | 0 | 0 | 0 | 0 | 0 | 0 | 0 |
| NS1 | X6R2L4 | RUVBL2 | 1 | 3.25 | 0 | 3.25 | 1.57 | 1.57 | 0.94 | 0 | 3 | 3 | 4 | 3 | 0 | 0 | 0 | 0 | 0 | 0 | 0 |
| NS1 | A0A3B3IT92 | MCM4 | 1 | 3 | 0 | 3 | 1.52 | 1.52 | 0.94 | 0 | 2 | 3 | 4 | 3 | 0 | 0 | 0 | 0 | 0 | 0 | 0 |
| NS1 | E9PNS3 | TOLLIP | 1 | 3 | 0 | 3 | 1.52 | 1.52 | 0.94 | 0 | 2 | 4 | 3 | 3 | 0 | 0 | 0 | 0 | 0 | 0 | 0 |
| NS1 | M0R257 | RAB3A | 1 | 2.75 | 0 | 2.75 | 1.48 | 1.48 | 0.94 | 0 | 2 | 3 | 3 | 3 | 0 | 0 | 0 | 0 | 0 | 0 | 0 |
| NS1 | E7ETR0 | RUVBL1 | 1 | 2.75 | 0 | 2.75 | 1.48 | 1.48 | 0.94 | 0 | 3 | 3 | 2 | 3 | 0 | 0 | 0 | 0 | 0 | 0 | 0 |
| NS1 | K7EKP8 | ACOT7 | 1 | 2.75 | 0 | 2.75 | 1.48 | 1.48 | 0.93 | 0 | 3 | 2 | 3 | 3 | 0 | 0 | 0 | 0 | 0 | 0 | 0 |
| NS1 | M0QXB4 | COPE | 1 | 2.5 | 0 | 2.5 | 1.44 | 1.43 | 0.93 | 0 | 2 | 3 | 3 | 2 | 0 | 0 | 0 | 0 | 0 | 0 | 0 |
| NS1 | A0A7P0TAG0 | MMP14 | 1 | 3 | 0 | 3 | 1.53 | 1.53 | 0.92 | 0 | 4 | 3 | 3 | 2 | 0 | 0 | 0 | 0 | 0 | 0 | 0 |
| NS1 | E9PEL3 | RPL37A | 1 | 2.5 | 0 | 2.5 | 1.44 | 1.43 | 0.92 | 0 | 2 | 3 | 3 | 2 | 0 | 0 | 0 | 0 | 0 | 0 | 0 |
| NS1 | H3BS86 | STUB1 | 1 | 2.5 | 0 | 2.5 | 1.44 | 1.44 | 0.91 | 0 | 3 | 3 | 2 | 2 | 0 | 0 | 0 | 0 | 0 | 0 | 0 |
| NS1 | H7C5Q2 | ACOT9 | 1 | 3.25 | 0 | 3.25 | 1.57 | 1.55 | 0.9 | 0 | 4 | 1 | 4 | 4 | 0 | 0 | 0 | 0 | 0 | 0 | 0 |
| NS1 | Q5T8D1 | YME1L1 | 1 | 2 | 0 | 2 | 1.35 | 1.35 | 0.9 | 0 | 2 | 2 | 2 | 2 | 0 | 0 | 0 | 0 | 0 | 0 | 0 |
| NS1 | B1AHC9 | XRCC6 | 1 | 3 | 0 | 3 | 1.52 | 1.52 | 0.89 | 0 | 3 | 2 | 3 | 4 | 0 | 0 | 0 | 0 | 0 | 0 | 0 |
| NS1 | E7EQV9 | RPL15 | 1 | 2.25 | 0 | 2.25 | 1.4 | 1.39 | 0.89 | 0 | 3 | 2 | 3 | 1 | 0 | 0 | 0 | 0 | 0 | 0 | 0 |
| NS1 | B3KY94 | CDIPT | 1 | 2 | 0 | 2 | 1.35 | 1.35 | 0.89 | 0 | 2 | 2 | 2 | 2 | 0 | 0 | 0 | 0 | 0 | 0 | 0 |
| NS1 | A0A087X2I1 | PSMC6 | 1 | 2.5 | 0 | 2.5 | 1.43 | 1.43 | 0.88 | 0 | 2 | 2 | 3 | 3 | 0 | 0 | 0 | 0 | 0 | 0 | 0 |
| NS1 | E5RGS4 | PFDN1 | 1 | 2.5 | 0 | 2.5 | 1.42 | 1.41 | 0.88 | 0 | 1 | 2 | 3 | 4 | 0 | 0 | 0 | 0 | 0 | 0 | 0 |
| NS1 | A0A499FIX8 | ESYT2 | 1 | 2.75 | 0 | 2.75 | 1.48 | 1.47 | 0.87 | 0 | 2 | 2 | 4 | 3 | 0 | 0 | 0 | 0 | 0 | 0 | 0 |
| NS1 | A0A7I2YQG2 | RPL19 | 1 | 2.75 | 0 | 2.75 | 1.47 | 1.47 | 0.87 | 0 | 2 | 2 | 3 | 4 | 0 | 0 | 0 | 0 | 0 | 0 | 0 |
| NS1 | E9PM69 | PSMC3 | 1 | 2 | 0 | 2 | 1.35 | 1.35 | 0.87 | 0 | 2 | 2 | 2 | 2 | 0 | 0 | 0 | 0 | 0 | 0 | 0 |
| NS1 | H0YGV8 | PSMD3 | 1 | 2.25 | 0 | 2.25 | 1.39 | 1.38 | 0.86 | 0 | 1 | 2 | 4 | 2 | 0 | 0 | 0 | 0 | 0 | 0 | 0 |
| NS1 | A0A087WX59 | ADRM1 | 1 | 2 | 0 | 2 | 1.35 | 1.35 | 0.86 | 0 | 2 | 2 | 2 | 2 | 0 | 0 | 0 | 0 | 0 | 0 | 0 |
| NS1 | R4GNA5 | YME1L1 | 1 | 2 | 0 | 2 | 1.34 | 1.34 | 0.86 | 0 | 2 | 1 | 2 | 3 | 0 | 0 | 0 | 0 | 0 | 0 | 0 |
| NS1 | E9PJH7 | SLC25A22 | 1 | 4.75 | 0 | 4.75 | 1.55 | 1.28 | 0.85 | 0 | 5 | 4 | 6 | 4 | 0 | 1 | 0 | 0 | 0 | 0 | 0 |

| BAIT | PROTID | GENE | preyFreq | preyAvg_AvgCounts | preySD_AvgCounts | NS2B_AVG_SPC | NS2B_FC_A | NS2B_FC_B | NS2B_SP | NS2B_iREF | IP_USP7_NS2B_R4 | IP_USP7_NS2B_R3 | IP_USP7_NS2B_R2 | IP_USP7_NS2B_R1 | IP_USP7_GFP_R8 | IP_USP7_GFP_R7 | IP_USP7_GFP_R6 | IP_USP7_GFP_R5 | IP_USP7_GFP_R4 | IP_USP7_GFP_R3 | IP_USP7_GFP_R1 |
|---|---|---|---|---|---|---|---|---|---|---|---|---|---|---|---|---|---|---|---|---|---|
| NS2B | E7EQR6 | TCP1 | 1 | 33.5 | 0 | 33.5 | 6.56 | 6.51 | 1 | 0 | 30 | 33 | 29 | 42 | 0 | 0 | 0 | 0 | 0 | 0 | 0 |
| NS2B | A0A2R8Y706 | DYNC1H1 | 1 | 26.5 | 0 | 26.5 | 5.4 | 5.37 | 1 | 0 | 25 | 30 | 21 | 30 | 0 | 0 | 0 | 0 | 0 | 0 | 0 |
| NS2B | A0A0D9SEI9 | SLC25A22 | 1 | 20.5 | 0 | 20.5 | 4.41 | 4.41 | 1 | 0 | 20 | 21 | 20 | 21 | 0 | 0 | 0 | 0 | 0 | 0 | 0 |
| NS2B | E9PJH7 | SLC25A22 | 1 | 23.75 | 0 | 23.75 | 4.17 | 3.45 | 1 | 0 | 24 | 23 | 21 | 27 | 0 | 1 | 0 | 0 | 0 | 0 | 0 |
| NS2B | A0A286YF22 | PHGDH | 1 | 22.25 | 0 | 22.25 | 3.96 | 3.28 | 1 | 0 | 22 | 21 | 22 | 24 | 0 | 1 | 0 | 0 | 0 | 0 | 0 |
| NS2B | C9JX88 | C9JX88 | 1 | 17.5 | 0 | 17.5 | 3.91 | 3.91 | 1 | 0 | 17 | 18 | 18 | 17 | 0 | 0 | 0 | 0 | 0 | 0 | 0 |
| NS2B | H7C5W9 | ATP2A2 | 1 | 15.75 | 0 | 15.75 | 3.61 | 3.6 | 1 | 0 | 14 | 17 | 14 | 18 | 0 | 0 | 0 | 0 | 0 | 0 | 0 |
| NS2B | A0A8I5KVV1 | AFG3L2 | 1 | 15.25 | 0 | 15.25 | 3.53 | 3.49 | 1 | 0 | 15 | 19 | 10 | 17 | 0 | 0 | 0 | 0 | 0 | 0 | 0 |
| NS2B | A0A2R8Y6Y7 | SUCLA2 | 1 | 14 | 0 | 14 | 3.33 | 3.33 | 1 | 0 | 14 | 13 | 14 | 15 | 0 | 0 | 0 | 0 | 0 | 0 | 0 |
| NS2B | K7ER00 | FARSA | 1 | 20.5 | 0 | 20.5 | 3.12 | 2.24 | 1 | 0 | 18 | 23 | 19 | 22 | 0 | 0 | 1 | 1 | 0 | 0 | 0 |
| NS2B | E9PCA1 | CCT5 | 1 | 12.75 | 0 | 12.75 | 3.11 | 3.03 | 1 | 0 | 10 | 10 | 10 | 21 | 0 | 0 | 0 | 0 | 0 | 0 | 0 |
| NS2B | H0YKF9 | ALDH1A3 | 1 | 12.5 | 0 | 12.5 | 3.08 | 3.06 | 1 | 0 | 14 | 15 | 11 | 10 | 0 | 0 | 0 | 0 | 0 | 0 | 0 |
| NS2B | A0A024QZP7 | CDC2 | 1 | 12.25 | 0 | 12.25 | 3.03 | 3.03 | 1 | 0 | 12 | 14 | 11 | 12 | 0 | 0 | 0 | 0 | 0 | 0 | 0 |
| NS2B | A0A6I8PTT9 | GFPT1 | 1 | 12 | 0 | 12 | 2.99 | 2.97 | 1 | 0 | 9 | 12 | 13 | 14 | 0 | 0 | 0 | 0 | 0 | 0 | 0 |
| NS2B | A0A1B0GV93 | HSD17B12 | 1 | 11.75 | 0 | 11.75 | 2.96 | 2.95 | 1 | 0 | 12 | 11 | 12 | 12 | 0 | 0 | 0 | 0 | 0 | 0 | 0 |
| NS2B | E9PK59 | SCYL1 | 1 | 11.5 | 0 | 11.5 | 2.91 | 2.9 | 1 | 0 | 10 | 12 | 11 | 13 | 0 | 0 | 0 | 0 | 0 | 0 | 0 |
| NS2B | A0A7P0TAG0 | MMP14 | 1 | 10.75 | 0 | 10.75 | 2.79 | 2.79 | 1 | 0 | 11 | 11 | 10 | 11 | 0 | 0 | 0 | 0 | 0 | 0 | 0 |
| NS2B | M0R2F8 | TIMM50 | 1 | 10.75 | 0 | 10.75 | 2.78 | 2.77 | 1 | 0 | 9 | 11 | 10 | 13 | 0 | 0 | 0 | 0 | 0 | 0 | 0 |
| NS2B | A0A087X2I1 | PSMC6 | 1 | 10.5 | 0 | 10.5 | 2.74 | 2.73 | 1 | 0 | 8 | 12 | 11 | 11 | 0 | 0 | 0 | 0 | 0 | 0 | 0 |
| NS2B | E9PRC8 | CCT3 | 1 | 10.5 | 0 | 10.5 | 2.74 | 2.74 | 1 | 0 | 9 | 11 | 10 | 12 | 0 | 0 | 0 | 0 | 0 | 0 | 0 |
| NS2B | A0A2R8Y891 | PFKM | 1 | 9.75 | 0 | 9.75 | 2.63 | 2.57 | 1 | 0 | 14 | 7 | 6 | 12 | 0 | 0 | 0 | 0 | 0 | 0 | 0 |
| NS2B | A0A0A0MTH3 | ILK | 1 | 9.5 | 0 | 9.5 | 2.57 | 2.55 | 1 | 0 | 7 | 11 | 8 | 12 | 0 | 0 | 0 | 0 | 0 | 0 | 0 |
| NS2B | E9PJH8 | SERPINH1 | 1 | 8.5 | 0 | 8.5 | 2.41 | 2.38 | 1 | 0 | 9 | 9 | 5 | 11 | 0 | 0 | 0 | 0 | 0 | 0 | 0 |
| NS2B | A0A1B0GVY6 | HSD17B12 | 1 | 8 | 0 | 8 | 2.33 | 2.32 | 1 | 0 | 9 | 6 | 7 | 10 | 0 | 0 | 0 | 0 | 0 | 0 | 0 |
| NS2B | E7EMC7 | SQSTM1 | 1 | 8 | 0 | 8 | 2.33 | 2.33 | 1 | 0 | 8 | 7 | 9 | 8 | 0 | 0 | 0 | 0 | 0 | 0 | 0 |
| NS2B | H0Y852 | SUCLG2 | 1 | 7.5 | 0 | 7.5 | 2.24 | 2.23 | 1 | 0 | 6 | 10 | 7 | 7 | 0 | 0 | 0 | 0 | 0 | 0 | 0 |
| NS2B | M0R3C3 | TECR | 1 | 7.5 | 0 | 7.5 | 2.24 | 2.23 | 1 | 0 | 6 | 10 | 7 | 7 | 0 | 0 | 0 | 0 | 0 | 0 | 0 |
| NS2B | J3QSA9 | PSMC5 | 1 | 6.75 | 0 | 6.75 | 2.12 | 2.11 | 1 | 0 | 7 | 5 | 7 | 8 | 0 | 0 | 0 | 0 | 0 | 0 | 0 |
| NS2B | B4DP62 | SLC25A1 | 1 | 6.75 | 0 | 6.75 | 2.12 | 2.11 | 1 | 0 | 7 | 6 | 5 | 9 | 0 | 0 | 0 | 0 | 0 | 0 | 0 |
| NS2B | A0A8I5KVL5 | ETFA | 1 | 6.5 | 0 | 6.5 | 2.08 | 2.06 | 1 | 0 | 7 | 8 | 4 | 7 | 0 | 0 | 0 | 0 | 0 | 0 | 0 |
| NS2B | E9PM69 | PSMC3 | 1 | 6.5 | 0 | 6.5 | 2.08 | 2.07 | 1 | 0 | 6 | 7 | 5 | 8 | 0 | 0 | 0 | 0 | 0 | 0 | 0 |
| NS2B | A0A669KB68 | STAT1 | 1 | 6.5 | 0 | 6.5 | 2.07 | 2.05 | 1 | 0 | 5 | 8 | 4 | 9 | 0 | 0 | 0 | 0 | 0 | 0 | 0 |
| NS2B | A0A2R8Y3X5 | OPA1 | 1 | 6.25 | 0 | 6.25 | 2.04 | 2.03 | 1 | 0 | 5 | 7 | 6 | 7 | 0 | 0 | 0 | 0 | 0 | 0 | 0 |
| NS2B | C9JNW5 | RPL24 | 1 | 6 | 0 | 6 | 2 | 1.99 | 1 | 0 | 5 | 7 | 7 | 5 | 0 | 0 | 0 | 0 | 0 | 0 | 0 |
| NS2B | M0R003 | TIMM50 | 1 | 5.75 | 0 | 5.75 | 1.96 | 1.95 | 1 | 0 | 6 | 6 | 5 | 6 | 0 | 0 | 0 | 0 | 0 | 0 | 0 |
| NS2B | H0YKD8 | RPL28 | 1 | 5.5 | 0 | 5.5 | 1.92 | 1.91 | 1 | 0 | 7 | 5 | 4 | 6 | 0 | 0 | 0 | 0 | 0 | 0 | 0 |
| NS2B | E9PKH2 | SERPINH1 | 1 | 5 | 0 | 5 | 1.83 | 1.83 | 1 | 0 | 6 | 5 | 4 | 5 | 0 | 0 | 0 | 0 | 0 | 0 | 0 |
| NS2B | J3QQY2 | TMCO1 | 1 | 4 | 0 | 4 | 1.67 | 1.66 | 1 | 0 | 4 | 3 | 4 | 5 | 0 | 0 | 0 | 0 | 0 | 0 | 0 |
| NS2B | A0A3B3IRI2 | CTPS1 | 1 | 14.75 | 0 | 14.75 | 3.25 | 2.99 | 0.99 | 0 | 13 | 19 | 11 | 16 | 0 | 0 | 0 | 0 | 0 | 0 | 1 |
| NS2B | A0A7I2V3Z0 | PFKP | 1 | 18.25 | 0 | 18.25 | 3.23 | 2.54 | 0.99 | 0 | 17 | 21 | 14 | 21 | 0 | 0 | 1 | 0 | 0 | 0 | 0 |
| NS2B | G3V0E5 | TFRC | 1 | 20 | 0 | 20 | 3.06 | 2.19 | 0.99 | 0 | 19 | 21 | 16 | 24 | 0 | 0 | 1 | 1 | 0 | 0 | 0 |
| NS2B | E9PDQ8 | SUCLG2 | 1 | 14.5 | 0 | 14.5 | 2.87 | 2.36 | 0.99 | 0 | 12 | 18 | 13 | 15 | 0 | 1 | 0 | 0 | 0 | 0 | 0 |
| NS2B | F5H2F4 | MTHFD1 | 1 | 13.25 | 0 | 13.25 | 2.73 | 2.27 | 0.99 | 0 | 13 | 16 | 12 | 12 | 0 | 0 | 0 | 0 | 1 | 0 | 0 |
| NS2B | K7EMW4 | NCLN | 1 | 7.75 | 0 | 7.75 | 2.29 | 2.29 | 0.99 | 0 | 7 | 8 | 8 | 8 | 0 | 0 | 0 | 0 | 0 | 0 | 0 |
| NS2B | G3V0E4 | PMPCB | 1 | 7 | 0 | 7 | 2.16 | 2.16 | 0.99 | 0 | 6 | 7 | 7 | 8 | 0 | 0 | 0 | 0 | 0 | 0 | 0 |
| NS2B | A0A7P0TA76 | ACSL3 | 1 | 6.75 | 0 | 6.75 | 2.12 | 2.1 | 0.99 | 0 | 8 | 7 | 4 | 8 | 0 | 0 | 0 | 0 | 0 | 0 | 0 |
| NS2B | E9PLK3 | NPEPPS | 1 | 6.75 | 0 | 6.75 | 2.11 | 2.09 | 0.99 | 0 | 4 | 6 | 7 | 10 | 0 | 0 | 0 | 0 | 0 | 0 | 0 |
| NS2B | A0A494C0G1 | PGM3 | 1 | 6.75 | 0 | 6.75 | 2.11 | 2.09 | 0.99 | 0 | 4 | 9 | 6 | 8 | 0 | 0 | 0 | 0 | 0 | 0 | 0 |
| NS2B | A0A2R8Y6P0 | MSH2 | 1 | 6 | 0 | 6 | 2 | 1.99 | 0.99 | 0 | 6 | 6 | 5 | 7 | 0 | 0 | 0 | 0 | 0 | 0 | 0 |
| NS2B | A0A8I5KUM4 | SPTLC1 | 1 | 6 | 0 | 6 | 2 | 1.97 | 0.99 | 0 | 7 | 8 | 3 | 6 | 0 | 0 | 0 | 0 | 0 | 0 | 0 |
| NS2B | K7EII7 | GALK1 | 1 | 5.5 | 0 | 5.5 | 1.91 | 1.9 | 0.99 | 0 | 5 | 8 | 4 | 5 | 0 | 0 | 0 | 0 | 0 | 0 | 0 |
| NS2B | B4DR61 | SEC61A1 | 1 | 5.25 | 0 | 5.25 | 1.87 | 1.87 | 0.99 | 0 | 5 | 5 | 5 | 6 | 0 | 0 | 0 | 0 | 0 | 0 | 0 |
| NS2B | A0A087WTV6 | PYCR2 | 1 | 5.25 | 0 | 5.25 | 1.87 | 1.86 | 0.99 | 0 | 5 | 4 | 5 | 7 | 0 | 0 | 0 | 0 | 0 | 0 | 0 |
| NS2B | H0YM31 | PCK2 | 1 | 5 | 0 | 5 | 1.83 | 1.83 | 0.99 | 0 | 5 | 6 | 4 | 5 | 0 | 0 | 0 | 0 | 0 | 0 | 0 |
| NS2B | S4R3D2 | ANKRD13A | 1 | 5 | 0 | 5 | 1.83 | 1.83 | 0.99 | 0 | 4 | 6 | 5 | 5 | 0 | 0 | 0 | 0 | 0 | 0 | 0 |
| NS2B | A0A087WW66 | PSMD1 | 1 | 4.75 | 0 | 4.75 | 1.79 | 1.78 | 0.99 | 0 | 5 | 6 | 3 | 5 | 0 | 0 | 0 | 0 | 0 | 0 | 0 |
| NS2B | H3BTB7 | EARS2 | 1 | 4.75 | 0 | 4.75 | 1.79 | 1.78 | 0.99 | 0 | 4 | 4 | 4 | 7 | 0 | 0 | 0 | 0 | 0 | 0 | 0 |
| NS2B | H0YGV8 | PSMD3 | 1 | 4.5 | 0 | 4.5 | 1.75 | 1.75 | 0.99 | 0 | 5 | 5 | 4 | 4 | 0 | 0 | 0 | 0 | 0 | 0 | 0 |
| NS2B | A0A6I8PIV1 | LARS1 | 1 | 4.5 | 0 | 4.5 | 1.75 | 1.74 | 0.99 | 0 | 4 | 4 | 4 | 6 | 0 | 0 | 0 | 0 | 0 | 0 | 0 |
| NS2B | A0A8I5KPP9 | AP2A2 | 1 | 4.25 | 0 | 4.25 | 1.71 | 1.7 | 0.99 | 0 | 6 | 5 | 3 | 3 | 0 | 0 | 0 | 0 | 0 | 0 | 0 |
| NS2B | A0A1B0GUL7 | A0A1B0GUL | 1 | 4.25 | 0 | 4.25 | 1.7 | 1.7 | 0.99 | 0 | 3 | 5 | 4 | 5 | 0 | 0 | 0 | 0 | 0 | 0 | 0 |
| NS2B | C9JAW5 | C9JAW5 | 1 | 4 | 0 | 4 | 1.67 | 1.67 | 0.99 | 0 | 4 | 4 | 4 | 4 | 0 | 0 | 0 | 0 | 0 | 0 | 0 |
| NS2B | H0Y5L2 | ECHDC1 | 1 | 4 | 0 | 4 | 1.67 | 1.66 | 0.99 | 0 | 5 | 4 | 3 | 4 | 0 | 0 | 0 | 0 | 0 | 0 | 0 |
| NS2B | A0A7I2V351 | KPNA2 | 1 | 3.75 | 0 | 3.75 | 1.62 | 1.61 | 0.99 | 0 | 3 | 3 | 3 | 6 | 0 | 0 | 0 | 0 | 0 | 0 | 0 |
| NS2B | F8VVM2 | SLC25A3 | 1 | 52.25 | 0 | 52.25 | 3.1 | 2.18 | 0.98 | 0 | 47 | 57 | 51 | 54 | 2 | 3 | 2 | 2 | 1 | 1 | 0 |
| NS2B | A0A7P0T895 | NAMPT | 1 | 12 | 0 | 12 | 2.39 | 1.88 | 0.98 | 0 | 8 | 15 | 13 | 12 | 0 | 0 | 1 | 0 | 0 | 0 | 0 |
| NS2B | M0QXB4 | COPE | 1 | 5.5 | 0 | 5.5 | 1.9 | 1.87 | 0.98 | 0 | 2 | 8 | 5 | 7 | 0 | 0 | 0 | 0 | 0 | 0 | 0 |
| NS2B | B1AHB1 | MCM5 | 1 | 5.25 | 0 | 5.25 | 1.86 | 1.84 | 0.98 | 0 | 3 | 6 | 4 | 8 | 0 | 0 | 0 | 0 | 0 | 0 | 0 |
| NS2B | A0A6Q8PHC2 | SEPTIN9 | 1 | 5 | 0 | 5 | 1.84 | 1.82 | 0.98 | 0 | 6 | 5 | 6 | 3 | 0 | 0 | 0 | 0 | 0 | 0 | 0 |
| NS2B | A0A0A0MSE2 | HADH | 1 | 5 | 0 | 5 | 1.83 | 1.83 | 0.98 | 0 | 5 | 5 | 5 | 5 | 0 | 0 | 0 | 0 | 0 | 0 | 0 |
| NS2B | E9PB90 | HK2 | 1 | 4.75 | 0 | 4.75 | 1.79 | 1.79 | 0.98 | 0 | 6 | 4 | 4 | 5 | 0 | 0 | 0 | 0 | 0 | 0 | 0 |
| NS2B | A0A8I5KP51 | RARS2 | 1 | 4.75 | 0 | 4.75 | 1.79 | 1.78 | 0.98 | 0 | 5 | 4 | 4 | 6 | 0 | 0 | 0 | 0 | 0 | 0 | 0 |
| NS2B | C9JA28 | SSR3 | 1 | 4.75 | 0 | 4.75 | 1.79 | 1.78 | 0.98 | 0 | 5 | 6 | 3 | 5 | 0 | 0 | 0 | 0 | 0 | 0 | 0 |
| NS2B | A0A0S2Z4Z4 | AP3B1 | 1 | 4.75 | 0 | 4.75 | 1.78 | 1.77 | 0.98 | 0 | 3 | 5 | 4 | 7 | 0 | 0 | 0 | 0 | 0 | 0 | 0 |
| NS2B | K7EJE8 | LONP1 | 1 | 4.5 | 0 | 4.5 | 1.75 | 1.74 | 0.98 | 0 | 3 | 4 | 6 | 5 | 0 | 0 | 0 | 0 | 0 | 0 | 0 |
| NS2B | A0A804HI56 | SLC25A25 | 1 | 4.5 | 0 | 4.5 | 1.74 | 1.74 | 0.98 | 0 | 3 | 6 | 5 | 4 | 0 | 0 | 0 | 0 | 0 | 0 | 0 |
| NS2B | A0A0A0MTJ1 | FKBP8 | 1 | 4.25 | 0 | 4.25 | 1.71 | 1.7 | 0.98 | 0 | 5 | 4 | 3 | 5 | 0 | 0 | 0 | 0 | 0 | 0 | 0 |
| NS2B | C9JP16 | CRTAP | 1 | 4.25 | 0 | 4.25 | 1.7 | 1.7 | 0.98 | 0 | 3 | 4 | 5 | 5 | 0 | 0 | 0 | 0 | 0 | 0 | 0 |
| NS2B | C9JW96 | PHB | 1 | 4 | 0 | 4 | 1.67 | 1.66 | 0.98 | 0 | 5 | 4 | 3 | 4 | 0 | 0 | 0 | 0 | 0 | 0 | 0 |
| NS2B | A0A087WVM4 | MTHFD1L | 1 | 4 | 0 | 4 | 1.66 | 1.66 | 0.98 | 0 | 4 | 4 | 3 | 5 | 0 | 0 | 0 | 0 | 0 | 0 | 0 |
| NS2B | A6NKB8 | RNPEP | 1 | 4 | 0 | 4 | 1.66 | 1.66 | 0.98 | 0 | 4 | 5 | 3 | 4 | 0 | 0 | 0 | 0 | 0 | 0 | 0 |
| NS2B | F5H6X2 | MGST1 | 1 | 3.75 | 0 | 3.75 | 1.62 | 1.61 | 0.98 | 0 | 3 | 5 | 2 | 5 | 0 | 0 | 0 | 0 | 0 | 0 | 0 |
| NS2B | J3KQL8 | APOL2 | 1 | 3.75 | 0 | 3.75 | 1.62 | 1.62 | 0.98 | 0 | 4 | 5 | 3 | 3 | 0 | 0 | 0 | 0 | 0 | 0 | 0 |
| NS2B | E7EQB8 | IDH3G | 1 | 3.75 | 0 | 3.75 | 1.62 | 1.62 | 0.98 | 0 | 4 | 4 | 4 | 3 | 0 | 0 | 0 | 0 | 0 | 0 | 0 |
| NS2B | M0R257 | RAB3A | 1 | 3.5 | 0 | 3.5 | 1.58 | 1.58 | 0.98 | 0 | 4 | 3 | 4 | 3 | 0 | 0 | 0 | 0 | 0 | 0 | 0 |
| NS2B | B3KY94 | CDIPT | 1 | 3.5 | 0 | 3.5 | 1.58 | 1.58 | 0.98 | 0 | 4 | 4 | 3 | 3 | 0 | 0 | 0 | 0 | 0 | 0 | 0 |
| NS2B | A0A804HHW9 | IARS1 | 1 | 3.5 | 0 | 3.5 | 1.58 | 1.57 | 0.98 | 0 | 3 | 4 | 2 | 5 | 0 | 0 | 0 | 0 | 0 | 0 | 0 |
| NS2B | C9K068 | NIPSNAP2 | 1 | 3.5 | 0 | 3.5 | 1.58 | 1.58 | 0.98 | 0 | 3 | 3 | 4 | 4 | 0 | 0 | 0 | 0 | 0 | 0 | 0 |
| NS2B | E9PKH6 | NDUFS8 | 1 | 3.25 | 0 | 3.25 | 1.54 | 1.53 | 0.98 | 0 | 2 | 3 | 3 | 5 | 0 | 0 | 0 | 0 | 0 | 0 | 0 |
| NS2B | F5H6I7 | ATL3 | 1 | 3.25 | 0 | 3.25 | 1.54 | 1.53 | 0.98 | 0 | 2 | 4 | 3 | 4 | 0 | 0 | 0 | 0 | 0 | 0 | 0 |
| NS2B | A0A087WVC4 | PRKACB | 1 | 3.25 | 0 | 3.25 | 1.54 | 1.54 | 0.98 | 0 | 4 | 3 | 4 | 2 | 0 | 0 | 0 | 0 | 0 | 0 | 0 |
| NS2B | E5RHK8 | DNM3 | 1 | 3.25 | 0 | 3.25 | 1.54 | 1.53 | 0.98 | 0 | 2 | 4 | 4 | 3 | 0 | 0 | 0 | 0 | 0 | 0 | 0 |
| NS2B | H0YHV5 | MARS | 1 | 3 | 0 | 3 | 1.5 | 1.5 | 0.98 | 0 | 4 | 3 | 2 | 3 | 0 | 0 | 0 | 0 | 0 | 0 | 0 |
| NS2B | E9PNW8 | FAR1 | 1 | 3 | 0 | 3 | 1.5 | 1.5 | 0.98 | 0 | 3 | 3 | 3 | 3 | 0 | 0 | 0 | 0 | 0 | 0 | 0 |
| NS2B | C9JBG0 | RAB34 | 1 | 3 | 0 | 3 | 1.5 | 1.49 | 0.98 | 0 | 3 | 3 | 2 | 4 | 0 | 0 | 0 | 0 | 0 | 0 | 0 |
| NS2B | A0A499FIX8 | ESYT2 | 1 | 2.75 | 0 | 2.75 | 1.46 | 1.46 | 0.98 | 0 | 3 | 3 | 2 | 3 | 0 | 0 | 0 | 0 | 0 | 0 | 0 |
| NS2B | F5H608 | ATP5H | 1 | 2.5 | 0 | 2.5 | 1.42 | 1.41 | 0.98 | 0 | 3 | 3 | 2 | 2 | 0 | 0 | 0 | 0 | 0 | 0 | 0 |
| NS2B | E7ENZ3 | CCT5 | 1 | 12 | 0 | 12 | 2.57 | 2.12 | 0.97 | 0 | 10 | 9 | 10 | 19 | 0 | 0 | 0 | 1 | 0 | 0 | 0 |
| NS2B | A0A1W2PNX8 | UNC45A | 1 | 5 | 0 | 5 | 1.83 | 1.83 | 0.97 | 0 | 5 | 6 | 4 | 5 | 0 | 0 | 0 | 0 | 0 | 0 | 0 |
| NS2B | G3V1J5 | DIS3 | 1 | 4 | 0 | 4 | 1.66 | 1.66 | 0.97 | 0 | 4 | 5 | 3 | 4 | 0 | 0 | 0 | 0 | 0 | 0 | 0 |
| NS2B | E9PGM4 | GBE1 | 1 | 3.75 | 0 | 3.75 | 1.63 | 1.61 | 0.97 | 0 | 5 | 4 | 2 | 4 | 0 | 0 | 0 | 0 | 0 | 0 | 0 |
| NS2B | B0YIW6 | ARCN1 | 1 | 3.75 | 0 | 3.75 | 1.62 | 1.61 | 0.97 | 0 | 4 | 3 | 2 | 6 | 0 | 0 | 0 | 0 | 0 | 0 | 0 |
| NS2B | B4E2Q0 | ATP2C1 | 1 | 3.75 | 0 | 3.75 | 1.62 | 1.62 | 0.97 | 0 | 3 | 3 | 4 | 5 | 0 | 0 | 0 | 0 | 0 | 0 | 0 |
| NS2B | M0R026 | ILVBL | 1 | 3.75 | 0 | 3.75 | 1.62 | 1.62 | 0.97 | 0 | 3 | 4 | 5 | 3 | 0 | 0 | 0 | 0 | 0 | 0 | 0 |
| NS2B | Q5SV16 | ATAD3A | 1 | 3.5 | 0 | 3.5 | 1.59 | 1.57 | 0.97 | 0 | 5 | 4 | 3 | 2 | 0 | 0 | 0 | 0 | 0 | 0 | 0 |
| NS2B | A0A804HJ08 | TUBGCP2 | 1 | 3.5 | 0 | 3.5 | 1.58 | 1.58 | 0.97 | 0 | 4 | 3 | 3 | 4 | 0 | 0 | 0 | 0 | 0 | 0 | 0 |
| NS2B | K7ERP6 | PRKACA | 1 | 3.25 | 0 | 3.25 | 1.54 | 1.54 | 0.97 | 0 | 4 | 3 | 3 | 3 | 0 | 0 | 0 | 0 | 0 | 0 | 0 |
| NS2B | A0A0J9YXC7 | LIMS4 | 1 | 3.25 | 0 | 3.25 | 1.54 | 1.53 | 0.97 | 0 | 3 | 4 | 2 | 4 | 0 | 0 | 0 | 0 | 0 | 0 | 0 |
| NS2B | H7C1H2 | PSMD2 | 1 | 3 | 0 | 3 | 1.5 | 1.5 | 0.97 | 0 | 3 | 3 | 3 | 3 | 0 | 0 | 0 | 0 | 0 | 0 | 0 |
| NS2B | F8WEJ5 | ASNS | 1 | 3 | 0 | 3 | 1.5 | 1.49 | 0.97 | 0 | 3 | 3 | 4 | 2 | 0 | 0 | 0 | 0 | 0 | 0 | 0 |
| NS2B | H3BRK9 | AMFR | 1 | 3 | 0 | 3 | 1.5 | 1.49 | 0.97 | 0 | 2 | 3 | 4 | 3 | 0 | 0 | 0 | 0 | 0 | 0 | 0 |
| NS2B | H0YCR7 | RNH1 | 1 | 3 | 0 | 3 | 1.5 | 1.5 | 0.97 | 0 | 3 | 3 | 3 | 3 | 0 | 0 | 0 | 0 | 0 | 0 | 0 |
| NS2B | A0A140T913 | HLA-A | 1 | 3 | 0 | 3 | 1.5 | 1.49 | 0.97 | 0 | 4 | 4 | 2 | 2 | 0 | 0 | 0 | 0 | 0 | 0 | 0 |
| NS2B | C9IZN5 | NDUFA5 | 1 | 3 | 0 | 3 | 1.49 | 1.49 | 0.97 | 0 | 2 | 4 | 2 | 4 | 0 | 0 | 0 | 0 | 0 | 0 | 0 |
| NS2B | A0A804HI36 | ACSL4 | 1 | 2.75 | 0 | 2.75 | 1.46 | 1.45 | 0.97 | 0 | 2 | 3 | 3 | 3 | 0 | 0 | 0 | 0 | 0 | 0 | 0 |

| | | | | | | | | | | | | | | | | | | | | | |
|---|---|---|---|---|---|---|---|---|---|---|---|---|---|---|---|---|---|---|---|---|---|
| NS2B | B7Z9C2 | NAP1L1 | 1 | 2.75 | 0 | 2.75 | 1.46 | 1.46 | 0.97 | 0 | 3 | 3 | 2 | 3 | 0 | 0 | 0 | 0 | 0 | 0 | 0 |
| NS2B | A0A140T912 | HLA-C | 1 | 2.75 | 0 | 2.75 | 1.46 | 1.45 | 0.97 | 0 | 4 | 3 | 2 | 2 | 0 | 0 | 0 | 0 | 0 | 0 | 0 |
| NS2B | F5GYJ8 | OTUB1 | 1 | 2.75 | 0 | 2.75 | 1.46 | 1.45 | 0.97 | 0 | 3 | 2 | 2 | 4 | 0 | 0 | 0 | 0 | 0 | 0 | 0 |
| NS2B | A0A087WZN1 | IDH3B | 1 | 9 | 0 | 9 | 2.14 | 1.78 | 0.96 | 0 | 8 | 9 | 6 | 13 | 0 | 0 | 0 | 1 | 0 | 0 | 0 |
| NS2B | F8VPD4 | CAD | 1 | 5 | 0 | 5 | 1.83 | 1.82 | 0.96 | 0 | 5 | 4 | 4 | 7 | 0 | 0 | 0 | 0 | 0 | 0 | 0 |
| NS2B | H0YML5 | PCK2 | 1 | 3.25 | 0 | 3.25 | 1.54 | 1.54 | 0.96 | 0 | 3 | 4 | 3 | 3 | 0 | 0 | 0 | 0 | 0 | 0 | 0 |
| NS2B | H0YIN8 | ANKRD13A | 1 | 3.25 | 0 | 3.25 | 1.54 | 1.53 | 0.96 | 0 | 2 | 4 | 3 | 4 | 0 | 0 | 0 | 0 | 0 | 0 | 0 |
| NS2B | H3BVG0 | NUP93 | 1 | 3.25 | 0 | 3.25 | 1.54 | 1.53 | 0.96 | 0 | 3 | 5 | 2 | 3 | 0 | 0 | 0 | 0 | 0 | 0 | 0 |
| NS2B | E9PK47 | PYGL | 1 | 3 | 0 | 3 | 1.5 | 1.5 | 0.96 | 0 | 3 | 3 | 3 | 3 | 0 | 0 | 0 | 0 | 0 | 0 | 0 |
| NS2B | I3L4J1 | I3L4J1 | 1 | 3 | 0 | 3 | 1.5 | 1.5 | 0.96 | 0 | 3 | 2 | 4 | 3 | 0 | 0 | 0 | 0 | 0 | 0 | 0 |
| NS2B | A0A0C4DGS1 | DDOST | 1 | 3 | 0 | 3 | 1.5 | 1.5 | 0.96 | 0 | 3 | 3 | 3 | 3 | 0 | 0 | 0 | 0 | 0 | 0 | 0 |
| NS2B | A0A7P0T8S8 | SNX9 | 1 | 3 | 0 | 3 | 1.5 | 1.49 | 0.96 | 0 | 3 | 3 | 2 | 4 | 0 | 0 | 0 | 0 | 0 | 0 | 0 |
| NS2B | A0A8I5KUS8 | FOXRED1 | 1 | 2.75 | 0 | 2.75 | 1.46 | 1.46 | 0.96 | 0 | 3 | 3 | 2 | 3 | 0 | 0 | 0 | 0 | 0 | 0 | 0 |
| NS2B | A0A140T953 | HLA-B | 1 | 2.75 | 0 | 2.75 | 1.46 | 1.45 | 0.96 | 0 | 3 | 4 | 2 | 2 | 0 | 0 | 0 | 0 | 0 | 0 | 0 |
| NS2B | J3KNQ3 | PSMD13 | 1 | 2.75 | 0 | 2.75 | 1.46 | 1.46 | 0.96 | 0 | 3 | 3 | 2 | 3 | 0 | 0 | 0 | 0 | 0 | 0 | 0 |
| NS2B | B7Z4L4 | RPN1 | 1 | 2.75 | 0 | 2.75 | 1.46 | 1.45 | 0.96 | 0 | 3 | 2 | 2 | 4 | 0 | 0 | 0 | 0 | 0 | 0 | 0 |
| NS2B | A0A1W2PNP0 | PIGT | 1 | 2.75 | 0 | 2.75 | 1.46 | 1.46 | 0.96 | 0 | 3 | 2 | 3 | 3 | 0 | 0 | 0 | 0 | 0 | 0 | 0 |
| NS2B | A0A2R8YGD1 | TPP1 | 1 | 2.75 | 0 | 2.75 | 1.45 | 1.45 | 0.96 | 0 | 2 | 4 | 2 | 3 | 0 | 0 | 0 | 0 | 0 | 0 | 0 |
| NS2B | H0YID3 | SDSL | 1 | 2.5 | 0 | 2.5 | 1.42 | 1.41 | 0.96 | 0 | 3 | 2 | 3 | 2 | 0 | 0 | 0 | 0 | 0 | 0 | 0 |
| NS2B | B0QZ43 | ERLIN1 | 1 | 2.5 | 0 | 2.5 | 1.42 | 1.41 | 0.96 | 0 | 3 | 2 | 3 | 2 | 0 | 0 | 0 | 0 | 0 | 0 | 0 |
| NS2B | H7C5Q2 | ACOT9 | 1 | 2.5 | 0 | 2.5 | 1.42 | 1.41 | 0.96 | 0 | 3 | 2 | 2 | 3 | 0 | 0 | 0 | 0 | 0 | 0 | 0 |
| NS2B | G3V126 | ATP6V1H | 1 | 2.5 | 0 | 2.5 | 1.42 | 1.41 | 0.96 | 0 | 3 | 3 | 2 | 2 | 0 | 0 | 0 | 0 | 0 | 0 | 0 |
| NS2B | E9PEL3 | RPL37A | 1 | 2.5 | 0 | 2.5 | 1.41 | 1.41 | 0.96 | 0 | 2 | 3 | 2 | 3 | 0 | 0 | 0 | 0 | 0 | 0 | 0 |
| NS2B | K7EPV0 | COA3 | 1 | 2.5 | 0 | 2.5 | 1.41 | 1.41 | 0.96 | 0 | 2 | 2 | 2 | 4 | 0 | 0 | 0 | 0 | 0 | 0 | 0 |
| NS2B | E9PL38 | PSMD13 | 1 | 2.25 | 0 | 2.25 | 1.38 | 1.37 | 0.96 | 0 | 3 | 2 | 2 | 2 | 0 | 0 | 0 | 0 | 0 | 0 | 0 |
| NS2B | B4E1G1 | DERL1 | 1 | 2.25 | 0 | 2.25 | 1.37 | 1.37 | 0.96 | 0 | 2 | 3 | 2 | 2 | 0 | 0 | 0 | 0 | 0 | 0 | 0 |
| NS2B | A0A590UJS2 | GNAS | 1 | 2.25 | 0 | 2.25 | 1.37 | 1.37 | 0.96 | 0 | 2 | 2 | 3 | 2 | 0 | 0 | 0 | 0 | 0 | 0 | 0 |
| NS2B | B8ZZW5 | AVL9 | 1 | 2.25 | 0 | 2.25 | 1.37 | 1.37 | 0.96 | 0 | 2 | 3 | 2 | 2 | 0 | 0 | 0 | 0 | 0 | 0 | 0 |
| NS2B | C9J7S3 | DARS | 1 | 2.25 | 0 | 2.25 | 1.37 | 1.37 | 0.96 | 0 | 2 | 2 | 2 | 3 | 0 | 0 | 0 | 0 | 0 | 0 | 0 |
| NS2B | I3L4C5 | UBE2Z | 1 | 2 | 0 | 2 | 1.33 | 1.33 | 0.96 | 0 | 2 | 2 | 2 | 2 | 0 | 0 | 0 | 0 | 0 | 0 | 0 |
| NS2B | D6R9X8 | ITGA3 | 1 | 2 | 0 | 2 | 1.33 | 1.33 | 0.96 | 0 | 2 | 2 | 2 | 2 | 0 | 0 | 0 | 0 | 0 | 0 | 0 |
| NS2B | I3L1D2 | MPDU1 | 1 | 2 | 0 | 2 | 1.33 | 1.33 | 0.96 | 0 | 2 | 2 | 2 | 2 | 0 | 0 | 0 | 0 | 0 | 0 | 0 |
| NS2B | A6NIC1 | RTCA | 1 | 2 | 0 | 2 | 1.33 | 1.33 | 0.96 | 0 | 2 | 2 | 2 | 2 | 0 | 0 | 0 | 0 | 0 | 0 | 0 |
| NS2B | H3BNX3 | H3BNX3 | 1 | 14.5 | 0 | 14.5 | 2.44 | 1.76 | 0.95 | 0 | 13 | 14 | 12 | 19 | 1 | 1 | 0 | 0 | 0 | 0 | 0 |
| NS2B | F5H4B6 | ALDH16A1 | 1 | 2.5 | 0 | 2.5 | 1.41 | 1.41 | 0.95 | 0 | 2 | 2 | 3 | 3 | 0 | 0 | 0 | 0 | 0 | 0 | 0 |
| NS2B | B1AR62 | HK1 | 1 | 2.25 | 0 | 2.25 | 1.37 | 1.37 | 0.95 | 0 | 2 | 3 | 2 | 2 | 0 | 0 | 0 | 0 | 0 | 0 | 0 |
| NS2B | H3BSM7 | C16ORF58 | 1 | 2.25 | 0 | 2.25 | 1.37 | 1.37 | 0.95 | 0 | 2 | 3 | 2 | 2 | 0 | 0 | 0 | 0 | 0 | 0 | 0 |
| NS2B | A0A804HIB9 | RAB18 | 1 | 2.25 | 0 | 2.25 | 1.37 | 1.37 | 0.95 | 0 | 2 | 2 | 2 | 3 | 0 | 0 | 0 | 0 | 0 | 0 | 0 |
| NS2B | F8WAG2 | NEK7 | 1 | 2 | 0 | 2 | 1.33 | 1.33 | 0.95 | 0 | 2 | 2 | 2 | 2 | 0 | 0 | 0 | 0 | 0 | 0 | 0 |
| NS2B | D6RFI0 | SFXN1 | 1 | 24 | 0 | 24 | 2.29 | 1.5 | 0.94 | 0 | 25 | 26 | 22 | 23 | 2 | 1 | 1 | 2 | 0 | 0 | 0 |
| NS2B | J3QQM1 | PSMC5 | 1 | 7.5 | 0 | 7.5 | 1.9 | 1.56 | 0.94 | 0 | 8 | 6 | 7 | 9 | 0 | 1 | 0 | 0 | 0 | 0 | 0 |
| NS2B | A0A2R8Y6J3 | RPL5 | 1 | 2.75 | 0 | 2.75 | 1.46 | 1.45 | 0.94 | 0 | 3 | 4 | 2 | 2 | 0 | 0 | 0 | 0 | 0 | 0 | 0 |
| NS2B | C9J1Z8 | ARF5 | 1 | 2.5 | 0 | 2.5 | 1.42 | 1.41 | 0.94 | 0 | 3 | 3 | 1 | 3 | 0 | 0 | 0 | 0 | 0 | 0 | 0 |
| NS2B | G3V1B3 | RPL21 | 1 | 2.5 | 0 | 2.5 | 1.42 | 1.4 | 0.94 | 0 | 2 | 3 | 4 | 1 | 0 | 0 | 0 | 0 | 0 | 0 | 0 |
| NS2B | H0Y659 | HUWE1 | 1 | 2.5 | 0 | 2.5 | 1.41 | 1.41 | 0.94 | 0 | 2 | 2 | 3 | 3 | 0 | 0 | 0 | 0 | 0 | 0 | 0 |
| NS2B | A0A5F9ZHD4 | ACAT1 | 1 | 2.25 | 0 | 2.25 | 1.37 | 1.37 | 0.94 | 0 | 2 | 2 | 3 | 2 | 0 | 0 | 0 | 0 | 0 | 0 | 0 |
| NS2B | B5MCT7 | PPM1F | 1 | 2 | 0 | 2 | 1.33 | 1.33 | 0.94 | 0 | 2 | 2 | 2 | 2 | 0 | 0 | 0 | 0 | 0 | 0 | 0 |
| NS2B | C9J236 | NDUFAF7 | 1 | 2 | 0 | 2 | 1.33 | 1.33 | 0.94 | 0 | 2 | 2 | 2 | 2 | 0 | 0 | 0 | 0 | 0 | 0 | 0 |
| NS2B | E7ERF2 | TCP1 | 1 | 35.25 | 0 | 35.25 | 3.03 | 1.99 | 0.93 | 0 | 31 | 33 | 33 | 44 | 2 | 2 | 1 | 1 | 0 | 0 | 1 |
| NS2B | H3BNP9 | SQOR | 1 | 7.5 | 0 | 7.5 | 1.93 | 1.62 | 0.93 | 0 | 7 | 6 | 8 | 9 | 0 | 0 | 0 | 1 | 0 | 0 | 0 |
| NS2B | Q5QNZ2 | ATP5F1 | 1 | 3.25 | 0 | 3.25 | 1.55 | 1.52 | 0.93 | 0 | 5 | 1 | 3 | 4 | 0 | 0 | 0 | 0 | 0 | 0 | 0 |
| NS2B | D6RGW0 | ERAP2 | 1 | 2.5 | 0 | 2.5 | 1.42 | 1.41 | 0.93 | 0 | 3 | 2 | 2 | 3 | 0 | 0 | 0 | 0 | 0 | 0 | 0 |
| NS2B | A0A5K1VW56 | ATAD3B | 1 | 2.5 | 0 | 2.5 | 1.42 | 1.41 | 0.93 | 0 | 3 | 2 | 2 | 3 | 0 | 0 | 0 | 0 | 0 | 0 | 0 |
| NS2B | H0Y7R7 | NT5E | 1 | 2.25 | 0 | 2.25 | 1.38 | 1.37 | 0.93 | 0 | 3 | 2 | 3 | 1 | 0 | 0 | 0 | 0 | 0 | 0 | 0 |
| NS2B | K7EPJ3 | DDX39A | 1 | 2.25 | 0 | 2.25 | 1.38 | 1.37 | 0.93 | 0 | 3 | 3 | 2 | 1 | 0 | 0 | 0 | 0 | 0 | 0 | 0 |
| NS2B | B4DEZ3 | NDUFA13 | 1 | 2.25 | 0 | 2.25 | 1.38 | 1.37 | 0.93 | 0 | 3 | 1 | 3 | 2 | 0 | 0 | 0 | 0 | 0 | 0 | 0 |
| NS2B | Q4VXZ8 | DECR2 | 1 | 2.25 | 0 | 2.25 | 1.38 | 1.37 | 0.93 | 0 | 3 | 2 | 3 | 1 | 0 | 0 | 0 | 0 | 0 | 0 | 0 |
| NS2B | E5RJM9 | RARS | 1 | 2.25 | 0 | 2.25 | 1.37 | 1.36 | 0.93 | 0 | 1 | 2 | 3 | 3 | 0 | 0 | 0 | 0 | 0 | 0 | 0 |
| NS2B | H7C5A7 | ACOT8 | 1 | 2 | 0 | 2 | 1.33 | 1.33 | 0.93 | 0 | 2 | 1 | 3 | 2 | 0 | 0 | 0 | 0 | 0 | 0 | 0 |
| NS2B | H0Y2X5 | ALDH1A3 | 1 | 20.25 | 0 | 20.25 | 2.64 | 1.82 | 0.92 | 0 | 22 | 23 | 17 | 19 | 0 | 1 | 1 | 1 | 0 | 0 | 1 |
| NS2B | F5GY37 | PHB2 | 1 | 6.25 | 0 | 6.25 | 1.75 | 1.47 | 0.92 | 0 | 5 | 7 | 6 | 7 | 0 | 0 | 0 | 1 | 0 | 0 | 0 |
| NS2B | A0A087WXC5 | NDUFA10 | 1 | 3.5 | 0 | 3.5 | 1.58 | 1.55 | 0.92 | 0 | 3 | 1 | 4 | 6 | 0 | 0 | 0 | 0 | 0 | 0 | 0 |
| NS2B | A0A6I8PIS1 | SORD | 1 | 2.5 | 0 | 2.5 | 1.41 | 1.4 | 0.92 | 0 | 1 | 4 | 3 | 2 | 0 | 0 | 0 | 0 | 0 | 0 | 0 |
| NS2B | E7EQV9 | RPL15 | 1 | 2.25 | 0 | 2.25 | 1.38 | 1.37 | 0.92 | 0 | 3 | 3 | 2 | 1 | 0 | 0 | 0 | 0 | 0 | 0 | 0 |
| NS2B | H0YAS8 | CLU | 1 | 2.25 | 0 | 2.25 | 1.38 | 1.37 | 0.92 | 0 | 3 | 1 | 2 | 3 | 0 | 0 | 0 | 0 | 0 | 0 | 0 |
| NS2B | F8VYZ3 | SDSL | 1 | 2.25 | 0 | 2.25 | 1.37 | 1.37 | 0.92 | 0 | 2 | 3 | 3 | 1 | 0 | 0 | 0 | 0 | 0 | 0 | 0 |
| NS2B | H7C4H2 | SRPRB | 1 | 2.25 | 0 | 2.25 | 1.37 | 1.36 | 0.92 | 0 | 2 | 1 | 2 | 4 | 0 | 0 | 0 | 0 | 0 | 0 | 0 |
| NS2B | A0A087WX59 | ADRM1 | 1 | 2 | 0 | 2 | 1.33 | 1.33 | 0.92 | 0 | 2 | 2 | 2 | 2 | 0 | 0 | 0 | 0 | 0 | 0 | 0 |
| NS2B | H3BR66 | MMP2 | 1 | 2 | 0 | 2 | 1.33 | 1.32 | 0.92 | 0 | 1 | 3 | 2 | 2 | 0 | 0 | 0 | 0 | 0 | 0 | 0 |
| NS2B | A0A140TA86 | C19ORF70 | 1 | 2 | 0 | 2 | 1.33 | 1.33 | 0.92 | 0 | 2 | 2 | 1 | 3 | 0 | 0 | 0 | 0 | 0 | 0 | 0 |
| NS2B | J3QSB2 | ERAL1 | 1 | 2 | 0 | 2 | 1.33 | 1.33 | 0.92 | 0 | 3 | 2 | 1 | 2 | 0 | 0 | 0 | 0 | 0 | 0 | 0 |
| NS2B | Q5TFJ7 | KPNA6 | 1 | 2 | 0 | 2 | 1.33 | 1.33 | 0.92 | 0 | 1 | 2 | 3 | 2 | 0 | 0 | 0 | 0 | 0 | 0 | 0 |
| NS2B | S4R3V3 | YTHDF2 | 1 | 1.75 | 0 | 1.75 | 1.29 | 1.29 | 0.92 | 0 | 2 | 2 | 1 | 2 | 0 | 0 | 0 | 0 | 0 | 0 | 0 |
| NS2B | A6NGW1 | YIF1A | 1 | 1.75 | 0 | 1.75 | 1.29 | 1.29 | 0.92 | 0 | 2 | 2 | 1 | 2 | 0 | 0 | 0 | 0 | 0 | 0 | 0 |
| NS2B | A8MX49 | PPM1F | 1 | 1.75 | 0 | 1.75 | 1.29 | 1.29 | 0.92 | 0 | 1 | 2 | 2 | 2 | 0 | 0 | 0 | 0 | 0 | 0 | 0 |
| NS2B | E9PIQ7 | HAX1 | 1 | 1.75 | 0 | 1.75 | 1.29 | 1.29 | 0.92 | 0 | 2 | 2 | 1 | 2 | 0 | 0 | 0 | 0 | 0 | 0 | 0 |
| NS2B | H7C1I0 | SLC35E1 | 1 | 1.75 | 0 | 1.75 | 1.29 | 1.29 | 0.92 | 0 | 2 | 2 | 1 | 2 | 0 | 0 | 0 | 0 | 0 | 0 | 0 |
| NS2B | K7EPK9 | DNM2 | 1 | 1.5 | 0 | 1.5 | 1.25 | 1.25 | 0.92 | 0 | 1 | 2 | 2 | 1 | 0 | 0 | 0 | 0 | 0 | 0 | 0 |
| NS2B | H7C2Q3 | PSMD2 | 1 | 1.5 | 0 | 1.5 | 1.25 | 1.24 | 0.92 | 0 | 1 | 2 | 1 | 2 | 0 | 0 | 0 | 0 | 0 | 0 | 0 |
| NS2B | B4DUR8 | CCT3 | 1 | 21.25 | 0 | 21.25 | 2.7 | 1.74 | 0.91 | 0 | 18 | 21 | 18 | 28 | 0 | 1 | 0 | 3 | 0 | 0 | 0 |
| NS2B | A0A494C0M1 | MSH6 | 1 | 3 | 0 | 3 | 1.49 | 1.47 | 0.91 | 0 | 1 | 4 | 2 | 5 | 0 | 0 | 0 | 0 | 0 | 0 | 0 |
| NS2B | C9JZR2 | CTNND1 | 1 | 2.25 | 0 | 2.25 | 1.37 | 1.37 | 0.91 | 0 | 2 | 2 | 2 | 3 | 0 | 0 | 0 | 0 | 0 | 0 | 0 |
| NS2B | Q5VWC4 | PSMD4 | 1 | 2 | 0 | 2 | 1.34 | 1.33 | 0.91 | 0 | 3 | 2 | 2 | 1 | 0 | 0 | 0 | 0 | 0 | 0 | 0 |
| NS2B | H0Y733 | SPTLC3 | 1 | 1.75 | 0 | 1.75 | 1.29 | 1.29 | 0.91 | 0 | 1 | 2 | 2 | 2 | 0 | 0 | 0 | 0 | 0 | 0 | 0 |
| NS2B | A0A087WTP2 | FDPS | 1 | 1.5 | 0 | 1.5 | 1.25 | 1.25 | 0.91 | 0 | 2 | 1 | 1 | 2 | 0 | 0 | 0 | 0 | 0 | 0 | 0 |
| NS2B | A6NLM8 | SSR4 | 1 | 6.5 | 0 | 6.5 | 1.76 | 1.45 | 0.9 | 0 | 7 | 6 | 6 | 7 | 0 | 1 | 0 | 0 | 0 | 0 | 0 |
| NS2B | A0A1B0GVU9 | QARS | 1 | 2.75 | 0 | 2.75 | 1.45 | 1.44 | 0.9 | 0 | 2 | 4 | 1 | 4 | 0 | 0 | 0 | 0 | 0 | 0 | 0 |
| NS2B | A0A7I2V2H0 | XPO1 | 1 | 2.25 | 0 | 2.25 | 1.38 | 1.37 | 0.9 | 0 | 3 | 3 | 1 | 2 | 0 | 0 | 0 | 0 | 0 | 0 | 0 |
| NS2B | A0A0C4DGQ8 | DHRS7B | 1 | 2 | 0 | 2 | 1.33 | 1.33 | 0.9 | 0 | 2 | 2 | 1 | 3 | 0 | 0 | 0 | 0 | 0 | 0 | 0 |
| NS2B | A6NEM5 | PIGK | 1 | 1.75 | 0 | 1.75 | 1.29 | 1.29 | 0.9 | 0 | 2 | 1 | 2 | 2 | 0 | 0 | 0 | 0 | 0 | 0 | 0 |
| NS2B | J3KPT4 | TRABD | 1 | 1.75 | 0 | 1.75 | 1.29 | 1.29 | 0.9 | 0 | 2 | 2 | 2 | 1 | 0 | 0 | 0 | 0 | 0 | 0 | 0 |
| NS2B | Q5JYR7 | RPN2 | 1 | 1.75 | 0 | 1.75 | 1.29 | 1.29 | 0.9 | 0 | 2 | 2 | 1 | 2 | 0 | 0 | 0 | 0 | 0 | 0 | 0 |
| NS2B | J3QR48 | KPNB1 | 1 | 1.5 | 0 | 1.5 | 1.25 | 1.25 | 0.9 | 0 | 1 | 2 | 2 | 1 | 0 | 0 | 0 | 0 | 0 | 0 | 0 |
| NS2B | H3BN55 | RAB27A | 1 | 1.5 | 0 | 1.5 | 1.25 | 1.25 | 0.9 | 0 | 1 | 1 | 2 | 2 | 0 | 0 | 0 | 0 | 0 | 0 | 0 |
| NS2B | E9PPB8 | FAR1 | 1 | 1.5 | 0 | 1.5 | 1.25 | 1.25 | 0.9 | 0 | 2 | 2 | 1 | 1 | 0 | 0 | 0 | 0 | 0 | 0 | 0 |
| NS2B | H3BN72 | COX4I1 | 1 | 1.5 | 0 | 1.5 | 1.25 | 1.25 | 0.9 | 0 | 1 | 1 | 2 | 2 | 0 | 0 | 0 | 0 | 0 | 0 | 0 |
| NS2B | A0A8Q3SIA0 | TIMM17B | 1 | 2.25 | 0 | 2.25 | 1.38 | 1.37 | 0.89 | 0 | 3 | 2 | 1 | 3 | 0 | 0 | 0 | 0 | 0 | 0 | 0 |
| NS2B | B7Z5P7 | LPXN | 1 | 1.75 | 0 | 1.75 | 1.29 | 1.28 | 0.89 | 0 | 1 | 3 | 1 | 2 | 0 | 0 | 0 | 0 | 0 | 0 | 0 |
| NS2B | D6RAA6 | TMEM33 | 1 | 1.75 | 0 | 1.75 | 1.29 | 1.28 | 0.89 | 0 | 2 | 1 | 1 | 3 | 0 | 0 | 0 | 0 | 0 | 0 | 0 |
| NS2B | M0R2K9 | FKBP8 | 1 | 1.5 | 0 | 1.5 | 1.25 | 1.25 | 0.89 | 0 | 2 | 2 | 1 | 1 | 0 | 0 | 0 | 0 | 0 | 0 | 0 |
| NS2B | Q5JXB7 | PLOD1 | 1 | 1.5 | 0 | 1.5 | 1.25 | 1.25 | 0.89 | 0 | 2 | 1 | 1 | 2 | 0 | 0 | 0 | 0 | 0 | 0 | 0 |
| NS2B | J3QR51 | RAB31 | 1 | 1.25 | 0 | 1.25 | 1.21 | 1.21 | 0.89 | 0 | 1 | 1 | 1 | 2 | 0 | 0 | 0 | 0 | 0 | 0 | 0 |
| NS2B | H3BMD2 | CPT1A | 1 | 1.25 | 0 | 1.25 | 1.21 | 1.21 | 0.89 | 0 | 1 | 1 | 1 | 2 | 0 | 0 | 0 | 0 | 0 | 0 | 0 |
| NS2B | A0A7P0T7Z1 | ACAD9 | 1 | 2.5 | 0 | 2.5 | 1.42 | 1.4 | 0.88 | 0 | 3 | 2 | 1 | 4 | 0 | 0 | 0 | 0 | 0 | 0 | 0 |
| NS2B | M0R0W6 | AXL | 1 | 2.25 | 0 | 2.25 | 1.38 | 1.36 | 0.88 | 0 | 3 | 4 | 1 | 1 | 0 | 0 | 0 | 0 | 0 | 0 | 0 |
| NS2B | A0A494C1M4 | ALDH1L2 | 1 | 1.75 | 0 | 1.75 | 1.29 | 1.29 | 0.88 | 0 | 2 | 2 | 1 | 2 | 0 | 0 | 0 | 0 | 0 | 0 | 0 |
| NS2B | Q5TDF0 | NTPCR | 1 | 1.75 | 0 | 1.75 | 1.29 | 1.28 | 0.88 | 0 | 1 | 3 | 1 | 2 | 0 | 0 | 0 | 0 | 0 | 0 | 0 |
| NS2B | A0A2R8Y653 | UCK2 | 1 | 1.75 | 0 | 1.75 | 1.29 | 1.29 | 0.88 | 0 | 2 | 2 | 1 | 2 | 0 | 0 | 0 | 0 | 0 | 0 | 0 |
| NS2B | E9PPM8 | DERA | 1 | 1.75 | 0 | 1.75 | 1.29 | 1.29 | 0.88 | 0 | 2 | 1 | 3 | 1 | 0 | 0 | 0 | 0 | 0 | 0 | 0 |
| NS2B | Q5QPD0 | NT5DC1 | 1 | 1.5 | 0 | 1.5 | 1.25 | 1.25 | 0.88 | 0 | 2 | 1 | 2 | 1 | 0 | 0 | 0 | 0 | 0 | 0 | 0 |
| NS2B | B4E128 | SLFN5 | 1 | 1.5 | 0 | 1.5 | 1.25 | 1.24 | 0.88 | 0 | 1 | 2 | 1 | 2 | 0 | 0 | 0 | 0 | 0 | 0 | 0 |
| NS2B | A0A087WYS6 | PSMA8 | 1 | 1.5 | 0 | 1.5 | 1.25 | 1.25 | 0.88 | 0 | 1 | 1 | 2 | 2 | 0 | 0 | 0 | 0 | 0 | 0 | 0 |
| NS2B | K7ESG8 | ACLY | 1 | 1.5 | 0 | 1.5 | 1.25 | 1.24 | 0.88 | 0 | 1 | 1 | 1 | 3 | 0 | 0 | 0 | 0 | 0 | 0 | 0 |
| NS2B | H3BNF0 | NUBP2 | 1 | 1.5 | 0 | 1.5 | 1.25 | 1.25 | 0.88 | 0 | 2 | 1 | 2 | 1 | 0 | 0 | 0 | 0 | 0 | 0 | 0 |
| NS2B | B4DIE3 | TMX3 | 1 | 1.25 | 0 | 1.25 | 1.21 | 1.21 | 0.88 | 0 | 1 | 1 | 1 | 2 | 0 | 0 | 0 | 0 | 0 | 0 | 0 |
| NS2B | U3KQG0 | TARS2 | 1 | 2 | 0 | 2 | 1.33 | 1.32 | 0.87 | 0 | 1 | 2 | 1 | 4 | 0 | 0 | 0 | 0 | 0 | 0 | 0 |
| NS2B | A0A0C4DFP4 | NGLY1 | 1 | 2 | 0 | 2 | 1.33 | 1.32 | 0.87 | 0 | 1 | 3 | 1 | 3 | 0 | 0 | 0 | 0 | 0 | 0 | 0 |
| NS2B | A0A087X0L7 | OXA1L | 1 | 1.75 | 0 | 1.75 | 1.29 | 1.29 | 0.87 | 0 | 1 | 2 | 2 | 2 | 0 | 0 | 0 | 0 | 0 | 0 | 0 |
| NS2B | A0A087WT48 | DNAJA2 | 1 | 1.25 | 0 | 1.25 | 1.21 | 1.2 | 0.87 | 0 | 1 | 2 | 1 | 1 | 0 | 0 | 0 | 0 | 0 | 0 | 0 |
| NS2B | E5RFV2 | DECR1 | 1 | 1.25 | 0 | 1.25 | 1.21 | 1.21 | 0.87 | 0 | 1 | 1 | 2 | 1 | 0 | 0 | 0 | 0 | 0 | 0 | 0 |
| NS2B | I3L0N3 | NSF | 1 | 12.25 | 0 | 12.25 | 2.18 | 1.58 | 0.86 | 0 | 12 | 12 | 11 | 14 | 1 | 1 | 0 | 0 | 0 | 0 | 0 |
| NS2B | F5H6E2 | MYO1C | 1 | 5 | 0 | 5 | 1.54 | 1.27 | 0.86 | 0 | 5 | 7 | 4 | 4 | 0 | 1 | 0 | 0 | 0 | 0 | 0 |
| NS2B | A0A7I2V381 | IGF2R | 1 | 2.25 | 0 | 2.25 | 1.37 | 1.35 | 0.86 | 0 | 2 | 5 | 1 | 1 | 0 | 0 | 0 | 0 | 0 | 0 | 0 |
| NS2B | A0A8Q3SIC5 | MYL6B | 1 | 1.75 | 0 | 1.75 | 1.29 | 1.29 | 0.86 | 0 | 2 | 2 | 2 | 1 | 0 | 0 | 0 | 0 | 0 | 0 | 0 |
| NS2B | A0A0S2Z4L5 | RAF1 | 1 | 1.75 | 0 | 1.75 | 1.29 | 1.29 | 0.86 | 0 | 2 | 2 | 2 | 1 | 0 | 0 | 0 | 0 | 0 | 0 | 0 |
| NS2B | F8WAM2 | CCT7 | 1 | 1.5 | 0 | 1.5 | 1.25 | 1.24 | 0.86 | 0 | 1 | 3 | 1 | 1 | 0 | 0 | 0 | 0 | 0 | 0 | 0 |
| NS2B | E7ET40 | PLAU | 1 | 1.5 | 0 | 1.5 | 1.25 | 1.25 | 0.86 | 0 | 2 | 1 | 1 | 2 | 0 | 0 | 0 | 0 | 0 | 0 | 0 |
| NS2B | X6R5E5 | UBAC2 | 1 | 1.5 | 0 | 1.5 | 1.25 | 1.25 | 0.86 | 0 | 2 | 2 | 1 | 1 | 0 | 0 | 0 | 0 | 0 | 0 | 0 |
| NS2B | H0YD17 | CD44 | 1 | 1.25 | 0 | 1.25 | 1.21 | 1.21 | 0.86 | 0 | 2 | 1 | 1 | 1 | 0 | 0 | 0 | 0 | 0 | 0 | 0 |
| NS2B | B4DLN1 | B4DLN1 | 1 | 1.5 | 0 | 1.5 | 1.25 | 1.25 | 0.85 | 0 | 1 | 2 | 2 | 1 | 0 | 0 | 0 | 0 | 0 | 0 | 0 |
| NS2B | A0A0B4J207 | PRPS1L1 | 1 | 1.25 | 0 | 1.25 | 1.21 | 1.21 | 0.85 | 0 | 1 | 1 | 2 | 1 | 0 | 0 | 0 | 0 | 0 | 0 | 0 |

| BAIT | PROTID | GENE | preyFreq | preyAvg_AvgCounts | preySD_AvgCounts | NS3_AVG_SPC | NS3_FC_A | NS3_FC_B | NS3_SP | NS3_iREF | IP_USP7_NS3_R4 | IP_USP7_NS3_R3 | IP_USP7_NS3_R2 | IP_USP7_NS3_R1 | IP_USP7_GFP_R8 | IP_USP7_GFP_R7 | IP_USP7_GFP_R6 | IP_USP7_GFP_R5 | IP_USP7_GFP_R4 | IP_USP7_GFP_R3 | IP_USP7_GFP_R1 |
|---|---|---|---|---|---|---|---|---|---|---|---|---|---|---|---|---|---|---|---|---|---|
| NS3 | A0A7I2V599 | HSPD1 | 1 | 75.75 | 0 | 75.75 | 16.83 | 16.26 | 1 | 0 | 88 | 93 | 75 | 47 | 0 | 0 | 0 | 0 | 0 | 0 | 0 |
| NS3 | E7EQR6 | TCP1 | 1 | 11 | 0 | 11 | 3.33 | 3.28 | 1 | 0 | 10 | 10 | 10 | 14 | 0 | 0 | 0 | 0 | 0 | 0 | 0 |
| NS3 | F8VPD4 | CAD | 1 | 7 | 0 | 7 | 2.44 | 2.37 | 0.99 | 0 | 7 | 7 | 3 | 11 | 0 | 0 | 0 | 0 | 0 | 0 | 0 |
| NS3 | A0A7P0TAQ9 | DNAJC10 | 1 | 6.25 | 0 | 6.25 | 2.26 | 2.22 | 0.98 | 0 | 8 | 5 | 3 | 9 | 0 | 0 | 0 | 0 | 0 | 0 | 0 |
| NS3 | E9PCA1 | CCT5 | 1 | 5 | 0 | 5 | 2.07 | 2.04 | 0.98 | 0 | 4 | 7 | 4 | 5 | 0 | 0 | 0 | 0 | 0 | 0 | 0 |
| NS3 | C9JNW5 | RPL24 | 1 | 5 | 0 | 5 | 2.03 | 1.96 | 0.97 | 0 | 5 | 4 | 2 | 9 | 0 | 0 | 0 | 0 | 0 | 0 | 0 |
| NS3 | H7C5W9 | ATP2A2 | 1 | 4.25 | 0 | 4.25 | 1.87 | 1.87 | 0.97 | 0 | 5 | 4 | 3 | 5 | 0 | 0 | 0 | 0 | 0 | 0 | 0 |
| NS3 | A0A0G2JLD8 | SSBP1 | 1 | 3.75 | 0 | 3.75 | 1.79 | 1.78 | 0.97 | 0 | 4 | 4 | 4 | 3 | 0 | 0 | 0 | 0 | 0 | 0 | 0 |
| NS3 | H0YKD8 | RPL28 | 1 | 5 | 0 | 5 | 2.05 | 1.93 | 0.96 | 0 | 4 | 2 | 3 | 11 | 0 | 0 | 0 | 0 | 0 | 0 | 0 |
| NS3 | S4R3D2 | ANKRD13A | 1 | 3.5 | 0 | 3.5 | 1.72 | 1.7 | 0.95 | 0 | 4 | 5 | 2 | 3 | 0 | 0 | 0 | 0 | 0 | 0 | 0 |
| NS3 | E7EMC7 | SQSTM1 | 1 | 3.25 | 0 | 3.25 | 1.66 | 1.66 | 0.94 | 0 | 4 | 3 | 2 | 4 | 0 | 0 | 0 | 0 | 0 | 0 | 0 |
| NS3 | A0A2R8Y706 | DYNC1H1 | 1 | 8.5 | 0 | 8.5 | 2.76 | 2.54 | 0.93 | 0 | 7 | 9 | 2 | 16 | 0 | 0 | 0 | 0 | 0 | 0 | 0 |
| NS3 | H7C4C8 | CCT8 | 1 | 2.75 | 0 | 2.75 | 1.57 | 1.57 | 0.92 | 0 | 3 | 3 | 2 | 3 | 0 | 0 | 0 | 0 | 0 | 0 | 0 |
| NS3 | A0A2R8Y6Y7 | SUCLA2 | 1 | 3 | 0 | 3 | 1.62 | 1.62 | 0.91 | 0 | 3 | 3 | 2 | 4 | 0 | 0 | 0 | 0 | 0 | 0 | 0 |
| NS3 | E7EQV9 | RPL15 | 1 | 2.25 | 0 | 2.25 | 1.46 | 1.46 | 0.89 | 0 | 3 | 2 | 2 | 2 | 0 | 0 | 0 | 0 | 0 | 0 | 0 |
| NS3 | M0R257 | RAB3A | 1 | 2 | 0 | 2 | 1.42 | 1.42 | 0.89 | 0 | 2 | 2 | 2 | 2 | 0 | 0 | 0 | 0 | 0 | 0 | 0 |
| NS3 | C9IZN5 | NDUFA5 | 1 | 2 | 0 | 2 | 1.42 | 1.42 | 0.88 | 0 | 2 | 2 | 2 | 2 | 0 | 0 | 0 | 0 | 0 | 0 | 0 |
| NS3 | A0A2R8Y6J3 | RPL5 | 1 | 2.75 | 0 | 2.75 | 1.58 | 1.56 | 0.87 | 0 | 2 | 2 | 2 | 5 | 0 | 0 | 0 | 0 | 0 | 0 | 0 |
| NS3 | H0YIN8 | ANKRD13A | 1 | 2.25 | 0 | 2.25 | 1.48 | 1.47 | 0.87 | 0 | 2 | 3 | 2 | 2 | 0 | 0 | 0 | 0 | 0 | 0 | 0 |

| BAIT | PROTID | GENE | preyFreq | preyAvg_AvgCounts | preySD_AvgCounts | NS4A_AVG_SPC | NS4A_FC_A | NS4A_FC_B | NS4A_SP | NS4A_iREF | IP_USP7_NS4A_R4 | IP_USP7_NS4A_R3 | IP_USP7_NS4A_R2 | IP_USP7_NS4A_R1 | IP_USP7_GFP_R8 | IP_USP7_GFP_R7 | IP_USP7_GFP_R6 | IP_USP7_GFP_R5 | IP_USP7_GFP_R4 | IP_USP7_GFP_R3 | IP_USP7_GFP_R1 |
|---|---|---|---|---|---|---|---|---|---|---|---|---|---|---|---|---|---|---|---|---|---|
| NS4A | J3KPX7 | PHB2 | 1 | 16.5 | 0 | 16.5 | 13.1 | 12.78 | 1 | 0 | 16 | 20 | 14 | 16 | 0 | 0 | 0 | 0 | 0 | 0 | 0 |
| NS4A | C9JZ20 | PHB | 1 | 13.25 | 0 | 13.25 | 10.03 | 9.17 | 1 | 0 | 12 | 16 | 12 | 13 | 0 | 0 | 0 | 0 | 0 | 0 | 1 |
| NS4A | E9PIE4 | MTCH2 | 1 | 10.5 | 0 | 10.5 | 8.29 | 8.25 | 1 | 0 | 6 | 12 | 12 | 12 | 0 | 0 | 0 | 0 | 0 | 0 | 0 |
| NS4A | F5GY37 | PHB2 | 1 | 11.5 | 0 | 11.5 | 7.92 | 6.57 | 1 | 0 | 9 | 16 | 10 | 11 | 0 | 0 | 0 | 1 | 0 | 0 | 0 |
| NS4A | C9JW96 | PHB | 1 | 9.5 | 0 | 9.5 | 7.86 | 7.8 | 1 | 0 | 8 | 10 | 10 | 10 | 0 | 0 | 0 | 0 | 0 | 0 | 0 |
| NS4A | J3QQY2 | TMCO1 | 1 | 7.75 | 0 | 7.75 | 6.63 | 6.55 | 1 | 0 | 7 | 9 | 7 | 8 | 0 | 0 | 0 | 0 | 0 | 0 | 0 |
| NS4A | C9JAW5 | C9JAW5 | 1 | 7.5 | 0 | 7.5 | 6.47 | 6.39 | 1 | 0 | 7 | 8 | 7 | 8 | 0 | 0 | 0 | 0 | 0 | 0 | 0 |
| NS4A | Q5T8U5 | SURF4 | 1 | 8 | 0 | 8 | 6.39 | 6.21 | 1 | 0 | 3 | 11 | 9 | 9 | 0 | 0 | 0 | 0 | 0 | 0 | 0 |
| NS4A | E9PLN8 | E9PLN8 | 1 | 5.25 | 0 | 5.25 | 4.73 | 4.72 | 1 | 0 | 4 | 6 | 5 | 6 | 0 | 0 | 0 | 0 | 0 | 0 | 0 |
| NS4A | B1APM4 | SOAT1 | 1 | 4.75 | 0 | 4.75 | 4.54 | 4.42 | 1 | 0 | 5 | 4 | 5 | 5 | 0 | 0 | 0 | 0 | 0 | 0 | 0 |
| NS4A | I3L0R8 | DERL2 | 1 | 3.75 | 0 | 3.75 | 3.8 | 3.7 | 0.99 | 0 | 4 | 3 | 4 | 4 | 0 | 0 | 0 | 0 | 0 | 0 | 0 |
| NS4A | A0A590UJA9 | TMUB1 | 1 | 3.75 | 0 | 3.75 | 3.64 | 3.58 | 0.99 | 0 | 3 | 3 | 3 | 6 | 0 | 0 | 0 | 0 | 0 | 0 | 0 |
| NS4A | B5MDE0 | RFT1 | 1 | 3.5 | 0 | 3.5 | 3.52 | 3.5 | 0.98 | 0 | 3 | 4 | 3 | 4 | 0 | 0 | 0 | 0 | 0 | 0 | 0 |
| NS4A | A6NP52 | PRAF2 | 1 | 3.5 | 0 | 3.5 | 3.5 | 3.46 | 0.98 | 0 | 3 | 3 | 3 | 5 | 0 | 0 | 0 | 0 | 0 | 0 | 0 |
| NS4A | E9PC15 | AGK | 1 | 4.5 | 0 | 4.5 | 4.32 | 4.05 | 0.97 | 0 | 5 | 2 | 4 | 7 | 0 | 0 | 0 | 0 | 0 | 0 | 0 |
| NS4A | H7C5W9 | ATP2A2 | 1 | 3.5 | 0 | 3.5 | 3.4 | 3.38 | 0.97 | 0 | 2 | 4 | 3 | 5 | 0 | 0 | 0 | 0 | 0 | 0 | 0 |
| NS4A | A0A1B0GV93 | HSD17B12 | 1 | 3 | 0 | 3 | 3.11 | 3.1 | 0.97 | 0 | 2 | 4 | 3 | 3 | 0 | 0 | 0 | 0 | 0 | 0 | 0 |
| NS4A | C9JI87 | VDAC1 | 1 | 4 | 0 | 4 | 3.71 | 3.38 | 0.96 | 0 | 4 | 5 | 3 | 4 | 0 | 0 | 0 | 0 | 0 | 0 | 1 |
| NS4A | B4E1G1 | DERL1 | 1 | 2.5 | 0 | 2.5 | 2.81 | 2.78 | 0.96 | 0 | 2 | 3 | 3 | 2 | 0 | 0 | 0 | 0 | 0 | 0 | 0 |
| NS4A | F8VVM2 | SLC25A3 | 1 | 11.25 | 0 | 11.25 | 2.94 | 2.05 | 0.95 | 0 | 10 | 11 | 12 | 12 | 2 | 3 | 2 | 2 | 1 | 1 | 0 |

| | | | | | | | | | | | | | | | | | | | | | |
|---|---|---|---|---|---|---|---|---|---|---|---|---|---|---|---|---|---|---|---|---|---|
| NS4A | A0A2R8Y5M6 | BCAP31 | 1 | 2.75 | 0 | 2.75 | 3.16 | 2.97 | 0.94 | 0 | 4 | 3 | 2 | 2 | 0 | 0 | 0 | 0 | 0 | 0 | 0 |
| NS4A | B7Z4M1 | RTN3 | 1 | 3 | 0 | 3 | 2.98 | 2.9 | 0.93 | 0 | 1 | 3 | 3 | 5 | 0 | 0 | 0 | 0 | 0 | 0 | 0 |
| NS4A | E9PNY1 | ZFPL1 | 1 | 2.25 | 0 | 2.25 | 2.64 | 2.61 | 0.93 | 0 | 2 | 3 | 2 | 2 | 0 | 0 | 0 | 0 | 0 | 0 | 0 |
| NS4A | C9JWM2 | SCAMP4 | 1 | 2.75 | 0 | 2.75 | 2.9 | 2.75 | 0.91 | 0 | 1 | 3 | 5 | 2 | 0 | 0 | 0 | 0 | 0 | 0 | 0 |
| NS4A | E9PNW8 | FAR1 | 1 | 2.75 | 0 | 2.75 | 2.85 | 2.79 | 0.91 | 0 | 1 | 4 | 3 | 3 | 0 | 0 | 0 | 0 | 0 | 0 | 0 |
| NS4A | M0R3C3 | TECR | 1 | 4.5 | 0 | 4.5 | 3.91 | 3.7 | 0.9 | 0 | 1 | 6 | 4 | 7 | 0 | 0 | 0 | 0 | 0 | 0 | 0 |
| NS4A | J3QRY4 | PSMD11 | 1 | 2 | 0 | 2 | 2.48 | 2.45 | 0.9 | 0 | 2 | 2 | 2 | 2 | 0 | 0 | 0 | 0 | 0 | 0 | 0 |
| NS4A | F8WBE5 | TFRC | 1 | 2.25 | 0 | 2.25 | 2.47 | 2.28 | 0.88 | 0 | 2 | 2 | 2 | 3 | 0 | 0 | 0 | 0 | 0 | 0 | 1 |
| NS4A | A0A0S2Z5D0 | CLN6 | 1 | 1.75 | 0 | 1.75 | 2.22 | 2.21 | 0.87 | 0 | 1 | 2 | 2 | 2 | 0 | 0 | 0 | 0 | 0 | 0 | 0 |
| NS4A | A0A087X1A9 | SEC22B | 1 | 2.5 | 0 | 2.5 | 2.28 | 1.89 | 0.86 | 0 | 1 | 3 | 2 | 4 | 0 | 0 | 0 | 1 | 0 | 0 | 0 |
| NS4A | B4DR61 | SEC61A1 | 1 | 1.75 | 0 | 1.75 | 2.22 | 2.21 | 0.85 | 0 | 1 | 2 | 2 | 2 | 0 | 0 | 0 | 0 | 0 | 0 | 0 |

| BAIT | PROTID | GENE | preyFreq | preyAvg_AvgCounts | preySD_AvgCounts | NS4B_AVG_SPC | NS4B_FC_A | NS4B_FC_B | NS4B_SP | NS4B_iREF | IP_USP7_NS4B_R4 | IP_USP7_NS4B_R3 | IP_USP7_NS4B_R2 | IP_USP7_NS4B_R1 | IP_USP7_GFP_R8 | IP_USP7_GFP_R7 | IP_USP7_GFP_R6 | IP_USP7_GFP_R5 | IP_USP7_GFP_R4 | IP_USP7_GFP_R3 | IP_USP7_GFP_R1 |
|---|---|---|---|---|---|---|---|---|---|---|---|---|---|---|---|---|---|---|---|---|---|
| NS4B | C9JI87 | VDAC1 | 1 | 14.5 | 0 | 14.5 | 8.1 | 7.14 | 1 | 0 | 15 | 9 | 20 | 14 | 0 | 0 | 0 | 0 | 0 | 0 | 1 |
| NS4B | M0R3C3 | TECR | 1 | 8.5 | 0 | 8.5 | 5.39 | 5.28 | 1 | 0 | 7 | 8 | 11 | 8 | 0 | 0 | 0 | 0 | 0 | 0 | 0 |
| NS4B | C9JAW5 | C9JAW5 | 1 | 7 | 0 | 7 | 4.62 | 4.55 | 1 | 0 | 7 | 7 | 6 | 8 | 0 | 0 | 0 | 0 | 0 | 0 | 0 |
| NS4B | K7EM90 | ENO1 | 1 | 5 | 0 | 5 | 3.61 | 3.51 | 1 | 0 | 4 | 4 | 7 | 5 | 0 | 0 | 0 | 0 | 0 | 0 | 0 |
| NS4B | C9JWM2 | SCAMP4 | 1 | 4.75 | 0 | 4.75 | 3.48 | 3.41 | 1 | 0 | 4 | 4 | 6 | 5 | 0 | 0 | 0 | 0 | 0 | 0 | 0 |
| NS4B | J3QQY2 | TMCO1 | 1 | 3.5 | 0 | 3.5 | 2.79 | 2.77 | 0.98 | 0 | 4 | 3 | 4 | 3 | 0 | 0 | 0 | 0 | 0 | 0 | 0 |
| NS4B | F8WBE5 | TFRC | 1 | 3.75 | 0 | 3.75 | 2.73 | 2.53 | 0.97 | 0 | 4 | 4 | 4 | 3 | 0 | 0 | 0 | 0 | 0 | 0 | 1 |
| NS4B | H7C5W9 | ATP2A2 | 1 | 6.75 | 0 | 6.75 | 4.45 | 4.11 | 0.96 | 0 | 11 | 3 | 9 | 4 | 0 | 0 | 0 | 0 | 0 | 0 | 0 |
| NS4B | K7EJ57 | TOMM40 | 1 | 3.75 | 0 | 3.75 | 2.99 | 2.86 | 0.94 | 0 | 5 | 2 | 3 | 5 | 0 | 0 | 0 | 0 | 0 | 0 | 0 |
| NS4B | I3L1D2 | MPDU1 | 1 | 2.25 | 0 | 2.25 | 2.16 | 2.14 | 0.94 | 0 | 2 | 2 | 3 | 2 | 0 | 0 | 0 | 0 | 0 | 0 | 0 |
| NS4B | B1AR62 | HK1 | 1 | 2.75 | 0 | 2.75 | 2.45 | 2.39 | 0.93 | 0 | 2 | 2 | 4 | 3 | 0 | 0 | 0 | 0 | 0 | 0 | 0 |
| NS4B | E7EQR6 | TCP1 | 1 | 2.75 | 0 | 2.75 | 2.43 | 2.38 | 0.93 | 0 | 4 | 2 | 2 | 3 | 0 | 0 | 0 | 0 | 0 | 0 | 0 |
| NS4B | H0YKD8 | RPL28 | 1 | 2.75 | 0 | 2.75 | 2.4 | 2.36 | 0.93 | 0 | 4 | 2 | 3 | 2 | 0 | 0 | 0 | 0 | 0 | 0 | 0 |
| NS4B | J3KQL8 | APOL2 | 1 | 2.75 | 0 | 2.75 | 2.41 | 2.37 | 0.91 | 0 | 3 | 2 | 4 | 2 | 0 | 0 | 0 | 0 | 0 | 0 | 0 |
| NS4B | H0YL72 | IDH3A | 1 | 3 | 0 | 3 | 2.14 | 1.8 | 0.9 | 0 | 4 | 4 | 2 | 2 | 0 | 0 | 0 | 0 | 0 | 1 | 0 |
| NS4B | D6RFI0 | SFXN1 | 1 | 7.75 | 0 | 7.75 | 2.26 | 1.46 | 0.87 | 0 | 11 | 7 | 7 | 6 | 2 | 1 | 1 | 2 | 0 | 0 | 0 |
| NS4B | F8VQE1 | LIMA1 | 1 | 2 | 0 | 2 | 2.03 | 2.02 | 0.86 | 0 | 2 | 2 | 2 | 2 | 0 | 0 | 0 | 0 | 0 | 0 | 0 |
| NS4B | A0A1B0GWA2 | AGPS | 1 | 2.25 | 0 | 2.25 | 2.15 | 2.14 | 0.85 | 0 | 3 | 2 | 2 | 2 | 0 | 0 | 0 | 0 | 0 | 0 | 0 |

| BAIT | PROTID | GENE | preyFreq | preyAvg_AvgCounts | preySD_AvgCounts | NS5_AVG_SPC | NS5_FC_A | NS5_FC_B | NS5_SP | NS5_iREF | IP_USP7_NS5_R4 | IP_USP7_NS5_R3 | IP_USP7_NS5_R2 | IP_USP7_NS5_R1 | IP_USP7_GFP_R8 | IP_USP7_GFP_R7 | IP_USP7_GFP_R6 | IP_USP7_GFP_R5 | IP_USP7_GFP_R4 | IP_USP7_GFP_R3 | IP_USP7_GFP_R1 |
|---|---|---|---|---|---|---|---|---|---|---|---|---|---|---|---|---|---|---|---|---|---|
| NS5 | H0YKD8 | RPL28 | 1 | 6 | 0 | 6 | 4.23 | 4.22 | 1 | 0 | 7 | 5 | 6 | 6 | 0 | 0 | 0 | 0 | 0 | 0 | 0 |
| NS5 | M0R181 | RPL21 | 1 | 3.75 | 0 | 3.75 | 2.98 | 2.94 | 0.97 | 0 | 6 | 3 | 3 | 3 | 0 | 0 | 0 | 0 | 0 | 0 | 0 |
| NS5 | A0A7I2V2L9 | RPL13A | 1 | 4.5 | 0 | 4.5 | 3.37 | 3.35 | 0.96 | 0 | 6 | 3 | 5 | 4 | 0 | 0 | 0 | 0 | 0 | 0 | 0 |
| NS5 | F2Z388 | RPL35 | 1 | 3 | 0 | 3 | 2.62 | 2.55 | 0.93 | 0 | 5 | 3 | 2 | 2 | 0 | 0 | 0 | 0 | 0 | 0 | 0 |
| NS5 | C9JAW5 | C9JAW5 | 1 | 2.75 | 0 | 2.75 | 2.52 | 2.49 | 0.93 | 0 | 3 | 3 | 2 | 3 | 0 | 0 | 0 | 0 | 0 | 0 | 0 |
| NS5 | D3YTB1 | RPL32 | 1 | 2.75 | 0 | 2.75 | 2.51 | 2.46 | 0.92 | 0 | 4 | 3 | 2 | 2 | 0 | 0 | 0 | 0 | 0 | 0 | 0 |
| NS5 | C9JNW5 | RPL24 | 1 | 3.25 | 0 | 3.25 | 2.61 | 2.55 | 0.88 | 0 | 5 | 1 | 3 | 4 | 0 | 0 | 0 | 0 | 0 | 0 | 0 |

**Table S5. Number and percentage of interacting proteins identified in SH-SY5Y and USP7 cells**

| | SH-SY5Y | Both | USP7 |
|---|---|---|---|
| NS1 | 72 (61.5%) | 9 (7.7%) | 36 (30.8%) |
| NS2B | 108 (32.2%) | 41 (12.2%) | 186 (55.5%) |
| NS3 | 44 (71.0%) | 2 (3.2%) | 16 (25.8%) |
| NS4A | 139 (82.7%) | 15 (8.9%) | 14 (8.3%) |
| NS4B | 47 (72.3%) | 5 (7.7%) | 13 (20.0%) |
| NS5 | 34 (82.9%) | 0 (0%) | 7 (17.1%) |

Note: The "Both" column refers to interacting proteins common to SH-SY5Y and USP7 cells. The "SH-SY5Y" and "USP7" columns refer to interacting proteins found exclusively in SH-SY5Y or USP7 cells, respectively. Numbers and percentages indicate the number of interacting proteins and their proportion relative to the total identified interacting proteins for each viral protein in SH-SY5Y and USP7 cells.
